# Architecture of the cytoplasmic face of the nuclear pore

**DOI:** 10.1101/2021.10.26.465790

**Authors:** Christopher J. Bley, Si Nie, George W. Mobbs, Stefan Petrovic, Anna T. Gres, Xiaoyu Liu, Somnath Mukherjee, Sho Harvey, Ferdinand M. Huber, Daniel H. Lin, Bonnie Brown, Aaron W. Tang, Emily J. Rundlet, Ana R. Correia, Shane Chen, Saroj G. Regmi, Mary Dasso, Alina Patke, Alexander F. Palazzo, Anthony A. Kossiakoff, André Hoelz

## Abstract

The nuclear pore complex (NPC) is the sole bidirectional gateway for nucleocytoplasmic transport. Despite recent progress in elucidating the NPC symmetric core architecture, the asymmetrically decorated cytoplasmic face, essential for mRNA export and a hotspot for nucleoporin-associated diseases, has remained elusive. Here, we report a composite structure of the entire human cytoplasmic face obtained by combining biochemical reconstitution, crystal structure determination, docking into cryo-electron tomographic reconstructions, and physiological validation, accounting for a third of the NPC’s mass. Whereas an evolutionarily conserved ∼540 kDa hetero-hexameric cytoplasmic filament nucleoporin complex is anchored by species-specific motifs above the central transport channel, attachment of the pentameric NUP358 bundles depends on the double-ring arrangement of the coat nucleoporin complex. Our results and the predictive power of our composite structure provide a rich foundation for elucidating the molecular basis of mRNA export and nucleoporin diseases.

**One sentence summary:** An interdisciplinary analysis established the near-atomic molecular architecture of the cytoplasmic face of the human nuclear pore complex.

## Introduction

The sequestration of genetic material in the nucleus represents one of the great hallmarks of evolution but creates the necessity for selective bidirectional transport across the nuclear envelope (*1–4*). The nuclear pore complex (NPC) is the sole gateway through which folded proteins and protein/nucleic acid complexes cross the nuclear envelope, making this transport organelle an essential machine for all eukaryotic life. Besides its direct role as a transport channel, the NPC serves as an organizer for nuclear and cytoplasmic processes that are essential for the flow of genetic information from DNA to RNA to protein, including transcription, spliceosome assembly, mRNA export, and ribosome assembly (*1–4*). Dysfunction of the NPC or its components represents a major cause of human disease (*2, 5, 6*).

Architecturally, the NPC consists of a central core with an 8-fold rotational symmetry across a nucleocytoplasmic axis and a two-fold rotational symmetry across the plane of the nuclear envelope, which links to compartment-specific asymmetric “cytoplasmic filaments” (CF) and a “nuclear basket” structure (Fig. 1A) (*1, 2*). The NPC is built from ∼34 different proteins, termed nucleoporins (nups) that are organized into distinct subcomplexes. Multiple copies of each nup in the NPC add up to an assembly that reaches an extraordinary molecular mass of ∼110 MDa in vertebrates. The symmetric core of the NPC is composed of an inner ring and two spatially segregated outer rings. The inner ring is embedded in nuclear envelope pores generated by the circumscribed fusion of the double membrane of the nuclear envelope. The diffusion barrier is formed by unstructured phenylalanine-glycine (FG) repeats that fill the central transport channel, imposing an ∼40 kDa limit on passive diffusion (*1–4*). Transport factors collectively termed karyopherins overcome the diffusion barrier by binding to FG repeats, thereby transporting cargo across the nuclear envelope (*7–9*). A significant fraction of the FG repeats in the inner ring is contributed by a hetero-trimeric channel nup complex (CNT) which is anchored by a single assembly sensor motif (*10–12*). The outer rings sit atop the nuclear envelope, sandwiching the inner ring from both sides. They are primarily formed by the Y-shaped coat nup complex (CNC) and serve as a platform for the asymmetric incorporation of the CF and nuclear basket nups.

**Fig. 1.**
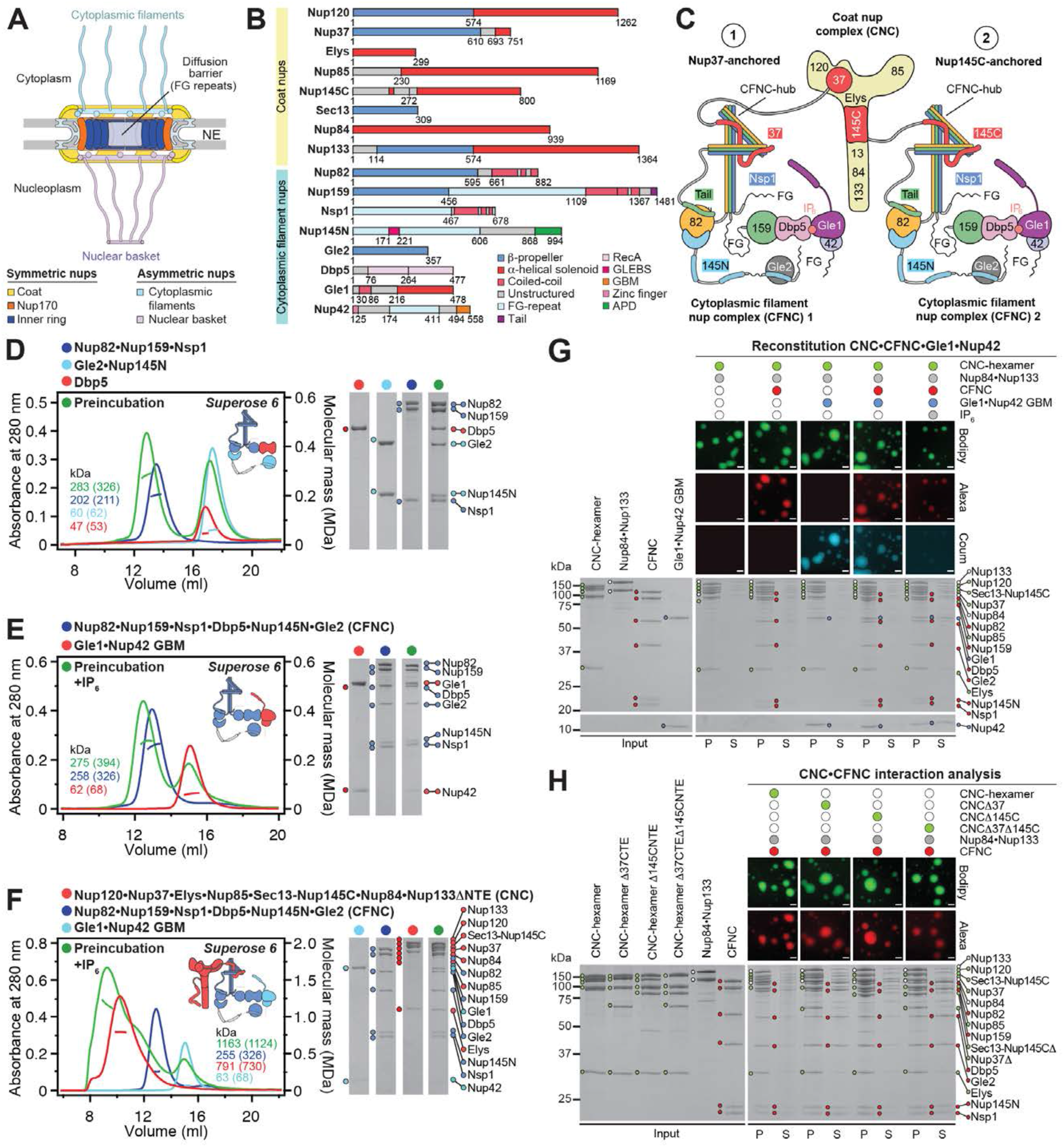
Reconstitution of a 16-protein *C. thermophilum* coat-cytoplasmic filament nup complex. (**A**) Cross sectional conceptual schematic of the fungal NPC architecture. (**B**) Domain structures of the coat and cytoplasmic filament nups. Domains are drawn as horizontal boxes with residue numbers indicated and their observed or predicted folds colored according to the legend. (**C**) Schematic representation summarizing the results of our biochemical reconstitution and dissection experiments with purified recombinant *C. thermophilum* nups, illustrating the cytoplasmic filament nup complex (CFNC) architecture and its attachment to the coat nup complex (CNC). The CNC harbors two assembly sensors, Nup37^CTE^ and Nup145C^NTE^, each anchoring a CFNC by binding to its central hub. (**D**-**F**) SEC-MALS interaction analyses, showing the stepwise biochemical reconstitution starting with (D) the ∼290 kDa CFNC (green) from Nup82•Nup159•Nsp1 (blue), Gle2•Nup145N (cyan), and Dbp5 (red), (E) CFNC•Gle1•Nup42^GBM^ (green) from CFNC (blue) and Gle1•Nup42^GBM^ (red), and culminating with (F) the ∼1.2 MDa 16-protein CNC•CFNC•Gle1•Nup42^GBM^ complex (green) from CNC (red), CFNC (blue), and Gle1•Nup42^GBM^ (cyan). SDS-PAGE gels strips of peak fractions are shown. Measured molecular masses are indicated, with respective theoretical masses provided in parentheses. (**G**, **H**) Liquid-liquid phase separation (LLPS) interaction assays, assessing (G) CFNC (red) and Gle1•Nup42^GBM^ (cyan) incorporation into CNC-LLPS (green), and (H) CFNC incorporation into CNC-LLPS, lacking either one or both Nup37^CTE^ and Nup145C^NTE^ assembly sensors. N-terminally fluorescently labeled CNC (Bodipy), CFNC (Alexa Fluor 647), and Gle1•Nup42^GBM^ (Coumarin) were visualized by fluorescence microscopy. Pelleted CNC condensate phase (P) and soluble (S) fractions were analyzed by SDS-PAGE and visualized by Coomassie brilliant blue staining. Scale bars are 10 μm.

Two decades ago, the atomic level characterization of the NPC began with individual nup domains and progressed to nup complexes of increasing size and complexity, culminating in the ∼400 kDa hetero-heptameric Y-shaped CNC (*11, 13–28*). Simultaneously, advances in the acquisition and processing of cryo-electron tomographic (cryo-ET) data gradually increased the resolution of intact NPC 3D reconstructions (*29*). Docking of the CNC into a ∼32 Å cryo-ET map of the intact human NPC demonstrated that two reticulated eight-membered CNC rings, linked by head-to-tail interactions, are present on each side of the nuclear envelope (*27, 30*). Moreover, this advance established that the resolution gap between high- and low-resolution structural methods can be overcome by combining biochemical reconstitution and X-ray crystallographic characterization of nups with cryo-ET reconstruction of the intact NPC. Expansion of this approach to the nine nups constituting the inner ring rapidly led to the reconstitution of two distinct ∼425 kDa inner ring complexes and the elucidation of their components’ structures (*10-12, 20, 31-38*). In turn, this advance enabled the determination of the near-atomic composite structure of the entire ∼56 MDa symmetric core of the human NPC, establishing the stoichiometry and placement of all 17 symmetric nups within a ∼23 Å cryo-ET reconstruction (*38, 39*). Subsequently, the architecture of the *Saccharomyces cerevisiae* NPC was determined with a similar approach, utilizing high-resolution nup crystal structures and ∼25 Å cryo-ET maps of either detergent purified or *in-situ* NPCs (*40, 41*). Compared to the human NPC, the *S. cerevisiae* NPC lacks the distal CNC ring and associated nups on both sides of the nuclear envelope, but the relative nup arrangement within the rest of the symmetric core remains essentially identical (*38, 39, 42*).

Projecting into the cytoplasm from the cytoplasmic face of the NPC, the CF nups recruit cargo•transport factor complexes for nucleocytoplasmic transport and orchestrate the export and remodeling of messenger ribonucleoprotein particles (mRNPs) in preparation for translation (*2, 43*). The nine-component CF nup machinery represents a major hotspot for human diseases ranging from degenerative brain disorders and cardiac diseases to cancer (*2, 5, 6*). Although linked to the human CF nups NUP358, NUP214, NUP62, NUP88, NUP98, GLE1, NUP42, RAE1, and DDX19, the pathophysiology and optimal therapeutic strategies for these conditions remain ill-defined.

Here, we present insight into the atomic and higher order architecture, function, and mechanism of action of the CF nups in the human NPC and the NPC of the thermophilic fungus *Chaetomium thermophilum*. First, we uncover a conserved modular architecture within the hetero-hexameric CF nup complex (CFNC) of both species: Holding the CFNC together is a coiled-coil hub built like the CNT, but formed by NUP62 with the C-terminal regions of NUP88 and NUP159, while their N-terminal β-propeller domains link to the mRNA export factors NUP98•RAE1 and the DEAD-box RNA helicase DDX19, respectively, which in turn recruit the remaining complex components. We further uncover evolutionary divergent mechanisms for the attachment of the intact CFNC at the cytoplasmic face of the NPC, which in *C. thermophilum* involves two distinct assembly sensors in the CNC that do not exist in humans. We are able to assemble the *C. thermophilum* CNC and CF nups into a ∼1.1 MDa 16-protein complex and find that it can be remodeled by inositol hexaphosphate (IP_6_). Towards dissecting the molecular mechanism of mRNA export, we systematically characterize the propensity of CF nups for RNA binding and find novel capabilities in two CFNC subcomplexes GLE1•NUP42 and NUP88•NUP214•NUP98 as well as different parts of the metazoan specific NUP358. To build a composite structure of the human NPC cytoplasmic face, we determine crystal structures of the NUP88^NTD^•NUP98^APD^ complex and all remaining structurally uncharacterized regions of NUP358, uncovering a hereto unobserved, S-shaped fold of three α-helical solenoids for the NUP358 N-terminal domain as well as a complex mechanism for NUP358 oligomerization. Docking of the novel structures along with previously characterized CF nups into a previously reported ∼23 Å and an unpublished ∼12 Å cryo-ET map of the intact human NPC (provided by the Beck group), as well as an ∼8 Å region of an anisotropic single particle cryo-EM composite map of the *Xenopus laevis* cytoplasmic NPC face accounts for all of the asymmetric density on the cytoplasmic NPC-side resolved in the maps (*44–46*). Validating our quantitative docking analysis in human cells engineered to enable rapid, inducible NUP358 depletion, we surprisingly find NUP358 to be dispensable for the architectural integrity of the assembled interphase NPC and mRNA export but having a general role in translation. The docking of the CFNC-hub in close proximity to a NUP93 fragment that, in the inner ring, acts as the assembly sensor for the CNT, allows us to predict and experimentally confirm that NUP93 also recruits the structurally related CFNC on the cytoplasmic face, thereby enabling identification of the elusive human CFNC NPC anchor. Thus, our near-atomic composite structure possesses predictive power, demonstrating its general utility for the mechanistic dissection of essential cellular events occurring on the cytoplasmic face of the human NPC.

## RESULTS

### Modular architecture of the evolutionarily conserved 6-protein cytoplasmic filament nup complex

Although pair-wise interactions between selected CF nups had previously been reported, comprehensive knowledge on the entire CF nup interaction network has remained unavailable to date (*11, 47–64*). We had previously found that nups from *C. thermophilum* exhibit superior biochemical stability, allowing us to overcome long-standing technical challenges and elucidate the interaction network of the seventeen symmetric core nups (*38*). We therefore first sought to establish the protein-protein interaction network and complex stoichiometry of the eight evolutionarily conserved *C. thermophilum* CF nups Nup82, Nup159, Nsp1, Nup145N, Gle2, Dbp5, Gle1, and Nup42 (*2*) (Fig. 1B and fig. S1). Most CF nups contain both structured and unstructured regions that can harbor multiple distinct binding sites and FG repeats (Fig. 1B). We established expression and purification protocols for the *C. thermophilum* CF nups, omitting FG-repeat regions as well as an unstructured linker region in Nup145N to improve solubility, and analyzed their binding by size-exclusion chromatography coupled with multiangle light scattering (SEC-MALS) (figs. S1 to S3 and tables S1 to S5).

Nup82 and Nup159 both contain N-terminal β-propeller domains and C-terminal tripartite coiled-coil segments (CCS1-3), which have previously been shown to complex with the similarly built C-terminal coiled-coil region of Nsp1 (*49, 65*). Another complex has previously been shown to be formed between the Gle2 β-propeller and the Gle2-binding sequence (GLEBS) motif of Nup145N (*48, 57*). Mixing of the Nup82•Nup159•Nsp1 and Gle2•Nup145N complexes with the DEAD-box RNA helicase Dbp5 resulted in a stoichiometric hetero-hexameric complex that we will refer to as the cytoplasmic filament nup complex (CFNC) (Fig. 1, C and D and fig. S4A and table S6). To further map CF nup interactions within this complex, we systematically removed individual fragments and domains from the CFNC and probed for remaining interactions (fig. S4). Consistent with previous characterizations of isolated CF nup pairs and trimers (*11, 47–64*), this analysis established that the N-terminal β-propeller domain of Nup82 binds an α-helical TAIL segment in Nup159 and the autoproteolytic domain (ADP) of Nup145N, which in turn recruits Gle2 (Fig. 1C and fig. S4). Analogously, the Nup159 N-terminal β-propeller domain provides the binding site for the Dbp5 (Fig. 1C and fig. S4). These results suggested that the CFNC must be held together by the C-terminal coiled-coil regions of Nup82, Nup159, and Nsp1. Indeed, when the CFNC is divided into four distinct stable subcomplexes, the Nup82•Nup159•Nsp1 C-terminal coiled-coil hetero-trimer, Nup82^NTD^•Nup159^TAIL^•Nup145N^APD^, Nup159^NTD^•Dbp5, and Gle2•Nup145N^GLEBS^, their mixing did not result in any additional interactions and the four subcomplexes did not bind each other, demonstrating that the entire CFNC is held together by interactions in the Nup82•Nup159•Nsp1 C-terminal coiled-coil heterotrimer (fig. S5). In addition to the CFNC, Nsp1 is also an essential part of the hetero-trimeric CNT, where it forms a parallel array of three coiled-coil segments together with Nup49 and Nup57 (*10, 11*). Given these similarities, we wondered whether the coiled-coil segments of Nup82•Nup159•Nsp1 also formed a parallel coiled-coil domain. Indeed, we were able to reconstitute separate stochiometric complexes of the CCS1 and the CCS2-3 coiled-coil regions of Nup82•Nup159•Nsp1, consistent with the CCS1-3 coiled-coil regions of Nup82, Nup159, and Nsp1 forming a CFNC-hub similar to the CNT (fig. S6).

Having characterized the protein-protein interactions within the CFNC, we next tested whether the reconstituted complex was capable of binding to the hetero-octameric CNC (Nup120•Nup37•Elys•Nup85•Sec13-Nup145C•Nup84•Nup133*^Δ^*^NTE^), its neighbor in the intact NPC (fig. S7). Indeed, the reconstituted CFNC and CNC formed a stoichiometric 1:1 complex that was dependent on the CFNC-hub (fig. S8). Because the structurally related CNT is recruited to NPCs through a 39-residue α-helical assembly sensor supplied by Nic96 (*11*), we next asked whether the CNC harbors equivalent assembly sensors for the CFNC-hub. Among the CNC subunits, Nup133, Nup145C, Nup85, and Nup37 each have N- or C-terminal primarily unstructured extensions (NTEs, CTEs) that could potentially contain assembly sensors and were tested for binding to the CFNC-hub. This systematic analysis identified Nup37^CTE^ and Nup145C^NTE^ as binding partners (fig. S9). Deletion of the CTE from Nup37 abolished its binding to the CFNC-hub showing that Nup37^CTE^ is both necessary and sufficient for binding (fig. S10). Moreover, removal of Nup37^CTE^ from the intact CNC substantially reduced its interactions with the CFNC (fig. S11A). While the removal of Nup145C^NTE^ alone from the CNC had no detectable effect, simultaneous elimination of both Nup37^CTE^ and Nup145C^NTE^ almost completely abolished the interaction with CFNC (fig. S11, B and C). Preferential CFNC-hub recruitment by Nup37^CTE^ persisted when the CNC was dissected into its three subcomplexes, with residual and stoichiometric interactions formed with Sec13•Nup145C and Nup120•Nup37•Elys•Nup85, respectively (fig. S12). Vestigial CFNC binding to a CNC lacking both Nup37^CTE^ and Nup145C^NTE^ was mapped to a weak CNC-interaction with Gle2-Nup145N^GLEBS^ (fig. S13).

In the CNT•Nic96 crystal structure, the 39-residue Nic96 assembly sensor forms two ∼10-residue α-helical segments joined by an ∼15-residue linker, with the C-terminal helix being necessary and sufficient for CNT binding (*11*). Secondary structure predictions for both Nup37^CTE^ and Nup145C^NTE^ suggest comparable arrangements of two shorter α-helices separated by a linker from a longer α-helix. Moreover, Nup145C^NTE^ interactions have previously been mapped to an equivalent region, encompassing the C-terminal helix followed by a short unstructured segment (*62*). Through systematic truncation of Nup37^CTE^, we mapped its minimal CFNC-hub binding site to the predicted C-terminal helix and a short unstructured region (fig S14). This demonstrates that the predicted C-terminal α-helix in both Nup37^CTE^ and Nup145C^NTE^ assembly sensors is necessary and sufficient for CFNC-hub binding, whereas the N-terminal helices likely adopt an organizational role in stabilizing a specific tertiary structure of the three coiled-coil segments, as previously observed for the CNT-Nic96 interaction (*11*).

Next, we asked whether Nup145C^NTE^ and Nup37^CTE^ function as *bona fide* assembly sensors for the CFNC-hub by recognizing intact hetero-trimeric coiled-coils but not isolated CFNC-hub constituents. Pulldown experiments probing Nup37^CTE^ or Nup145C^NTE^ with bacterial lysates containing individual CFNC-hub components or a mixture of all three, revealed that both fragments bound exclusively to the assembled hetero-trimeric CFNC-hub, but failed to recognize individual component nups (fig. S15, A and B). Moreover, splitting the CFNC-hub into CCS1 and CCS2-3 portions revealed that both Nup37^CTE^ and Nup145C^NTE^ bind preferentially to CCS2-3, albeit weaker than to the intact CFNC-hub CCS1-3 (figs. S6 and S15C).

Because our SEC-MALS analysis indicated that the CNC only binds to a single CFNC copy, we investigated whether Nup37^CTE^ and Nup145C^NTE^ can simultaneously bind to the CFNC-hub. We found that Nup37^CTE^ and Nup145C^NTE^ bind the CFNC-hub in a mutually exclusive manner, via a hydrophobic interface stable in a high salt (1 M NaCl) buffer (figs. S15D and S16). In fact, Nup37^CTE^ outcompetes Nup145C^NTE^ for CFNC-hub binding, consistent with both our intact CNC and CFNC interaction experiments (figs. S12 and S17). Although we conducted our SEC-MALS analyses at the highest feasible concentrations, we acknowledge the apparent discrepancy arising from the detection of a single CFNC binding to an intact CNC that harbored two assembly sensor binding sites, yet at permissible concentrations Nup145C^NTE^ interactions are unstable.

In addition to the hetero-hexameric CFNC analyzed thus far, Gle1 and Nup42 locate to the cytoplasmic face of the NPC, where they are involved in the export and remodeling of mRNPs (*63*). Gle1 can interact with Dbp5 in an IP_6_-dependent manner and also binds Nup42 at its Gle1-binding motif (GBM) (*53, 58, 63, 66*). To further characterize nup complex interactions at the cytoplasmic NPC side, we tested for incorporation of Gle1•Nup42^GBM^ into the CFNC or CNC. Despite binding only weakly to its known interaction partner Dbp5 in isolation, Gle1•Nup42^GBM^ formed a stable, stochiometric, hetero-octameric complex with the CFNC that cannot be mapped to any of its sub-complexes, indicating a distributed binding site (Fig. 1E and figs. S18 to S21). Furthermore, Gle1•Nup42^GBM^ alone does not interact with the CNC, but it can be incorporated into the CNC•CFNC complex, forming a stoichiometric ∼1.2 MDa 16-protein nup assembly (Fig. 1F and figs. S22 and S23A). Because the Gle1-Dbp5 interaction is IP_6_-dependent (*53, 66*), we carried out all SEC-MALS interaction experiments involving Gle1•Nup42^GBM^ in the presence of IP_6_. However, when Gle1•Nup42^GBM^, CNC, and CFNC were mixed in the absence of IP_6_, the resulting complex displayed a molecular mass of ∼2.0 MDa, consistent with the formation of a stochiometric dimer of the 16-protein complex (fig. S23B). We systematically mapped this biochemical behavior and identified a novel interaction between Gle1^CTD^•Nup42^GBM^ and the CNC, establishing that the CNC-CF nup interaction network can be remodeled by IP_6_ (figs. S23 to S25).

For the Y-shaped CNC, we had previously observed liquid-liquid phase separation (LLPS) when subcomplexes representing its upper arms (Nup120•Nup37•Elys•Nup85•Sec13•Nup145C, CNC-hexamer) and base (Nup84•Nup133) were mixed (*38*). This LLPS formation depended on the N-terminal extension (NTE) of Nup133, which mediates head-to-tail *trans*-interactions between CNCs (*21*). In fact, interaction analyses involving CNCs by SEC-MALS require prevention of LLPS formation by removal of Nup133^NTE^ and are also limited by an inherent CNC solubility limit of 8 μM. To rule out having missed interactions between coat and CF nups in our SEC-MALS analyses because of these limitations, we re-tested binding behaviors by measuring the incorporation of the respective CF nup complexes into a CNC LLPS condensate. Importantly, these interaction analyses performed at higher concentrations corroborated all of our SEC-MALS findings (Fig. 1, G and H, and fig. S26).

Given the special importance of the CF nups for human disease, we next tested whether the molecular architecture of the CFNC is evolutionarily conserved from *C. thermophilum* to humans. The human CFNC is comprised of NUP88 (*ct*Nup82), NUP214 (*ct*Nup159), NUP62 (*ct*Nsp1), DDX19 (*ct*Dbp5), RAE1 (*ct*Gle2), and NUP98 (*ct*Nup145N). Apart from a rearrangement of the FG-repeat and coiled-coil regions in NUP214, the domain organization of the human CFNC nups is identical to that of the *C. thermophilum* orthologs (Fig. 2, A and B and fig. S27). As in *C. thermophilum*, we developed expression and purification protocols omitting FG-repeat and unstructured regions and analyzed CFNC nup interactions by SEC-MALS. Indeed, mixing of the NUP88•NUP214•NUP62 hetero-trimer with RAE1•NUP98 and DDX19 resulted in a monodisperse stoichiometric ∼337 kDa *H. sapiens* CFNC hetero-hexamer (Fig. 2C and figs. S28 and S29). To establish whether the *hs*CFNC adopts the same modular architecture as in *C. thermophilum*, we dissected the human CF nups to generate CFNC sub-complexes and carried out a systematic pairwise SEC-MALS interaction analysis (Fig. 2D and figs. S30 to S39). Showing complete evolutionary conservation, the human CFNC could be divided into the same four subcomplexes we found in our analysis of the *C. thermophilum* CF nups: NUP88^NTD^•NUP98^APD^•NUP214^TAIL^, NUP214^NTD^•DDX19, RAE1•NUP98^GLEBS^, and the NUP88•NUP214•NUP62 coiled-coil complex (*hs*CFNC-hub) (Fig. 2D and figs. S30 to S39). Because the identified Nup37 and Nup145C coat nup assembly sensors for the intact CFNC in *C. thermophilum* each contain ∼60-residue regions with three short α-helices, we conducted secondary structure predictions of all ten human coat nups. This analysis confirmed the presence of extended unstructured regions in NUP96, NUP107, and NUP133, yet analogous assembly sensor motifs could not be identified.

**Fig. 2.**
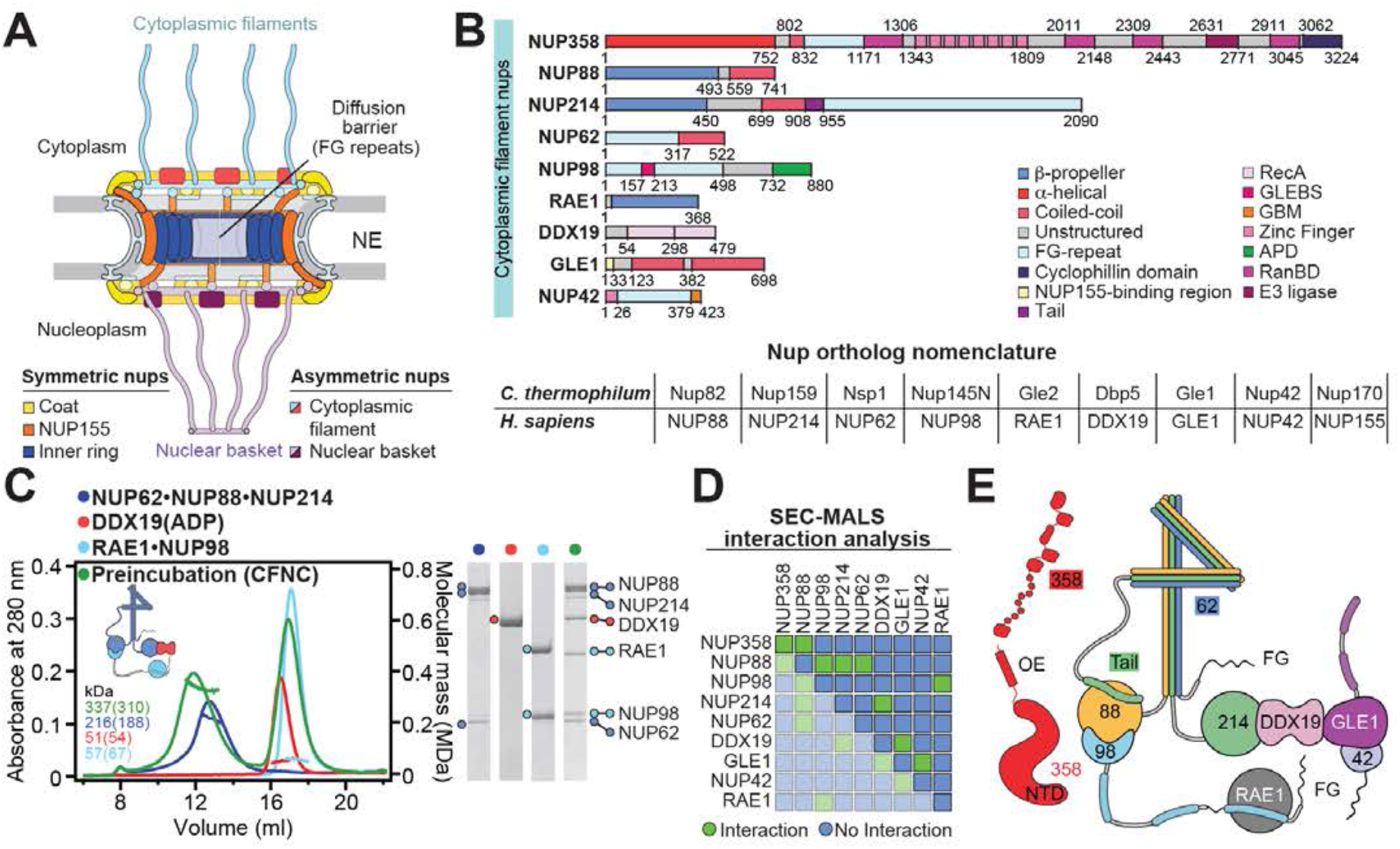
Evolutionarily conserved modular architecture of the hetero-hexameric human CFNC. (**A**) Cross sectional conceptual schematic of the human NPC architecture. (**B**) Domain structures of human cytoplasmic filament nups. Domains are drawn as horizontal boxes with residue numbers indicated and their observed or predicted folds colored according to the legend. For reference, the nomenclature of *H. sapiens* and *C. thermophilum* nup orthologs is indicated. (**C**) Biochemical reconstitution of the ∼310 kDa hetero-hexameric human CFNC. SEC-MALS interaction analysis of NUP88•NUP214•NUP62 (blue), DDX19 (ADP) (red), RAE1•NUP98 (cyan), and their preincubation (green). Measured molecular masses are indicated, with theoretical masses provided in parentheses. SDS-PAGE gel strips of peak fractions visualized by Coomassie brilliant blue staining are shown. (**D**) Summary of pairwise SEC-MALS interaction analyses between human cytoplasmic filament nups. (**E**) Schematic summary of the human CFNC architecture and the cytoplasmic filament nup interaction network.

Together, our data establish that the CF nups form an evolutionarily conserved 6-protein complex that is held together by an extensive parallel coiled-coil hub generated by the C-terminal regions of Nup82/NUP88, Nup159/NUP214 and Nsp1/NUP62, which shares architectural similarities with the heterotrimeric Nsp1/NUP62•Nup49/NUP58•Nup57/NUP54 CNT (*11*). The Nup82/NUP88 N-terminal β-propeller domain is attached by an interaction between the C-terminal α-helical TAIL fragment of Nup159/NUP214 and provides a binding site for the Nup145N/NUP98 APD, which in turn recruits Gle2/RAE1 to the NPC. Analogously, the Nup159/NUP214 N-terminal β-propeller domain provides a binding site for the DEAD-box helicase Dbp5/DDX19. In *C. thermophilum*, the CFNC-hub is anchored to the CNC by two distinct assembly sensors in Nup37^CTE^ and Nup145C^NTE^, similar to the anchoring of the CNT by the Nic96^R1^/NUP93^R1^ assembly sensor in the inner ring. In contrast, the human CNC lacks comparable assembly sensor motifs, suggesting alternative mechanisms for anchoring CF nups at the cytoplasmic face of the human NPC.

### RNA interactions of human cytoplasmic filament nups

Given their essential roles in mRNA export, we next sought to identify which of the human CF nups possessed RNA-binding capabilities (*50, 67–70*). Until now, the RNA binding properties of CF nups have been addressed by disparate methods, utilizing a wide but inconsistent selection of RNA and DNA probes, with several protein constituents remaining untested. Previous work established DDX19 and RAE1•NUP98^GLEBS^ binding to U_10_ single stranded ssRNA, degenerate ssRNA, poly(A), poly(C), poly(G) RNA, as well as ssDNA and double stranded dsDNA across a variety of assays (*50, 56, 57, 63, 71*).

Taking advantage of our complete set of purified human CF nup domains and sub-complexes, we carried out a comprehensive electrophoretic mobility shift assay (EMSA) screen to systematically assess binding against a consistent set of ss/dsRNA probes (Fig. 3). We selected a 12 bp GC-rich stem loop capped by a UUCG tetraloop dsRNA and U_20_ ssRNA as generic probes. Positive RNA interactions where subsequently validated by titrating the protein concentration in EMSAs to establish approximate apparent binding constants (*K_D_*s) (fig. S40). Additionally, to establish whether the observed binding was RNA-specific, we tested all CF nups for binding to ss/dsDNA using a U1 3’ box ssDNA oligonucleotide in the absence and presence of a complimentary ssDNA oligonucleotide.

**Fig. 3.**
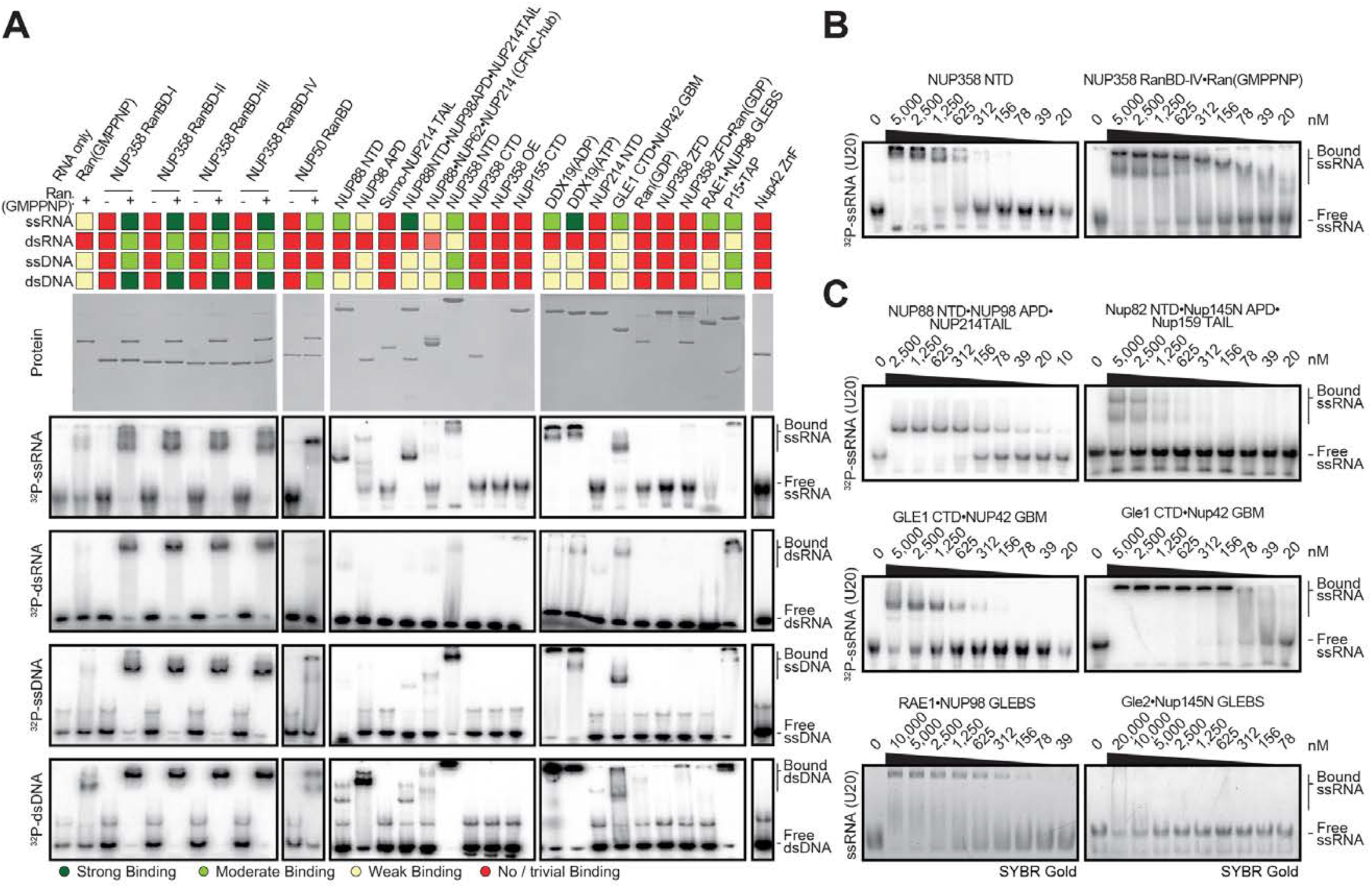
Evolutionary conservation of cytoplasmic filament RNA binding properties. (**A**) Human cytoplasmic filament nup domains and complexes were assayed for binding to single stranded (ss) and double stranded (ds) RNA and DNA probes by electrophoretic mobility shift assay (EMSA). Input proteins resolved by SDS-PAGE were visualized by Coomassie brilliant blue staining. Qualitative assessment of nucleic acid binding is denoted by color-coded boxes. Initial analysis was performed with 5 μM protein and ^32^P-5’ end labeled probes; ssRNA (U_20_), dsRNA (28-nt hairpin), ssDNA (34-nt DNA oligonucleotide), dsDNA (34-bp hybridized DNA oligonucleotides). (**B**, **C**) EMSAs with ssRNA titrated against (B) metazoan-specific NUP358^NTD^ and NUP358^RanBD-IV^•Ran(GMPPNP), and (C) the indicated *H. sapiens* CFNC subcomplexes and their *C. thermophilum* orthologs. RAE1•NUP98^GLEBS^ and Gle2•Nup145C^GLEBS^ EMSAs were performed with unlabeled RNA and visualized with SYBR Gold stain.

First, we assayed the CFNC nups for nucleic acid binding. Both RAE1•NUP98^GLEBS^ and DDX19 bound ss/dsRNA, as previously shown (fig. S40, B and C) (*50, 57*). We discovered moderate binding of NUP88^NTD^ and GLE1^CTD^•NUP42^GBM^ to ssRNA (Fig. 3C and fig. S40A). In the context of the NUP88^NTD^•NUP98^APD^•NUP214^TAIL^ complex, NUP88^NTD^ binding was specifically enhanced to ssRNA, exhibiting a *K_D_*≈150 nM compared to *K_D_*s >5 μM for dsRNA and ss/dsDNA probes (fig. S40A). In contrast, GLE1^CTD^•NUP42^GBM^ bound both ssRNA and ssDNA weakly with *K_D_*s >2 μM. We found that the orthologous *C. thermophilum* Nup82^NTD^•Nup145C^APD^•Nup159^TAIL^, Gle1^CTD^•Nup42^GBM^, and Gle2•Nup145C^GLEBS^ complexes bound ssRNA, demonstrating that RNA binding is an evolutionarily conserved feature of CFNC nups (Fig. 3C and fig. S40D).

Next, we analyzed the RNA-binding properties of the metazoan-specific NUP358, taking advantage of structurally well-defined domain boundaries for the N-terminal domain (NTD), oligomerization element (OE), four distinct Ran-binding domains (RanBDs), a zinc-finger domain (ZFD) comprising eight C4-type zinc-finger motifs (ZnF) arranged in tandem, and the C-terminal prolyl isomerase domain (CTD) (see below) (*72–75*). Previously, NUP358^ZFD^ was shown to bind to a 63 nt RNA fragment encoding the signal sequence coding region (SSCR) of insulin, an interaction that was abolished in the presence of Ran(GDP/GTP) (*76*). Using our four nucleic acid probes, we did not detect RNA binding for NUP358^OE^, NUP358^ZFD^, or NUP358^CTD^ at the assayed concentrations (Fig. 3A). However, we detected moderate RNA and DNA binding for NUP358^NTD^ (Fig. 3B and fig. S40G). Because the previously identified insulin SSCR RNA binding partner for NUP358 was sensitive to degradation, we systematically tested RNAs encoding the SSCRs of other secretory proteins. This analysis identified the 63 nt placental alkaline phosphatase (ALPP) SSCR RNA as most amenable to biochemical studies (fig. S40I). Whereas we only observed an insubstantial shift of ALPP^SSCR^ RNA with NUP358^ZFD^, NUP358^NTD^ robustly bound to ALPP^SSCR^ RNA with a K_D_≈150 nM, ∼4-fold tighter than our generic ssRNA or dsRNA probes (fig. S40I). Unexpectedly, we also found that the four NUP358^RanBD^•Ran(GMPPNP) complexes preferentially bound ssRNA, with a K_D_≈500 nM, over dsRNA and ss/dsDNA (Fig. 3B and fig. S40E). To test whether RNA binding is a general property of RanBDs when bound to Ran, we assayed the related nuclear basket NUP50^RanBD^•Ran(GMPPNP) complex, but detected weaker binding to RNA and DNA (fig. S40, E and F).

Together, our systematic CF nup analysis confirmed previously established RNA binding sites in DDX19, p15•TAP, and RAE1•NUP98^GLEBS^, whilst uncovering novel sites in NUP88^NTD^•NUP214^TAIL^•NUP98^APD^, GLE1^CTD^•NUP42^GBM^, NUP358^NTD^, and the four NUP358 RanBD•Ran(GMPPNP) complexes. Future studies need to delineate whether these RNA binding sites present sequence specific RNA affinity, and what the implications of such specificity would be in the overall mRNA export pathway.

### Structural and biochemical analyses of NUP358

Composed of a 3,224-residue polypeptide, NUP358 is a metazoan-specific additional CF component and the largest constituent of the NPC (*72, 75, 77, 78*). Previous studies established that its N-terminal ∼900-residue α-helical region is necessary for nuclear envelope recruitment (*79*). Within this region, the first 145 residues have been biochemically and structurally characterized, shown to form three tetratricopeptide repeats (TPR) (*80*). Guided by secondary structure predictions, we systematically screened expression constructs for solubility, identifying three fragments: NUP358^NTD^*^Δ^*^TPR^ (residues 145-752), NUP358^NTD^ (residues 1-752), and an extended region spanning residues 1-832 (NUP358^1-832^). Subsequent purifications revealed that the NUP358^NTD^ and NUP358^1-832^ fragments behave differently, with the latter forming amorphous precipitates in buffer NaCl concentrations below 300 mM. Therefore, we characterized these NUP358 fragments in both high salt (350 mM NaCl) and low salt (100 mM NaCl) buffers, wherever possible.

NUP358^NTD^ exhibited concentration-dependent homo-dimerization in low salt buffer, with measured molecular masses between values corresponding to monomeric and dimeric species but existed as a monomeric species in high salt buffer at all NUP358^NTD^ concentrations tested (Fig. 4, A and B and fig. S41). Conversely, NUP358^NTD^*^Δ^*^TPR^ was exclusively monomeric, suggesting the TPR mediates homo-dimerization (Fig. 4B and fig. S41). Furthermore, the extended NUP358^1-832^ fragment forms oligomers with measured molecular masses between those of a tetramer and a pentamer (Fig. 4C and fig. S41). Subsequent C-terminal mapping revealed an oligomerization element (OE) located between residues 802-832. NUP358^OE^ formed concentration-dependent oligomers of measured molecular mass ranging between those expected for dimeric and tetrameric species in both low and high salt buffers (Fig. 4G and fig. S42). Thus, NUP358 oligomerization is mediated by the TPR and OE regions, each located on opposite sides of the N-terminal α-helical region.

**Fig. 4.**
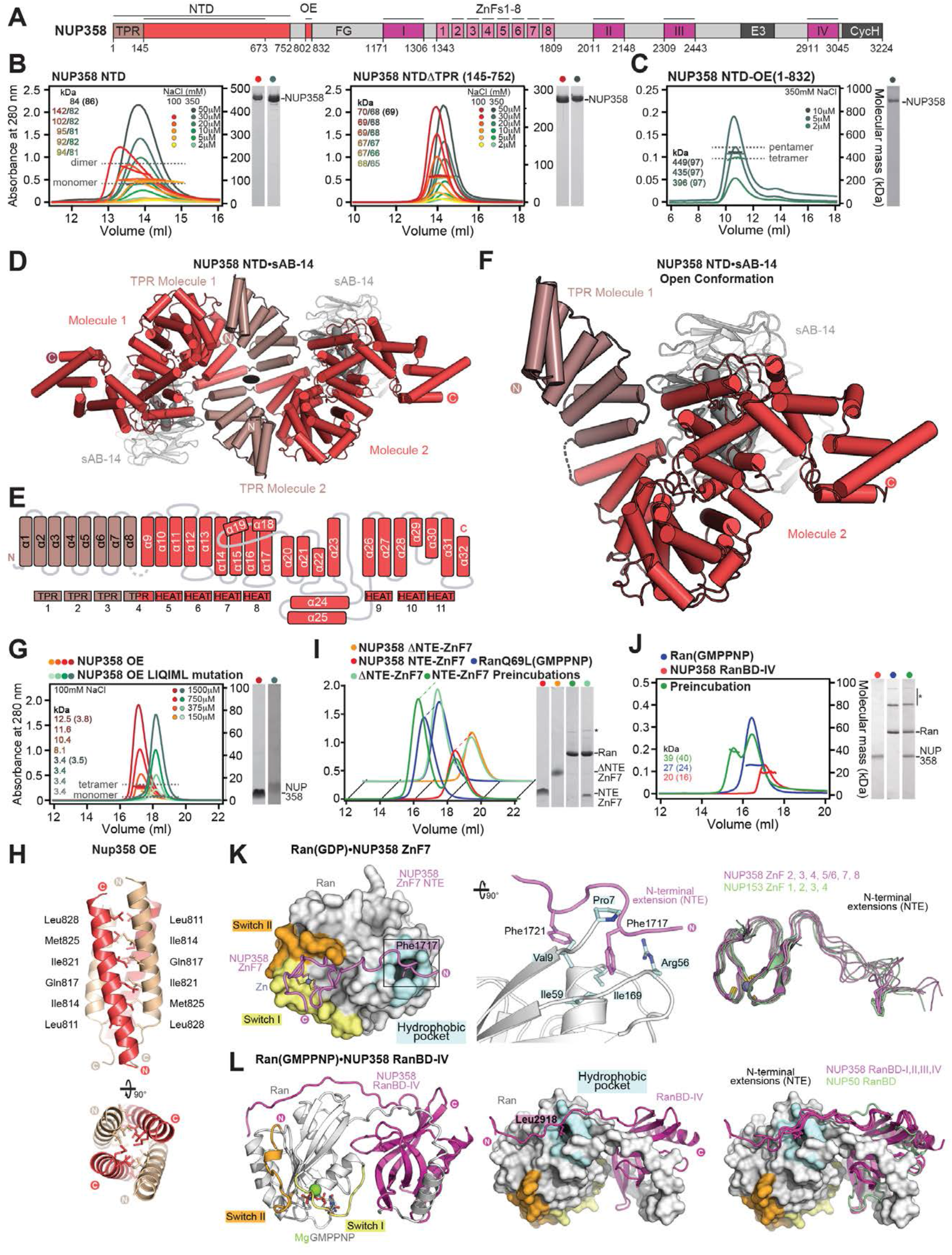
Structural analysis and biochemical characterization of NUP358. (**A**) Domain structure of NUP358 drawn as horizontal boxes with residue numbers indicated. Black lines indicate the boundaries of the crystallized fragments. (**B**, **C**) SEC-MALS analysis of the oligomeric behavior of (B) NUP358^NTD^ and NUP358^NTDΔTPR^, and (C) NUP358^NTD-OE^, performed at the indicated protein concentrations. Measured molecular masses are indicated, with theoretical masses in parentheses. SDS-PAGE gel strips of peak fractions are shown and visualized by Coomassie brilliant blue staining. (**D**) Cartoon representation of the NUP358^NTD^•sAB-14 co-crystal structure dyad of the P6_5_22 lattice, illustrating the NUP358^NTD^ dimer between symmetry-related molecules. (**E**) Schematic of the NUP358^NTD^ structure and structural motifs. (**F**) TPR of molecule 1 complements α-helical stacking of the N-terminal solenoid of the symmetry-related molecule 2, generating the open conformation of NUP358^NTD^. (**G**) SEC-MALS analysis of the oligomeric behavior of NUP358^OE^ and the NUP358^OE^ LIQIML mutant performed at the indicated protein concentrations. (**H**) Cartoon representation of the homo-tetrameric NUP358^OE^ crystal structure with hydrophobic core residues shown in ball-and-stick representation. (**I**) SEC interaction analysis of NUP358^ZnF7^ and NUP358^ZnF7ΔNTE^ binding to Ran(GTP). (**J**) SEC-MALS interaction analysis of NUP358^RanBD-IV^ binding to Ran(GMPPNP). (**K**) Co-crystal structure of NUP358^ZnF7^•Ran(GDP), shown in cartoon and surface representation (*left*). The inset indicates the location of the magnified and 90° rotated view of the Ran hydrophobic pocket in which NUP358 residue phenylalanine-1717 is buried (*middle*). Superposition of the six NUP358^ZnF^•Ran(GDP) and four NUP153^ZnF^•Ran(GDP) co-crystal structures with the Zn^2+^-coordinating cysteines and Ran-burying NTE hydrophobic residues shown as sticks (*right*). (**L**) Co-crystal structure of NUP358^RanBD-IV^•Ran(GMPPNP) with NUP358^RanBD-IV^ shown in cartoon and Ran(GMPPNP) shown in cartoon (*left*) or surface (*middle*) representation. Superposition of Ran(GMPPNP) bound to NUP358^RanBD-I^, NUP358^RanBD-II^, NUP358^RanBD-III^, NUP358^RanBD-IV^, and NUP50^RanBD^, illustrating the burial of NUP358^RanBD^ NTE conserved hydrophobic residues into the Ran hydrophobic pocket (*right*).

Despite extensive efforts, we were initially unable to generate well-diffracting NUP358^NTD^ crystals. However, we could obtain crystals of NUP358^145-673^, allowing for structure determination by single-wavelength anomalous dispersion (SAD) using seleno-L-methionine (SeMet)-labeled protein (tables S7 to S9). To aid the crystallization of the entire NUP358^NTD^, we generated high-affinity, synthetic antibody fragments (sABs) by phage display selection (*81*). By systematically screening the generated 62 sABs as crystallization chaperones, we identified a NUP358^NTD^•sAB-14 complex that crystallized, enabling *de novo* structure determination of the entire NUP358^NTD^ at a 3.95 Å resolution (table S10). To unambiguously assign the Nup358^NTD^ sequence register, we crystallized 17 additional SeMet-labeled methionine mutants (fig. S43 and table S11 and S12).

The asymmetric unit contained two copies of the NUP358^NTD^•sAB14 complex, in one of which the first three and a half TPR repeats are not resolved. The second copy forms extensive interactions with a symmetry related molecule (Fig. 4, D to F, and fig. S44). This NUP358 NTD dimer reveals two alternative TPR conformations, in which the TPR either forms a continuous N-terminal solenoid (open), or folds back, separating TPR4 and forming electrostatic interactions with HEAT repeats 5-7 of the N-terminal solenoid (closed) (Fig. 4F and fig. S44C). Toggling between these two states provides a molecular explanation for the salt-sensitive, concentration-dependent, dimerization behavior of NUP358^NTD^ (Fig. 4B). Because the open confirmation is identified in the intact NPC (see below), we focus our description on this state.

The open conformation of NUP358^NTD^ can be divided in three sections: an N-terminal α-helical solenoid composed of four TPRs and four <u>H</u>untingtin, <u>e</u>longation factor 3 (EF3), protein phosphatase 2<u>A</u> (PP2A), and the yeast kinase <u>T</u>OR1 (HEAT) repeats, a central α-helical wedge domain, and a short C-terminal α-helical solenoid formed by three HEAT repeats (fig. S44D). The N- and C-terminal TPR and HEAT repeats are capped by solvating helices. Inserted between helix α17 of the N-terminal solenoid and helix α20 of the wedge domain is a ∼50-residue loop that wraps around the convex face of the N-terminal solenoid. The N-terminal solenoid and the wedge domain form a large composite concave surface with a striking overall positive charge (figs. S45 and S46). The central wedge domain, composed of six α-helices, makes extensive hydrophobic contacts with the sides of the N- and C-terminal solenoids, generating a non-canonical S-shaped overall architecture (fig. S44D). Indeed, a Dali 3D search of the Protein Data Bank revealed that the NUP358^NTD^ architecture is novel (*82*).

Our biochemical analysis revealed that NUP358^NTD^ interacts weakly with NUP88^NTD^ and possesses RNA-binding activity, both of which were susceptible to elevated buffer salt concentration (figs. S47 and S48). By splitting NUP358^NTD^ into two fragments, NUP358^TPR^ and NUP358*^Δ^*^TPR^, we show that both halves of NUP358^NTD^ are necessary, yet insufficient, for either NUP88^NTD^ or RNA binding (figs. S47 and S48). To further map these interactions, we performed a saturating NUP358^NTD^ surface mutagenesis, screening 106 mutants for NUP88^NTD^ and ALPP^SSCR^ RNA binding (fig. S49). We found that positively charged residues in the concave surface compositely formed by the N-terminal solenoid and the wedge domain mediate binding to both NUP88^NTD^ and RNA. Mutations that abolished NUP88^NTD^-binding clustered exclusively on the N-terminal solenoid, whereas RNA disruption required additional mutations in the wedge domain. By systematically combining individual alanine substitutions, we identified a NUP358^NTD^ 2R5K mutation (K34A, K40A, R58A, K61A, R64A, K500A, and K502A) which abolished both interactions (fig. S50).

Next, we determined the crystal structure of NUP358^OE^ at 1.1 Å resolution (table S13). NUP358^OE^ is a small α-helical element that homo-tetramerizes to form an anti-parallel four-helix bundle (Fig. 4, G and H, and fig. S42A). The core of the α-helical bundle is lined with hydrophobic residues that coordinate oligomeric inter-helical packing, demonstrated by the monomeric form assumed by the NUP358^OE^ LIQIML mutant (L811A, I814A, Q817A, I821A, M825A, and L828A) (Fig. 4G and fig. S42, B to E). To validate our NUP358^OE^ structure, we tested the effect of introducing the LIQIML mutation into the larger NUP358^NTD-OE^ fragment. Whereas wildtype NUP358^NTD-OE^ formed higher order oligomeric species, the oligomerization profile of the LIQIML NUP358^NTD-OE^ mutant matched that of the OE-less NUP358^NTD^, presenting concentration-dependent dimerization in low salt buffer but persisting in a monodisperse monomeric state in high salt buffer (fig. S42, F and G).

Our data show that NUP358^NTD^ is composed of three distinct α-helical solenoids that interact in a novel manner, adopting a unique overall S-shaped architecture with a propensity to form domain-swapped homo-dimers. Connected to NUP358^NTD^ by a ∼50-residue linker is an oligomerization element that forms homo-tetramers/pentamers in solution. These dual modes of homo-oligomerization provide a plausible explanation for NUP358’s propensity to form phase separation, as observed during NPC assembly in *Drosophila melanogaster* oocytes (*83*).

### Ran interactions with human asymmetric nups

Nucleocytoplasmic transport depends on Karyopherin transport factors (Kaps) and a gradient in the state of the small GTPase Ran from the Ran(GDP)-high cytoplasm to the Ran(GTP)-high nucleus (*2, 7, 8, 10*). Import Kap•cargo complexes assembled in the cytoplasm enter the central transport channel of the NPC and are disassembled upon arrival in the nucleus by Kap binding to Ran(GTP), which triggers the release of the bound cargo. Conversely, Ran(GTP) is an obligate component of nuclear export Kap•cargo complexes, which are disassembled upon activation of Ran’s GTPase activity at the cytoplasmic face of the NPC. Multiple Ran binding sites are distributed among the asymmetric nups at the cytoplasmic and nuclear sides of the NPC in the form of distinct Ran-binding domains (RanBDs) and Zn^2+^-finger (ZnF) modules. On the cytoplasmic face, NUP358 contains four dispersed Ran-binding domains (RanBDs) and a central zinc finger domain (ZFD) with a tandem array of eight ZnFs (Fig. 4A) (*84*). On the nuclear side, NUP153 and NUP50 contain a central ZFD with four ZnFs and a solitary C-terminal RanBD, respectively (fig. S51) (*85, 86*).

Testing the Ran(GDP/GTP)-binding activity of all 17 domains by SEC-MALS, we confirmed that all domains bound to Ran as expected except for NUP358^ZnF1^ (Fig. 4, I and J, and figs. S51 to S55). Consistent with previous reports, the RanBDs of NUP358 and NUP50 only bound Ran(GTP), whereas the ZnFs in NUP358 and NUP153 bound Ran in both nucleotide states but showed a preference for Ran(GDP) (figs. S52 and S54) (*84, 87–89*). To clarify the molecular basis for the differential binding behaviors, we determined the co-crystal structures of all 16 domains bound to Ran in their preferred nucleotide-bound state at 1.8 Å-2.45 Å resolutions (tables S14 to S16 and figs. S51 and S55).

All eleven NUP358/NUP153 ZnFs form small, ∼36-residue spherical domains stabilized by a central Zn^2+^ ion coordinated by the sulfhydryl groups of four conserved cysteines (Fig. 4K and fig. S51). The primary interface between the ZnFs and Ran(GDP) is a flat hydrophobic composite surface formed by Ran’s switch-I and -II regions, as previously described (Fig. 4K and fig. S51) (*88, 89*). However, an additional ZnF-Ran interaction was also observed in all structures, which was formed between a phenylalanine or leucine in a ∼10-residue unstructured extension N-terminal to the ZnF module and a nucleotide state-independent hydrophobic pocket on the Ran surface (Fig. 4K and fig. S51). Removal of this extension from ZnFs eliminated Ran(GTP)-binding, showing that the secondary interaction we observed is responsible for the weak Ran(GTP)-binding capacity of NUP358/NUP153 ZnFs (fig. S52). Structural elucidation of the ZnF-Ran interfaces also showed that NUP358^ZnF1^ harbors sequence variations at important sites, explaining its weakened Ran(GDP) binding we observed biochemically (figs. S51 and S52). Furthermore, the NUP358 ZnF structures revealed an absence of conserved residues mediating canonical RNA binding, providing a molecular explanation for the surprising lack of RNA binding we observed biochemically contrary to previous reports (Fig. 3A and fig. S53) (*76, 90, 91*).

The crystal structures of the five NUP358/NUP50 RanBD•Ran(GMPPNP) complexes show that they have the same architecture as previously observed for the NUP358^RanBD-I^•Ran complex (Fig. 4L and fig. S55 and table S16) (*73*). In addition to the previously described interactions between the pleckstrin-homology-fold of RanBDs and Ran(GMPPNP), we observed an extensive positively charged surface patch in all five RanBDs that sequesters Ran’s acidic C-terminal DEDDDL motif and is necessary for tight binding (*92*). Ran’s C-terminal α-helix binds to a conserved groove on the surface of all five RanBDs, but shows some plasticity in the exact molecular interactions (fig. S55E). Interestingly, the same hydrophobic Ran pocket bound by ZnF N-terminal extensions (NTEs) in the ZnF•Ran(GDP) complex structures is occupied by a proline or leucine residue in the ∼20-residue NTEs of all four NUP358 RanBDs, but not the NUP50 RanBD (fig. S55C).

Together, our data establish that the human CF and nuclear basket nups NUP358, NUP153, and NUP50 harbor a total of 16 distinct Ran binding sites that, considering their stoichiometry in the NPC, could together recruit up to several hundred Ran molecules. Considering the substantial size difference between metazoan and *S. cerevisiae* cells, it is conceivable that additional Ran binding sites provided by the metazoan-specific asymmetric nups NUP358 and NUP153 help ensure high enough Ran-concentrations in the NPC vicinity to enable nucleocytoplasmic transport, as has been previously suggested (*89*).

### Docking of NUP358^NTD^ into the cytoplasmic unassigned density cluster I

Nup358 is known to reside on the cytoplasmic face of the NPC. However, due to lack of sufficient structure information, its exact location in the NPC could so far only be inferred from the differential absence of unassigned density in an ∼38 Å cryo-ET map of the human NPC that was derived from NUP358-depleted cells (*44*). In the accompanying manuscript, the quantitative docking of residue-level resolution structures into the symmetric core of a ∼12 Å cryo-ET map of the intact human NPC led to a novel assignment of 16 copies of the symmetric nups NUP205 and NUP93 in the cytoplasmic outer ring, as well as the identification of two clusters (I and II) of unassigned density (Fig. 5A), of which the first corresponds to the previously observed Nup358-dependent density (*42, 46*). Because of the large size and unique fold of our newly determined NUP358^NTD^ crystal structure, we sought to directly determine its position in the intact human NPC (Fig. 5A).

**Fig. 5.**
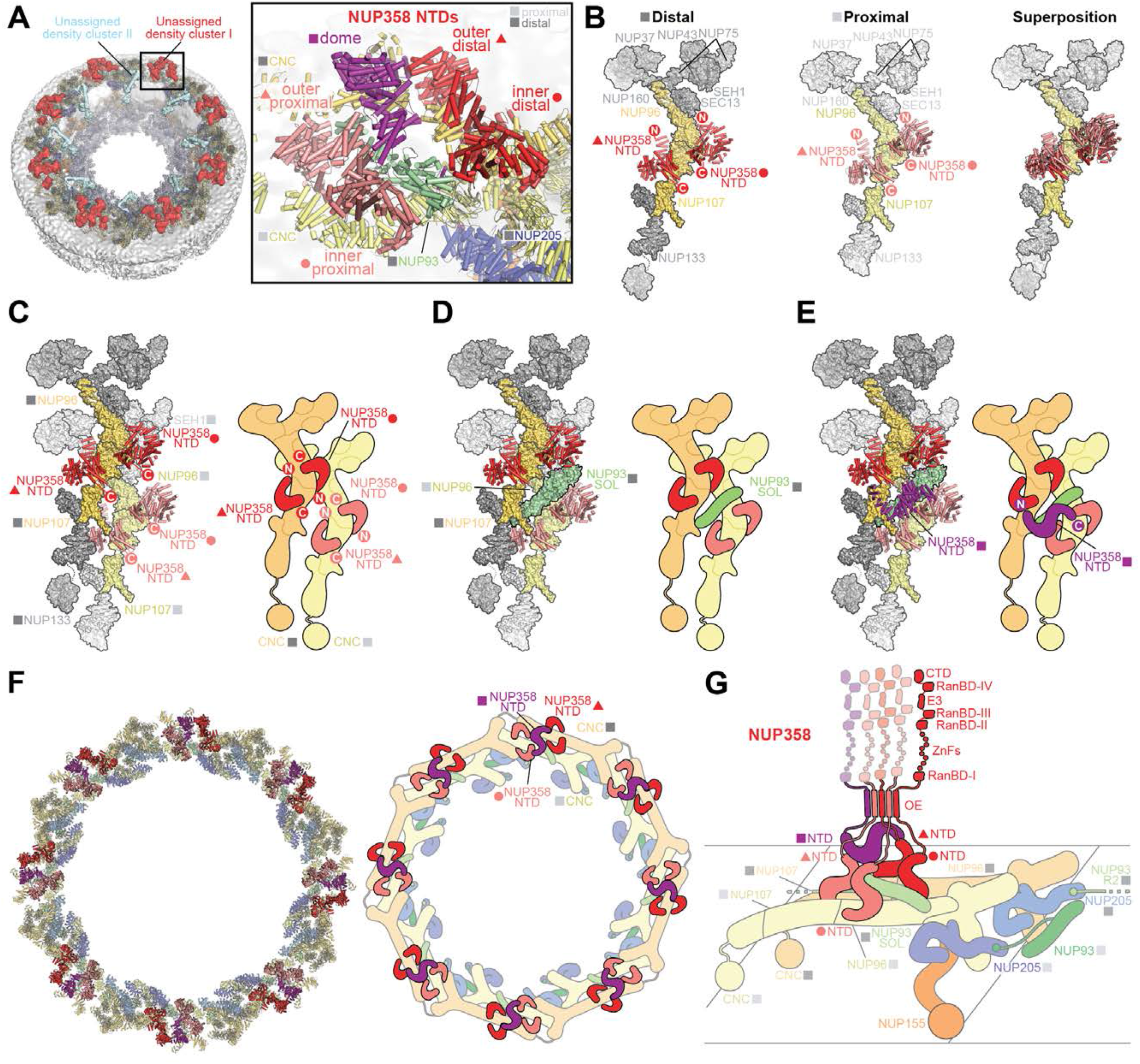
Docking of NUP358^NTD^ on the cytoplasmic face of the NPC. (**A**) Overview of the NPC cytoplasmic face with isosurface rendering of unexplained density clusters I (red) and II (cyan) of the ∼12 Å cryo-ET map of the intact human NPC. The inset indicates the location of the magnified view showing cartoon representations of five copies of NUP358^NTD^ docked in unassigned density cluster I of a single spoke. An inner and outer copy of the NUP358^NTD^ interface with each distal and proximal stalk of the Y-shaped CNCs is shown. The distal NUP93^SOL^ is collocated at the center of the NUP358^NTD^ quartet and the fifth dome NUP358^NTD^ copy lies on top. (**B**) Comparison of the binding of two NUP358^NTD^ copies to distal and proximal CNCs. NUP358^NTD^ copies are rendered as cartoons and CNCs as surfaces. (**C**-**E**) Architecture of the pentameric NUP358^NTD^ bundle attachment site on a cytoplasmic outer ring spoke with cartoon- and surface-represented structures (*left*) and schematics (*right*), sequentially illustrating the placement of (C) four copies of NUP358^NTD^ around NUP96 and NUP107 interfaces on the stalks of tandem-arranged Y-shaped CNCs, (D) a distal copy of NUP93^SOL^ collocated at the center of the NUP358^NTD^ bundle, interfacing with both proximal and distal CNC stalks, and (E) the NUP358^NTD^ dome copy interfacing with the stalk-attached NUP358^NTD^ quartet beneath. (**F**) Overview of the entire cytoplasmic face of the NPC in cartoon representation and as schematic, illustrating the distribution of 40 NUP358^NTD^ copies anchored as pentameric bundles across the eight NPC spokes. (**G**) Schematic of NUP358 attachment to the cytoplasmic outer ring spoke. Four copies of NUP358^NTD^ wrapped around the stalks of the tandem arranged Y-shaped CNCs and the fifth dome NUP358^NTD^ are linked together by interactions between oligomerization elements (OEs). Anchored by NUP358^NTD^, the rest of NUP358 domains are linked by unstructured linker sequences and are expected to freely project from the cytoplasmic face of the NPC. Distal and proximal positions are labeled according to the legend in (A).

In our docking analysis, we calculated correlations between a new ∼12 Å cryo-ET map of the intact human NPC provided by the Beck group and one million randomly placed and locally fit-optimized resolution-matched densities simulated from the open and closed conformations of the NUP358^NTD^ crystal structure (*46*). Unlike for the closed NUP358^NTD^ conformation, docking scores for five placements of the open NUP358^NTD^ conformation segregated to high confidence of placement in the previously unassigned density cluster I, leaving no unexplained density (fig. S56). We found four copies of NUP358^NTD^ to be interfaced with the α-helical solenoid folds of the CNC components NUP96, NUP107, and the distal copy of NUP93^SOL^, wrapping around the stalks of the tandem arranged Y-shaped CNCs in pairs, at equivalent distal and proximal positions (Fig. 5, B and C). As identified in the docking analysis of the symmetric core reported in the accompanying manuscript, the distal NUP93^SOL^ bisects the stalks of the tandem-arranged Y-shaped CNCs by interfacing with the distal NUP107 and the proximal NUP96 α-helical solenoids, cloistered between the four NUP358^NTD^ copies that wrap around the CNC stalks (Fig. 5D) (*42*). Lastly, the fifth NUP358^NTD^ that we refer to as the dome copy was placed above the other four NUP358^NTD^ copies and the distal NUP93^SOL^, with its N- and C-termini oriented towards the C-termini of the outer distal and inner proximal copies of NUP358^NTD^, respectively (Fig. 5E). Though unexpected, the placement of the dome NUP358^NTD^ was the second most confident docking solution into the ∼12 Å cryo-ET map of the intact human NPC, as well as into a previously reported ∼23 Å cryo-ET map of the intact human NPC (*44*), with which we repeated the unbiased docking as further reference and identified all the same placements, albeit with significantly lower confidence (fig. S57). Finally, we successfully placed the composite structure of the entire cytoplasmic outer ring protomer, including all five NUP358^NTD^ copies, into an anisotropic ∼7 Å region of a composite single particle cryo-EM map of the *X. laevis* NPC cytoplasmic outer ring protomer (figs. S58 and S59) (*45*). This analysis enabled by our novel residue-level resolution crystal structures allowed us to rectify an incorrect assignment of density previously referred to as the ‘U-domain’ and the ‘bridge domain’ of the ‘NUP358 complex’ to the dome copy of NUP358^NTD^ and the distal copy of NUP93^SOL^, respectively. Considering the unusual triple α-helical solenoid architecture of NUP358^NTD^, it is unsurprising that *de novo* atomic model building and chain tracing in a featureless distorted map led to incorrect density interpretation (fig. S59).

The arrangement of five NUP358^NTD^ copies in each spoke places their C-termini in proximity of each other, projecting the rest of the NUP358 domains towards the cytoplasm. Consequently, the oligomerization domains of the five NUP358^NTD^ copies are constrained to form a homomeric assembly within the same spoke. (Fig. 5, F and G). The oligomerization of NUP358 observed in the NUP358^OE^ crystal structure and SEC-MALS analysis would boost the avidity of NUP358 attachment to the cytoplasmic face of the NPC (Figs. 4, G and H, and 5G).

### NUP358 is dispensable for NPC integrity during interphase

Our quantitative docking showed that NUP358^NTD^ is the primary attachment point for NUP358 at the cytoplasmic face of the NPC. To validate this result physiologically, we sought to determine the subcellular localization of structure-guided NUP358 fragments in intact cells. To prevent default localization of ectopically expressed fragments at the nuclear envelope because of homo-oligomerization with the Nup358^OE^ of the endogenous proteins, we generated an inducible NUP358 knockout cell line, in which an N-terminal auxin-inducible degron (AID) tag was inserted into both genomic *NUP358* loci (*AID*::*NUP358* HCT116) (fig. S60). We selected HCT116 cells for their amenability to subcellular imaging and stable diploid karyotype (*93*). Addition of 1 mM auxin resulted in the rapid, selective, and complete degradation of endogenous NUP358 within three hours, confirmed by the loss of immunohistochemical nuclear envelope rim staining and western blot analysis of cellular NUP358 protein levels (Fig. 6A and figs. S60 and S61).

**Fig. 6.**
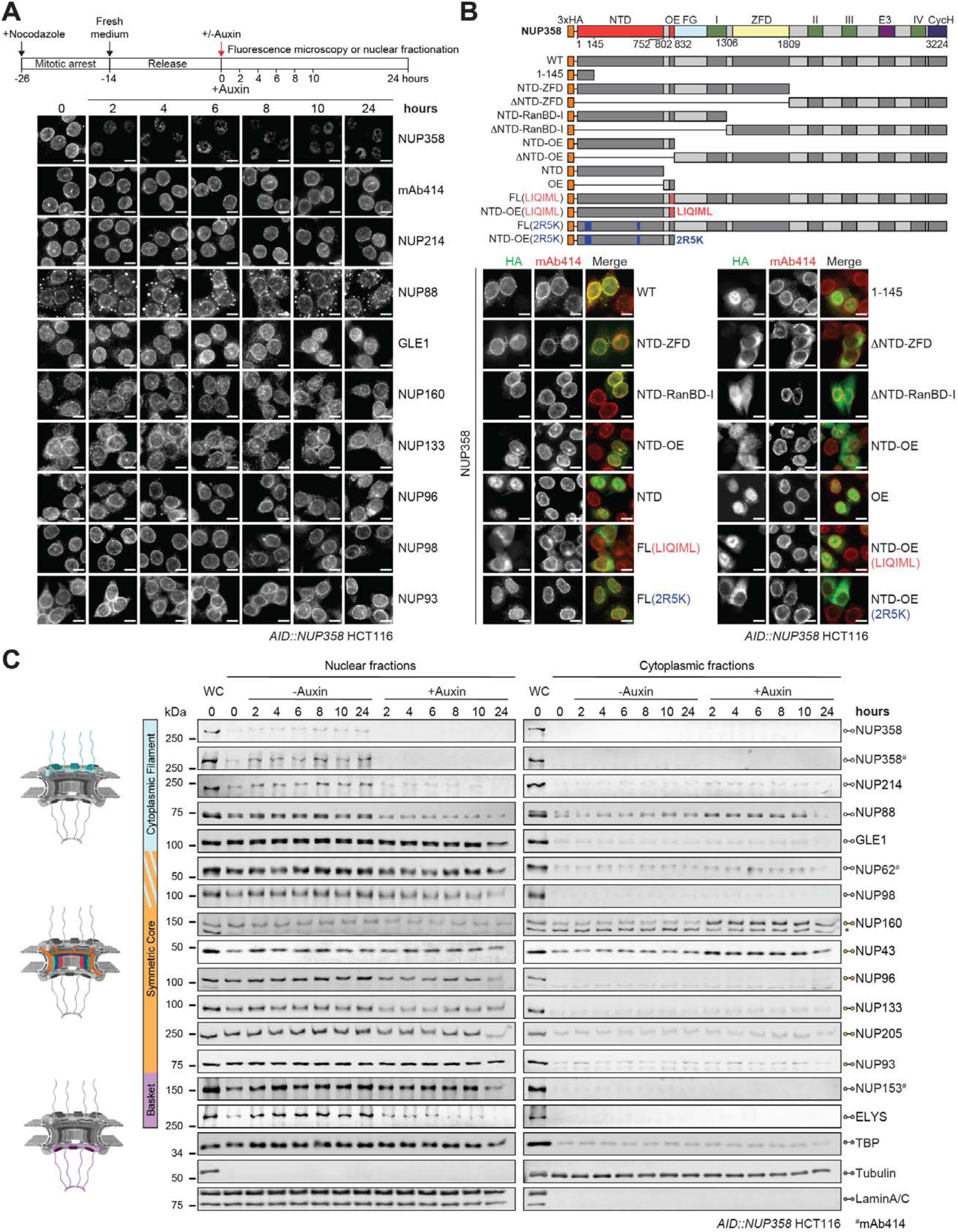
NUP358 is dispensable for NPC integrity in interphase cells. (**A**) Subcellular localization analysis by immunofluorescent staining of endogenous nups in synchronized *AID*::*NUP358* HCT116 cells at the indicated time points following auxin-induced NUP358 depletion. Experimental timeline is shown (*above*). (**B**) Subcellular localization analysis of N-terminally 3×HA-tagged NUP358 variants in *AID*::*NUP358* HCT116 cells by immunofluorescence microscopy. mAb414 staining served as reference for the nuclear envelope rim location. The domain structure of the transfected NUP358 variants is shown (*above*). (**C**) Western blot analysis of nup levels in cytoplasmic and nuclear fractions of synchronized *AID*::*NUP358* HCT116 cells upon auxin-induced NUP358 depletion, according to the experimental timeline outlined in (A). Control cells were not treated with auxin. A whole cell extract (WC) control is included in the first lane of each blot. Colored regions in the schematics indicate nup locations within the NPC (*left*). Asterisk indicates a non-specific band. mAb414 antibody recognizes nups containing FG repeats including NUP62, NUP153, NUP214, and NUP358. Scale bars are 10 μm.

To identify the minimal NUP358 region necessary and sufficient for nuclear envelope targeting, we generated a systematic series of HA-tagged N- and C-terminal fragments, splitting the protein into two pieces after the NTD, OE, RanBD-I, or ZFD, and determined their subcellular localization by immunofluorescence microscopy. Consistent with our previous results, NUP358 targeting to the nuclear envelope in the absence of auxin required both NTD and OE regions, whereas all other domains were dispensable (Fig. 6B and fig. S62). When NUP358^NTD^ or NUP358^OE^ were tested in isolation, neither was found to be sufficient, with both domains exhibiting strong nuclear staining (Fig. 6B). Our docking analysis revealed that NUP358^NTD^ contacts the CNC via the same concave surface that we had found to mediate interactions with NUP88^NTD^ and RNA. We therefore asked whether introduction of the NUP358 2R5K mutation that abolishes NUP88^NTD^ and RNA binding and significantly alters local surface charge would also influence NUP358 subcellular localization. Indeed, the 2R5K mutation either abolished or severely reduced nuclear envelope rim staining when introduced into HA-NUP358^NTD-OE^ or HA-NUP358^FL^, respectively (Fig. 6B). Analogously, we tested whether NUP358 oligomerization is required for localization to the nuclear envelope. Indeed, introduction of the LIQIML mutation, which abolishes NUP358^NTD-OE^ oligomerization *in vitro*, eliminated nuclear envelope rim staining of both HA-NUP358^NTD-OE^ and HA-NUP358^FL^ (Fig. 6B). To rule out confounding effects from ectopically expressed HA-NUP358 variants hetero-oligomerizing with endogenous NUP358, we repeated the fluorescence microscopy analysis after auxin-induced depletion of endogenous NUP358 and obtained identical results (fig. S63).

The previous ∼38 Å cryo-ET map of the human NPC from purified nuclear envelopes of cells in which NUP358 had been knocked down by *RNAi* showed loss of the distal cytoplasmic CNC ring and potentially other CF nups in the absence of NUP358, leading to the conclusion that NUP358 is required for the integrity of the interphase NPC (*44*). To determine whether the architectural stability of the interphase NPC depends on NUP358, we sought to analyze the effect of auxin-induced NUP358 depletion on the subcellular localization of eight nups representative of all NPC sub-complexes by immunofluorescence microscopy (Fig. 6A). Because previous studies had shown that NUP358 depletion results in cell cycle arrest at the G2/M transition (*94*), we first monitored the levels of cell cycle markers cyclin-B1, cyclin-D1, cyclin-E, and serine 10-phosphorylated histone H3 in uninduced, nocodazole-synchronized *AID*::*NUP358* HCT116 cells, thus determining an approximate cell cycle length of ∼14-hour consistent with previous reports for HCT cells (fig. S64) (*95*). With this knowledge, we then induced NUP358-degradation in nocodazole-synchronized cells and imaged nups at various timepoints prior to cells entering mitosis with the exception of the last timepoint at 24 hours. Although NUP358 nuclear envelope rim staining was rapidly lost two hours after induction of degradation and remained absent throughout the remaining timepoints, all eight representative nups continued to display robust nuclear envelope rim staining throughout the course of the experiment (Fig. 6A). This suggests that NPC integrity is not dependent on NUP358 attachment to the NPC and also demonstrates the specificity of the auxin-induced NUP358 knockout in our AID cell line. To reconcile the apparent conflict between our results and the prominent nup loss from the NPC in the absence of NUP358 observed by cryo-ET, we investigated whether NUP358 depletion led to release of nups from the nuclear fraction during cellular fractionation. Indeed, we observed an auxin-dependent leakage of NUP214, NUP88, and NUP160 from the nuclear to the cytoplasmic fraction (Fig. 6C). Curiously, we also consistently observed a marked reduction of the nuclear basket nup ELYS in the nuclear fraction upon NUP358 depletion (Fig. 6C).

Taken together, these data confirm the quantitative docking of NUP358^NTD^, validate the physiological relevance of NUP358^OE^-mediated bundling, and establish that NUP358 is dispensable for the architectural integrity of the assembled interphase NPC, although its depletion made the structural integrity of the cytoplasmic face of the NPC susceptible to the biochemical stresses inherent to cell fractionation. Future studies need to establish the extent of NUP358’s role in the formation of the double CNC-ring architecture during NPC assembly.

### NUP358 plays a general role in translation of exported mRNAs

Export of mRNA from the nucleus to the cytoplasm is an essential step in the expression of eukaryotic proteins. mRNA maturation and export are a result of complex multistep processes that involve 5’-capping, splicing, 3’-polyadenylation, formation of export-competent mRNPs, and translocation across the NPC along with subsequent remodeling at the cytoplasmic face (Fig. 7A) (*43*). Our biochemical analysis revealed that NUP358 has multiple RNA-binding domains distributed throughout the protein, suggesting a potential role in RNA export and mRNP remodeling (Fig. 3B). The docking of five NUP358^NTD^ copies in the intact human NPC revealed occlusion of the RNA/NUP88^NTD^-binding surfaces of the dome and inner distal NUP358 copies by the CNC stalk, but exposure of the remaining copies’ binding sites (fig. S65). Thus, some NUP358^NTD^ copies could potentially be simultaneously attached to the NPC and dynamically interacting with RNA/NUP88^NTD^.

**Fig. 7.**
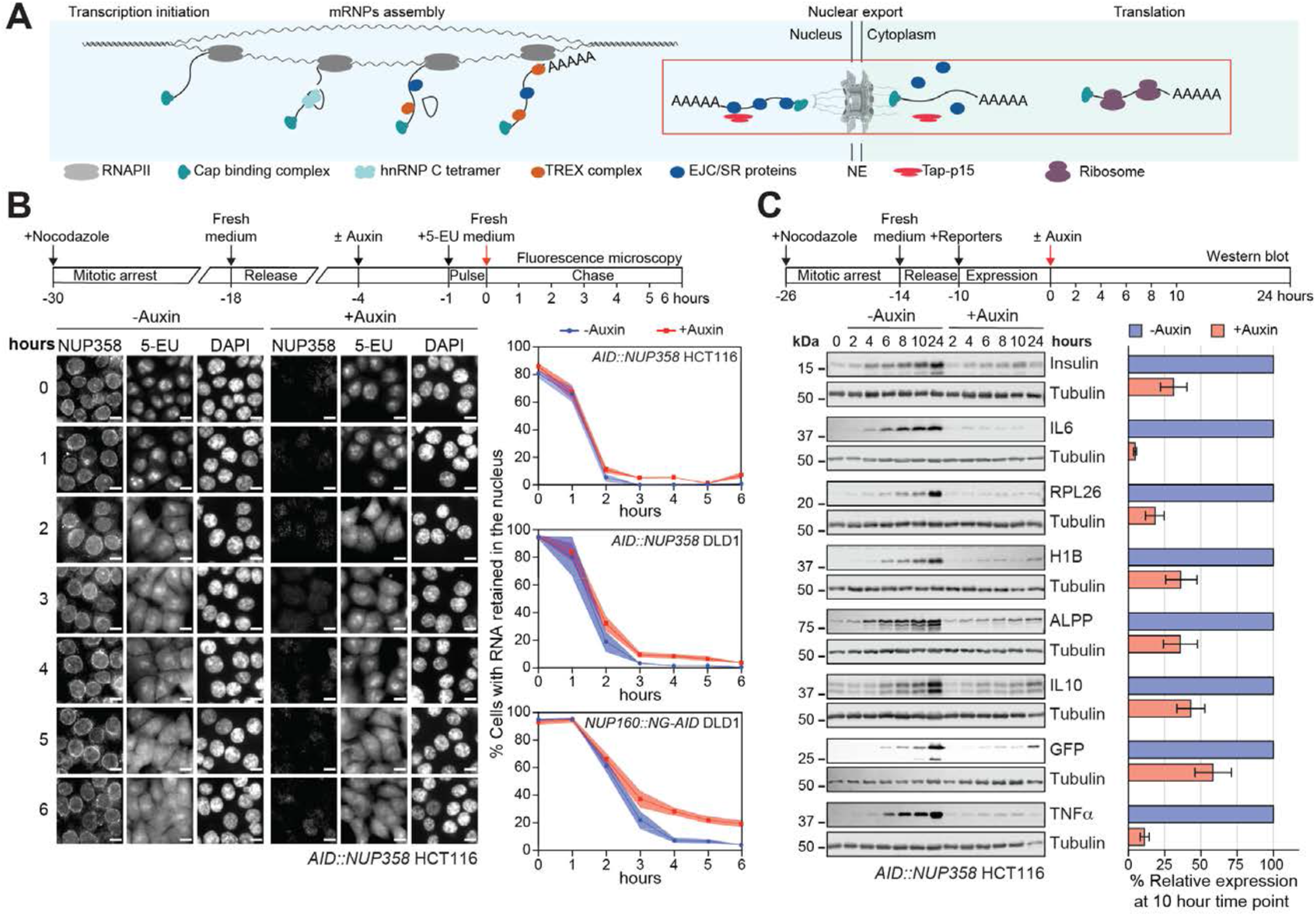
NUP358 plays a general role in translation of exported mRNA. (**A**) Schematic illustrating the life cycle of mRNA, from transcription, maturation of an export-competent messenger ribonucleoprotein particle (mRNP) in the nucleus, nuclear export and mRNP remodeling at the cytoplasmic face of the NPC, to translation in the cytoplasm. Steps associated with the NPC are highlighted by a red box. (**B**) Time-resolved analysis of RNA nuclear retention in synchronized *AID*::*NUP358* HCT116 cells upon auxin-induced NUP358 depletion. Control cells were not treated with auxin. Nascent RNA was metabolically pulse-labeled with 5-ethynyl uridine (5-EU) and its subcellular localization determined by Alexa Fluor 594 azide conjugation and fluorescence microscopy at the indicated time points. Experimental timeline is shown (*above*). Representative images (*left*) and quantitation of the proportion of cells (n>200) with nuclear RNA retention at each timepoint are shown (*right*). Experiments were performed in triplicate, and mean values with the respective standard error are indicated by squares and shaded areas, respectively. Also shown is the quantitation of the same analysis repeated in unsynchronized *AID*::*NUP358* DLD1 and *NUP160*::*NG-AID* DLD1 cells. Scale bars are 10 μm. (**C**) Time-resolved Western blot analysis of the expression of C-terminally 3×FLAG-tagged reporter proteins in synchronized *AID*::*NUP358* HCT116 cells upon auxin-induced NUP358 depletion. Experimental timeline is shown (*above*). Control cells were not treated with auxin. Quantitated reporter expression in auxin-treated cells was normalized to expression in control cells, at the 10-hour timepoint. Experiments were performed in triplicate, and the mean with the associated standard error reported.

Previous studies had found that efficient translation of secretory proteins requires NUP358 binding to the ∼63 nucleotide GC-rich signal sequence coding region (SSCR) of mRNAs encoding secretory proteins (*76*). NUP358 knockdown by lentivirally delivered short hairpin (sh) RNAs was shown to prevent the translation of various secretory protein reporters, but had no effect on the distribution of mRNA in the cell (*76*). These RNA interference (*RNAi*) experiments involved extended incubation periods and achieved only a partial NUP358 knockdown, potentially allowing secondary phenotypes to emerge from prolonged NPC disruption, non-NPC related NUP358 effects, or defective postmitotic NPC re-assembly. With the ability to rapidly deplete NUP358 in our AID cell line, we sought to examine whether NUP358 is directly involved in mRNA export or mRNPs remodeling events by monitoring the subcellular distribution of pulse-labeled, newly synthesized RNA upon loss of Nup358. Specifically, synchronized *AID*::*NUP358* HCT116 cells were metabolically labeled with 5-ethynyl uridine (5-EU) for one hour starting three hours after auxin treatment, followed by the start of the chase with fresh media. *In situ* labeled RNA was visualized by fluorescence microscopy at hourly intervals during the chase for up to 6 hours (Fig. 7B). At early time points, 90-100% of cells displayed a strong nuclear 5-EU-labeled RNA signal that decreased over time with a concomitant increase in the cytoplasmic signal, indicative of RNA being exported. After six hours, only ∼5% of NUP358-depleted cells exhibited nuclear retention of labeled RNA, compared to <1% of control cells (Fig. 7B). Because of this rather subtle effect, we sought to confirm the result in *AID*::*NUP358* DLD1 cells, another cell line with inducible NUP358-depletion (Fig. 7B and figs. S66 and S67). Because DLD1 cells do not respond to nocodazole, the assay was performed in non-synchronized cells with a fraction of the cells undergoing mitosis during the 7-hour pulse-chase experiment. Similar to the *AID*::*NUP358* HCT116 cell results, only ∼3% of NUP358-depleted DLD1 cells exhibited nuclear RNA retention after a 6-hour chase (Fig. 7B and fig. S68). By contrast, nuclear RNA retention was present in ∼20% of *NUP160*::*NG-AID* DLD1 cells after depletion of NUP160, whose knockout causes RNA retention in *S. cerevisiae*, demonstrating the principle suitability of our experimental approach (Fig. 7B and fig. S68) (*96*). Our observed intact mRNA export despite NUP358 loss is also in agreement with previous reports of lentiviral NUP358 *RNAi* knockdown in U2OS cells not resulting in elevated nuclear retention of either poly(A)^+^ RNA or the stable *fushi tarazu* (*ftz*) reporter mRNA, four days post infection (*76*).

Next, we analyzed the fate of the genetic message downstream of mRNA export by examining the dependence of cellular protein expression on NUP358 in *AID*::*NUP358* HCT116 cells using eight different reporter constructs. Synchronized cells were transfected with C-terminally FLAG-tagged reporter constructs prior to NUP358 depletion, and the amount of reporter in whole cell extracts was determined by Western blot analysis (Fig. 7C and fig. S69). First, we focused our analysis on representative secretory protein reporter constructs including insulin, anti-inflammatory cytokine interleukin-10 (IL-10), pro-inflammatory cytokine IL-6, tumor necrosis factor alpha (TNFα), and membrane-bound ALPP. NUP358 depletion significantly reduced the cellular levels of all five reporters. To test whether NUP358 depletion specifically affects the expression of secretory proteins, we carried out the same analysis with non-secretory protein reporters. Contrary to the previous observation of secretory protein-effect specificity, we found that NUP358 depletion also significantly reduced the cellular levels of non-secretory protein reporters, including large ribosomal subunit protein 26 (RPL26), green fluorescence protein (GFP), and histone 1B (H1B) (Fig. 7C and fig. S69).

In summary, our data confirm that NUP358 depletion does not result in a marked nuclear RNA accumulation, but nevertheless affects the efficient translation of secreted and membrane-bound proteins, as previously proposed (*76*). However, our findings also demonstrate that the observed translational defect is not restricted to secretory proteins, suggesting a more general role of NUP358 in mRNP remodeling events that occur at the cytoplasmic face of the NPC after mRNA export.

### Characterization of NUP358 harboring ANE1 mutations

Acute necrotizing encephalopathy (ANE) is an autoimmune disease in which previously healthy children experience a cytokine storm in response to common viral infections, causing massive brain inflammation and rapid deterioration from seizures to coma and that can ultimately be fatal (*97*). ANE1, the familial and recurring form of ANE, has been associated with four distinct NUP358 mutations: T585M, T653I, I656V, and W681C (*97, 98*). All four ANE1 mutations map to the C-terminal α-helical solenoid of NUP358^NTD^ (Fig. 8A). Apart from T585, which is exposed on the surface, the ANE1 mutations locate to helices α28 and α30 in the closely packed hydrophobic core (Fig. 8B). To elucidate whether the ANE1 mutations induce conformational changes, we determined co-crystal structures of NUP358^NTD^ harboring the individual ANE1 mutations T585M, T653I, and I656V in complex with sAB-14. Consistent with the conservative nature of the ANE1 substitutions, the structures revealed no substantial structural changes, with root-mean square deviation (RMSD) calculated over 746 C_α_ atoms of ∼0.5 Å (Fig. 8B and table S10). Next, we assessed whether individual ANE1 mutations affected the known cellular functions of NUP358 but were unable to detect differences in nuclear envelope rim staining or binding to NUP88^NTD^ and ALPP SSCR RNA (Fig. 8, C and D, and fig. S70). Because we previously found that GLE1 mutations associated with various neurodegenerative diseases did not affect its function but rather decreased its thermostability by up to ∼10 °C (*63*), we tested the effect of ANE1 mutations on NUP358^NTD^ thermostability. Indeed, we found that W681C, T653I, and I656V mutants exhibited marked decreases in thermostability compared to the wildtype protein, with the *in vitro* thermostability of mutants reduced below body temperature (Fig. 8E and fig. S71). Notably, we observed that binding to sAB-14 increased the thermostability of all three ANE1 mutants beyond wildtype levels (fig. S72).

**Fig. 8.**
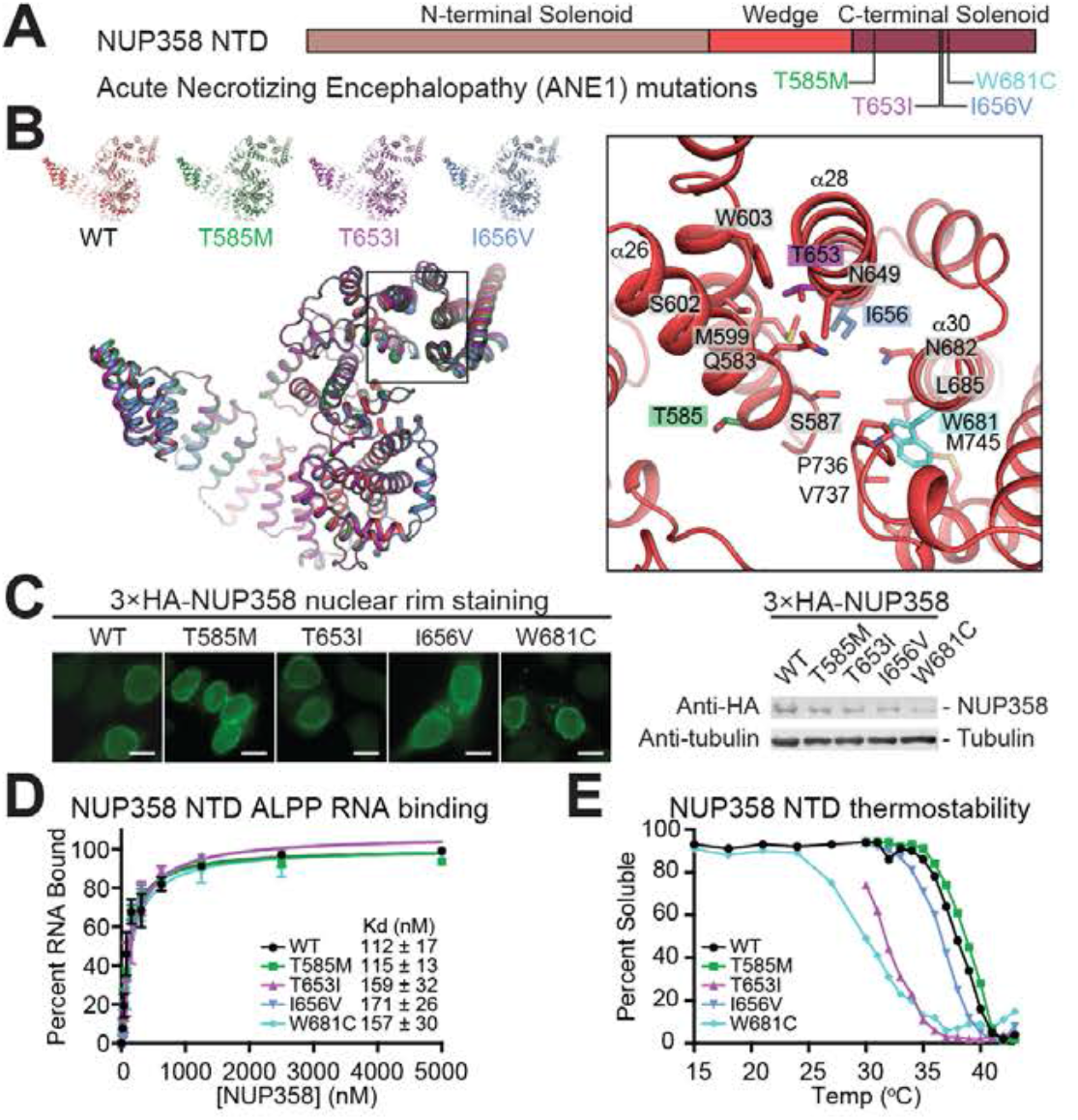
ANE1 mutations decrease NUP358^NTD^ thermostability. (**A**) NUP358^NTD^ domain architecture with location of ANE1 mutations indicated. (**B**) Crystal structures of wildtype NUP358^NTD^ and three distinct ANE1 mutants (T585M, T653I, or I656V) are shown individually and superposed in cartoon representation. A magnified view of the inset region shows the local environment of the four ANE1-associated mutations (*right*). (**C**) Subcellular localization analysis by immunofluorescence microscopy of N-terminally 3×HA-tagged NUP358 ANE1 mutants in *AID*::*NUP358* HCT116 cells. Scale bars are 10 μm. Expression of NUP358 mutants was verified by western blot of whole cell extracts. (**D**) Binding of wildtype and ANE1 mutant NUP358^NTD^ to ALPP^SSCR^ RNA measured by EMSA. Shown is the quantitation of triplicate experiments with mean values and standard errors. Apparent dissociation constants (*K_D_*) were estimated by fitting single-site binding models to the data. (**E**) Thermostability of wildtype and ANE1 mutant purified NUP358^NTD^ measured by pelleting assay.

Together, our results indicate that ANE1 mutations neither directly disturb the overall fold nor affect the known cellular functions of NUP358. Our observation of a substantially reduced thermostability of NUP358 ANE1 mutants is interesting, considering that the sudden onset of symptoms appears to require a fever-inducing trigger such as a viral infection (*97*). Future studies will need to systematically assess whether ANE1 mutations affect unknown cellular functions of NUP358.

### Structural and biochemical analysis of NUP88^NTD^**•**NUP98^APD^**•**NUP214^TAIL^

Besides NUP358, the NUP88^NTD^•NUP98^APD^•NUP214^TAIL^ complex had up to now been another CF component for which atomic level structural information had remained unavailable. Through extensive screening of crystallization fragments and conditions, we were able to solve the structure of the hetero-dimeric NUP88^NTD^•NUP98^APD^ at 2.0 Å resolution (table S17). NUP88^NTD^ forms a seven-bladed β-propeller domain, which is extensively decorated by a series of extended loops and insertions, with NUP98^APD^ bound to the fourth and fifth blades (Fig. 9, A and B). NUP88^NTD^ and NUP98^APD^ form a bipartite interaction through a cross-handshake exchange of loops. NUP98^APD^ extends the K/R loop to present residue K814 to hydrogen bond with the NUP88^NTD^ 4D5A loop backbone carbonyls of N273 and G308 (Fig. 9C). The tight K814 coordination forms a robust interaction, which is dependent on the conformation of NUP88^NTD^ 4D5A loop. Indeed, severing requires an aggressive combination mutant of four 4D5A loop residues (E275A, M299A, N306A, and Y307A; EMNY) (Fig. 9D and fig. S73). The FGL loop of NUP88^NTD^ reaches over to bind into a hydrophobic groove of NUP98^APD^ (Fig. 9E). Removal of the FGL loop leads to weakened interactions between NUP88^NTD^ and NUP98^APD^ (Fig. 9F and fig. S73). Despite low sequence homology, the overall architecture of the NUP88^NTD^•NUP98^APD^ complex is conserved from fungi to humans, although the orientation of NUP98^APD^ relative to NUP88^NTD^ varies between the co-crystal structures of human and fungal orthologs by as much as ∼20° (figs. S74 to S76) (*11, 59*).

**Fig. 9.**
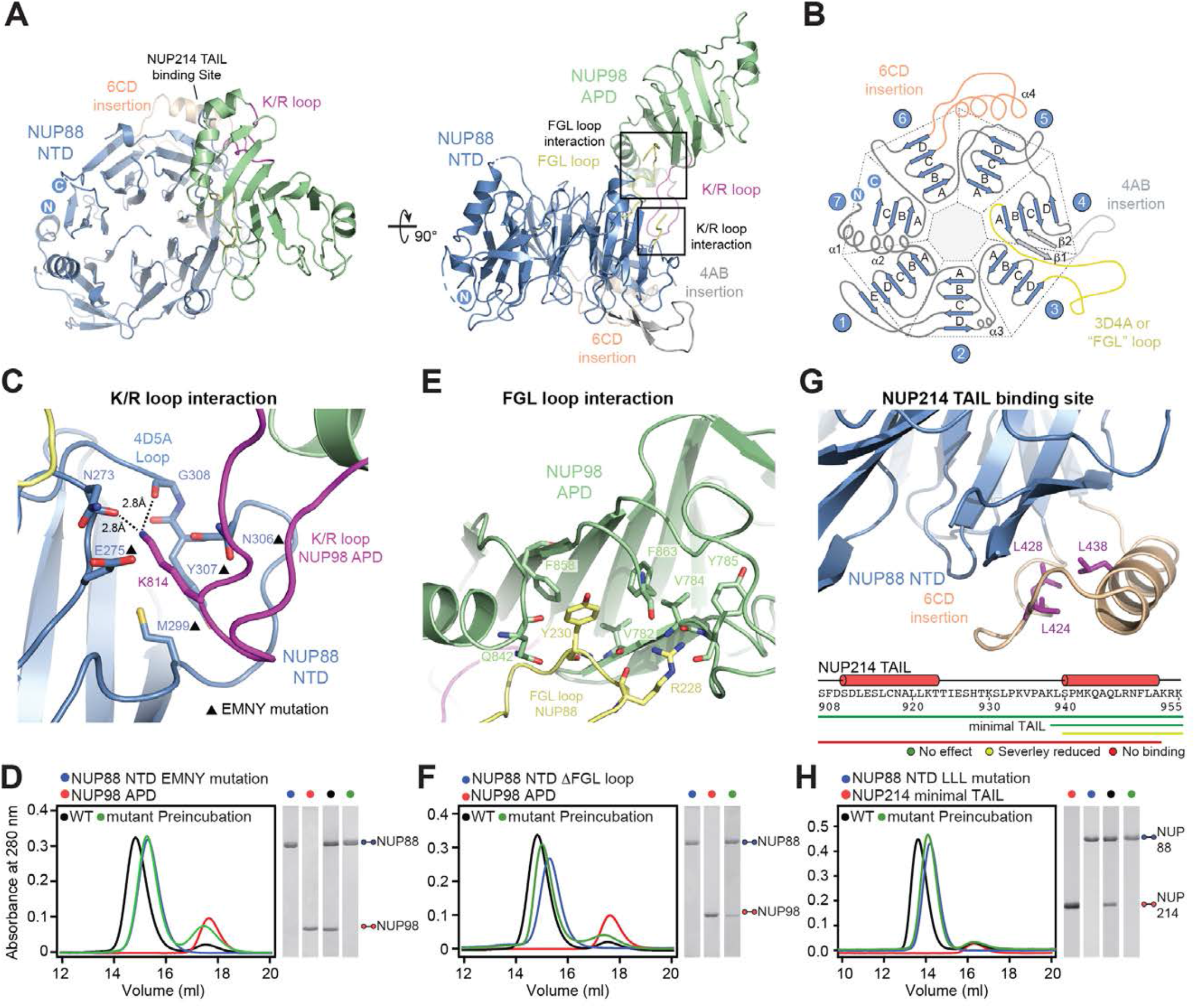
Structure of the NUP88^NTD^•NUP98^APD^ heterodimer. (**A**) Cartoon representation of the NUP88^NTD^•NUP98^APD^ crystal structure shown in different orientations. (**B**) Schematic of the seven-bladed NUP88^NTD^ β-propeller domain, indicating prominent loop insertions. (**C**) Magnified view of the NUP98 K/R loop interaction with NUP88^NTD^. Black triangles indicate alanine substitutions in the EMNY mutant. (**D**) SEC interaction analysis of NUP98^APD^ with NUP88^NTD^ EMNY mutant. (**E**) Magnified view of NUP88^NTD^ FGL loop interaction with NUP98^APD^. NUP88^NTD^ tyrosine-230 slots into a hydrophobic groove of NUP98^APD^. (**F**) SEC interaction analysis of NUP98^APD^ with NUP88^NTD^ lacking the FGL loop. (**G**) Magnified view of the NUP214^TAIL^ binding site on NUP88^NTD^ highlighting the residues of the LLL mutant (purple), which abolishes the interaction (*top*). Truncation mapping of minimal NUP214^TAIL^ fragment binding NUP88^NTD^ (*bottom*). (**H**) SEC interaction analysis of SUMO-tagged minimal NUP214^TAIL^ with NUP88^NTD^ LLL mutant. SDS-PAGE gel strips of peak fractions are shown and visualized by Coomassie brilliant blue staining.

Because the NUP214^TAIL^-NUP88^NTD^ interaction was crystallographically intractable, we mapped a minimal NUP88^NTD^-binding region spanning NUP214 residues 938-955 by systematic truncation (Fig. 9G and figs. S77 and S78). NUP214^TAIL^ forms a hydrophobic interaction with NUP88^NTD^ at the 6CD insertion, which was abolished by a combined NUP88^NTD^ LLL mutation (L424A, L428A, and L438A), analogous to a mutation we had previously shown to abolish the interaction between the *S. cerevisiae* orthologs Nup159^TAIL^ and Nup82^NTD^ (fig. S78) (*59*). Interestingly, this NUP88 LLL mutation straddles a naturally occurring D434Y mutation in NUP88 linked to a fatal disorder called fetal akinesia deformation sequence (FADS), which is associated with congenital malformations and impaired fetal movement (fig. S79) (*99*). Given its location, the D434Y mutation is expected to interfere with the NUP214^TAIL^ interaction.

Complementing our earlier NUP358^NTD^ surface mutagenesis, we also performed structure-driven mutagenesis of NUP88^NTD^ to identify the NUP358^NTD^ binding site, focusing on acidic surface residues expected to bind to the basic NUP358^NTD^ concave surface. Indeed, we identified several alanine substitutions of acidic residues at the tip of the 6CD insertion and in the adjacent 5C5D extended loop that abolished NUP358^NTD^ binding (fig. S80).

Combined, our structural and biochemical data reveal the molecular details of the interactions between NUP88, NUP214, and NUP98, uncovering that despite the divergence of individual nup sequences, their shape, mode of interaction, and the overall architecture of the nup complex they give rise to are evolutionarily conserved from fungi to humans.

### Docking of the cytoplasmic filament nup complex into unassigned cluster II

Following the placement of five NUP358^NTD^ copies into unassigned density cluster I on the cytoplasmic face of the NPC, we reasoned that the remaining unassigned density cluster II would be composed of the CFNC and that the density might be interpretable by docking structures of rigid subunits.

Unassigned density cluster II exhibits a distinctive shape reminiscent of a cargo crane that projects its arm into the central transport channel from the rim of the cytoplasmic outer ring. It is composed of two near-perpendicular tube-like segments that bisect the NUP75 arms of the distal and proximal Y-shaped CNCs, a globular segment lodged between the base of the long tube-like segment and the proximal NUP75 arm, and a dumbbell-shaped globular density that projects toward the central transport channel (Fig. 10A). Due to the small size and lack of distinctive shape features, the quantitative docking of NUP88^NTD^•NUP98^APD^, NUP214^NTD^•DDX19, GLE1^CTD^•NUP42^GBM^, GLE1^CTD^•NUP42^GBM^•DDX19, RAE1•NUP98^GLEBS^, NUP358^RanBD^•Ran(GMPPNP), NUP358^ZnF^•Ran(GMPPNP) and NUP358^CTD^, into the ∼12 Å cryo-ET map of the intact human NPC from which all the hereto explained density had been subtracted did not result in high confidence solutions (fig. S81). We therefore took the less objective approach of manual placement based on shape complementarity and biochemical restraints, followed by local rigid body refinement.

**Fig. 10.**
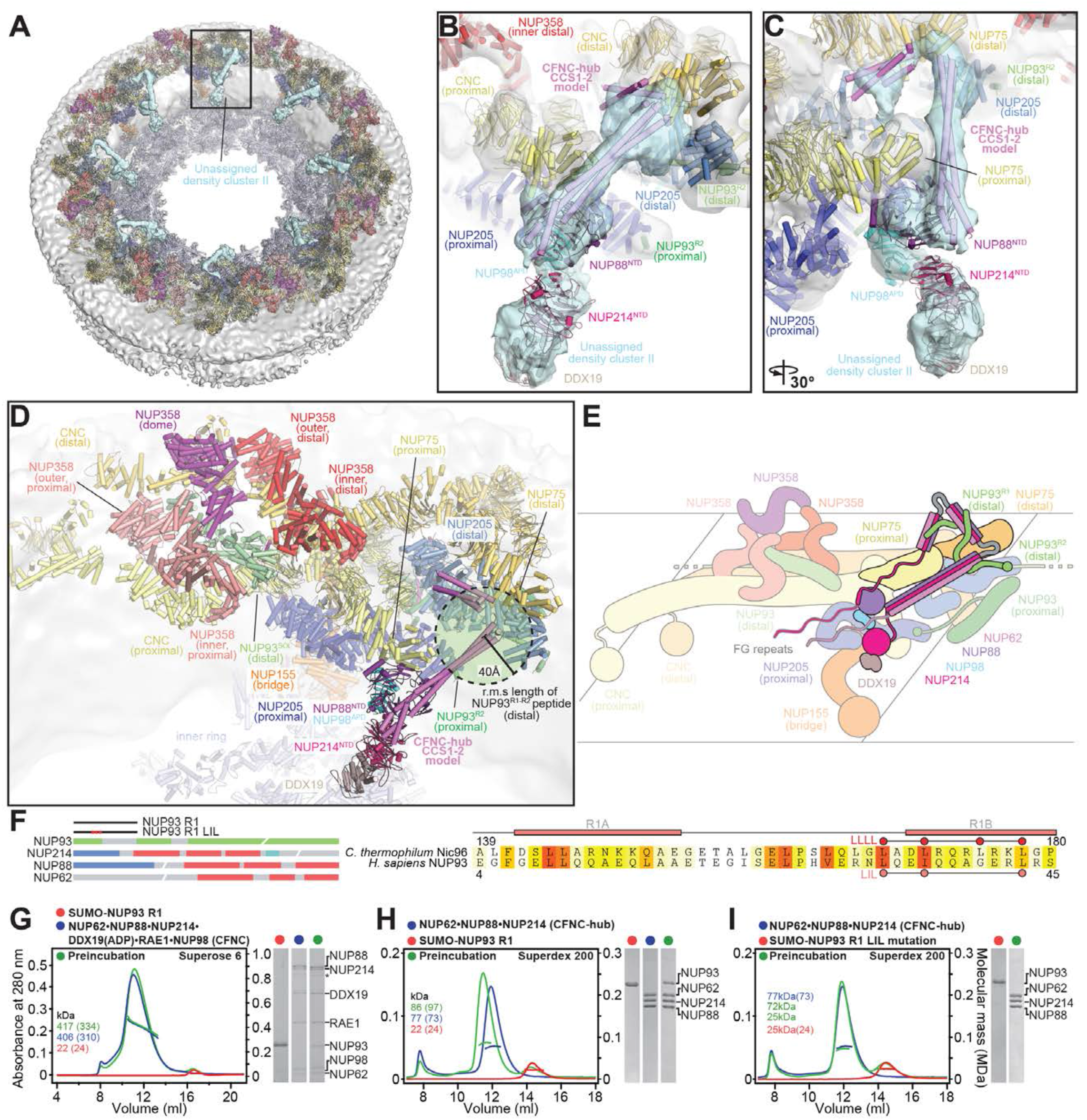
Docking analysis reveals that NUP93 anchors the CFNC to the cytoplasmic outer ring. (**A**) Overview of the NPC cytoplasmic face with isosurface rendering of unexplained density cluster II (cyan) of the ∼12 Å cryo-ET map of the intact human NPC. The inset indicates the location of the magnified view in (B). (**B**, **C**) Two views of manually placed poly-alanine models of CFNC-hub segments CCS1 and CCS2, as well as of the tentatively placed NUP88^NTD^•NUP98^APD^ and NUP214^NTD^•DDX19 co-crystal structures shown in cartoon representation, into unassigned density cluster II. (**D**) Cartoon representation of a cytoplasmic face spoke illustrating the relative positioning of the cytoplasmic filament nups. The root-mean square (r.m.s.) end-to-end length estimate for the NUP93^R1-R2^ linker defines a ∼40 Å radius for the expected location of the distal NUP93^R1^ region. (**E**) Schematic of a cytoplasmic face spoke illustrating CFNC-hub anchoring by the distal NUP93^R1^ positioned by the distal NUP205-bound NUP93^R2^. (**F**) Domain architectures of NUP214, NUP88, NUP62, and NUP93. The location of the NUP93 LIL mutation is indicated (red dots). An alignment of *C. thermophilum* Nic96 and *H. sapiens* NUP93 sequences shows the conservation of residues targeted by the LLLL and LIL mutations that abolish binding to the CNT. Residues are colored according to a multispecies sequence alignment from white (less than 55% similarity), to yellow (55% similarity), to red (100% identity), using the BLOSUM62 weighting algorithm. (**G**, **H**) Mapping of the NUP93^R1^ interaction site to the CFNC-hub. SEC-MALS interaction analysis showing the binding of SUMO-NUP93^R1^ to (G) the hetero-hexameric CFNC and (H) the CFNC-hub. (**I**) SEC-MALS analysis illustrating the abolished binding of the SUMO-NUP93^R1^ LIL mutant to the CFNC-hub. Proteins were visualized by Coomassie brilliant blue staining.

We used the *C. thermophilum* and *X. laevis* CNT crystal structures, the latter containing NUP62, as a template for the polyalanine model of the CCS1 and CCS2 coiled-coil segments of the CFNC-hub (*10, 11*). Remarkably, the CCS1 and CCS2 models based on CNT structures matched the shape and dimensions of two near-perpendicular segments of tube-like density, suggesting that the CFNC-hub coiled-coil architecture is similar to that of the CNT (Fig. 10, B and C, and fig. S82A). NUP88^NTD^•NUP98^APD^ fit best at the base of the long CCS1 segment, interfacing with the NUP75 arm of the proximal CNC. The tentative placement would be consistent with the biochemically mapped interaction between NUP88^NTD^•NUP98^APD^ and NUP214^TAIL^ segment expected to emanate from the C-terminal base of the CCS3 segment and thus restrain NUP88^NTD^•NUP98^APD^ near the CFNC-hub base (Fig. 10, B, C, and E). A dumbbell-shaped density interfacing with NUP88^NTD^•NUP98^APD^ and extending towards the central transport channel is consistent with the shape and size of the NUP214^NTD^•DDX19 crystal structure, although it could be explained by other more transiently tethered components of the nucleocytoplasmic transport machinery or cargo (Fig. 10, B and C, and fig. S82A). However, the NUP214^NTD^•DDX19 complex forms tighter interactions when DDX19 is in its ADP-bound state and would therefore be expected to exist in the ATP-depleted environment of the purified nuclear envelope (*50, 63*). The placements of the CFNC-hub model and NUP88^NTD^•NUP98^APD^ were further supported by manual docking into an anisotropic ∼8 Å region of a composite single particle cryo-EM map of the *X. laevis* cytoplasmic outer ring protomer, though the map masking excluded the region comprising the dumbbell-shaped density (fig. S87B) (*45*).

Placement of the CFNT-hub into the tube-like density puts CCS1, CCS2, and likely the unresolved CCS3, within reach of the ∼40 Å r.m.s length of the linker that tethers NUP93^R1^ to the NUP205-bound NUP93^R2^ (Fig. 10D and fig. S83). In the accompanying paper, we demonstrated that the NUP93^R1^ fragment (residues 2-93), like the orthologous fungal Nic96^R1^ assembly sensor, binds to the CNT complex of the inner ring (*42*). The proximity of the CFNT-hub coiled-coil segments to the expected NUP93^R1^ location suggested that NUP93^R1^ might act as assembly sensor for the CFNT-hub, as well. Indeed, the NUP93^R1^ fragment formed stable complexes with the intact CFNC and CFNC-hub (Fig. 10, E to H, and fig. S84). Strikingly, the NUP93^R1^ LIL (L33A, I36A, L43A) mutation that abolished CNT binding (*42*) also abolished the interaction with the CFNT-hub (Fig. 10I). To ensure we did not miss an interaction of the *C. thermophilum* CFNC (*ct*CFNC), we tested whether Nic96^R1^ could bind the *ct*CFNC-hub. In fact, Nic96^R1^ did not bind to *ct*CFNC-hub, consistent with our reconstitution results that identified two distinct assembly sensors for the *ct*CFNC in the CNC (fig. S85). These data indicate that the long elusive anchoring point of the human CFNC is not on the Y-shaped CNC, but rather provided by the NUP205-positioned NUP93^R1^.

A second NUP93^R1^ assembly sensor emanating from the proximal NUP205-positioned NUP93^R2^ represents a potential anchoring site for a second, proximal CFNC (fig. S83). The lack of density corresponding to a proximal CFNC in the ∼12 Å cryo-ET map of the intact human NPC does not rule out the existence of a flexibly attached proximal CFNC. The placement of 16 copies of the CFNC on the cytoplasmic face of the NPC, half of which are unresolved in the ∼12 Å cryo-ET map, is consistent with the previously established stoichiometry of the intact human NPC from purified nuclear envelopes (*100*). An *in situ ∼*37 Å cryo-ET map of the dilated human NPC (*101*) presents an unexplained elongated density near the expected location of the proximal NUP93^R1^, but could not be further interpreted at the current resolution of the map (fig. S86).

Finally, we tentatively placed the human CF nup GLE1^CTD^•NUP42^GBM^ crystal structure into a region of unexplained density in the ∼12 Å cryo-ET map of the intact human NPC between the cytoplasmic bridge NUP155 and the cytoplasmic face of the nuclear envelope, consistent with our previous analysis (fig. S87) (*63*).

### Steric occlusion is insufficient to explain asymmetric decoration of the NPC

Having assigned all cytoplasmic density of cluster I and cluster II to NUP358 pentameric bundles and CFNCs, respectively, we next wondered whether any structural features prevent NUP358 or CFNC mis-localization at the nuclear face of the NPC. Examining the density remaining on the nuclear face of the ∼12 Å cryo-ET human NPC map, we found that the hereto unexplained density adjacent to the NUP160 arms of the proximal and distal Y-shaped CNCs could be assigned to 16 copies of the structured N-terminal ELYS domains (fig. S88) (*25*). The placed ELYS domains did not overlap with nuclear regions equivalent to the sites occupied by NUP358 and CFNC on the cytoplasmic NPC side, thereby excluding the possibility that steric competition with ELYS prevents NUP358 or CFNC mis-localization on the nuclear side (Fig. 11, A and B). On the contrary, the ∼12 Å cryo-ET map revealed rod-shaped unassigned densities atop the nuclear outer ring in regions equivalent to NUP358 sites on the cytoplasmic face (Fig. 11C). Analogously, we asked if recruitment of the CFNC to the nuclear face was prevented by steric hindrance from a nuclear basket component. Although the NUP205-NUP93^R2^ attachment site from which NUP93^R1^ is flexibly projected remains unencumbered, an unassigned rod-shaped cryo-ET density present on the nuclear face overlaps with an area equivalent to the CFNC-hub CCS1-2 docking sites on the cytoplasmic face (Fig. 11C). These findings suggest that mechanisms other than steric competition alone, such as active nuclear transport of asymmetric nups, are key determinants of the exclusively cytoplasmic and nuclear localizations of NUP358 or the CFNC and ELYS, respectively.

**Fig. 11.**
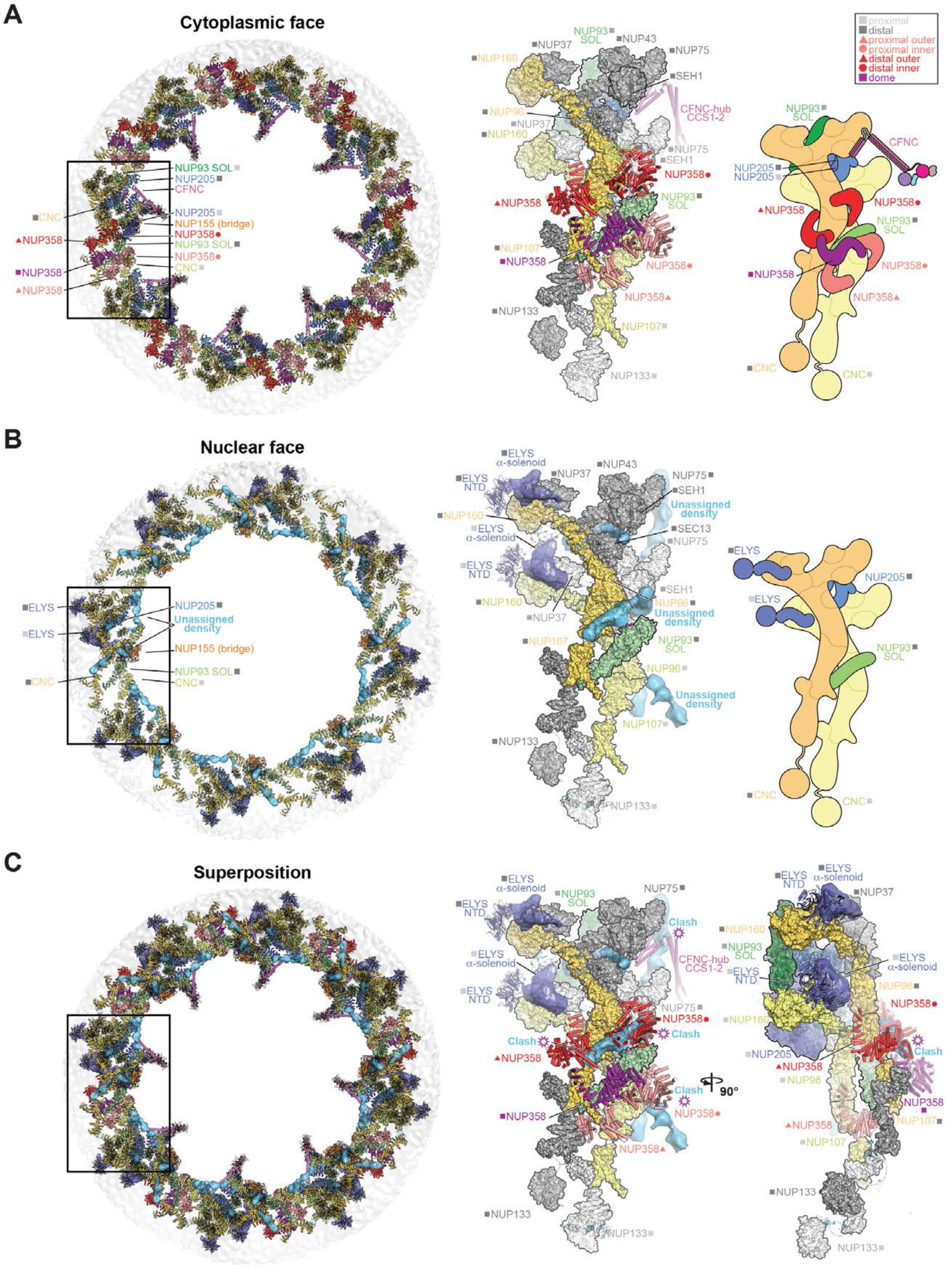
Comparison of cytoplasmic and nuclear faces of the human NPC. Overall top view (*left*), single spoke protomer with symmetric core nups in surface, docked asymmetric nups in cartoon, and unexplained density of the ∼12 Å cryo-ET map in isosurface representation (*middle*), and schematic (*right*) of the (**A**) cytoplasmic and (**B**) nuclear face of the intact human NPC. (**C**) Superpositions of the overall view (*left*) and two orthogonal views of single spoke protomers (*middle* and *right*) of the nuclear and cytoplasmic faces, with hypothetical steric clashes between the cytoplasmic filaments in cartoon representation, and the unassigned asymmetric nuclear density (cyan), indicated. Distal and proximal positions are labeled according to the legend.

Together, our data complete the near-atomic composite structure of the symmetric and cytoplasmic asymmetric portions of the human NPC (Fig. 12).

**Fig. 12.**
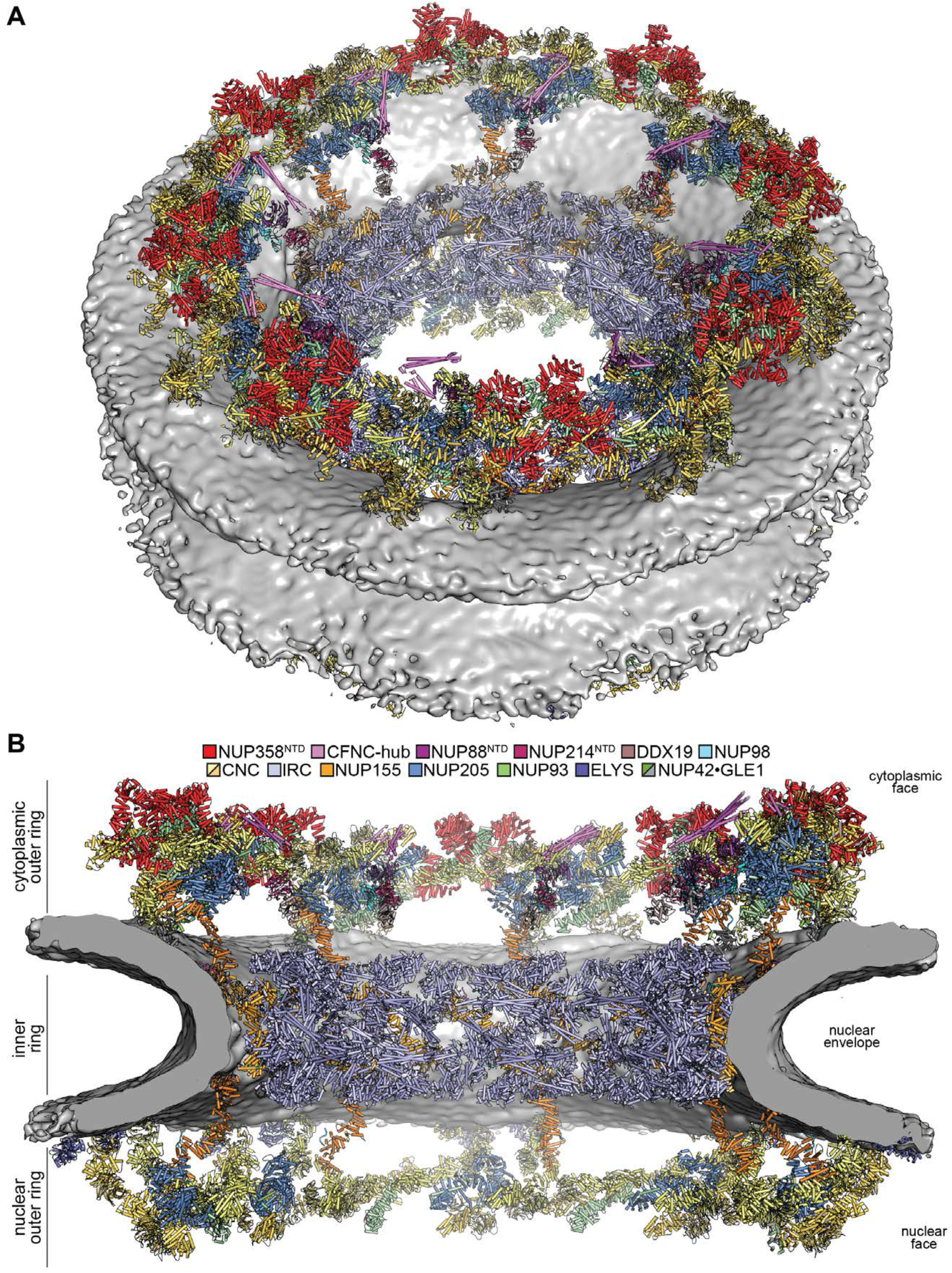
Architecture of the human NPC cytoplasmic face. Near-atomic composite structure of the human NPC generated by docking individual nup and nup complex crystal and single particle cryo-EM structures into a ∼12 Å cryo-ET map of the intact human NPC, viewed from (**A**) the cytoplasmic face, and (**B**) the central transport channel, as a cross-section. Newly placed asymmetric nucleoporin and nucleoporin complex structures include the quantitatively docked NUP358^NTD^ and the manually docked NUP88^NTD^•NUP98^APD^, NUP214^NTD^•DDX19, GLE1^CTD^•NUP42^GBM^, ELYS^NTD^, as well as a model of CFNC-hub. The nuclear envelope is rendered as a grey isosurface, whereas nups are shown in cartoon representation and colored according to the legend.

### Conclusions

Situated on the cytoplasmic face of the NPC, CF nups remodel mRNPs as they emerge from the central transport channel, ensuring directional transport of mRNA and preparing it for downstream translation. Given this essential cellular function, it is unsurprising that CF nups are a hotspot for mutations associated with currently incurable diseases, ranging from neurodegenerative and auto-immune disorders to aggressive cancers. Through a comprehensive analysis combining *in vitro* complex reconstitution, crystal structure determination, quantitative docking, and *in vivo* validation, we established a near-atomic composite structure of the cytoplasmic face of the human NPC.

Our biochemical reconstitution highlights the evolutionary conservation of the CFNC modular assembly, which consists of a central heterotrimeric coiled-coil hub that tethers two separate mRNP remodeling complexes together. Despite the divergence in attachment mechanisms, the anchoring of two copies of the CFNC module to each of the eight NPC spokes appears to be an evolutionarily conserved architectural outcome: The *C. thermophilum* NPC presents two distinct assembly sensor motifs for the CFNC-hub in the Nup37 and Nup145C subunits of each CNC. The human NPC reuses the NUP93 sensor for the assembly and anchoring of the CNT in the inner ring as an anchor for the CFNC in the cytoplasmic outer ring by intercalating two NUP205•NUP93 copies among the tandem-arranged CNCs of each spoke. Previous studies have also shown that the *S. cerevisiae* NPC incorporates a P-shaped CFNC dimer (*61*) to a single site within each of the eight outer ring spokes (*40, 41*).

In addition to the CFNC, the asymmetric CF nup decoration of the human NPC cytoplasmic face includes NUP358. Conjoined by an oligomerization element, pentameric bundles of the uniquely folded NUP358^NTD^ envelop the tandem-arrayed stalks of a CNC pair in each of the eight spokes. Each attached NUP358^NTD^ anchors an extensive ∼2,400-residue C-terminal region that harbors 14 different domains connected by unstructured linkers, thereby reaching as far away as ∼60 nm from the NPC (*102*). Our placement of the CNC and CF nups explains the entirety of the observed cytoplasmic face cryo-ET density, accounting for ∼23 MDa of structured mass. The more flexibly attached regions of the CF nups that are not captured by the current sub-tomogram averaged cryo-ET map account for an additional ∼19 MDa of mass.

We found that the interactions between the *C. thermophilum* CF nups and the CNC are modulated by the small molecule IP_6_, the presence of which is required for mRNA export. Future studies need to address the concerted role of post-translational modifications, second messengers and other small molecules, and macromolecular factors in regulating the assembly and functions of the NPC cytoplasmic face in mRNA export.

The integral membrane proteins and nuclear basket portions of the NPC represent an outstanding challenge for structural determination. Nevertheless, our analysis has already identified that competition for binding sites could play a role in the segregation of CF and nuclear basket nups to opposite faces of the NPC. However, steric occlusion alone is insufficient to deterministically establish NPC polarity, whereby the correct asymmetric nups are segregated to either the cytoplasmic or the nuclear face, or the proximal NUP93 and NUP205 copies are excluded from the nuclear outer ring. Nuclear and cytoplasmic eviction mediated by the nucleocytoplasmic transport machinery is perhaps the most obvious candidate for a mechanism that maintains the polar subcellular segregation of asymmetric nups.

The data presented provide a comprehensive biochemical foundation and a structural framework for the design of future experiments aimed at elucidating the multiple mechanistic steps involved in mRNP export and remodeling. This mechanistic insight will be vital for illuminating disease mechanisms associated with CF nup genetic variants and mechanisms by which viral virulence factors, e.g. SARS-CoV2 ORF6, hijack the functions of the NPC (*103*).

Our results represent a significant step towards the complete *in vitro* reconstitution of the NPC and establish a near-atomic composite structure of the entire cytoplasmic face of the human NPC. More broadly, they illustrate the effectiveness of our divide-and-conquer approach in successfully elucidating the near-atomic architecture of an assembly as large and complex as the NPC, serving as a paradigm for studying similar macromolecular machines, which remains a major frontier in structural cell biology.

## Supporting information

Hoelz_NPCCytoplasmicFace_SI

## Acknowledgements

We thank the members of the Hoelz laboratory for insightful discussions, Martin Beck for sharing of the ∼12Å cryoET reconstruction of the intact human NPC prior to publication, Valerie Doye, Ulrike Kutay, Ed Hurt, and Iain Mattaj for providing material, Alex Cohen, Jacqueline Chou, Reeti Gulati, Young Jeon, Claudia Jette, Hannah Margolis, Taylor Stevens, Evelyn Stuwe, Tobias Stuwe, and Jimmy Thai for experimental support, and Valerie Altounian for the preparation of NPC schematics. We acknowledge Jens Kaiser and the scientific staff of the Stanford Synchrotron Radiation Laboratory (SSRL) Beamline 12-2 and the National Institute of General Medical Sciences and National Cancer Institute Structural Biology Facility (GM/CA) at the Advanced Photon Source (APS) for their support with X-ray diffraction measurements. The Molecular Observatory at the California Institute of Technology (Caltech) is supported by the Gordon and Betty Moore Foundation, the Beckman Institute, and the Sanofi-Aventis Bioengineering Research Program. The operations at the SSRL and APS are supported by the U.S. Department of Energy and the National Institutes of Health (NIH). GM/CA has been funded in whole or in part with federal funds from the National Cancer Institute (ACB-12002) and the National Institute of General Medical Sciences (AGM-12006). D.H.L. was supported by an NIH Research Service Award (5 T32 GM07616) and Amgen Graduate Fellowship through the Caltech-Amgen Research Collaboration. S.P and F.M.H. were supported by a Ph.D. fellowship of the Boehringer Ingelheim Fonds. A.A.K. was supported by NIH grants R01-GM117372 and P50-GM082545. A.H. was supported by a Camille-Dreyfus Teacher Scholar Award and NIH grants R01-GM117360 and R01-GM111461, is an Investigator of the Heritage Medical Research Institute, and a Faculty Scholar of the Howard Hughes Medical Institute. The coordinates and structure factors have been deposited in the Protein Data Bank with accession codes 7MNJ (NUP358^145-673^), 7MNK (NUP358^OE^), 7MNI (NUP88^NTD^•NUP98^APD^), 7MNL (NUP358^NTD^•Fab14), 7MNM (NUP358^NTD^ T585M•sAB-14), 7MNN (NUP358^NTD^ T653I•Fab14), 7MNO (NUP358^NTD^ I656V•Fab14), 7MNP (NUP358-ZnF2•Ran(GDP)), 7MNQ (NUP358-ZnF2•Ran(GDP)), 7MNR (NUP358-ZnF3•Ran(GDP)), 7MNS (NUP358-ZnF4•Ran(GDP)), 7MNT (NUP358-ZnF5/6•Ran(GDP)), 7MNU (NUP358-ZnF7•Ran(GDP)),7MNV (NUP358-ZnF8•Ran(GDP)), 7MNW (NUP358-RanBD-I•Ran(GMPPNP)), 7MNX (Nup358-RanBD-II•Ran(GMPPNP)), 7MNY (NUP358-RanBD-III•Ran(GMPPNP)), 7MNZ (NUP358-RanBD-IV^QNYDNKQV^•Ran(GPPNHP)), 7MO0 (NUP50-RanBD•Ran(GMPPNP)), 7MO1 (NUP153-ZnF1•Ran(GDP)), 7MO2 (NUP153-ZnF2•Ran(GDP)), 7MO3 (NUP153-ZnF3•Ran(GDP), 2.05Å), 7MO4 (NUP153-ZnF3•Ran(GDP), 2.4Å), 7MO5 (NUP153-ZnF4•Ran(GDP)). PyMol and Chimera sessions containing the composite structure of the human NPC cytoplasmic face can be obtained from our webpage (http://ahweb.caltech.edu), as well as deposited on PDB-dev (XXXX) and CaltechDATA (XXXX). The authors declare no financial conflicts of interest.

## Supplementary Materials

Materials and Methods

Figs. S1-S88

Tables S1-S17

References (*104–145*)

## Notes

### Competing Interest Statement

The authors have declared no competing interest.

