## Supplementary material for "Architecture of the cytoplasmic face of the nuclear pore": Hoelz_NPCCytoplasmicFace_SI

**This PDF file includes:**

Materials and Methods
Figs. S1 to S88
Tables S1 to S17

#### Materials and Methods

##### Materials and reagents

A detailed summary of materials and reagents is provided in Table S1.

##### Bacterial expression constructs

*C. thermophilum* cDNA was generated using a previously established protocol (*104*). Briefly, *C. thermophilum (DSM*-1495; gift of Ed Hurt, University of Heidelberg, Germany) mycelium was grown in combined-carbon source media (CCM). After harvesting, washing, and drying, the mycelium was frozen in liquid nitrogen and lysed in a Retsch mill. Total RNA was isolated using the SV Total RNA Isolation System (Promega). cDNA was generated using Superscript III Reverse Transcriptase (Invitrogen) and an oligo d($T_{20}$) primer prior to purification with a QIAquick PCR Purification Kit (QIAGEN). The generation of cDNAs encoding Nsp1, Dbp5, Nup145N, Nup42, Nic96, Nup120, Nup37, Elys, Nup85, Sec13, Nup145C, Nup84, and Nup133 was previously reported (*11, 38, 63*). NUP98 cDNA was a gift of Beatriz Fontoura (UT Southwestern Medical Center, USA) (*105*), and cDNAs encoding NUP42 and NUP155 were a gift of Susan Wente (Vanderbilt University, USA) (*106*). NUP62 and NUP88 cDNAs were obtained commercially (Open Biosystems). To improve bacterial expression of NUP88, the sequence encoding residues 1 to 493 was replaced with a synthetic codon-optimized DNA fragment (GenScript). The generation of cDNAs encoding NUP358 (*80*), NUP214 (*54*), DDX19 (*55*), GLE1 (*63*), and RAE1 (*57*) was previously described.

DNA fragments were amplified by PCR using *C. thermophilum* and *H. sapiens* cDNA and cloned into either a modified pET28a vector encoding an N-terminal $His_6$-tag followed by a PreScission protease cleavage site (pET28a-PreS) (*107*), a modified pET28a vector (Novagen) encoding an N-terminal $His_6$-tag followed by a small ubiquitin-like modifier (SUMO) with or without an uncleavable C-terminal $His_6$-tag (pET28a-SUMO) (*108*), a modified pET-Duet1 vector (Novagen) encoding an N-terminal $His_6$-SUMO-tag in the first expression site and an untagged protein in the second site (pET-Duet1-SUMO), a modified pET-Duet1 vector encoding N-terminal SUMO-tags in both expression sites (pET-Duet1-SUMO2), a modified pET-MCN vector (*109*) encoding an N-terminal $His_6$-tag followed by a PreScission protease cleavage site (pET-MCN-PreS), a modified pET-MCN vector encoding an N-terminal $His_6$-tag followed by SUMO (pET-MCN-SUMO) (*11*), an unmodified pET24a vector (Novagen) encoding an uncleavable N-terminal $His_6$-tagged protein, or pGEX-6P1 vector (GE Healthcare) encoding an N-terminal GST-tag followed by a PreScission protease cleavage site. pET28a-SUMO-Avi and pET-Duet1-SUMO-Avi expression constructs were generated by subsequently cloning an Avi-tag-encoding DNA cassette into the BamHI site. pET28a-$His_6$-PreS-Avi-SUMO constructs were generated by first cloning DNA encoding proteins to pET28a-SUMO using the BamHI and XhoI restriction sites, and subsequently subcloning the SUMO-insert cassette using NdeI and BamHI restriction sites into the previously described pET28a-PreS-Avi vector (*27*). The variable complementarity determining region (CDR) of synthetic antibodies (sAB) was cloned into the pRH2.2 vector (Sidhu lab, University of Toronto, Canada), using the HindIII and SalI restriction sites (*110*). Mutants were generated by QuikChange mutagenesis using PfuUltra II Fusion HotStart DNA Polymerase (Agilent Technologies) and confirmed by DNA sequencing. Details of bacterial expression constructs are summarized in Table S2.

##### Bacterial protein expression

Bacterial protein expression was carried out in *E. coli* BL21-CodonPlus(DE3)-RIL cells in Luria-Bertani (LB) media and induced at an OD 600 of 0.8 with 0.5 mM isopropyl-β-D-thiogalactopyranoside (IPTG). Seleno-L-methionine (SeMet)-labeled proteins were expressed by a methionine pathway inhibition protocol, as previously described (*111*). For details of the expression times and temperatures, see Table S2. Cells were harvested by centrifugation and re-suspended in a buffer containing 20 mM TRIS (pH 8.0), 300 mM NaCl (for cells expressing GST-tagged proteins) or 500 mM NaCl (for cells expressing $His_6$-tagged proteins), 5 mM 2-mercaptoethanol (β-ME), supplemented with complete EDTA-free protease inhibitor cocktail (Roche), 2 μM bovine lung aprotinin (Sigma), and 1 mM phenylmethylsulfonyl fluoride (PMSF, Gold Biotechnology). For purification, the cell suspensions of all proteins and protein complexes were supplemented with 1 mg DNase I and lysed using a cell disruptor (Avestin). Cell lysates were cleared by centrifugation at 30,000 x *g* for 1 hour. The supernatants were filtered through a 0.45 μm filter (Millipore) and purified using standard chromatography methods.

#### Baculoviral protein expression

The baculoviral expression constructs of RAE1 and RAE1•NUP98$^{GLEBS}$ were described previously (*57*). Briefly, a DNA fragment encompassing the entire coding region of RAE1 was cloned into a pFastbac-Dual vector (Gibco), using SmaI and KpnI restriction sites, expressing RAE1 with a non-cleavable C-terminal $His_7$-tag. A DNA fragment encoding NUP98$^{GLEBS}$ (residues 157-213) with an N-terminal tobacco etch virus (TEV) protease cleavable N-terminal GST-tag was cloned into the pFastbac-Dual vector that harbored RAE1 in the second expression site, using BamHI and EcoRI restriction sites. For *C. thermophilum* Gle2, a DNA fragment encompassing the entire coding region of the protein was cloned into the pFastBac-HTB vector (Gibco), using BamHI and XhoI restriction sites, expressing Gle2 with a TEV-cleavable N-terminal $His_6$-tag. Details of Baculoviral expression constructs are summarized in Table S3. Baculoviral stocks were generated using the Bac-to-Bac baculovirus expression system (Gibco) according to the manufacturer's guidelines. Briefly, pFastBac vectors were transformed into DH10Bac cells, bacmids were then extracted by precipitation in 50% (v/v) isopropanol after cell lysis using the QIAprep miniprep kit (QIAGEN). P0 virus was generated by transfection of 10 μl bacmid DNA with 20 μl CellFectin-II (Gibco) into 4 ml of $0.6\times10^6$/ml *S. frugiperda* Sf9 cells grown in Sf-900 III SFM medium (Gibco) incubated at 27 ºC for 5-6 days until ~25% viability. P1 virus was generated by infecting 40 ml of $1.5\times10^6$/ml Sf9 cells in suspension with 135 μl of P0 virus and incubating at 27 ºC for 3-4 days until ~60% viability. Protein expression was carried out in 1 L cultures of $2.5\times10^6$/ml *T. ni* Hi5 cells grown in suspension and infected with 1% (v/v) P1 virus to give an approximate multiplicity of infection (MOI) of 1. Hi5 cells were grown in ESF 921 insect cell culture medium (Expression systems) at 27 ºC, supplemented after 24 hours with 1.5% (v/v) production boost additive (Expression systems). The cells were harvested 2-3 days post infection at ~80% viability through centrifugation and re-suspended in a buffer, containing 20 mM TRIS (pH 8.0), 500 mM NaCl, and 5 mM β-ME, supplemented with Complete EDTA-free protease inhibitor cocktail, 2 μM bovine lung aprotinin, and 1 mM PMSF. For purification, the cell suspensions were supplemented with 1 mg DNase I and lysed using a cell disruptor (Avestin), and 2×30 second cycles of sonication. Cell lysates were cleared by centrifugation at 30,000 × *g* for 1 hour. The supernatants were filtered through a 5 μm filter (Millipore) and purified using standard chromatography methods.

#### Protein purification

A detailed summary of the chromatography steps and associated buffer conditions for all protein purifications is provided in Table S5. The purification and reconstitution of protein complexes and sABs are detailed below.

*Purification of sABs.* The expression of sABs was carried out in *E. coli* BL21-Gold cells (Agilent Technologies) and induced at an OD 600 of 1.0 with 0.5 mM IPTG and grown at 37 °C for 3 hours. The lysate was incubated at 65 °C for 30 minutes and then cooled on ice for 15 minutes before centrifugation at 30,000 × *g* for 1 hour. The supernatant was filtered through a 0.45 μm filter (Millipore) and loaded onto a 5 ml HiTrap MabSelect SuRe column (GE Healthcare), equilibrated in a buffer containing 20 mM TRIS (pH 8.0) and 500 mM NaCl and eluted using a linear gradient of an elution buffer containing 0.1 M acetic acid. The eluted fractions were dialyzed against a buffer containing 20 mM TRIS (pH 8.0) and 100 mM NaCl and concentrated to ~10 mg/ml for biochemical interaction experiments, complex reconstitution, and crystallization.

*Reconstitution of RAE1•NUP98$^{GLEBS\text{-}APD}$.* Purified RAE1 was mixed with a 1.2-fold molar excess of NUP98$^{GLEBS\text{-}APD}$ and incubated on ice for 30 minutes. The mixture was then loaded onto a Superdex 75 10/300 GL gel filtration column (GE Healthcare), equilibrated in buffer containing 20 mM TRIS (pH 8.0), 150 mM NaCl, and 5 mM DTT. Peak fractions containing a stoichiometric RAE1•NUP98$^{GLEBS\text{-}APD}$ complex were pooled and concentrated to ~10 mg/ml for biochemical studies.

*Reconstitution of NUP62•NUP88•NUP214•DDX19(ADP)•RAE1•NUP98$^{GLEBS\text{-}APD}$ (CFNC).* Purified NUP62•NUP88•NUP214 was mixed with a 1.5-fold molar excess of purified DDX19(ADP) and RAE1•NUP98$^{GLEBS\text{-}APD}$ and incubated on ice for 30 minutes. The mixture was then loaded onto a Superose 6 10/300 GL gel filtration column (GE Healthcare), equilibrated in a buffer containing 20 mM TRIS (pH 8.0), 100 mM NaCl, 5 mM DTT, and 5% (v/v) glycerol. Peak fractions containing CFNC were pooled and concentrated to ~7 mg/ml for biochemical studies.

*Reconstitution of NUP88$^{NTD}$•NUP98$^{APD}$.* Purified NUP88$^{NTD}$ was mixed with a 1.2-fold molar excess of NUP98$^{APD}$ and incubated on ice for 1 hour. The mixture was then loaded onto a HiLoad Superdex 200 16/60 PG gel filtration column (GE Healthcare), equilibrated in a buffer containing 20 mM TRIS (pH 8.0), 100 mM NaCl, and 5 mM DTT. Peak fractions containing a stoichiometric NUP88$^{NTD}$•NUP98$^{APD}$ complex were pooled and concentrated to ~20 mg/ml for biochemical studies and crystallization.

*Purification of NUP358$^{NTD}$•sAB-14.* Purified NUP358$^{NTD}$ was mixed with a 1.5-fold molar excess of sAB14 and incubated on ice for 1 hour. The mixture was then loaded onto a HiLoad Superdex 200 16/60 PG gel filtration column equilibrated in buffer containing 20 mM TRIS (pH 8.0), 100 mM NaCl, and 5 mM DTT. Peak fractions containing a stoichiometric NUP358$^{NTD}$•sAB-14 complex were pooled and concentrated to ~10 mg/ml for biochemical studies and crystallization.

*Purification of NUP358/NUP50 RanBDs in complex with Ran(GMPPNP).* Ran nucleotide exchanged from bound GDP to the non-hydrolysable GTP analogue 5'-guanosine-[β,γ-imido]-triphosphate (GMPPNP) was achieved by incubating Ran(GDP) with a two-fold excess of GMPPNP and 2.5 units of alkaline phosphatase (Roche) per mg protein overnight at 4 °C in buffer containing 20 mM potassium phosphate (pH 7.4), 200 mM ammonium sulfate, 0.1 mM $ZnCl_2$, and 1 mM DTT, as previously described (*73*). After nucleotide exchange, protein was buffer exchanged using a 5 ml HiTrap Desalting column (GE Healthcare), equilibrated in a buffer containing 20 mM TRIS (pH 8.0), 100 mM NaCl, 5 mM DTT, and 2 mM $MgCl_2$. The purified RanBD protein was mixed with a 1.5-fold molar excess of Ran(GMPPNP) and incubated on ice for 1 hour. The mixture was injected onto a 5 ml HiTrap QHP column (GE Healthcare) and eluted via a NaCl gradient (100-400 mM). The complex-containing fractions were pooled, concentrated, and further purified by size-exclusion chromatography using a Superdex 200 16/60 PG gel filtration column equilibrated in a buffer containing 20 mM TRIS (pH 8.0), 100 mM NaCl, 5 mM DTT, and 2 mM $MgCl_2$. For Nup358$^{RanBD-II}$ and Nup50$^{RanBD}$, the HiTrap QHP column step was omitted. Peak fractions containing stoichiometric RanBD•Ran(GMPPNP) complex were pooled and concentrated to ~15 mg/ml for biochemical studies and crystallization.

*Reconstitution of Nup82•Nup159•Nsp1•Dpb5•Gle2•Nup145N$^{GLEBS-APD}$ (CFNC).* Purified Nup82•Nup159•Nsp1 was mixed with a 1.5-fold molar excess of purified Dpb5 and Gle2•Nup145N$^{GLEBS-APD}$ and incubated on ice for 1 hour before injection onto a Superose 6 10/300 GL gel filtration column equilibrated in a buffer containing 20 mM TRIS (pH 8.0), 100 mM NaCl, 5 mM DTT, and 5% (v/v) glycerol. Peak fractions containing stoichiometric CFNC were pooled and concentrated to ~10 mg/ml for biochemical studies.

*Reconstitution of Nup120•Nup37•Elys•Nup85•Sec13-Nup145C (CNC-hexamer).* Purified Nup120•Nup37•Elys•Nup85 hetero-tetramer was mixed with a 1.2-fold molar excess of purified Sec13-Nup145C, incubated for 1 hour on ice, and injected onto a Superose 6 10/300 GL gel filtration column equilibrated in a buffer containing 20 mM TRIS (pH 8.0), 100 mM NaCl, 5 mM DTT, and 5% (v/v) glycerol. Peak fractions containing stoichiometric CNC-hexamer were pooled and concentrated to ~10 mg/ml for biochemical studies and further reconstitutions.

*Reconstitution of Nup120•Nup37$^{\Delta CTE}$•Elys•Nup85•Sec13-Nup145C (CNC-hexamer$^{\Delta 37CTE}$).* Purified Nup120•Nup37$^{\Delta CTE}$•Elys•Nup85 hetero-tetramer was mixed with a 1.2-fold molar excess of purified Sec13-Nup145C and incubated for 1 hour on ice. The mixture was loaded onto a Superose 6 10/300 GL gel filtration column equilibrated in a buffer containing 20 mM TRIS (pH 8.0), 100 mM NaCl, 5 mM DTT, and 5% (v/v) glycerol. Peak fractions containing stoichiometric CNC-hexamer$^{\Delta 37CTE}$ were pooled and concentrated to ~10 mg/ml for biochemical studies and further reconstitutions.

*Reconstitution of Nup120•Nup37•Elys•Nup85•Sec13-Nup145C$^{\Delta NTE}$ (CNC-hexamer$^{\Delta 145CNTE}$).* Purified Nup120•Nup37•Elys•Nup85 hetero-tetramer was mixed with a 1.2-fold molar excess of purified Sec13-Nup145C$^{\Delta NTE}$ and incubated for 1 hour on ice. The mixture was loaded onto a Superose 6 10/300 GL gel filtration column equilibrated in a buffer containing 20 mM TRIS (pH 8.0), 100 mM NaCl, 5 mM DTT, and 5% (v/v) glycerol. Peak fractions containing stoichiometric CNC-hexamer$^{\Delta 145CNTE}$ were pooled and concentrated to ~10 mg/ml for biochemical studies and further reconstitutions.

*Reconstitution of Nup120•Nup37$^{\Delta CTE}$•Elys•Nup85•Sec13-Nup145C$^{\Delta NTE}$ (CNC-hexamer$^{\Delta 37CTE \Delta 145CNTE}$).* Purified Nup120•Nup37$^{\Delta CTE}$•Elys•Nup85 hetero-tetramer was mixed with a 1.2-fold molar excess of purified Sec13-Nup145C$^{\Delta NTE}$ and incubated for 1 hour on ice. The mixture was loaded onto a Superose 6 10/300 GL gel filtration column equilibrated in a buffer containing 20 mM TRIS (pH 8.0), 100 mM NaCl, 5 mM DTT, and 5% (v/v) glycerol. Peak fractions containing stoichiometric CNC-hexamer$^{\Delta 37CTE \Delta 145CNTE}$ were pooled and concentrated to ~10 mg/ml for biochemical studies and further reconstitutions.

*Reconstitution of Nup120•Nup37•Elys•Nup85•Sec13-Nup145C•Nup84•Nup133$^{\Delta NTE}$ (CNC).* Purified Nup120•Nup37•Elys•Nup85•Sec13-Nup145C (CNC-hexamer) was mixed with a 1.2-fold molar excess of purified Nup84•Nup133$^{\Delta NTE}$ hetero-dimer, incubated for 1 hour on ice, and injected onto a Superose 6 10/300 GL gel filtration column, equilibrated in a buffer containing 20 mM TRIS (pH 8.0), 100 mM NaCl, 5 mM DTT, and 5% (v/v) glycerol. Peak fractions containing stoichiometric CNC were pooled and concentrated to ~10 mg/ml for biochemical studies and further reconstitutions. CNC variants lacking either Nup37$^{CTE}$, Nup145C$^{NTE}$, or both were reconstituted using the corresponding CNC-hexamer variants.

*Reconstitution of Nup82$^{NTD}$•Nup145N$^{APD}$•Nup159$^{TAIL}$.* Purified Nup82$^{NTD}$•Nup159$^{TAIL}$ heterodimer was mixed with a 1.5-fold molar excess of purified Nup145N$^{APD}$, incubated for 1 hour on ice, and injected onto a HiLoad Superdex 200 16/60 PG gel filtration column, equilibrated in a buffer containing 20 mM TRIS (pH 8.0), 100 mM NaCl, and 5 mM DTT. Peak fractions containing a stoichiometric Nup82$^{NTD}$•Nup145N$^{APD}$•Nup159$^{TAIL}$ were pooled and concentrated to ~30 mg/ml for biochemical studies.

*Reconstitution of Nup159$^{NTD}$• Dbp5.* Purified Dbp5 was mixed with a 1.5-fold molar excess of Nup159$^{NTD}$ and incubated on ice for 1 hour. The mixture was then loaded onto a HiLoad Superdex 200 16/60 PG gel filtration column, equilibrated in a buffer containing 20 mM TRIS (pH 8.0), 100 mM NaCl, and 5 mM DTT. Peak fractions containing a stoichiometric Nup159$^{NTD}$•Dbp5 were pooled and concentrated to ~20 mg/ml for biochemical studies.

*Reconstitution of Gle2•Nup145$^{GLEBS-APD}$.* Purified Gle2 was mixed with a 1.5-fold molar excess of Nup145N$^{GLEBS-APD}$ and incubated on ice for 1 hour. The mixture was then loaded onto a HiLoad Superdex 75 16/60 PG gel filtration column, equilibrated in a buffer containing 20 mM TRIS (pH 8.0), 100 mM NaCl, and 5 mM DTT. Peak fractions containing a stoichiometric Gle2•Nup145N$^{GLEBS-APD}$ were pooled and concentrated to ~10 mg/ml for biochemical studies.

*Reconstitution of Gle2•Nup145N$^{GLEBS}$.* Purified Gle2 was mixed with a 1.5-fold molar excess of Nup145N$^{GLEBS}$ and incubated on ice for 1 hour. The mixture was then loaded onto a HiLoad Superdex 75 16/60 PG gel filtration column equilibrated in a buffer containing 20 mM TRIS (pH 8.0), 100 mM NaCl, and 5 mM DTT. Peak fractions containing a stoichiometric Gle2•Nup145N$^{GLEBS}$ were pooled and concentrated to ~20 mg/ml for biochemical studies.

**Multi-angle light scattering coupled to analytical size-exclusion chromatography (SEC-MALS)**

Purified proteins and complex formation were characterized by inline multi-angle light scattering following separation on Superdex 200 10/300 GL, Superdex 200 Increase 10/300 GL, or Superose 6 10/300 GL columns (*112*). *C. thermophilum* proteins and complexes were primarily analyzed in a buffer containing 20 mM TRIS (pH 8.0), 100 mM NaCl, 5 mM DTT, and 5% (v/v) glycerol. The stability of the CFNC-hub interactions with Nup37$^{CTE}$ and Nup145C$^{NTE}$ was assessed in a series of buffers containing 20 mM TRIS (pH 8.0), 5 mM DTT, 5% (v/v) glycerol, and increasing NaCl concentrations (100 mM, 200 mM, 300 mM, 500 mM, or 1000 mM). Reconstitution of the human CFNC hexamer was also tested in a buffer containing 20 mM TRIS (pH 8.0), 100 mM NaCl, 5 mM DTT, and 5% (v/v) glycerol. Whereas the human CFNC pairwise interaction network was conducted in a buffer containing 20 mM TRIS (pH 8.0), 100 mM NaCl, and 5 mM DTT. The oligomerization behavior of NUP358$^{NTD}$, NUP358$^{OE}$, and the NUP358$^{NTD-OE}$ LIQIML mutant was analyzed in a 20 mM TRIS (pH 8.0), 5 mM DTT buffer, containing either 100 mM or 350 mM NaCl. Because of solubility restrictions, NUP358$^{NTD-OE}$ was analyzed exclusively in a buffer containing 350 mM NaCl. The chromatography system was connected in series with an 18-angle light scattering detector (DAWN HELEOS II, Wyatt Technology), a dynamic light scattering detector (DynaPro Nanostar, Wyatt Technology), and a refractive index detector (Optilab t-rEX, Wyatt Technology). Data were collected every 1 second at a flow rate of 0.5 ml/min (Superdex 200 10/300 GL, Superdex 200 Increase

10/300 GL) or 0.4 ml/min (Superose 6 10/300 GL) at 25 ºC and analyzed using ASTRA 6 software, yielding molar mass and mass distribution (polydispersity) of the samples. For interaction studies, proteins were mixed and incubated on ice for 30 minutes prior to being applied to the gel filtration column. Protein-containing fractions were analyzed by SDS-PAGE followed by Coomassie brilliant blue staining. Theoretical and experimental masses of all proteins and complexes measured by SEC-MALS are listed in Table S6.

**Liquid-liquid phase separation (LLPS) protein-protein interaction assay**

Proteins were labeled under conditions that were selective for the N-terminal amino group, as previously described (*38*). Prior to labeling, purified protein complexes at concentrations of 10-20 μM were dialyzed in a buffer containing 100 mM sodium bicarbonate (pH 8.0), 100 mM NaCl, 5% (v/v) glycerol, and 5 mM DTT. For the labeling reactions three fluorescent dyes were utilized, Bodipy (boron-dipyrromethene) NHS ester (succinimidyl ester; Invitrogen), Coumarin (7-hydroxycoumarin-3-carboxylic acid) NHS ester (Sigma-Aldrich), or Alexa Fluor 647 NHS ester (Invitrogen), each dissolved in dimethyl sulfoxide (DMSO; Sigma-Aldrich) and added at equimolar ratios to the dialyzed protein samples, then incubated for 50 minutes at room temperature. Subsequently, the labeling reaction was quenched by buffer exchange using a 5 ml HiTrap Desalting column, equilibrated in a buffer containing 25 mM TRIS (pH 8.0), 100 mM NaCl, 5% (v/v) glycerol, and 5 mM DTT. Fluorescently labeled proteins were concentrated to 10-20 mg/ml for use in LLPS interaction assays. CNC-hexamer variants were labeled with Bodipy, the CFNC and its subcomplexes Dbp5•Nup159$^{NTD}$, Nup82$^{NTD}$•Nup145N$^{APD}$•Nup159$^{TAIL}$, Gle2•Nup145N$^{GLEBS}$, and CFNC-hub were labeled with Alexa Fluor 647, and Gle1•Nup42$^{GBM}$, Gle1$^{NTD}$, and Gle1$^{CTD}$•Nup42$^{GBM}$ were labeled with Coumarin.

LLPS droplet formation was analyzed by fluorescence microscopy with a Carl Zeiss AxioImagerZ.1 microscope equipped with a Hamamatsu C10600 Orca-R2 camera and a 63×oil immersion objective. Fluorescently labeled proteins were mixed directly on a glass coverslip in a total volume of 10 μl in a buffer containing 25 mM TRIS (pH 8.0), 100 mM NaCl, 5% (v/v) glycerol, and 5 mM DTT. CNC-LLPS formation was triggered by combining Bodipy-labeled CNC-hexamer with a 1.2-fold molar excess of unlabeled Nup84•Nup133 with final concentrations of 8 μM and 9.6 μM, respectively. In a stepwise manner, the Alexa Fluor 647- and Coumarin-labeled cytoplasmic filament nups were added to the CNC-LLPS in 1.2-fold molar excess with final concentrations of 9.6 μM. In a subset of experiments, $IP_6$ was added at a 200 μM final concentration. These LLPS mixtures were incubated for 10 minutes at room temperature prior to imaging. Analogously, an SDS-PAGE analysis of proteins present in the LLPS condensate was conducted with unlabeled proteins. The CNC-hexamer at a final concentration of 8 μM was mixed with a 1.1-fold molar excess of Nup84•Nup133 and the cytoplasmic filament nups in a total volume of 40 μl. These mixtures were incubated on ice for 10 minutes prior to centrifugation at 20,000 × *g* for 10 minutes. The pellet was resuspended in 55 μl 1.5×SDS-PAGE loading dye, while the soluble fraction was transferred to a fresh tube, to which 15 μl 4×SDS-PAGE loading dye was added. Subsequently, the pellet and soluble fractions were analyzed by SDS-PAGE and visualized by Coomassie brilliant blue staining.

**Biotinylation of Avi-tagged protein**

Biotinylation was carried out with 40 μM purified Avi-tagged proteins in a buffer containing 50 mM bicine (pH 8.3), 100 mM biotin, 10 mM ATP, 10 mM magnesium acetate, and 30 μg biotin ligase (BirA) at 30 °C for 2 hours in a total volume of 2 ml. After labeling, proteins were buffer exchanged using a 5 ml HiTrap Desalting column equilibrated in a buffer containing 20 mM TRIS (pH 8.0), 100 mM NaCl, and 5 mM DTT. The extent of biotinylation and efficiency of capture were tested by incubating 10 μg protein with 50 μl of Streptavidin MagneSphere Paramagnetic Particles (Promega), followed by a single washing step with 50 μl buffer containing 20 mM TRIS (pH 8.0), 100 mM NaCl, and 5 mM DTT. The bound proteins were resolved by SDS-PAGE and visualizing by Coomassie brilliant blue staining.

**Streptavidin pulldowns of CFNC-hub with Nup37 and Nup145C assembly sensors**

Streptavidin magnetic beads were prebound with biotinylated Nup37$^{CTE}$ or Nup145C$^{NTE}$ bait by incubating 50 μl of resuspended streptavidin beads with 10 μM bait in 50 μl buffer containing 20 mM TRIS (pH 8.0), 300 mM NaCl, and 4 mM β-ME for 30 minutes at room temperature. Bait-bound streptavidin beads were washed three times with 50 μl buffer, discarding the supernatant. Beads were incubated on ice for 30 minutes, with 30 μl of clarified lysate for the individually expressed Nup82, Nup159 and Nsp1 coiled-coil segments either in isolation, or 10 μl of each lysate mixed together, then diluted to a total volume of 75 μl with buffer. Beads were washed twice with 50 μl buffer discarding the supernatant and then resuspended in 50 μl buffer and supplemented with SDS loading dye. Competition experiments were performed by mixing 150 μM unlabeled $Nup37^{CTE}$ or $Nup145C^{NTE}$ into lysate before incubation with bait-bound streptavidin magnetic beads. Competitor protein was added with an ~10-fold excess relative to prey. Bound proteins were analyzed by SDS-PAGE and visualized by Coomassie brilliant blue staining.

##### *In vitro* RNA transcription and purification

$U_{20}$ single stranded ssRNA (5'-GGGUUUUUUUUUUUUUUUUUUUU-3'), $A_{20}$ ssRNA (5'-GGGAAAAA AAAAAAAAAAAAAAA-3'), double stranded dsRNA (5'-GGCCGGGCCCGGTTCGCCGGGCCCGGC C-3'), CTE1/2 RNA (5'-GUGACAUUUUCACUAACCUAAGACAGGAGGGCCGUCAGAGCUACUGCC UAAUCCAAAGACGGUAAAGUGAUAAAAAUGUAUCAC-3'), and RNA encoding the placental alkaline phosphatase signal sequence coding region ($ALPP^{SSCR}$, 5'-GGACCAUGCUGGGGCCCUGCAUGCUG CUGCUGCUGCUGCUGCUGGGCCUGAGGCUACAGCCUCGA-3') were prepared by *in vitro* transcription using T7 RNA polymerase in a 1 ml run-off transcription reaction containing 40 mM TRIS (pH 8.0), 20 mM $MgCl_2$, 2 mM spermidine-HCl, 10 mM DTT, 7.5 mM of each NTP, 15 U inorganic pyrophosphatase from baker's yeast (Sigma-Aldrich), 20 U SUPERase-In (Invitrogen) RNase inhibitor, and 25 μg PCR generated template. After a 4-hour incubation at 37 °C, the RNA was resolved on a 0.5×TBE (44.5 mM TRIS-borate (pH 8.3), 1 mM EDTA), 10% denaturing polyacrylamide-urea gel and visualized by UV shadowing. For purification, the RNA band was excised and passively eluted overnight at 4 °C in a buffer containing 25 mM TRIS (pH 8.0), 300 mM sodium acetate, and 0.5 mM EDTA (pH 8.0), followed by extraction with an acid-phenol:chloroform:isoamyl alcohol (125:134:1) mixture (Invitrogen) and ethanol precipitation. Pelleted RNA was resuspended in a 10 mM TRIS (pH 8.5) buffer. The RNA concentration was measured by UV absorbance at 260 nm.

##### Radioactive 5'-end labeling of RNA and DNA

Prior to radiolabeling, 600 ng RNA was dephosphorylated with 5 U calf intestinal alkaline phosphatase (New England Biolabs) and 10 U SUPERase-In (Invitrogen) for 1 hour at 37 °C in 1×NEB Buffer 4 (New England Biolabs) containing 50 mM potassium acetate, 20 mM TRIS-acetate, and 10 mM magnesium acetate (pH 7.9). Following dephosphorylation, RNA was extracted twice with an acid-phenol:chloroform:isoamyl alcohol (125:134:1) mixture. After ethanol precipitation, the pelleted RNA was resuspended in 25 μl of a 10 mM TRIS (pH 8.5) buffer. Radioactive $^{32}P$ 5'-end labeling of 600 ng of dephosphorylated RNA or DNA oligonucleotides was carried out in a 30 μl reaction mixture volume, composed of 1×PNK buffer (New England Biolabs) containing 70 mM TRIS-HCl (pH 7.6), 10 mM $MgCl_2$, and 5 mM DTT, supplemented with 0.5μl $\gamma$-$^{32}P$-ATP (3,000 Ci/mmol, 10 μCi/μl; Perkin Elmer), 5 U T4 polynucleotide kinase (New England Biolabs), and 10 U SUPERase-In, and incubated for 1 hour at 37 °C. Free nucleotide was removed with a 0.5 ml Zeba Spin Desalting Column (Thermo Scientific), following the manufacturer's protocol. For future experiments, radiolabeled RNAs and DNAs were stored at -80 °C at a concentration of ~5 ng/μl.

##### Electrophoretic mobility shift assay (EMSA)

Prior to use, $U_{20}$ ssRNA, $A_{20}$ ssRNA, dsRNA, CTE1/2 RNA, and $ALPP^{SSCR}$ RNA and ssDNA were diluted in 10 mM TRIS (pH8.0) heated to 80 °C for 2 minutes and snap cooled on ice and diluted into EMSA buffer containing 20 mM TRIS (pH 8.0), 100 mM NaCl, 1 mM $MgCl_2$, and 5 mM DTT. A DNA oligonucleotide encoding U1 3' box (5'-ACTTTCTGGAGTTTCAAAAACAGACTGTACGCCA-3') was used as a ssDNA probe, which was hybridized with a complimentary oligonucleotide (5'-TGGCGTACAGTCTGTTTTTGAAACTCCAGAAAGT-3') to form a dsDNA probe, as previously described (*113*). dsDNA was prepared by heating to 80 °C and slowly cooled on the benchtop. Protein was serial diluted in EMSA buffer and mixed with an equal volume of 25 nM RNA/DNA or 2.5 nM RNA/DNA for unlabeled and $^{32}P$ radiolabeled probes, respectively. DDX19 binding reactions were supplemented with 5% (v/v) glycerol. The protein-RNA/DNA mixture was incubated on ice for 30 minutes and resolved on a pre-run 0.25×TBE, 5% polyacrylamide gel run at a constant voltage of 10 V/cm for

35 minutes. Unlabeled gels were incubated with SYBR Gold (Invitrogen) stain (1:10,000 dilution in 0.5×TBE) for 3 minutes and imaged on a FluorChem HD2 imager. Radioactive $^{32}P$-end labeled gels were dried, exposed to storage phosphor screen and imaged on a Typhoon FLA 9000 (GE Healthcare).

RAE1•NUP98$^{GLEBS}$ and Gle2•Nup145N$^{GLEBS}$ gel shifts were performed as previously described for RAE1•NUP98$^{GLEBS}$ with minor modifications (*57*). Protein was serial diluted in 20 mM TRIS (pH 8.5), 150 mM NaCl, and 1 mM DTT and mixed with an equal volume of nucleic acid diluted in the same buffer. The protein-RNA/DNA mixture was incubated 5 minutes at room temperature before being loaded on a pre-run 0.5×TBE (pH 8.5), 6% polyacrylamide native gel run at a constant voltage of 3 V/cm for 3 hours at 4 °C. The gel was stained with SYBR Gold for 3 minutes and imaged.

**Generation of high affinity synthetic antibodies (sABs) by phage display**

sABs were obtained by phage display selection using library E and a biotinylated NUP358$^{145\text{-}673}$ bait, following published protocols (*81, 114*). In the first round, 1 ml of phage library E containing $10^{12}$-$10^{13}$ virions was added to streptavidin magnetic beads pre-coated with 200 nM of biotinylated NUP358$^{145\text{-}673}$ and incubated for 30 minutes. Beads containing bound virions after extensive washing were used to infect freshly grown log-phase *E. coli* XL1-Blue cells (Agilent Technologies). Phages were amplified overnight in 2×YT media with 50 μg/ml ampicillin and $10^9$ plaque-forming units/ml M13-KO7 helper phage (New England Biolabs). To increase the stringency of selection, three additional rounds of sorting were performed, using the amplified pool of virions of the preceding round as the input and decreasing the NUP358$^{145\text{-}673}$ concentration to 100 nM, 50 nM and 10 nM in the second, third, and fourth rounds, respectively. Sorting from the second to fourth rounds was done on a Kingfisher instrument. From the second round onward, a 0.1 M glycine (pH 2.7) buffer was used to elute the bound phage from immobilized NUP358$^{145\text{-}673}$. To eliminate potential streptavidin binders, the amplified phage pool after each round was negatively selected against streptavidin magnetic beads before using as input for the next round. After four rounds of selection, the enrichment ratio, defined as phage titer of the NUP358$^{145\text{-}673}$ specific binders over the background, was >50.

A single-point competitive phage ELISA was used to rapidly estimate the affinities of the obtained sABs in phage format (*115*). Colonies of *E. coli* XL1-Blue harboring phagemids were inoculated directly into 500 μl of 2×YT media supplemented with 100 μg/ml ampicillin and M13-KO7 helper phage. The cultures were grown at 37 °C for 16-20 hours in a 96 deep-well block plate shaken at 280 rpm. 25 nM biotinylated NUP358$^{145\text{-}673}$ was immobilized on ELISA plates (Greiner Bio-One) coated with neutravidin (ThermoFisher) and blocked with 0.5% (w/v) bovine serum albumin (BSA, Sigma-Aldrich) in TRIS-buffered saline (TBS) buffer containing 50 mM TRIS (pH 7.5) and 150 mM NaCl. Culture supernatants containing sAB-phage were diluted ten-fold in TBS-T buffer (TBS buffer supplemented with 0.05% (v/v) Tween-20) in the absence or presence of 50 nM NUP358$^{145\text{-}673}$ protein as soluble competitor in a 96-well plate. After 15-minute incubation at room temperature, the mixtures were transferred to ELISA plates, containing the immobilized NUP358$^{145\text{-}673}$. The ELISA plates were incubated with the phage-competitor mixture for another 15 minutes and then washed with TBS-T buffer. The washed ELISA plates were incubated with a mouse monoclonal HRP conjugated anti-M13 phage coat protein antibody (Invitrogen, 1:5,000 dilution in TBS-T buffer) for 30 minutes. After a final TBS-T buffer wash, the plates were developed with 3,3′,5,5′-tetramethylbenzidine (TMB) substrate (Pierce), quenched with 1.0 M HCl, and the absorbance determined spectrophotometrically at 450 nm ($A_{450}$). For each clone, the ratio of $A_{450}$ in the presence or absence of 50 nM competitor gives the fraction of Fab-phage un-complexed with soluble competitor (competition ratio). DNA of the clones with high ELISA signal and low competition ratio were sequenced to identify the number of unique binders. For Nup358$^{145\text{-}673}$, 62 unique binders were obtained. Binders were ranked on basis of high ELISA signal and low competition ratio then reformatted as sAB in the expression vector pRH2.2 to generate untagged antibody constructs for subsequent expression and purification.

**Crystallization and structure determination**

Crystallization trials for all proteins were conducted using hanging drop vapor diffusion at 21 °C with drops containing 1 μl of protein and 1 μl of reservoir solution. Details regarding protein crystallization and cryoprotection conditions are supplied in Table S7. X-ray diffraction data were collected at 100 K, on GM/CA-CAT beamlines 23ID-B and 23ID-D at the Advanced Photon Source (APS), and beamline BL12-2 at the Stanford Synchrotron Radiation Source (SSRL). X-ray diffraction data were either processed in HKL2000 (*116*), or integrated with XDS (*117*) or DIALS (*118*) and scaled with AIMLESS (*119*). Initial phases were calculated in PHASER (*120*) or SHARP (*121*) by single anomalous dispersion (SAD), using anomalous X-ray diffraction data collected at the peak wavelength. The positions of the anomalous scatterers were determined using SHELXD (*122*) or HYSS (*123*). The inclusion of additional native and anomalous inflection and high energy remote wavelength X-ray diffraction data in subsequent phase calculations was systematically assessed for their ability to help in the identification of anomalous scatterer positions and to improve the obtained electron density maps. Solvent flattening and NCS averaging were performed in RESOLVE (*124*) or DM (*125*), yielding improved phases and electron density maps of excellent quality. Initial models were improved by iterative rounds of model building and refinement using Coot (*126*) and PHENIX (*127*), respectively. The final coordinates of all crystal structures exhibit excellent stereochemistry and geometry, as determined by MolProbity (*128*). Details of data processing and refinement statistics are provided in Tables S8 to S17.

*Crystal structure determination of NUP358$^{145\text{-}673}$*. NUP358$^{145\text{-}673}$ was crystalized at 21 °C with the hanging drop method using 1 μl of protein solution (20 mg/ml) and 1 μl of reservoir solution, composed of 0.1 M HEPES (pH 7.0), 0.8 M succinic acid (pH 7.0), and 1% (w/v) PEG 2,000 MME. Crystals were cryoprotected by exchange into reservoir solution containing 20% (v/v) ethylene glycol, prior to plunge freezing in liquid nitrogen. Native data were integrated with DIALS (*118*) and scaled with AIMLESS (*119*) to 3.8 Å resolution. Anomalous SeMet X-ray diffraction data were collected at three wavelengths, corresponding to the selenium peak, inflection and high energy remote energies, and processed in HKL2000 to 4.2 Å resolution (*116*). Initial phases were calculated in SHARP (*121*) by multiple isomorphous replacement with anomalous scattering (MIRAS), identifying 20 selenium sites with 3 molecules in the asymmetric unit (ASU). Density modification in DM with solvent flattening, non-crystallographic symmetry (NCS) averaging, and histogram matching, yielded an electron density map of sufficient quality for *de novo* building of a complete initial model. Iterative rounds of model building and refinement were performed using the 3.8 Å resolution native X-ray diffraction data in Coot (*126*) and PHENIX (*127*), respectively. Refinement was performed with NCS restraints, manually assigned for three sections, encompassing residues 148-215, 228-502, and 512-672. The unambiguous assignment of the sequence register was achieved by anomalous difference Fourier map calculations using anomalous X-ray diffraction data, obtained from a series of SeMet-labeled crystals, each harboring methionine substitutions (I169M, I372M, I385M, I401M, I481M, I599M, I615M and I652M). The final structure possessed excellent stereochemistry and geometry and was refined to 3.8 Å resolution with $R_{work}$ and $R_{free}$ values of 20.8% and 24.2%, respectively. For data processing and refinement statistics, see Tables S8 and S9.

*Crystal structure determination of NUP358$^{NTD}$•sAB-14.* Despite extensive efforts, NUP358$^{NTD}$ did not yield well diffracting crystals. This hurdle was overcome by systematically employing high affinity, conformation specific, synthetic antibodies (sABs) as crystallization chaperones. Crystals of NUP358$^{NTD}$•sAB-14 were obtained at 21 °C with the hanging drop method using 1 μl of protein solution (5 mg/ml) and 1 μl of reservoir solution, composed of 0.1 M sodium citrate (pH 4.6), 0.15 M sodium acetate, and 3% (w/v) PEG 4,000. Crystals were cryoprotected by exchange into reservoir solution containing 25% (v/v) ethylene glycol, prior to plunge freezing in liquid nitrogen. Native data was integrated with XDS (*117*) and scaled with AIMLESS (*119*) to 3.95 Å resolution. The NUP358$^{NTD}$•sAB-14 structure was determined by molecular replacement with PHASER (*120*), using the coordinates of NUP358$^{TPR}$ (PDB ID 4GA0) (*80*), sAB-158 (PDB ID 5CWS) (*11*) and NUP358$^{145\text{-}673}$ as search models, identifying two NUP358$^{NTD}$•sAB-14 complexes per ASU. Model building was aided by a single isomorphous replacement anomalous scattering (SIRAS) experimental map, combining SeMet-peak SAD measurements with native X-ray diffraction data. Refinement was performed using torsion NCS restraints with PHENIX (*127*). The unambiguous assignment of the NUP358$^{NTD}$ sequence register was further aided by anomalous difference Fourier map calculations from seventeen distinct SeMet mutant NUP358$^{NTD}$ crystals, introducing eighteen additional selenium sites (T92M, Q137M, D140M, D152M, D171M, I322M, L401M, I427M, I453M, I539M, I559M, I599M, I615M, I626M, I652M, V668M, D693M and N709M), providing a sequence register marker for every α-helix in NUP358$^{NTD}$, apart from the TPR sub-domain, for which we had previously determined a 0.95 Å resolution structure (*80*). The final structure possessed excellent stereochemistry and geometry and was refined to 3.95 Å resolution with $R_{work}$ and $R_{free}$ values of 19.2% and 24.1%, respectively. For data processing and refinement statistics, see Tables S10 to S12.

*Crystal structure determination of NUP358$^{NTD}$•sAB-14 ANE1 mutants*. Structures of NUP358$^{NTD}$•sAB-14 harboring the ANE1 T585M, T653I or I656V mutant were individually determined. The three NUP358$^{NTD}$•sAB-14 ANE1 mutant complexes were crystallized at 21 °C with the hanging drop method using 1 μl of protein solution (5 mg/ml) and 1 μl of reservoir solution, composed of 0.1 M sodium citrate (pH 4.6), 0.15 M sodium acetate, and 3.5% (w/v) PEG 4,000. Crystals were cryoprotected by exchange into reservoir solution containing 20% (v/v) ethylene glycol, prior to plunge freezing in liquid nitrogen. Native X-ray diffraction data were integrated with XDS (*117*) and scaled with AIMLESS (*119*) to 4.7 Å, 6.7 Å, and 6.7 Å resolution for the NUP358$^{NTD}$•sAB-14 T585M, T653I, and I656V ANE1 mutant, respectively. All three NUP358$^{NTD}$•sAB-14 ANE1 mutant structures were determined by molecular replacement with PHASER (*120*), using the coordinates of wildtype NUP358$^{NTD}$•sAB-14 as a search model, identifying two complexes per ASU. Refinement was performed using torsion NCS restraints with PHENIX (*127*). The final structures possessed excellent stereochemistry and geometry and were refined to $R_{work}/R_{free}$ values of 20.4/25.4% (T585M), 21.7/26.3% (T653I), and 20.5/25.4% (I656). For data processing and refinement statistics, see Table S10.

*Crystal structure determination of NUP358$^{OE}$*. NUP358$^{OE}$ was crystalized at 21 °C with the hanging drop method using 1 μl of protein solution (20 mg/ml) and 1 μl of reservoir solution, composed of 2.0 M ammonium sulfate and 0.1 M citric acid (pH 3.5). Crystals were cryoprotected by gradual exchange into reservoir solution containing 20% (v/v) ethylene glycol in 5% steps, prior to plunge freezing in liquid nitrogen. The NUP358$^{OE}$ structure was solved in PHASER (*120*) using highly redundant 1.45 Å resolution anomalous sulfur-SAD X-ray diffraction data, identifying 19 sulfur sites and four molecules per ASU. An initial model was built *de novo* into the experimental electron density map and was iteratively refined using 1.1 Å resolution native X-ray diffraction data in PHENIX (*127*). The final structure possessed ideal stereochemistry and geometry and was refined to 1.1 Å resolution with $R_{work}$ and $R_{free}$ values of 15.5% and 16.7%, respectively. For data processing and refinement statistics, see Table S13.

*Crystal structure determination of NUP358$^{ZnF}$•Ran(GDP) complexes*. NUP358$^{ZFD}$ is composed of eight ZnF modules, ZnF1 does not bind to Ran and ZnF5 and ZnF6 are identical in sequence. Initially, we determined the crystal structures of ZnF2, ZnF3, ZnF4, ZnF5/6, ZnF7, and ZnF8 in complex with Ran(GDP), containing an N-terminal $His_6$-thrombin cleavage site. Purified NUP358$^{ZnF}$ was mixed with equimolar Ran(GDP) and co-crystallized in similar conditions, containing 0.1 M Bis-TRIS (pH 6.4-7.0) and 17-22% (w/v) PEG 3,350. Crystals were cryoprotected by exchange into reservoir solution containing 20% (v/v) glycerol, prior to plunge freezing in liquid nitrogen. X-ray diffraction data were collected at the zinc edge, with data integrated using XDS (*117*) or DIALS (*118*) and scaled with AIMLESS (*119*) to resolutions in the range of 1.8-2.45 Å. The NUP358$^{ZnF}$•Ran(GDP) structures were all solved by zinc-SAD using PHASER (*120*). In the NUP358$^{ZnF2}$•Ran(GDP) structure, the N-terminal $His_6$-thrombin-tag occupied Ran's nucleotide-state independent hydrophobic surface pocket. To structurally characterize NUP358$^{ZnF2}$ interactions at this pocket, we re-crystalized NUP358$^{ZnF2}$ with Ran(GDP), using a Ran construct lacking the N-terminal $His_6$-thrombin-tag, in a reservoir solution, containing 0.1 M Bis-TRIS (pH 6.4) and 17% (w/v) PEG 3,350. All six NUP358$^{ZnF}$•Ran(GDP) structures were iteratively refined using torsion NCS restraints in PHENIX (*127*). The final structures possess excellent refinement statistics and exhibit excellent geometry. For data processing and refinement statistics, see Table S14.

*Crystal structure determination of NUP153$^{ZnF}$•Ran(GDP) complexes.* NUP153$^{ZFD}$ is composed of four ZnF modules. We determined NUP153 ZnF1, ZnF2, ZnF3, and ZnF4 in complex with Ran(GDP) containing a thrombin cleavable N-terminal $His_6$-tag. Purified NUP153$^{ZnF}$ was mixed with equimolar Ran(GDP) and co-crystallized in similar conditions, containing 0.1 M Bis-TRIS (pH 7.5) and 19% (w/v) PEG 3,350. Crystals were cryoprotected by exchange into reservoir solution, containing 20% (v/v) glycerol, prior to plunge freezing in liquid nitrogen. Anomalous X-ray diffraction data were collected at the zinc edge, integrated using XDS (*117*) and scaled with AIMLESS (*119*) to resolutions between 1.6-1.9 Å. The NUP153$^{ZnF}$•Ran(GDP) structures were all solved by zinc-SAD using PHASER (*120*). As previously observed with NUP358$^{ZnF2}$•Ran(GDP), all four NUP153$^{ZnF}$•Ran(GDP) structures determined exhibited a $His_6$-thrombin-tag bound to Ran's nucleotide-state independent hydrophobic surface pocket. To structurally characterize NUP153$^{ZnF}$ interactions with this pocket, we re-crystalized all four NUP153$^{ZnF}$ domains in complex with Ran(GDP), using a Ran construct lacking the N-terminal thrombin cleavable His$_6$-tag, in reservoir solution, containing 0.1 M Bis-TRIS (pH 6.0-7.0) and 21% (w/v) PEG 3,350. Crystals were cryoprotected by exchange into reservoir solution containing 20% (v/v) glycerol, prior to plunge freezing in liquid nitrogen. The structures were determined by molecular replacement with the equivalent NUP153$^{ZnF}$•Ran(GDP) structure containing the His$_6$-thrombin-tag as a search model using PHASER (*120*). We report two NUP153$^{ZnF3}$•Ran(GDP) structures, one at 2.05 Å resolution with data completeness of 93.2%, where specific flexible regions of the structure could not be modeled accurately, and a complete 2.4 Å resolution dataset with additional electron density features. All five NUP153$^{ZnF}$•Ran(GDP) structures were iteratively refined using torsion NCS restraints in PHENIX (*127*). The final structures exhibit excellent refinement statistics and stereochemistry. For data processing and refinement statistics, see Table S15.

*Structure determination of NUP50$^{RanBD}$•Ran(GMPPNP) complex.* SeMet-labeled Nup50$^{RBD}$ was complexed with nucleotide exchanged Ran(GMPPNP) and crystalized at 21 °C with the hanging drop method using 1 μl of protein solution (20 mg/ml) and 1 μl of reservoir solution, containing 0.1 M HEPES (pH 7.5), 0.2 M ammonium sulfate, and 20% (w/v) PEG 3,350. Crystals were cryoprotected by exchanging the crystallization buffer with reservoir solution containing 20% (v/v) ethylene glycol, prior to plunge freezing in liquid nitrogen. Anomalous X-ray diffraction data was collected at the selenium edge and processed to 2.45 Å resolution with XDS (*117*) and AIMLESS (*119*). The NUP50$^{RBD}$•Ran(GMPPNP) structure was solved by selenium-SAD using PHASER (*120*), identifying 14 selenium sites and two complexes per ASU. Refinement was performed using torsion NCS restraints with PHENIX (*127*). The final structure possessed excellent stereochemistry and geometry and was refined to 2.45 Å resolution with $R_{work}$ and $R_{free}$ values of 21.2% and 23.9%, respectively. For data processing and refinement statistics, see Table S16.

*Crystal structure determination of NUP358$^{RanBD}$•Ran(GMPPNP) complexes.* NUP358 contains four RanBD domains NUP358$^{RanBD-I}$, NUP358$^{RanBD-II}$, NUP358$^{RanBD-III}$, and NUP358$^{RanBD-IV}$. Crystals of NUP358$^{RanBD-I}$•Ran(GMPPNP) were grown in a reservoir solution containing 0.2 M potassium sodium tartrate tetrahydrate and 20% (w/v) PEG 3,350, and were cryoprotected by transfer in reservoir solution supplemented with 20% (v/v) glycerol. Crystals of NUP358$^{RanBD-II}$•Ran(GMPPNP) were grown in 125 mM magnesium formate and 15% (w/v) PEG 3,350, and were cryoprotected by transfer in reservoir solution supplemented with 20% (v/v) ethylene glycol. Crystals of NUP358$^{RanBD-III}$•Ran(GMPPNP) were grown in 0.2 M sodium formate and 20% (w/v) PEG 3,350, and were cryoprotected by transfer in reservoir solution supplemented with 25% (v/v) ethylene glycol. All crystals were frozen by plunge freezing in liquid nitrogen. Despite extensive efforts NUP358$^{RanBD-IV}$ initially failed to yield well diffracting crystals. A sequence alignment of all NUP358 RanBD identified a poorly conserved 8-residue surface loop in NUP358$^{RanBD-IV}$ (residues 2962-2969; WHTMKNYY) that was mutated to the corresponding region of NUP358$^{RanBD-III}$ (residues 2359-2366; QNYDNKQV). The modified NUP358$^{RanBD-IV}$•Ran(GMPPNP) complex crystallized in a reservoir solution, containing 0.1 M HEPES (pH 7.0), 0.1 M ammonium sulfate, and 20% (w/v) PEG 3,350. Crystals were cryoprotected by transfer in reservoir solution supplemented with 20% (v/v) ethylene glycol, prior to plunge freezing in liquid nitrogen. Native X-ray diffraction data was processed with XDS/AIMLESS (*117, 119*) for RanBD-I and RanBD-IV at 2.4 Å and 2.7 Å resolution, and with DIALS/AIMLESS (*118, 119*) for RanBD-II and RanBD-III at 2.4 Å and 2.7 Å resolution, respectively. The NUP358$^{RanBD}$•Ran(GMPPNP) structures were determined by molecular replacement using the coordinates of NUP50$^{RanBD}$•Ran(GMPPNP) as a search model in PHASER (*120*). Refinement was performed using torsion NCS restraints with PHENIX (*127*). The final structures possess excellent refinement statistics and exhibit excellent geometry. For data processing and refinement statistics, see Table S16.

*Crystal structure determination of NUP88$^{NTD}$•NUP98$^{APD}$*. NUP88$^{NTD}$•NUP98$^{APD}$ was crystalized at 21 °C by the hanging drop method using 1 μl of protein solution (15 mg/ml) and 1 μl of reservoir solution, consisting of 0.1 M TRIS (pH 8.3), 0.22 M NaCl, and 20% (w/v) PEG 4,000. Crystals were cryoprotected by exchanging the crystallization buffer with paratone-N, prior to plunge freezing in liquid nitrogen. X-ray diffraction data were collected at beamline 23ID-B at the Advanced Photon Source (APS). Native data was integrated with XDS (*117*) and scaled with AIMLESS (*119*) to 2.0 Å resolution. The

NUP88$^{NTD}$•NUP98$^{APD}$ structure was determined by molecular replacement using the coordinates of the *C. thermophilum* Nup82$^{NTD}$ (PDB ID 5CWW) (*11*) and human NUP98$^{APD}$ (PDB ID 1KO6) (*13*) as search models in PHASER (*120*). Refinement was performed using torsion NCS restraints in PHENIX (*127*). The final model possessed excellent stereochemistry and geometry and was refined to 2.0 Å resolution with $R_{work}$ and $R_{free}$ values of 17.5% and 20.8%, respectively. For data processing and refinement statistics, see Table S17.

**Quantitative docking of NUP358$^{NTD}$ into intact human NPC cryo-ET maps**

NUP358$^{NTD}$ open and closed conformation structures were searched for in the ~12 Å cryo-ET map of the intact human NPC (*46*), as part of an incremental docking procedure outlined in the accompanying manuscript (*42*). Briefly, we used the Chimera fitmap tool to generate 1 million random placements of volumes simulated from the NUP358$^{NTD}$ crystal structures and low pass filtered to the resolution of the cryo-ET map (*129*). Placements were limited to an asymmetric subunit of the C8 symmetry averaged cryo-ET map from which nuclear envelope regions had been eliminated by segmentation, and a volume overlap cutoff of 20% was employed to limit the random placements to map density. To evaluate the docking within a statistical framework, an analysis similar to previous statistical approaches, docking residue-level resolution structures into lower resolution cryo-ET maps, was applied (*30*). After local rigid body optimization, correlations about the mean (Pearson correlations) were calculated for every placed volume and the map, and then Fisher z-transformed and normalized to approximate a normal sampling distribution. Because correct placements are rare events, we used the Fisher z-score distribution as a null distribution from which to calculate one-tailed p-values for the top-scoring placements.

Five copies of the NUP358$^{NTD}$ open conformation were identified as top scoring placements per human NPC spoke, for a total of 40 copies per NPC. Whereas the 1st, 3rd, 4th, and 5th highest scoring solutions corresponded to NUP358$^{NTD}$ copies that wrap around the stalk of the Y-shaped CNCs, the 2nd highest scoring solution identified the dome NUP358$^{NTD}$ copy that sits on top of the other four NUP358$^{NTD}$ copies. All docking solutions identified for the NUP358$^{NTD}$ closed conformation fell below the Pearson correlation score and p-value of the accepted NUP358$^{NTD}$ open conformation placements.

For reference, we performed a similar analysis by docking a resolution-matched simulated density of the NUP358$^{NTD}$ open conformation into a previously available ~23 Å cryo-ET map of an intact human NPC (*44*). NUP358$^{NTD}$ placements from the docking in the ~12 Å cryo-ET map corresponded to top scoring solutions (1st, 3rd, 4th, and 5th), except for the placement at the proximal inner position, which was scored 53$^{rd}$ highest. Overall, the placement confidence as estimated by p-values was considerably lower for the NUP358$^{NTD}$ open conformation docked in the ~23 Å cryo-ET map compared to the ~12 Å cryo-ET map.

**Placement of cytoplasmic filaments into intact human NPC cryo-ET map**

After docking of five copies of NUP358$^{NTD}$ into the unexplained density cluster I, we attempted to explain unexplained density cluster II by quantitatively docking NUP88$^{NTD}$•NUP98$^{APD}$, NUP214$^{NTD}$•DDX19 (PDB ID 3FMO) (*55*), GLE1$^{CTD}$•NUP42$^{GBM}$ (PDB ID 6B4F) (*63*), RAE1•NUP98$^{GLEBS}$ (PDB ID 3MMY) (*57*), NUP358$^{RanBD-II}$•Ran(GMPPNP), NUP358$^{ZnF3}$•Ran(GTP), and NUP358$^{CTD}$ (PDB ID 4I9Y) (45) structures into the ~12 Å cryo-ET map of the intact human NPC after all the hereto explained density has been subtracted, but could not confidently assign either to the density. We proceeded to interpret the two nearly perpendicular segments of tube-shaped density by manually placing individual coiled-coil segments of a poly-alanine model derived from the *X. laevis* NUP54•NUP58•NUP62 (PDB ID 5C3L) (*73*) and *C. thermophilum* Nup49•Nup57•Nsp1•Nic96 (PDB ID 5CWS) (*78*) complexes, in lieu of an experimental structure of the CFNC-hub. Based on shape complementarity and the distance restraint imposed by its interaction with NUP214$^{TAIL}$, we placed the NUP88$^{NTD}$•NUP98$^{APD}$ into the disk-shaped density at the base of the long tube-like density assigned to the N-terminal segment of the CFNC-hub coiled-coil and adjacent to the NUP75 arm of the proximal CNC. We placed the NUP214$^{NTD}$•DDX19 crystal structure into the dumbbell-shaped density that protrudes into the central transport channel, thus explaining all the density in the unexplained density cluster II. All docked structures were locally rigid body fit into the cryo-ET density, but the accuracy of the orientation is inherently limited by the lack of shape features in the structures and the resolution of the cryo-ET map.

**Estimation of the root-mean square length of flexible NUP93 linker**

The linker spanning the region between the NUP205-binding $NUP93^{R2}$ region and the CFNC-hub-binding $NUP93^{R1}$ region is ~45 residues in length and predicted to be devoid of secondary structure. To estimate the expected distance at which the $NUP93^{R1}$ region is found, we calculated the root-mean square (r.m.s) end-to-end length of a linear polymer chain according to the Gaussian chain model developed by Flory, with a characteristic ratio parameter of ~3, typical for unstructured polypeptide linkers (*130, 131*).

**Placement of $GLE1^{CTD}$•$NUP42^{GBM}$ into the ~12 Å intact human NPC cryo-ET map**
Due to the small size and lack of distinct shape features, the quantitative docking of the *H. sapiens* $GLE1^{CTD}$•$NUP42^{GBM}$ crystal structure (PDB 6B4F) (*63*) into the ~12 Å cryo-ET map of the intact human NPC after all the hereto explained density has been subtracted did not result in high confidence placement. Nevertheless, we identified unexplained density between the cytoplasmic NUP155 bridge and the cytoplasmic face of the nuclear envelope that could accommodate the $GLE1^{CTD}$•$NUP42^{GBM}$ structure. The placement is supported by the proximity of a mutually exclusive binding site for $NUP98^{R3}$ and $GLE1^{NTD}$ on the bridge NUP155 surface (*63*).

**Placement of ELYS into the ~12 Å cryo-ET map of the intact human NPC**
Due to the small size and lack of distinct shape features, the quantitative docking of the *M. musculus* $ELYS^{NTD}$ domain (PDB 4I0O) (*25*) into the ~12 Å cryo-ET map of the intact human NPC after all the hereto explained density has been subtracted did not result in high confidence placements. Nevertheless, we identified unexplained density in the shape of a crescent with a disk at the base in the nuclear outer rings alongside NUP160. We assigned $ELYS^{NTD}$ $\beta$-propeller domains in the disk-like shape, seemingly interfacing with the nuclear envelope at the base of proximal and distal NUP160 copies. The crescent-like densities are consistent with the $\alpha$-solenoid ELYS domain (residues 495-1018). The anchoring of 32 copies of ELYS by binding to NUP160 in the nuclear outer ring of the human NPC is supported by biochemical and *in vivo* evidence of interaction between ELYS and NUP160 homologs from different species (*24, 25, 132*).

**Docking into the anisotropic single particle cryo-EM map of the *X. laevis* cytoplasmic outer ring**
As in the accompanying manuscript, the anisotropic 5.5-7.9 Å composite single particle cryo-EM map of the *X. laevis* cytoplasmic outer ring protomer (EMD-0909) (*45*) was superposed to the ~12 Å cryo-ET map of the intact human NPC (EMD-XXXX) (*46*). A cytoplasmic outer ring protomer composite structure was then further refined by rigid-body fitting the individual nup and nup complex structures into the 5.5-7.9 Å composite single particle cryo-EM map of the *X. laevis* cytoplasmic outer ring protomer. The map did not include regions corresponding to the proximal copy of $NUP93^{SOL}$•$NUP53^{R2}$ and $NUP214^{NTD}$•DDX19, presumably due to omission by the masking applied during map processing.

**Docking into the dilated ~37 Å *in situ* cryo-ET map of the human NPC**
As in the accompanying manuscript, a composite structure of a spoke of the cytoplasmic outer ring built into the constricted ~12 Å cryo-ET sub-tomogram averaged human NPC map (EMD-XXXX) (*46*), including the placed asymmetric nucleoporins, was manually placed in the dilated *in situ* ~37 Å cryo-ET sub-tomogram averaged human NPC map (EMD-11967) (*42, 101*). The structure fit into the dilated human NPC ~37 Å cryo-ET map was further locally refined as two rigid bodies: all distal subunits, and all proximal subunits together with the CFNC-hub model, $NUP88^{NTD}$•$NUP98^{APD}$, and $NUP214^{NTD}$•DDX19 structures. A C8 symmetry operation was applied to place the docked spoke to the rest of the dilated human NPC ~37 Å cryo-ET map.

**Plasmid construction for AID-tagged NUP358 and NUP160**
The CRISPR/Cas9 gene-editing technique was utilized to tag the N-terminus of NUP358 and the C-terminus of NUP160 at the endogenous loci. Two gRNA plasmids were generated per construct and integrated into pX330 vectors, as previously described (*133*). Guide RNAs were selected using CRISPR Design Tools (*133*). For AID-tagged NUP358, the guide RNA primers: 5'-caccgCCACCCCGTCGCCTC GAC-3', 5'-aaacGTCGAGGCGACGGGGTGGc-3', 5'-caccgAGCAAGGCTGACGTGGAG-3', 5'-aaacCT CCACCAGCCTTGCTc-3' were used. To induce the desired DNA sequence into CRISPR/Cas9-cleaved genomic regions via homology-mediated recombination, the donor vector, pCassette-Hyg-P2A-microAID-NUP358 was built from pCassette (*134*). Briefly, the homology arms are amplified from genomic DNA extract from HCT116 and DLD1 cells. The DNA sequences of hygromycin, P2A, and

micro-AID degron encoding residues 71-114 from plant AID sequence were cloned into pCassette vector between Asc1 and Sbf1 restriction sites. For AID-tagged NUP160, the gRNA plasmids were generated, as previously described (*135*). The donor vector pCassette-NUP160-NeonGreen-AID-P2A-Hyg was built from pCassette (*135*). The sequences of homology arms were amplified from genomic DNA extracted from DLD1 cells. The DNA sequence of NeonGreen fluorescent protein, hygromycin, P2A, and three copies of reduced AID tag (65-132 amino acids) were used. All PCR reactions were performed using Platinum SuperFi DNA Polymerase (ThermoFisher).

**Construction of cell lines**

Ten thousand cells per well were plated in 24-well plates one day before transfection. Plasmids for transfection were purified using the NucleoSpin buffer set (Clontech) and Nucleic acid miniprep columns (VitaScientifc). Plasmids were not linearized before transfection. Cells were transfected with 500 ng of donor and sgRNA plasmids in ratio 1:1 using ViaFect (Promega) transfection reagent according to the manufacturer's instructions. Cells were seeded on 100 mm cell culture dishes with the selective antibiotics (hygromycin 200 μg/ml) in complete medium after 72 hours post-transfection and cultured until clones were visualized on a plate. Clones were picked and propagated in regular complete media without selective antibiotics. *AID*::*NUP358* HCT116, *AID*::*NUP358* DLD1, and *NUP160*::*NG-AID* DLD1 cell lines were generated by CRISPR/Cas9. The insertion of the microAID tag was verified by genomic PCR using oligos: 5'-TGTGGT CTTTTTCATTATGCAGTTC-3' and 5'-AACTGGCCCCAAATACCCAG-3'. The insertion of plant E3 ubiquitin ligase, TIR1, in the *AID*::*NUP358* HCT116, *AID*::*NUP358* DLD1, and *NUP160*::*NG-AID* DLD1 cell lines was confirmed, as previously described (*134*). Briefly, using CRISPR/Cas9, C-terminus of RCC1 was tagged with sequences encoding infra-red fluorescent protein (iRFP670) and TIR1-9Myc, separated by a P2A sequence. The fusion protein is cleaved during translation to yield $RCC1^{iRFP670}$ and TIR1-9myc. Localization and expression of targeted NUP358 and NUP160 were validated using immunofluorescence microscopy and western blot.

**Cell culture and synchronization**

*AID*::*NUP358* HCT116 cells were grown in McCoy's 5A medium containing 10% fetal bovine serum (FBS), 100 U/ml penicillin, and 100 μg/ml streptomycin. *AID*::*NUP358* DLD1 and *NUP160*::*NG-AID* DLD1 cells were grown in Dulbecco's Modified Eagle Medium (DMEM) containing 10% FBS, 100 U/ml penicillin, and 100 μg/ml streptomycin. Cells were cultured in 100 mm cell culture dishes at 37 °C in 5% $CO_2$/95% humidified incubator. Cell cultures were passaged every 3-4 days using a 0.25% trypsin and 0.05% EDTA solution in phosphate buffered saline (PBS), containing 137 mM NaCl, 2.7 mM KCl, 8 mM sodium hydrogen phosphate and 2 mM potassium dihydrogen phosphate. For NUP358 and NUP160 depletion, the media was supplemented with 1 mM 3-indoleacetic acid (auxin; Sigma-Aldrich), added from a 250 mM auxin-solution in PBS. For synchronization, *AID*::*NUP358* HCT116 cells were seeded in 24-well plates and grown to ~25% confluency. Nocodazole (Sigma-Aldrich) was then supplemented to the cells with a final concentration of 50 ng/ml. Following 12 hours of incubation, nocodazole-containing medium was replaced with fresh 37 °C prewarmed medium. Cell populations at this point were mitotically enriched.

**Immunofluorescence microscopy**

For immunofluorescence, different NUP358 variants were cloned into a modified pcDNA3.1 vector (Invitrogen) containing an N-terminal 3×HA-tag using BamHI and NotI restriction sites. For a summary of all mammalian expression constructs, see Table S4. *AID*::*NUP358* HCT116 cells were grown on Poly-L-lysine coated slides until ~50% confluency. Transfection was performed with Lipofectamine 2000 transfection reagent (Invitrogen) according to the manufacturer's instructions. After 10 hours, the media was removed, and cells are washed in PBS, and fixed in PBS supplemented with 2% (w/v) paraformaldehyde (PFA) for 5 minutes at room temperature. After two washes with PBS, the cells were permeabilized with PBS containing 0.1% (v/v) TritonX-100 (Sigma-Aldrich) for 10 minutes at room temperature. The cells were then washed in PBS and blocked in PBS supplemented with 10% (v/v) FBS for 20 minutes at room temperature. For nuclear envelope staining, the cells were incubated with a 1:1,000 dilution of the monoclonal antibody mAb414 in PBS buffer, supplemented with 0.1% (w/v) saponin and 10% (v/v) FBS for 1 hour at room temperature, followed by three washes with PBS supplemented with 10% (v/v) FBS for 10 minutes at room temperature. Secondary antibody incubation was performed with a 1:3,000 dilution of Alexa Fluor 568-labeled goat anti-mouse antibody (Invitrogen) in PBS, supplemented with 0.1% (w/v) saponin and 10% (v/v) FBS for 1 hour at room temperature, followed by three washes with PBS supplemented with 10% (v/v) FBS for 10 minutes at room temperature. For detection of the HA-tagged proteins, the cells were incubated with a mouse monoclonal anti-HA Fluor488 conjugate antibody (Invitrogen) at 1:2,000 dilution for 16 hours at 4°C, followed by three washes with PBS supplemented with 10% (v/v) FBS for 10 minutes at room temperature. The cells were mounted onto coverslips with ProLong Gold Antifade reagent with DAPI (Invitrogen). Slides were examined by fluorescence microscopy on a Carl Zeiss AxioImagerZ.1 equipped with an AxioCamMRm camera.

The synchronized *AID*::*NUP358* HCT116 cells were treated with 1 mM auxin at 37 °C. The localization of different nups was examined by immunofluorescence after 0, 2, 4, 6, 8, 10, and 24 hours. The immunofluorescence analysis was performed with a rabbit polyclonal anti-NUP358 antibody (Bethyl Laboratories; 1:500 dilution), a rabbit polyclonal anti-NUP214 antibody (Abcam; 1:1,000 dilution), a mouse monoclonal anti-NUP88 antibody (Santa Cruz; 1:250 dilution), a rat monoclonal anti-NUP98 antibody (Abcam; 1:500 dilution), a rabbit polyclonal anti-GLE1 antibody (Abcam; 1:500 dilution), a mouse monoclonal mAb414 antibody (BioLegend; 1:1,000 dilution), a rabbit polyclonal anti-NUP160 antibody (St John's Lab; 1:500 dilution), a rabbit polyclonal anti-NUP96 antibody (Bethyl; 1:500 dilution), a rabbit polyclonal anti-NUP133 antibody (Bethyl; 1:500 dilution), and a mouse monoclonal anti-NUP93 antibody (Santa Cruz; 1:200 dilution). Secondary antibody incubation was performed with an Alexa Fluor 488-labeled donkey anti-rabbit antibody (Invitrogen, 1:3,000 dilution), an Alexa Fluor 568-labeled goat anti-mouse antibody (Invitrogen, 1:3,000 dilution), or an Alexa Fluor 488-labeled goat anti-rat antibody (Invitrogen, 1:3,000 dilution).

**Cell fractionation**

Synchronized *AID*::*NUP358* HCT116 cells were treated with 1 mM auxin at 37 °C and after 0, 2, 4, 6, 8, 10 and 24 hours cells were trypsinized, washed twice with cold PBS, and harvested by centrifugation at 400 × *g* for 5 minutes at 4 ºC. $5\times10^5$ cells were pelleted and resuspended in 35 μL lysis buffer, containing 10 mM TRIS (pH 8.0), 320 mM sucrose, 3 mM calcium chloride, 2 mM magnesium acetate, 0.1 mM EDTA, 0.05% Triton X-100, 1 mM DTT, and Complete protease inhibitor cocktail for 15 minutes on ice and spun at 500 × *g* for 5 minutes at 4 ºC. The supernatant was kept as the cytoplasmic fraction and then spun at 30,000 × *g* for 15 minutes to clarify, diluted with SDS-PAGE loading dye. Pelleted nuclei were washed with a buffer containing 10 mM TRIS (pH 8.0), 320 mM sucrose, 3 mM calcium chloride, 2 mM magnesium acetate, 0.1 mM EDTA, 1 mM DTT, and Complete protease inhibitor cocktail and spun at 500 × *g* for 5 minutes at 4 ºC. The supernatant was aspirated, and the pelleted nuclei were resuspended in SDS-PAGE loading dye. Cytoplasmic and nuclear fractions were boiled at 95 °C for 10 minutes before SDS-PAGE analysis.

**Reporter expression analysis**

The generation of cDNAs encoding insulin, green fluorescence protein (GFP), and histone 1B (H1B) was previously described (*76*). The cDNAs encoding anti-inflammatory cytokine interleukin-10 (IL-10), pro-inflammatory cytokine IL-6, tumor necrosis factor alpha (TNFα), membrane-bound alkaline phosphatase placental (ALPP), and large ribosomal subunit protein 26 (RPL26) were obtained commercially (DNASU, Arizona State University; Horizon Discovery Biosciences). DNA fragments encompassing the entire coding regions of insulin, GFP, H1B, IL-10, IL-6, TNFα, ALPP, and RPL26 were amplified by PCR and cloned into a modified pcDNA3.1 vector (Invitrogen) containing a C-terminal 3×FLAG-tag using the KpnI restriction site (*136*). To avoid transfection differences arising from auxin treatment, we transfected reporter constructs prior to auxin induced degradation of NUP358. Specifically, four hours after release from mitotic arrest synchronized *AID*::*NUP358* HCT116 cells were transfected with reporter constructs using Lipofectamine 2000, according to the manufacturer's instructions. After 10 hours, the cells were treated with 1 mM auxin in fresh media at 37 °C. After 0, 2, 4, 6, 8, 10, or 24 hours, cells were trypsinized, washed twice with cold PBS, and harvested by centrifugation at 400 × *g* for 5 minutes at 4 ºC. Cell pellets were resuspended in 2×SDS-PAGE loading dye and boiled at 95 °C for 10 minutes prior to loading on an SDS-PAGE gel. The relative expression of reporter from auxin treated or untreated cells was quantitated at the 10-hour time point. Bands were quantitated in Fiji and plotted as a percentage against untreated cells (*137*). The statistical analysis was carried out using three sets of independent experiments for every analyzed sample. Error bars represent standard error.

**Western blot analysis**

Expression levels of 3×HA-NUP358, and C-terminal 3×FLAG tagged reporter constructs were assessed by western blot analysis. The transfected cells were trypsinized, washed twice with cold PBS, and harvested by centrifugation at 400 × *g* for 5 minutes at 4 ºC. Cell pellets were resuspended in 2×SDS-PAGE loading dye and boiled at 95 °C for 10 minutes prior to loading on an SDS-PAGE gel. Western blot analyses were performed with a mouse monoclonal anti-$\alpha$-tubulin (Sigma-Aldrich; 1:5,000 dilution), a mouse monoclonal anti-HA (Covance, 1:4,000 dilution), and a mouse monoclonal anti-FLAG (Sigma-Aldrich, 1:4,000 dilution) antibodies, then visualized with and a goat anti-mouse antibody fused to an IR800 fluorescent protein (Licor, 1:5,000 dilution). The imaging was carried out with an IR imager (Li-Cor Odyssey) using the 800 nm channels in a single scan at 169 $\mu$m resolution and a scan intensity of 5. Antibodies were diluted in Superblock-PBS blocking buffer (Thermo Fisher) and washes were carried out in PBS supplemented with 0.05% (v/v) Tween-20 (PBS-T).

Both cytoplasmic and nuclear fractions were sampled from auxin treated synchronized *AID*::*NUP358* HCT116 cells at different times, as described above. The expression levels of different nups were examined by western blot, using a rabbit polyclonal anti-NUP358 (Bethyl Laboratories; 1:1,000 dilution), a rabbit polyclonal anti-NUP214 (Abcam; 1:2,000 dilution), a mouse monoclonal anti-NUP88 (Santa Cruz; 1:500 dilution), a rat monoclonal anti-NUP98 (Abcam; 1:1,000 dilution), a rabbit anti-GLE1 (Abcam; 1:1,000 dilution), a mouse monoclonal mAb414 (BioLegend; 1:3,000 dilution), a rabbit polyclonal anti-NUP160 (St John's Lab; 1:500 dilution), a rabbit polyclonal anti-NUP96 (Bethyl; 1: 1,000 dilution), a rabbit polyclonal anti-NUP133 (Bethyl; 1:1,000 dilution), a rabbit polyclonal anti-NUP43 (gift of Valerie Doye, CRNS Paris, France; 1:1,000 dilution), a rabbit polyclonal anti-ELYS (gift of Iain Mattaj, EMBL Heidelberg, Germany; 1:3,000 dilution), a rabbit polyclonal anti-NUP205 (gift of Ulrike Kutay, ETH Zurich, Switzerland; 1:2,000 dilution), a mouse polyclonal anti-NUP93 (Santa Cruz; 1:200 dilution), a rabbit polyclonal anti-LaminA/C (Abcam; 1:5,000 dilution), a rabbit polyclonal anti-TBP (Abcam; 1:1,000 dilution), and a mouse monoclonal anti-$\alpha$-tubulin (Sigma-Aldrich; 1:5,000 dilution) antibodies, visualized with goat anti-mouse, goat anti-rabbit, and goat anti-rat secondary antibodies fused to an IR800 fluorescent protein (Licor, 1:5,000 dilution). Antibodies were diluted in Superblock-PBS and washes were carried out in PBS-T.

**Pulse-chase RNA export assay**

For NUP358/NUP160 depletion, the media of synchronized *AID*::*NUP358* HCT116, and unsynchronized *AID*::*NUP358 DLD1* or *NUP160*::*NG-AID* DLD1 cells was supplemented with 1 mM auxin, added from a 250 mM auxin-solution in PBS. Control cells were supplemented with PBS buffer only. After 3 hours, the media was exchanged for DMEM-10% (v/v) FBS, supplemented with 0.5 mM 5-ethynyl uridine (5-EU; Invitrogen) and 1 mM auxin. After 1-hour pulse-labeling of newly synthesized RNA, the media was exchanged for fresh DMEM-10% (v/v) FBS, supplemented with auxin, and either fixed immediately, using a 4% (w/v) PFA solution in PBS, or fixed after an additional 1-, 2-, 3-, 4-, or 5-hour chase. The fixed cells were washed with PBS and permeabilized using 0.1% (v/v) Triton X-100 for 15 minutes at room temperature. 5-EU labeling of cells was visualized according to the manufacturer's instructions (Invitrogen, Click-iT RNA imaging kits). Briefly, the samples were incubated for 30 minutes at room temperature with a 1×Click-iT reaction cocktail, containing Alexa Fluor 594 azide and copper sulfate. After removal of the reaction cocktail, cells were washed once with Click-iT reaction rinse buffer. After this step, samples were processed further for NUP358 or NUP160 antibody staining. The statistical analysis was carried out using three sets of independent images with at least 200 cells each for every analyzed sample. Error bars represent standard error.

**Thermostability assay**

10 $\mu$g of purified NUP358$^{NTD}$, wildtype or harboring ANE1 mutations, were incubated in a total volume of 50 $\mu$l for 20 minutes at 30-43 ºC, increased in 1 ºC increments. An additional titration was performed for NUP358$^{NTD}$ ANE1 W681C and wildtype protein from 15-33 ºC, increased in 3 ºC increments. Soluble and pellet fractions were separated by centrifugation at 30,000 × *g* for 15 minutes at 4 ºC, resolved by SDS-PAGE, and visualized by Coomassie brilliant blue staining. Bands were quantitated in Fiji and

plotted as a percentage of the soluble fraction per temperature point (*137*).

**Illustrations and figures**

Gel filtration profiles and MALS graphs were generated in IGOR (WaveMetrics) and assembled in Illustrator (Adobe). Sequence alignments were generated using ClustalX (*138*) and colored with ALSCRIPT (*139*). Electrostatic potentials were calculated with APBS (Adaptive Poisson-Blotzmann Solver) software (*140*). Structure figures were generated with PyMOL (www.pymol.org).

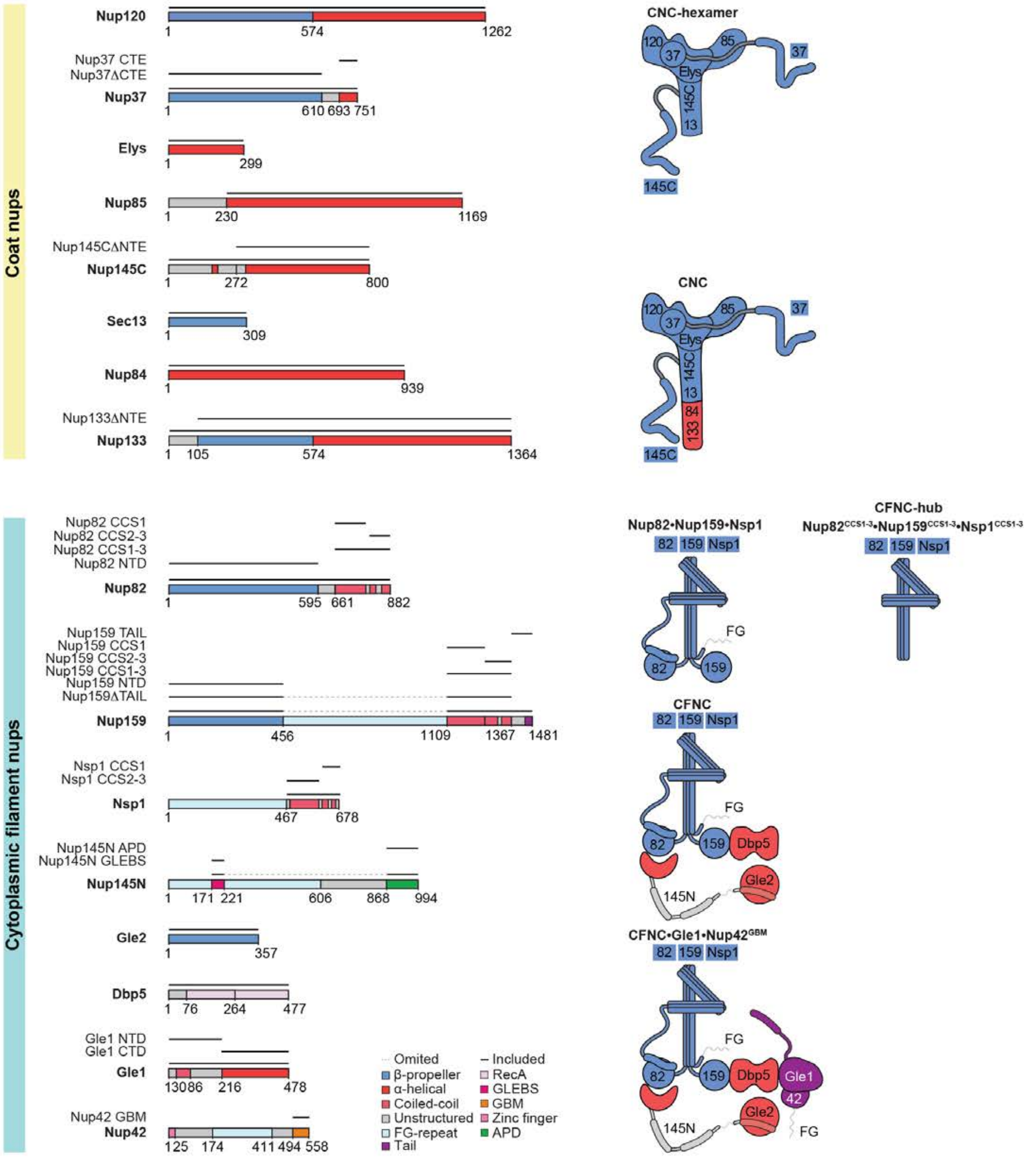
Coat nups
Nup120
1 574 1262
Nup37 CTE
Nup37ΔCTE
Nup37
1 610 693 751
Elys
1 299
Nup85
1 230 1169
Nup145CΔNTE
Nup145C
1 272 800
Sec13
1 309
Nup84
1 939
Nup133ΔNTE
Nup133
1 105 574 1364
CNC-hexamer
CNC
Cytoplasmic filament nups
Nup82 CCS1
Nup82 CCS2-3
Nup82 CCS1-3
Nup82 NTD
Nup82
1 595 661 882
Nup159 TAIL
Nup159 CCS1
Nup159 CCS2-3
Nup159 CCS1-3
Nup159 NTD
Nup159ΔTAIL
Nup159
1 456 1109 1367 1481
Nsp1 CCS1
Nsp1 CCS2-3
Nsp1
1 467 678
Nup145N APD
Nup145N GLEBS
Nup145N
1 171 221 606 868 994
Gle2
1 357
Dbp5
1 76 264 477
Gle1 NTD
Gle1 CTD
Gle1
1 30 86 216 478
Nup42 GBM
Nup42
1 25 174 411 494 558
Omited
β-propeller
α-helical
Coiled-coil
Unstructured
FG-repeat
Tail
Included
RecA
GLEBS
GBM
Zinc finger
APD
Nup82•Nup159•Nsp1
CFNC-hub
Nup82CCS1-3•Nup159CCS1-3•Nsp1CCS1-3
CFNC
CFNC•Gle1•Nup42GBM

**Fig. S1. *C. thermophilum* nucleoporin fragments.** Domain architectures of the *C. thermophilum* coat and cytoplasmic filament nups. Domains are drawn as horizontal boxes with residue numbers indicated and their observed or predicted folds colored according to the legend. The black lines indicate the fragments used throughout the text. Dashed lines indicate regions excluded from expression constructs. As a reference, schematics of nup complexes are shown on the right.

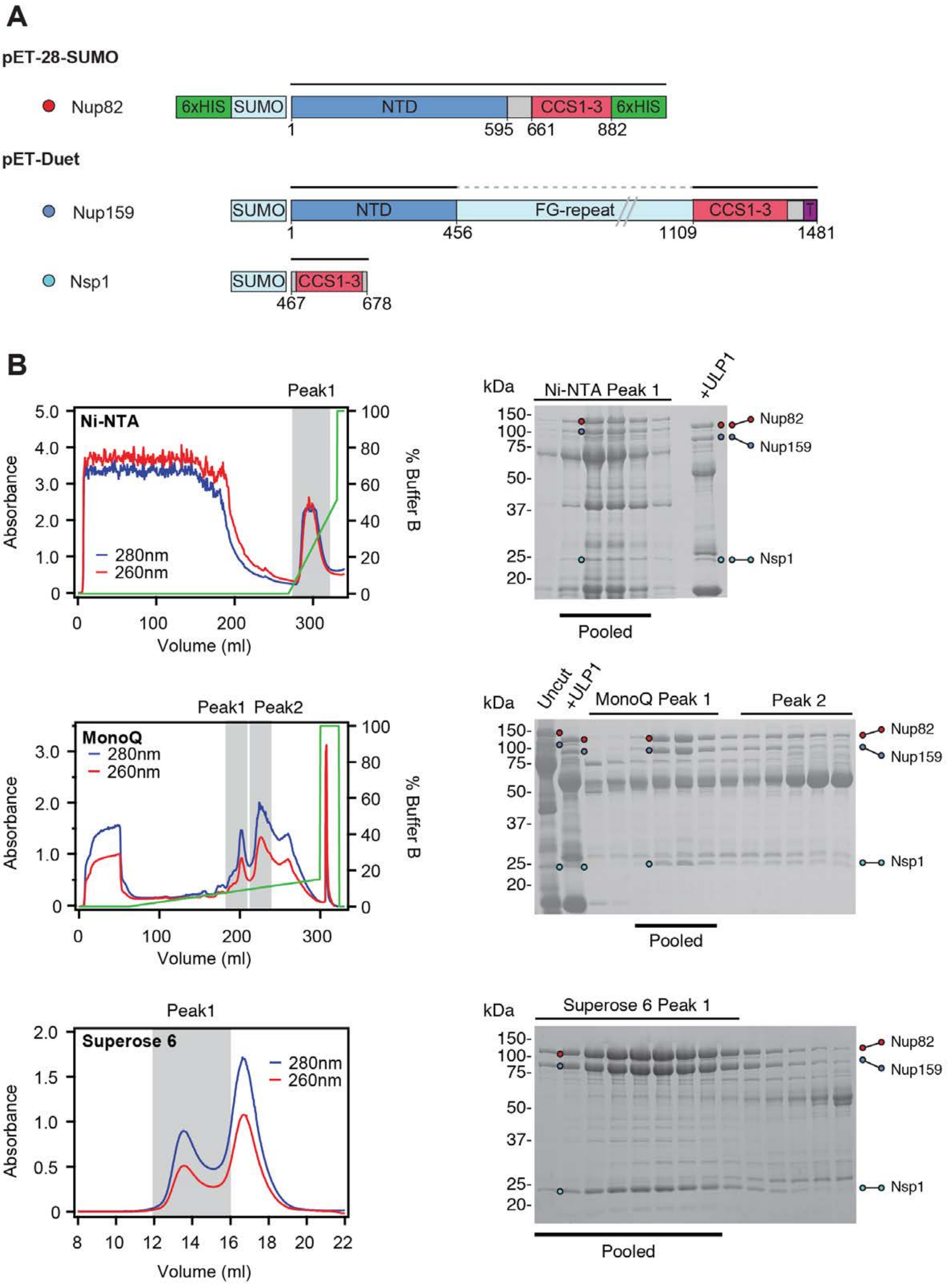
A
pET-28-SUMO
Nup82
6xHIS
SUMO
NTD
CCS1-3
6xHIS
1
595
661
882
pET-Duet
Nup159
SUMO
NTD
FG-repeat
CCS1-3
T
1
456
1109
1481
Nsp1
SUMO
CCS1-3
467
678
B
Peak1
Ni-NTA
Absorbance
% Buffer B
280nm
260nm
Volume (ml)
kDa
Ni-NTA Peak 1
+ULP1
Nup82
Nup159
Nsp1
Pooled
Peak1 Peak2
MonoQ
280nm
260nm
Absorbance
% Buffer B
Volume (ml)
kDa
Uncut
+ULP1
MonoQ Peak 1
Peak 2
Nup82
Nup159
Nsp1
Pooled
Peak1
Superose 6
280nm
260nm
Absorbance
Volume (ml)
kDa
Superose 6 Peak 1
Nup82
Nup159
Nsp1
Pooled

**Fig. S2. Purification of the *C. thermophilum* Nup82•Nup159•Nsp1 hetero-trimer.** (**A**) Domain boundaries of the purified nups are shown with black lines indicating the construct boundaries of Nup82, Nup159, and Nsp1 in our bacterial expression constructs. The dashed line indicates a central FG-repeat region in Nup159 that was excluded. (**B**) Nup82•Nup159•Nsp1 hetero-trimer purification. Sequential chromatography purification steps are shown from top to bottom with the employed columns indicated. The gray boxes indicate fractions that were resolved on SDS-PAGE gels and visualized by Coomassie staining. Pooled fractions are indicated with a black bar below the SDS-PAGE gels. For details of the buffer conditions see Table S4.

**Reconstitution and biochemical dissection of the cytoplasmic filament nup complex (CFNC)**

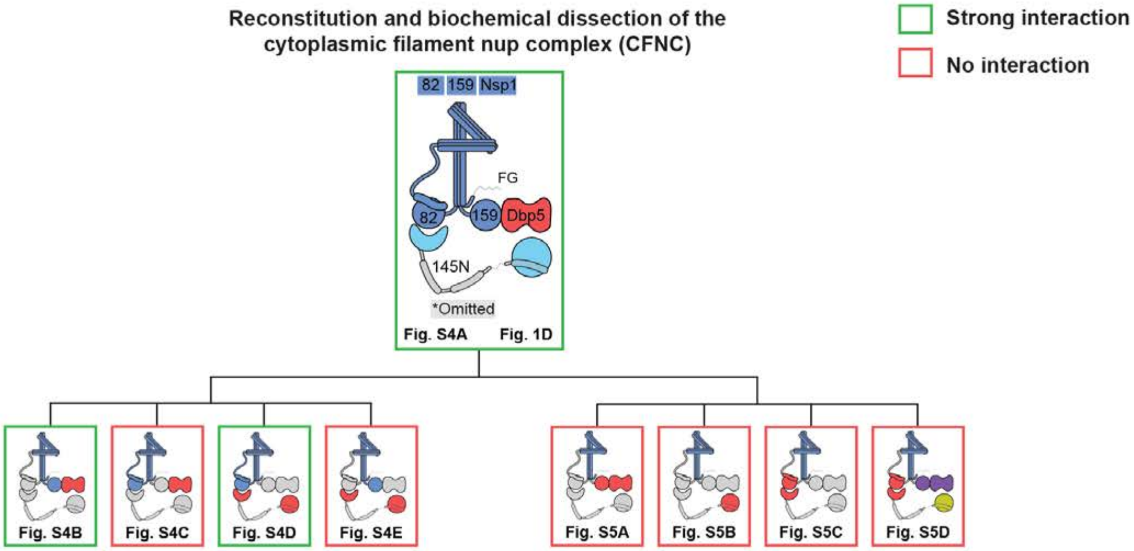

**Sup Fig. S3**

**Hoelz et al., 2021b**

**Fig. S3. Biochemical dissection of the CFNC architecture**. Summary of the SEC-MALS interaction analyses of the indicated figure panels. Each box in the flow chart represents a SEC-MALS interaction experiment, involving individually purified CFNC sub-complexes that were tested for their ability to form stochiometric complexes, colored by the experimental outcome; strong interaction (green), no interaction (red). The inset schematics are colored to indicate the CFNC sub-complexes of each SEC-MALS interaction experiment.

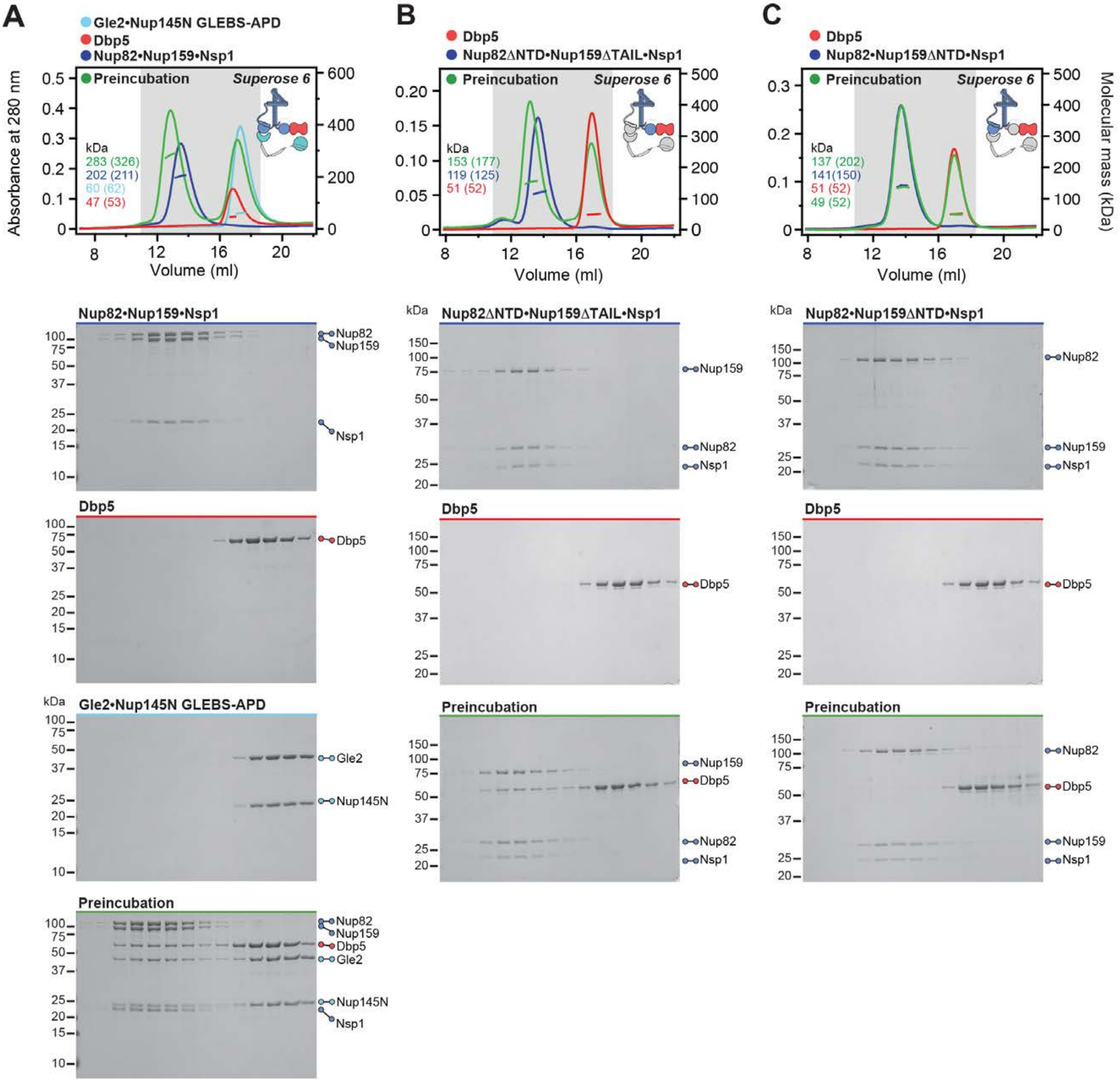

A
Gle2•Nup145N GLEBS-APD
Dbp5
Nup82•Nup159•Nsp1
Preincubation
Superose 6
Absorbance at 280 nm
kDa
283 (326)
202 (211)
60 (62)
47 (53)
Volume (ml)
B
Dbp5
Nup82ΔNTD•Nup159ΔTAIL•Nsp1
153 (177)
119 (125)
51 (52)
C
Nup82•Nup159ΔNTD•Nsp1
137 (202)
141(150)
51 (52)
49 (52)
Molecular mass (kDa)
Nup82
Nup159
Nsp1
Dbp5
Gle2
Nup145N

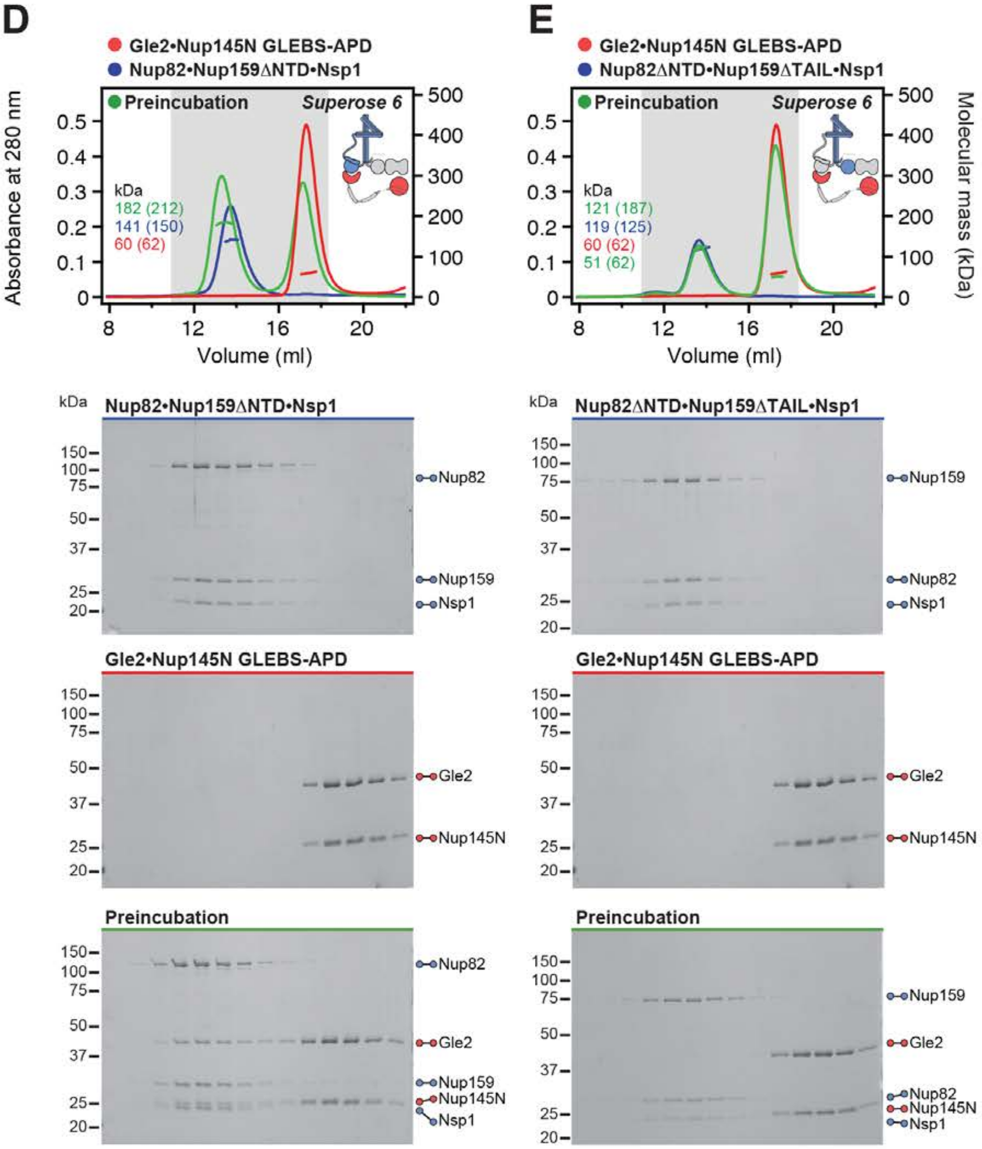
D
Gle2•Nup145N GLEBS-APD
Nup82•Nup159ΔNTD•Nsp1
Preincubation
Superose 6
Absorbance at 280 nm
kDa
182 (212)
141 (150)
60 (62)
Volume (ml)
Nup82•Nup159ΔNTD•Nsp1
Nup82
Nup159
Nsp1
Gle2•Nup145N GLEBS-APD
Gle2
Nup145N
Preincubation
Nup82
Gle2
Nup159
Nup145N
Nsp1
E
Gle2•Nup145N GLEBS-APD
Nup82ΔNTD•Nup159ΔTAIL•Nsp1
Preincubation
Superose 6
Molecular mass (kDa)
kDa
121 (187)
119 (125)
60 (62)
51 (62)
Volume (ml)
Nup82ΔNTD•Nup159ΔTAIL•Nsp1
Nup159
Nup82
Nsp1
Gle2•Nup145N GLEBS-APD
Gle2
Nup145N
Preincubation
Nup159
Gle2
Nup82
Nup145N
Nsp1

**Fig. S4. Reconstitution of the CFNC and elucidation of the Dbp5 and Gle2•Nup145N binding sites.** (**A**) SEC-MALS interaction analysis showing the biochemical reconstitution of the ~290 kDa hetero-hexameric CFNC from Nup82•Nup159•Nsp1, Gle2•Nup145N, and Dbp5 (corresponding to Fig. 1D). (**B, C**) SEC-MALS interaction analyses probing the interaction of Dbp5 with (B) Nup82$^{\Delta NTD}$•Nup159$^{\Delta TAIL}$•Nsp1 and (C) Nup82•Nup159$^{\Delta NTD}$•Nsp1 truncated CFNC variants. (**D, E**) SEC-MALS interaction analyses probing the interaction of Gle2•Nup145N$^{GLEBS\text{-}APD}$ with (D) Nup82•Nup159$^{\Delta NTD}$•Nsp1 and (E) Nup82$^{\Delta NTD}$•Nup159$^{\Delta TAIL}$•Nsp1 truncated CFNC variants. SEC-MALS profiles of nup complexes are shown individually (red, blue, and cyan) and after preincubation (green). SEC profiles were obtained using a Superose 6 10/300 GL column. Measured molecular masses are indicated, with the respective theoretical masses shown in parentheses. Inset schematics display tested protein samples colored to match the corresponding chromatogram trace. Gray boxes indicate fractions resolved on SDS-PAGE gels and visualized by Coomassie brilliant blue staining.

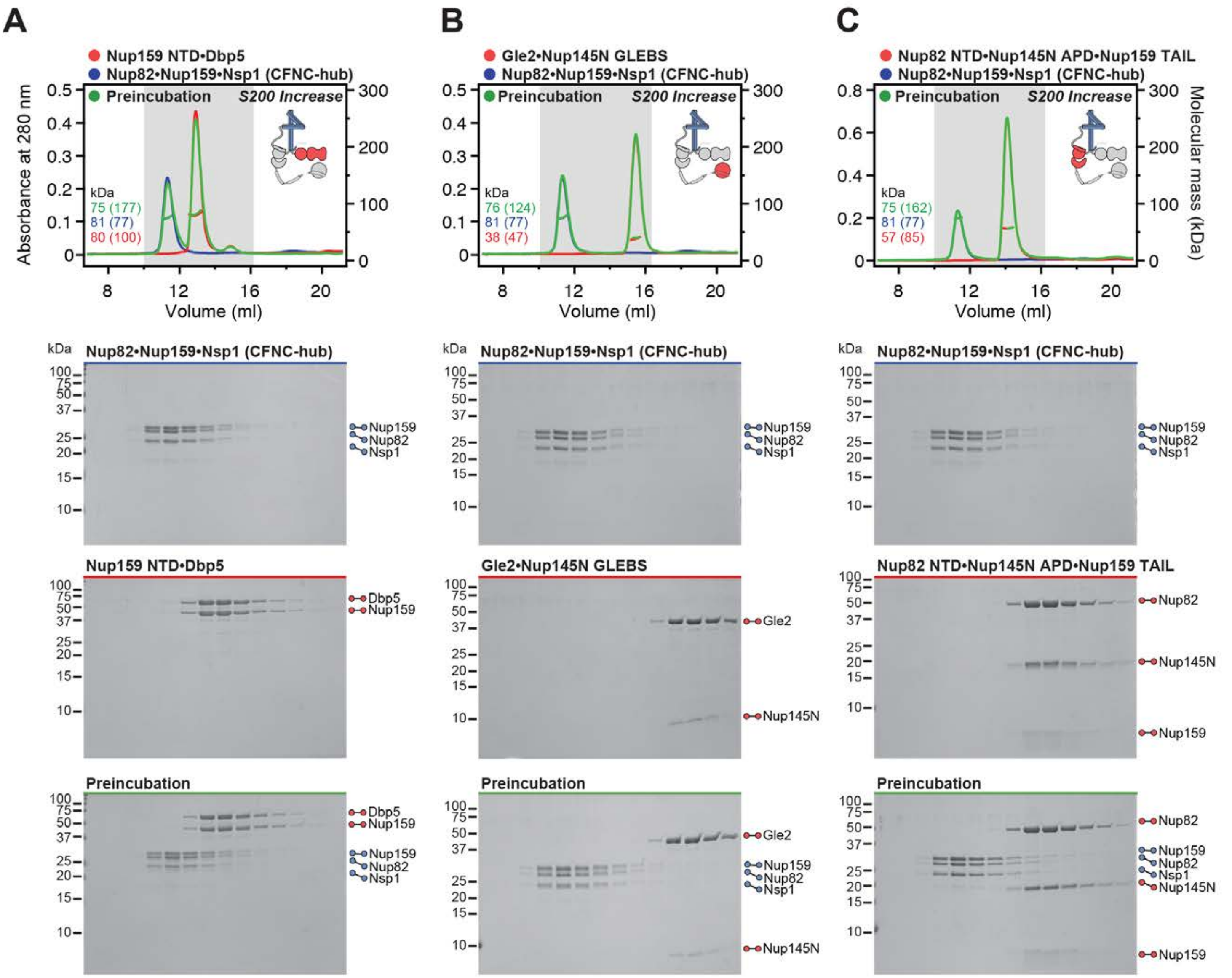

A
Nup159 NTD•Dbp5
Nup82•Nup159•Nsp1 (CFNC-hub)
Preincubation
S200 Increase
Absorbance at 280 nm
kDa
75 (177)
81 (77)
80 (100)
Volume (ml)
B
Gle2•Nup145N GLEBS
Nup82•Nup159•Nsp1 (CFNC-hub)
Preincubation
S200 Increase
kDa
76 (124)
81 (77)
38 (47)
Volume (ml)
C
Nup82 NTD•Nup145N APD•Nup159 TAIL
Nup82•Nup159•Nsp1 (CFNC-hub)
Preincubation
S200 Increase
Molecular mass (kDa)
kDa
75 (162)
81 (77)
57 (85)
Volume (ml)
Nup82•Nup159•Nsp1 (CFNC-hub)
Nup159
Nup82
Nsp1
Nup159 NTD•Dbp5
Dbp5
Nup159
Gle2•Nup145N GLEBS
Gle2
Nup145N
Nup82 NTD•Nup145N APD•Nup159 TAIL
Nup82
Nup145N
Nup159
Preincubation

**D**

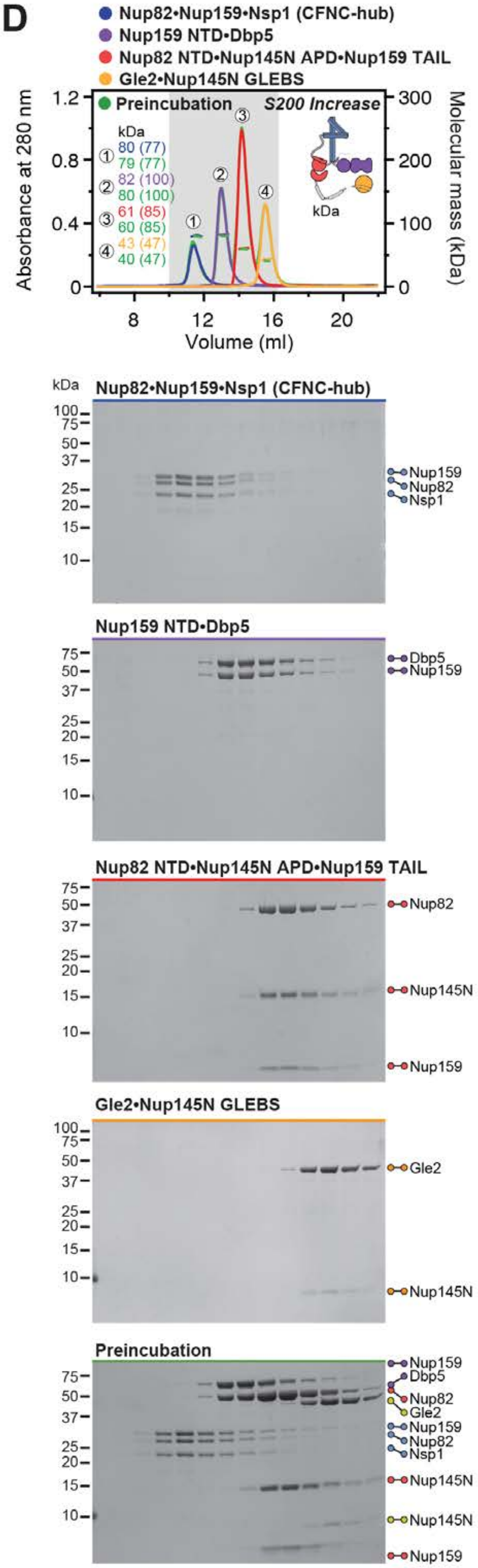

Nup82•Nup159•Nsp1 (CFNC-hub)
Nup159 NTD•Dbp5
Nup82 NTD•Nup145N APD•Nup159 TAIL
Gle2•Nup145N GLEBS
Preincubation
S200 Increase
kDa
① 80 (77) 79 (77)
② 82 (100) 80 (100)
③ 61 (85) 60 (85)
④ 43 (47) 40 (47)
Absorbance at 280 nm
Molecular mass (kDa)
Volume (ml)
Nup82•Nup159•Nsp1 (CFNC-hub)
Nup159
Nup82
Nsp1
Nup159 NTD•Dbp5
Dbp5
Nup159
Nup82 NTD•Nup145N APD•Nup159 TAIL
Nup82
Nup145N
Nup159
Gle2•Nup145N GLEBS
Gle2
Nup145N
Preincubation
Nup159
Dbp5
Nup82
Gle2
Nup159
Nup82
Nsp1
Nup145N
Nup145N
Nup159

**Fig. S5. Biochemical dissection of the CFNC into four distinct non-interacting sub-complexes.** SEC-MALS interaction analysis of the Nup82•Nup159•Nsp1 CCS1-3 (CFNC-hub) with (**A**) Nup159$^{NTD}$•Dbp5, (**B**) Gle2•Nup145N$^{GLEBS}$, (**C**) Nup82$^{NTD}$•Nup145N$^{APD}$•Nup159$^{TAIL}$, and (**D**) a mixture of all four subcomplexes. SEC-MALS profiles of nup complexes are shown individually (red, blue, purple and orange) and after preincubation (green). SEC profiles were obtained using a Superdex 200 Increase 10/300 GL column. Measured molecular masses are indicated, with the respective theoretical masses shown in parentheses. Inset schematics display tested protein samples colored to match the corresponding chromatogram trace. Gray boxes indicate fractions resolved on SDS-PAGE gels and visualized by Coomassie brilliant blue staining.

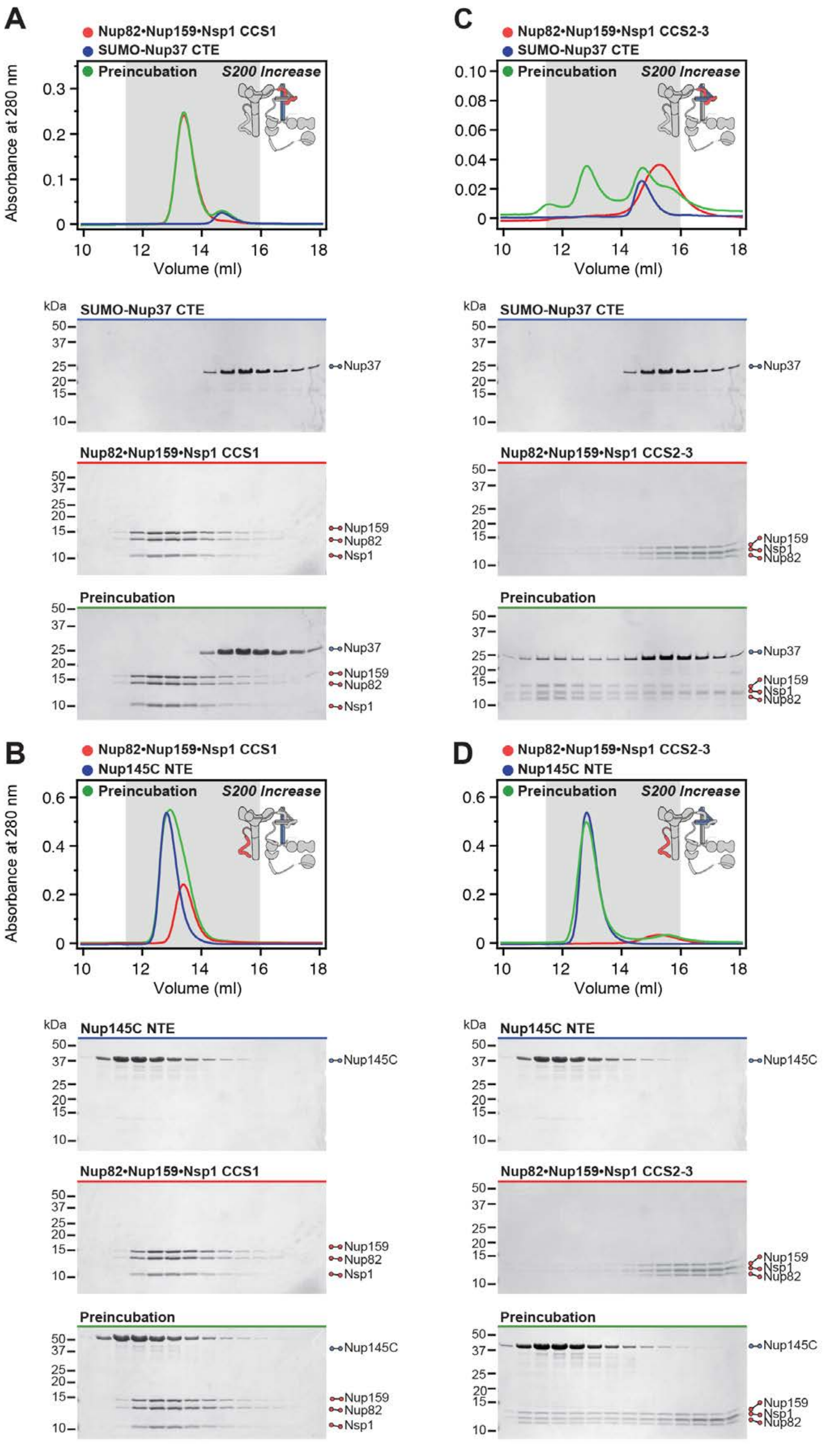

Sup Fig. S6

Hoelz et al., 2021b

**Fig. S6. Biochemical analysis of the Nup37/Nup145C-CFNC-hub interaction.** (**A**, **B**) SEC interaction analyses of Nup82•Nup159•Nsp1 CCS1 with (A) SUMO-Nup37$^{CTE}$ and (B) Nup145C$^{NTE}$, showing that neither form a stable complex. (**C**, **D**) SEC interaction analyses of Nup82•Nup159•Nsp1 CCS2-3 with (C) SUMO-Nup37$^{CTE}$ and (D) Nup145C$^{NTE}$, demonstrating that both form stable interactions. SEC profiles of nup complexes are shown individually (red, blue) and after preincubation (green). SEC profiles were obtained using a Superdex 200 Increase 10/300 GL column. Inset schematics display tested protein samples colored to match the corresponding chromatogram trace. Gray boxes indicate fractions resolved on SDS-PAGE gels and visualized by Coomassie brilliant blue staining.

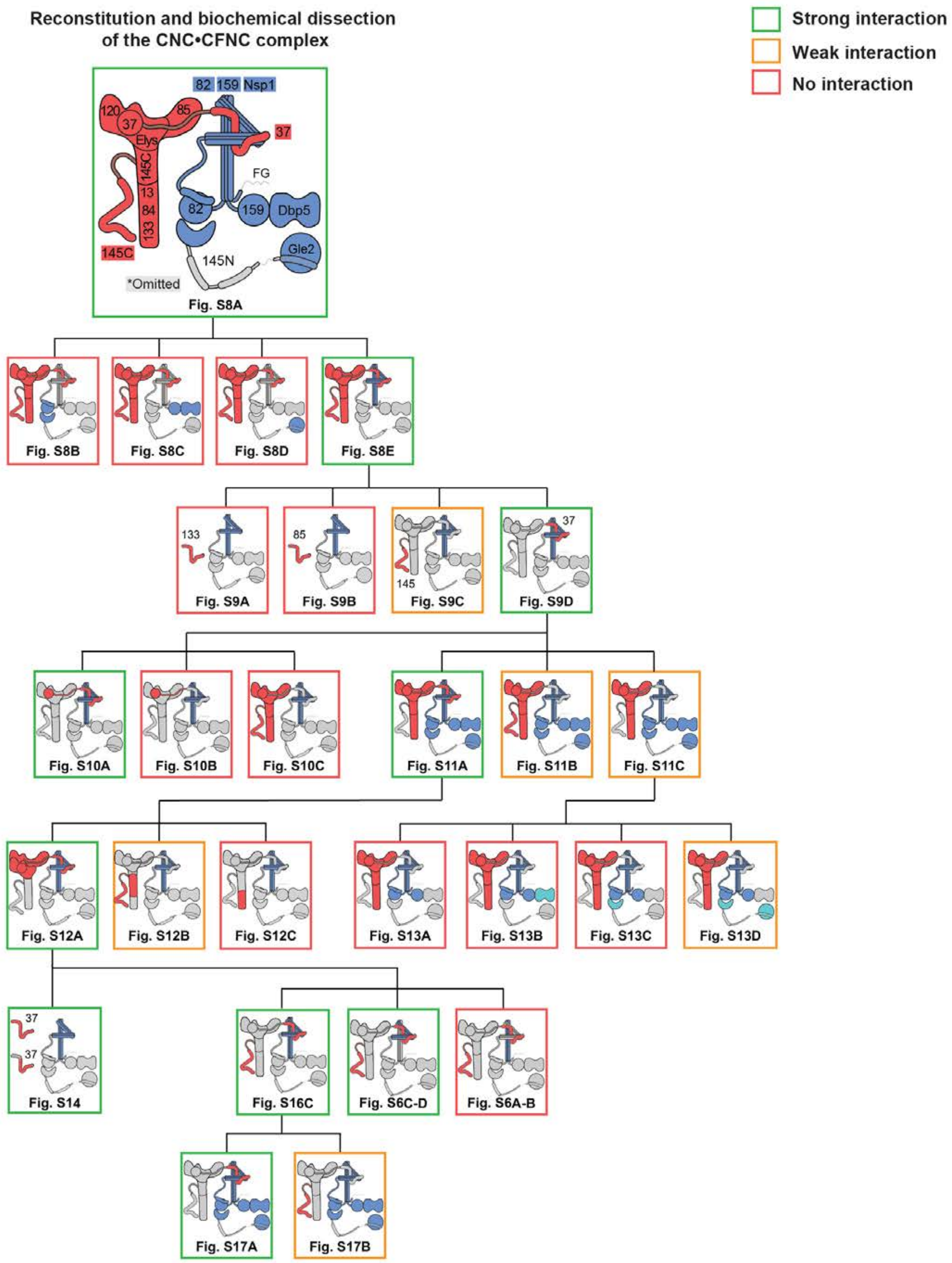

Reconstitution and biochemical dissection of the CNC•CFNC complex
Strong interaction
Weak interaction
No interaction
82 159 Nsp1
120
37
85
Elys
145C
13
84
133
37
FG
82
159
Dbp5
145C
145N
Gle2
*Omitted
Fig. S8A
Fig. S8B
Fig. S8C
Fig. S8D
Fig. S8E
133
Fig. S9A
85
Fig. S9B
145
Fig. S9C
37
Fig. S9D
Fig. S10A
Fig. S10B
Fig. S10C
Fig. S11A
Fig. S11B
Fig. S11C
Fig. S12A
Fig. S12B
Fig. S12C
Fig. S13A
Fig. S13B
Fig. S13C
Fig. S13D
37
37
Fig. S14
Fig. S16C
Fig. S6C-D
Fig. S6A-B
Fig. S17A
Fig. S17B

**Fig. S7. Reconstitution and dissection strategy for the 14-protein CNC•CFNC complex.** Flow chart detailing reconstitution and biochemical dissection of the 14-protein *C. thermophilum* CNC•CFNC complex. Each box in the flow chart represents a SEC-MALS interaction experiment, involving individually purified CNC and CNFC sub-complexes that were tested for their ability to form stochiometric complexes, colored by the experimental outcome; strong interaction (green), weak interaction (orange), no interaction (red). The inset schematics are colored to indicate the CNC and CFNC sub-complexes of each SEC-MALS interaction experiment, gray regions stipulate proteins omitted from each analysis.

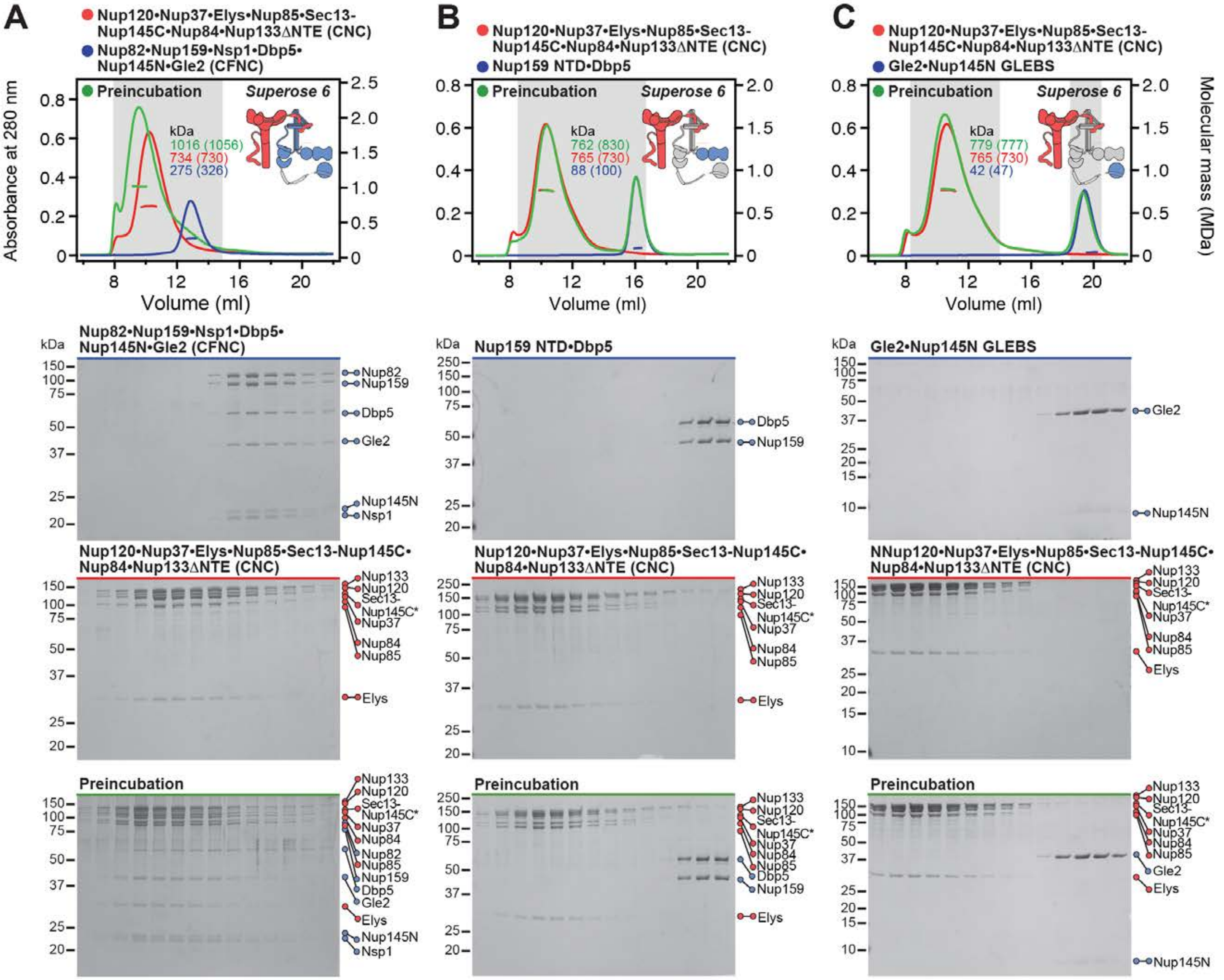
A
Nup120•Nup37•Elys•Nup85•Sec13-Nup145C•Nup84•Nup133ΔNTE (CNC)
Nup82•Nup159•Nsp1•Dbp5•Nup145N•Gle2 (CFNC)
Preincubation
Superose 6
kDa
1016 (1056)
734 (730)
275 (326)
Absorbance at 280 nm
Volume (ml)
B
Nup120•Nup37•Elys•Nup85•Sec13-Nup145C•Nup84•Nup133ΔNTE (CNC)
Nup159 NTD•Dbp5
762 (830)
765 (730)
88 (100)
C
Nup120•Nup37•Elys•Nup85•Sec13-Nup145C•Nup84•Nup133ΔNTE (CNC)
Gle2•Nup145N GLEBS
779 (777)
765 (730)
42 (47)
Molecular mass (MDa)
Nup82•Nup159•Nsp1•Dbp5•Nup145N•Gle2 (CFNC)
Nup82
Nup159
Dbp5
Gle2
Nup145N
Nsp1
Nup159 NTD•Dbp5
Gle2•Nup145N GLEBS
Nup120•Nup37•Elys•Nup85•Sec13-Nup145C•Nup84•Nup133ΔNTE (CNC)
NNup120•Nup37•Elys•Nup85•Sec13-Nup145C•Nup84•Nup133ΔNTE (CNC)
Nup133
Nup120
Sec13-
Nup145C*
Nup37
Nup84
Nup85
Elys
Preincubation

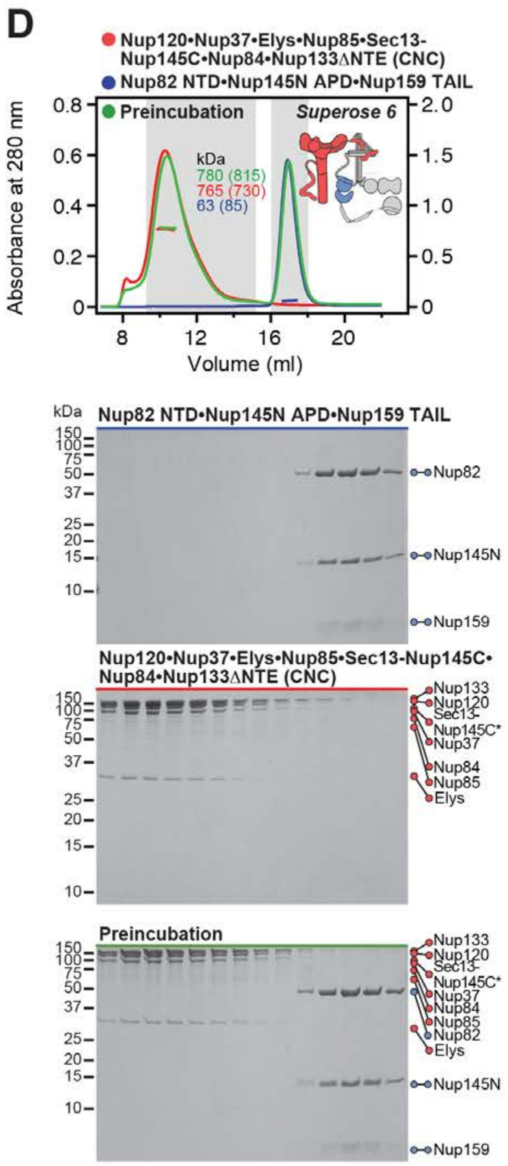
D
Nup120•Nup37•Elys•Nup85•Sec13-Nup145C•Nup84•Nup133ΔNTE (CNC)
Nup82 NTD•Nup145N APD•Nup159 TAIL
Preincubation
Superose 6
kDa
780 (815)
765 (730)
63 (85)
Absorbance at 280 nm
Volume (ml)
Nup82 NTD•Nup145N APD•Nup159 TAIL
Nup82
Nup145N
Nup159
Nup120•Nup37•Elys•Nup85•Sec13-Nup145C•Nup84•Nup133ΔNTE (CNC)
Nup133
Nup120
Sec13-
Nup145C*
Nup37
Nup84
Nup85
Elys
Preincubation
Nup82

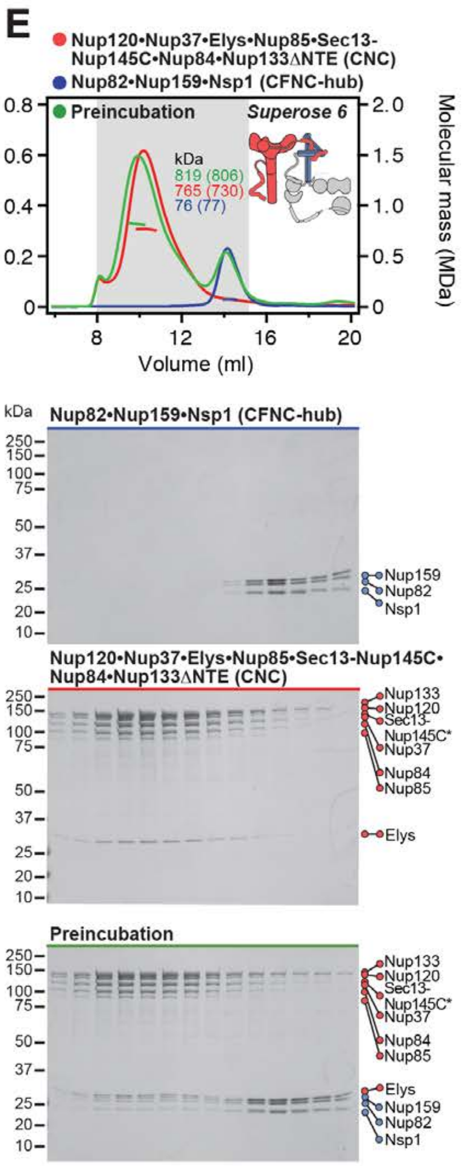
E
Nup120•Nup37•Elys•Nup85•Sec13-Nup145C•Nup84•Nup133ΔNTE (CNC)
Nup82•Nup159•Nsp1 (CFNC-hub)
Preincubation
Superose 6
kDa
819 (806)
765 (730)
76 (77)
Molecular mass (MDa)
Volume (ml)
Nup82•Nup159•Nsp1 (CFNC-hub)
Nup159
Nup82
Nsp1
Nup120•Nup37•Elys•Nup85•Sec13-Nup145C•Nup84•Nup133ΔNTE (CNC)
Nup133
Nup120
Sec13-
Nup145C*
Nup37
Nup84
Nup85
Elys
Preincubation

**Fig. S8. Reconstitution and dissection of the 14-protein CNC-CFNC interaction.** (**A**) SEC-MALS interaction analysis of the Nup120•Nup37•Elys•Nup85•Sec13-Nup145C•Nup84•Nup133$^{\Delta NTE}$ (CNC) with the intact CFNC, demonstrating the formation of a ~1,016 kDa monodisperse, stochiometric 14-protein CNC•CFNC complex. (**B**-**E**) SEC-MALS interaction analysis of the intact CNC with (**B**) Nup159$^{NTD}$•Dbp5, (**C**) Gle2•Nup145N$^{GLEBS}$, (**D**) Nup82$^{NTD}$•Nup145N$^{APD}$•Nup159$^{TAIL}$, and (**E**) CFNC-hub, showing that of the four distinct CFNC subcomplexes only the CFNC-hub forms a stable interaction with the CNC. SEC-MALS profiles of nup complexes are shown individually (red, blue) and after preincubation (green). SEC profiles were obtained using a Superose 6 10/300 GL column. Measured molecular masses are indicated, with the respective theoretical masses shown in parentheses. Inset schematics display tested protein samples colored to match the corresponding chromatogram trace. Gray boxes indicate fractions resolved on SDS-PAGE gels and visualized by Coomassie brilliant blue staining.

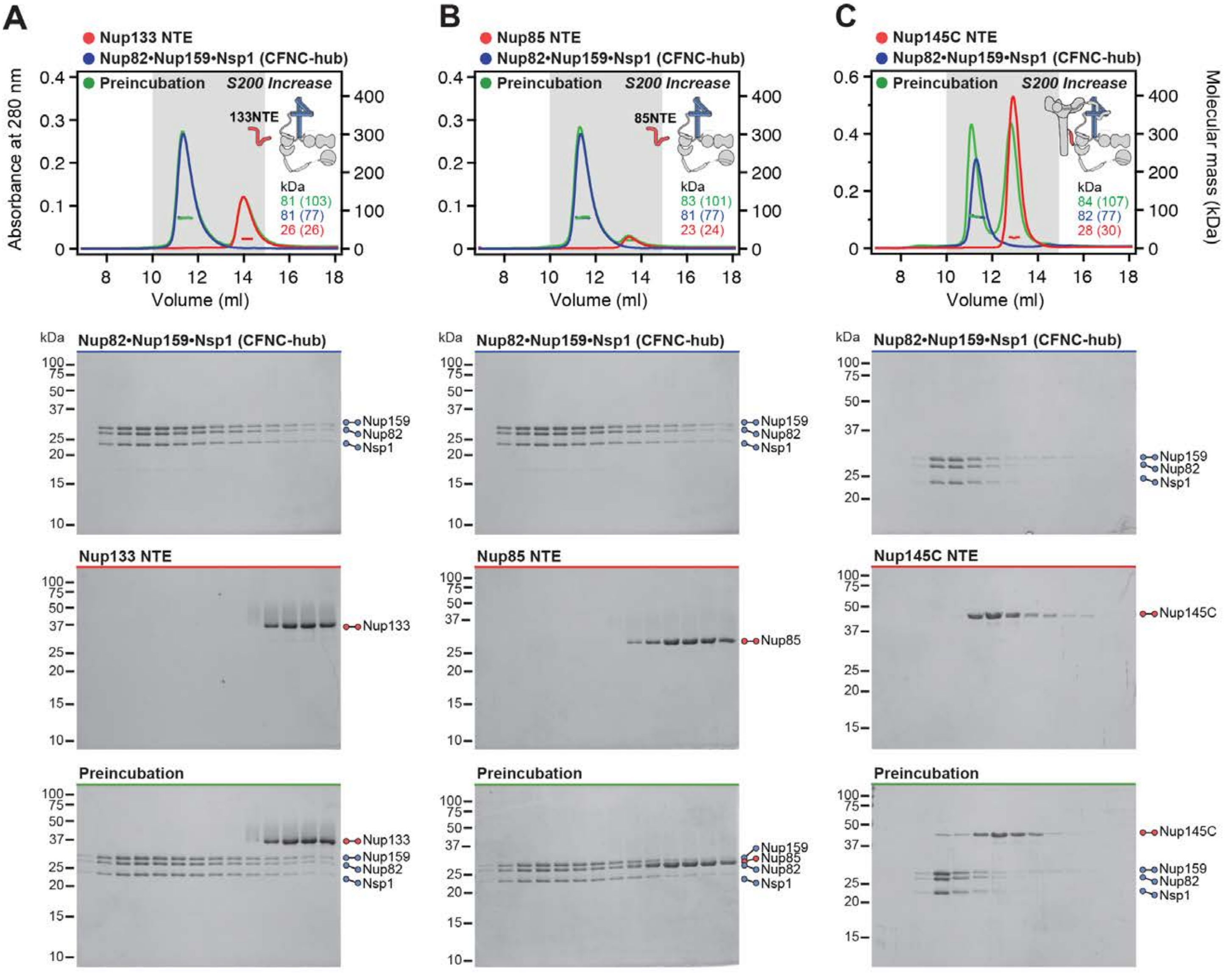
A
Nup133 NTE
Nup82•Nup159•Nsp1 (CFNC-hub)
Preincubation
S200 Increase
133NTE
kDa
81 (103)
81 (77)
26 (26)
Absorbance at 280 nm
Volume (ml)
B
Nup85 NTE
Nup82•Nup159•Nsp1 (CFNC-hub)
Preincubation
S200 Increase
85NTE
kDa
83 (101)
81 (77)
23 (24)
C
Nup145C NTE
Nup82•Nup159•Nsp1 (CFNC-hub)
Preincubation
S200 Increase
kDa
84 (107)
82 (77)
28 (30)
Molecular mass (kDa)
Nup82•Nup159•Nsp1 (CFNC-hub)
Nup159
Nup82
Nsp1
Nup133 NTE
Nup133
Nup85 NTE
Nup85
Nup145C NTE
Nup145C
Preincubation

D

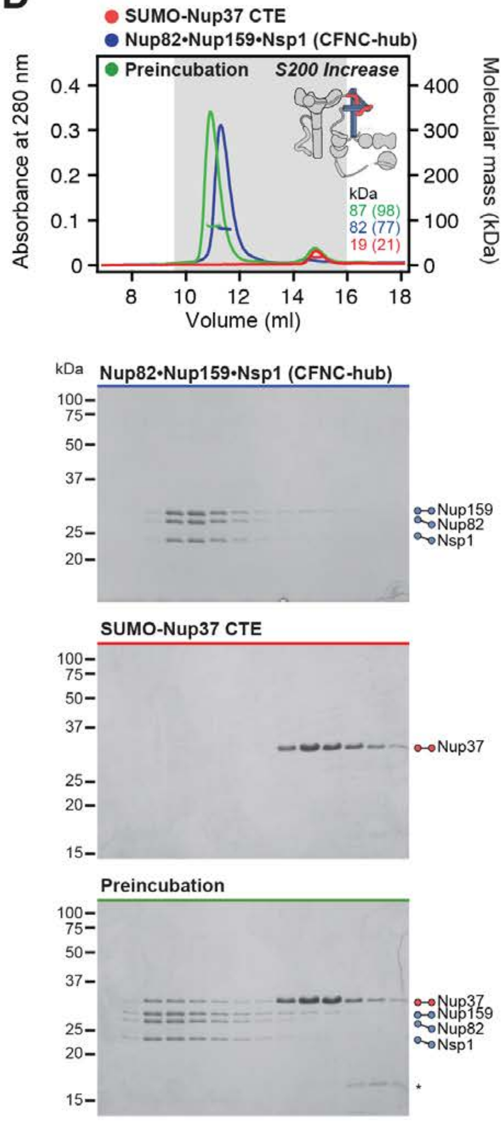

SUMO-Nup37 CTE
Nup82•Nup159•Nsp1 (CFNC-hub)
Preincubation
S200 Increase
Absorbance at 280 nm
Molecular mass (kDa)
Volume (ml)
kDa
87 (98)
82 (77)
19 (21)
Nup82•Nup159•Nsp1 (CFNC-hub)
Nup159
Nup82
Nsp1
SUMO-Nup37 CTE
Nup37
Preincubation
Nup37
Nup159
Nup82
Nsp1

**Fig. S9. Anchoring of the CNFC-hub to the CNC is mediated by Nup145C$^{NTE}$ and Nup37$^{CTE}$.** SEC-MALS interaction analyses of CFNC-hub with (**A**) Nup133$^{NTE}$, (**B**) Nup85$^{NTE}$, (**C**) Nup145C$^{NTE}$, and (**D**) SUMO-Nup37$^{CTE}$, showing that Nup145C and Nup37 bind to the CFNC-hub. SEC-MALS profiles of nup complexes are shown individually (red, blue) and after preincubation (green). SEC profiles were obtained using a Superdex 200 Increase 10/300 GL column. Measured molecular masses are indicated, with the respective theoretical masses shown in parentheses. Inset schematics display tested protein samples colored to match the corresponding chromatogram trace. Gray boxes indicate fractions resolved on SDS-PAGE gels and visualized by Coomassie brilliant blue staining.

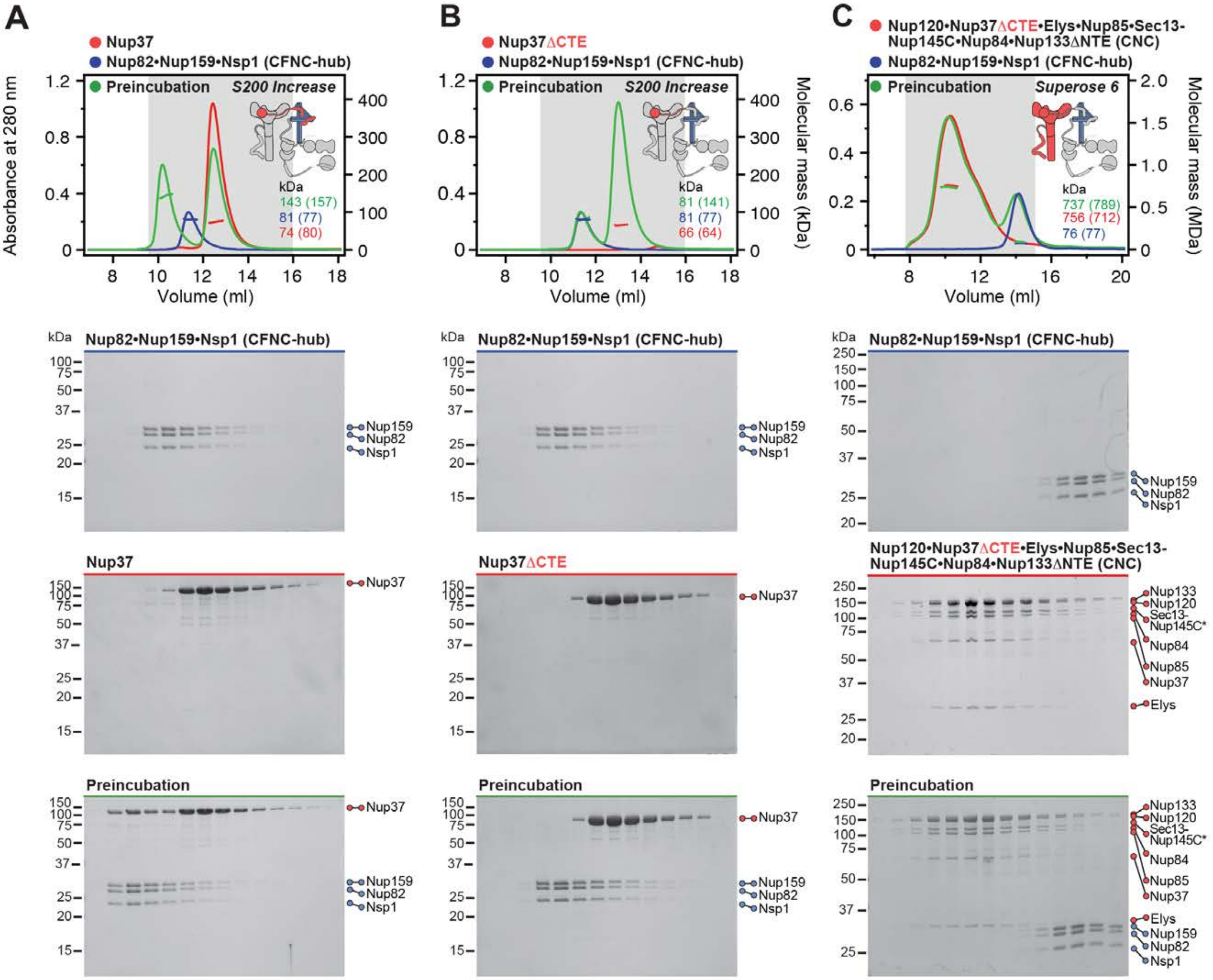

A
Nup37
Nup82•Nup159•Nsp1 (CFNC-hub)
Preincubation
S200 Increase
Absorbance at 280 nm
Volume (ml)
kDa
143 (157)
81 (77)
74 (80)
B
Nup37ΔCTE
Nup82•Nup159•Nsp1 (CFNC-hub)
Preincubation
S200 Increase
Molecular mass (kDa)
81 (141)
81 (77)
66 (64)
C
Nup120•Nup37ΔCTE•Elys•Nup85•Sec13-Nup145C•Nup84•Nup133ΔNTE (CNC)
Nup82•Nup159•Nsp1 (CFNC-hub)
Preincubation
Superose 6
Molecular mass (MDa)
737 (789)
756 (712)
76 (77)
Nup82•Nup159•Nsp1 (CFNC-hub)
Nup159
Nup82
Nsp1
Nup37
Nup37ΔCTE
Nup120•Nup37ΔCTE•Elys•Nup85•Sec13-Nup145C•Nup84•Nup133ΔNTE (CNC)
Nup133
Nup120
Sec13-Nup145C*
Nup84
Nup85
Elys
Preincubation

**Fig. S10. Nup37$^{CTE}$ is necessary and sufficient for CFNC-hub binding.** SEC-MALS interaction analysis of CFNC-hub with (**A**) Nup37, (**B**) Nup37$^{\Delta CTE}$, (**C**) Nup120•Nup37$^{\Delta CTE}$•Elys•Nup85•Sec13-Nup145C•Nup84•Nup133$^{\Delta NTE}$ (CNC$^{\Delta 37CTE}$). SEC-MALS profiles of nup complexes are shown individually (red, blue) and after preincubation (green). SEC profiles were obtained using (A, B) Superose 6 10/300 GL and (C) Superdex 200 10/300 GL columns. Measured molecular masses are indicated, with the respective theoretical masses shown in parentheses. Inset schematics display tested protein samples colored to match the corresponding chromatogram trace. Gray boxes indicate fractions resolved on SDS-PAGE gels and visualized by Coomassie brilliant blue staining.

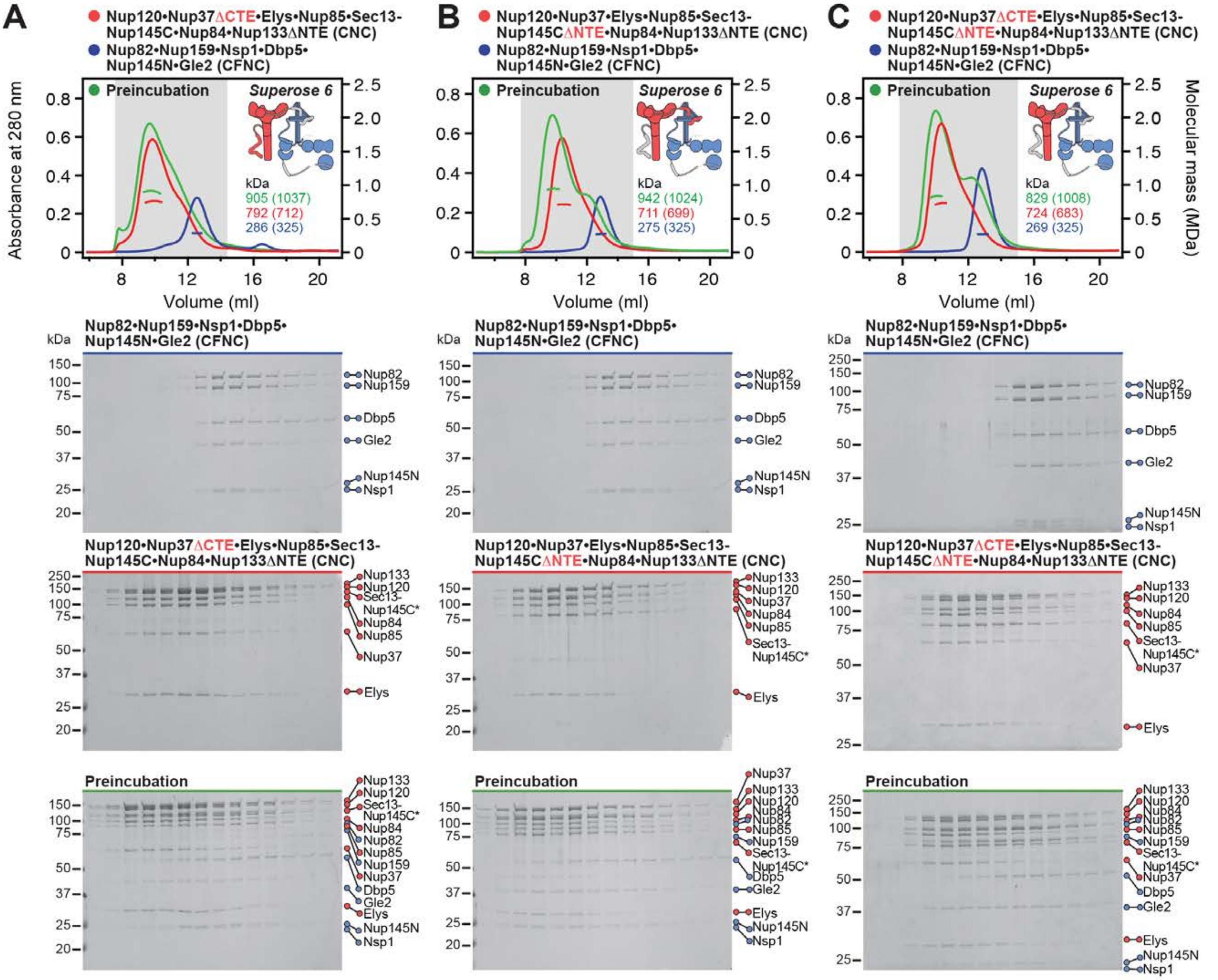
A
Nup120•Nup37ΔCTE•Elys•Nup85•Sec13-Nup145C•Nup84•Nup133ΔNTE (CNC)
Nup82•Nup159•Nsp1•Dbp5•Nup145N•Gle2 (CFNC)
Preincubation
Superose 6
kDa
905 (1037)
792 (712)
286 (325)
Absorbance at 280 nm
Volume (ml)
B
Nup120•Nup37•Elys•Nup85•Sec13-Nup145CΔNTE•Nup84•Nup133ΔNTE (CNC)
Nup82•Nup159•Nsp1•Dbp5•Nup145N•Gle2 (CFNC)
Preincubation
Superose 6
kDa
942 (1024)
711 (699)
275 (325)
Volume (ml)
C
Nup120•Nup37ΔCTE•Elys•Nup85•Sec13-Nup145CΔNTE•Nup84•Nup133ΔNTE (CNC)
Nup82•Nup159•Nsp1•Dbp5•Nup145N•Gle2 (CFNC)
Preincubation
Superose 6
kDa
829 (1008)
724 (683)
269 (325)
Molecular mass (MDa)
Volume (ml)
Nup82•Nup159•Nsp1•Dbp5•Nup145N•Gle2 (CFNC)
Nup82
Nup159
Dbp5
Gle2
Nup145N
Nsp1
Nup120•Nup37ΔCTE•Elys•Nup85•Sec13-Nup145C•Nup84•Nup133ΔNTE (CNC)
Nup120•Nup37•Elys•Nup85•Sec13-Nup145CΔNTE•Nup84•Nup133ΔNTE (CNC)
Nup120•Nup37ΔCTE•Elys•Nup85•Sec13-Nup145CΔNTE•Nup84•Nup133ΔNTE (CNC)
Nup133
Nup120
Sec13-Nup145C*
Nup84
Nup85
Nup37
Elys
Preincubation

**Fig. S11. Each intact CNC contains two distinct CFNC binding sites.** SEC-MALS interaction analysis of intact CFNC with (**A**) CNC$^{\Delta 37CTE}$, (**B**) CNC$^{\Delta 145CNTE}$, and (**C**) CNC$^{\Delta 37CTE \Delta 145CNTE}$, showing that the simultaneous removal of both Nup37$^{CTE}$ and Nup145C$^{NTE}$ disrupts the CFNC interaction, whereas their individual removal only weakens the CNC-CFNC interaction. SEC-MALS profiles of nup complexes are shown individually (red, blue) and after preincubation (green). SEC profiles were obtained using a Superose 6 10/300 GL column. Measured molecular masses are indicated, with the respective theoretical masses shown in parentheses. Inset schematics display tested protein samples colored to match the corresponding chromatogram trace. Gray boxes indicate fractions resolved on SDS-PAGE gels and visualized by Coomassie brilliant blue staining.

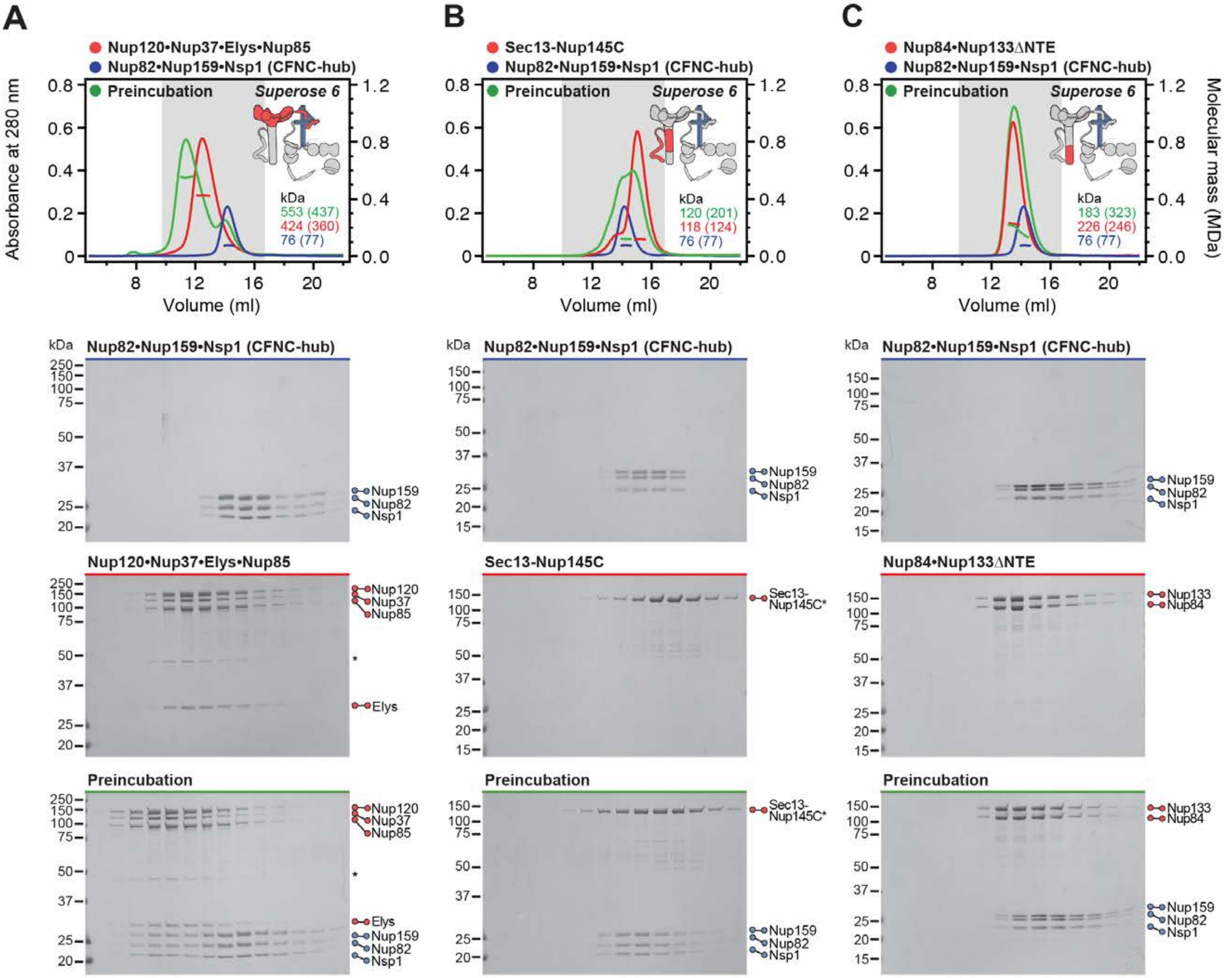
A
Nup120•Nup37•Elys•Nup85
Nup82•Nup159•Nsp1 (CFNC-hub)
Preincubation
Superose 6
Absorbance at 280 nm
Volume (ml)
kDa
553 (437)
424 (360)
76 (77)
B
Sec13-Nup145C
Nup82•Nup159•Nsp1 (CFNC-hub)
Preincubation
Superose 6
120 (201)
118 (124)
76 (77)
C
Nup84•Nup133ΔNTE
Nup82•Nup159•Nsp1 (CFNC-hub)
Preincubation
Superose 6
183 (323)
226 (246)
76 (77)
Molecular mass (MDa)
Nup82•Nup159•Nsp1 (CFNC-hub)
Nup159
Nup82
Nsp1
Nup120•Nup37•Elys•Nup85
Nup120
Nup37
Nup85
Elys
Sec13-Nup145C
Sec13-Nup145C*
Nup84•Nup133ΔNTE
Nup133
Nup84
Preincubation

**Fig. S12. Two CNC sub-complexes bind the CFNC-hub.** SEC-MALS interaction analysis of CFNC-hub with (**A**) Nup120•Nup37•Elys•Nup85, (**B**) Sec13-Nup145C, and (**C**) Nup84•Nup133$^{\Delta NTE}$. SEC-MALS profiles of nup complexes are shown individually (red, blue) and after preincubation (green). SEC profiles were obtained using a Superose 6 10/300 GL column. Measured molecular masses are indicated, with the respective theoretical masses shown in parentheses. Inset schematics display tested protein samples colored to match the corresponding chromatogram trace. Gray boxes indicate fractions resolved on SDS-PAGE gels and visualized by Coomassie brilliant blue staining.

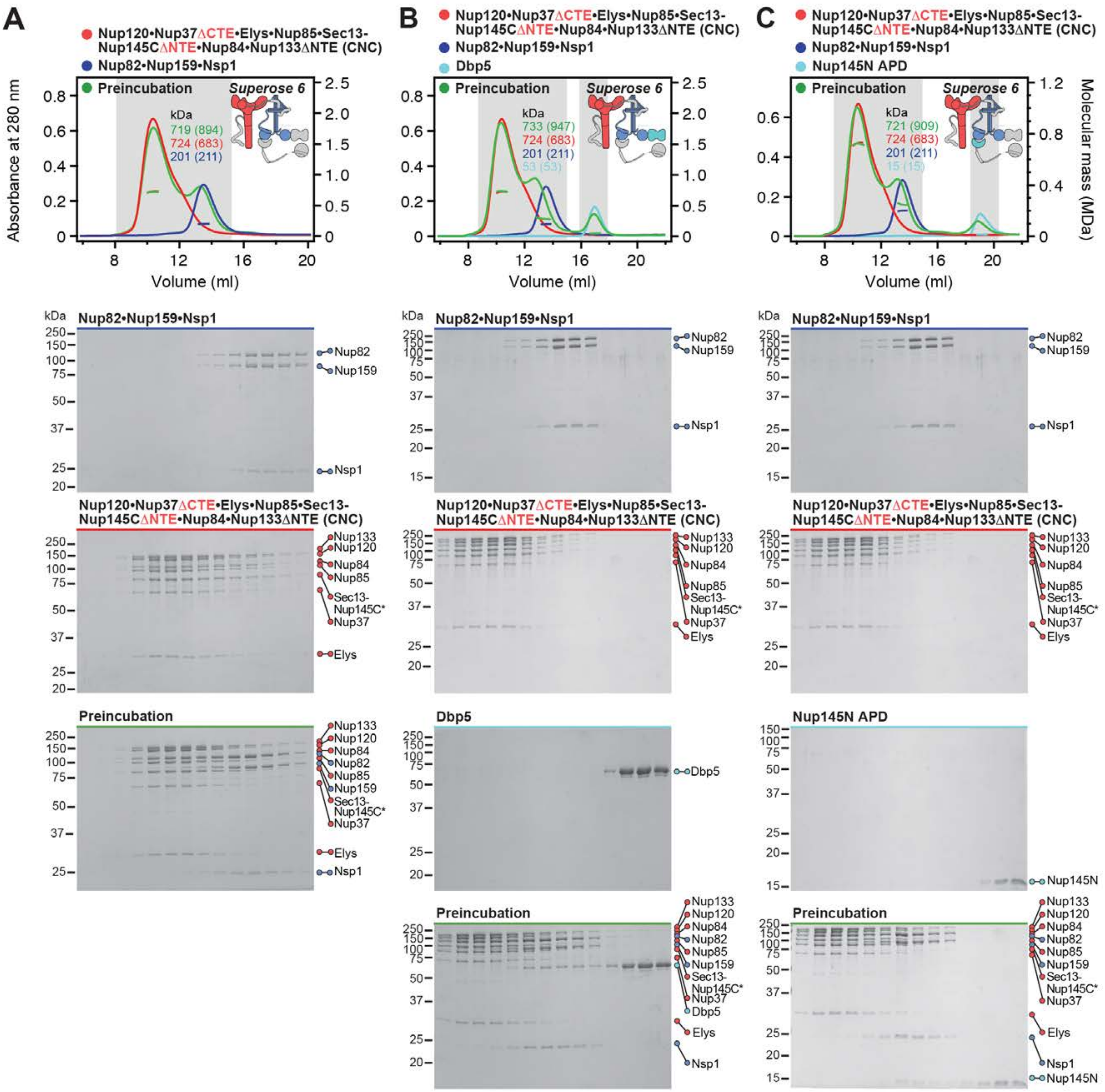
A
B
C
Nup120•Nup37ΔCTE•Elys•Nup85•Sec13-Nup145CΔNTE•Nup84•Nup133ΔNTE (CNC)
Nup82•Nup159•Nsp1
Dbp5
Nup145N APD
Preincubation
Superose 6
Absorbance at 280 nm
Molecular mass (MDa)
Volume (ml)
kDa
719 (894)
724 (683)
201 (211)
733 (947)
53 (53)
721 (909)
15 (15)
Nup133
Nup120
Nup84
Nup82
Nup85
Nup159
Sec13-Nup145C*
Nup37
Elys
Nsp1
Dbp5
Nup145N

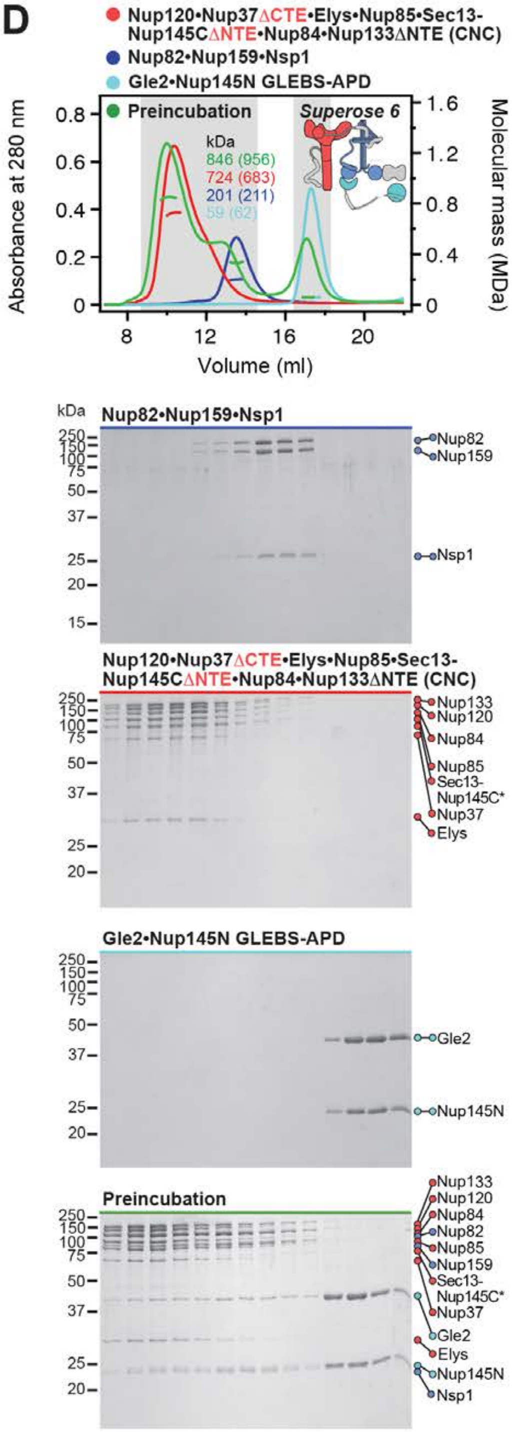
D
Nup120•Nup37ΔCTE•Elys•Nup85•Sec13-Nup145CΔNTE•Nup84•Nup133ΔNTE (CNC)
Nup82•Nup159•Nsp1
Gle2•Nup145N GLEBS-APD
Preincubation
Superose 6
kDa
846 (956)
724 (683)
201 (211)
59 (62)
Absorbance at 280 nm
Molecular mass (MDa)
Volume (ml)
Nup82•Nup159•Nsp1
Nup82
Nup159
Nsp1
Nup120•Nup37ΔCTE•Elys•Nup85•Sec13-Nup145CΔNTE•Nup84•Nup133ΔNTE (CNC)
Nup133
Nup120
Nup84
Nup85
Sec13-Nup145C*
Nup37
Elys
Gle2•Nup145N GLEBS-APD
Gle2
Nup145N
Preincubation
Nup133
Nup120
Nup84
Nup82
Nup85
Nup159
Sec13-Nup145C*
Nup37
Gle2
Elys
Nup145N
Nsp1

**Fig. S13. Gle2•Nup145N contributes weakly to the CNC-CFNC interaction.** SEC-MALS interaction analysis of $CNC^{\Delta 37CTE \Delta 145CNTE}$ with (**A**) Nup82•Nup159•Nsp1, (**B**) Nup82•Nup159•Nsp1 and Dbp5, (**C**) Nup82•Nup159•Nsp1 and $Nup145N^{APD}$, and (**D**) Nup82•Nup159•Nsp1 and $Gle2•Nup145N^{GLEBS\text{-}APD}$, showing that in the absence of both $Nup37^{CTE}$ and $Nup145C^{NTE}$ CFNC binding sites, residual CNC binding is $Gle2•Nup145N^{GLEBS}$-dependent. SEC-MALS profiles of nup complexes are shown individually (red, blue, cyan) and after preincubation (green). SEC profiles were obtained using a Superose 6 10/300 GL column. Measured molecular masses are indicated, with the respective theoretical masses shown in parentheses. Inset schematics display tested protein samples colored to match the corresponding chromatogram trace. Gray boxes indicate fractions resolved on SDS-PAGE gels and visualized by Coomassie brilliant blue staining.

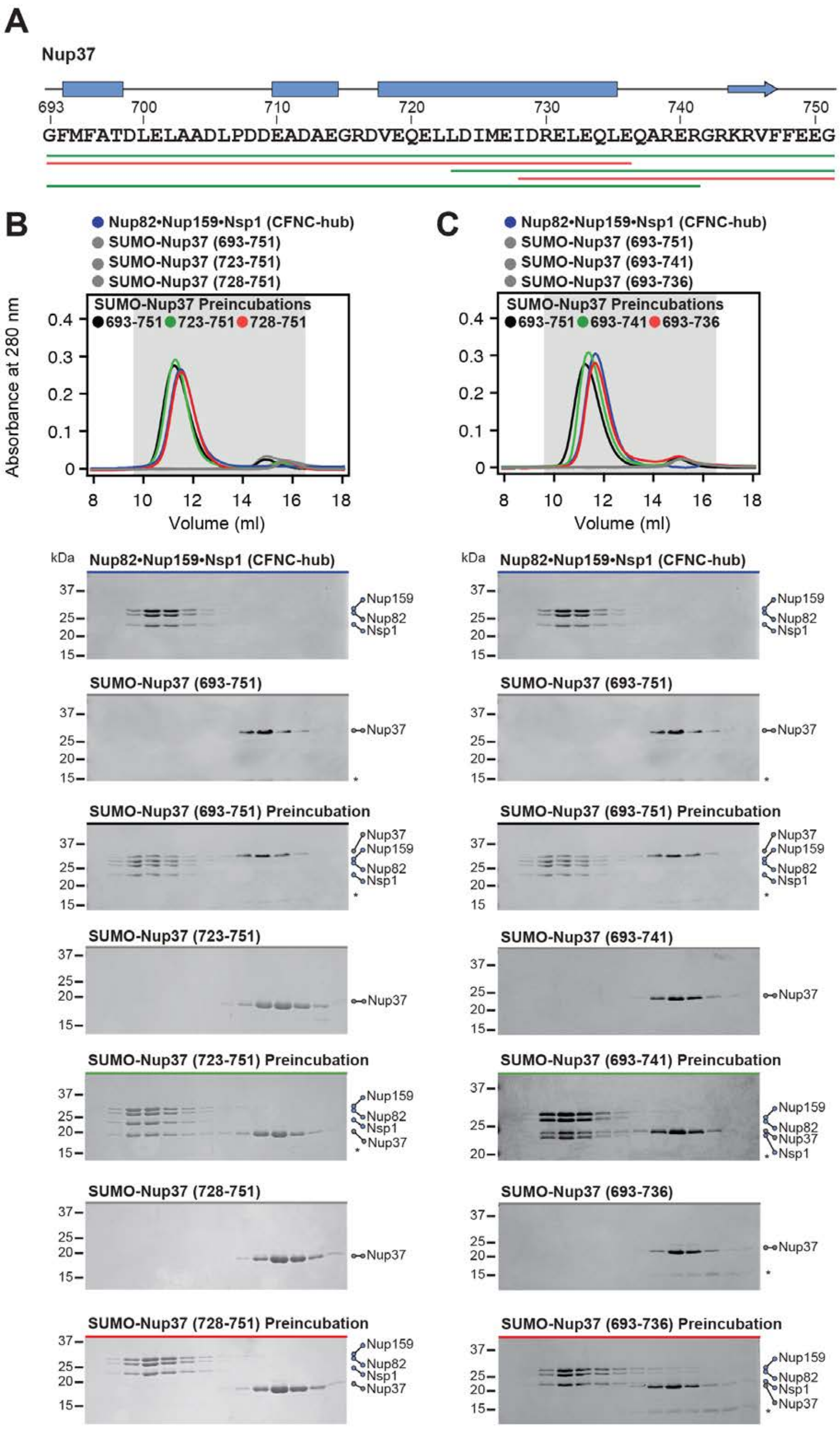
A
Nup37
693 700 710 720 730 740 750
GFMFATDLELAADLPDDEADAEGRDVEQELLDIMEIDRELEQLEQARERGRKRVFFEEG
B
Nup82•Nup159•Nsp1 (CFNC-hub)
SUMO-Nup37 (693-751)
SUMO-Nup37 (723-751)
SUMO-Nup37 (728-751)
SUMO-Nup37 Preincubations
693-751 723-751 728-751
Absorbance at 280 nm
Volume (ml)
C
Nup82•Nup159•Nsp1 (CFNC-hub)
SUMO-Nup37 (693-751)
SUMO-Nup37 (693-741)
SUMO-Nup37 (693-736)
SUMO-Nup37 Preincubations
693-751 693-741 693-736
Absorbance at 280 nm
Volume (ml)
kDa
Nup82•Nup159•Nsp1 (CFNC-hub)
Nup159
Nup82
Nsp1
SUMO-Nup37 (693-751)
Nup37
SUMO-Nup37 (693-751) Preincubation
SUMO-Nup37 (723-751)
SUMO-Nup37 (723-751) Preincubation
SUMO-Nup37 (728-751)
SUMO-Nup37 (728-751) Preincubation
SUMO-Nup37 (693-741)
SUMO-Nup37 (693-741) Preincubation
SUMO-Nup37 (693-736)
SUMO-Nup37 (693-736) Preincubation

**Fig. S14. Identification of a minimal Nup37$^{CTE}$ CFNC-hub binding region.** (**A**) Primary sequence of Nup37$^{CTE}$ with predicted secondary structure elements. Colored lines indicate Nup37$^{CTE}$ N- and C-terminal truncation fragments that were systematically tested for their ability to interact with the CNFC-hub, colored in green and red to indicate binding and no binding, respectively. Secondary structure predictions were conducted using Jpred4 (*141*). (**B**-**C**) SEC interaction analyses of CFNC-hub with (B) N-terminal SUMO-Nup37$^{CTE}$ truncations, or (C) C-terminal SUMO-Nup37$^{CTE}$ truncations, mapping a minimal binding site to residues 723-741. SEC profiles of nup complexes are shown individually (blue, gray) and after preincubation (black, red and green). SEC profiles were obtained using a Superdex 200 10/300 GL column. Gray boxes indicate fractions resolved on SDS-PAGE gels and visualized by Coomassie brilliant blue staining.

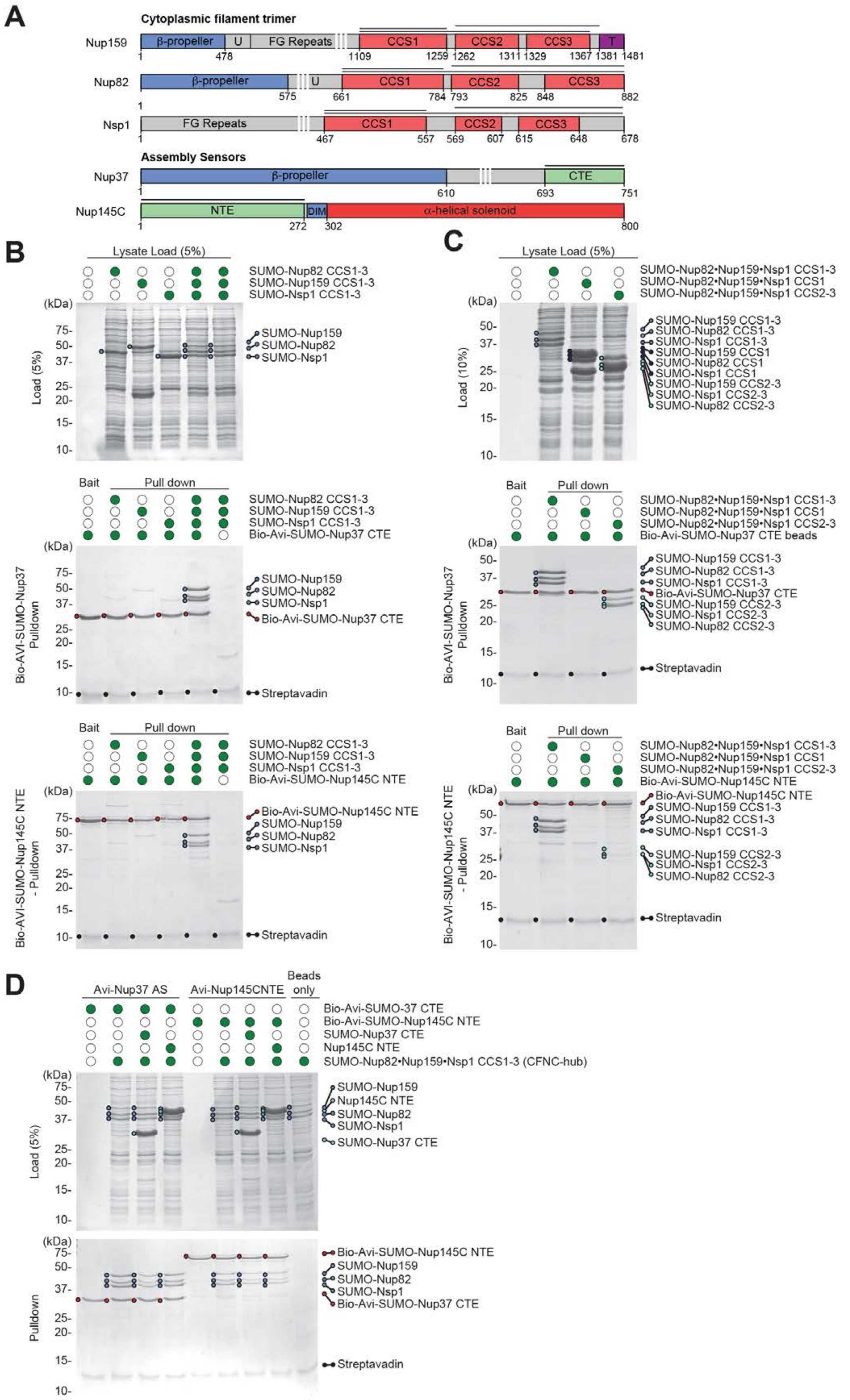
A
Cytoplasmic filament trimer
Nup159
β-propeller
U
FG Repeats
CCS1
CCS2
CCS3
T
Nup82
Nsp1
Assembly Sensors
Nup37
CTE
Nup145C
NTE
DIM
α-helical solenoid
B
Lysate Load (5%)
SUMO-Nup82 CCS1-3
SUMO-Nup159 CCS1-3
SUMO-Nsp1 CCS1-3
Load (5%)
SUMO-Nup159
SUMO-Nup82
SUMO-Nsp1
Bait
Pull down
Bio-Avi-SUMO-Nup37 CTE
Bio-AVI-SUMO-Nup37 Pulldown
Streptavadin
Bio-Avi-SUMO-Nup145C NTE
Bio-AVI-SUMO-Nup145C NTE - Pulldown
C
SUMO-Nup82•Nup159•Nsp1 CCS1-3
SUMO-Nup82•Nup159•Nsp1 CCS1
SUMO-Nup82•Nup159•Nsp1 CCS2-3
Load (10%)
SUMO-Nup159 CCS1-3
SUMO-Nup82 CCS1-3
SUMO-Nsp1 CCS1-3
SUMO-Nup159 CCS1
SUMO-Nup82 CCS1
SUMO-Nsp1 CCS1
SUMO-Nup159 CCS2-3
SUMO-Nsp1 CCS2-3
SUMO-Nup82 CCS2-3
Bio-Avi-SUMO-Nup37 CTE beads
D
Avi-Nup37 AS
Avi-Nup145CNTE
Beads only
Bio-Avi-SUMO-37 CTE
Bio-Avi-SUMO-Nup145C NTE
SUMO-Nup37 CTE
Nup145C NTE
SUMO-Nup82•Nup159•Nsp1 CCS1-3 (CFNC-hub)
Pulldown

**Fig. S15. Nup37$^{CTE}$ and Nup145C$^{NTE}$ encode assembly sensors for the CFNC-hub.** (**A**) Domain architectures of Nup159, Nup82, Nsp1, Nup37, and Nup145C. Domains are drawn as horizontal boxes with residue numbers indicated and their observed or predicted folds colored. The black lines indicate the fragments used in the pull-down analysis. (**B**-**D**) Streptavidin pulldowns using Bio-Avi-SUMO-Nup37$^{CTE}$ or Bio-Avi-SUMO-Nup145C$^{NTE}$ baits with (B) bacterial lysates, containing either Nup82, Nup159, and Nsp1 CCS1-3 fragments individually or a mixture of all three, demonstrating exclusive binding to the hetero-trimeric CFNC-hub, (C) bacterial lysates containing either the Nup82•Nup159•Nsp1 CCS1-3 (CFNC-hub), or hetero-trimeric CCS1 and CCS2-3 segments, demonstrating binding to CCS2-3 albeit to a lesser extent, and (D) bacterial lysates containing CFNC-hub and a ~10 fold excess of purified SUMO-Nup37$^{CTE}$ or Nup145C$^{NTE}$ competitor, demonstrating mutually exclusive binding events. SDS-PAGE gels are visualized by Coomassie brilliant blue staining.

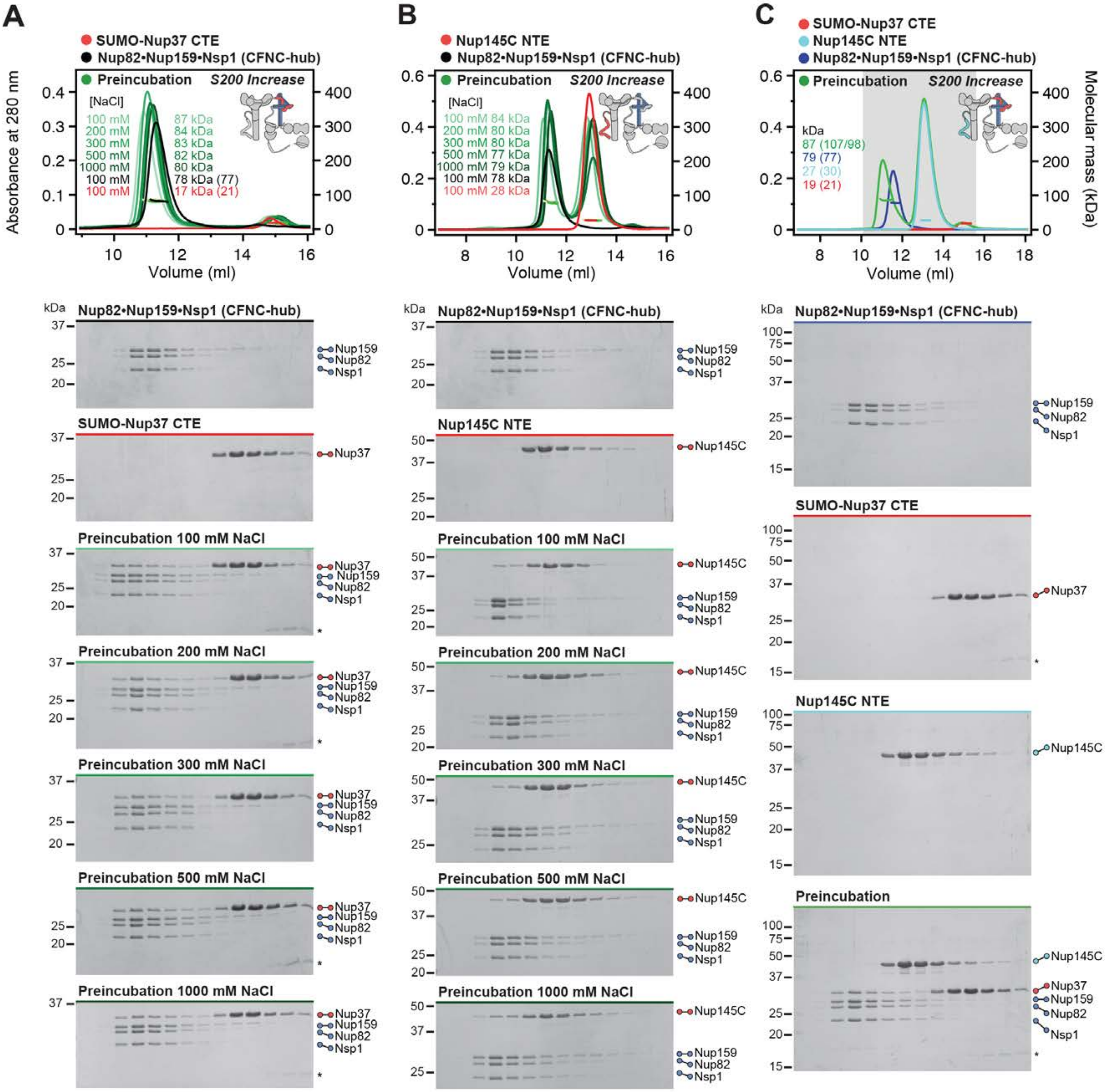
A
SUMO-Nup37 CTE
Nup82•Nup159•Nsp1 (CFNC-hub)
Preincubation
S200 Increase
[NaCl]
100 mM 87 kDa
200 mM 84 kDa
300 mM 83 kDa
500 mM 82 kDa
1000 mM 80 kDa
100 mM 78 kDa (77)
100 mM 17 kDa (21)
Absorbance at 280 nm
Volume (ml)
B
Nup145C NTE
Nup82•Nup159•Nsp1 (CFNC-hub)
Preincubation
S200 Increase
[NaCl]
100 mM 84 kDa
200 mM 80 kDa
300 mM 80 kDa
500 mM 77 kDa
1000 mM 79 kDa
100 mM 78 kDa
100 mM 28 kDa
Volume (ml)
C
SUMO-Nup37 CTE
Nup145C NTE
Nup82•Nup159•Nsp1 (CFNC-hub)
Preincubation
S200 Increase
kDa
87 (107/98)
79 (77)
27 (30)
19 (21)
Volume (ml)
Molecular mass (kDa)
kDa
Nup82•Nup159•Nsp1 (CFNC-hub)
SUMO-Nup37 CTE
Nup145C NTE
Preincubation
Preincubation 100 mM NaCl
Preincubation 200 mM NaCl
Preincubation 300 mM NaCl
Preincubation 500 mM NaCl
Preincubation 1000 mM NaCl
Nup159
Nup82
Nsp1
Nup37
Nup145C
*

**Fig. S16. Nup37$^{CTE}$ and Nup145C$^{NTE}$ bind CFNC-hub in a mutually exclusive fashion.** SEC-MALS interaction analysis of the CFNC-hub with (**A**) SUMO-Nup37$^{CTE}$ and (**B**) Nup145C$^{NTE}$ across increasing buffer NaCl concentrations, establishing the hydrophobic nature of both interactions. (**C**) SEC-MALS competition analysis of SUMO-Nup37$^{CTE}$ and Nup145C$^{NTE}$ binding to CFNC-hub, establishing mutually exclusive overlapping binding sites. SEC-MALS profiles of nup complexes are shown individually (red, blue, cyan, and black) and after preincubation (green). SEC profiles were obtained using a Superdex 200 Increase 10/300 GL column. Measured molecular masses are indicated, with the respective theoretical masses shown in parentheses. Inset schematics display tested protein samples colored to match the corresponding chromatogram trace. Gray boxes indicate fractions resolved on SDS-PAGE gels and visualized by Coomassie brilliant blue staining.

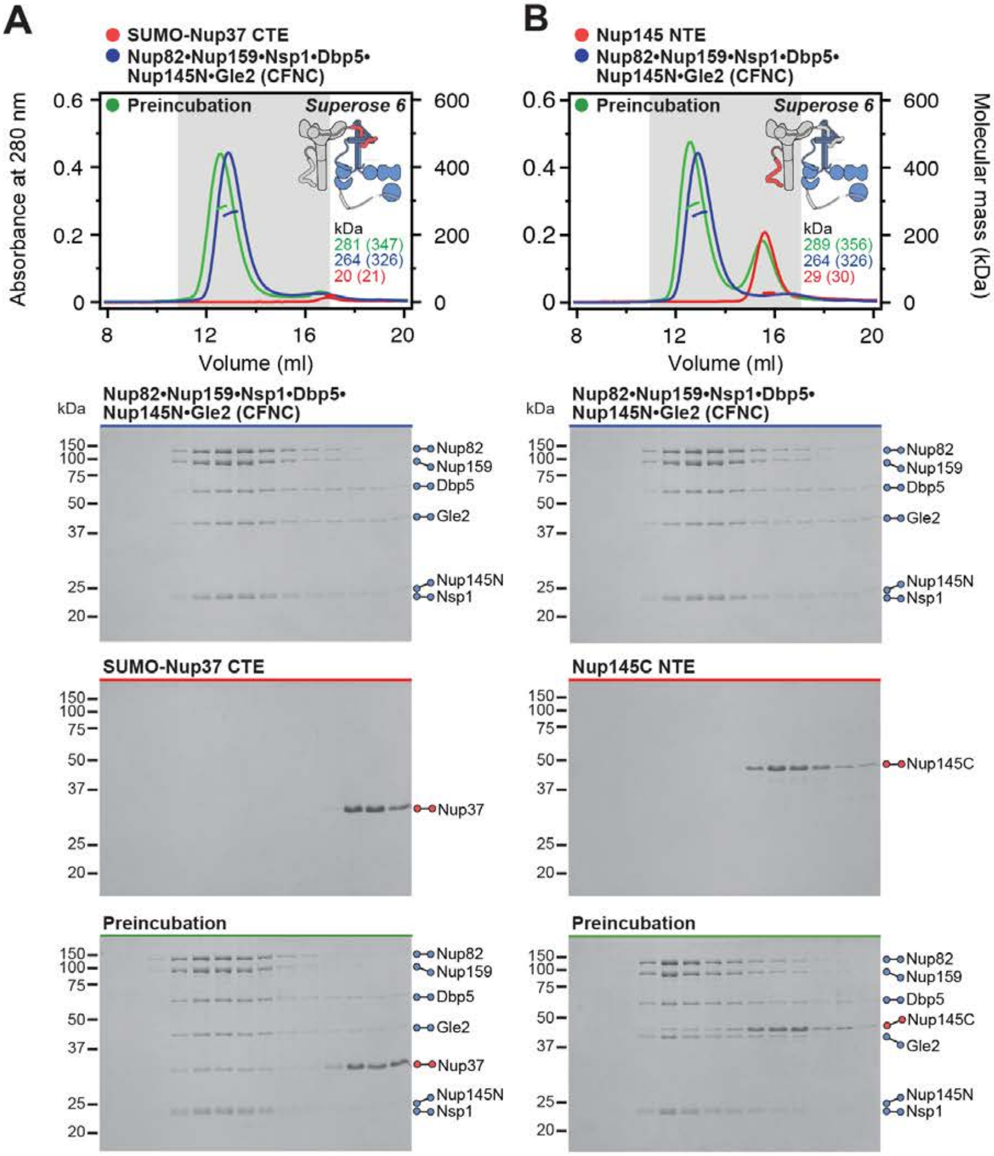

A
SUMO-Nup37 CTE
Nup82•Nup159•Nsp1•Dbp5•
Nup145N•Gle2 (CFNC)
Preincubation
Superose 6
Absorbance at 280 nm
kDa
281 (347)
264 (326)
20 (21)
Volume (ml)
Nup82•Nup159•Nsp1•Dbp5•
Nup145N•Gle2 (CFNC)
Nup82
Nup159
Dbp5
Gle2
Nup145N
Nsp1
SUMO-Nup37 CTE
Nup37
Preincubation
B
Nup145 NTE
Nup82•Nup159•Nsp1•Dbp5•
Nup145N•Gle2 (CFNC)
Preincubation
Superose 6
Molecular mass (kDa)
kDa
289 (356)
264 (326)
29 (30)
Volume (ml)
Nup145C NTE
Nup145C
Preincubation

**Fig. S17. Nup37$^{CTE}$ and Nup145C$^{NTE}$ bind the intact CFNC hetero-hexamer.** SEC-MALS interaction analysis of CFNC with (**A**) SUMO-Nup37$^{CTE}$ and (**B**) Nup145C$^{NTE}$. SEC-MALS profiles of nup complexes are shown individually (red, blue) and after preincubation (green). SEC profiles were obtained using a Superose 6 10/300 GL column. Measured molecular masses are indicated, with the respective theoretical masses shown in parentheses. Inset schematics display tested protein samples colored to match the corresponding chromatogram trace. Gray boxes indicate fractions resolved on SDS-PAGE gels and visualized by Coomassie brilliant blue staining.

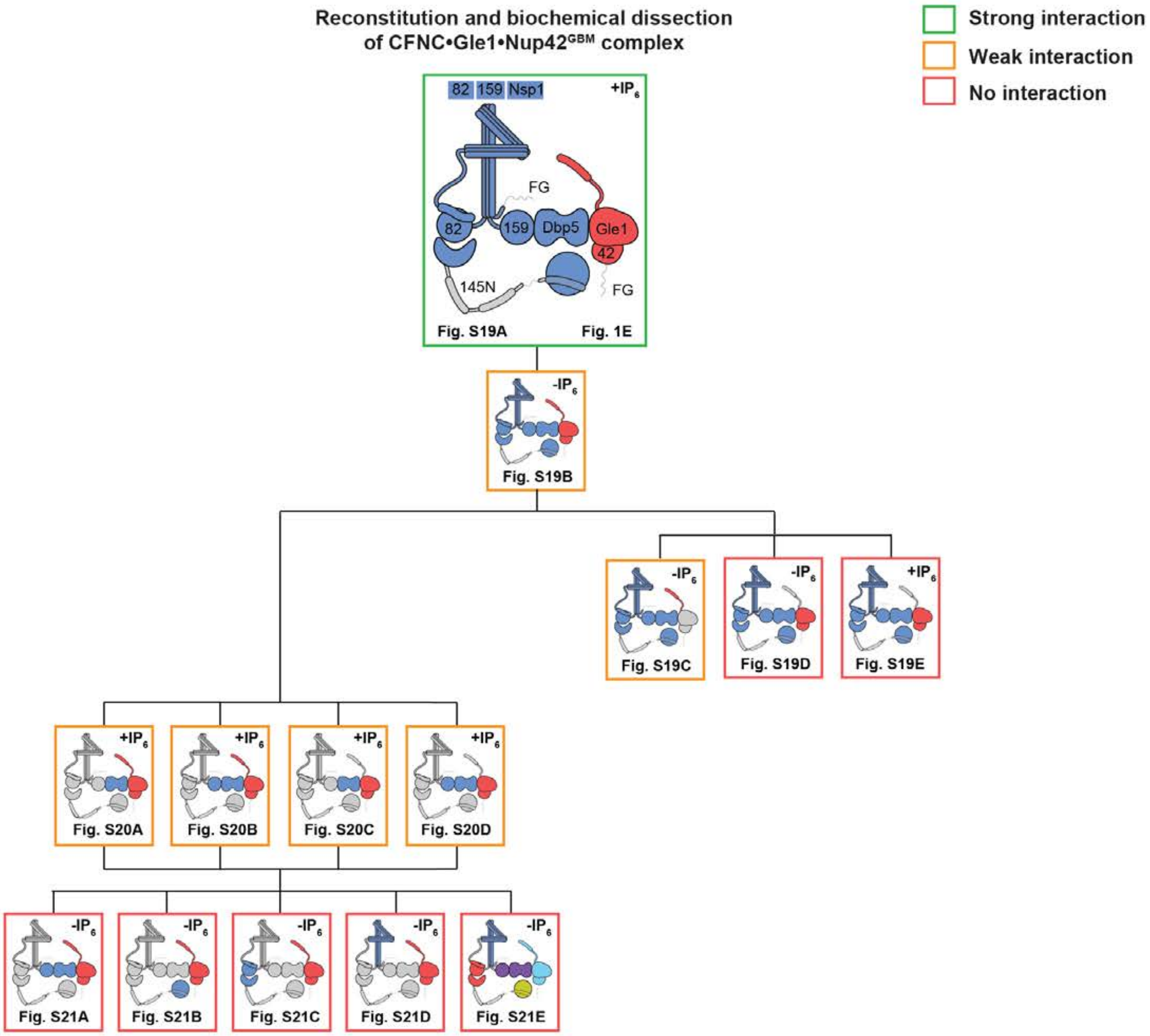
Reconstitution and biochemical dissection of CFNC•Gle1•Nup42GBM complex
Strong interaction
Weak interaction
No interaction
82
159
Nsp1
+IP6
FG
82
159
Dbp5
Gle1
42
145N
FG
Fig. S19A
Fig. 1E
-IP6
Fig. S19B
-IP6
Fig. S19C
-IP6
Fig. S19D
+IP6
Fig. S19E
+IP6
Fig. S20A
+IP6
Fig. S20B
+IP6
Fig. S20C
+IP6
Fig. S20D
-IP6
Fig. S21A
-IP6
Fig. S21B
-IP6
Fig. S21C
-IP6
Fig. S21D
-IP6
Fig. S21E

**Fig. S18. Reconstitution and dissection strategy for the CFNC•Gle1•Nup42$^{GBM}$ hetero-octamer.** Summary of the SEC-MALS interaction analyses of the indicated figure panels. Each box in the flow chart represents a SEC-MALS interaction experiment, involving individually purified Gle1•Nup42$^{GBM}$ and CNFC sub-complexes that were tested for their ability to form stochiometric complexes, colored by the experimental outcome; strong interaction (green), weak interaction (orange), no interaction (red). The inset schematics are colored to indicate the individually purified Gle1•Nup42$^{GBM}$ and CFNC sub-complexes of each SEC-MALS interaction experiment. Gray regions stipulate proteins omitted from each analysis and the presence and absence of $IP_6$ in the interaction analyses are indicated.

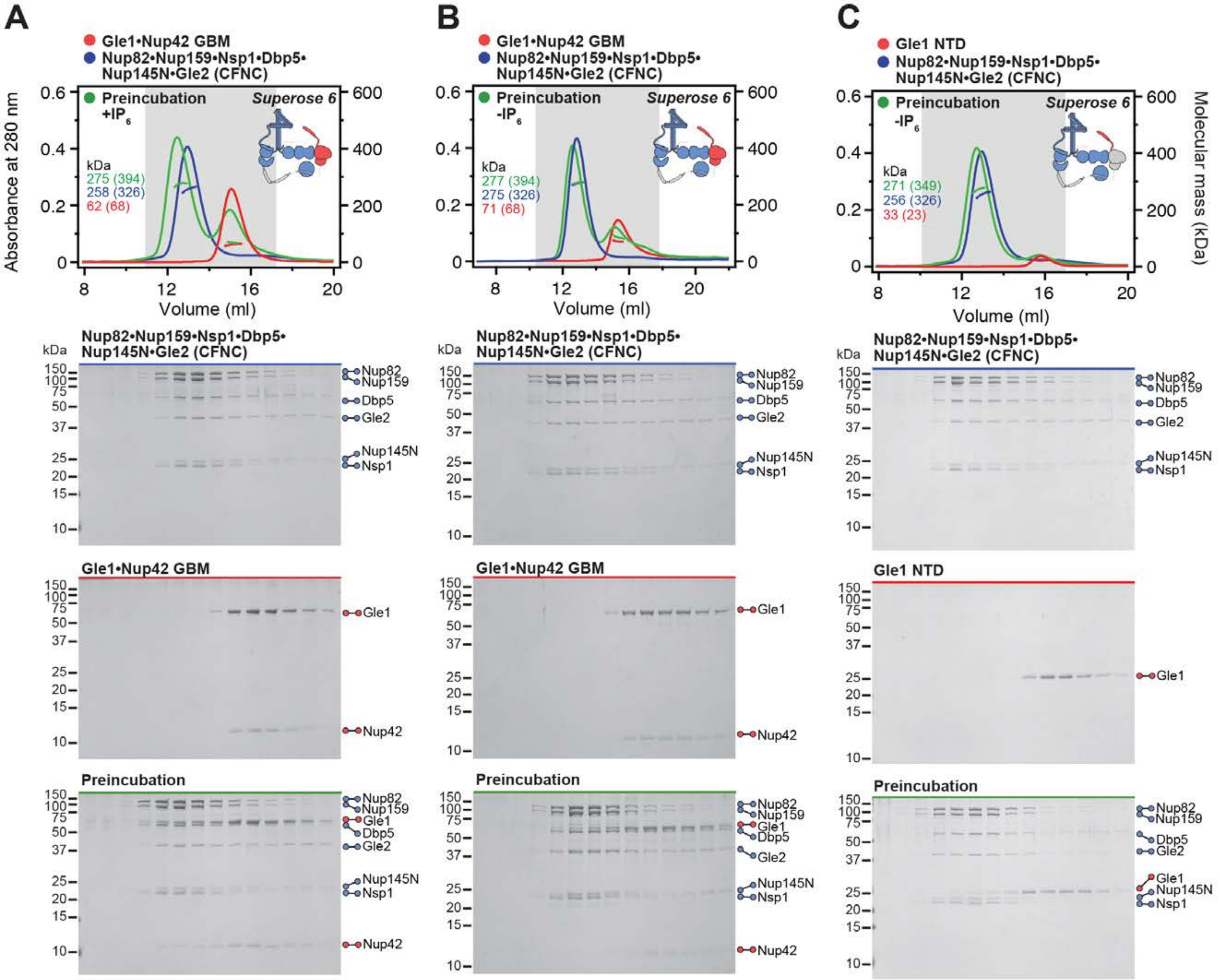

A
B
C
Gle1•Nup42 GBM
Nup82•Nup159•Nsp1•Dbp5•Nup145N•Gle2 (CFNC)
Preincubation +IP6
Preincubation -IP6
Gle1 NTD
Superose 6
Absorbance at 280 nm
Molecular mass (kDa)
Volume (ml)
kDa
275 (394)
258 (326)
62 (68)
277 (394)
275 (326)
71 (68)
271 (349)
256 (326)
33 (23)
Nup82
Nup159
Dbp5
Gle2
Nup145N
Nsp1
Gle1
Nup42
Preincubation

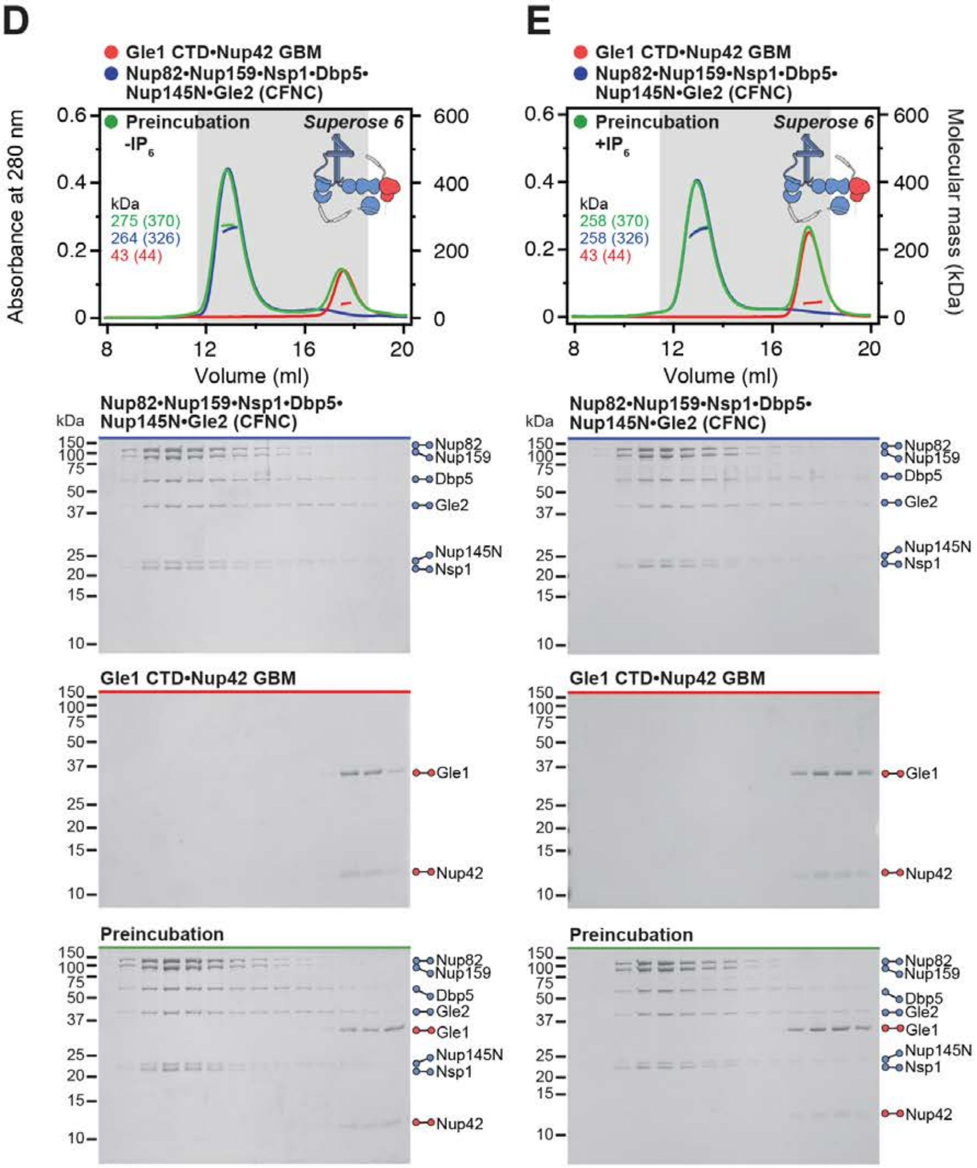
D
Gle1 CTD•Nup42 GBM
Nup82•Nup159•Nsp1•Dbp5•Nup145N•Gle2 (CFNC)
Preincubation -IP6
Superose 6
kDa
275 (370)
264 (326)
43 (44)
Absorbance at 280 nm
Volume (ml)
Nup82•Nup159•Nsp1•Dbp5•Nup145N•Gle2 (CFNC)
Nup82
Nup159
Dbp5
Gle2
Nup145N
Nsp1
Gle1 CTD•Nup42 GBM
Gle1
Nup42
Preincubation
E
Gle1 CTD•Nup42 GBM
Nup82•Nup159•Nsp1•Dbp5•Nup145N•Gle2 (CFNC)
Preincubation +IP6
Superose 6
kDa
258 (370)
258 (326)
43 (44)
Molecular mass (kDa)
Volume (ml)
Nup82•Nup159•Nsp1•Dbp5•Nup145N•Gle2 (CFNC)
Gle1 CTD•Nup42 GBM
Preincubation

**Fig. S19. Gle1•Nup42$^{GBM}$ binds the CFNC via a distributed binding site.** (**A**, **B**) SEC-MALS interaction analysis of Gle1•Nup42$^{GBM}$ and CFNC in the (A) presence and (B) absence of $IP_6$, demonstrating that formation of a stable stoichiometric CFNC•Gle1•Nup42$^{GBM}$ hetero-octamer is $IP_6$-dependent (corresponding to Fig. 1E). (**C**, **D**) SEC-MALS interaction analyses of CFNC in the absence of $IP_6$ with (C) Gle1$^{NTD}$ and (D) Gle1$^{CTD}$•Nup42$^{GBM}$, demonstrating that Gle1$^{NTD}$ contributes to CNFC binding. (**E**) SEC-MALS interaction analysis of CNFC in the presence of $IP_6$ with Gle1$^{CTD}$•Nup42$^{GBM}$, showing that Gle1$^{CTD}$ is insufficient for CFNC binding. SEC-MALS profiles of nup complexes are shown individually (red and blue) and after preincubation (green). SEC profiles were obtained using a Superose 6 10/300 GL column. Measured molecular masses are indicated, with the respective theoretical masses shown in parentheses. Inset schematics display tested protein samples colored to match the corresponding chromatogram trace. Gray boxes indicate fractions resolved on SDS-PAGE gels and visualized by Coomassie brilliant blue staining.

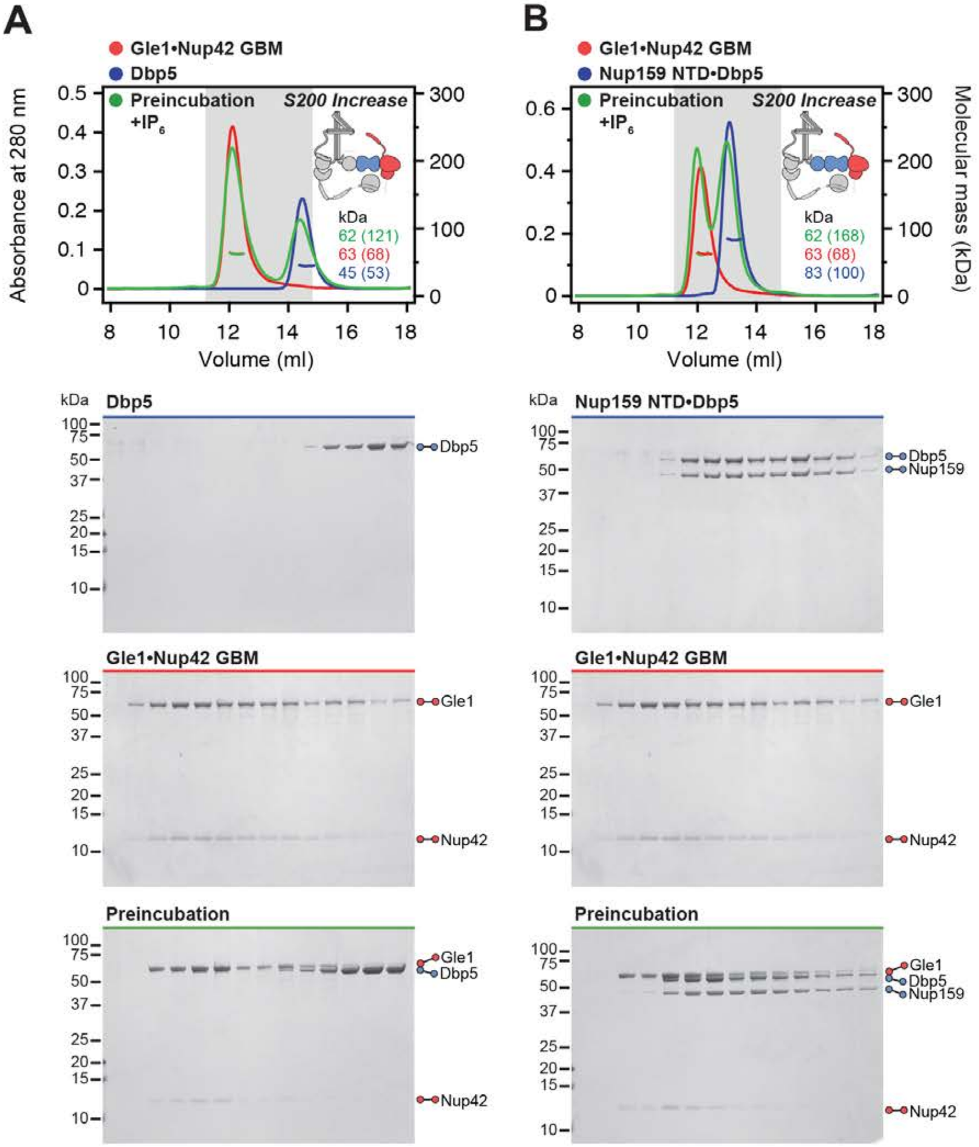
A
Gle1•Nup42 GBM
Dbp5
Preincubation
+IP6
S200 Increase
Absorbance at 280 nm
kDa
62 (121)
63 (68)
45 (53)
Volume (ml)
B
Gle1•Nup42 GBM
Nup159 NTD•Dbp5
Preincubation
+IP6
S200 Increase
Molecular mass (kDa)
kDa
62 (168)
63 (68)
83 (100)
Volume (ml)
Dbp5
Nup159 NTD•Dbp5
Nup159
Gle1•Nup42 GBM
Gle1
Nup42
Preincubation

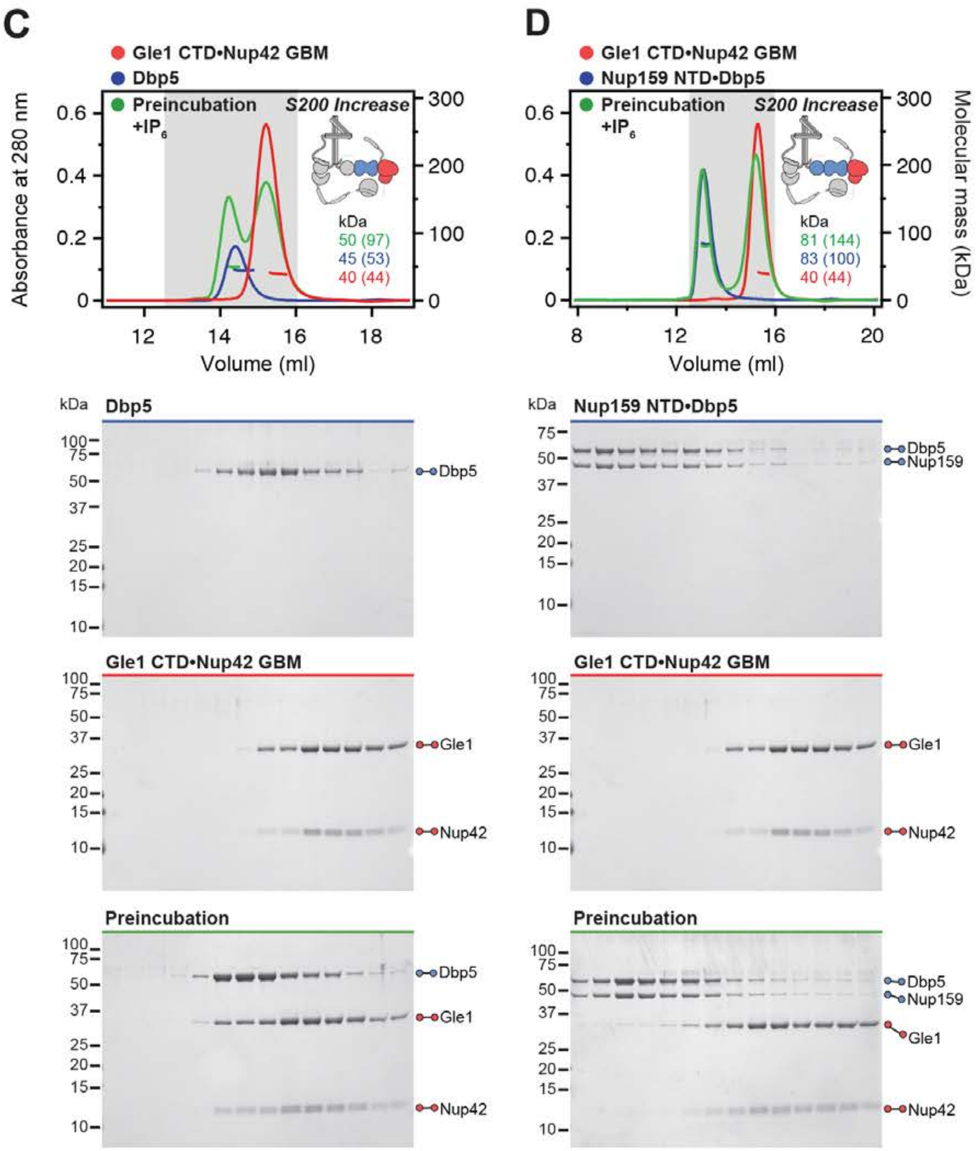

Sup Fig. S20

Hoelz et al., 2021b

**Fig. S20. Both Nup159$^{NTD}$ and Gle1$^{NTD}$ are dispensable for the Gle1•Nup42$^{GBM}$-Dbp5 interaction.** (**A**, **B**) SEC-MALS interaction analyses of Gle1•Nup42$^{GBM}$ in the presence of $IP_6$ with (A) Dbp5 and (B) Nup159$^{NTD}$•Dbp5, showing that Nup159$^{NTD}$ is dispensable for the Gle1•Nup42$^{GBM}$-Dbp5 interaction. (**C**, **D**) SEC-MALS interaction analyses of Gle1$^{CTD}$•Nup42$^{GBM}$ in the presence of $IP_6$ with (C) Dbp5 and (D) Nup159$^{NTD}$•Dbp5, showing that Gle1$^{NTD}$ is dispensable for the Gle1•Nup42$^{GBM}$-Dbp5 interaction**.** SEC-MALS profiles of nup complexes are shown individually (red, blue) and after preincubation (green). SEC profiles were obtained using a Superdex 200 Increase 10/300 GL column. Measured molecular masses are indicated, with the respective theoretical masses shown in parentheses. Inset schematics display tested protein samples colored to match the corresponding chromatogram trace. Gray boxes indicate fractions resolved on SDS-PAGE gels and visualized by Coomassie brilliant blue staining.

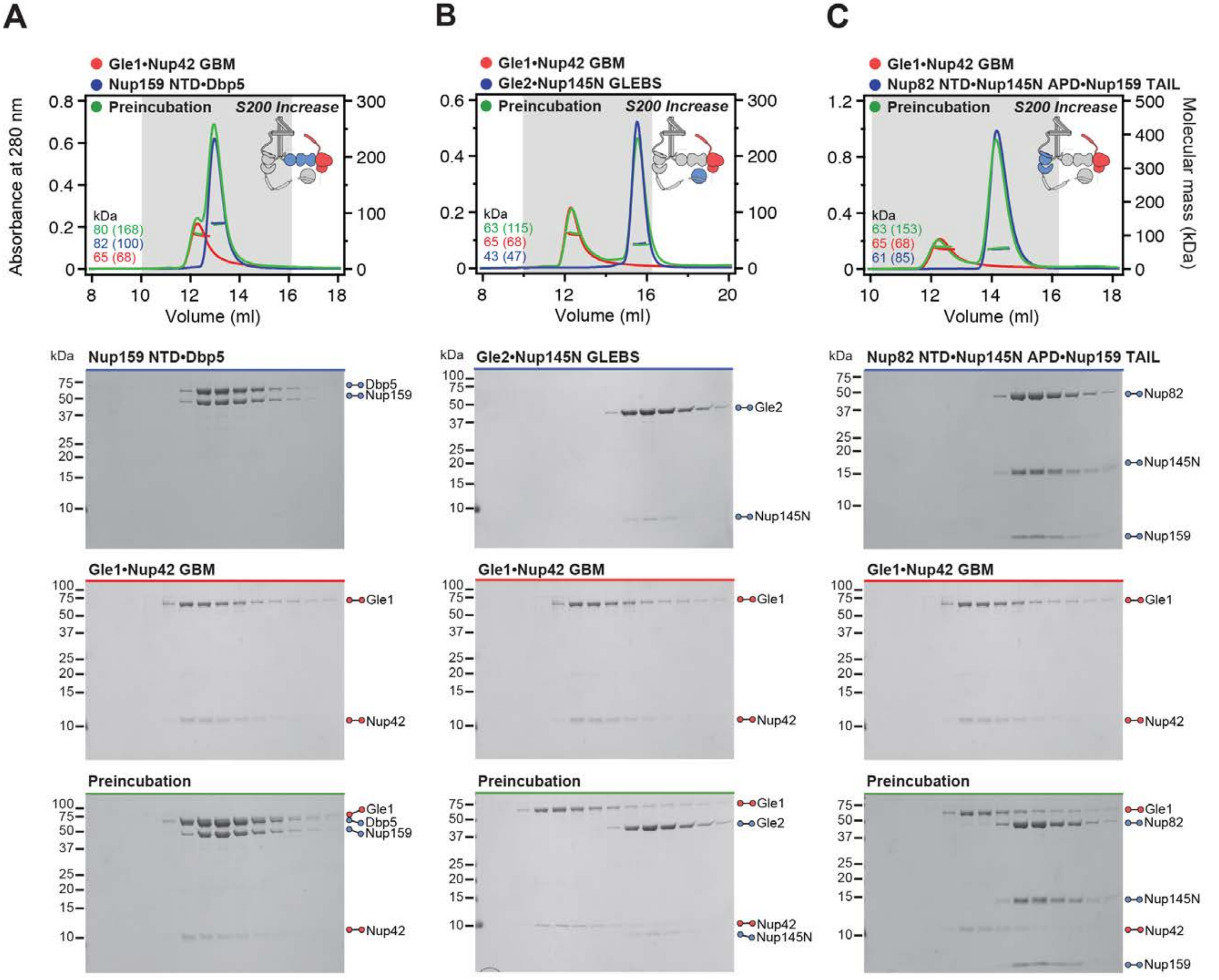
A
Gle1•Nup42 GBM
Nup159 NTD•Dbp5
Preincubation
S200 Increase
Absorbance at 280 nm
kDa
80 (168)
82 (100)
65 (68)
Volume (ml)
Nup159 NTD•Dbp5
Dbp5
Nup159
Gle1•Nup42 GBM
Gle1
Nup42
Preincubation
Gle1
Dbp5
Nup159
Nup42
B
Gle1•Nup42 GBM
Gle2•Nup145N GLEBS
Preincubation
S200 Increase
kDa
63 (115)
65 (68)
43 (47)
Volume (ml)
Gle2•Nup145N GLEBS
Gle2
Nup145N
Gle1•Nup42 GBM
Gle1
Nup42
Preincubation
Gle1
Gle2
Nup42
Nup145N
C
Gle1•Nup42 GBM
Nup82 NTD•Nup145N APD•Nup159 TAIL
Preincubation
S200 Increase
Molecular mass (kDa)
kDa
63 (153)
65 (68)
61 (85)
Volume (ml)
Nup82 NTD•Nup145N APD•Nup159 TAIL
Nup82
Nup145N
Nup159
Gle1•Nup42 GBM
Gle1
Nup42
Preincubation
Gle1
Nup82
Nup145N
Nup42
Nup159

D

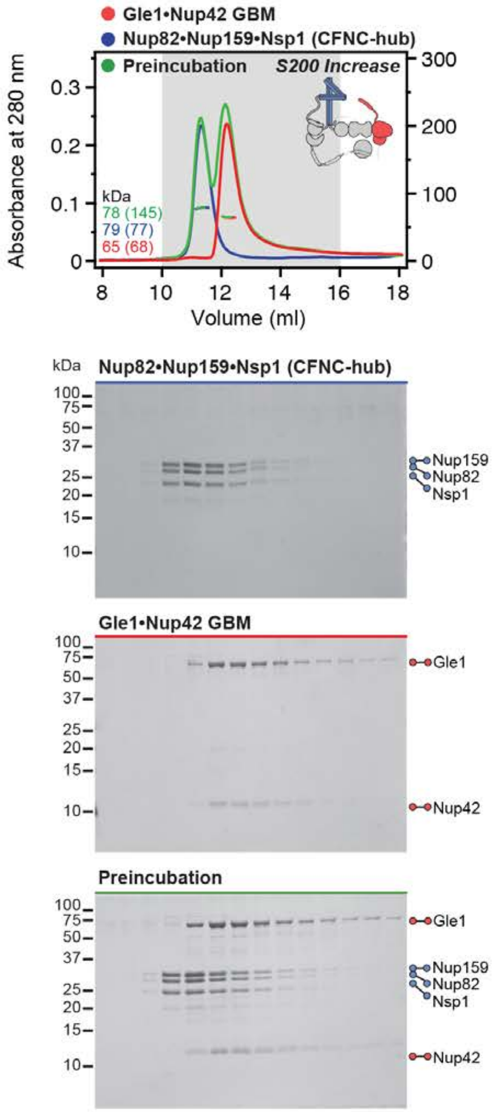

Gle1•Nup42 GBM
Nup82•Nup159•Nsp1 (CFNC-hub)
Preincubation
S200 Increase
Absorbance at 280 nm
Volume (ml)
kDa
78 (145)
79 (77)
65 (68)
Nup82•Nup159•Nsp1 (CFNC-hub)
Nup159
Nup82
Nsp1
Gle1•Nup42 GBM
Gle1
Nup42
Preincubation
Gle1
Nup159
Nup82
Nsp1
Nup42

E

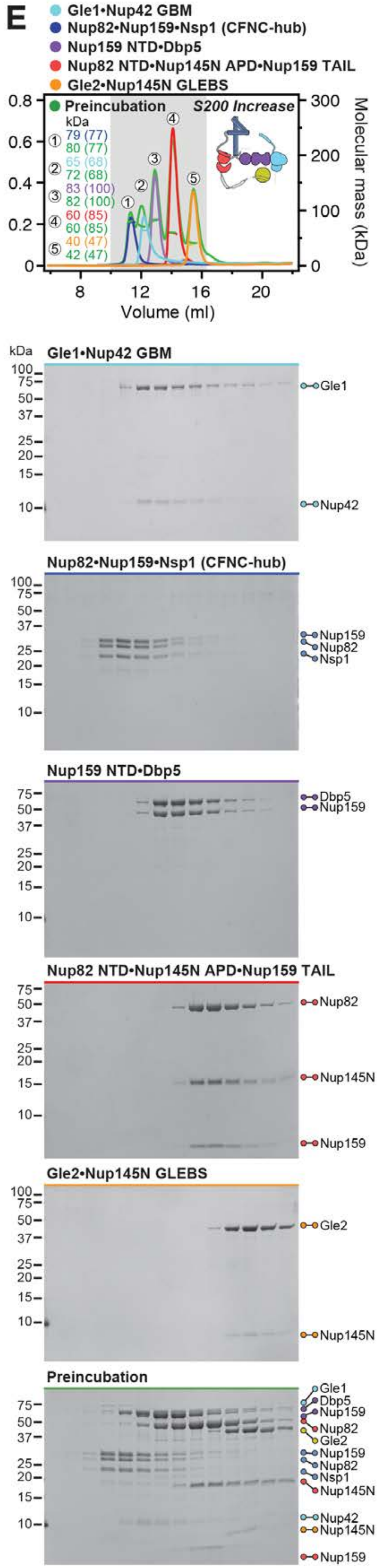

Gle1•Nup42 GBM
Nup82•Nup159•Nsp1 (CFNC-hub)
Nup159 NTD•Dbp5
Nup82 NTD•Nup145N APD•Nup159 TAIL
Gle2•Nup145N GLEBS
Preincubation
S200 Increase
kDa
79 (77)
80 (77)
65 (68)
72 (68)
83 (100)
82 (100)
60 (85)
60 (85)
40 (47)
42 (47)
Molecular mass (kDa)
Volume (ml)
Gle1•Nup42 GBM
Gle1
Nup42
Nup82•Nup159•Nsp1 (CFNC-hub)
Nup159
Nup82
Nsp1
Nup159 NTD•Dbp5
Dbp5
Nup159
Nup82 NTD•Nup145N APD•Nup159 TAIL
Nup82
Nup145N
Nup159
Gle2•Nup145N GLEBS
Gle2
Nup145N
Preincubation
Gle1
Dbp5
Nup159
Nup82
Gle2
Nup159
Nup82
Nsp1
Nup145N
Nup42
Nup145N
Nup159

**Fig. S21. A distributed CNFC binding site mediates the Gle1•Nup42$^{GBM}$ interaction.** (**A**-**E**) SEC-MALS interaction analyses of Gle1•Nup42$^{GBM}$ in the absence of $IP_6$ with (A) Nup159$^{NTD}$•Dbp5, (B) Gle2•Nup145N$^{GLEBS}$, (C) Nup82$^{NTD}$•Nup145N$^{APD}$•Nup159$^{TAIL}$, (D) CFNC-hub, and (E) a mixture of all four CNFC subcomplexes, demonstrating that none of the four CNFC sub-complexes recapitulates the binding of intact CFNC. SEC-MALS profiles of nup complexes are shown individually (red, blue, purple and yellow) and after preincubation (green). SEC profiles were obtained using a Superdex 200 Increase 10/300 GL column. Measured molecular masses are indicated, with the respective theoretical masses shown in parentheses. Inset schematics display tested protein samples colored to match the corresponding chromatogram trace. Gray boxes indicate fractions resolved on SDS-PAGE gels and visualized by Coomassie brilliant blue staining.

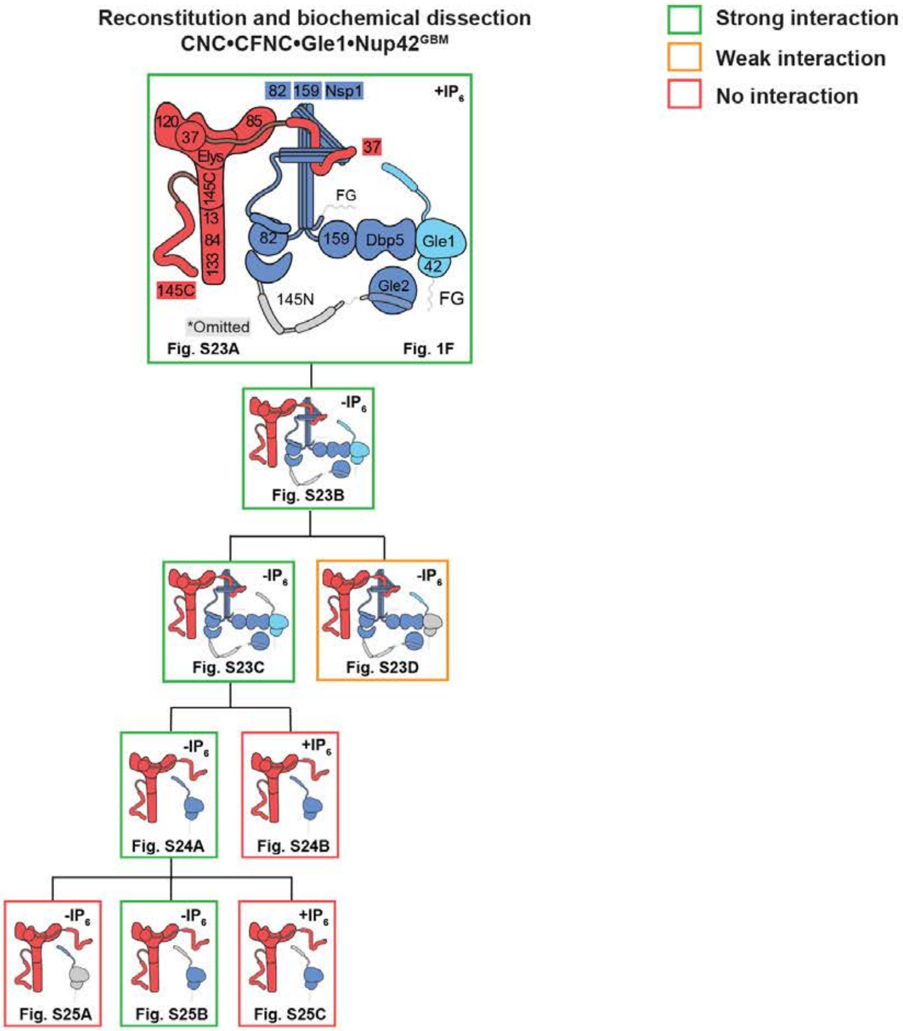

Reconstitution and biochemical dissection
CNC•CFNC•Gle1•Nup42GBM
Strong interaction
Weak interaction
No interaction
82 159 Nsp1
+IP6
120
37
85
Elys
145C
13
84
133
37
FG
82
159
Dbp5
Gle1
42
145C
145N
Gle2
FG
*Omitted
Fig. S23A
Fig. 1F
-IP6
Fig. S23B
-IP6
Fig. S23C
-IP6
Fig. S23D
-IP6
Fig. S24A
+IP6
Fig. S24B
-IP6
Fig. S25A
-IP6
Fig. S25B
+IP6
Fig. S25C

**Fig. S22. Reconstitution and dissection strategy for the 16-protein CNC•CFNC•Gle1•Nup42$^{GBM}$.** Summary of the SEC-MALS interaction analyses of the indicated figure panels. Each box in the flow chart represents a SEC-MALS interaction experiment, involving individually purified Gle1•Nup42$^{GBM}$, CNFC, and CNC sub-complexes that were tested for their ability to form stochiometric complexes, colored by the experimental outcome; strong interaction (green), weak interaction (orange), no interaction (red). The inset schematics are colored to indicate the individually purified Gle1•Nup42$^{GBM}$, CFNC, and CNC sub-complexes of each SEC-MALS interaction experiment. Gray regions stipulate proteins omitted from each analysis and the presence and absence of $IP_6$ in the interaction analyses are indicated.

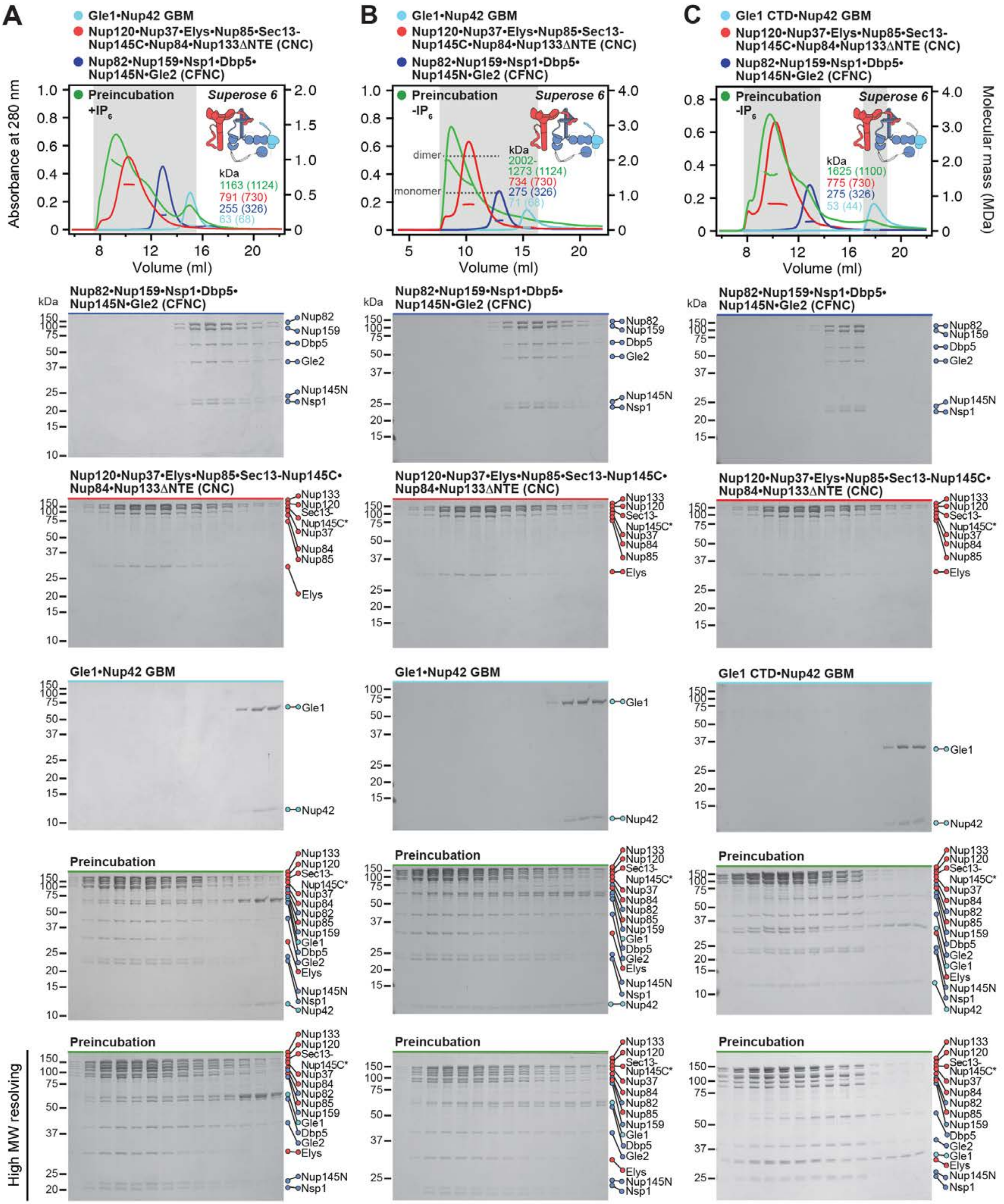
A
B
C
Gle1•Nup42 GBM
Gle1 CTD•Nup42 GBM
Nup120•Nup37•Elys•Nup85•Sec13-Nup145C•Nup84•Nup133ΔNTE (CNC)
Nup82•Nup159•Nsp1•Dbp5•Nup145N•Gle2 (CFNC)
Preincubation +IP6
Preincubation -IP6
Superose 6
Absorbance at 280 nm
Molecular mass (MDa)
Volume (ml)
kDa
1163 (1124)
791 (730)
255 (326)
63 (68)
dimer
monomer
2002-1273 (1124)
734 (730)
275 (326)
71 (68)
1625 (1100)
775 (730)
275 (326)
53 (44)
Nup82
Nup159
Dbp5
Gle2
Nup145N
Nsp1
Nup133
Nup120
Sec13-Nup145C*
Nup37
Nup84
Nup85
Elys
Gle1
Nup42
Preincubation
High MW resolving

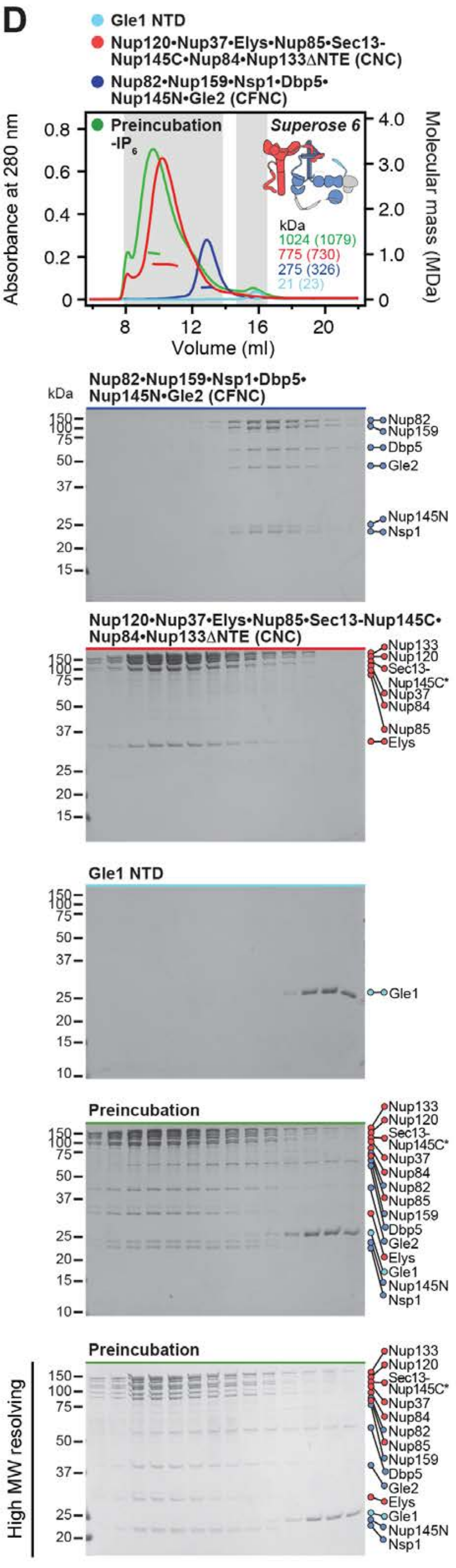

D
Gle1 NTD
Nup120•Nup37•Elys•Nup85•Sec13-Nup145C•Nup84•Nup133ΔNTE (CNC)
Nup82•Nup159•Nsp1•Dbp5•Nup145N•Gle2 (CFNC)
Preincubation -IP6
Superose 6
Absorbance at 280 nm
Molecular mass (MDa)
Volume (ml)
kDa
1024 (1079)
775 (730)
275 (326)
21 (23)
Nup82•Nup159•Nsp1•Dbp5•Nup145N•Gle2 (CFNC)
Nup82
Nup159
Dbp5
Gle2
Nup145N
Nsp1
Nup120•Nup37•Elys•Nup85•Sec13-Nup145C•Nup84•Nup133ΔNTE (CNC)
Nup133
Nup120
Sec13-Nup145C*
Nup37
Nup84
Nup85
Elys
Gle1 NTD
Gle1
Preincubation
High MW resolving

**Fig. S23. Reconstitution of a 16-protein coat-cytoplasmic filament nup complex.** (**A**) SEC-MALS interaction analysis of CNC, CFNC, and Gle1•Nup42$^{GBM}$ in the presence of $IP_6$, demonstrating the formation of an ~1,163 kDa monodisperse, stoichiometric 16-protein CNC•CFNC complex (corresponding to Fig. 1F). (**B**) SEC-MALS interaction analysis of CNC, CFNC, and Gle1•Nup42$^{GBM}$ in the absence of $IP_6$, demonstrating the formation of a peak corresponding to a monomeric and a dimeric assembly state of the 16-protein CNC•CFNC complex with molecular masses ranging from ~2,002 to ~1,273 kDa. (**C, D**) SEC-MALS analysis of the CNC•CFNC interaction with (C) Gle1$^{CTD}$•Nup42$^{GBM}$ and (D) Gle1$^{NTD}$ in the absence of $IP_6$, demonstrating Gle1$^{CTD}$• Nup42$^{GBM}$ incorporation into the CNC-CFNC complex and partial dimer formation, whereas the Gle1$^{NTD}$ exhibits barely detectable binding. SEC-MALS profiles of nup complexes are shown individually (red, blue and cyan) and after preincubation (green). SEC profiles were obtained using a Superose 6 10/300 GL column. Measured molecular masses are indicated, with the respective theoretical masses shown in parentheses. Inset schematics display tested protein samples colored to match the corresponding chromatogram trace. Gray boxes indicate fractions resolved on SDS-PAGE gels and visualized by Coomassie brilliant blue staining.

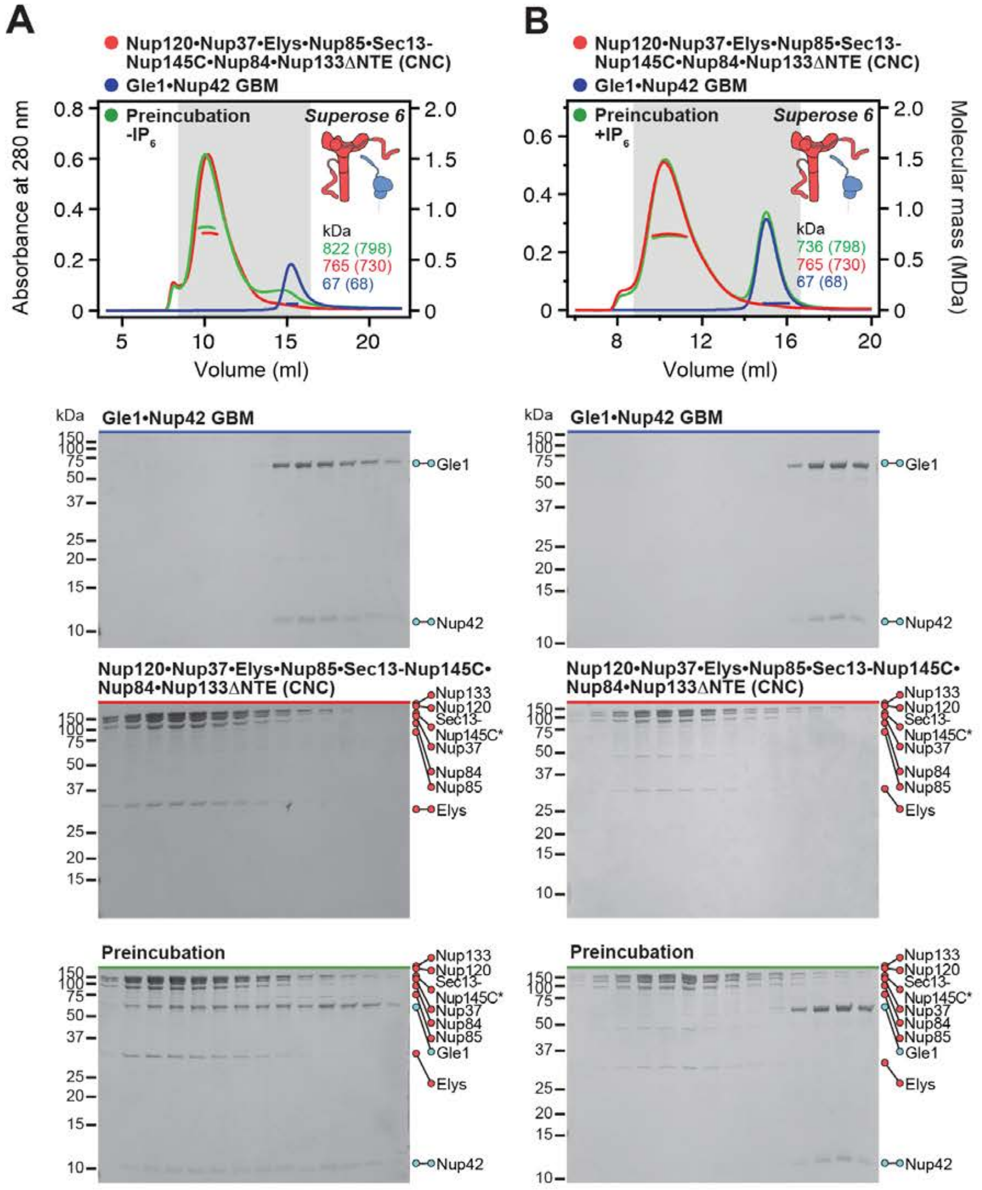
A
B
Nup120•Nup37•Elys•Nup85•Sec13-Nup145C•Nup84•Nup133ΔNTE (CNC)
Gle1•Nup42 GBM
Preincubation -IP6
Preincubation +IP6
Superose 6
kDa
822 (798)
765 (730)
67 (68)
736 (798)
Absorbance at 280 nm
Molecular mass (MDa)
Volume (ml)
Gle1
Nup42
Nup133
Nup120
Sec13-Nup145C*
Nup37
Nup84
Nup85
Elys
Preincubation

**Fig. S24. Gle1•Nup42$^{GBM}$ binds to the CNC in an $IP_6$-dependent fashion. (A, B)** SEC-MALS analyses of the interaction between Gle1•Nup42$^{GBM}$ and CNC (A) in the absence and (B) presence of $IP_6$, demonstrating the formation of a stoichiometric monodisperse ~822 kDa Gle1•Nup42$^{GBM}$•CNC complex exclusively in the absence of $IP_6$. SEC-MALS profiles of nup complexes are shown individually (red, blue) and after preincubation (green). SEC profiles were obtained using a Superose 6 10/300 GL column. Measured molecular masses are indicated, with the respective theoretical masses shown in parentheses. Inset schematics display tested protein samples colored to match the corresponding chromatogram trace. Gray boxes indicate fractions resolved on SDS-PAGE gels and visualized by Coomassie brilliant blue staining.

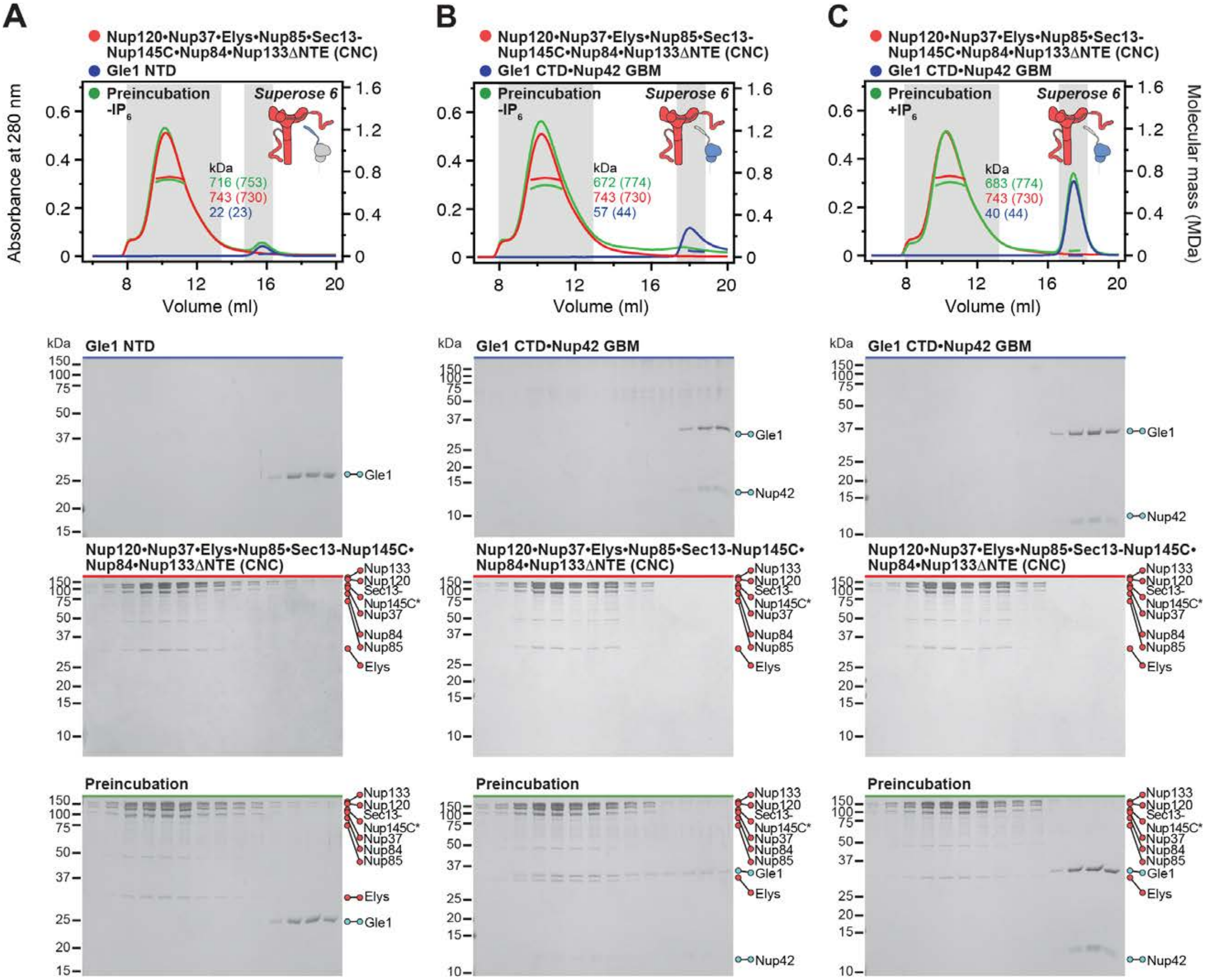
A
Nup120•Nup37•Elys•Nup85•Sec13-Nup145C•Nup84•Nup133ΔNTE (CNC)
Gle1 NTD
Preincubation -IP6
Superose 6
kDa
716 (753)
743 (730)
22 (23)
Absorbance at 280 nm
Volume (ml)
B
Gle1 CTD•Nup42 GBM
672 (774)
743 (730)
57 (44)
C
Preincubation +IP6
683 (774)
40 (44)
Molecular mass (MDa)
Gle1
Nup42
Preincubation
Nup133
Nup120
Sec13-
Nup145C*
Nup37
Nup84
Nup85
Elys

**Fig. S25. Gle1$^{NTD}$ is dispensable for the Gle1•Nup42$^{GBM}$-CNC interaction.** (**A**, **B**) SEC-MALS interaction analyses of the CNC in the absence of $IP_6$ with (A) Gle1$^{NTD}$ and (B) Gle1$^{CTD}$•Nup42$^{GBM}$, showing that Gle1$^{CTD}$•Nup42$^{GBM}$ binds the CNC, whereas no binding is detected for Gle1$^{NTD}$. (**C**) SEC-MALS interaction analysis of the CNC in the presence of $IP_6$ with Gle1$^{CTD}$•Nup42$^{GBM}$, demonstrating that the presence of $IP_6$ prevents binding of Gle1$^{CTD}$•Nup42 $^{GBM}$ to CNC. SEC-MALS profiles of nup complexes are shown individually (red, blue) and after preincubation (green). SEC profiles were obtained using a Superose 6 10/300 GL column. Measured molecular masses are indicated, with the respective theoretical masses shown in parentheses. Inset schematics display tested protein samples colored to match the corresponding chromatogram trace. Gray boxes indicate fractions resolved on SDS-PAGE gels and visualized by Coomassie brilliant blue staining.

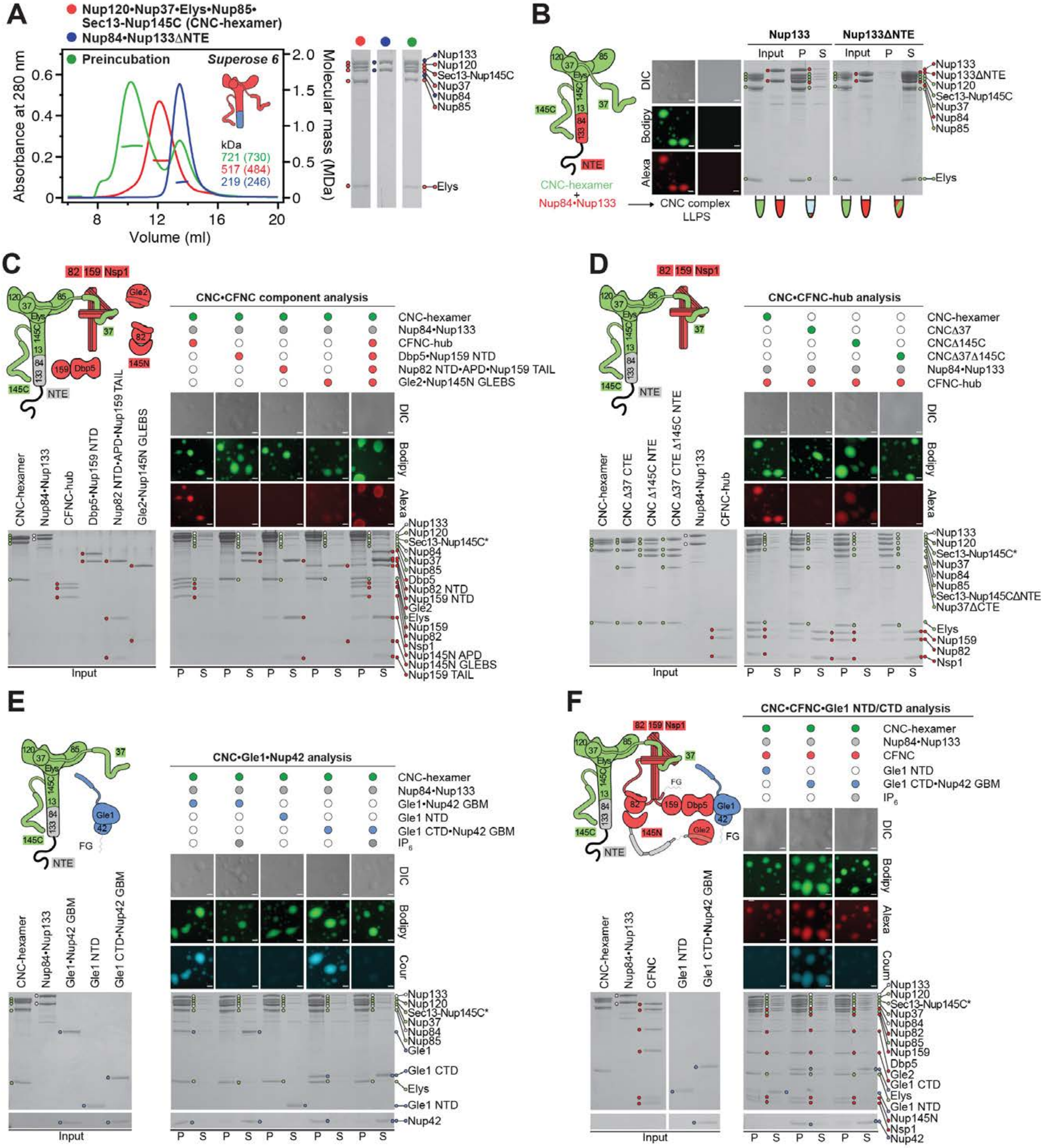

A
Nup120•Nup37•Elys•Nup85•Sec13-Nup145C (CNC-hexamer)
Nup84•Nup133ΔNTE
Preincubation
Superose 6
Absorbance at 280 nm
Molecular mass (MDa)
Volume (ml)
kDa
721 (730)
517 (484)
219 (246)
Nup133
Nup120
Sec13-Nup145C
Nup37
Nup84
Nup85
Elys
B
Nup133
Nup133ΔNTE
Input P S
CNC-hexamer + Nup84•Nup133
CNC complex LLPS
DIC
Bodipy
Alexa
Nup133ΔNTE
C
CNC•CFNC component analysis
CNC-hexamer
Nup84•Nup133
CFNC-hub
Dbp5•Nup159 NTD
Nup82 NTD•APD•Nup159 TAIL
Gle2•Nup145N GLEBS
Input
Sec13-Nup145C*
Dbp5
Nup82 NTD
Nup159 NTD
Gle2
Nup159
Nup82
Nsp1
Nup145N APD
Nup145N GLEBS
Nup159 TAIL
D
CNC•CFNC-hub analysis
CNCΔ37
CNCΔ145C
CNCΔ37Δ145C
CNC Δ37 CTE
CNC Δ145C NTE
CNC Δ37 CTE Δ145C NTE
Sec13-Nup145CΔNTE
Nup37ΔCTE
E
CNC•Gle1•Nup42 analysis
Gle1•Nup42 GBM
Gle1 NTD
Gle1 CTD•Nup42 GBM
IP6
Cour
Gle1
Gle1 CTD
Gle1 NTD
Nup42
F
CNC•CFNC•Gle1 NTD/CTD analysis
CFNC
Nup145N

**Fig. S26. Liquid-liquid phase separation analysis of coat-cytoplasmic filament nup interactions.** (**A**) SEC-MALS interaction analysis of Nup120•Nup37•Elys•Nup85•Sec13-Nup145C (CNC-hexamer; red) with Nup84•Nup133$^{\Delta NTE}$ (blue), and their preincubation (green), showing the formation of a stoichiometric monodisperse CNC complex in the absence of Nup133$^{NTE}$. SEC profiles were obtained using a Superose 6 10/300 GL column. Measured molecular masses are indicated, with the respective theoretical masses shown in parentheses. SDS-PAGE gels strips of the peak fractions are shown. (**B**) Mixing CNC-hexamer (green) with Nup84•Nup133 (red) results in formation of liquid-liquid phase separation (LLPS) condensates, dependent on the presence of Nup133$^{NTE}$. (**C**) LLPS interaction assay, assessing the incorporation of the four CNFC subcomplexes (CFNC-hub, Dbp5•Nup159$^{NTD}$, Nup82$^{NTD}$•Nup145N$^{APD}$•Nup159$^{TAIL}$, and Gle2•Nup145N$^{GLEBS}$) into the CNC-LLPS condensate, identifying that only CFNC-hub enters. (**D**) LLPS interaction assay, assessing CFNC-hub incorporation into LLPS condensates, generated by CNC variants lacking either Nup37$^{CTE}$, Nup145C$^{NTE}$, or both assembly sensors, demonstrating that CFNC-hub entry is dependent on both assembly sensors. (**E**) LLPS interaction assay, assessing the incorporation of Gle1•Nup42$^{GBM}$, Gle1$^{NTD}$, and Gle1$^{CTD}$•Nup42$^{GBM}$, into the CNC-LLPS condensate. Whereas Gle1$^{NTD}$ fails to incorporate, both Gle1•Nup42$^{GBM}$ and Gle1$^{CTD}$•Nup42$^{GBM}$ enter the condensate, but only in the absence of $IP_6$. (**F**) LLPS interaction assay, assessing Gle1$^{NTD}$ and Gle1$^{CTD}$•Nup42$^{GBM}$ incorporation into CNC-CFNC-LLPS condensates. As for the CNC-LLPS, Gle1$^{NTD}$ fails to incorporate, whereas the condensate incorporation of Gle1$^{CTD}$•Nup42$^{GBM}$ occurs only in the absence of $IP_6$. These results establish that the CFNC-CNC interaction is mediated by the Nup37$^{CTE}$ and Nup145C$^{NTE}$ assembly sensors and show that Gle1•Nup42$^{GMB}$ incorporation requires both Gle1$^{NTD}$ and Gle1$^{CTD}$•Nup42$^{GMB}$ in the presence of $IP_6$. The fluorescence microscopy analyses utilized dye labeled proteins, CNC-hexamer and its variants (Bodipy; green), CFNC and its sub-complexes (Alexa Fluor 647; red), and Gle1•Nup42$^{GBM}$ and its fragments (Coumarin; cyan). Pelleted CNC condensate phase (P) and soluble (S) fractions were analyzed by SDS-PAGE and visualized by Coomassie brilliant blue staining. All experiments were repeated at least three times. Scale bars are 10 $\mu$m.

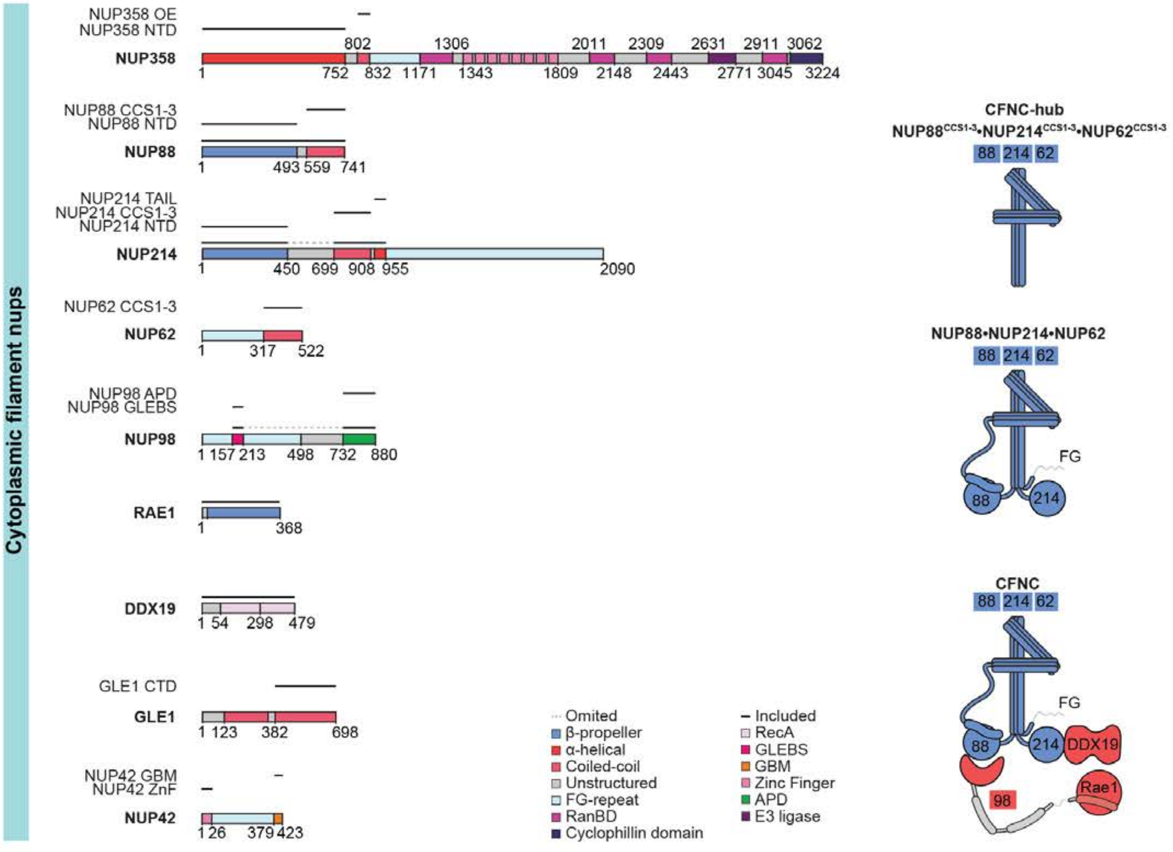
Cytoplasmic filament nups
NUP358 OE
NUP358 NTD
NUP358
1 752 802 832 1171 1306 1343 1809 2011 2148 2309 2443 2631 2771 2911 3045 3062 3224
NUP88 CCS1-3
NUP88 NTD
NUP88
1 493 559 741
NUP214 TAIL
NUP214 CCS1-3
NUP214 NTD
NUP214
1 450 699 908 955 2090
NUP62 CCS1-3
NUP62
1 317 522
NUP98 APD
NUP98 GLEBS
NUP98
1 157 213 498 732 880
RAE1
1 368
DDX19
1 54 298 479
GLE1 CTD
GLE1
1 123 382 698
NUP42 GBM
NUP42 ZnF
NUP42
1 26 379 423
Omited
β-propeller
α-helical
Coiled-coil
Unstructured
FG-repeat
RanBD
Cyclophillin domain
Included
RecA
GLEBS
GBM
Zinc Finger
APD
E3 ligase
CFNC-hub
NUP88CCS1-3•NUP214CCS1-3•NUP62CCS1-3
88 214 62
NUP88•NUP214•NUP62
88 214 62
FG
88
214
CFNC
88 214 62
FG
88
214
DDX19
98
Rae1

**Fig. S27. *H. sapiens* nucleoporin fragments.** Domain architectures of the human cytoplasmic filament nups. Domains are drawn as horizontal boxes with residue numbers indicated and their observed or predicted folds colored according to the legend. The black lines indicate the fragments used throughout the text. Dashed lines indicate regions excluded from expression constructs. As a reference, schematics of nup complexes are shown on the right.

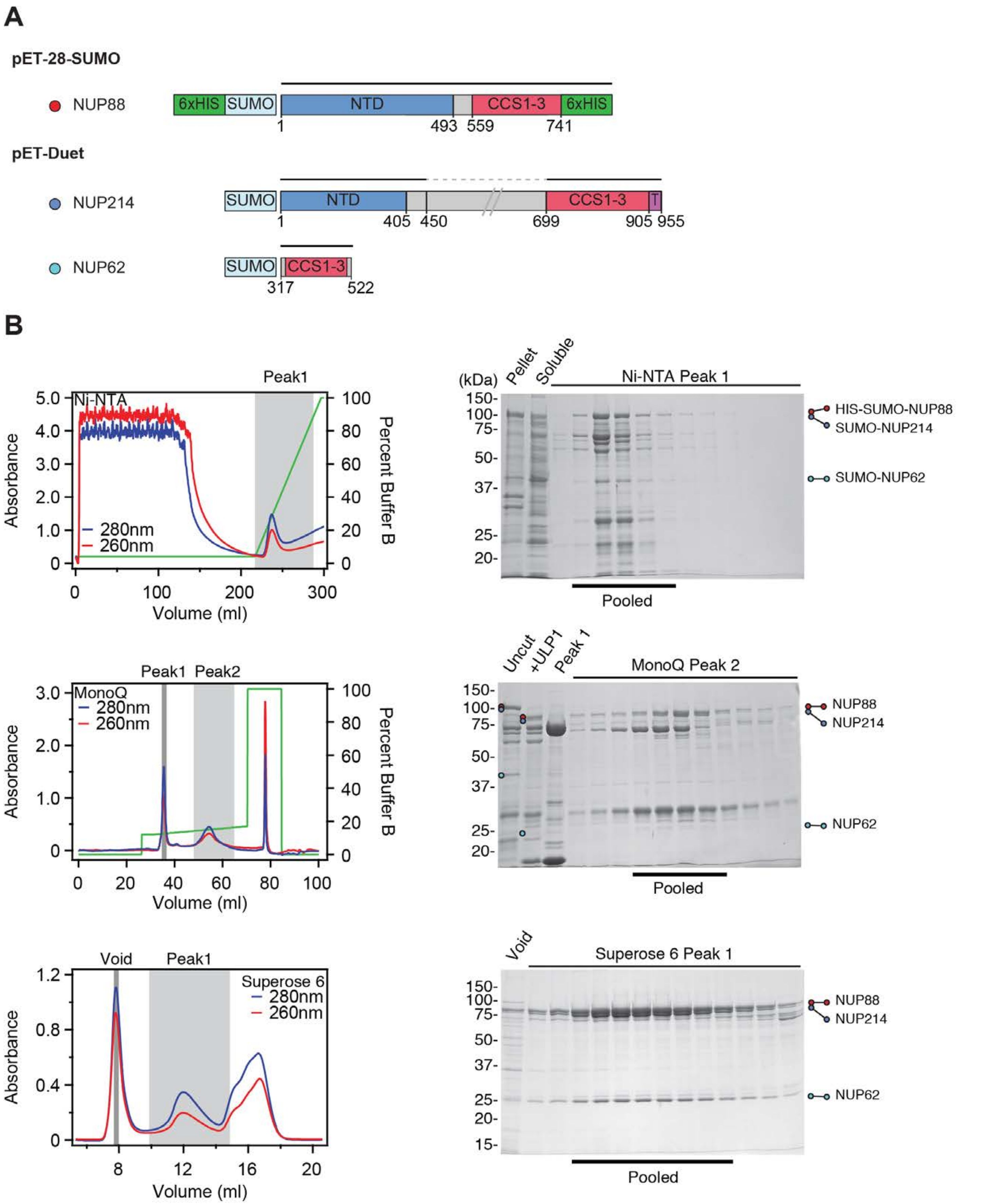
A
pET-28-SUMO
NUP88
6xHIS
SUMO
NTD
CCS1-3
6xHIS
1
493
559
741
pET-Duet
NUP214
SUMO
NTD
CCS1-3
T
1
405
450
699
905
955
NUP62
SUMO
CCS1-3
317
522
B
Peak1
Ni-NTA
Absorbance
Percent Buffer B
280nm
260nm
Volume (ml)
(kDa)
Pellet
Soluble
Ni-NTA Peak 1
HIS-SUMO-NUP88
SUMO-NUP214
SUMO-NUP62
Pooled
Peak1
Peak2
MonoQ
280nm
260nm
Absorbance
Percent Buffer B
Volume (ml)
Uncut
+ULP1
Peak 1
MonoQ Peak 2
NUP88
NUP214
NUP62
Pooled
Void
Peak1
Superose 6
280nm
260nm
Absorbance
Volume (ml)
Void
Superose 6 Peak 1
NUP88
NUP214
NUP62
Pooled

**Fig. S28. Purification of the human NUP88•NUP214•NUP62 hetero-trimer.** (**A**) Domain boundaries of the purified nups are shown with black lines indicating the construct boundaries of NUP88, NUP214, and NUP62 in our bacterial expression constructs. The dashed line indicates a central unstructured region in NUP214 that was excluded. (**B**) NUP88•NUP214•NUP62 hetero-trimer purification. Sequential chromatography purification steps are shown from top to bottom with the employed columns indicated. The gray boxes indicate fractions that were resolved on SDS-PAGE gels and visualized by Coomassie staining. Pooled fractions are indicated with a black bar below the SDS-PAGE gels. For details of the buffer conditions see Table S4.

**A**

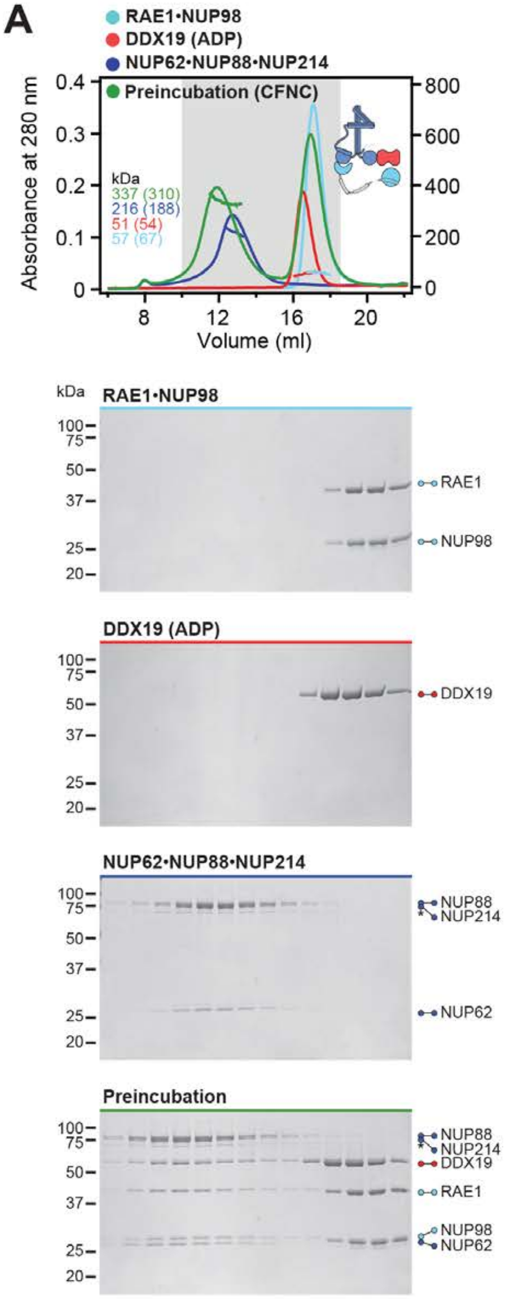
RAE1•NUP98
DDX19 (ADP)
NUP62•NUP88•NUP214
Preincubation (CFNC)
Absorbance at 280 nm
Volume (ml)
kDa
337 (310)
216 (188)
51 (54)
57 (67)
RAE1•NUP98
RAE1
NUP98
DDX19 (ADP)
DDX19
NUP62•NUP88•NUP214
NUP88
NUP214
NUP62
Preincubation
NUP88
NUP214
DDX19
RAE1
NUP98
NUP62

**B**

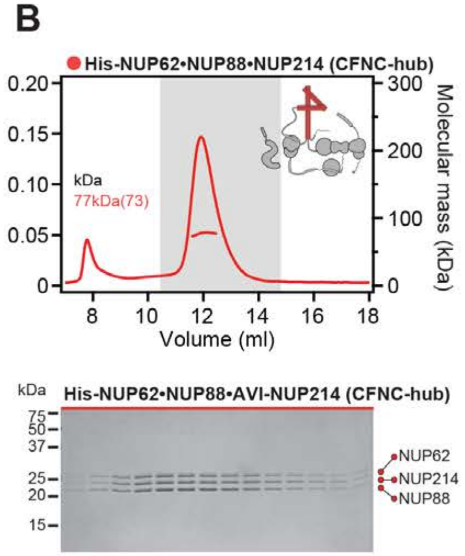
His-NUP62•NUP88•NUP214 (CFNC-hub)
Molecular mass (kDa)
Volume (ml)
kDa
77kDa(73)
His-NUP62•NUP88•AVI-NUP214 (CFNC-hub)
NUP62
NUP214
NUP88

**Fig. S29. Reconstitution of the evolutionarily conserved human CFNC hetero-hexamer.** (**A**) SEC-MALS interaction analysis showing the biochemical reconstitution of the ~330 kDa hetero-hexameric CFNC (green) from NUP62•NUP88•NUP214 (blue), RAE1•NUP98 (cyan), and DDX19(ADP) (red) (corresponding to Fig. 2C). SEC profiles were obtained on a Superose 6 10/300 GL column. (**B**) SEC-MALS analysis of the the hetero-trimeric NUP62•NUP88•NUP214 CCS1-3 complex (CFNC-hub). SEC profile was obtained on a Superdex 200 10/300 GL column. Measured molecular masses are indicated, with the respective theoretical masses shown in parentheses. The inset schematics are colored to indicate the CFNC sub-complexes of each SEC-MALS interaction experiment, with gray domains and regions omitted from the experiment. Gray boxes indicate fractions resolved on SDS-PAGE gels and visualized by Coomassie brilliant blue staining.

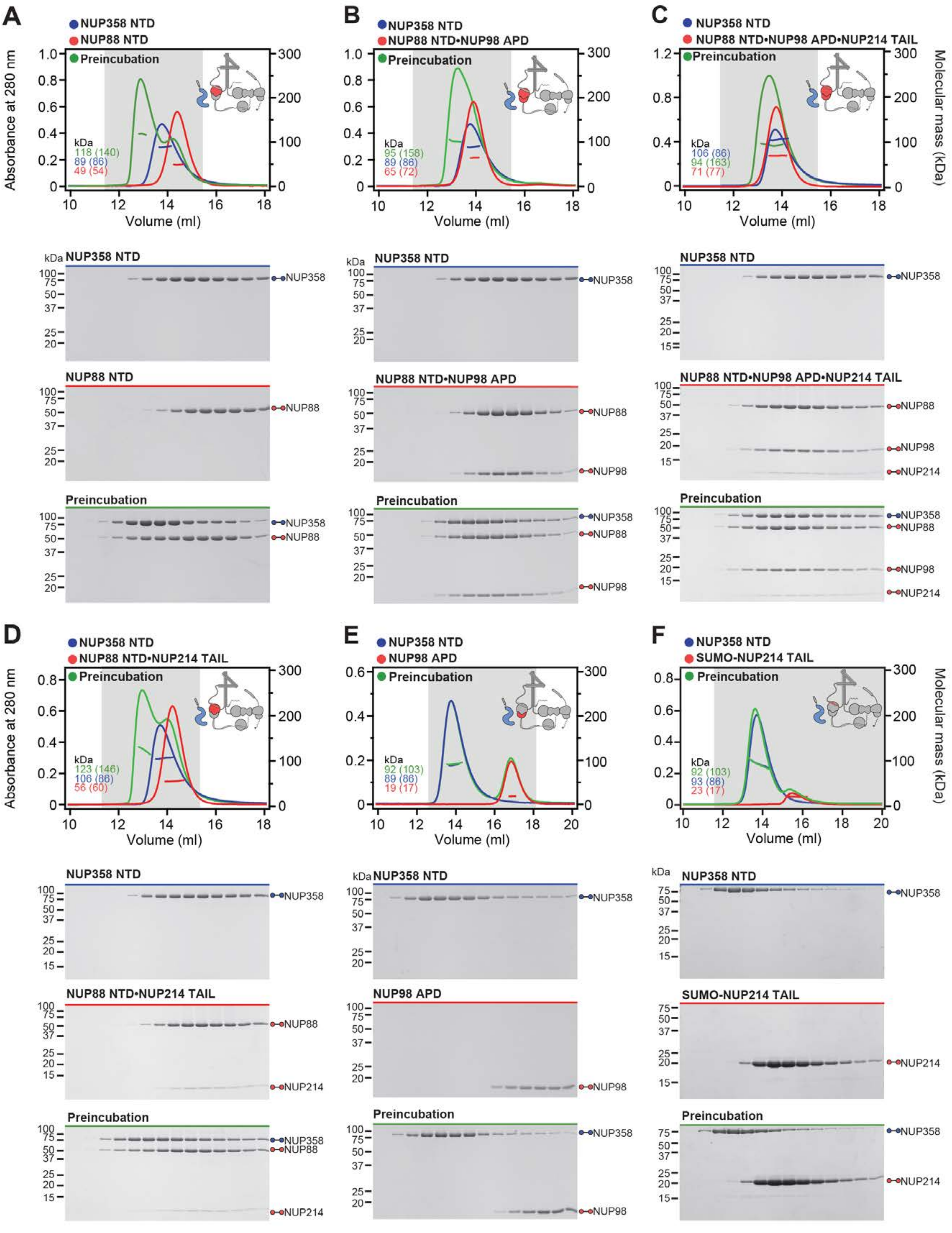
A
NUP358 NTD
NUP88 NTD
Preincubation
Absorbance at 280 nm
Volume (ml)
kDa
118 (140)
89 (86)
49 (54)
B
NUP358 NTD
NUP88 NTD•NUP98 APD
Preincubation
95 (158)
89 (86)
65 (72)
C
NUP358 NTD
NUP88 NTD•NUP98 APD•NUP214 TAIL
Preincubation
106 (86)
94 (163)
71 (77)
Molecular mass (kDa)
NUP358 NTD
NUP88 NTD
NUP88 NTD•NUP98 APD
NUP88 NTD•NUP98 APD•NUP214 TAIL
Preincubation
NUP358
NUP88
NUP98
NUP214
D
NUP358 NTD
NUP88 NTD•NUP214 TAIL
Preincubation
123 (146)
106 (86)
56 (60)
E
NUP358 NTD
NUP98 APD
Preincubation
92 (103)
89 (86)
19 (17)
F
NUP358 NTD
SUMO-NUP214 TAIL
Preincubation
92 (103)
93 (86)
23 (17)
NUP88 NTD•NUP214 TAIL
NUP98 APD
SUMO-NUP214 TAIL

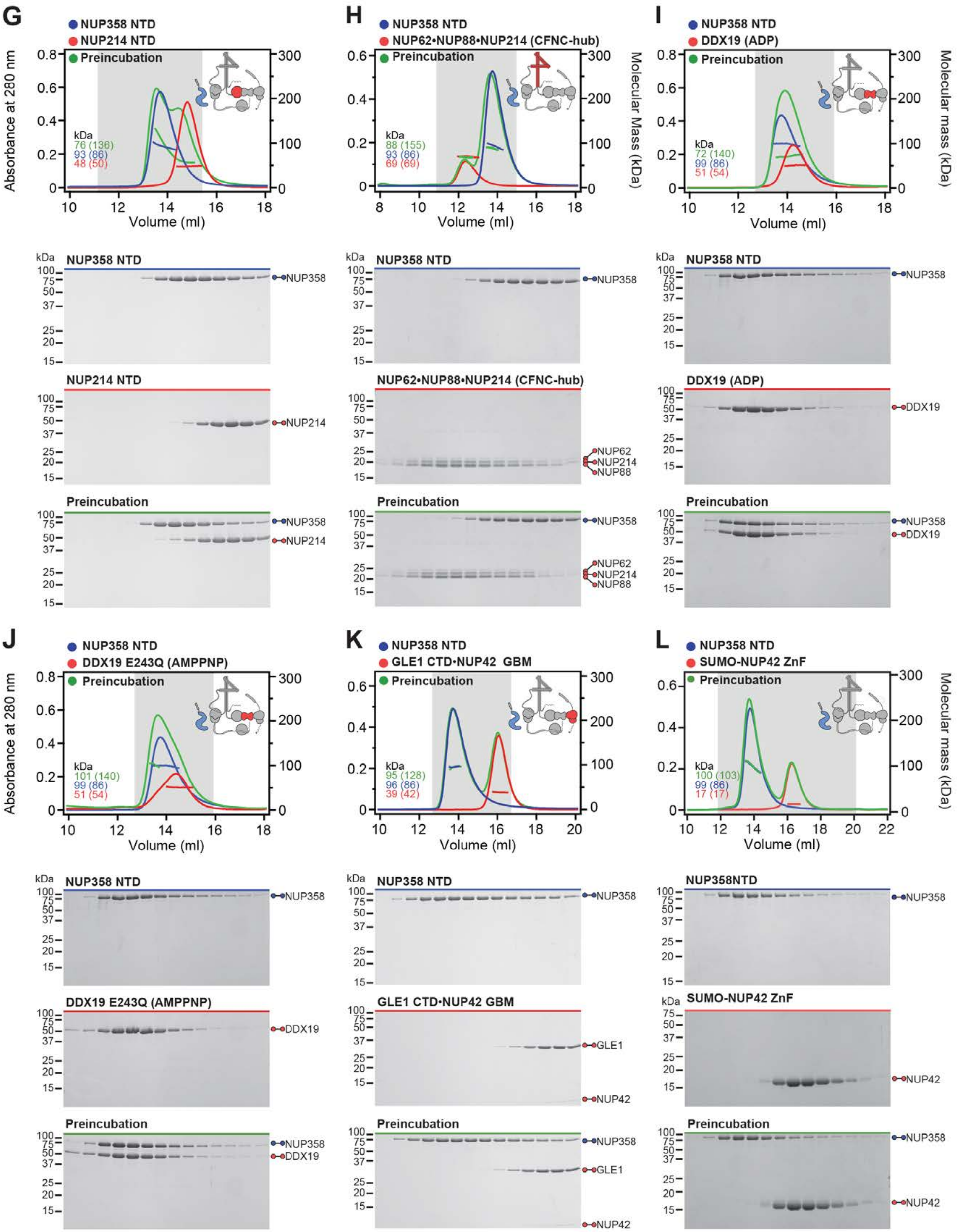
G
NUP358 NTD
NUP214 NTD
Preincubation
Absorbance at 280 nm
Volume (ml)
kDa
76 (136)
93 (86)
48 (50)
NUP358 NTD
NUP358
NUP214 NTD
NUP214
Preincubation
H
NUP358 NTD
NUP62•NUP88•NUP214 (CFNC-hub)
Preincubation
Molecular Mass (kDa)
88 (155)
93 (86)
69 (69)
NUP62•NUP88•NUP214 (CFNC-hub)
NUP62
NUP214
NUP88
I
NUP358 NTD
DDX19 (ADP)
Preincubation
Molecular mass (kDa)
72 (140)
99 (86)
51 (54)
DDX19
J
NUP358 NTD
DDX19 E243Q (AMPPNP)
Preincubation
101 (140)
99 (86)
51 (54)
K
NUP358 NTD
GLE1 CTD•NUP42 GBM
Preincubation
95 (128)
96 (86)
39 (42)
GLE1
NUP42
L
NUP358 NTD
SUMO-NUP42 ZnF
Preincubation
100 (103)
99 (86)
17 (17)
NUP358NTD

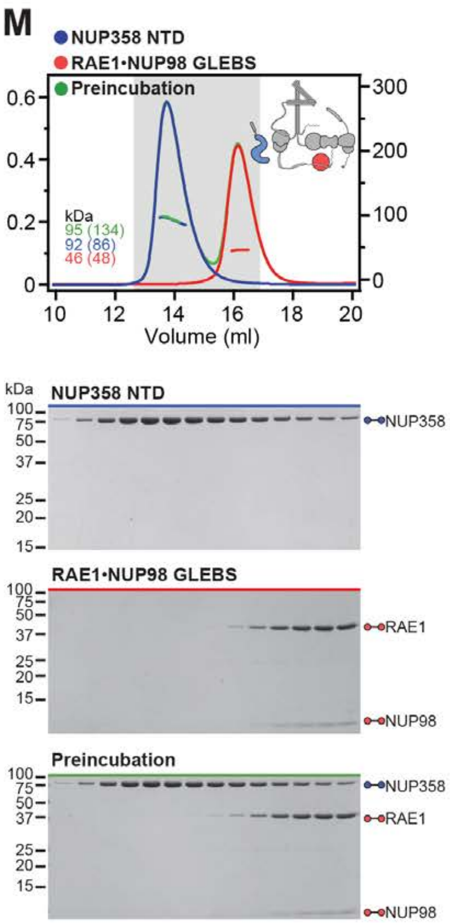
M
NUP358 NTD
RAE1•NUP98 GLEBS
Preincubation
kDa
95 (134)
92 (86)
46 (48)
Volume (ml)
NUP358 NTD
RAE1•NUP98 GLEBS
Preincubation
NUP358
RAE1
NUP98

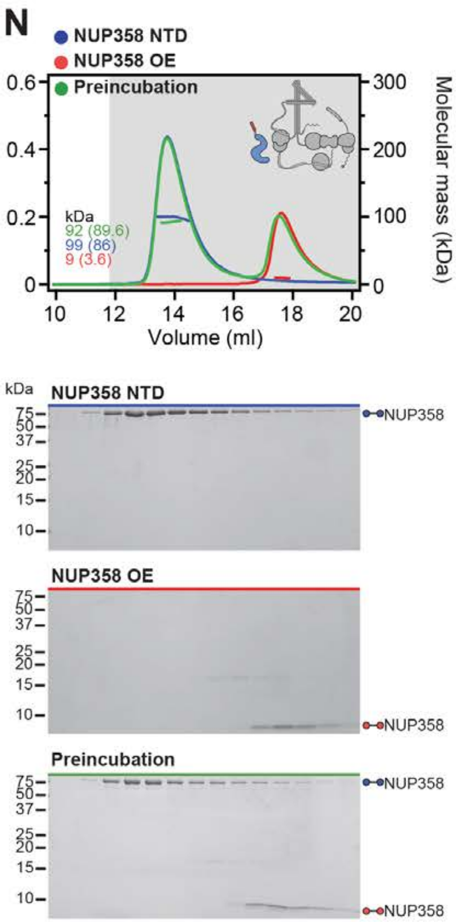
N
NUP358 NTD
NUP358 OE
Preincubation
kDa
92 (89.6)
99 (86)
9 (3.6)
Volume (ml)
Molecular mass (kDa)
NUP358 NTD
NUP358 OE
Preincubation
NUP358

**Fig. S30. Systematic interaction analyses of NUP358$^{NTD}$ with human CF nup fragments.** SEC-MALS interaction analyses of NUP358$^{NTD}$ with (**A**) Nup88$^{NTD}$, (**B**) NUP88$^{NTD}$•NUP98$^{APD}$, (**C**) NUP88$^{NTD}$•NUP98$^{APD}$•NUP214$^{TAIL}$, (**D**) NUP88$^{NTD}$•NUP214$^{TAIL}$, (**E)** NUP98$^{APD}$, (**F**) SUMO-NUP214$^{TAIL}$, (**G**) NUP214$^{NTD}$, (**H**) CFNC-hub, (**I**) DDX19 (ADP), (**J**) DDX19 E243Q (AMPPNP), (**K**) GLE1$^{CTD}$•NUP42, (**L**) SUMO-NUP42$^{ZnF}$, (**M**) RAE1•NUP98$^{GLEBS}$, and (**N**) NUP358$^{OE}$. Interactions were observed between Nup358$^{NTD}$ and Nup88$^{NTD}$, Nup88$^{NTD}$-Nup98$^{APD}$, and Nup88$^{NTD}$-Nup98$^{APD}$-Nup214$^{TAIL}$. SEC-MALS profiles of nup complexes are shown individually (red and blue) and after preincubation (green). SEC profiles were obtained using a Superdex 200 10/300 GL column. Measured molecular masses are indicated, with the respective theoretical masses shown in parentheses. Inset schematics display tested protein samples colored to match the corresponding chromatogram trace. Gray boxes indicate fractions resolved on SDS-PAGE gels and visualized by Coomassie brilliant blue staining.

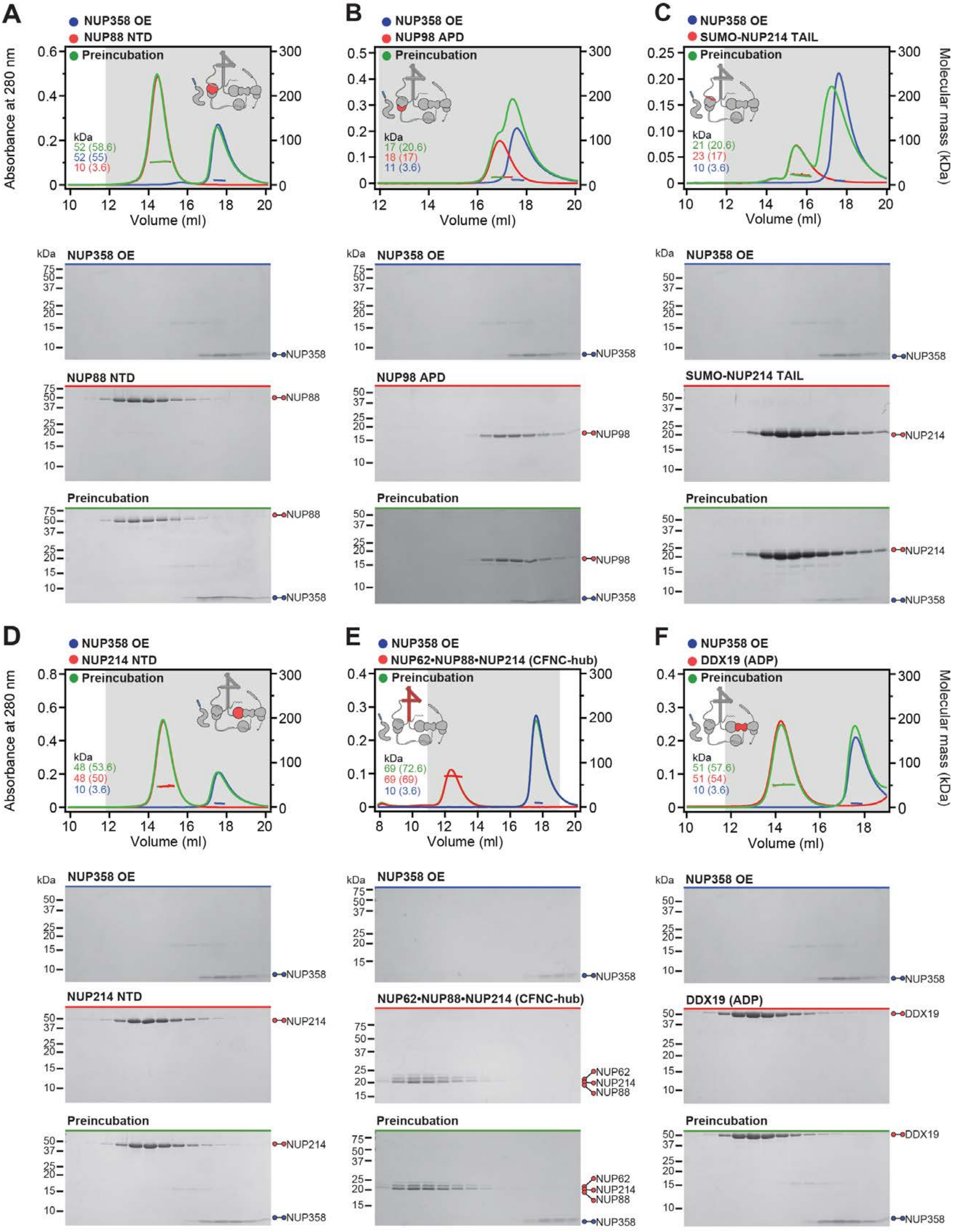
A
NUP358 OE
NUP88 NTD
Preincubation
Absorbance at 280 nm
kDa
52 (58.6)
52 (55)
10 (3.6)
Volume (ml)
NUP358 OE
NUP358
NUP88 NTD
NUP88
Preincubation
NUP88
NUP358
B
NUP358 OE
NUP98 APD
Preincubation
kDa
17 (20.6)
18 (17)
11 (3.6)
Volume (ml)
NUP358 OE
NUP358
NUP98 APD
NUP98
Preincubation
NUP98
NUP358
C
NUP358 OE
SUMO-NUP214 TAIL
Preincubation
Molecular mass (kDa)
kDa
21 (20.6)
23 (17)
10 (3.6)
Volume (ml)
NUP358 OE
NUP358
SUMO-NUP214 TAIL
NUP214
Preincubation
NUP214
NUP358
D
NUP358 OE
NUP214 NTD
Preincubation
Absorbance at 280 nm
kDa
48 (53.6)
48 (50)
10 (3.6)
Volume (ml)
NUP358 OE
NUP358
NUP214 NTD
NUP214
Preincubation
NUP214
NUP358
E
NUP358 OE
NUP62•NUP88•NUP214 (CFNC-hub)
Preincubation
kDa
69 (72.6)
69 (69)
10 (3.6)
Volume (ml)
NUP358 OE
NUP358
NUP62•NUP88•NUP214 (CFNC-hub)
NUP62
NUP214
NUP88
Preincubation
NUP62
NUP214
NUP88
NUP358
F
NUP358 OE
DDX19 (ADP)
Preincubation
Molecular mass (kDa)
kDa
51 (57.6)
51 (54)
10 (3.6)
Volume (ml)
NUP358 OE
NUP358
DDX19 (ADP)
DDX19
Preincubation
DDX19
NUP358

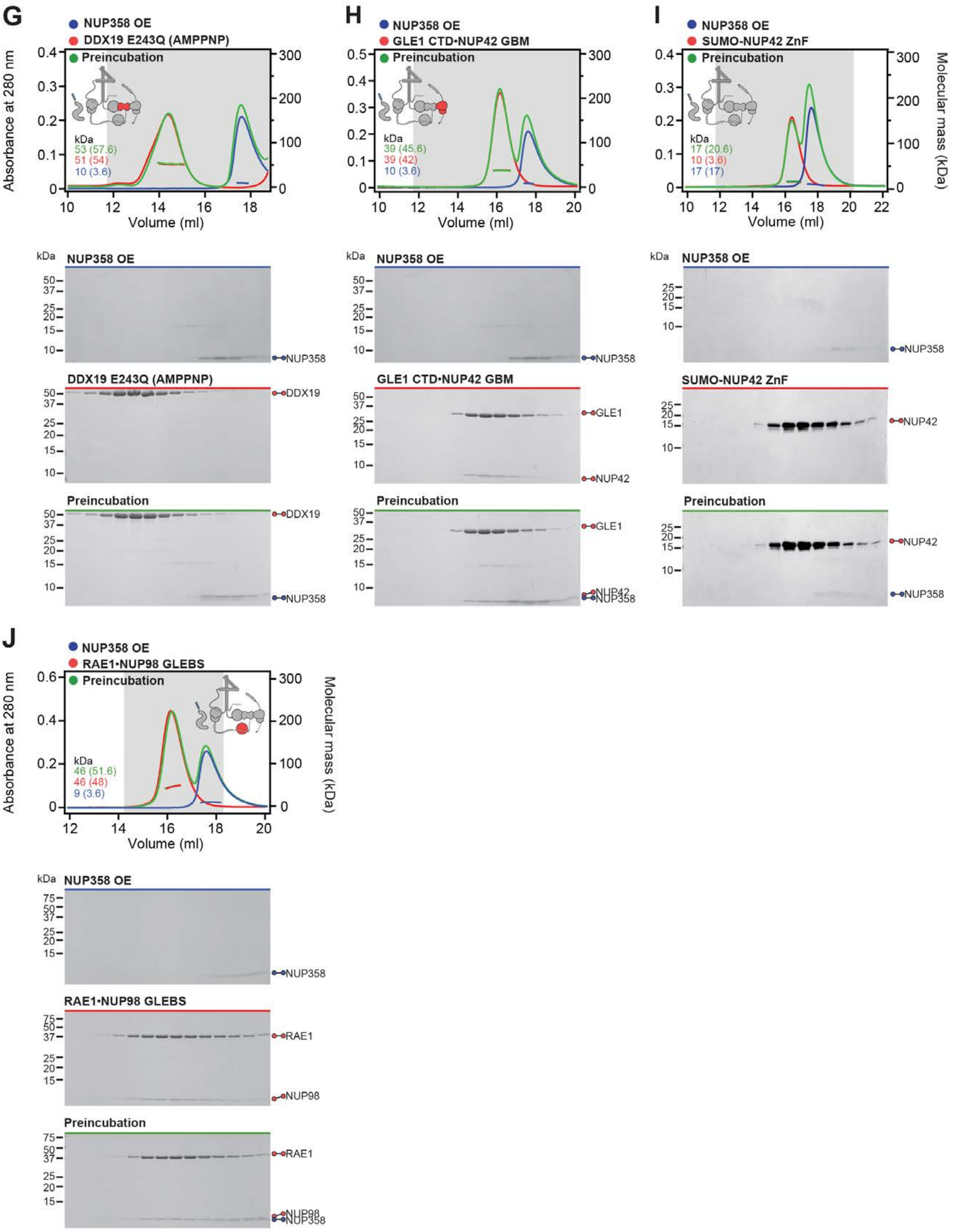
G
NUP358 OE
DDX19 E243Q (AMPPNP)
Preincubation
Absorbance at 280 nm
Volume (ml)
kDa
53 (57.6)
51 (54)
10 (3.6)
NUP358 OE
NUP358
DDX19 E243Q (AMPPNP)
DDX19
Preincubation
DDX19
NUP358
H
NUP358 OE
GLE1 CTD•NUP42 GBM
Preincubation
Volume (ml)
kDa
39 (45.6)
39 (42)
10 (3.6)
NUP358 OE
NUP358
GLE1 CTD•NUP42 GBM
GLE1
NUP42
Preincubation
GLE1
NUP42
NUP358
I
NUP358 OE
SUMO-NUP42 ZnF
Preincubation
Molecular mass (kDa)
Volume (ml)
kDa
17 (20.6)
10 (3.6)
17 (17)
NUP358 OE
NUP358
SUMO-NUP42 ZnF
NUP42
Preincubation
NUP42
NUP358
J
NUP358 OE
RAE1•NUP98 GLEBS
Preincubation
Absorbance at 280 nm
Molecular mass (kDa)
Volume (ml)
kDa
46 (51.6)
46 (48)
9 (3.6)
NUP358 OE
NUP358
RAE1•NUP98 GLEBS
RAE1
NUP98
Preincubation
RAE1
NUP98
NUP358

**Fig. S31. Systematic interaction analyses of NUP358$^{OE}$ with human CF nup fragments.** SEC-MALS interaction analyses of NUP358$^{OE}$ with (**A**) Nup88$^{NTD}$, (**B**) NUP98$^{APD}$, (**C**) SUMO-NUP214$^{TAIL}$, (**D**) NUP214$^{NTD}$, (**E**) CFNC-hub, (**F**) DDX19(ADP), (**G**) DDX19(AMPPNP) E243Q, (**H**) GLE1$^{CTD}$•NUP42, (**I**) SUMO-NUP42$^{ZnF}$, and (**J**) RAE1•NUP98$^{GLEBS}$. SEC-MALS profiles of nup complexes are shown individually (red and blue) and after preincubation (green). SEC profiles were obtained using a Superdex 200 10/300 GL column. Measured molecular masses are indicated, with the respective theoretical masses shown in parentheses. Inset schematics display tested protein samples colored to match the corresponding chromatogram trace. Gray boxes indicate fractions resolved on SDS-PAGE gels and visualized by Coomassie brilliant blue staining.

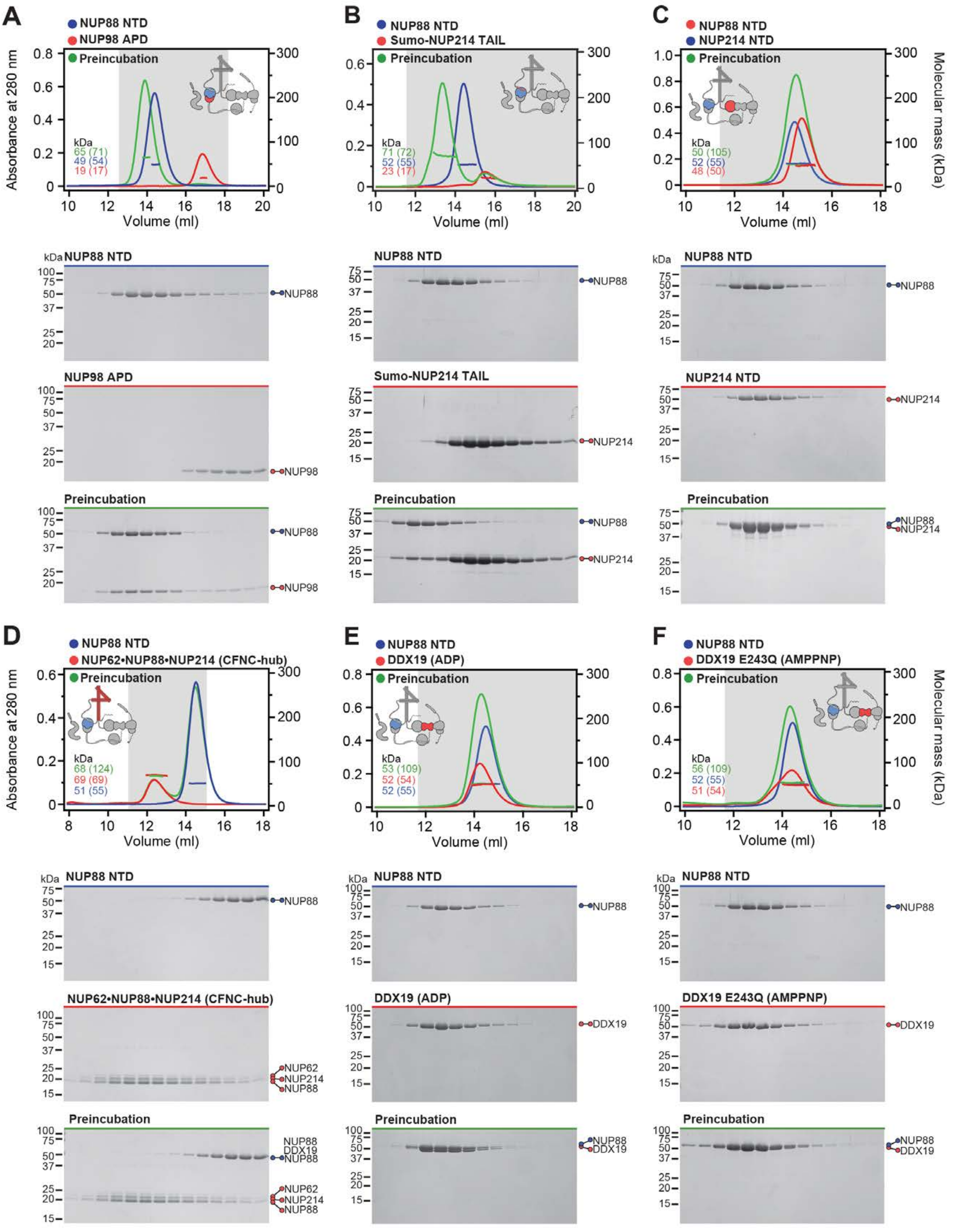

A
NUP88 NTD
NUP98 APD
Preincubation
Absorbance at 280 nm
Volume (ml)
kDa
65 (71)
49 (54)
19 (17)
NUP88
NUP98
B
NUP88 NTD
Sumo-NUP214 TAIL
Preincubation
71 (72)
52 (55)
23 (17)
NUP214
C
NUP88 NTD
NUP214 NTD
Preincubation
Molecular mass (kDa)
50 (105)
52 (55)
48 (50)
D
NUP88 NTD
NUP62•NUP88•NUP214 (CFNC-hub)
Preincubation
68 (124)
69 (69)
51 (55)
NUP62
E
NUP88 NTD
DDX19 (ADP)
Preincubation
53 (109)
52 (54)
52 (55)
DDX19
F
NUP88 NTD
DDX19 E243Q (AMPPNP)
Preincubation
56 (109)
52 (55)
51 (54)

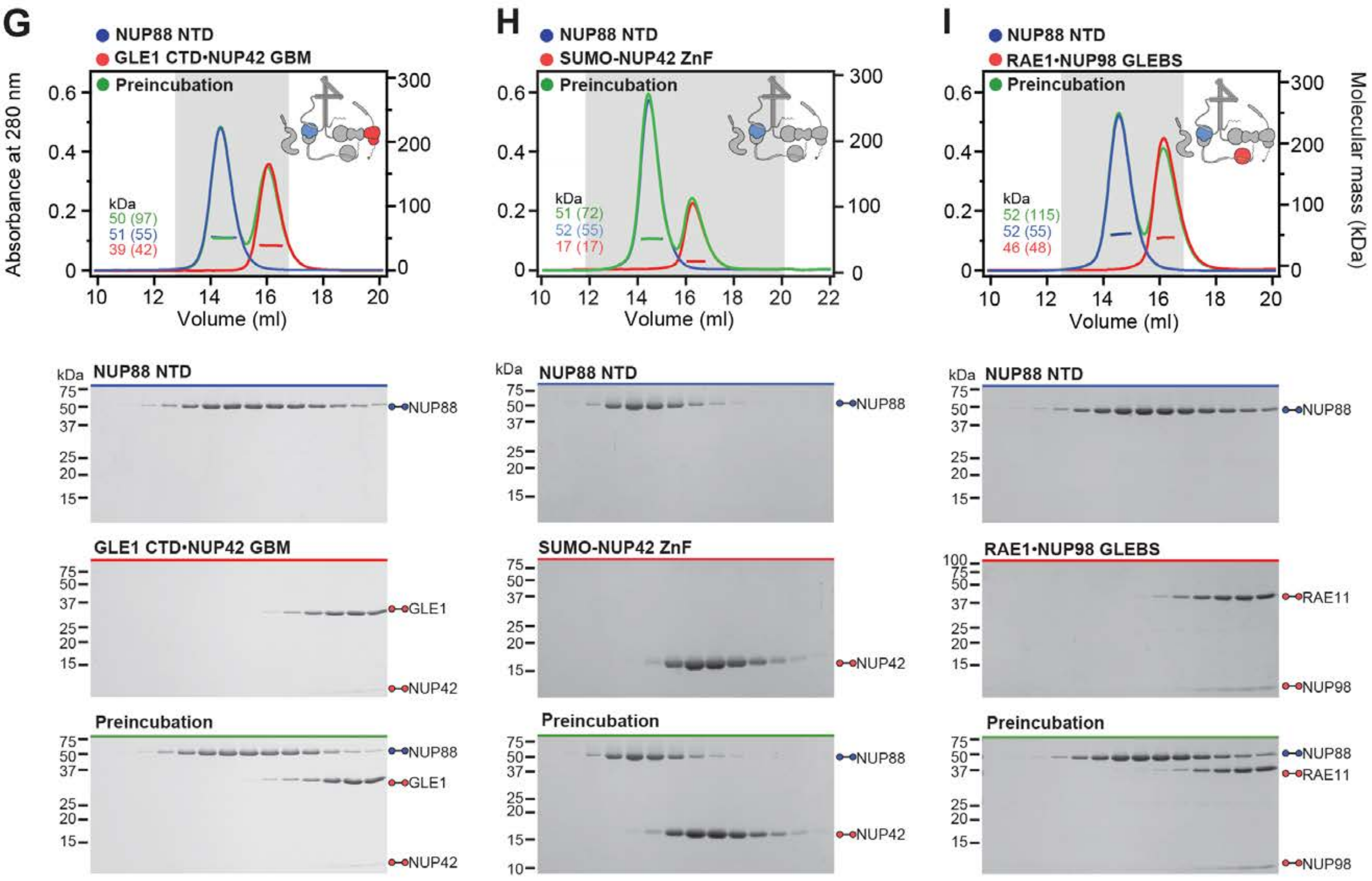

G
NUP88 NTD
GLE1 CTD•NUP42 GBM
Preincubation
Absorbance at 280 nm
kDa
50 (97)
51 (55)
39 (42)
Volume (ml)
NUP88 NTD
GLE1 CTD•NUP42 GBM
Preincubation
NUP88
GLE1
NUP42
H
NUP88 NTD
SUMO-NUP42 ZnF
Preincubation
kDa
51 (72)
52 (55)
17 (17)
Volume (ml)
NUP88 NTD
SUMO-NUP42 ZnF
Preincubation
NUP88
NUP42
I
NUP88 NTD
RAE1•NUP98 GLEBS
Preincubation
Molecular mass (kDa)
kDa
52 (115)
52 (55)
46 (48)
Volume (ml)
NUP88 NTD
RAE1•NUP98 GLEBS
Preincubation
NUP88
RAE11
NUP98

**Fig. S32 Systematic interaction analyses of NUP88$^{NTD}$ with human CF nup fragments.** SEC-MALS interaction analyses of NUP88$^{NTD}$ with (**A**) NUP98$^{APD}$, (**B**) SUMO-NUP214$^{TAIL}$, (**C**) NUP214$^{NTD}$, (**D**) CFNC-hub, (**E**) DDX19(ADP), (**F**) DDX19(AMPPNP) E243Q, (**G**) GLE1$^{CTD}$•NUP42, (**H**) SUMO-NUP42$^{ZnF}$, and (**I**) RAE1•NUP98$^{GLEBS}$. SEC-MALS profiles of nup complexes are shown individually (red and blue) and after preincubation (green). SEC profiles were obtained using a Superdex 200 10/300 GL column. Measured molecular masses are indicated, with the respective theoretical masses shown in parentheses. Inset schematics display tested protein samples colored to match the corresponding chromatogram trace. Gray boxes indicate fractions resolved on SDS-PAGE gels and visualized by Coomassie brilliant blue staining.

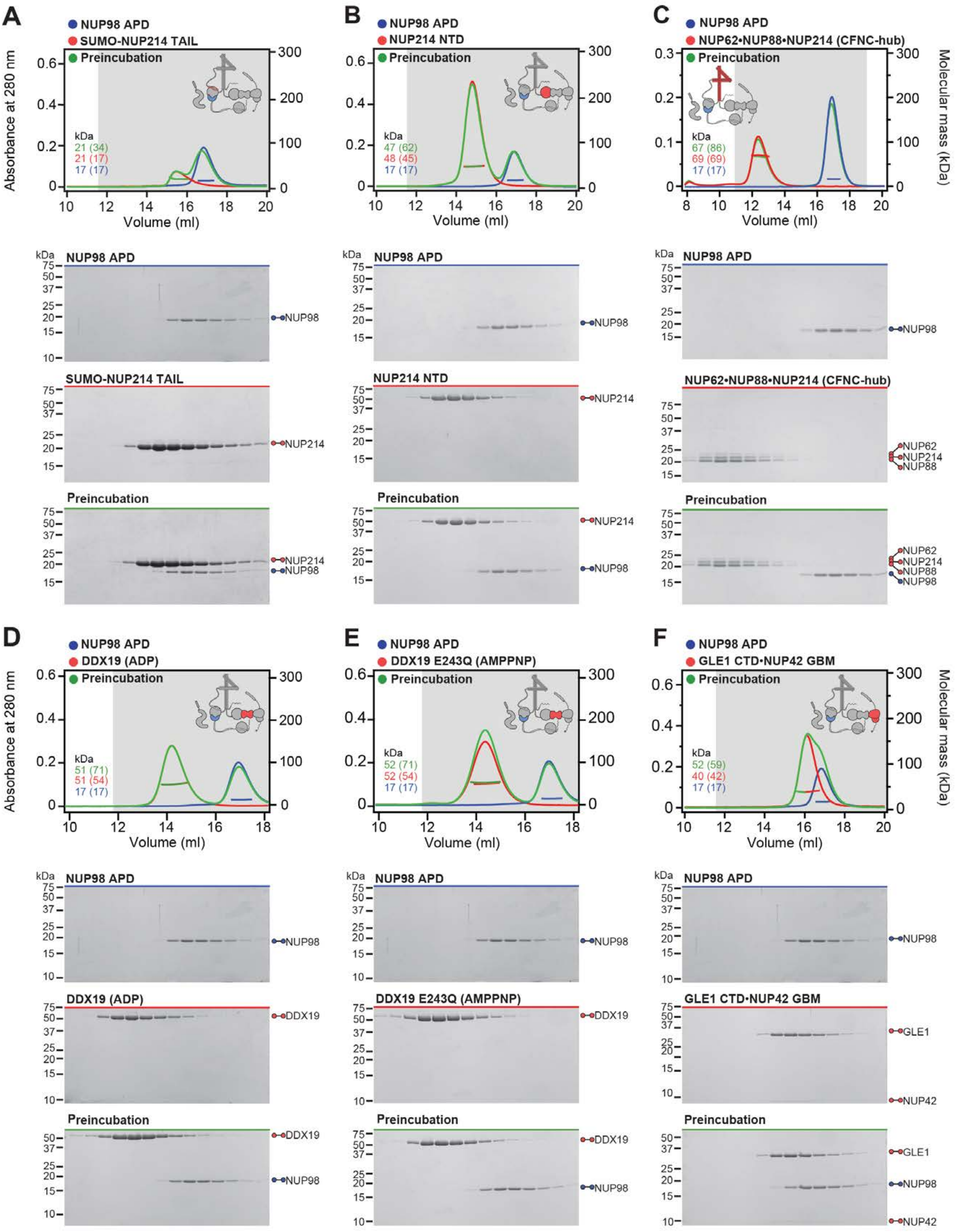

A
NUP98 APD
SUMO-NUP214 TAIL
Preincubation
kDa
21 (34)
21 (17)
17 (17)
Absorbance at 280 nm
Volume (ml)
NUP98 APD
NUP98
SUMO-NUP214 TAIL
NUP214
Preincubation
NUP214
NUP98
B
NUP98 APD
NUP214 NTD
Preincubation
kDa
47 (62)
48 (45)
17 (17)
Volume (ml)
NUP98 APD
NUP98
NUP214 NTD
NUP214
Preincubation
NUP214
NUP98
C
NUP98 APD
NUP62•NUP88•NUP214 (CFNC-hub)
Preincubation
kDa
67 (86)
69 (69)
17 (17)
Volume (ml)
Molecular mass (kDa)
NUP98 APD
NUP98
NUP62•NUP88•NUP214 (CFNC-hub)
NUP62
NUP214
NUP88
Preincubation
NUP62
NUP214
NUP88
NUP98
D
NUP98 APD
DDX19 (ADP)
Preincubation
kDa
51 (71)
51 (54)
17 (17)
Absorbance at 280 nm
Volume (ml)
NUP98 APD
NUP98
DDX19 (ADP)
DDX19
Preincubation
DDX19
NUP98
E
NUP98 APD
DDX19 E243Q (AMPPNP)
Preincubation
kDa
52 (71)
52 (54)
17 (17)
Volume (ml)
NUP98 APD
NUP98
DDX19 E243Q (AMPPNP)
DDX19
Preincubation
DDX19
NUP98
F
NUP98 APD
GLE1 CTD•NUP42 GBM
Preincubation
kDa
52 (59)
40 (42)
17 (17)
Volume (ml)
Molecular mass (kDa)
NUP98 APD
NUP98
GLE1 CTD•NUP42 GBM
GLE1
NUP42
Preincubation
GLE1
NUP98
NUP42

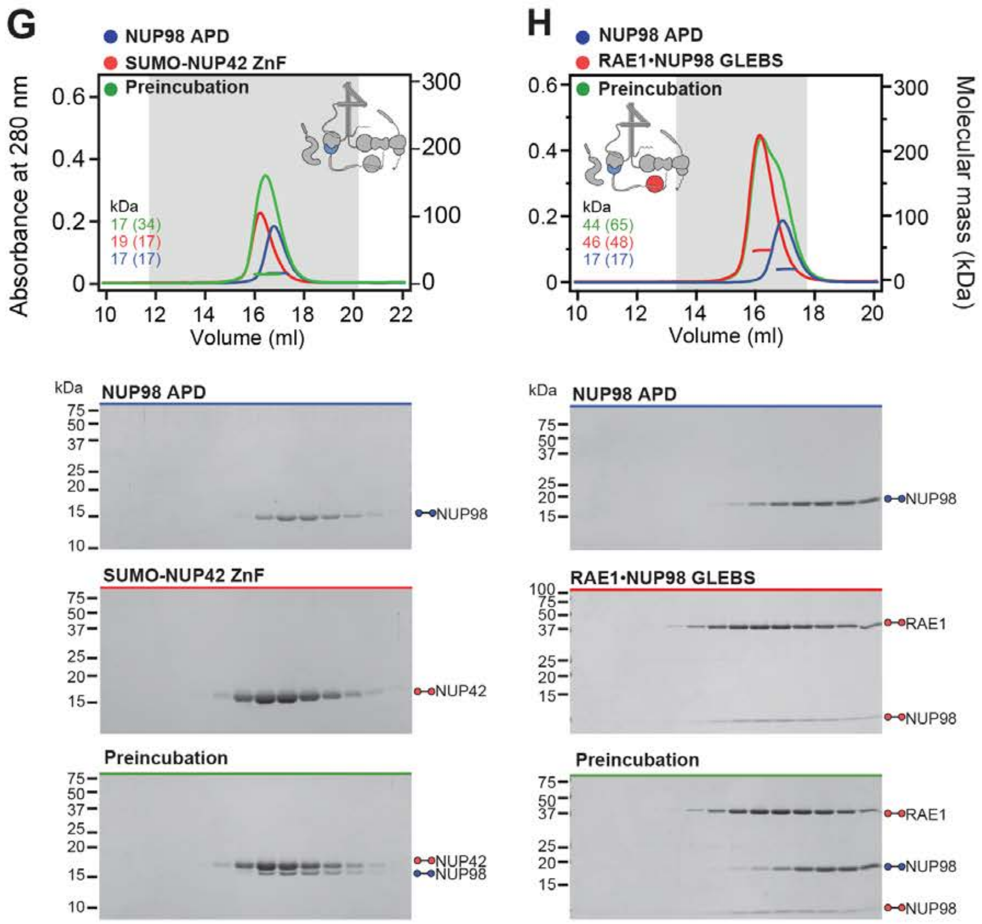

G
NUP98 APD
SUMO-NUP42 ZnF
Preincubation
Absorbance at 280 nm
Volume (ml)
kDa
17 (34)
19 (17)
17 (17)
NUP98 APD
SUMO-NUP42 ZnF
Preincubation
NUP98
NUP42
NUP42
NUP98
H
NUP98 APD
RAE1•NUP98 GLEBS
Preincubation
Molecular mass (kDa)
Volume (ml)
kDa
44 (65)
46 (48)
17 (17)
NUP98 APD
RAE1•NUP98 GLEBS
Preincubation
NUP98
RAE1
NUP98
RAE1
NUP98
NUP98

**Fig. S33 Systematic interaction analyses of NUP98$^{APD}$ with human CF nup fragments.** SEC-MALS interaction analyses of NUP98$^{APD}$ with (**A**) SUMO-NUP214$^{TAIL}$, (**B**) NUP214$^{NTD}$, (**C**) CFNC-hub, (**D**) DDX19(ADP), (**E**) DDX19(AMPPNP) E243Q, (**F**) GLE1$^{CTD}$•NUP42, (**G**) SUMO-NUP42$^{ZnF}$, and (**H**) RAE1•NUP98$^{GLEBS}$. SEC-MALS profiles of nup complexes are shown individually (red and blue) and after preincubation (green). SEC profiles were obtained using a Superdex 200 10/300 GL column. Measured molecular masses are indicated, with the respective theoretical masses shown in parentheses. Inset schematics display tested protein samples colored to match the corresponding chromatogram trace. Gray boxes indicate fractions resolved on SDS-PAGE gels and visualized by Coomassie brilliant blue staining.

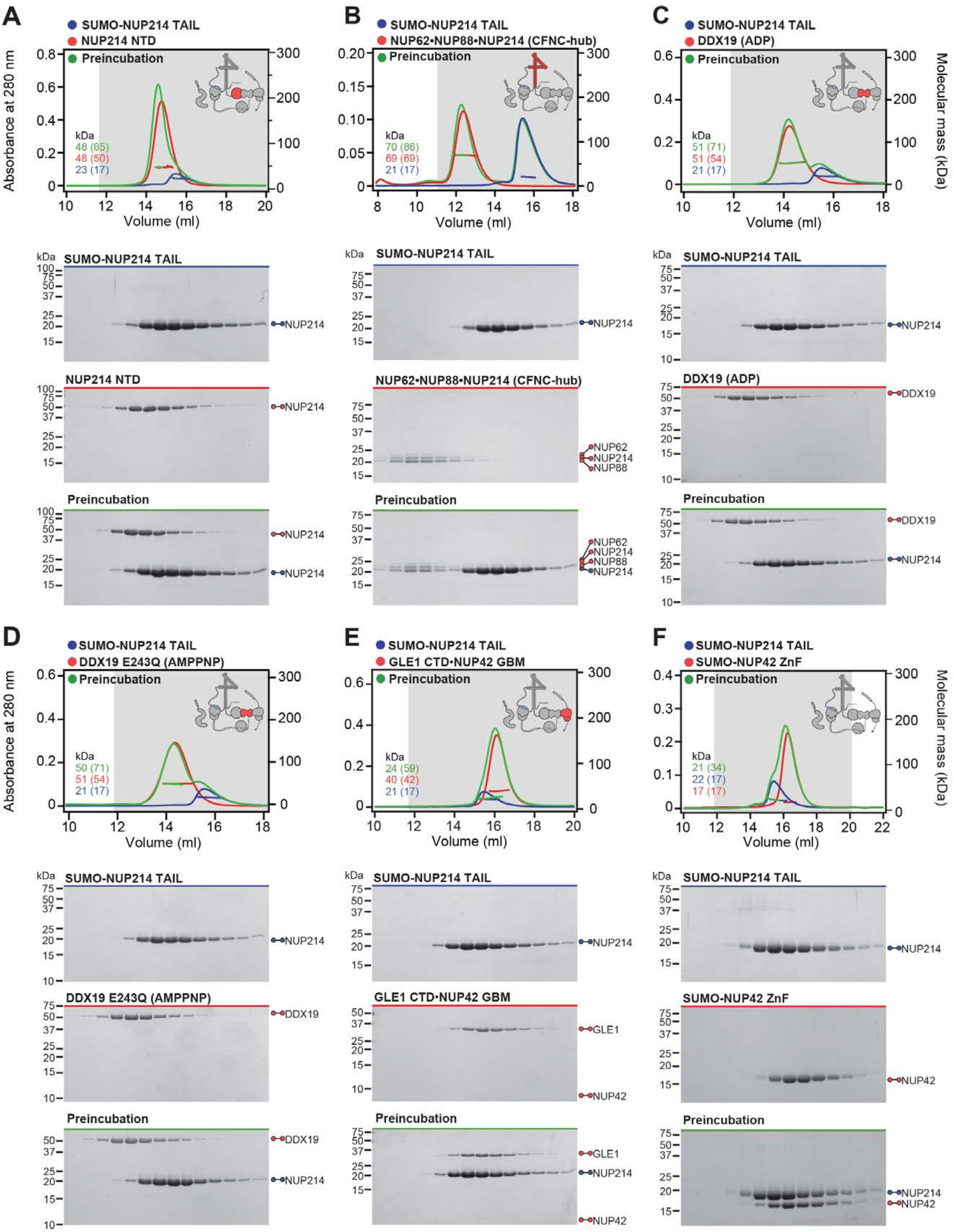
A
SUMO-NUP214 TAIL
NUP214 NTD
Preincubation
Absorbance at 280 nm
kDa
48 (65)
48 (50)
23 (17)
Volume (ml)
B
SUMO-NUP214 TAIL
NUP62•NUP88•NUP214 (CFNC-hub)
Preincubation
70 (86)
69 (69)
21 (17)
C
SUMO-NUP214 TAIL
DDX19 (ADP)
Preincubation
51 (71)
51 (54)
21 (17)
Molecular mass (kDa)
NUP214
NUP62
NUP88
DDX19
D
SUMO-NUP214 TAIL
DDX19 E243Q (AMPPNP)
Preincubation
50 (71)
51 (54)
21 (17)
E
SUMO-NUP214 TAIL
GLE1 CTD•NUP42 GBM
Preincubation
24 (59)
40 (42)
21 (17)
GLE1
NUP42
F
SUMO-NUP214 TAIL
SUMO-NUP42 ZnF
Preincubation
21 (34)
22 (17)
17 (17)

G

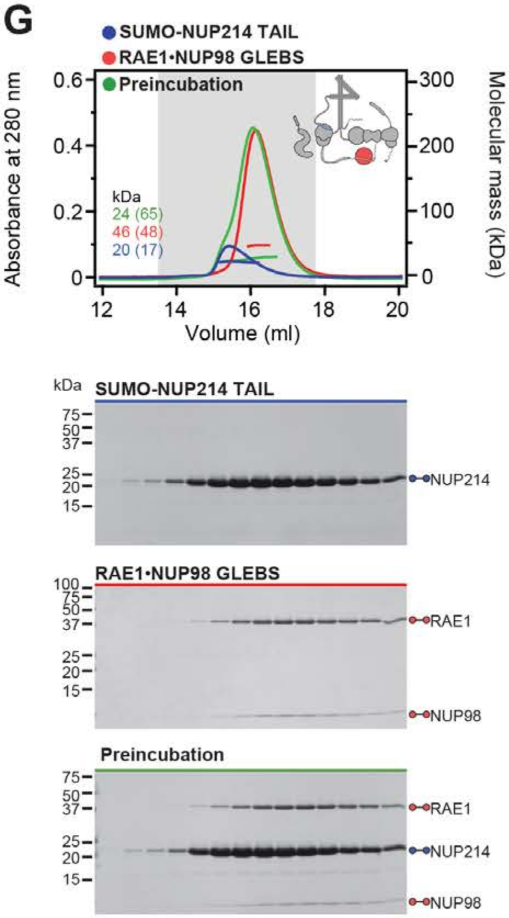

SUMO-NUP214 TAIL
RAE1•NUP98 GLEBS
Preincubation
Absorbance at 280 nm
Molecular mass (kDa)
kDa
24 (65)
46 (48)
20 (17)
Volume (ml)
kDa
SUMO-NUP214 TAIL
NUP214
RAE1•NUP98 GLEBS
RAE1
NUP98
Preincubation
RAE1
NUP214
NUP98

**Fig. S34. Systematic interaction analyses of NUP214$^{TAIL}$ with human CF nup fragments.** SEC-MALS interaction analyses of SUMO-NUP214$^{TAIL}$ with (**A**) NUP214$^{NTD}$, (**B**) CFNC-hub, (**C**) DDX19(ADP), (**D**) DDX19(AMPPNP) E243Q, (**E**) GLE1$^{CTD}$•NUP42, (**F**) SUMO-NUP42$^{ZnF}$, and (**G**) RAE1•NUP98$^{GLEBS}$. SEC-MALS profiles of nup complexes are shown individually (red and blue) and after preincubation (green). SEC profiles were obtained using a Superdex 200 10/300 GL column. Measured molecular masses are indicated, with the respective theoretical masses shown in parentheses. Inset schematics display tested protein samples colored to match the corresponding chromatogram trace. Gray boxes indicate fractions resolved on SDS-PAGE gels and visualized by Coomassie brilliant blue staining.

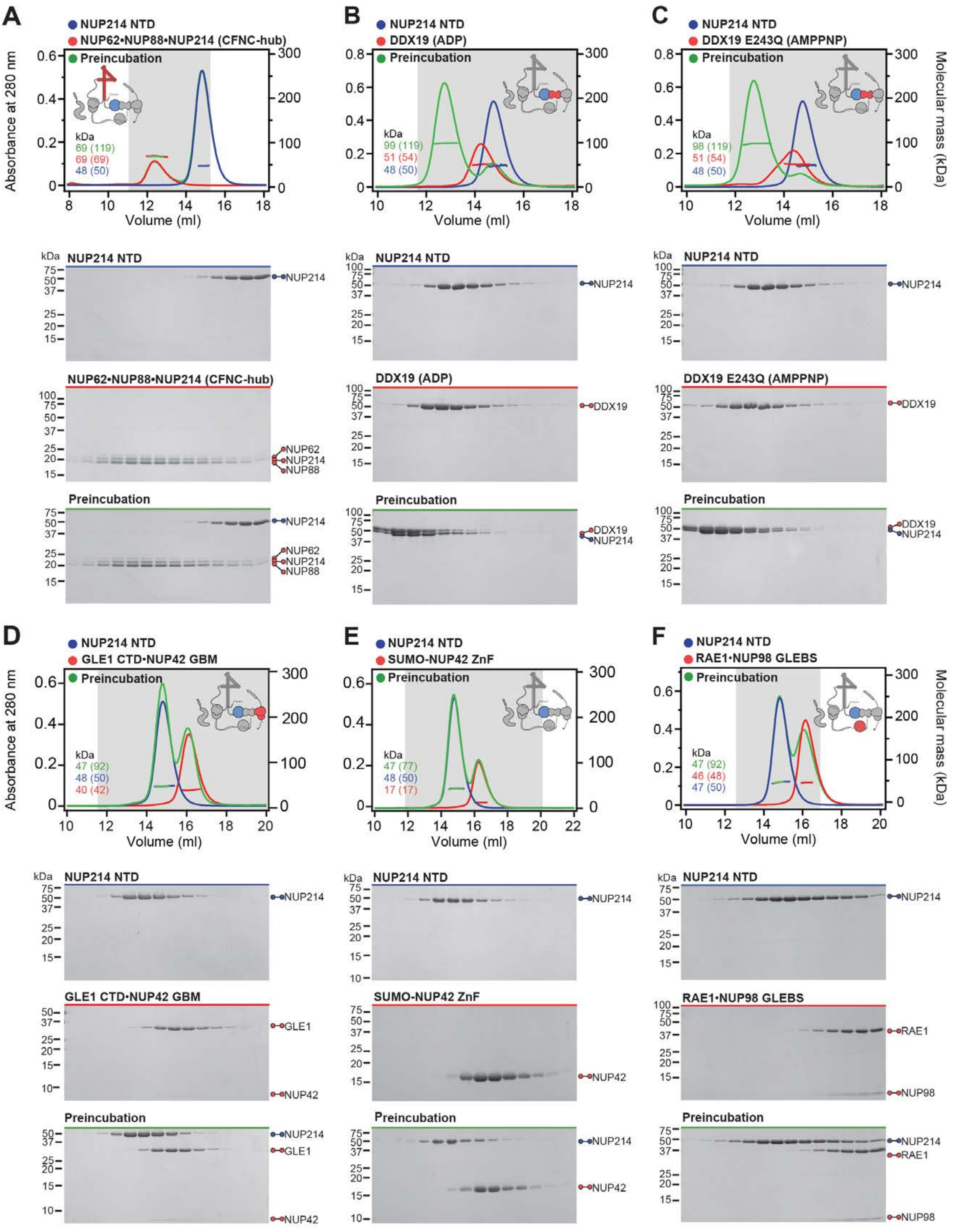

A
NUP214 NTD
NUP62•NUP88•NUP214 (CFNC-hub)
Preincubation
Absorbance at 280 nm
Volume (ml)
kDa
69 (119)
69 (69)
48 (50)
B
NUP214 NTD
DDX19 (ADP)
Preincubation
kDa
99 (119)
51 (54)
48 (50)
C
NUP214 NTD
DDX19 E243Q (AMPPNP)
Preincubation
Molecular mass (kDa)
kDa
98 (119)
51 (54)
48 (50)
NUP214 NTD
NUP62•NUP88•NUP214 (CFNC-hub)
Preincubation
NUP214
NUP62
NUP88
DDX19 (ADP)
DDX19
DDX19 E243Q (AMPPNP)
D
NUP214 NTD
GLE1 CTD•NUP42 GBM
Preincubation
kDa
47 (92)
48 (50)
40 (42)
E
NUP214 NTD
SUMO-NUP42 ZnF
Preincubation
kDa
47 (77)
48 (50)
17 (17)
F
NUP214 NTD
RAE1•NUP98 GLEBS
Preincubation
kDa
47 (92)
46 (48)
47 (50)
GLE1
NUP42
RAE1
NUP98

**Fig. S35. Systematic interaction analyses of NUP214$^{TAIL}$ with human CF nup fragments.** SEC-MALS interaction analyses of NUP214$^{NTD}$ with (**A**) CFNC-hub, (**B**) DDX19(ADP), (**C**) DDX19(AMPPNP) E243Q, (**D**) GLE1$^{CTD}$•NUP42, (**E**) SUMO-NUP42$^{ZnF}$, and (**F**) RAE1•NUP98$^{GLEBS}$. SEC-MALS profiles of nup complexes are shown individually (red and blue) and after preincubation (green). SEC profiles were obtained using a Superdex 200 10/300 GL column. Measured molecular masses are indicated, with the respective theoretical masses shown in parentheses. Inset schematics display tested protein samples colored to match the corresponding chromatogram trace. Gray boxes indicate fractions resolved on SDS-PAGE gels and visualized by Coomassie brilliant blue staining.

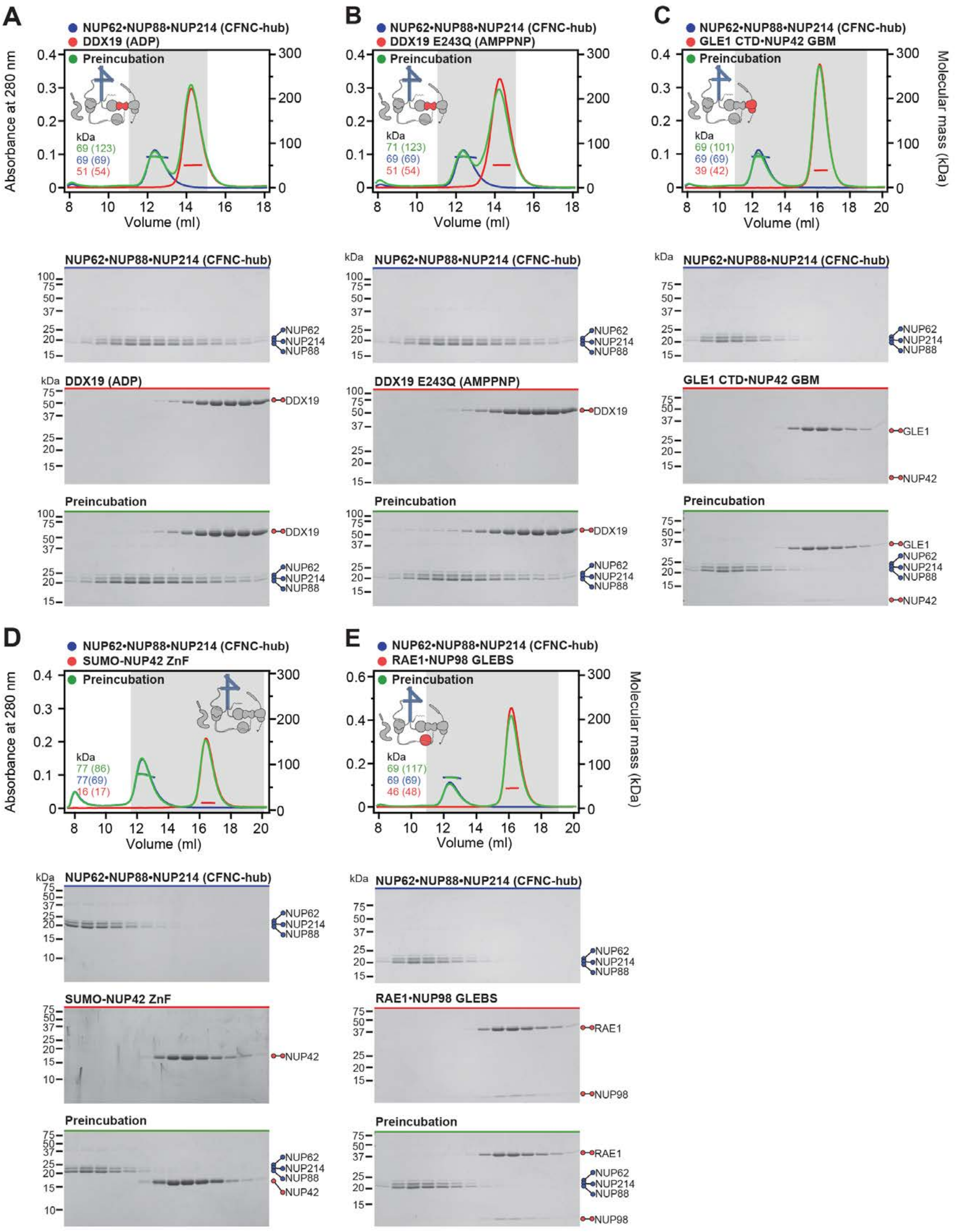

A
NUP62•NUP88•NUP214 (CFNC-hub)
DDX19 (ADP)
Preincubation
Absorbance at 280 nm
Volume (ml)
kDa
69 (123)
69 (69)
51 (54)
B
DDX19 E243Q (AMPPNP)
71 (123)
69 (69)
51 (54)
C
GLE1 CTD•NUP42 GBM
Molecular mass (kDa)
69 (101)
69 (69)
39 (42)
D
SUMO-NUP42 ZnF
77 (86)
77(69)
16 (17)
E
RAE1•NUP98 GLEBS
69 (117)
69 (69)
46 (48)
NUP62
NUP214
NUP88
DDX19
GLE1
NUP42
RAE1
NUP98

**Fig. S36. Systematic interaction analyses of NUP214$^{TAIL}$ with human CF nup fragments.** SEC-MALS interaction analyses of CFNC-hub with (**A**) DDX19 (ADP), (**B**) DDX19(AMPPNP) E243Q, (**C**) GLE1$^{CTD}$•NUP42, (**D**) SUMO-NUP42$^{ZnF}$, and (**E**) RAE1•NUP98$^{GLEBS}$. SEC-MALS profiles of nups complexes are shown individually (red and blue) and after preincubation (green). SEC profiles were obtained using a Superdex 200 10/300 GL column. Measured molecular masses are indicated, with the respective theoretical masses shown in parentheses. Inset schematics display tested protein samples colored to match the corresponding chromatogram trace. Gray boxes indicate fractions resolved on SDS-PAGE gels and visualized by Coomassie brilliant blue staining.

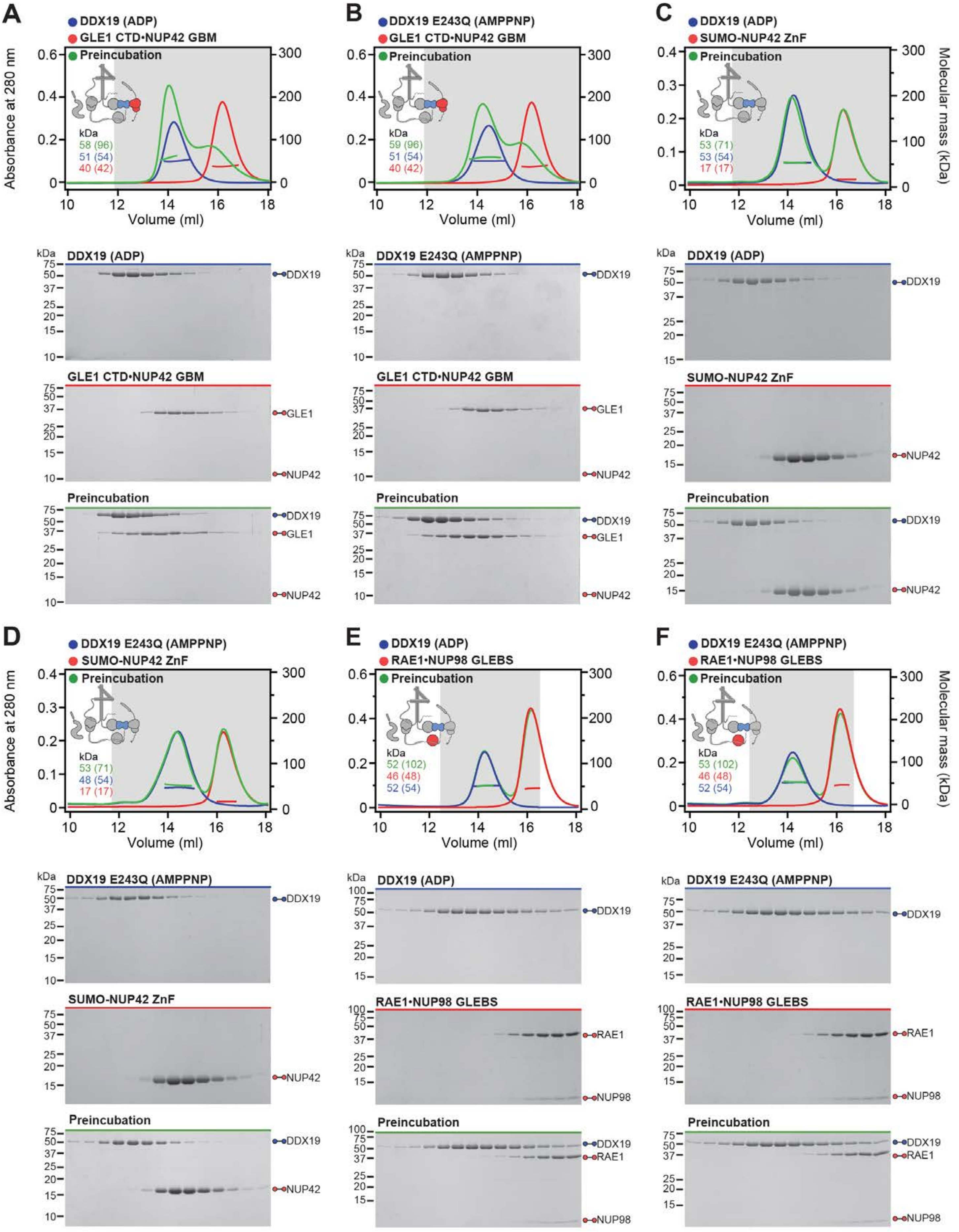
A
DDX19 (ADP)
GLE1 CTD•NUP42 GBM
Preincubation
kDa
58 (96)
51 (54)
40 (42)
Absorbance at 280 nm
Volume (ml)
B
DDX19 E243Q (AMPPNP)
GLE1 CTD•NUP42 GBM
Preincubation
kDa
59 (96)
51 (54)
40 (42)
C
DDX19 (ADP)
SUMO-NUP42 ZnF
Preincubation
kDa
53 (71)
53 (54)
17 (17)
Molecular mass (kDa)
DDX19 (ADP)
GLE1 CTD•NUP42 GBM
Preincubation
DDX19
GLE1
NUP42
DDX19 E243Q (AMPPNP)
SUMO-NUP42 ZnF
D
DDX19 E243Q (AMPPNP)
SUMO-NUP42 ZnF
Preincubation
kDa
53 (71)
48 (54)
17 (17)
E
DDX19 (ADP)
RAE1•NUP98 GLEBS
Preincubation
kDa
52 (102)
46 (48)
52 (54)
F
DDX19 E243Q (AMPPNP)
RAE1•NUP98 GLEBS
Preincubation
kDa
53 (102)
46 (48)
52 (54)
RAE1
NUP98

**Fig. S37. Systematic interaction analyses of DDX19 with human CF nup fragments.** SEC-MALS interaction analyses of DDX19 (ADP) or DDX19(AMPPNP) E243Q with (**A, B**) $GLE1^{CTD}$•NUP42, (**C, D**) $SUMO\text{-}NUP42^{ZnF}$, and (**E, F**) RAE1•$NUP98^{GLEBS}$. SEC-MALS profiles of nup complexes are shown individually (red and blue) and after preincubation (green). SEC profiles were obtained using a Superdex 200 10/300 GL column. Measured molecular masses are indicated, with the respective theoretical masses shown in parentheses. Inset schematics display tested protein samples colored to match the corresponding chromatogram trace. Gray boxes indicate fractions resolved on SDS-PAGE gels and visualized by Coomassie brilliant blue staining.

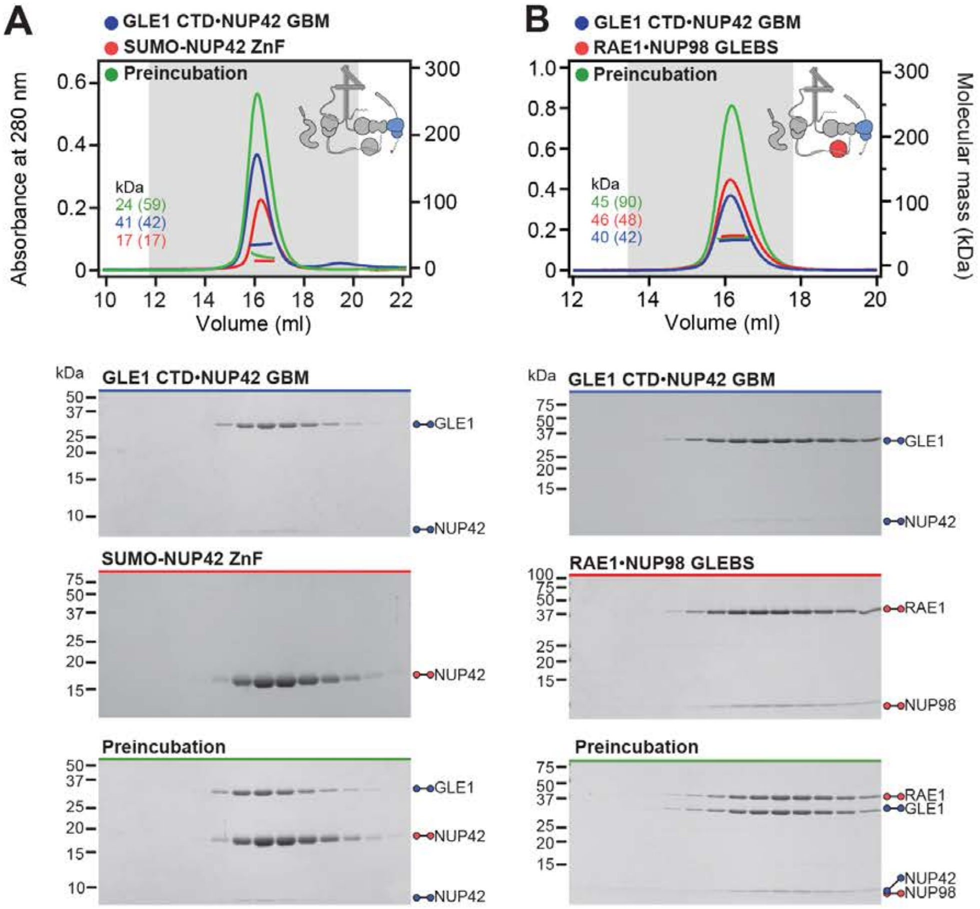

Sup Fig. S38

Hoelz et al., 2021b

**Fig. S38. Systematic interaction analyses of GLE1$^{CTD}$•NUP42 with human CF nup fragments.** SEC-MALS interaction analyses of GLE1$^{CTD}$•NUP42 with (**A**) SUMO-NUP42$^{ZnF}$, and (**B**) RAE1•NUP98$^{GLEBS}$. SEC-MALS profiles of nup complexes are shown individually (red and blue) and after preincubation (green). SEC profiles were obtained using a Superdex 200 10/300 GL column. Measured molecular masses are indicated, with the respective theoretical masses shown in parentheses. Inset schematics display tested protein samples colored to match the corresponding chromatogram trace. Gray boxes indicate fractions resolved on SDS-PAGE gels and visualized by Coomassie brilliant blue staining.

A

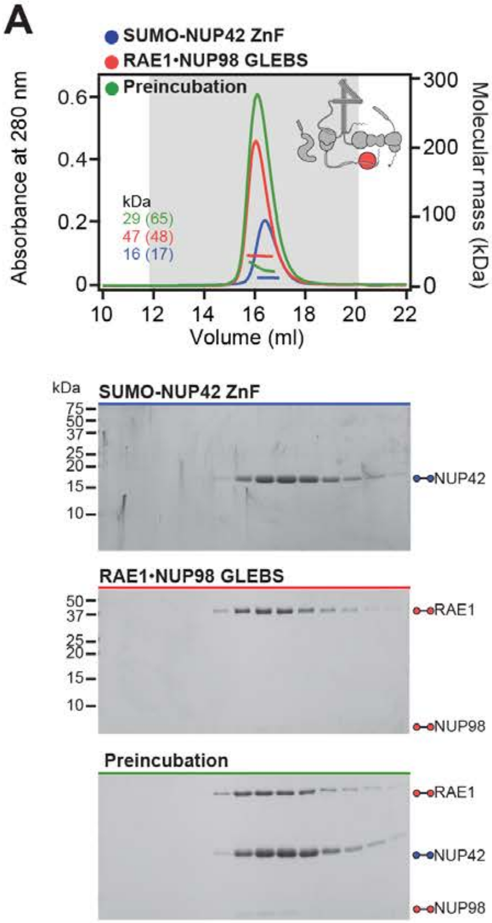
SUMO-NUP42 ZnF
RAE1•NUP98 GLEBS
Preincubation
Absorbance at 280 nm
Molecular mass (kDa)
kDa
29 (65)
47 (48)
16 (17)
Volume (ml)
SUMO-NUP42 ZnF
NUP42
RAE1•NUP98 GLEBS
RAE1
NUP98
Preincubation
RAE1
NUP42
NUP98

**Fig. S39. Systematic interaction analyses of NUP42$^{ZnF}$ and RAE1•NUP98$^{GLEBS}$ with human CF nup fragments.** SEC-MALS interaction analysis of SUMO-NUP42$^{ZnF}$ with (**A**) RAE1•NUP98$^{GLEBS}$. SEC-MALS profiles of SUMO-NUP42$^{ZnF}$ and RAE1•NUP98$^{GLEBS}$ are shown individually (blue and red, respectively) and after preincubation (green). SEC profiles were obtained using a Superdex 200 10/300 GL column. Measured molecular masses are indicated, with the respective theoretical masses shown in parentheses. Inset schematics display tested protein samples colored to match the corresponding chromatogram trace. Gray boxes indicate fractions resolved on SDS-PAGE gels and visualized by Coomassie brilliant blue staining.

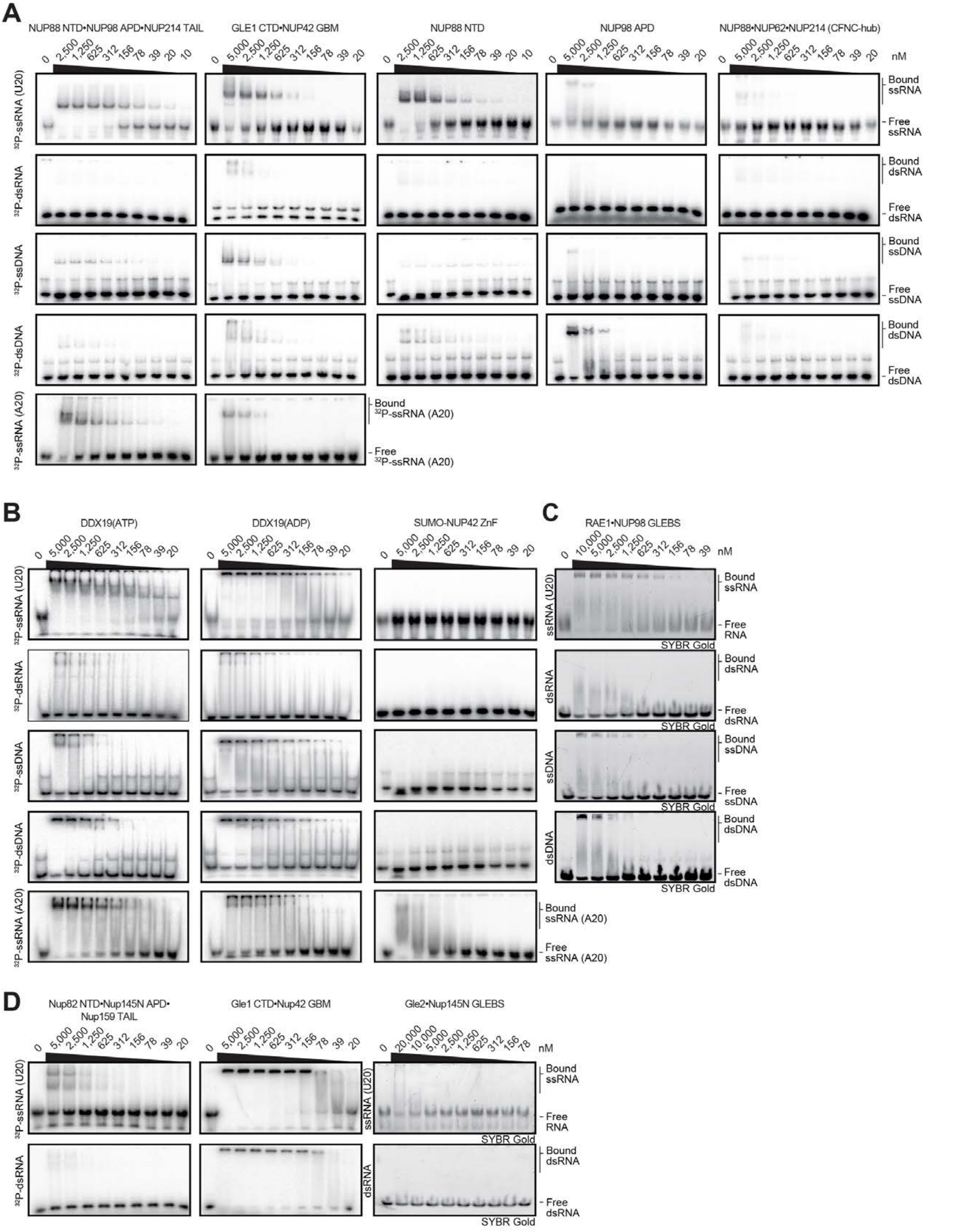
A
NUP88 NTD•NUP98 APD•NUP214 TAIL
GLE1 CTD•NUP42 GBM
NUP88 NTD
NUP98 APD
NUP88•NUP62•NUP214 (CFNC-hub)
nM
Bound ssRNA
Free ssRNA
Bound dsRNA
Free dsRNA
Bound ssDNA
Free ssDNA
Bound dsDNA
Free dsDNA
Bound 32P-ssRNA (A20)
Free 32P-ssRNA (A20)
B
DDX19(ATP)
DDX19(ADP)
SUMO-NUP42 ZnF
Bound ssRNA (A20)
Free ssRNA (A20)
C
RAE1•NUP98 GLEBS
SYBR Gold
Free RNA
D
Nup82 NTD•Nup145N APD•
Nup159 TAIL
Gle1 CTD•Nup42 GBM
Gle2•Nup145N GLEBS

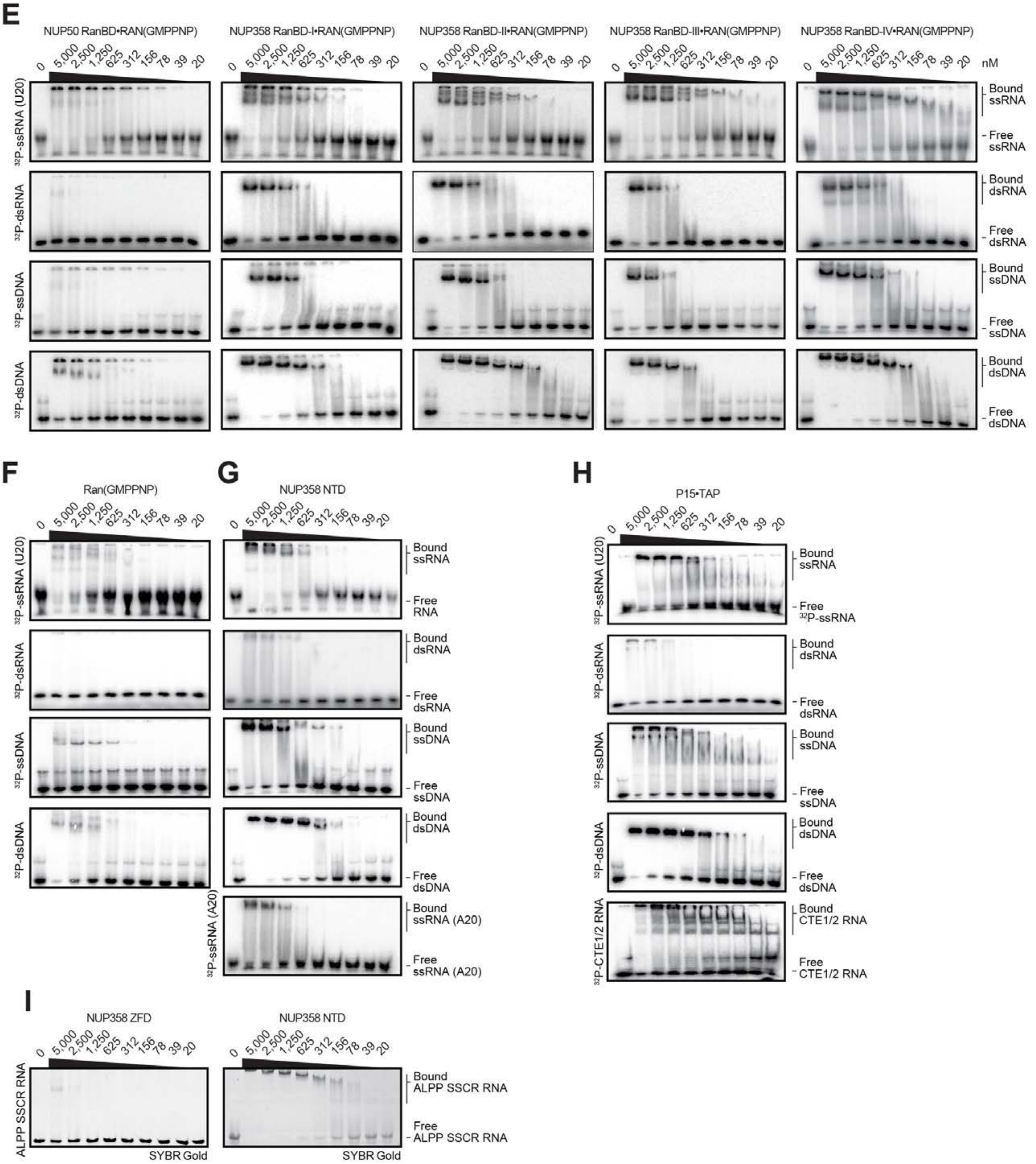
E
NUP50 RanBD•RAN(GMPPNP)
NUP358 RanBD-I•RAN(GMPPNP)
NUP358 RanBD-II•RAN(GMPPNP)
NUP358 RanBD-III•RAN(GMPPNP)
NUP358 RanBD-IV•RAN(GMPPNP)
0 5,000 2,500 1,250 625 312 156 78 39 20 nM
32P-ssRNA (U20)
32P-dsRNA
32P-ssDNA
32P-dsDNA
Bound ssRNA
Free ssRNA
Bound dsRNA
Free dsRNA
Bound ssDNA
Free ssDNA
Bound dsDNA
Free dsDNA
F
Ran(GMPPNP)
G
NUP358 NTD
Free RNA
32P-ssRNA (A20)
Bound ssRNA (A20)
Free ssRNA (A20)
H
P15•TAP
Free 32P-ssRNA
32P-CTE1/2 RNA
Bound CTE1/2 RNA
Free CTE1/2 RNA
I
NUP358 ZFD
NUP358 NTD
ALPP SSCR RNA
Bound ALPP SSCR RNA
Free ALPP SSCR RNA
SYBR Gold

**Fig. S40. Evolutionary conservation of cytoplasmic filament nup RNA binding properties.** Cytoplasmic filament nup domains and complexes were assayed for binding to single stranded (ss) and double stranded (ds) RNA and DNA probes by electrophoretic mobility shift assay (EMSA); ssRNA ($U_{20}$), dsRNA (28-nt hairpin), ssDNA (34-nt DNA oligonucleotide), dsDNA (34-bp hybridized DNA oligonucleotides) and ssRNA ($A_{20}$). Final concentrations of protein and complexes are indicated. EMSA titration of (**A**-**C**) human CFNC nup domains and subcomplexes, (**D**) *C. thermophilum* CFNC subcomplexes, (**E**) NUP358 and NUP50 RanBDs pre-complexed with Ran(GMPPNP), (**F**) Ran(GMPPMP), and (**G**) NUP358$^{NTD}$. (**H**) EMSA titration of mRNA export complex P15•TAP with ss/ds RNA and DNA and CTE1/2 RNA, a commonly used viral RNA probe for P15•TAP. (**I**) EMSA titrations of NUP358$^{NTD}$ and NUP358$^{ZFD}$ with ALPP$^{SSCR}$ RNA.

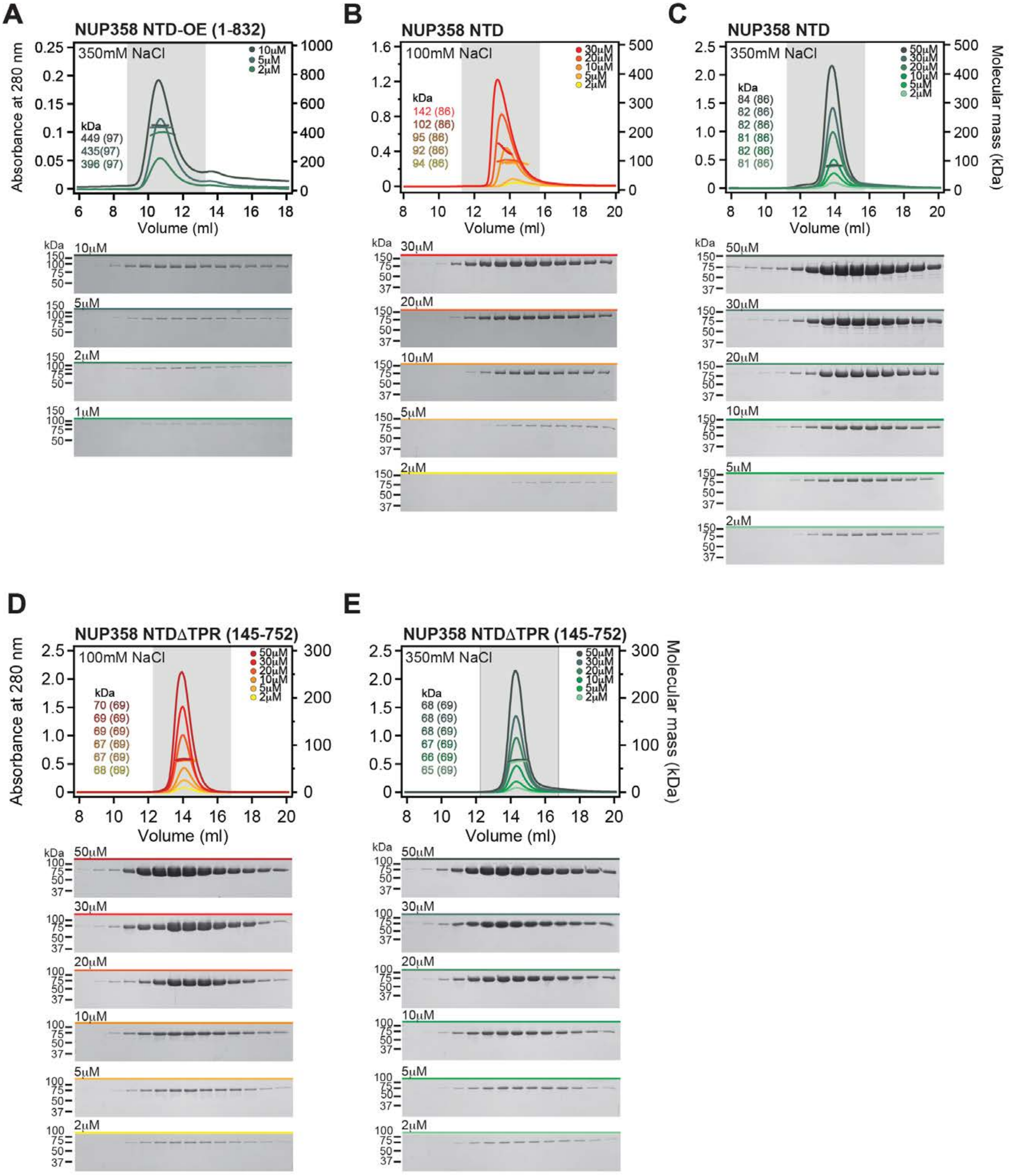

A
NUP358 NTD-OE (1-832)
350mM NaCl
kDa
449 (97)
435(97)
396 (97)
B
NUP358 NTD
100mM NaCl
kDa
142 (86)
102 (86)
95 (86)
92 (86)
94 (86)
C
NUP358 NTD
350mM NaCl
kDa
84 (86)
82 (86)
82 (86)
81 (86)
82 (86)
81 (86)
D
NUP358 NTDΔTPR (145-752)
100mM NaCl
kDa
70 (69)
69 (69)
69 (69)
67 (69)
67 (69)
68 (69)
E
NUP358 NTDΔTPR (145-752)
350mM NaCl
kDa
68 (69)
68 (69)
68 (69)
67 (69)
66 (69)
65 (69)
Absorbance at 280 nm
Molecular mass (kDa)
Volume (ml)

**Fig. S41. Biochemical analysis of oligomerization properties of NUP358 fragments.** (**A**) SEC-MALS analysis of NUP358$^{NTD\text{-}OE}$ at the indicated increasing protein concentrations. The measured molecular masses increase from ~400 kDa to ~450 kDa, indicating the formation of a concentration-dependent oligomer, ranging from a homo-tetramer or -pentamer. (**B**, **C**) SEC-MALS oligomerization analyses of NUP358$^{NTD}$ at the indicated increasing protein concentrations and buffer conditions. NUP358$^{NTD}$ forms a concentration-dependent monomer-dimer mixture that is interrupted by 350 mM NaCl. (**D**, **E**) SEC-MALS oligomerization analyses of NUP358$^{NTD\Delta TPR}$ at the indicated increasing protein concentrations and buffer conditions. NUP358$^{NTD\Delta TPR}$ is monomeric in all buffer conditions tested, establishing that NUP358$^{TPR}$ contributes to NUP358$^{NTD}$ dimerization. Measured molecular masses are indicated, with the respective theoretical masses shown in parentheses. Gray boxes indicate fractions resolved on SDS-PAGE gels and visualized by Coomassie brilliant blue staining.

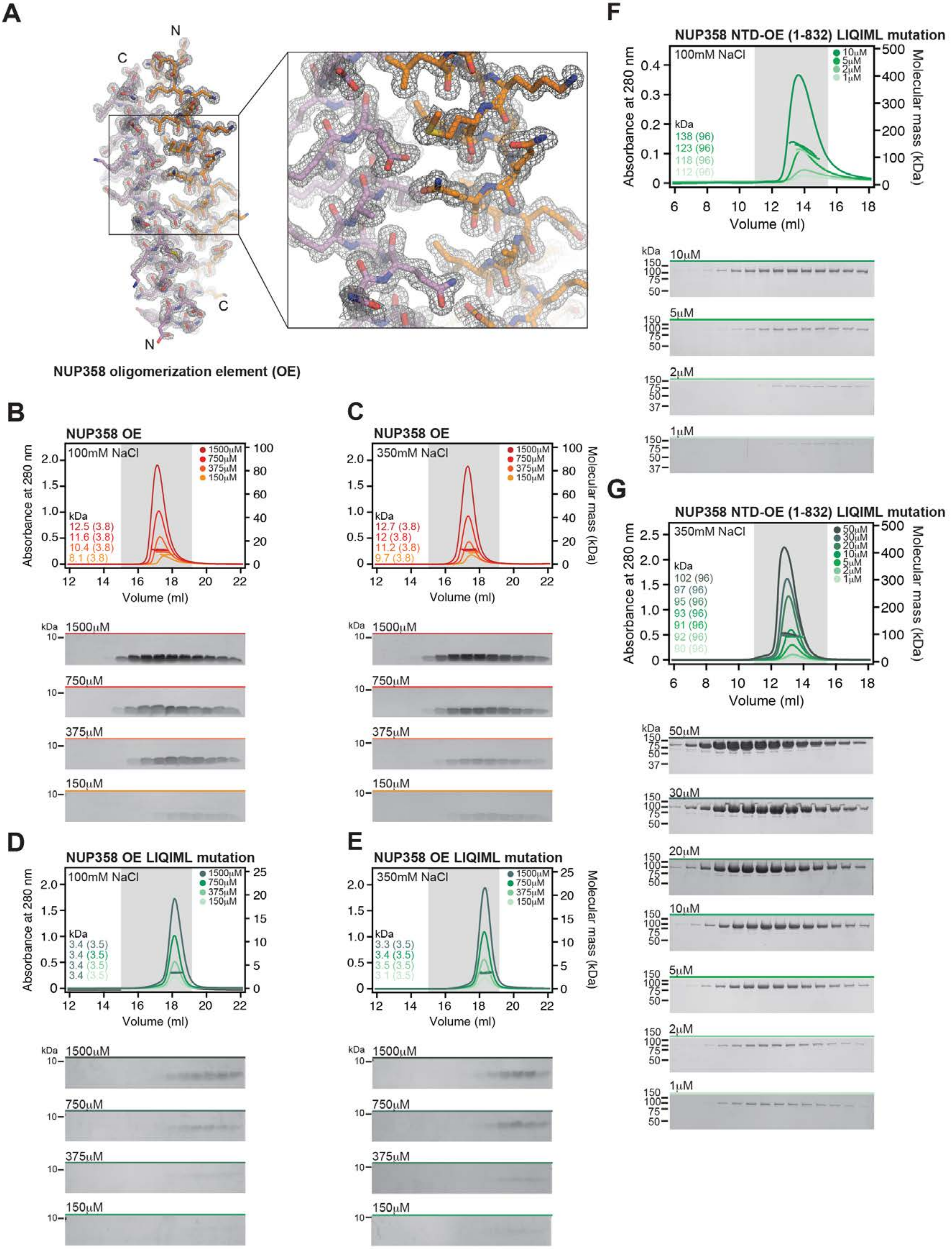
A
N
C
C
N
NUP358 oligomerization element (OE)
B
NUP358 OE
100mM NaCl
1500μM
750μM
375μM
150μM
kDa
12.5 (3.8)
11.6 (3.8)
10.4 (3.8)
8.1 (3.8)
Absorbance at 280 nm
Volume (ml)
C
NUP358 OE
350mM NaCl
12.7 (3.8)
12 (3.8)
11.2 (3.8)
9.7 (3.8)
Molecular mass (kDa)
D
NUP358 OE LIQIML mutation
100mM NaCl
3.4 (3.5)
3.4 (3.5)
3.4 (3.5)
3.4 (3.5)
E
NUP358 OE LIQIML mutation
350mM NaCl
3.3 (3.5)
3.4 (3.5)
3.5 (3.5)
3.1 (3.5)
F
NUP358 NTD-OE (1-832) LIQIML mutation
100mM NaCl
10μM
5μM
2μM
1μM
138 (96)
123 (96)
118 (96)
112 (96)
G
NUP358 NTD-OE (1-832) LIQIML mutation
350mM NaCl
50μM
30μM
20μM
10μM
5μM
2μM
1μM
102 (96)
97 (96)
95 (96)
93 (96)
91 (96)
92 (96)
90 (96)

**Fig. S42. Structural and biochemical characterization of NUP358$^{OE}$.** (**A**) Experimental electron density map after density modification of NUP358$^{OE}$ phased with the sulfur-SAD anomalous X-ray diffraction data, contoured at 1.0 $\sigma$. (**B**, **C**) SEC-MALS oligomerization analysis of NUP358$^{OE}$, at the indicated increasing protein concentrations and buffer conditions. NUP358$^{OE}$ forms oligomeric states ranging from homo-dimer to -tetramer independent of the buffer condition. (**D**, **E**) SEC-MALS oligomerization analysis of NUP358$^{OE}$ LIQIML mutant, at the indicated increasing protein concentrations and buffer conditions. The measured molecular masses at all proteins concentrations and both buffer conditions correspond to a monomeric species, demonstrating that the LIQIML mutation abolishes the oligomeric behavior of NUP358$^{OE}$. (**F**, **G**) SEC-MALS oligomerization analysis of NUP358$^{NTD\text{-}OE}$ LIQIML mutant, at the indicated increasing protein concentrations and buffer conditions. The LIQIML mutation abolishes the homo-tetramer or -pentamer formation, yielding a dynamic monomer-dimer equilibrium, as observed for NUP358$^{NTD}$. In the 350 mM NaCl buffer, the NUP358$^{NTD\text{-}OE}$ LIQIML mutant is a monodisperse monomer at all protein concentrations tested. SEC profiles were obtained using a Superdex 200 10/300 GL column. Measured molecular masses are indicated, with the respective theoretical masses shown in parentheses. Gray boxes indicate fractions resolved on SDS-PAGE gels and visualized by Coomassie brilliant blue staining.

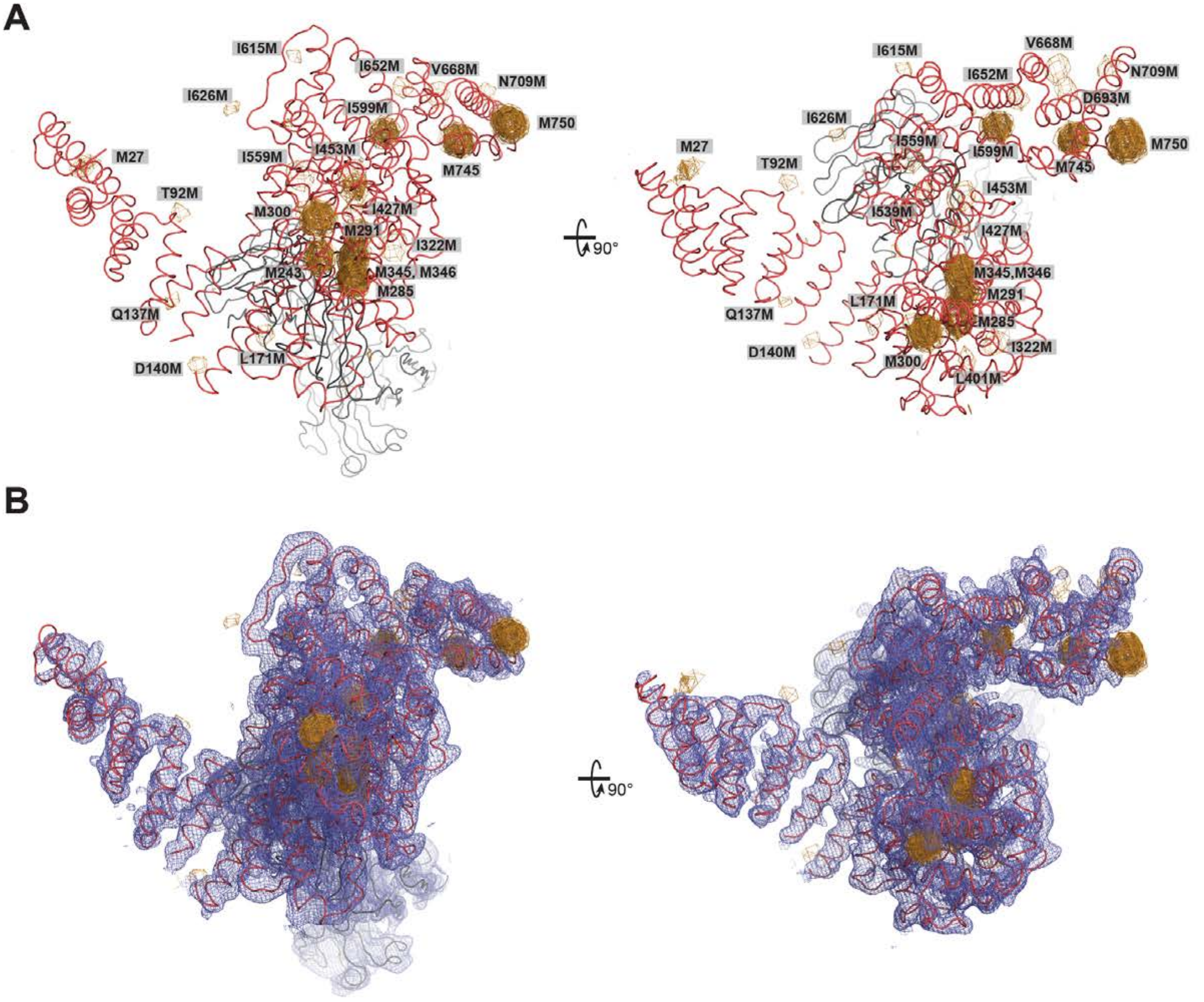
A
I615M
I652M
V668M
N709M
I626M
I599M
M750
M27
I559M
I453M
M745
T92M
M300
I427M
M291
I322M
M243
M345, M346
M285
Q137M
D140M
L171M
90°
I615M
V668M
I652M
N709M
D693M
I626M
M27
I559M
I599M
M750
T92M
M745
I453M
I539M
I427M
M345,M346
L171M
M291
Q137M
M285
D140M
I322M
M300
L401M
B
90°

**Fig. 43. Determination of the NUP358$^{NTD}$•sAB-14 crystal structure and sequence assignment.** (**A**) The NUP358$^{NTD}$•sAB-14 structure is shown in coil representation and overlayed with anomalous difference Fourier maps (contoured at 3.5 $\sigma$), calculated from SeMet-SAD X-ray diffraction data obtained from SeMet-labeled crystals of 17 different methionine mutants. Together with the endogenous methionines of the wildtype complex, this analysis identified a total of 26 selenium positions, providing a marker for the unambiguous assignment of the NUP358$^{NTD}$ sequence register. A 90º-rotated view is shown on the right. For clarity, the second NUP358$^{NTD}$•sAB-14 copy in the asymmetric unit is omitted. (**B**) Final 2|Fo|-|Fc| electron density map contoured at 1.0 $\sigma$.

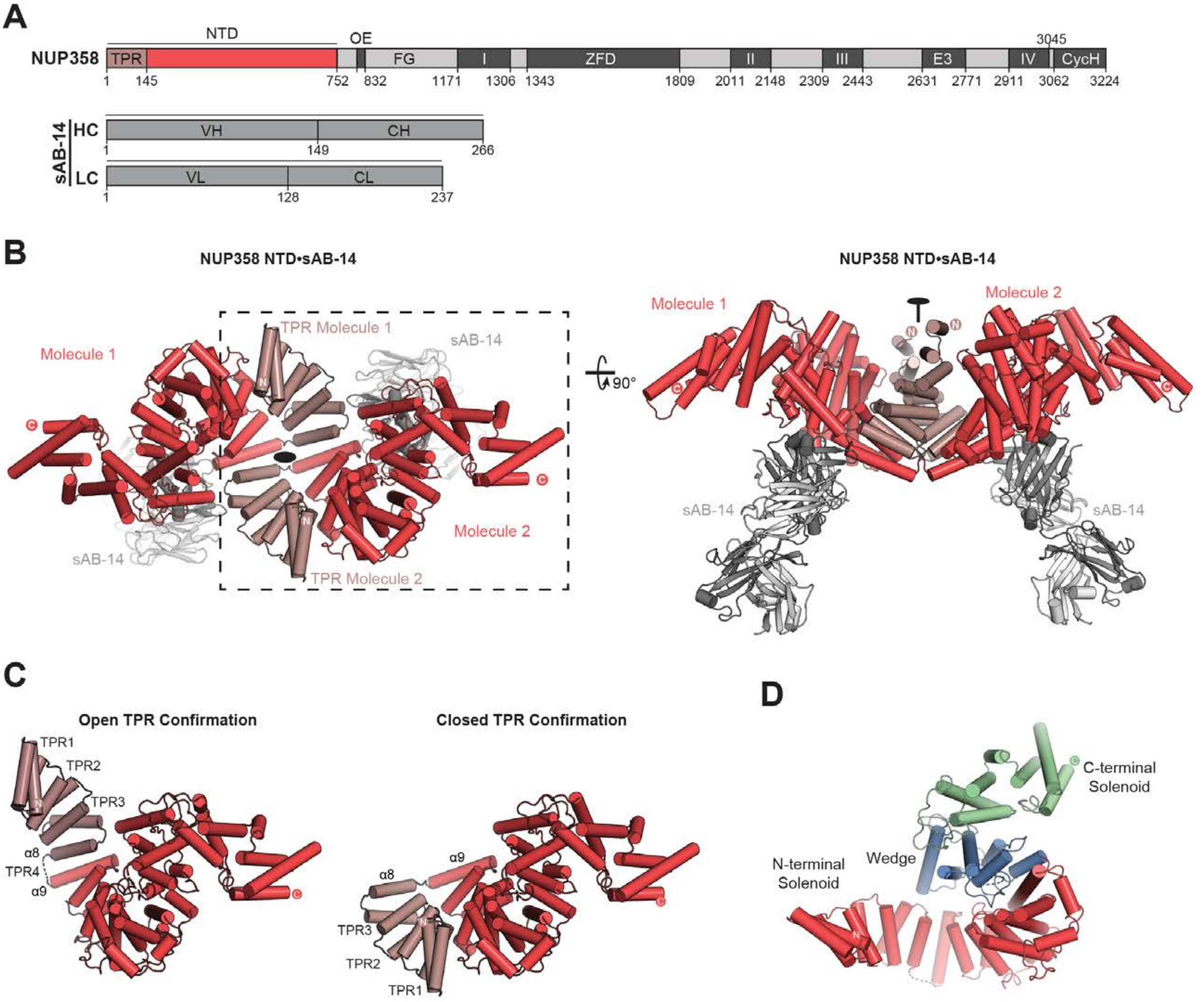
A
NUP358
NTD
OE
TPR
FG
I
ZFD
II
III
E3
IV
CycH
3045
1
145
752
832
1171
1306
1343
1809
2011
2148
2309
2443
2631
2771
2911
3062
3224
sAB-14
HC
VH
CH
1
149
266
LC
VL
CL
1
128
237
B
NUP358 NTD•sAB-14
Molecule 1
TPR Molecule 1
sAB-14
Molecule 2
TPR Molecule 2
90°
NUP358 NTD•sAB-14
Molecule 1
Molecule 2
sAB-14
sAB-14
C
Open TPR Confirmation
TPR1
TPR2
TPR3
α8
TPR4
α9
Closed TPR Confirmation
α9
α8
TPR3
TPR2
TPR1
D
C-terminal Solenoid
Wedge
N-terminal Solenoid

**Fig. S44. Crystal structure of NUP358$^{NTD}$ bound by sAB-14.** (**A**) Domain structure of NUP358 and sAB-14 light and heavy chains drawn as horizontal boxes with residue numbers indicated. Black lines indicate the boundaries of the crystallized fragments. (**B**) Cartoon representation of the NUP358$^{NTD}$•sAB-14 co-crystal structure dyad of the $P6_522$ lattice, illustrating the swap of the N-terminal TPR subdomain between symmetry related NUP358$^{NTD}$ molecules. A 90°-rotated view highlights sAB-14 binding to the convex surface of NUP358$^{NTD}$. (**C**) Comparison of open and closed conformations of NUP358$^{NTD}$. The crystallographic dimer reveals two alternative conformations of NUP358$^{NTD}$, in which NUP358$^{TPR}$ either forms a continuous solenoid (open), or folds back, separating TPR4 and forming electrostatic interactions with HEAT repeats 5-7 of the N-terminal solenoid (closed). The open conformation adopts an overall S-shaped architecture, whereas the closed conformation is a compact particle. (**D**) Cartoon representation colored to indicate the three $\alpha$-helical solenoids that compose the NUP358$^{NTD}$ giving rise to an overall S-shape molecule; N-terminal $\alpha$-helical solenoid (red), $\alpha$-helical wedge (blue), C-terminal $\alpha$-helical solenoid (green).

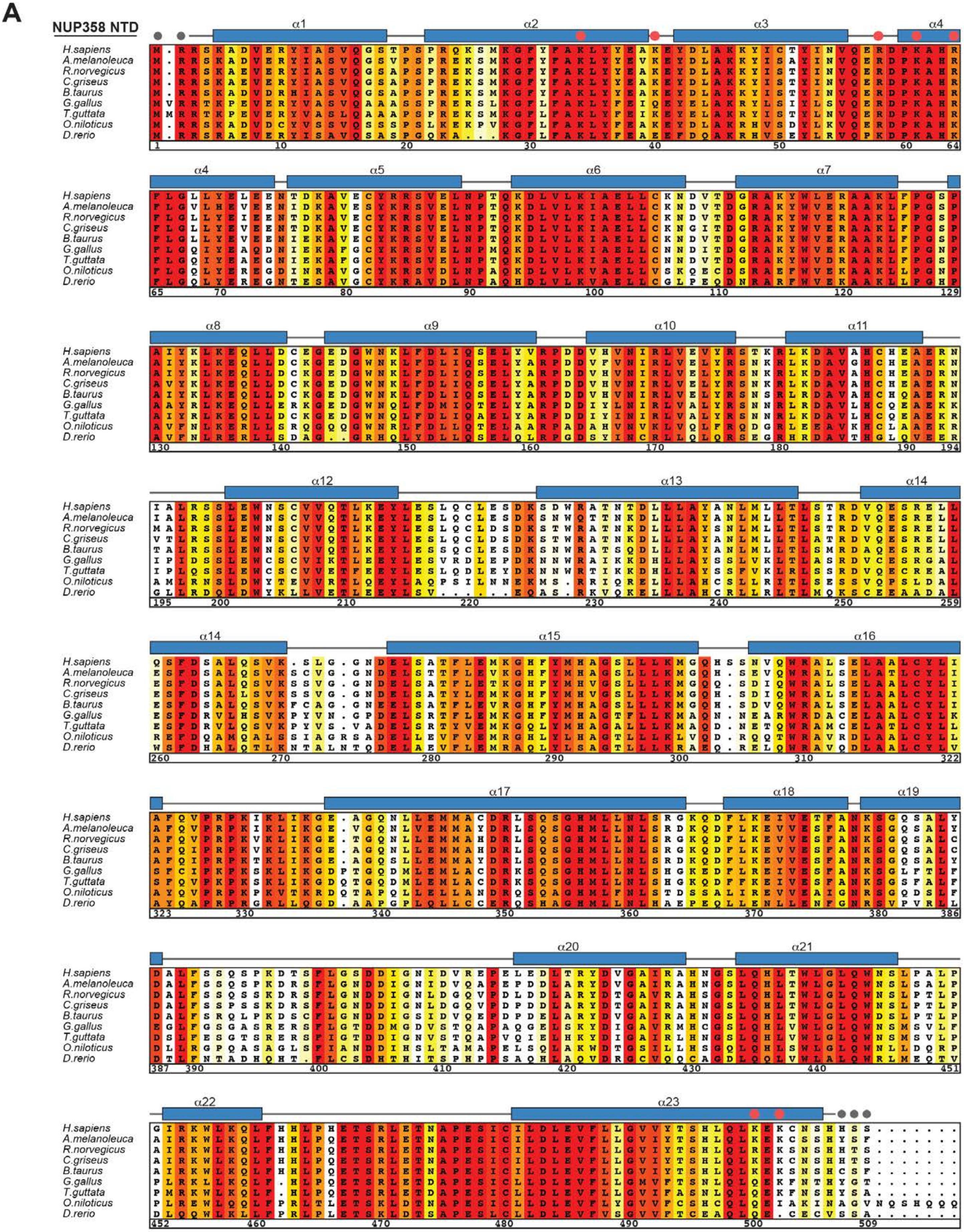
A
NUP358 NTD
H.sapiens
A.melanoleuca
R.norvegicus
C.griseus
B.taurus
G.gallus
T.guttata
O.niloticus
D.rerio

**NUP358 NTD**

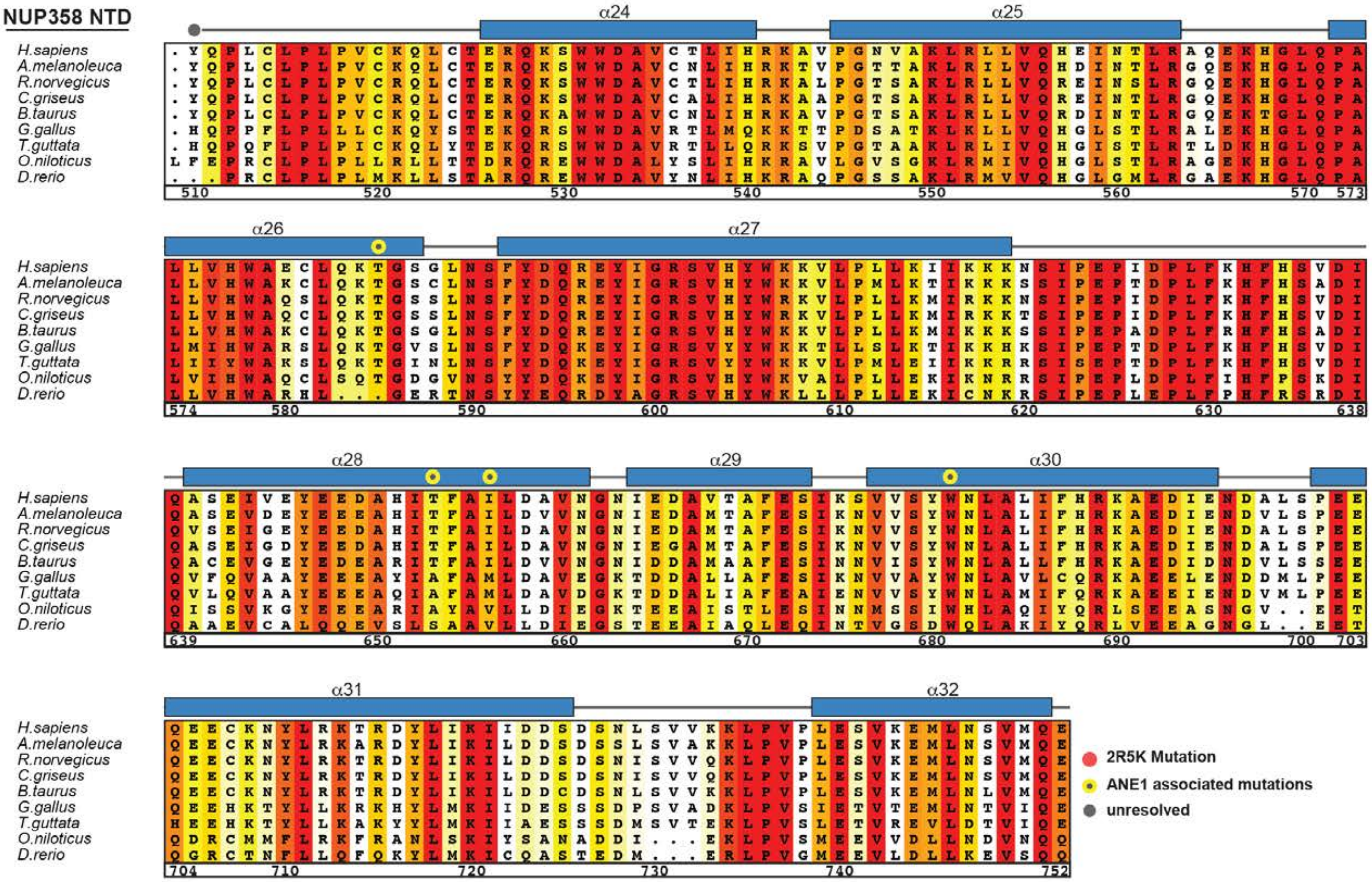
α24
α25
α26
α27
α28
α29
α30
α31
α32
H.sapiens
A.melanoleuca
R.norvegicus
C.griseus
B.taurus
G.gallus
T.guttata
O.niloticus
D.rerio
2R5K Mutation
ANE1 associated mutations
unresolved

**Fig. S45. Multispecies sequence alignment of NUP358$^{NTD}$.** Sequences of nine metazoan species were aligned and colored according to similarity using the BLOSUM62 matrix from white (less than 40% similarity), through yellow (60% similarity), to red (100% identity). The numbering is according to the *H. sapiens* protein. The secondary structure is indicated above the sequences as rectangles ($\alpha$-helices) and lines (unstructured regions), residues unresolved in the crystal structure are indicated by gray dots. Yellow dots indicate residues found in sequence variants associated with acute necrotizing encephalopathy 1 (ANE1). Red dots indicate the seven residues that are substituted by alanine in the 2R5K mutation that abolishes the NUP88$^{NTD}$ and ALPP$^{SSCR}$ RNA interactions.

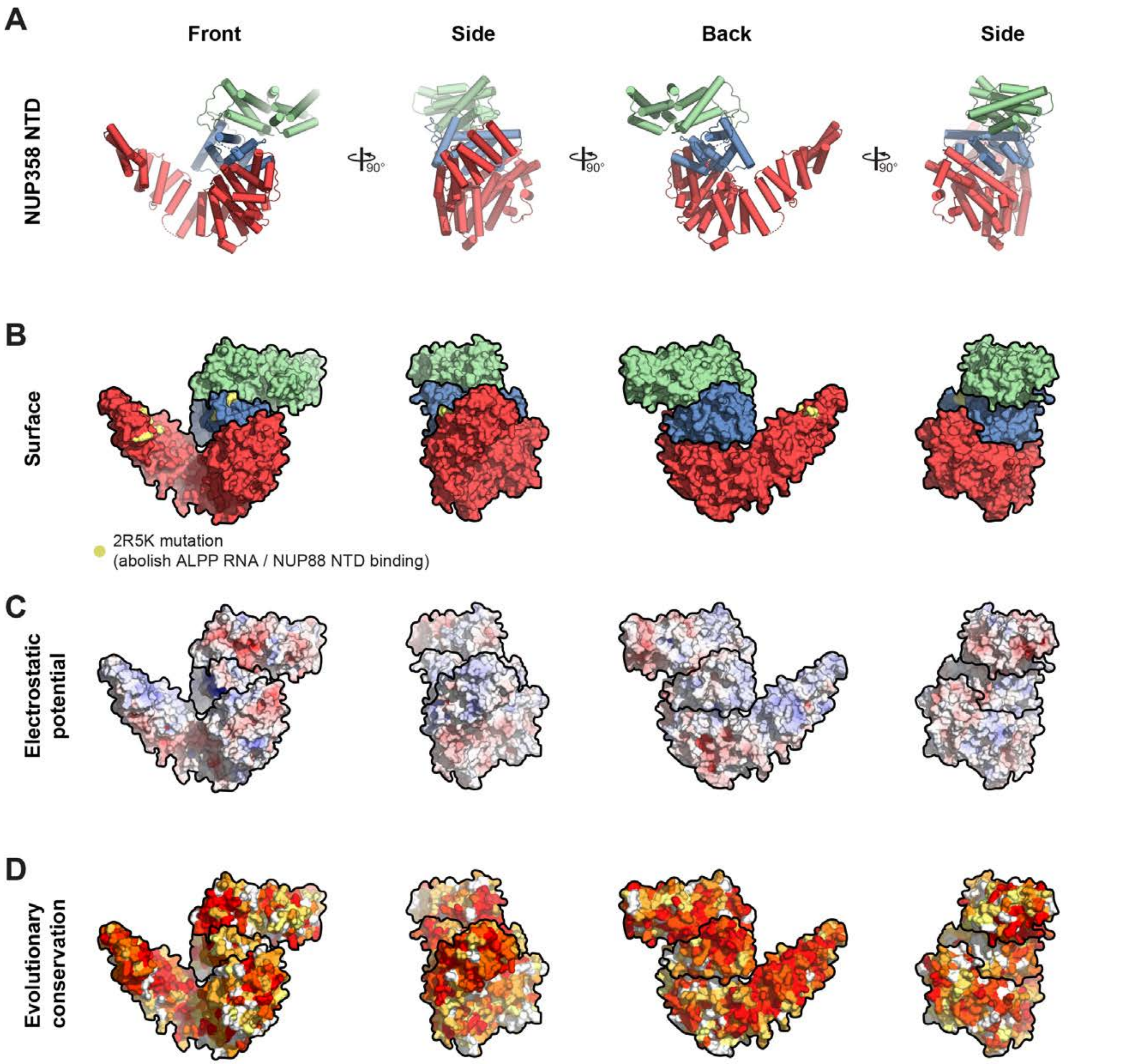
A
Front
Side
Back
Side
NUP358 NTD
90°
90°
90°
B
Surface
2R5K mutation
(abolish ALPP RNA / NUP88 NTD binding)
C
Electrostatic
potential
D
Evolutionary
conservation

**Fig. S46. Surface properties of NUP358$^{NTD}$.** (**A**) Cartoon representation of NUP358$^{NTD}$ with N-terminal $\alpha$-helical solenoid in red, $\alpha$-helical wedge domain in blue, and C-terminal $\alpha$-helical solenoid in green. (**B**) Surface representation of NUP358$^{NTD}$ colored as in (A). Residues mutated to alanine in the 2R5K mutation are indicated in yellow (**C**) Surface representation colored according to electrostatic potential from red (-10 $k_BT/e$) to blue (+10 $k_BT/e$). (**D**) Surface representation colored according to sequence conservation based on alignment in fig. S45. The NUP358$^{NTD}$ structure is shown from four different orientations related by 90° rotations around the vertical axis.

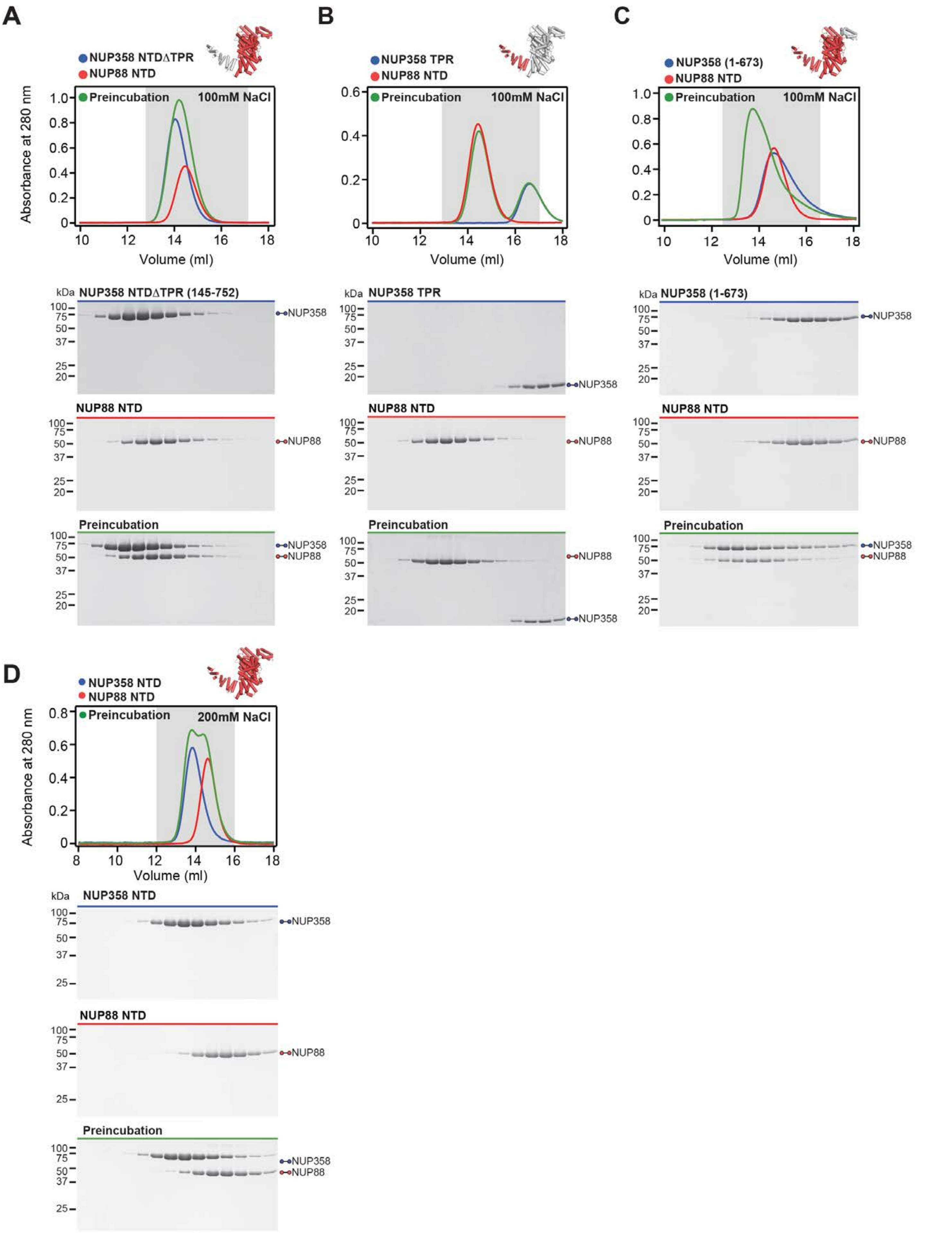

A
NUP358 NTDΔTPR
NUP88 NTD
Preincubation
100mM NaCl
Absorbance at 280 nm
Volume (ml)
kDa
NUP358 NTDΔTPR (145-752)
NUP358
NUP88 NTD
NUP88
Preincubation
B
NUP358 TPR
NUP88 NTD
Preincubation
100mM NaCl
Volume (ml)
NUP358 TPR
C
NUP358 (1-673)
NUP88 NTD
Preincubation
100mM NaCl
Volume (ml)
NUP358 (1-673)
D
NUP358 NTD
NUP88 NTD
Preincubation
200mM NaCl
Absorbance at 280 nm
Volume (ml)
NUP358 NTD

**Fig. S47. Biochemical characterization of the NUP358$^{NTD}$-NUP88$^{NTD}$ interaction.** (**A**-**C**) SEC interaction analyses of NUP88$^{NTD}$ with (A) NUP358$^{NTD\Delta TPR}$, (B) NUP358$^{TPR}$, and (C) NUP358$^{1\text{-}673}$, demonstrating that NUP358$^{TPR}$ is necessary but insufficient for NUP88$^{NTD}$ binding. (**D**) SEC analysis of NUP88$^{NTD}$ with NUP358$^{NTD}$ in a buffer containing 200 mM NaCl, revealing the salt-sensitive nature of the interaction. SEC profiles of NUP358$^{NTD}$ variants (blue) and NUP88$^{NTD}$ (red) are shown individually and after their preincubation (green). SEC profiles were obtained using a Superdex 200 10/300 GL column. Gray boxes indicate fractions resolved on SDS-PAGE gels and visualized by Coomassie brilliant blue staining.

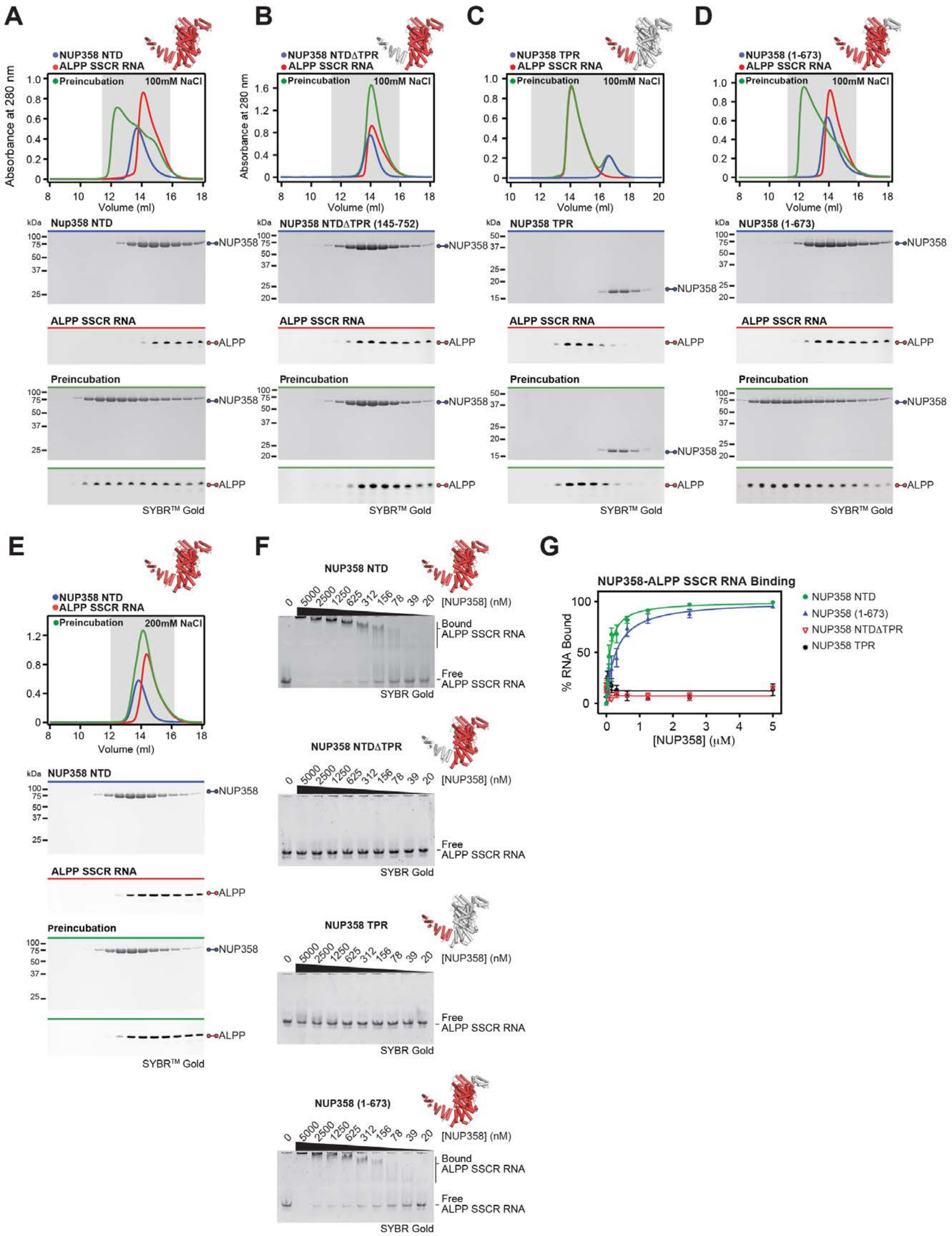

A
NUP358 NTD
ALPP SSCR RNA
Preincubation
100mM NaCl
Absorbance at 280 nm
Volume (ml)
B
NUP358 NTDΔTPR
C
NUP358 TPR
D
NUP358 (1-673)
Nup358 NTD
NUP358 NTDΔTPR (145-752)
NUP358 TPR
NUP358 (1-673)
kDa
NUP358
ALPP
SYBR™ Gold
E
200mM NaCl
F
NUP358 NTD
[NUP358] (nM)
Bound ALPP SSCR RNA
Free ALPP SSCR RNA
SYBR Gold
NUP358 NTDΔTPR
NUP358 TPR
NUP358 (1-673)
G
NUP358-ALPP SSCR RNA Binding
% RNA Bound
[NUP358] (μM)
NUP358 NTD
NUP358 (1-673)
NUP358 NTDΔTPR
NUP358 TPR

**Fig. S48. Biochemical characterization of the NUP358$^{NTD}$-ALPP$^{SSCR}$ RNA interaction.** (**A**-**D**) SEC interaction analyses of ALPP$^{SSCR}$ RNA with (A) NUP358$^{NTD}$, (B) NUP358$^{NTD\Delta TPR}$, (C) NUP358$^{TPR}$, and (D) NUP358$^{1\text{-}673}$, demonstrating that NUP358$^{TPR}$ is necessary but insufficient for ALPP$^{SSCR}$ RNA binding. (**E**) SEC analysis of ALPP$^{SSCR}$ RNA with NUP358$^{NTD}$ in a buffer containing 200 mM NaCl, revealing the salt-sensitive nature of the interaction. SEC profiles of NUP358 variants (blue) and ALPP$^{SSCR}$ RNA (red) are shown individually and after their preincubation (green). SEC profiles were obtained using a Superdex 200 10/300 GL column. Gray boxes indicate fractions resolved on SDS-PAGE and 0.5X TBE urea-PAGE denaturing gels and visualized by Coomassie brilliant blue and SYBR Gold staining, respectively. (**F**) EMSA titrations against ALPP$^{SSCR}$ RNA are shown for each NUP358 variant used in (A-D), corroborating the SEC interaction results. (**G**) Quantitation of RNA binding of three independent experiment and plotted against protein concentration. Error bars represent standard error.

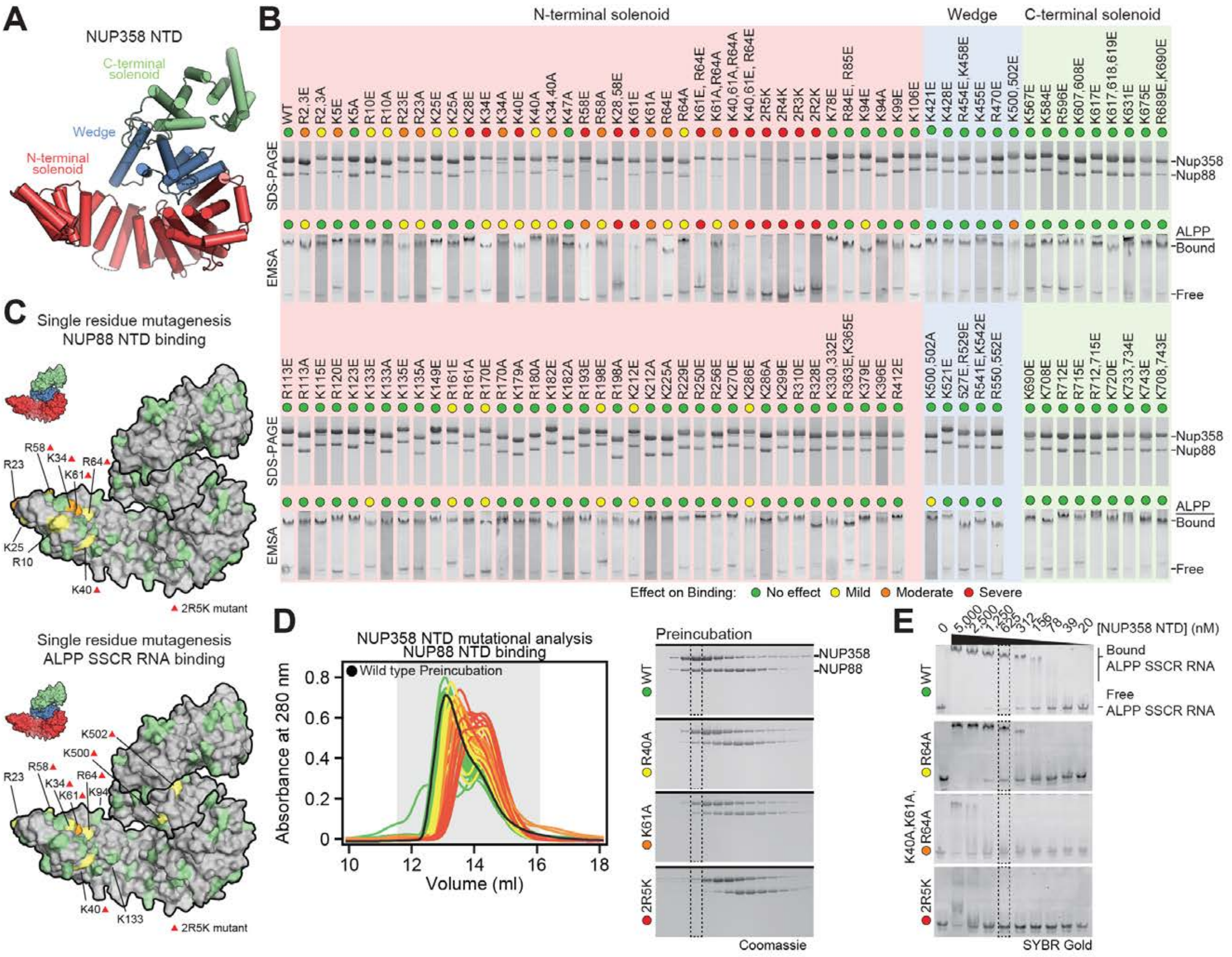
A
NUP358 NTD
C-terminal solenoid
Wedge
N-terminal solenoid
B
N-terminal solenoid
Wedge
C-terminal solenoid
WT R2,3E R2,3A K5E K5A R10E R10A R23E R23A K25E K25A K28E K34E K34A K40E K40A K34,40A K47A R58E R58A K28,58E K61E K61A R64E R64A K61E, R64E K61A,R64A K40,61A,R64A K40,61E, R64E 2R5K 2R4K 2R3K 2R2K K78E R84E, R85E K94E K94A K99E K106E
K421E K428E R454E,K458E K455E R470E K500,502E
K567E K584E R596E K607,608E K617E K617,618,619E K631E K675E R689E,K690E
SDS-PAGE
EMSA
Nup358
Nup88
ALPP
Bound
Free
R113E R113A K115E R120E K123E K133E K133A K135E K135A K149E R161E R161A R170E R170A K179A R180A K182E K182A R193E R198E R198A K212E K212A K225A R229E R250E R256E K270E K286E K286A K299E R310E R328E K330,332E R363E,K365E K379E K396E R412E
K500,502A K521E 527E, R529E R541E,K542E R550,552E
K690E K708E R712E R715E R712,715E K720E K733,734E K743E K708,743E
Effect on Binding: No effect Mild Moderate Severe
C
Single residue mutagenesis
NUP88 NTD binding
R58 K34 R64 R23 K61 K25 R10 K40
2R5K mutant
Single residue mutagenesis
ALPP SSCR RNA binding
K502 K500 R23 R58 R64 K34 K94 K61 K40 K133
2R5K mutant
D
NUP358 NTD mutational analysis
NUP88 NTD binding
Wild type Preincubation
Absorbance at 280 nm
0 0.2 0.4 0.6 0.8
10 12 14 16 18
Volume (ml)
Preincubation
WT
R40A
K61A
2R5K
NUP358
NUP88
Coomassie
E
0 5,000 2,500 1,250 625 312 156 78 39 20 [NUP358 NTD] (nM)
Bound
ALPP SSCR RNA
Free
ALPP SSCR RNA
WT
R64A
K40A,K61A, R64A
2R5K
SYBR Gold

**Fig. S49. Identification of NUP88$^{NTD}$ and ALPP$^{SSCR}$ RNA binding sites on the NUP358$^{NTD}$ surface.** (**A**) Cartoon representation of NUP358$^{NTD}$. (**B**) 106 NUP358$^{NTD}$ surface mutants were tested for their interaction with NUP88$^{NTD}$ and ALPP$^{SSCR}$ RNA by SEC and EMSA, respectively. SDS-PAGE gel strips from each NUP358$^{NTD}$ mutant, corresponding to the peak fraction from the wildtype SEC interaction, were visualized by Coomassie brilliant blue staining. EMSA gel strips for each NUP358$^{NTD}$ mutant at 625 nM concentration, were visualized by SYBR Gold staining. The colored dots indicate the effect on binding of the mutations. (**C**) Surface representations of NUP358$^{NTD}$ with saturating mutagenesis residues colored according to their effect on NUP88$^{NTD}$ and ALPP$^{SSCR}$ RNA binding. Red triangles indicate residues of the 2R5K combination mutant, which abolishes the interactions with both NUP88$^{NTD}$ and ALPP$^{SSCR}$ RNA. (**D**) SEC profiles of NUP88$^{NTD}$ interactions with NUP358$^{NTD}$ mutants are overlayed and colored according to their effect on NUP88$^{NTD}$ binding. Gray boxes indicate fractions resolved on SDS-PAGE gels for representative interactions in each category of binding. (**E**) Full EMSA titrations are shown for representative interactions in each category of binding. Dashed lines indicate SDS-PAGE and EMSA lanes shown in panel (B). Combination mutations are referred to as 2R5K (K34A, K40A, R58A, K61A, R64A, K500A, and K502A), 2R4K (K40A, R58A, K61A, R64A, K500A, and K502A), 2R3K (K34A, K40A, R58A, K61A, and R64A), and 2R2K (K40A, R58A, K61A, and R64A).

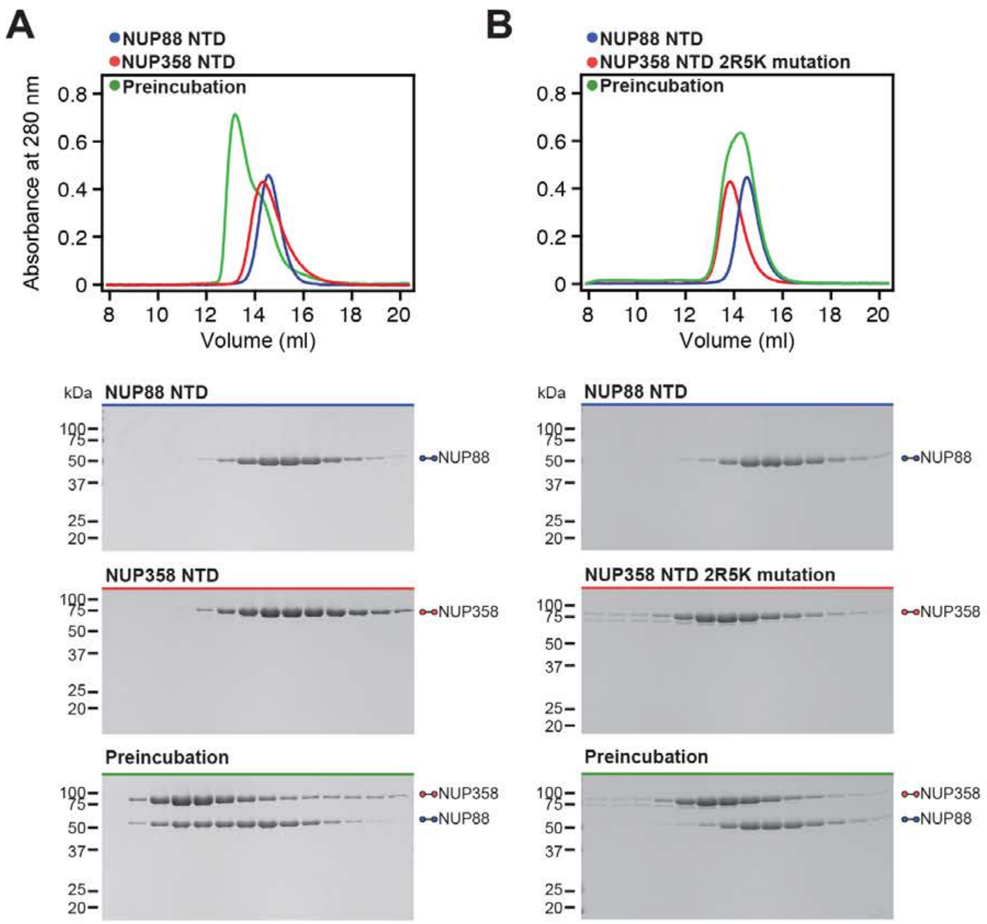
A
NUP88 NTD
NUP358 NTD
Preincubation
Absorbance at 280 nm
0.8
0.6
0.4
0.2
0
8
10
12
14
16
18
20
Volume (ml)
kDa
NUP88 NTD
100
75
50
37
25
20
NUP88
NUP358 NTD
NUP358
Preincubation
NUP358
NUP88
B
NUP88 NTD
NUP358 NTD 2R5K mutation
Preincubation
Volume (ml)
kDa
NUP88 NTD
NUP88
NUP358 NTD 2R5K mutation
NUP358
Preincubation
NUP358
NUP88

**Fig. S50. NUP358$^{NTD}$ 2R5K combination mutant abolishes NUP88$^{NTD}$ binding.** (**A**, **B**) SEC interaction analysis of NUP88$^{NTD}$ with (**A**) NUP358$^{NTD}$ and (**B**) the NUP358$^{NTD}$ 2R5K combination mutant. The 2R5K combination mutation abolishes the NUP358$^{NTD}$-NUP88$^{NTD}$ interaction. SEC profiles of NUP358$^{NTD}$ variants (blue) and NUP88$^{NTD}$ (red) are shown individually and after their preincubation (green). SEC profiles were obtained using a Superdex 200 10/300 GL column. Gray boxes indicate fractions resolved on SDS-PAGE gels and visualized by Coomassie brilliant blue staining.

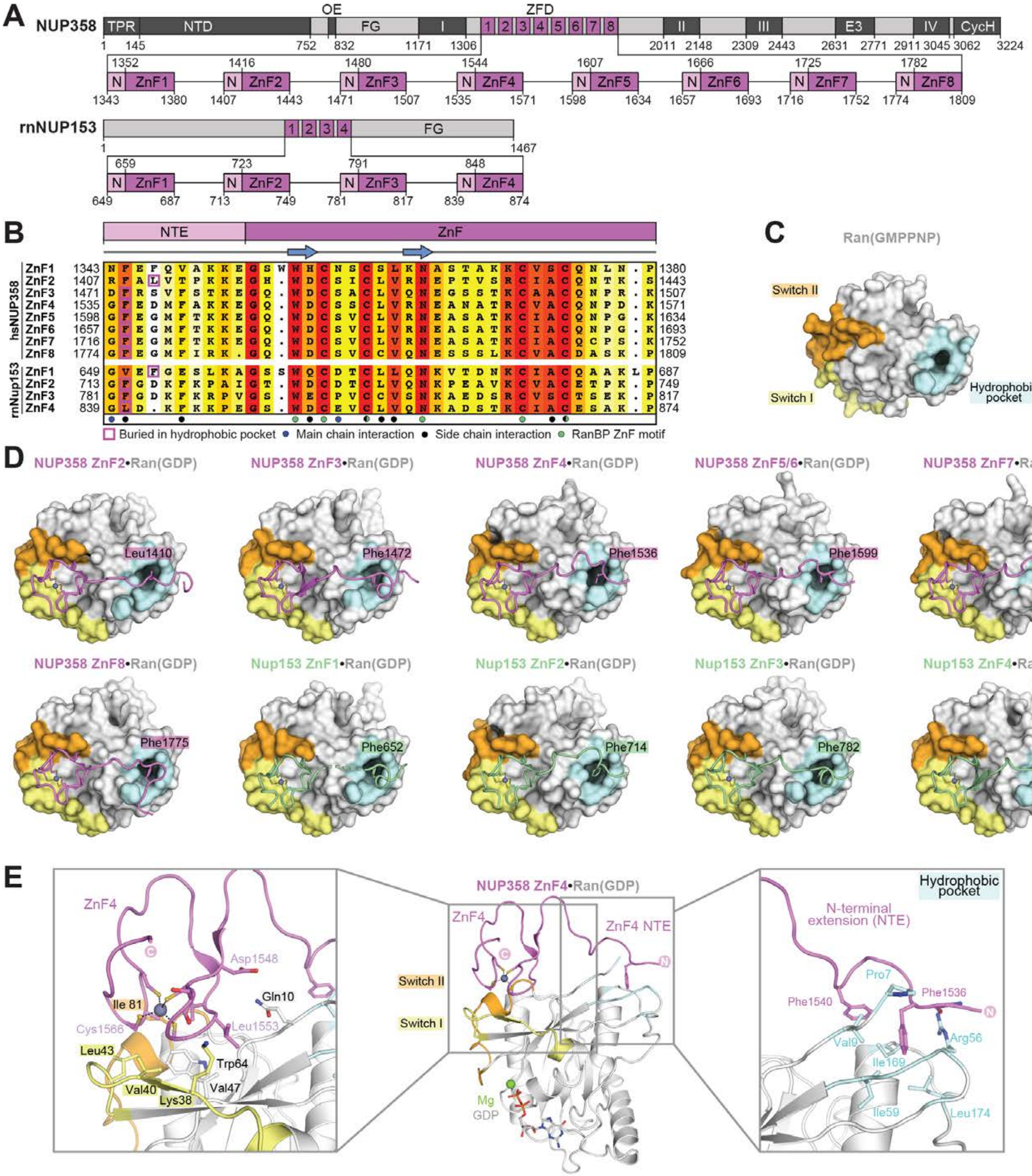

A
NUP358
TPR
NTD
OE
FG
I
ZFD
1 2 3 4 5 6 7 8
II
III
E3
IV
CycH
1 145 752 832 1171 1306 2011 2148 2309 2443 2631 2771 2911 3045 3062 3224
1352 1416 1480 1544 1607 1666 1725 1782
N ZnF1 N ZnF2 N ZnF3 N ZnF4 N ZnF5 N ZnF6 N ZnF7 N ZnF8
1343 1380 1407 1443 1471 1507 1535 1571 1598 1634 1657 1693 1716 1752 1774 1809
rnNUP153
1 2 3 4
FG
1 1467
659 723 791 848
N ZnF1 N ZnF2 N ZnF3 N ZnF4
649 687 713 749 781 817 839 874
B
NTE
ZnF
hsNUP358
ZnF1 ZnF2 ZnF3 ZnF4 ZnF5 ZnF6 ZnF7 ZnF8
rnNup153
ZnF1 ZnF2 ZnF3 ZnF4
Buried in hydrophobic pocket
Main chain interaction
Side chain interaction
RanBP ZnF motif
C
Ran(GMPPNP)
Switch II
Switch I
Hydrophobic pocket
D
NUP358 ZnF2•Ran(GDP)
Leu1410
NUP358 ZnF3•Ran(GDP)
Phe1472
NUP358 ZnF4•Ran(GDP)
Phe1536
NUP358 ZnF5/6•Ran(GDP)
Phe1599
NUP358 ZnF7•Ran(GDP)
Phe1717
NUP358 ZnF8•Ran(GDP)
Phe1775
Nup153 ZnF1•Ran(GDP)
Phe652
Nup153 ZnF2•Ran(GDP)
Phe714
Nup153 ZnF3•Ran(GDP)
Phe782
Nup153 ZnF4•Ran(GDP)
Leu840
E
ZnF4
Asp1548
Gln10
Ile 81
Cys1566
Leu1553
Leu43
Trp64
Val40
Val47
Lys38
NUP358 ZnF4•Ran(GDP)
ZnF4 NTE
Switch II
Switch I
Mg
GDP
Hydrophobic pocket
N-terminal extension (NTE)
Pro7
Phe1540
Phe1536
Val9
Arg56
Ile169
Ile59
Leu174

**Fig. S51. Crystal structures of NUP358$^{ZnF}$•Ran(GDP) and NUP153$^{ZnF}$•Ran(GDP) complexes.** (**A**) Domain structures of NUP358 and NUP153 drawn as horizontal boxes with residue numbers indicated. (**B**) Sequence alignment of the eight NUP358 and four *R. norvegicus* rnNUP153 ZnFs colored according to the BLOSUM62 weighting algorithm from white (less than 40% similarity), through yellow (60% similarity), to red (100% identity). (**C**) Surface representation of Ran(GMPPNP) from the NUP358$^{RanBD-IV}$•Ran crystal structure, highlighting the switch I (yellow), switch II (orange), and the nucleotide-independent hydrophobic pocket (cyan). (**D**) Co-crystal structures of six NUP358$^{ZnF}$•Ran(GDP) and four rnNUP153$^{ZnF}$•Ran(GDP) complexes are shown. The ZnFs and Ran(GDP) are shown in cartoon and surface representation, respectively. In all structures, a hydrophobic residue of each NTE occupies Ran's nucleotide-independent hydrophobic pocket. (**E**) Magnified views of NUP358$^{ZnF4}$•Ran(GDP) illustrating Ran's interactions with the ZnF motif and NTE.

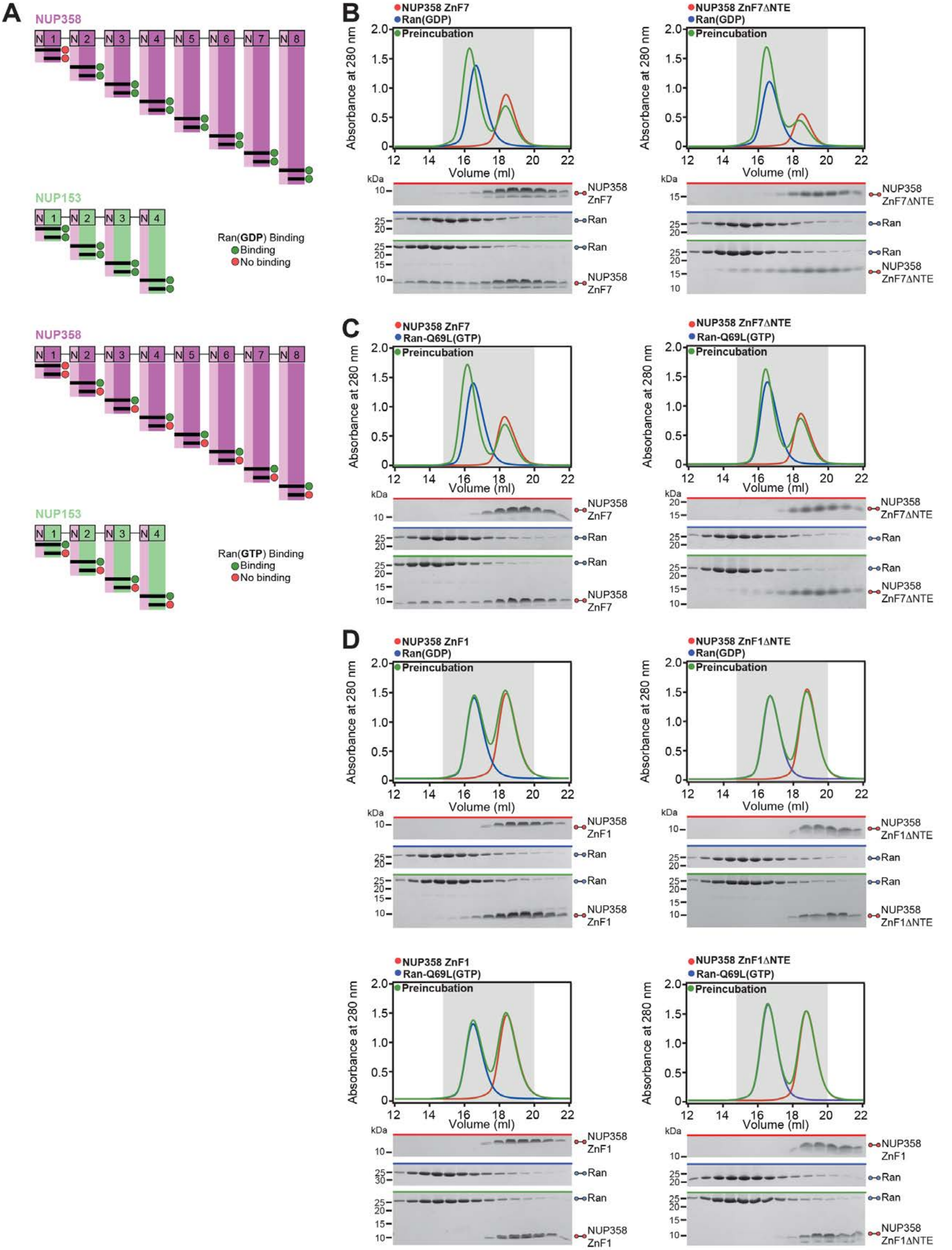

A
NUP358
NUP153
Ran(GDP) Binding
Binding
No binding
Ran(GTP) Binding
B
NUP358 ZnF7
Ran(GDP)
Preincubation
Absorbance at 280 nm
Volume (ml)
kDa
NUP358 ZnF7
Ran
NUP358 ZnF7ΔNTE
C
Ran-Q69L(GTP)
D
NUP358 ZnF1
NUP358 ZnF1ΔNTE

**Fig. S52. Biochemical analysis of the Ran interaction with NUP358/NUP153 ZnF modules.** (**A**) Summary of the Ran(GDP) binding properties of the twelve NUP358/NUP153 ZnF modules. Apart from NUP358$^{ZnF1}$, for which no binding is detected, the remaining eleven ZnF modules bind Ran(GDP) irrespective of the presence of the NTE. (**B**) Representative SEC interaction analysis of RAN(GDP) with NUP358$^{ZnF7}$ in the absence or presence of the NTE. (**C**) Summary of the Ran(GTP) binding properties of the twelve NUP358/NUP153 ZnF modules. Apart from NUP358$^{ZnF1}$, for which no binding is detected, Ran(GTP) binding of the remaining eleven ZnF modules is dependent on the presence of the NTE. (**D**) SEC interaction analyses of NUP358$^{ZnF1}$ with Ran(GDP) and Ran(GTP), showing no binding irrespective of the presence of the NTE.

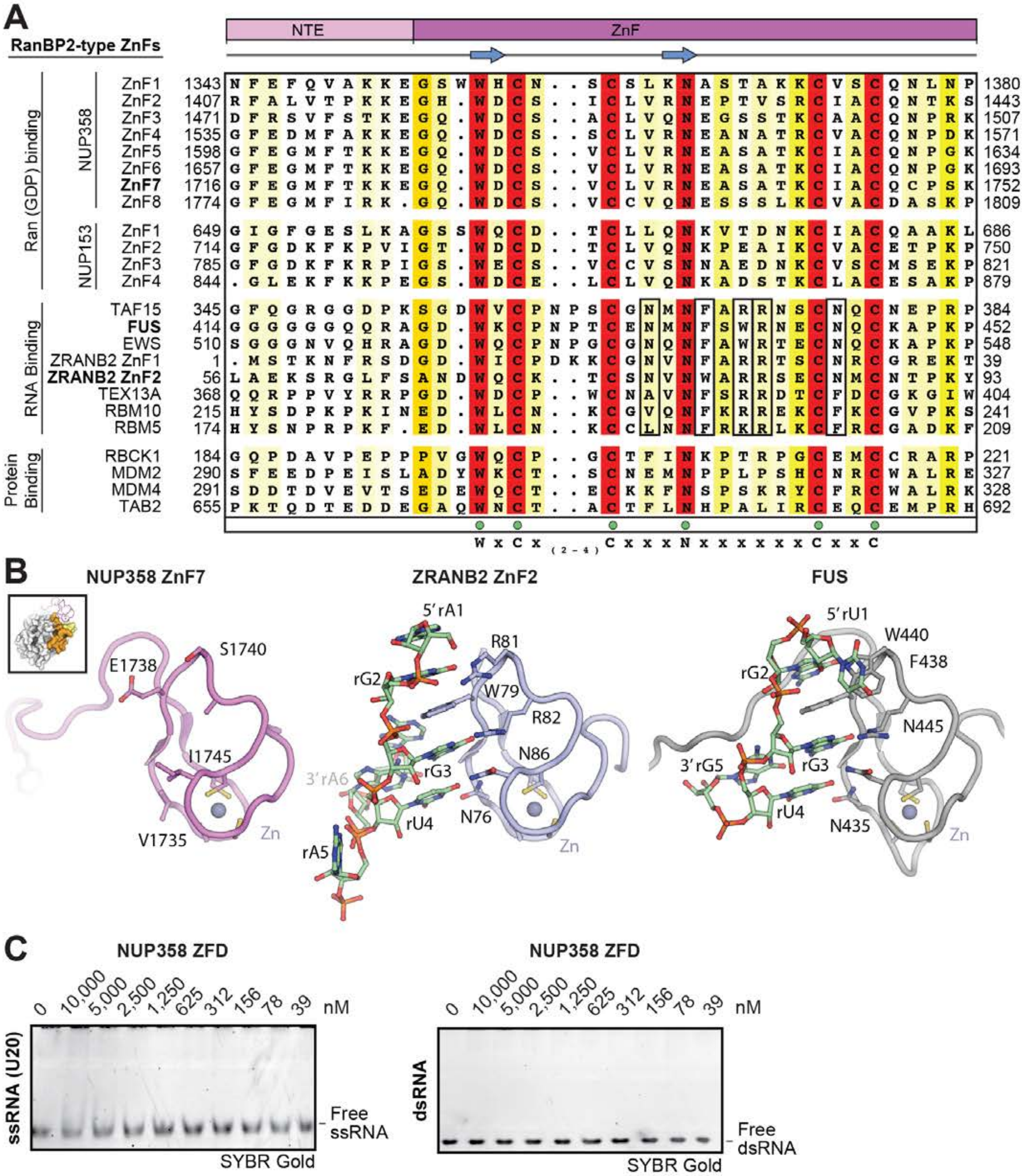
A
RanBP2-type ZnFs
NTE
ZnF
Ran (GDP) binding
NUP358
ZnF1 1343 1380
ZnF2 1407 1443
ZnF3 1471 1507
ZnF4 1535 1571
ZnF5 1598 1634
ZnF6 1657 1693
ZnF7 1716 1752
ZnF8 1774 1809
NUP153
ZnF1 649 686
ZnF2 714 750
ZnF3 785 821
ZnF4 844 879
RNA Binding
TAF15 345 384
FUS 414 452
EWS 510 548
ZRANB2 ZnF1 1 39
ZRANB2 ZnF2 56 93
TEX13A 368 404
RBM10 215 241
RBM5 174 209
Protein Binding
RBCK1 184 221
MDM2 290 327
MDM4 291 328
TAB2 655 692
W x C x (2-4) C x x x N x x x x x x C x x C
B
NUP358 ZnF7
E1738
S1740
I1745
V1735
Zn
ZRANB2 ZnF2
5′rA1
R81
rG2
W79
R82
N86
3′rA6
rG3
rU4
N76
rA5
Zn
FUS
5′rU1
W440
F438
rG2
N445
3′rG5
rG3
rU4
N435
Zn
C
NUP358 ZFD
0 10,000 5,000 2,500 1,250 625 312 156 78 39 nM
ssRNA (U20)
Free ssRNA
SYBR Gold
NUP358 ZFD
0 10,000 5,000 2,500 1,250 625 312 156 78 39 nM
dsRNA
Free dsRNA
SYBR Gold

**Fig. S53. NUP358 ZFD does not bind to single- or double-stranded RNA.** (**A**) Sequence alignment of the twelve NUP358/NUP153 ZnF modules with representative members of the RanBP2 ZnF family with established RNA or protein binding properties, colored according to the BLOSUM62 weighting algorithm from white (less than 40% similarity) through yellow (60% similarity) to red (100% identity). Essential RNA binding residues are highlighted by a black box. RanBP2-type zinc finger motif residues are indicated by green dots. (**B**) Structures of $NUP358^{ZnF7}$, $ZRANB2^{ZnF}$•ssRNA (PDB ID 3G9Y) (*90*), and $FUS^{ZnF}$•ssRNA (PDB ID 6G99) (*91*) are shown, illustrating the absence of residues required for RNA binding. Inset indicates $NUP358^{ZnF7}$ orientation in comparison to Ran(GDP). (**C**) EMSA titration of $NUP358^{ZFD}$ with ssRNA and dsRNA visualized with SYBR Gold stain, demonstrating the absence of RNA binding properties.

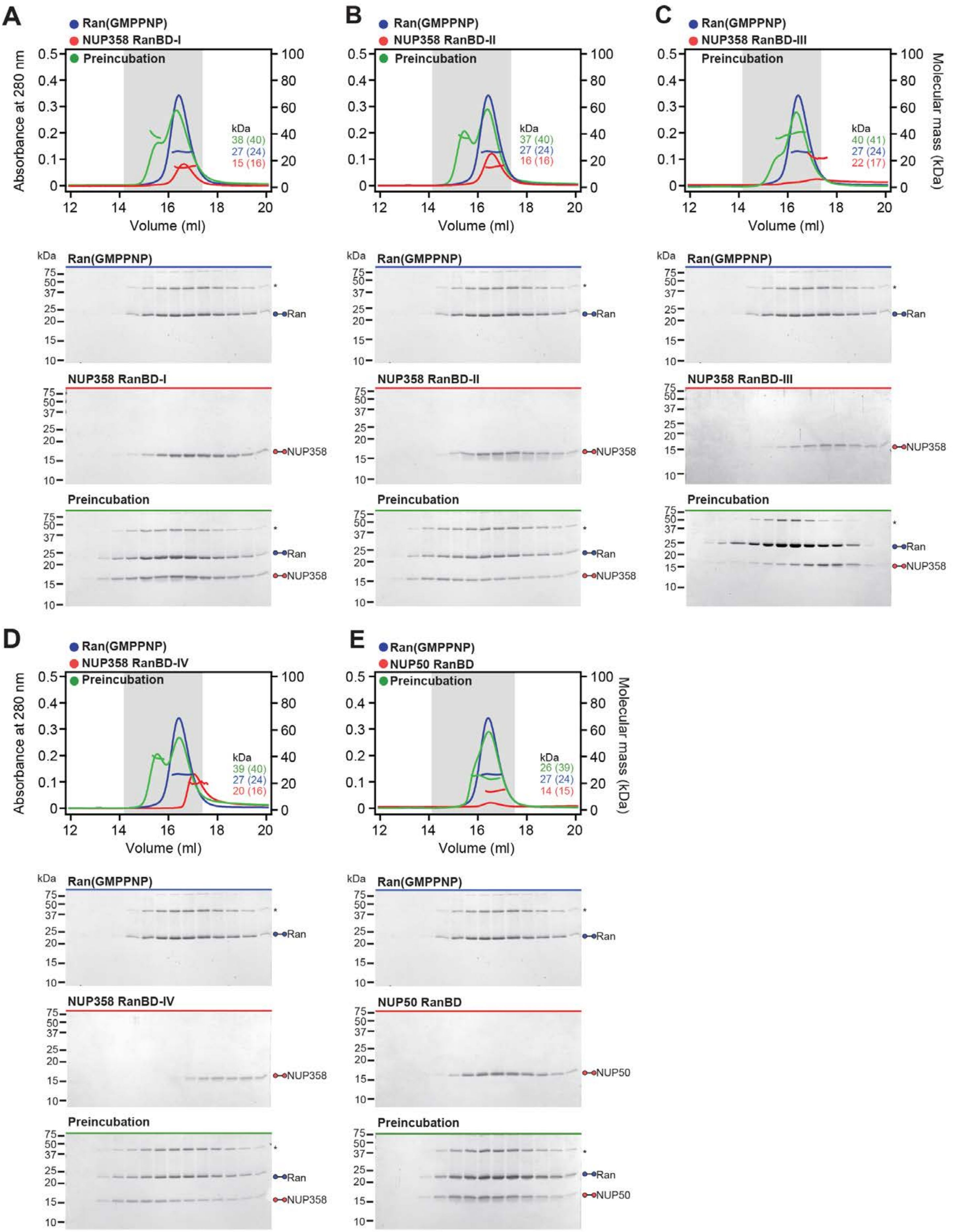

A
Ran(GMPPNP)
NUP358 RanBD-I
Preincubation
kDa
38 (40)
27 (24)
15 (16)
B
NUP358 RanBD-II
37 (40)
27 (24)
16 (16)
C
NUP358 RanBD-III
40 (41)
27 (24)
22 (17)
D
NUP358 RanBD-IV
39 (40)
27 (24)
20 (16)
E
NUP50 RanBD
26 (39)
27 (24)
14 (15)
Absorbance at 280 nm
Molecular mass (kDa)
Volume (ml)
Ran
NUP358
NUP50

**Fig. S54. Biochemical analysis of the Ran interaction with NUP358/NUP50 RanBDs.** (**A**-**E**) SEC-MALS interaction analyses of Ran(GMPPNP) with (A) $NUP358^{RanBD\text{-}I}$, (B) $NUP358^{RanBD\text{-}II}$, (C) $NUP358^{RanBD\text{-}III}$, (D) $NUP358^{RanBD\text{-}IV}$, and (E) $NUP50^{RanBD}$. SEC profiles of RanBDs (blue) and Ran(GMPPNP) red are shown individually and after their preincubation (green). Measured molecular masses are indicated, with the respective theoretical masses shown in parentheses. Gray boxes indicate fractions resolved on SDS-PAGE gels and visualized by Coomassie brilliant blue staining. Asterisks indicate oligomeric Ran species, arising from boiling in SDS-loading dye.

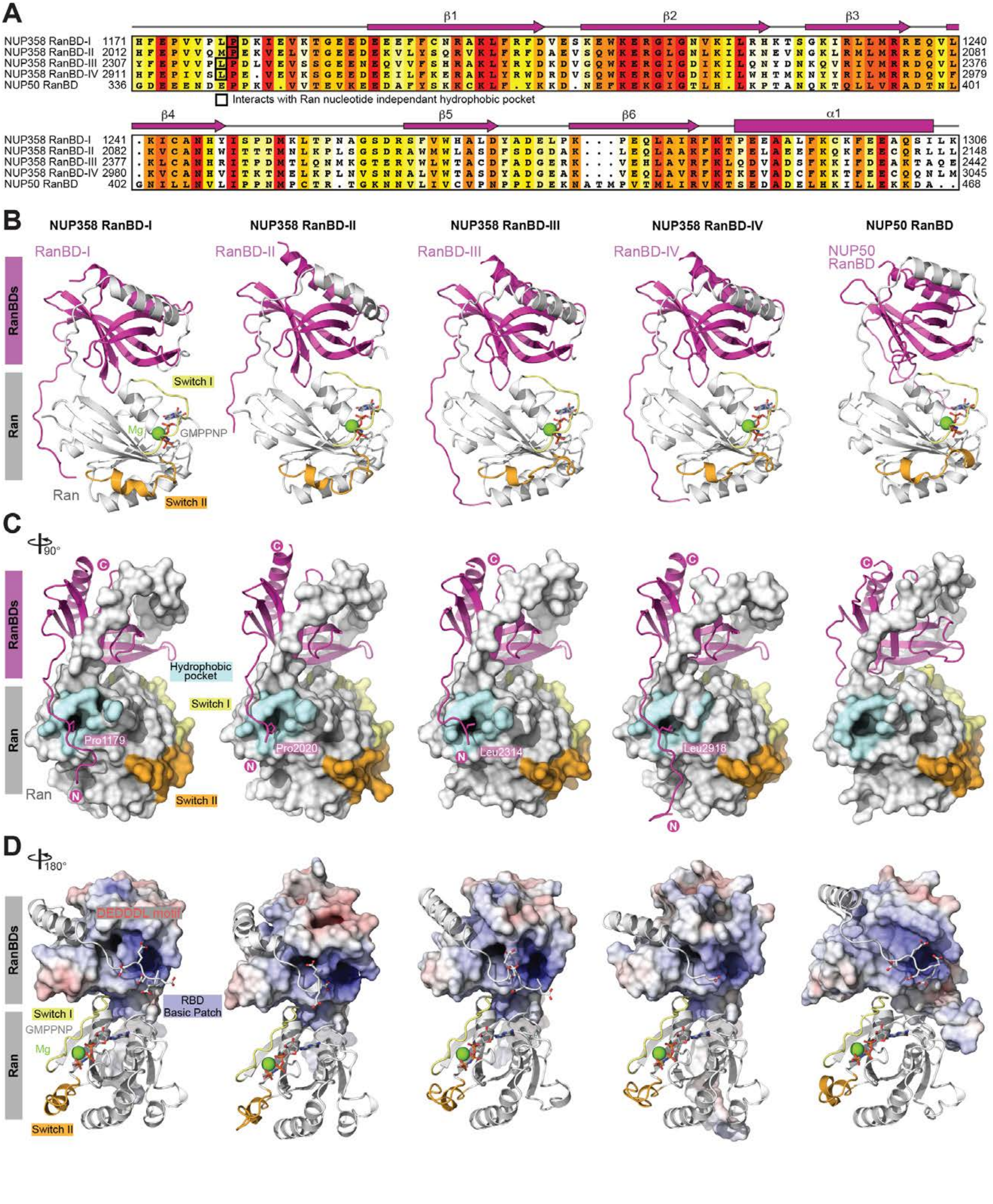

A
β1
β2
β3
NUP358 RanBD-I 1171 HFEPVVPLPDKIEVKTGEEDEEEFFCNRAKLFRFDVESKEWKERGIGNVKILRHKTSGKIRLLMRREQVL 1240
NUP358 RanBD-II 2012 HFEPVVQMPEKVELVTGEEDEKVLYSQRVKLFRFDAEVSQWKERGLGNLKILKNEVNGKLRMLMRREQVL 2081
NUP358 RanBD-III 2307 YFEPVVPLPDLVEVSSGEENEQVVFSHRAKLYRYDKDVGQWKERGIGDIKILQNYDNKQVRIVMRRDQVL 2376
NUP358 RanBD-IV 2911 HFEPIVSLPE.VEVKSGEEDEEILFKERAKLYRWDRDVSQWKERGVGDIKILWHTMKNYYRILMRRDQVF 2979
NUP50 RanBD 336 GDEEENDEPPKV.VVTEVKEEDAFYSKKCKLF.YKKD.NEFKEKGIGTLH.LKPTANQKTQLLVRADTNL 401
Interacts with Ran nucleotide independant hydrophobic pocket
β4
β5
β6
α1
NUP358 RanBD-I 1241 .KICANHYISPDMKLTPNAGSDRSFVWHALDYADELPK...PEQLAIRFKTPEEAALFKCKFEEAQSILK 1306
NUP358 RanBD-II 2082 .KVCANHWITTTMNLKPLSGSDRAWMWLASDFSDGDAK...LEQLAAKFKTPELAEEFKQKFEECQRLLL 2148
NUP358 RanBD-III 2377 .KLCANHRITPDMTLQNMKGTERVWLWTACDFADGERK...VEHLAVRFKLQDVADSFKKIFDEAKTAQE 2442
NUP358 RanBD-IV 2980 .KVCANHVITKTMELKPLNVSNNALVWTASDYADGEAK...VEQLAVRFKTKEVADCFKKTFEECQQNLM 3045
NUP50 RanBD 402 GNILLNVLIPPNMPCTR.TGKNNVLIVCVPNPPIDEKNATMPVTMLIRVKTSEDADELHKILLEKKDA.. 468
B
NUP358 RanBD-I
NUP358 RanBD-II
NUP358 RanBD-III
NUP358 RanBD-IV
NUP50 RanBD
RanBD-I
RanBD-II
RanBD-III
RanBD-IV
NUP50 RanBD
RanBDs
Ran
Switch I
Mg
GMPPNP
Ran
Switch II
C
90°
C
RanBDs
Hydrophobic pocket
Switch I
Ran
Pro1179
Pro2020
Leu2314
Leu2918
N
Ran
Switch II
D
180°
RanBDs
DEDDDL motif
RBD Basic Patch
Switch I
GMPPNP
Mg
Ran
Switch II

**Fig. S55. Crystal structures of NUP358/NUP50$^{RanBD}$•Ran(GMPPNP).** (**A**) Sequence alignment of the NUP358/NUP50 RanBD, colored according to the BLOSUM62 weighting algorithm from white (less than 40% similarity), through yellow (60% similarity), to red (100% identity). The secondary structure is indicated above the sequences as rectangles (a-helices), arrows ($\beta$-strands), and lines (unstructured regions). (**B**) Co-crystal structures of the five NUP358/NUP50$^{RanBD}$•Ran(GMPPNP) complexes shown in cartoon representation. (**C**) 90º-rotated views of all five complexes illustrating the location of Ran's nucleotide-independent hydrophobic pocket that accommodates a hydrophobic residue of all four NUP358$^{RanBD}$, an interaction not observed in the NUP50$^{RanBD}$ complex. (**D**) 180º-rotated views of all five complexes highlighting the interaction of the DEDDDL motif of Ran with the basic patch found on each RanBD. The RanBD surface representation is colored according to electrostatic potential from red (-10 $k_BT/e$) to blue (+10 $k_BT/e$).

**A**

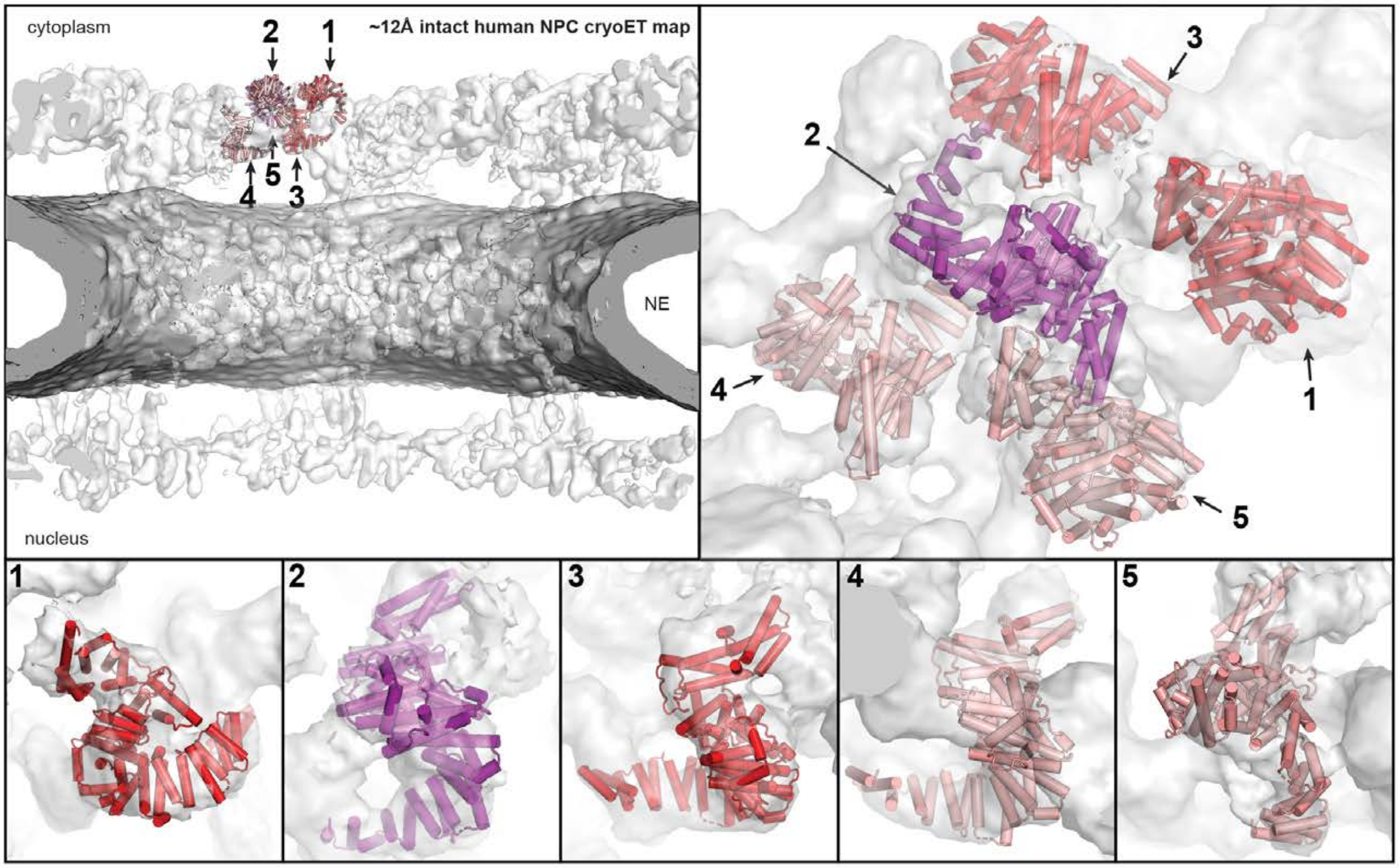

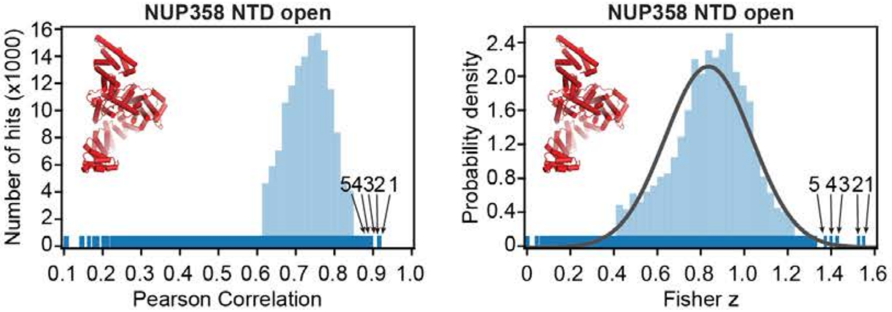

**Cryo-ET map fitting statistics**

| solution rank | Pearson correlation | Fisher z | p-value |
|---|---|---|---|
| 1 | 0.913 | 1.54 | 1.7 e-4 |
| 2 | 0.909 | 1.52 | 2.6 e-4 |
| 3 | 0.892 | 1.43 | 1.5 e-3 |
| 4 | 0.891 | 1.43 | 1.5 e-3 |
| 5 | 0.890 | 1.40 | 2.3 e-3 |

**B**

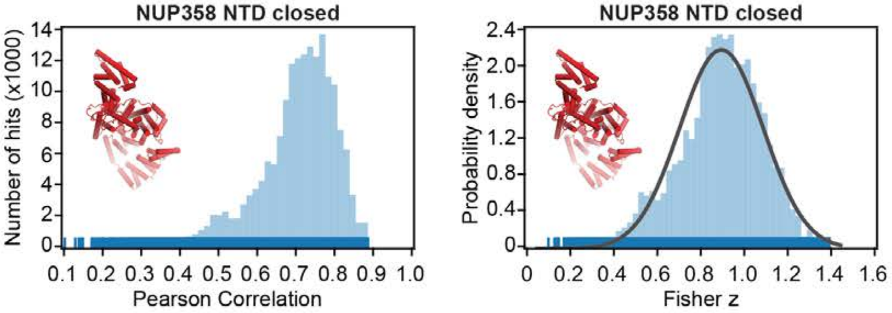

**Cryo-ET map fitting statistics**

| solution rank | Pearson correlation | Fisher z | p-value |
|---|---|---|---|
| 1 | 0.882 | 1.38 | 4.9 e-3 |
| 2 | 0.880 | 1.38 | 5.6 e-3 |
| 3 | 0.878 | 1.37 | 6.2 e-3 |
| 4 | 0.877 | 1.36 | 6.7 e-3 |
| 5 | 0.875 | 1.35 | 7.6 e-3 |

**Fig. S56. Docking of the NUP358$^{NTD}$ structure into an ~12Å cryo-ET map of the intact human NPC.** (**A**) The open conformation NUP358$^{NTD}$ crystal structure was quantitatively docked into the ~12 Å sub-tomogram averaged cryo-ET map of the intact human NPC (*46*). Placed unique solutions are shown in cartoon representation with their rank indicated, whereas the cryo-ET map is rendered as a transparent isosurface. Solutions are viewed from a central cross-section of the intact NPC (*left*), from a closeup view from above the cytoplasmic face (*right*), or individually (*bottom*). Isosurfaces of the nuclear envelope and the NPC are colored dark and light gray, respectively. Rug plots (blue) and histograms (light blue) of Pearson correlation scores and derived Fisher z scores, fit with a normalized Gaussian curve (black), from a global search with 1 million random initial placements, are shown. Arrows and corresponding numbers indicate the rank of accepted solutions. A table summary of the accepted solutions fitting statistics, along with one-tailed p-values calculated from the Fisher z score distribution, is shown. (**B**) The same analysis as in (A) was applied to the closed conformation NUP358$^{NTD}$ crystal structure. Unlike with the open conformation, the docking solutions identified for the closed conformation of NUP358$^{NTD}$ crystal structure did not segregate to high confidence.

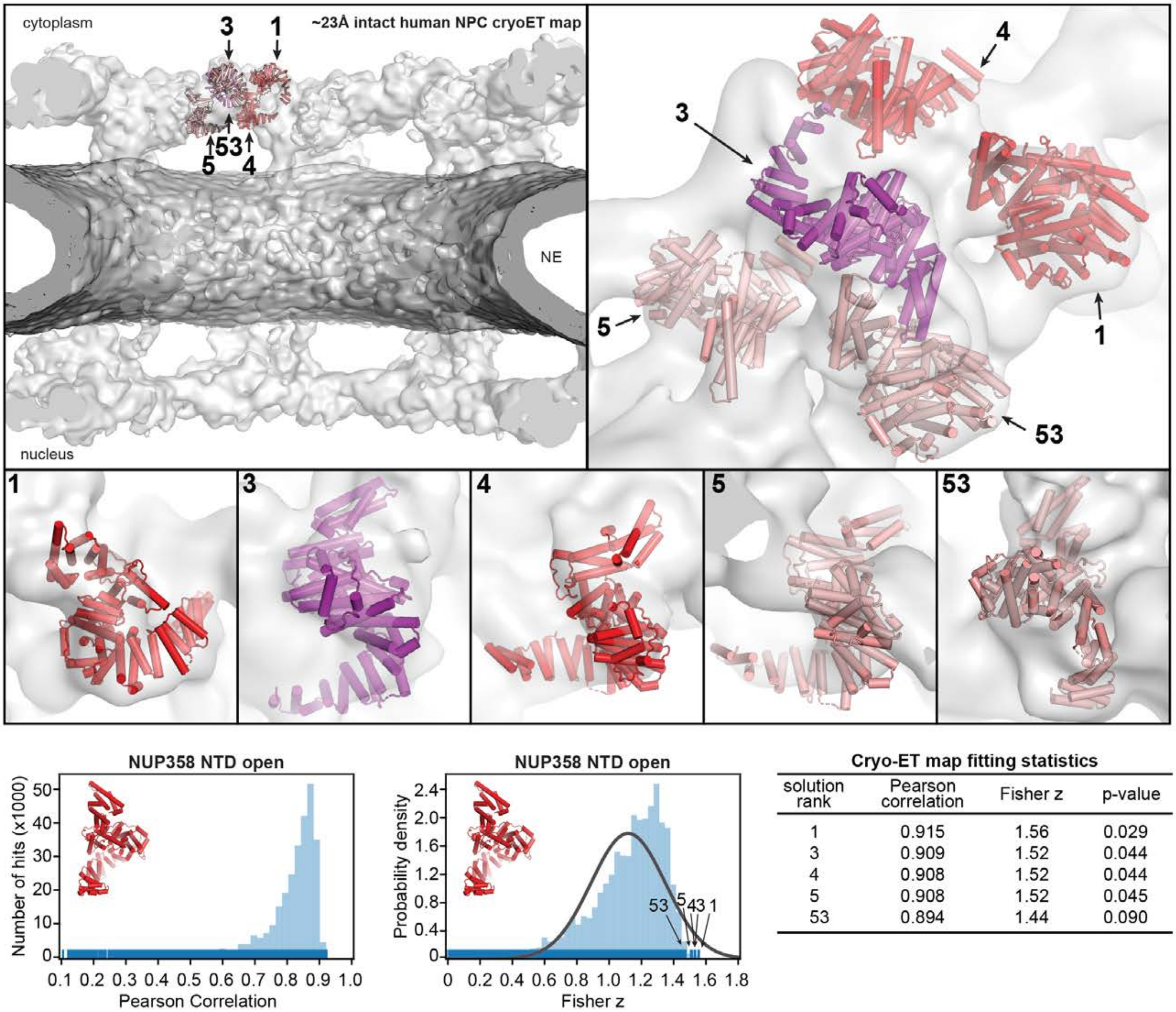

**Cryo-ET map fitting statistics**

| solution rank | Pearson correlation | Fisher z | p-value |
|---|---|---|---|
| 1 | 0.915 | 1.56 | 0.029 |
| 3 | 0.909 | 1.52 | 0.044 |
| 4 | 0.908 | 1.52 | 0.044 |
| 5 | 0.908 | 1.52 | 0.045 |
| 53 | 0.894 | 1.44 | 0.090 |

**Fig. S57. Docking of the NUP358$^{NTD}$ structure into an ~23Å cryo-ET map of the intact human NPC**. The open conformation NUP358$^{NTD}$ crystal structure was quantitatively docked into the ~23 Å sub-tomogram averaged cryo-ET map of the intact human NPC (*44*). Placed unique solutions are shown in cartoon representation with their rank indicated, whereas the cryo-ET map is rendered as a transparent isosurface. Solutions are viewed from a central cross-section of the intact NPC (*left*), from a closeup view from above the cytoplasmic face (*right*), or individually (*bottom*). Isosurfaces of the nuclear envelope and the NPC are colored dark and light gray, respectively. Rug plots (blue) and histograms (light blue) of Pearson correlation scores and derived Fisher z scores, fit with a normalized Gaussian curve (black), from a global search with 1 million random initial placements, are shown. Arrows and corresponding numbers indicate the rank of accepted solutions. A table summary of the accepted solutions fitting statistics, along with one-tailed p-values calculated from the Fisher z score distribution, is shown.

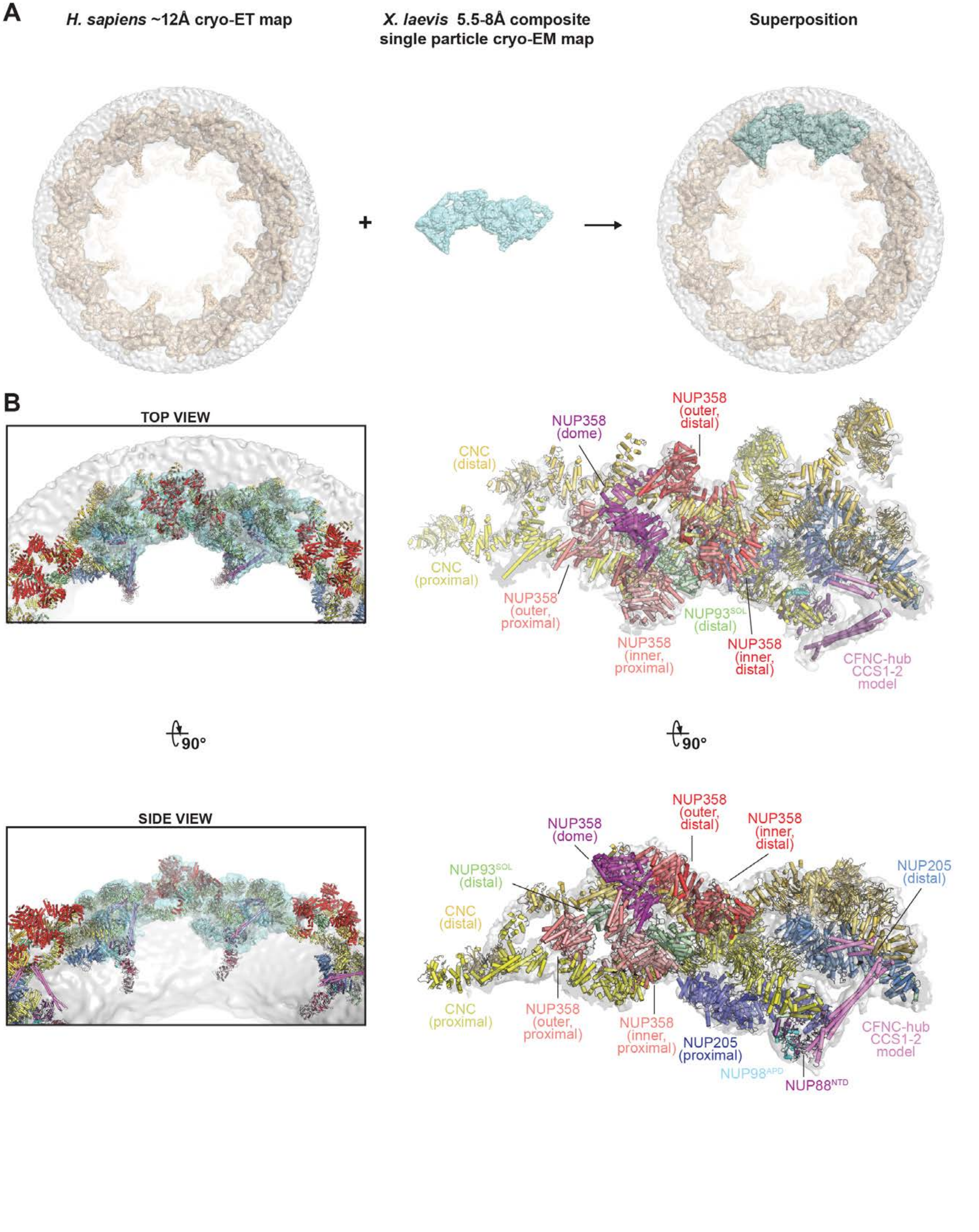

A
H. sapiens ~12Å cryo-ET map
X. laevis 5.5-8Å composite single particle cryo-EM map
Superposition
+
B
TOP VIEW
SIDE VIEW
90°
90°
NUP358 (outer, distal)
NUP358 (dome)
CNC (distal)
CNC (proximal)
NUP358 (outer, proximal)
NUP93SOL (distal)
NUP358 (inner, proximal)
NUP358 (inner, distal)
CFNC-hub CCS1-2 model
NUP358 (outer, distal)
NUP358 (inner, distal)
NUP358 (dome)
NUP93SOL (distal)
CNC (distal)
NUP205 (distal)
CNC (proximal)
NUP358 (outer, proximal)
NUP358 (inner, proximal)
NUP205 (proximal)
CFNC-hub CCS1-2 model
NUP98APD
NUP88NTD

**Fig. S58. Docking of the CF nups into the anisotropic composite single particle cryo-EM map of the *X. laevis* cytoplasmic face protomer.** (**A**) An anisotropic ~8 Å composite single particle cryo-EM map of the *X. laevis* cytoplasmic face protomer (light cyan; EMD-0909) (*45*) was superposed to the ~12 Å cryo-ET map of the intact human NPC (wheat; EMD-XXXX) (*46*). Maps and nuclear envelope (gray) are rendered as isosurfaces. (**B**) The superposition of the two maps placed the docked cytoplasmic outer ring and cytoplasmic face asymmetric nup structures, shown in cartoon representation, into the composite single particle cryo-EM map of the *X. laevis* cytoplasmic face protomer (*left*). The fit of the structures was subsequently locally optimized against the fully explained *X. laevis* cryo-EM map (*right*).

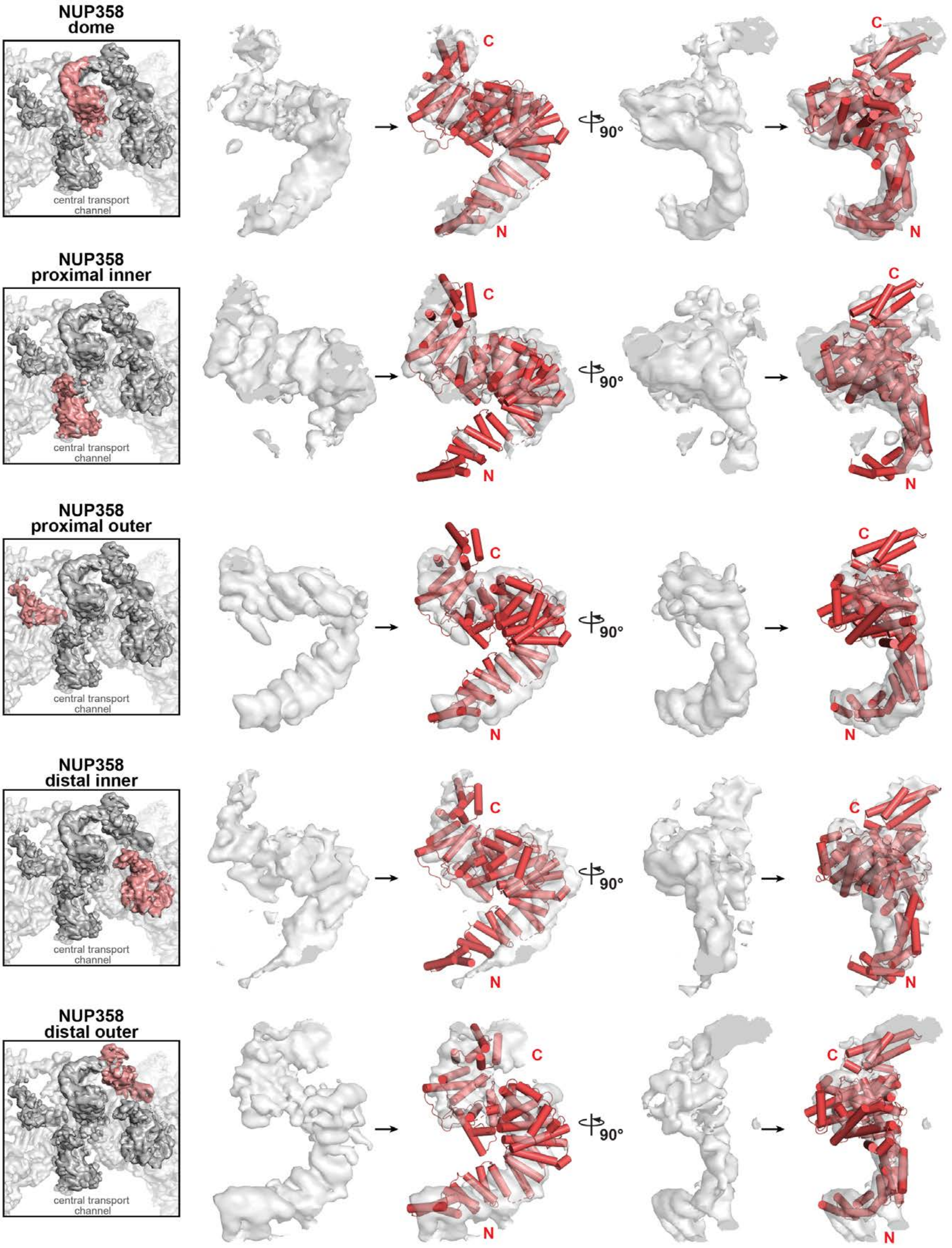
NUP358
dome
central transport
channel
C
N
90°
C
N
NUP358
proximal inner
central transport
channel
C
N
90°
C
N
NUP358
proximal outer
central transport
channel
C
N
90°
C
N
NUP358
distal inner
central transport
channel
C
N
90°
C
N
NUP358
distal outer
central transport
channel
C
N
90°
C
N

**Fig. S59. Identifying five NUP358$^{NTD}$ copies in the *X. laevis* cytoplasmic face protomer.** The NUP358$^{NTD}$ open conformation crystal structure was globally placed by superposition of the ~12 Å cryo-ET map of the intact human NPC and locally rigid body refined at five positions (*left*) in the ~7 Å region of an anisotropic composite single particle cryo-EM map of the *X. laevis* cytoplasmic face protomer (light cyan; EMD-0909) (*45*). Two orthogonal views (*middle* and *right*) of carved out cryo-EM density rendered as transparent isosurfaces are shown superposed to NUP358$^{NTD}$ structures in cartoon representation.

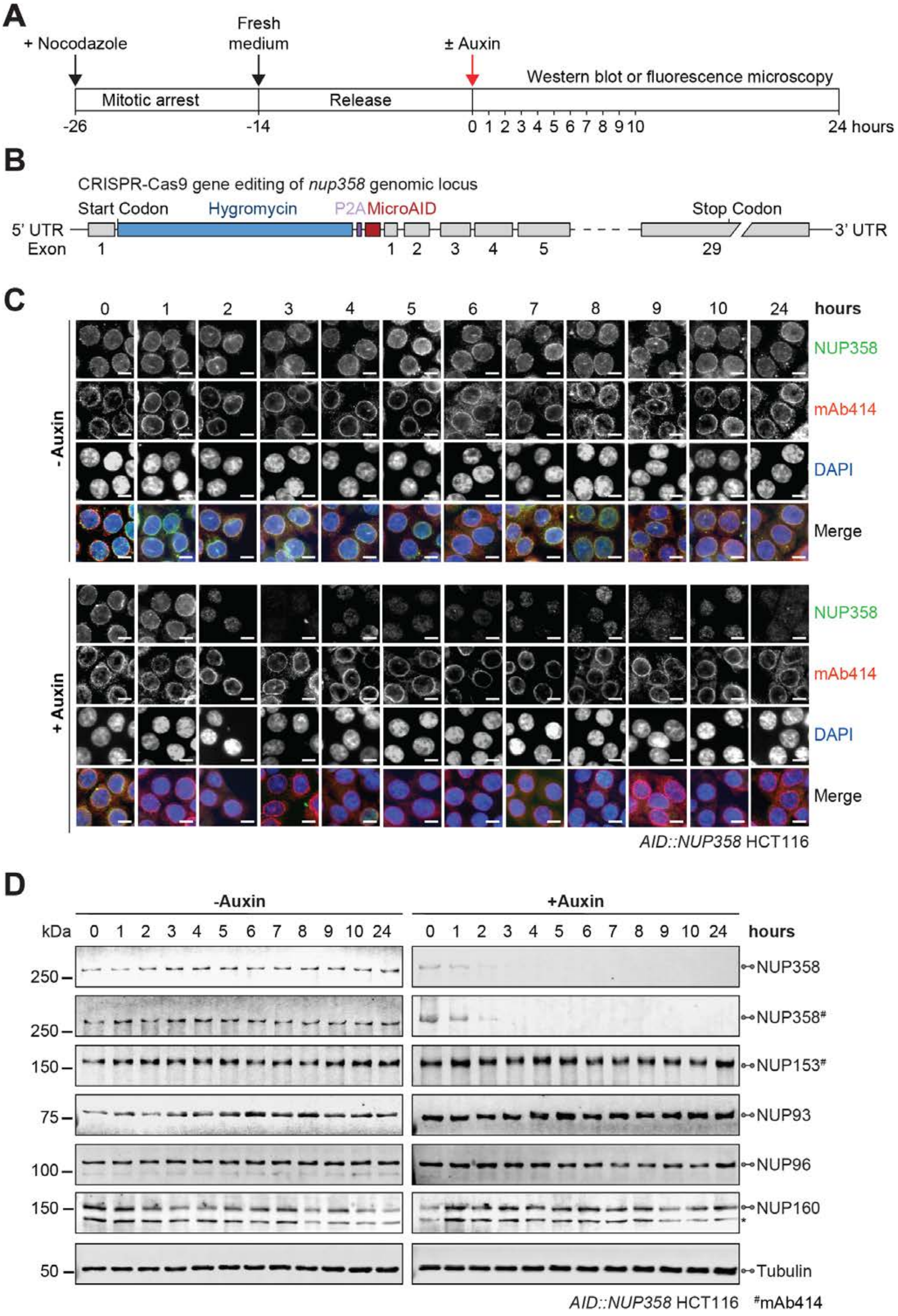

A
+ Nocodazole
Fresh medium
± Auxin
Mitotic arrest
Release
Western blot or fluorescence microscopy
-26
-14
0 1 2 3 4 5 6 7 8 9 10
24 hours
B
CRISPR-Cas9 gene editing of nup358 genomic locus
Start Codon
Hygromycin
P2A
MicroAID
Stop Codon
5' UTR
3' UTR
Exon
1
1 2 3 4 5
29
C
0 1 2 3 4 5 6 7 8 9 10 24 hours
- Auxin
+ Auxin
NUP358
mAb414
DAPI
Merge
AID::NUP358 HCT116
D
-Auxin
+Auxin
kDa
0 1 2 3 4 5 6 7 8 9 10 24
hours
250
150
75
100
50
NUP358
NUP358#
NUP153#
NUP93
NUP96
NUP160
*
Tubulin
AID::NUP358 HCT116
#mAb414

**Fig. S60. Validation of auxin-induced NUP358 degradation in *AID*::*NUP358* HCT116 cells.** (**A**) Experimental timeline. (**B**) Schematic representation of CRISPR/Cas9-mediated genomic tagging of *nup358* alleles. (**C**) Time-resolved subcellular localization analysis of AID-tagged NUP358 by immunofluorescence microscopy upon auxin treatment. The mAb414 antibody (red) and DAPI (blue) staining were used as a reference for the nuclear envelope and nucleus, respectively. Scale bars are 10 μm. (**D**) Time-resolved western blot analysis of AID-tagged NUP358 expression levels upon auxin treatment. The specificity of auxin induced NUP358 depletion was assessed by the indicated antibodies. Asterisks indicate a nonspecific band detected by the anti-NUP160 antibody. Control cells were not treated with auxin.

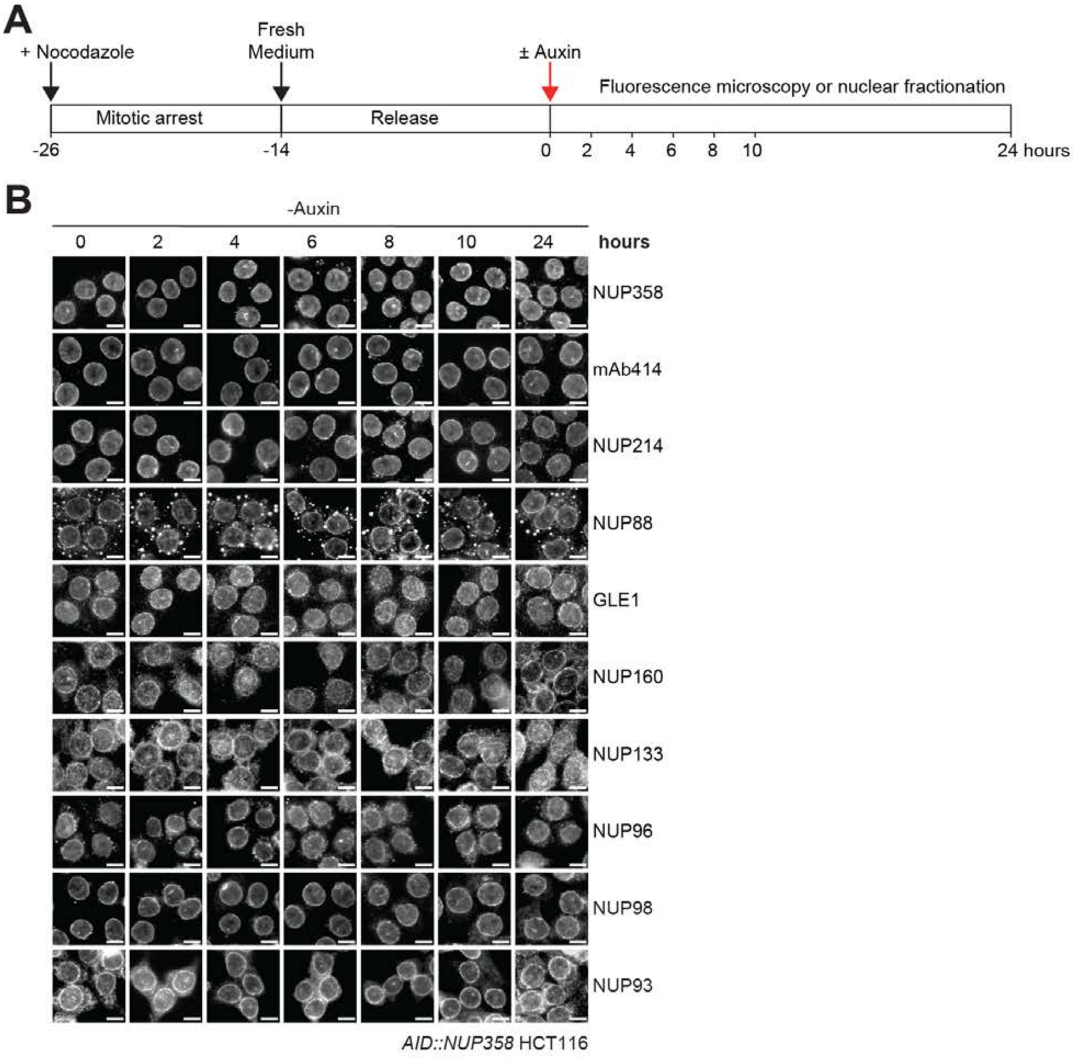
A
+ Nocodazole
Fresh Medium
± Auxin
Fluorescence microscopy or nuclear fractionation
Mitotic arrest
Release
-26
-14
0
2
4
6
8
10
24 hours
B
-Auxin
0
2
4
6
8
10
24
hours
NUP358
mAb414
NUP214
NUP88
GLE1
NUP160
NUP133
NUP96
NUP98
NUP93
AID::NUP358 HCT116

**Fig. S61. Time-resolved localization analysis of endogenous nups in *AID*::*NUP358* HCT116 cells.** (**A**) Experimental timeline. (**B**) Time-resolved subcellular localization analysis by immunofluorescence microscopy of endogenous nups in synchronized *AID*::*NUP358* HCT116 cells without auxin treatment, as a control for Fig. 6A. Scale bars are 10 μm.

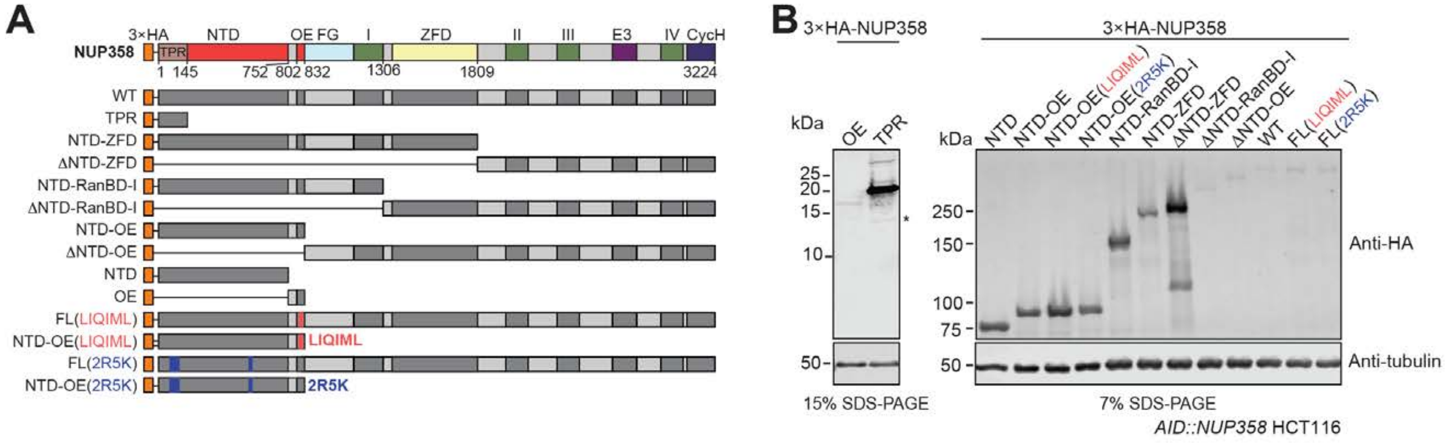
A
3×HA
NTD
OE
FG
I
ZFD
II
III
E3
IV
CycH
NUP358
TPR
1
145
752
802
832
1306
1809
3224
WT
TPR
NTD-ZFD
ΔNTD-ZFD
NTD-RanBD-I
ΔNTD-RanBD-I
NTD-OE
ΔNTD-OE
NTD
OE
FL(LIQIML)
NTD-OE(LIQIML)
LIQIML
FL(2R5K)
NTD-OE(2R5K)
2R5K
B
3×HA-NUP358
kDa
OE
TPR
25
20
15
10
50
15% SDS-PAGE
3×HA-NUP358
kDa
NTD
NTD-OE
NTD-OE(LIQIML)
NTD-OE(2R5K)
NTD-RanBD-I
NTD-ZFD
ΔNTD-ZFD
ΔNTD-RanBD-I
ΔNTD-OE
WT
FL(LIQIML)
FL(2R5K)
250
150
100
75
50
Anti-HA
Anti-tubulin
7% SDS-PAGE
AID::NUP358 HCT116

**Fig. S62. Expression levels of 3×HA-tagged NUP358 variants in *AID*::*NUP358* HCT116 cells.** (**A**) Schematics indicating the domain boundaries of the transfected N-terminally 3×HA-tagged NUP358 variants. (**B**) Western blot expression analysis of the 3×HA-NUP358 variants in *AID*::*NUP358* HCT116 cells (corresponding to localization in Fig. 6B).

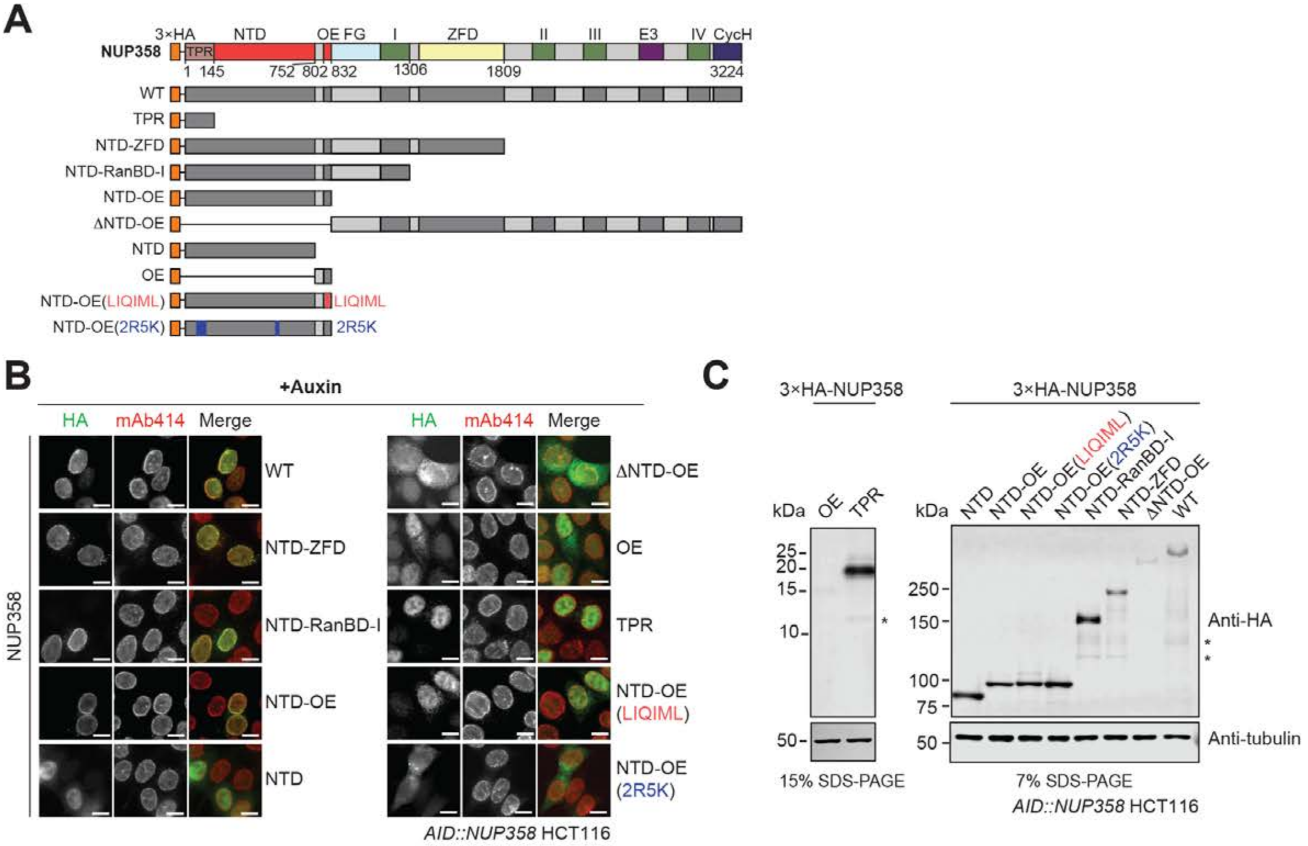
A
3×HA
NTD
OE
FG
I
ZFD
II
III
E3
IV
CycH
NUP358
TPR
1
145
752
802
832
1306
1809
3224
WT
TPR
NTD-ZFD
NTD-RanBD-I
NTD-OE
ΔNTD-OE
NTD
OE
NTD-OE(LIQIML)
LIQIML
NTD-OE(2R5K)
2R5K
B
+Auxin
HA
mAb414
Merge
NUP358
WT
NTD-ZFD
NTD-RanBD-I
NTD-OE
NTD
ΔNTD-OE
OE
TPR
NTD-OE (LIQIML)
NTD-OE (2R5K)
AID::NUP358 HCT116
C
3×HA-NUP358
3×HA-NUP358
kDa
OE
TPR
25
20
15
10
50
15% SDS-PAGE
NTD
NTD-OE
NTD-OE(LIQIML)
NTD-OE(2R5K)
NTD-RanBD-I
NTD-ZFD
ΔNTD-OE
WT
250
150
100
75
50
Anti-HA
Anti-tubulin
7% SDS-PAGE
AID::NUP358 HCT116

**Fig. S63. Localization analysis of 3×HA-tagged NUP358 variants in NUP358 depleted cells.** (**A**) Schematics indicating the domain boundaries of the transfected N-terminally 3×HA-tagged NUP358 variants. (**B**) Immunofluorescence microscopy localization analysis of 3×HA-NUP358 variants in *AID*::*NUP358* HCT116 cells after 3 hours of auxin treatment. mAb414 antibody staining was used as reference for the nuclear envelope. Scale bars are 10 μm. (**C**) Western blot expression analysis of 3×HA-tagged NUP358 variants in *AID*::*NUP358* HCT116 cells after 3 hours of auxin treatment.

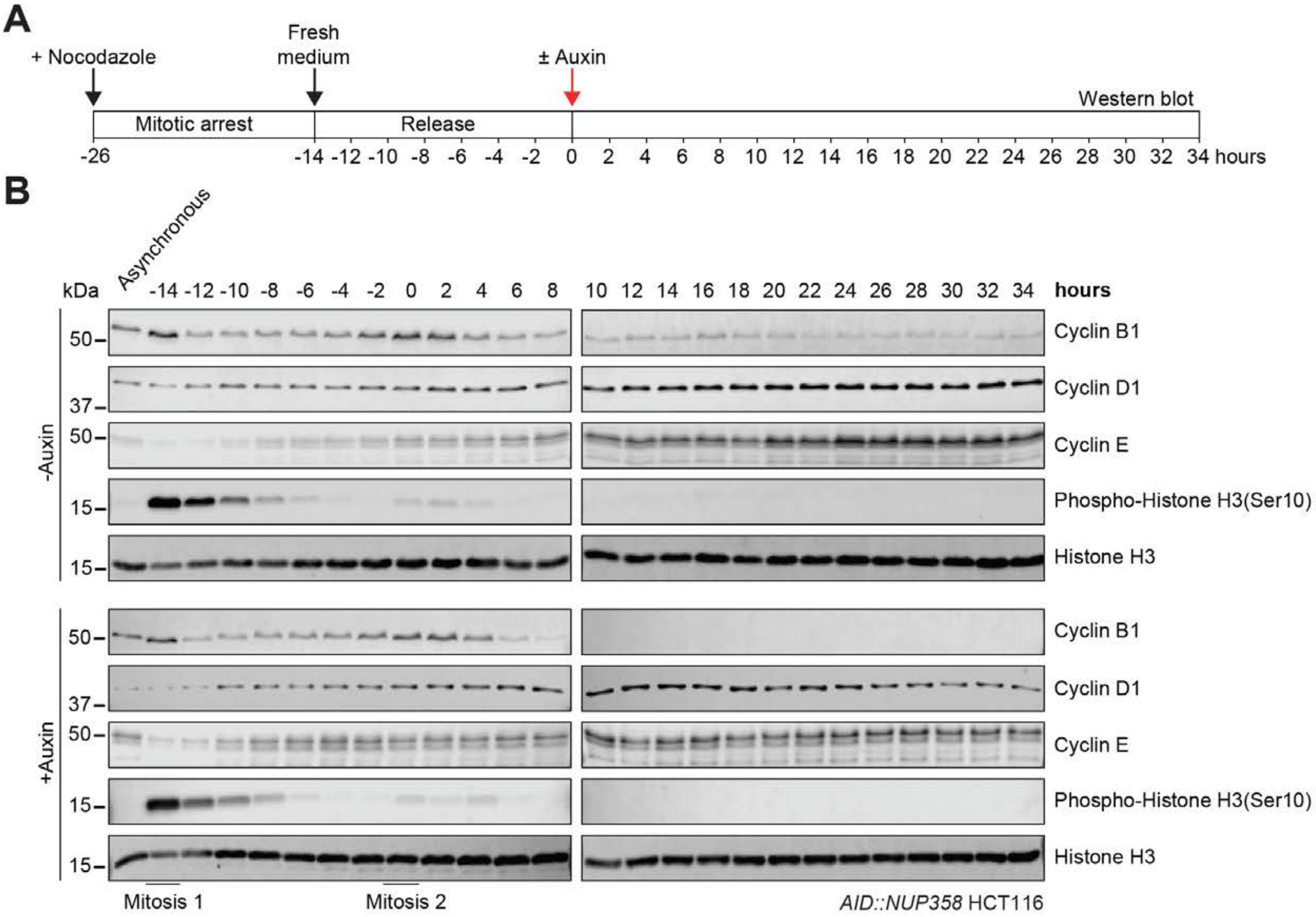

Sup Fig. S64

Hoelz et al., 2021b

**Fig. S64. Analysis of the cell cycle in nocodazole synchronized *AID*::*NUP358* HCT116 cells.** (**A**) Experimental timeline. (**B**) Time resolved western blot analysis of the indicated cell cycle markers in *AID*::*NUP358* HCT116 cells post release from nocodazole block, establishing an ~14 hour cell cycle. Histone H3 serves as a loading control.

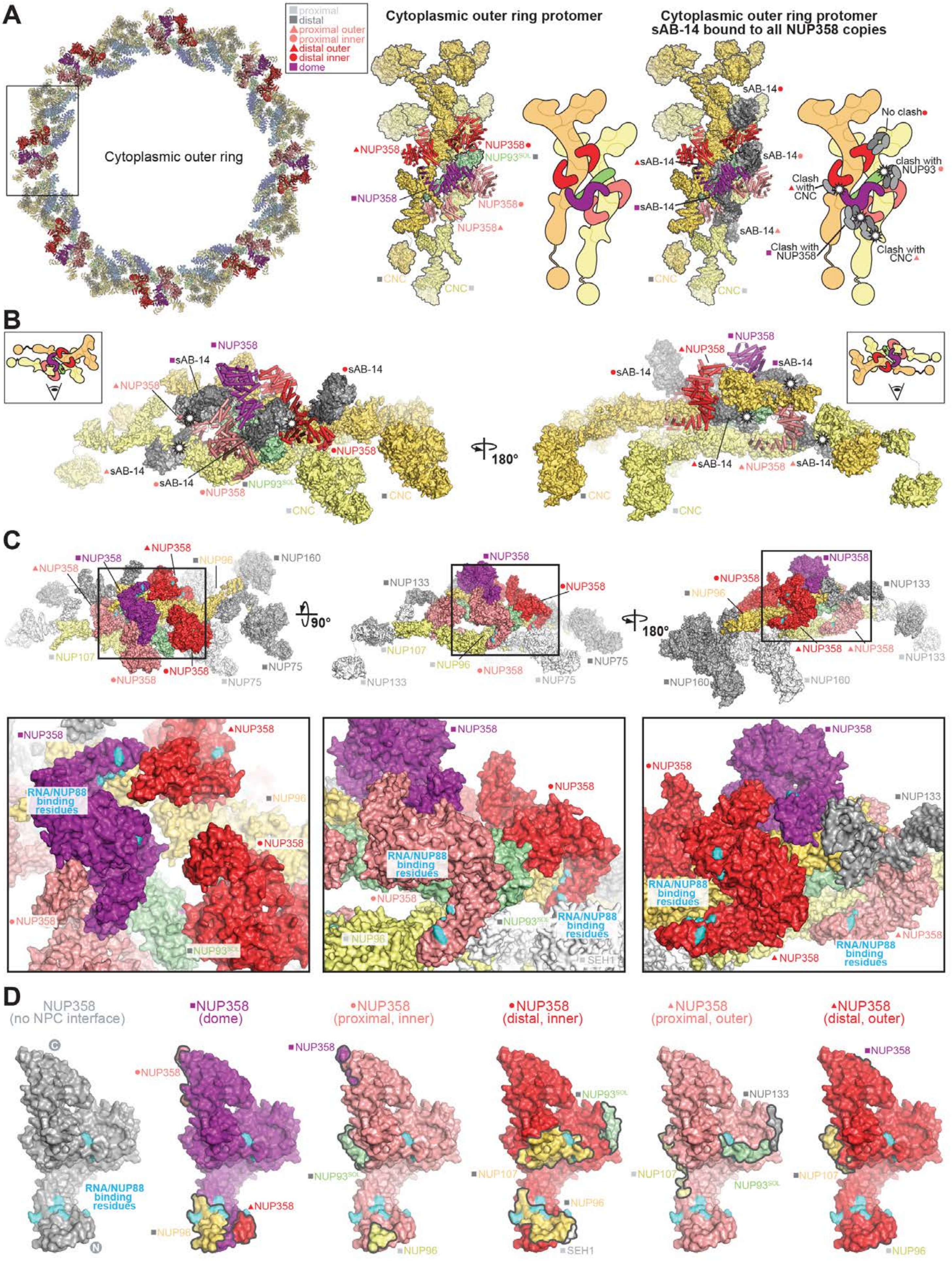
A
proximal
distal
proximal outer
proximal inner
distal outer
distal inner
dome
Cytoplasmic outer ring
Cytoplasmic outer ring protomer
Cytoplasmic outer ring protomer sAB-14 bound to all NUP358 copies
NUP358
NUP93SOL
CNC
sAB-14
No clash
clash with NUP93
Clash with CNC
Clash with NUP358
Clash with CNC
B
180°
C
90°
180°
NUP160
NUP107
NUP96
NUP75
NUP133
RNA/NUP88 binding residues
SEH1
D
NUP358 (no NPC interface)
NUP358 (dome)
NUP358 (proximal, inner)
NUP358 (distal, inner)
NUP358 (proximal, outer)
NUP358 (distal, outer)

**Fig. S65. Accessibility of sAB-14 and RNA/NUP88$^{NTD}$ binding sites in NPC-attached NUP358$^{NTD}$.** (**A**) Top view of the cytoplasmic face of the human NPC with NUP358$^{NTD}$ docked (*left*). A closeup view of the region from the inset indicates a single spoke protomer with tandem-arranged CNCs (distal, gold; proximal, pale yellow), and distal NUP93$^{SOL}$ (pale green) shown in surface representation, as well as five NUP358$^{NTD}$ copies (salmon, red, and purple) shown as cartoons (*middle*). An equivalent protomer is shown with NUP358$^{NTD}$•sAB-14 crystal structures superposed to the docked NUP358$^{NTD}$ copies. The sAB-14 heavy chain (light gray) and light chain (dark gray) are shown in surface representation. White stars indicate potential clashes that would occur if sAB-14 bound to a given NPC-attached NUP358$^{NTD}$. Schematics are shown to facilitate the interpretation of the rendered cartoons and surfaces. Distal and proximal positions are labeled according to the legend. (**B**) Two side views orthogonal to the cytoplasmic face protomer superposed with NUP358$^{NTD}$•sAB-14 in (A). (**C**) Three views of a surface representation of the cytoplasmic face protomer with closeup views of the regions in the insets. Residues identified as involved in RNA and NUP88$^{NTD}$ binding by surface mutagenesis of NUP358$^{NTD}$ are colored cyan. (**D**) Surface representation (gray) of NUP358$^{NTD}$ with RNA/NUP88$^{NTD}$-binding residues colored cyan (*left*). Five NUP358$^{NTD}$ copies found at the five unique positions in the cytoplasmic face protomer and colored accordingly (salmon, red, and purple) are shown in the same orientation, with RNA/NUP88$^{NTD}$-binding residues colored cyan, and surface patches colored according to the color assigned to the interfacing nup.

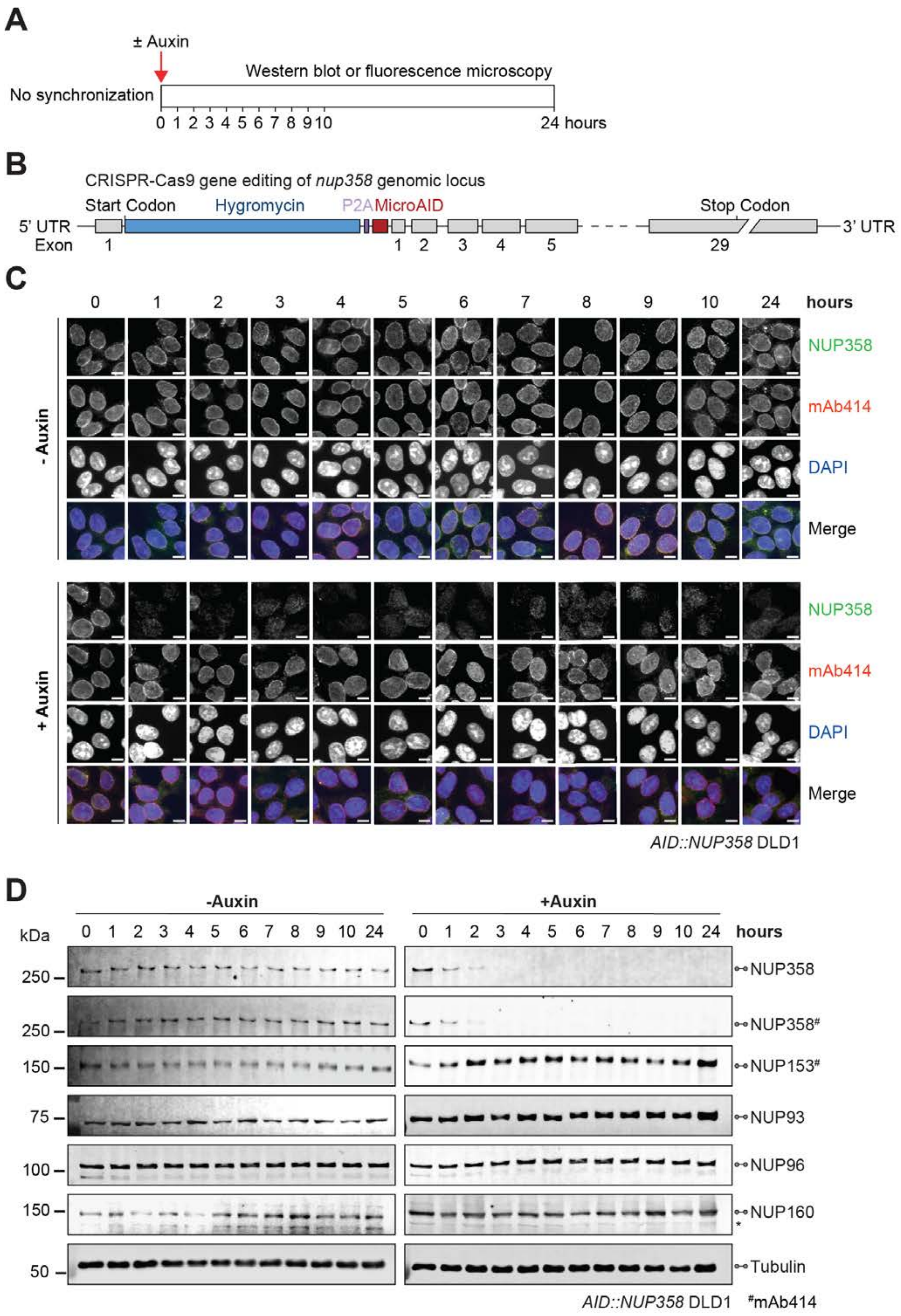
A
± Auxin
Western blot or fluorescence microscopy
No synchronization
0 1 2 3 4 5 6 7 8 9 10
24 hours
B
CRISPR-Cas9 gene editing of nup358 genomic locus
Start Codon
Hygromycin
P2A
MicroAID
Stop Codon
5' UTR
3' UTR
Exon
1
1 2 3 4 5
29
C
0 1 2 3 4 5 6 7 8 9 10 24 hours
- Auxin
+ Auxin
NUP358
mAb414
DAPI
Merge
AID::NUP358 DLD1
D
-Auxin
+Auxin
kDa
0 1 2 3 4 5 6 7 8 9 10 24
0 1 2 3 4 5 6 7 8 9 10 24 hours
250
250
150
75
100
150
50
NUP358
NUP358#
NUP153#
NUP93
NUP96
NUP160
*
Tubulin
AID::NUP358 DLD1
#mAb414

**Fig. S66. Validation of auxin-induced NUP358 degradation in *AID*::*NUP358* DLD1 cells.** (**A**) Experimental timeline. (**B**) Schematic representation of CRISPR/Cas9-mediated genomic tagging of *nup358* alleles. (**C**) Time-resolved subcellular localization analysis of AID-tagged NUP358 by immunofluorescence microscopy upon auxin treatment. The mAb414 antibody (red) and DAPI (blue) staining were used as a reference for the nuclear envelope and nucleus, respectively. Scale bars are 10 μm. (**D**) Time-resolved western blot analysis of AID-tagged NUP358 expression levels upon auxin treatment. The specificity of auxin induced NUP358 depletion was assessed by the indicated antibodies. Asterisks indicate a nonspecific band detected by the anti-NUP160 antibody. Control cells were not treated with auxin.

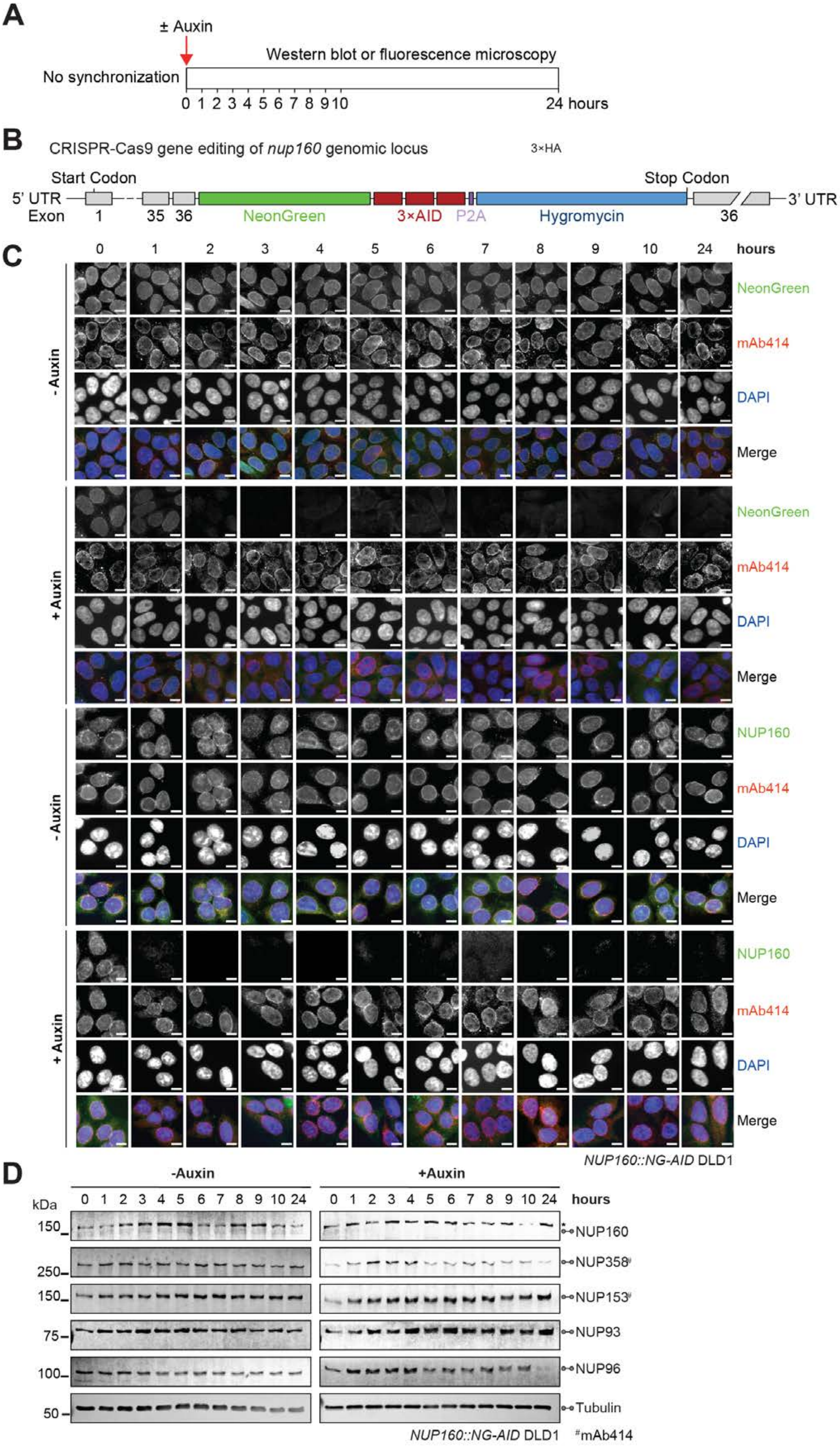
A
± Auxin
Western blot or fluorescence microscopy
No synchronization
0 1 2 3 4 5 6 7 8 9 10
24 hours
B
CRISPR-Cas9 gene editing of nup160 genomic locus
3×HA
Start Codon
Stop Codon
5' UTR
3' UTR
Exon
1
35
36
NeonGreen
3×AID
P2A
Hygromycin
36
C
0 1 2 3 4 5 6 7 8 9 10 24 hours
- Auxin
+ Auxin
NeonGreen
mAb414
DAPI
Merge
NUP160
NUP160::NG-AID DLD1
D
-Auxin
+Auxin
kDa
0 1 2 3 4 5 6 7 8 9 10 24
0 1 2 3 4 5 6 7 8 9 10 24 hours
150
250
150
75
100
50
NUP160
NUP358#
NUP153#
NUP93
NUP96
Tubulin
NUP160::NG-AID DLD1
#mAb414

**Fig. S67. Validation of auxin-induced NUP160 degradation in *NUP160*::*NG-AID* DLD1 cells.** (**A**) Experimental timeline. (**B**) Schematic representation of CRISPR/Cas9-mediated genomic tagging of *nup160* alleles. (**C**) Time-resolved subcellular localization analysis of NeonGreen (NG)-AID-tagged NUP160 by immunofluorescence microscopy upon auxin treatment. The NG-AID-tagged NUP160 was detected by an anti-NUP160 antibody (green) or NeonGreen fluorescence (green). The mAb414 antibody (red) and DAPI (blue) staining were used as a reference for the nuclear envelope and nucleus, respectively. Scale bars are 10 $\mu$m. (**D**) Time-resolved western blot analysis of NG-AID-tagged NUP160 expression levels upon auxin treatment. The specificity of auxin induced NUP160 depletion was assessed by the indicated antibodies. Asterisks indicate a nonspecific band detected by the anti-NUP160 antibody. Control cells were not treated with auxin.

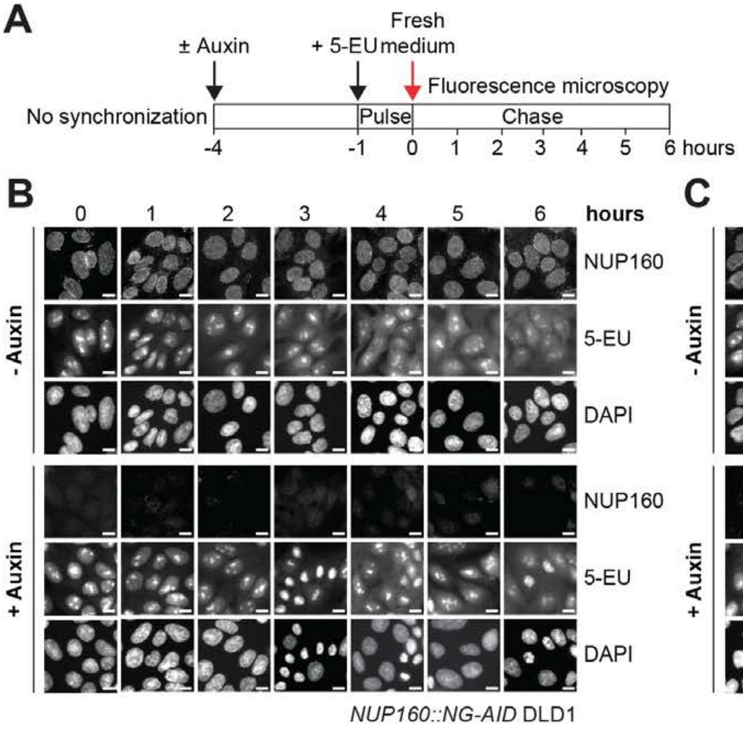

A
± Auxin
+ 5-EU
Fresh medium
Fluorescence microscopy
No synchronization
Pulse
Chase
-4
-1
0
1
2
3
4
5
6 hours
B
0
1
2
3
4
5
6
hours
- Auxin
+ Auxin
NUP160
5-EU
DAPI
NUP160
5-EU
DAPI
NUP160::NG-AID DLD1
C
0
1
2
3
4
5
6
hours
- Auxin
+ Auxin
NUP358
5-EU
DAPI
NUP358
5-EU
DAPI
AID::NUP358 DLD1

**Fig. S68. Time-resolved analysis of nuclear RNA retention in NUP160/NUP358 depleted DLD1 cells.** (**A**) Experimental timeline. (**B, C**) Time-resolved fluorescence microscopy analysis of RNA nuclear retention in (B) *NUP160*::*NG-AID* or (C) *AID*::*NUP358* DLD1 cells upon auxin treatment. Nascent RNA was metabolically pulse-labeled with 5-ethynyl uridine (5-EU) and its subcellular localization determined by Alexa Fluor 594 azide conjugation and fluorescence microscopy at the indicated time points. Scale bars are 10 $\mu$m.

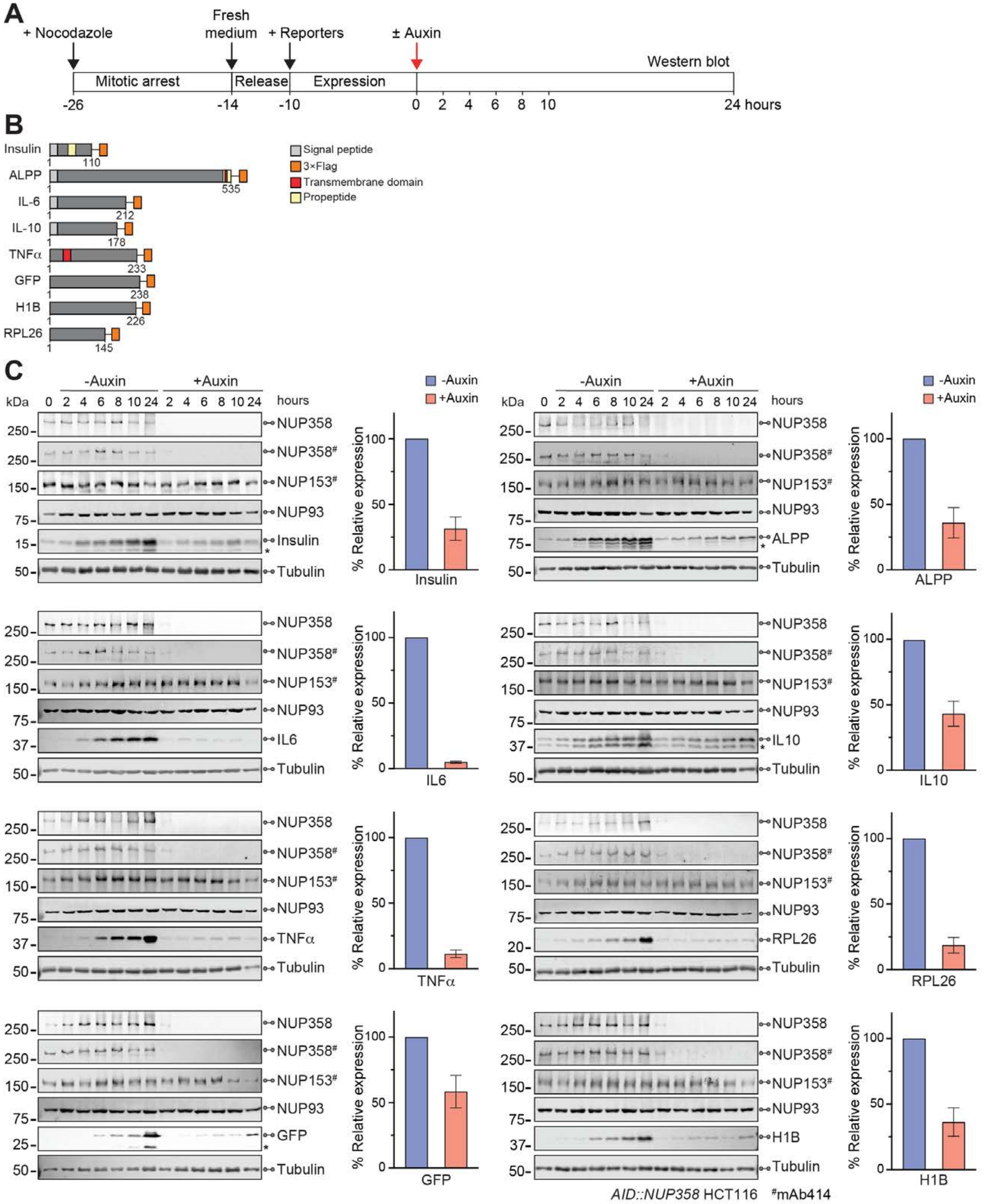
A
+ Nocodazole
Fresh medium
+ Reporters
± Auxin
Mitotic arrest
Release
Expression
Western blot
-26
-14
-10
0 2 4 6 8 10
24 hours
B
Insulin
ALPP
IL-6
IL-10
TNFα
GFP
H1B
RPL26
Signal peptide
3×Flag
Transmembrane domain
Propeptide
C
-Auxin
+Auxin
hours
kDa
NUP358
NUP358#
NUP153#
NUP93
Insulin
ALPP
IL6
IL10
TNFα
RPL26
GFP
H1B
Tubulin
% Relative expression
AID::NUP358 HCT116
#mAb414

**Fig. S69. Depletion of NUP358 reduces reporter protein expression.** (**A**) Experimental timeline. (**B**) Domain structures of the transfected C-terminally 3×FLAG-tagged reporters. (**C**) Time-resolved western blot expression analysis of the 3×FLAG-tagged reporter proteins in synchronized *AID*::*NUP358* HCT116 cells upon auxin-induced NUP358 depletion. Control cells were not treated with auxin. Quantitated reporter expression in auxin-treated cells was normalized to expression in control cells, at the 10-hour timepoint. Experiments were performed in triplicate, and the mean with the associated standard error reported. Asterisks indicate degradation bands detected by the anti-FLAG antibody.

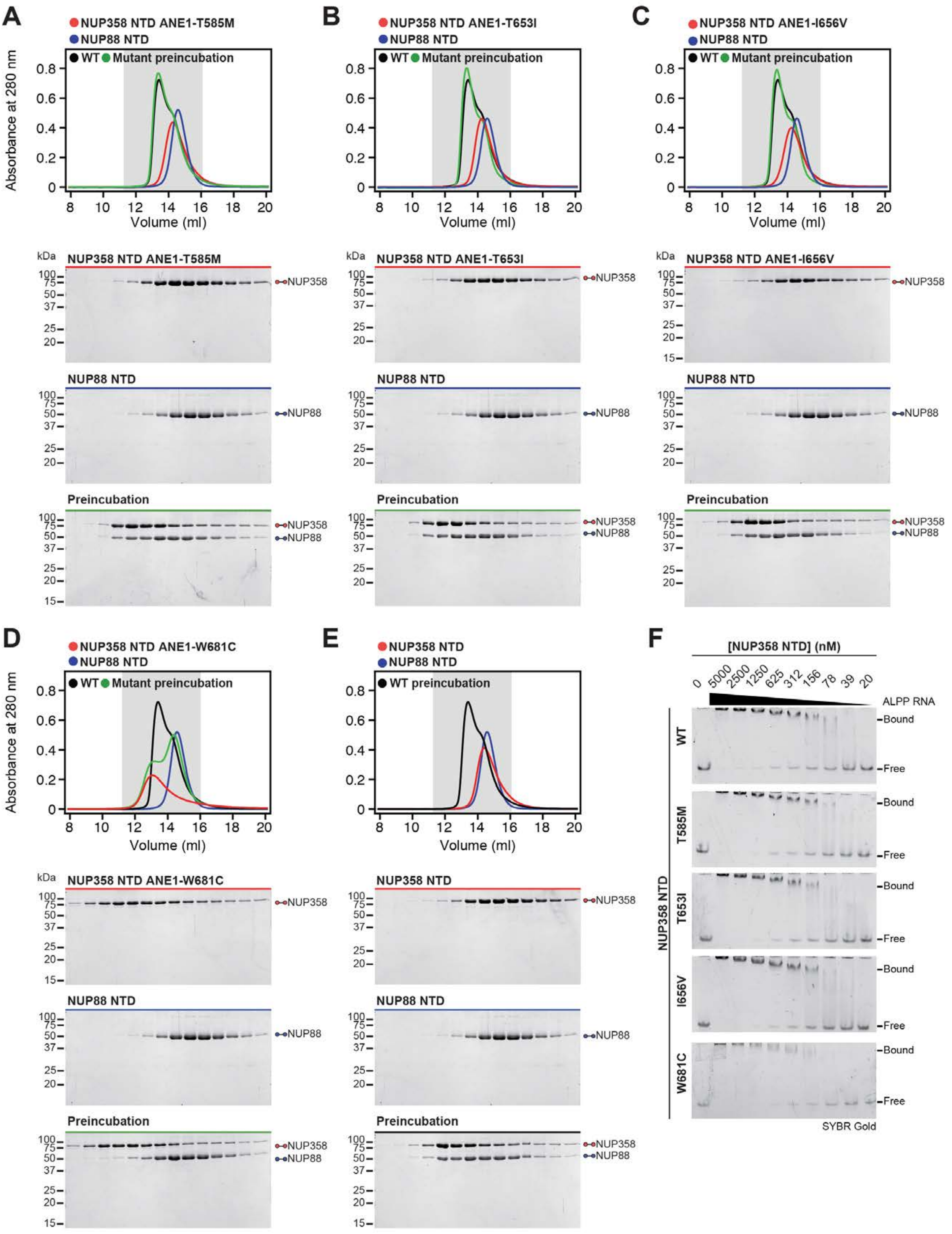
A
NUP358 NTD ANE1-T585M
NUP88 NTD
WT Mutant preincubation
Absorbance at 280 nm
Volume (ml)
kDa
NUP358 NTD ANE1-T585M
NUP358
NUP88 NTD
NUP88
Preincubation
B
NUP358 NTD ANE1-T653I
NUP88 NTD
WT Mutant preincubation
Volume (ml)
NUP358 NTD ANE1-T653I
C
NUP358 NTD ANE1-I656V
NUP88 NTD
WT Mutant preincubation
Volume (ml)
NUP358 NTD ANE1-I656V
D
NUP358 NTD ANE1-W681C
NUP88 NTD
WT Mutant preincubation
NUP358 NTD ANE1-W681C
E
NUP358 NTD
NUP88 NTD
WT preincubation
F
[NUP358 NTD] (nM)
0 5000 2500 1250 625 312 156 78 39 20
ALPP RNA
NUP358 NTD
WT
T585M
T653I
I656V
W681C
Bound
Free
SYBR Gold

**Fig. S70. Interaction analysis of NUP358$^{NTD}$ ANE1 mutants with NUP88$^{NTD}$ and ALPP$^{SSCR}$ RNA.** (**A**-**E**) SEC interaction analysis of NUP88$^{NTD}$ with (A) NUP358$^{NTD}$ T585M, (B) NUP358$^{NTD}$ T653I, (C) NUP358$^{NTD}$ I656V, (D) NUP358$^{NTD}$ W681C, and (E) wildtype NUP358$^{NTD}$. SEC profiles of NUP358$^{NTD}$ variants (red) and NUP88$^{NTD}$ (blue) are shown individually and after their preincubation (green). SEC profiles were obtained using a Superdex 200 10/300 GL column. Gray boxes indicate fractions resolved on SDS-PAGE gels and visualized by Coomassie brilliant blue staining. (**F**) EMSA titrations of NUP358$^{NTD}$ variants used in (A-E) against ALPP$^{SSCR}$ RNA (corresponding to Fig. 8D). Representative titrations of triplicate are shown. EMSAs visualized with SYBR Gold stain.

A

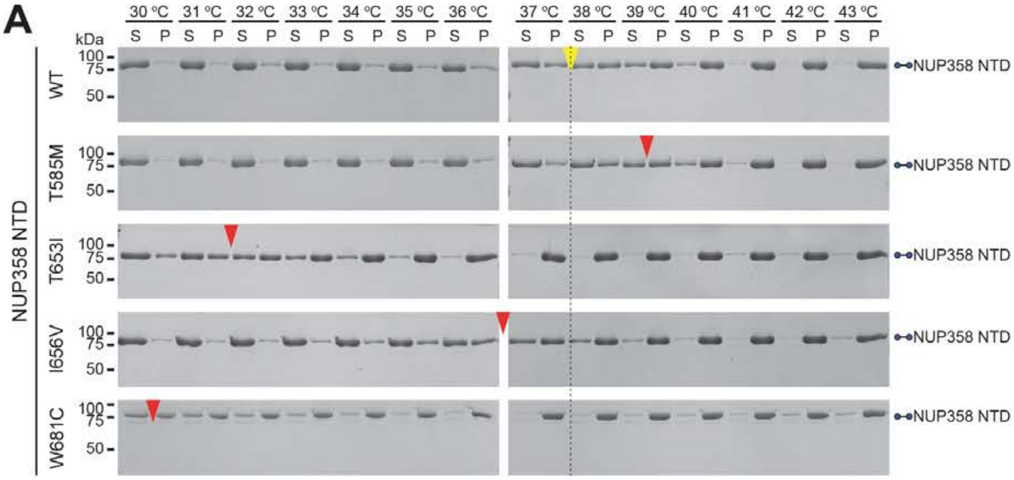
30 °C
31 °C
32 °C
33 °C
34 °C
35 °C
36 °C
37 °C
38 °C
39 °C
40 °C
41 °C
42 °C
43 °C
S
P
kDa
100
75
50
WT
T585M
T653I
I656V
W681C
NUP358 NTD
NUP358 NTD

B

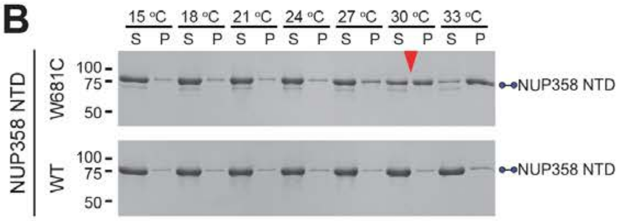
15 °C
18 °C
21 °C
24 °C
27 °C
30 °C
33 °C
S
P
100
75
50
W681C
WT
NUP358 NTD
NUP358 NTD

**Fig. S71. Thermostability assay of NUP358 harboring ANE1 mutants.** (**A**) Wildtype and NUP358$^{NTD}$ mutants associated with ANE1 (T585M, T653I, I656V, and W681C) were incubated for 20 minutes at the indicated temperatures prior to centrifugation. Pelleted (P) and soluble (S) fractions were analyzed by SDS-PAGE and visualized by Coomassie brilliant blue staining. (**B**) Lower temperature range thermostability pelleting assay for highly unstable NUP358$^{NTD}$ W681C and wildtype Nup358$^{NTD}$. Red arrows indicate 50% solubility temperature for NUP358$^{NTD}$ ANE1 mutants. For reference, a yellow arrow indicates 50% solubility temperature of wildtype NUP358$^{NTD}$.

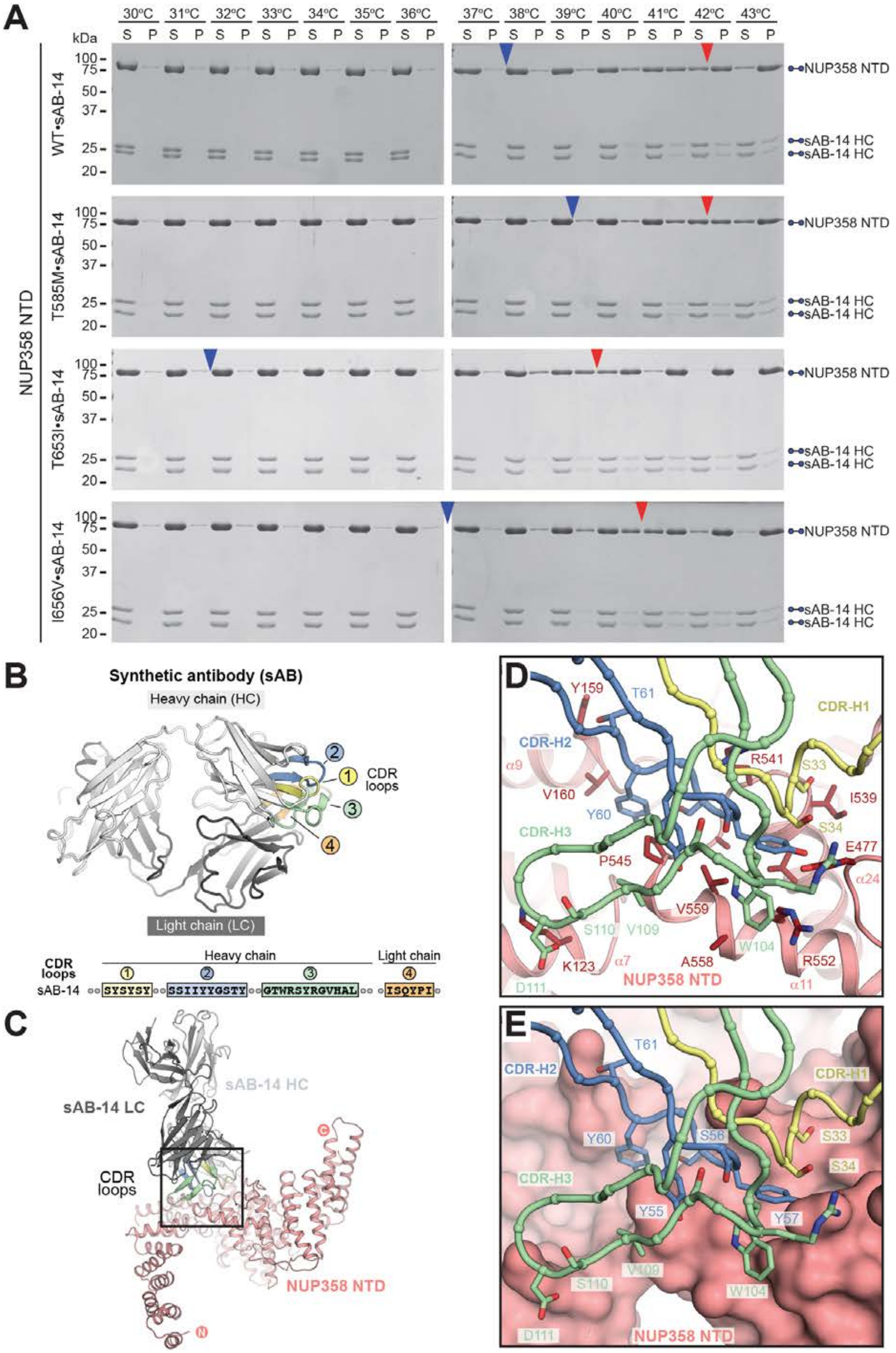
A
30ºC 31ºC 32ºC 33ºC 34ºC 35ºC 36ºC 37ºC 38ºC 39ºC 40ºC 41ºC 42ºC 43ºC
S P
kDa
100
75
50
37
25
20
NUP358 NTD
WT•sAB-14
T585M•sAB-14
T653I•sAB-14
I656V•sAB-14
sAB-14 HC
B
Synthetic antibody (sAB)
Heavy chain (HC)
Light chain (LC)
CDR loops
Heavy chain
Light chain
sAB-14
SYSYSY
SSIIYYGSTY
GTWRSYRGVHAL
ISQYPI
C
sAB-14 HC
sAB-14 LC
CDR loops
NUP358 NTD
D
Y159
T61
CDR-H1
CDR-H2
R541
S33
I539
V160
Y60
S34
CDR-H3
E477
P545
V559
S110
V109
W104
K123
A558
R552
D111
NUP358 NTD
E
S56
Y55
Y57

**Fig. S72. Thermostability of NUP358$^{NTD}$ ANE1 mutations is increased by sAB-14 binding.** (**A**) Thermostability assay of NUP358 harboring ANE1 mutants (T585M, T653I, I656V, and W681C) in the presence of sAB-14. Wildtype NUP358$^{NTD}$ and mutants associated with ANE1 were pre-complexed with sAB-14 and purified by SEC prior to performing the protein thermostability pelleting assay. Complexes were incubated at the indicated temperatures for 20 minutes prior to centrifugation. Pelleted (P) and soluble (S) fractions were analyzed by SDS-PAGE and visualized by Coomassie brilliant blue staining. The 50% solubility temperature for proteins in isolation and in presence of sAB-14 are indicated by blue and red arrows, respectively. (**B**) sAB-14 structure shown in cartoon representation. The sequences of the four complementarity determining region (CDR) loops are provided and their location in the structure indicated. (**C**) Structure of NUP358$^{NTD}$•sAB-14 shown in carton representation. The sAB-14 binding site is located on the convex surface of NUP358$^{NTD}$ at the interface between the N-terminal solenoid and central wedge domain. The inset indicates the region magnified in panels (D) and (E). (**D**) Magnified view of the sAB-14-NUP358$^{NTD}$ interface, illustrating the tight interactions formed by the four sAB-14 CDR loops with an exposed hydrophobic patch on the NUP358NTD surface. (**E**) Magnified view as (D) with NUP358$^{NTD}$ shown in surface representation highlighting the shape complementarity of the sAB-14 CDR loops.

**A**

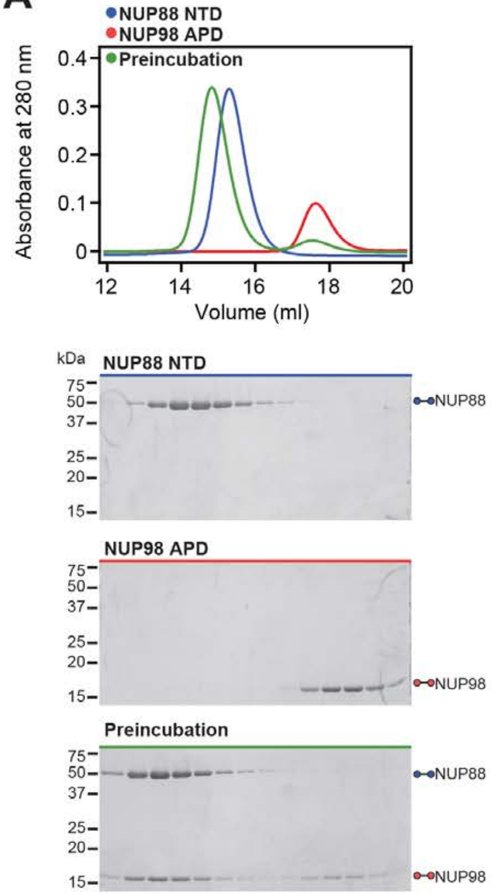
NUP88 NTD
NUP98 APD
Preincubation
Absorbance at 280 nm
0.4
0.3
0.2
0.1
0
12
14
16
18
20
Volume (ml)
kDa
NUP88 NTD
75
50
37
25
20
15
NUP88
NUP98 APD
NUP98
Preincubation
NUP88
NUP98

**B**

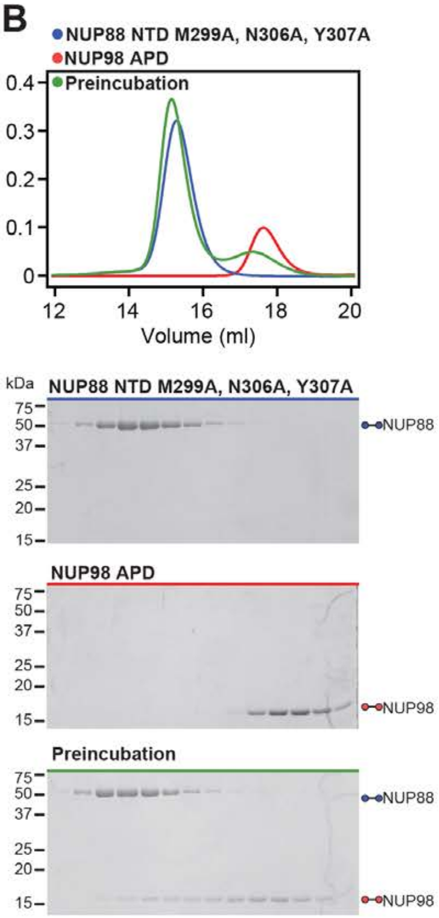
NUP88 NTD M299A, N306A, Y307A
NUP98 APD
Preincubation
0.4
0.3
0.2
0.1
0
12
14
16
18
20
Volume (ml)
kDa
NUP88 NTD M299A, N306A, Y307A
75
50
37
25
20
15
NUP88
NUP98 APD
NUP98
Preincubation
NUP88
NUP98

**C**

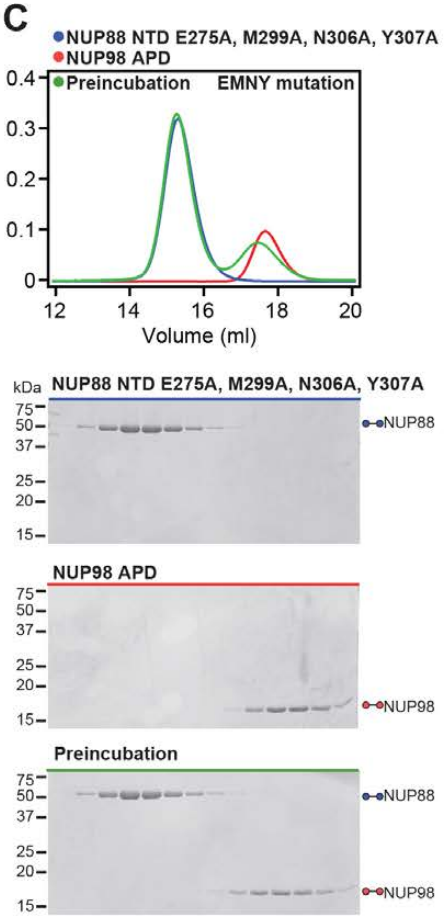
NUP88 NTD E275A, M299A, N306A, Y307A
NUP98 APD
Preincubation
EMNY mutation
0.4
0.3
0.2
0.1
0
12
14
16
18
20
Volume (ml)
kDa
NUP88 NTD E275A, M299A, N306A, Y307A
75
50
37
25
20
15
NUP88
NUP98 APD
NUP98
Preincubation
NUP88
NUP98

**D**

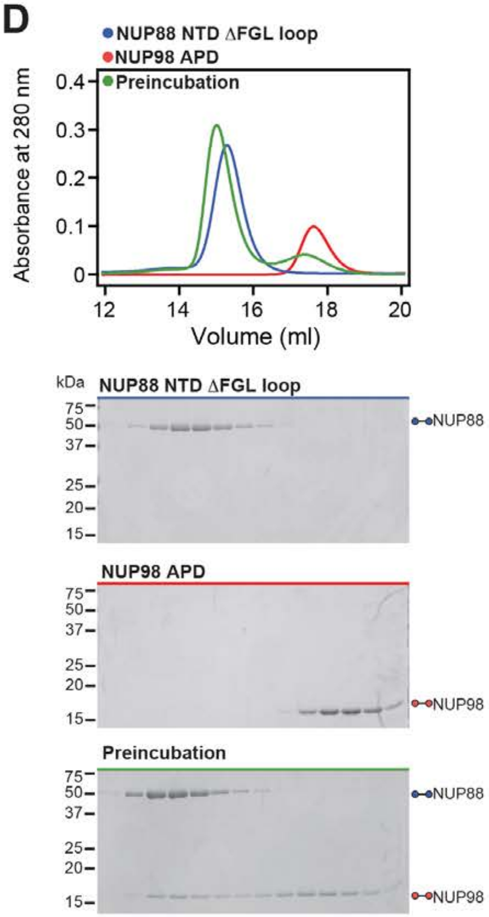
NUP88 NTD ΔFGL loop
NUP98 APD
Preincubation
Absorbance at 280 nm
0.4
0.3
0.2
0.1
0
12
14
16
18
20
Volume (ml)
kDa
NUP88 NTD ΔFGL loop
75
50
37
25
20
15
NUP88
NUP98 APD
NUP98
Preincubation
NUP88
NUP98

**Fig. S73. Mutational analysis of the NUP88$^{NTD}$-Nup98$^{APD}$ interaction.** (**A**-**D**) SEC interaction analyses (corresponding to Fig. 9, E and G) of NUP98$^{APD}$ with (A) wildtype NUP88$^{NTD}$, (B) the NUP88$^{NTD}$ M2999A/N306A/Y307A mutant, (C) the NUP88$^{NTD}$ E275A/M2999A/N306A/Y307A mutant, and (D) the NUP88$^{NTD}$ ∆FGL-loop mutant. SEC profiles of wildtype and mutant NUP88$^{NTD}$ (blue) and NUP98$^{APD}$ (red) are shown individually and after their preincubation (green). SEC profiles were obtained using a Superdex 200 10/300 GL column. Gray boxes indicate fractions resolved on SDS-PAGE gels and visualized by Coomassie brilliant blue staining.

**A**

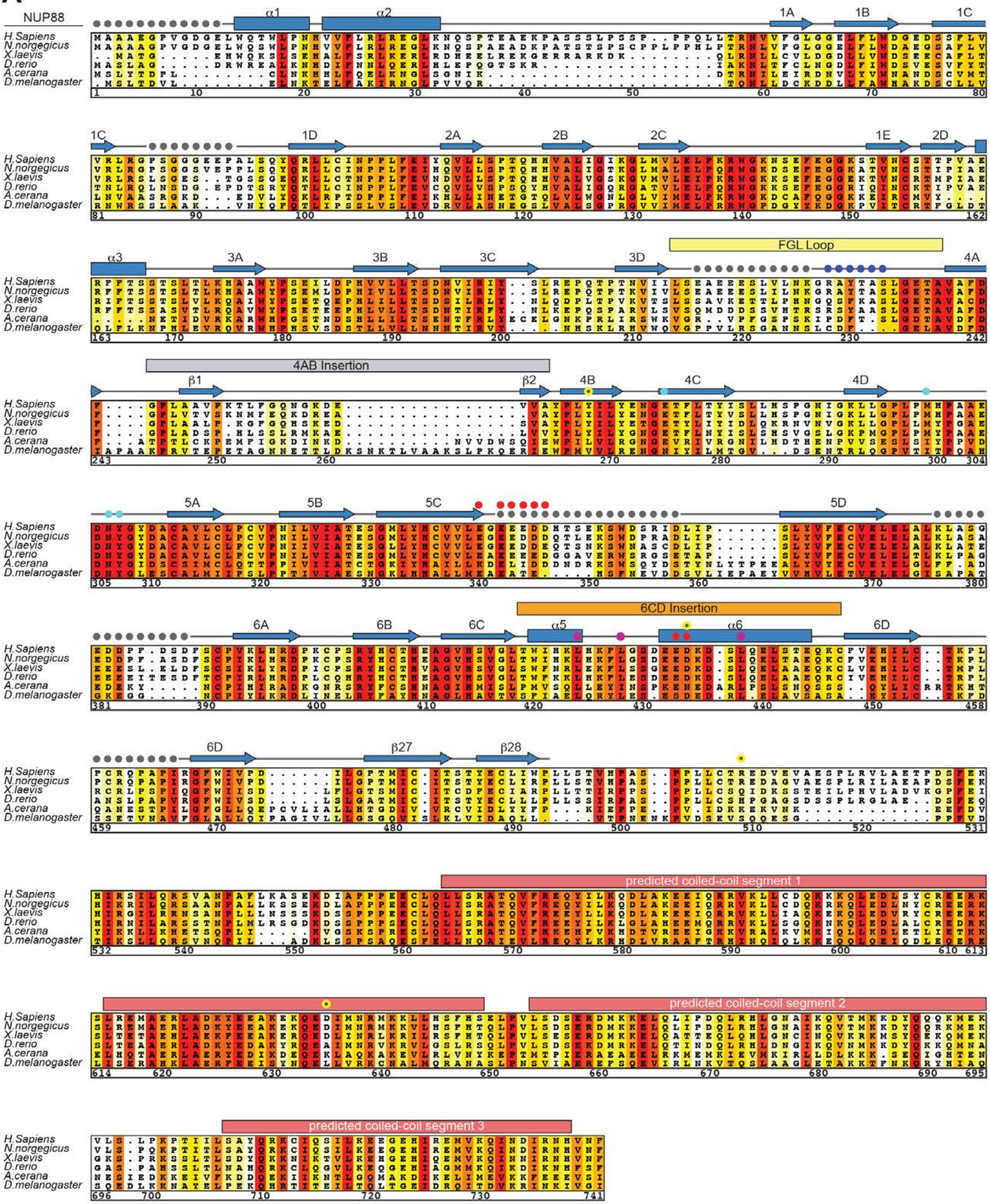

- fetal akinesia deformation sequence (FADS) associated mutation sites
- LLL mutation (NUP214 interaction)
- 5E3D muation (NUP358 interaction)
- ΔFGL mutation (NUP98 interaction)
- EMNY mutation (NUP98 interaction)
- unresolved

**Fig. S74. Multispecies sequence alignment of NUP88.** Sequences of NUP88 from six species were aligned and colored according to similarity using the BLOSUM62 matrix from white (less than 40% similarity), through yellow (60% similarity), to red (100% identity). The numbering is according to the *H. sapiens* protein. For $NUP88^{NTD}$, the secondary structure is indicated above the sequences as rectangles ($\alpha$-helices), arrows ($\beta$-strands), and lines (unstructured regions). Unresolved residues are indicated by gray dots. Secondary structure elements are colored according to Fig. 9. For $NUP88^{CTD}$, the secondary structure elements were predicted by JPred4 (*141*). Residues of sequence variations associated with fetal akinesia deformation sequence disease (*99*) are indicated by yellow dots. Residues substituted by alanine in the NUP88NTD 5E3D, LLL, and EMNY combination mutants are indicated by red, magenta, and cyan dots, respectively. Residues removed in the $\Delta$FGL loop mutation are indicated by blue dots.

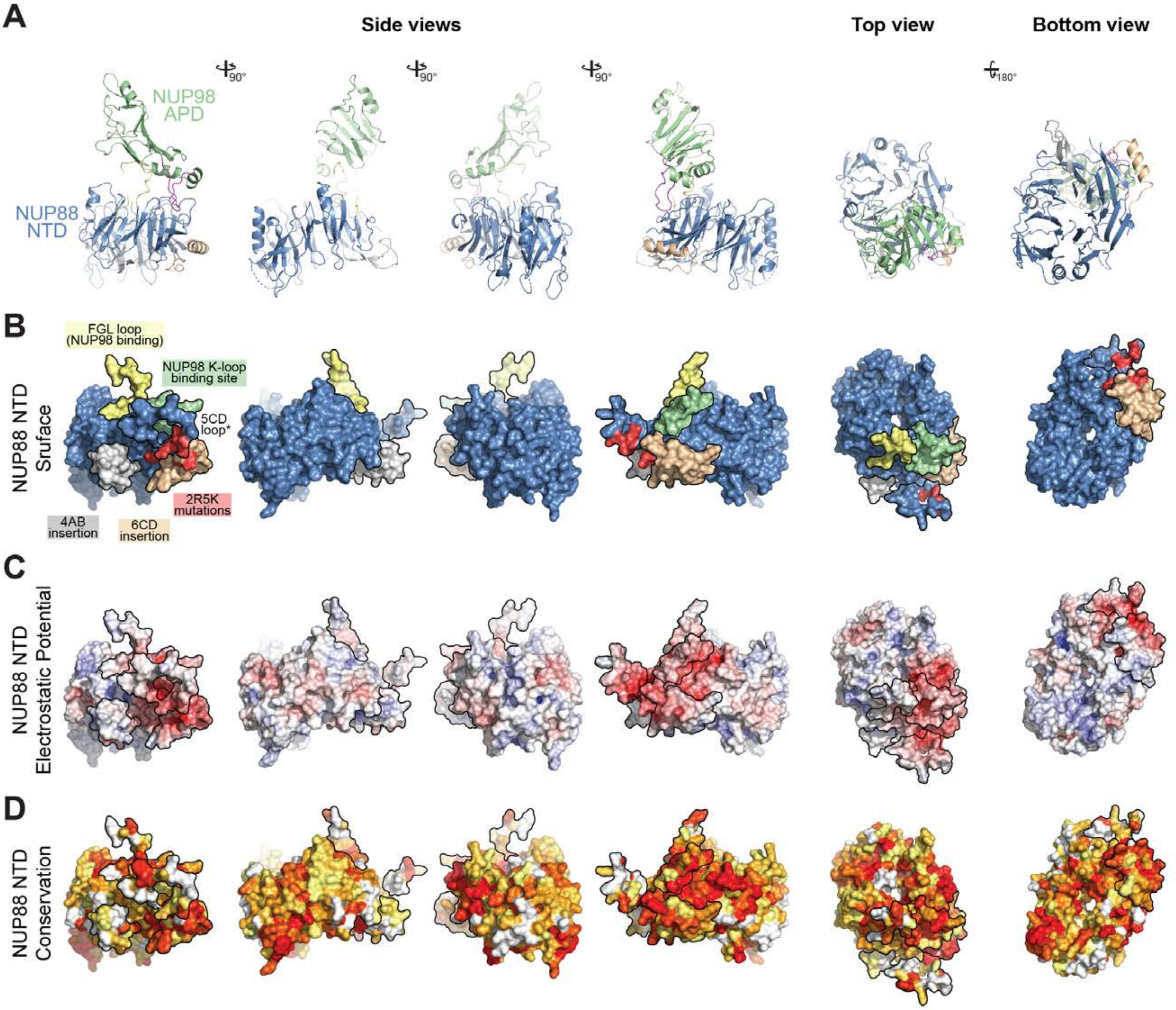
A
Side views
Top view
Bottom view
90°
90°
90°
180°
NUP98
APD
NUP88
NTD
B
FGL loop
(NUP98 binding)
NUP98 K-loop
binding site
5CD
loop*
2R5K
mutations
4AB
insertion
6CD
insertion
NUP88 NTD
Sruface
C
NUP88 NTD
Electrostatic Potential
D
NUP88 NTD
Conservation

**Fig. S75. Surface properties of NUP88$^{NTD}$.** (**A**) Cartoon representation of the NUP88$^{NTD}$•NUP98$^{APD}$ crystal structure. (**B**) Surface representation of NUP88$^{NTD}$ colored as in (A). The binding site of NUP98 (green) and the approximate location of the residues of the 2R5K combination mutant (yellow), which are located in the unresolved 5CD loop and abolish NUP358NTD binding, are indicated. (**C**) Surface representation colored according to electrostatic potential from red (-10 $k_BT/e$) to blue (+10 $k_BT/e$). (**D**) Surface representation colored according to sequence identity based on the alignment in fig. S74.The NUP98$^{APD}$ binding site is strongly negatively charged.

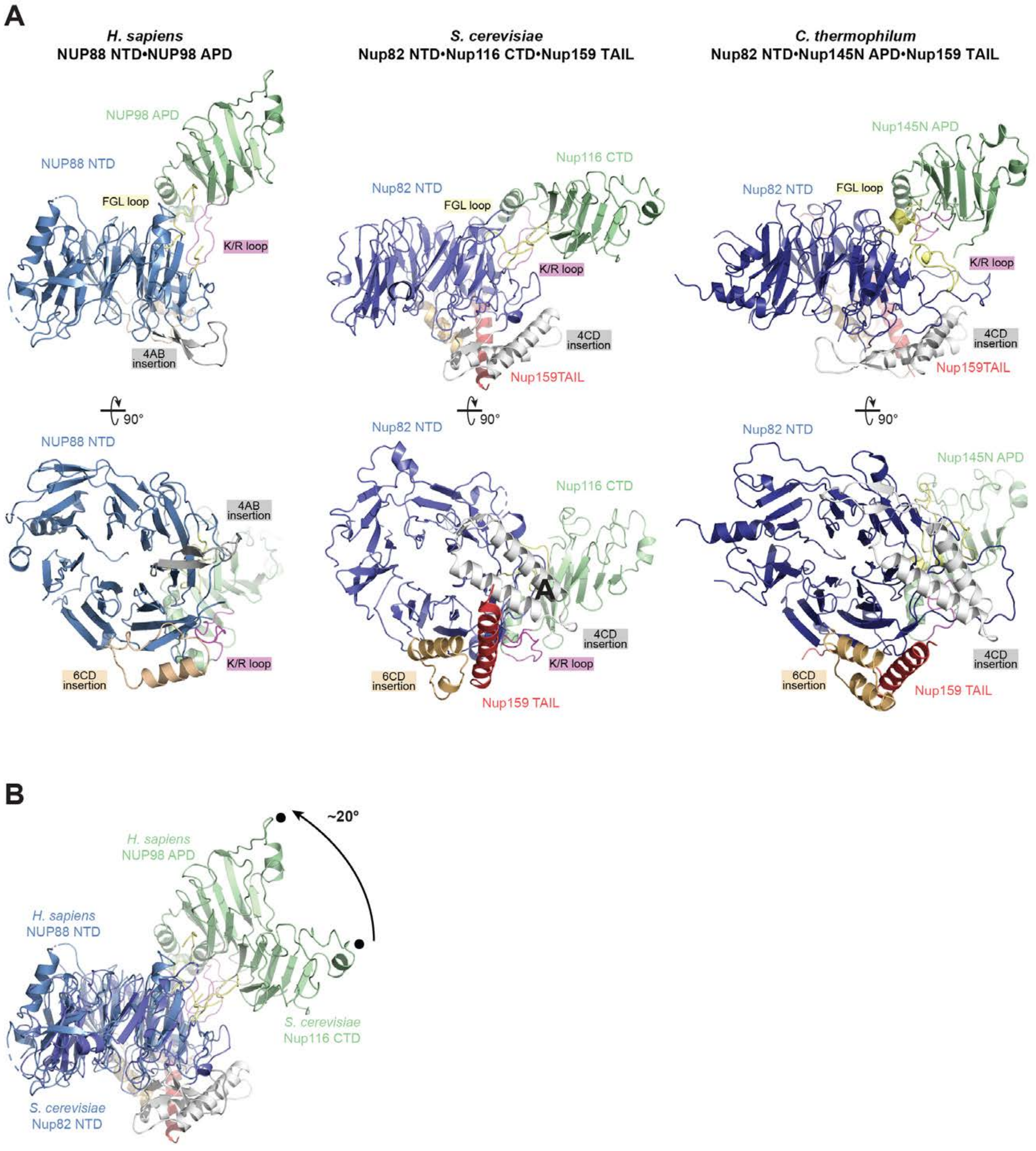
A
H. sapiens
NUP88 NTD•NUP98 APD
S. cerevisiae
Nup82 NTD•Nup116 CTD•Nup159 TAIL
C. thermophilum
Nup82 NTD•Nup145N APD•Nup159 TAIL
NUP98 APD
NUP88 NTD
FGL loop
K/R loop
4AB insertion
Nup116 CTD
Nup82 NTD
4CD insertion
Nup159TAIL
Nup145N APD
90°
6CD insertion
Nup159 TAIL
B
~20°
H. sapiens
NUP98 APD
H. sapiens
NUP88 NTD
S. cerevisiae
Nup116 CTD
S. cerevisiae
Nup82 NTD

**Fig. S76. Evolutionary conservation of the NUP88$^{NTD}$•NUP98$^{APD}$•NUP214$^{TAIL}$ architecture.** (**A**) Cartoon representations of the NUP88$^{NTD}$•NUP98$^{APD}$ co-crystal structure and the previously determined crystal structures of *S. cerevisiae* Nup82$^{NTD}$•Nup116$^{CTD}$•Nup159$^{TAIL}$ (PDB ID 3PBP) (*59*) and *C. thermophilum* Nup82$^{NTD}$•Nup145N$^{APD}$•Nup159$^{TAIL}$ (PDB ID 5CWW) (*11*), illustrating the evolutionary conservation of the NUP88/Nup82 FGL, 6CD, and 4AB/4CD loops and the NUP98/Nup116/Nup145N K/R loop (*top*). 90º-rotated views of all three complexes illustrating the location of the NUP88/Nup82$^{NTD}$ 6CD, and 4AB/4CD loop insertions that form the evolutionarily conserved binding site for NUP214/Nup159$^{TAIL}$ (*bottom*). (**D**) Superposition of *H. sapiens* and *S. cerevisiae* structures indicating the ~20° rotation of NUP98$^{APD}$/Nup116$^{CTD}$.

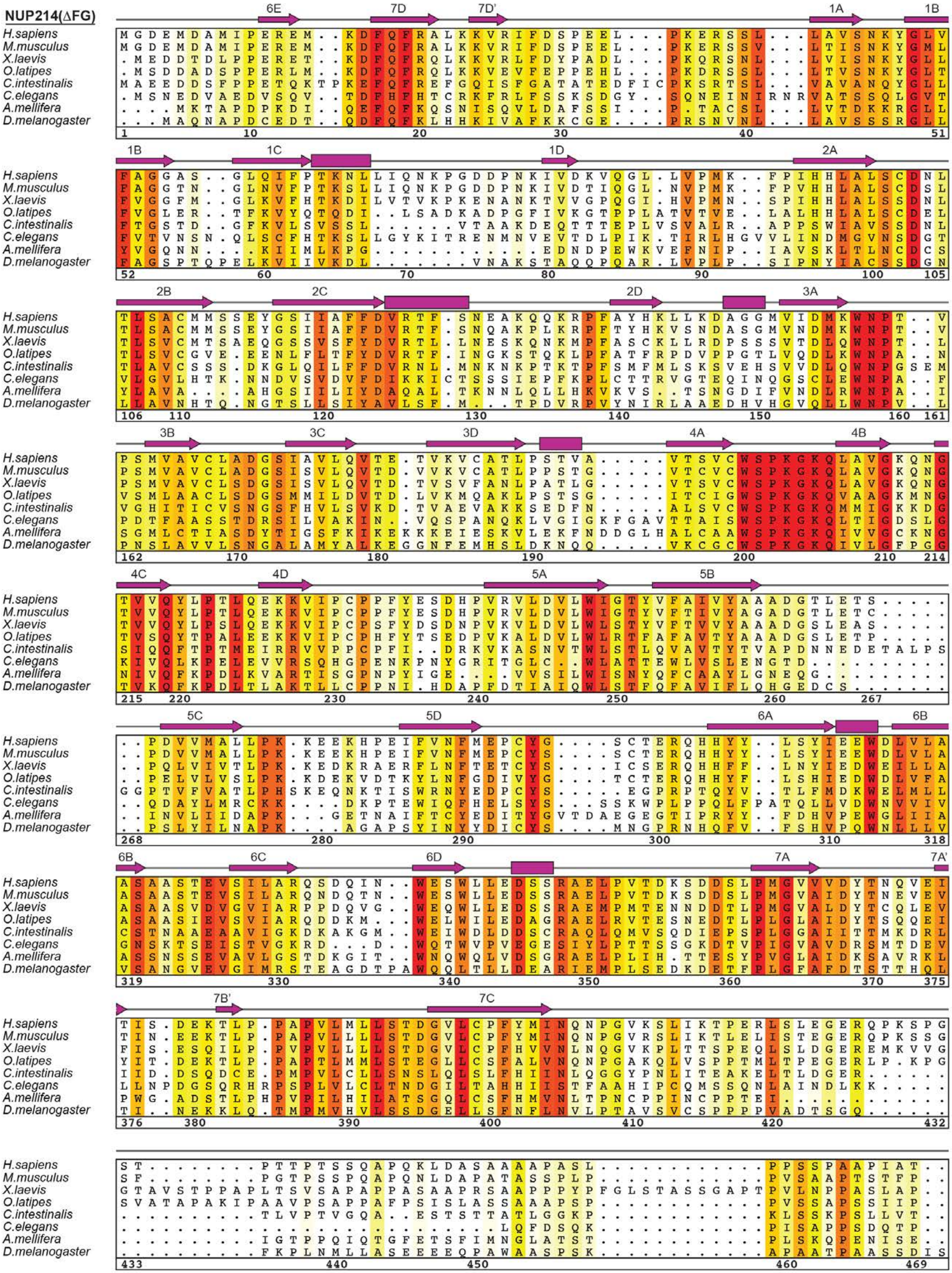

Sup Fig. S77A

Hoelz et al., 2021b

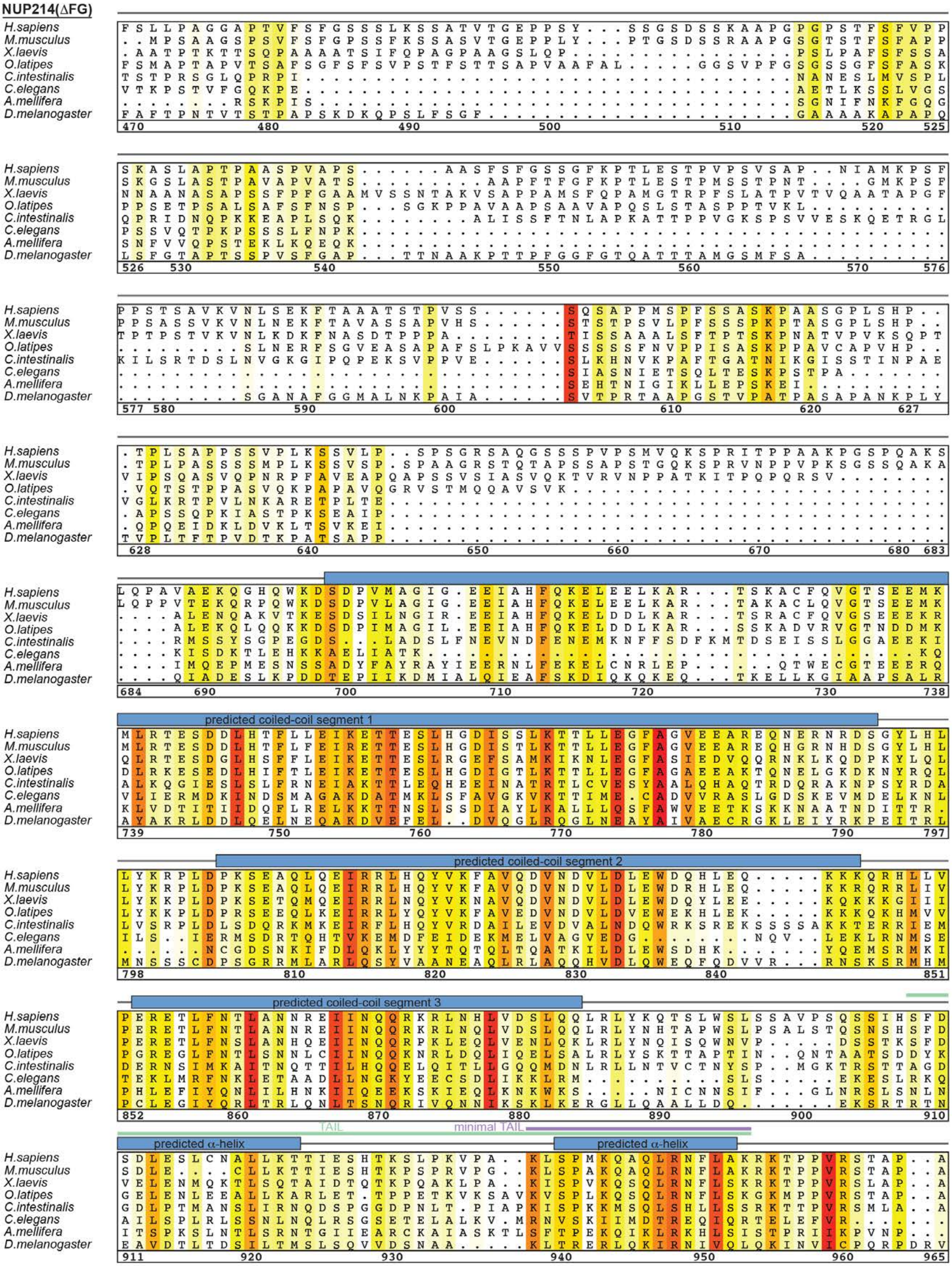

NUP214(ΔFG)
H.sapiens
M.musculus
X.laevis
O.latipes
C.intestinalis
C.elegans
A.mellifera
D.melanogaster
predicted coiled-coil segment 1
predicted coiled-coil segment 2
predicted coiled-coil segment 3
TAIL
minimal TAIL
predicted α-helix
predicted α-helix

**Fig. S77. Multispecies sequence alignment of NUP214.** Sequences of NUP214 from eight species were aligned and colored according to similarity using the BLOSUM62 matrix from white (less than 40% similarity), through yellow (60% similarity), to red (100% identity). The numbering is according to the *H. sapiens* protein. For $NUP214^{NTD}$, the secondary structure is indicated above the sequences as rectangles ($\alpha$-helices, magenta), arrows ($\beta$-strands, magenta), and lines (unstructured regions, gray) and was derived from the previously reported crystal structure (PDB ID 2OIT) (*54*). The secondary structure elements of the structurally uncharacterized C-terminal coiled-coil region is indicated as rectangles ($\alpha$-helices, blue) and was determined with JPred4 (*141*). The C-terminal ~1,100-residue FG-repeat region was omitted.

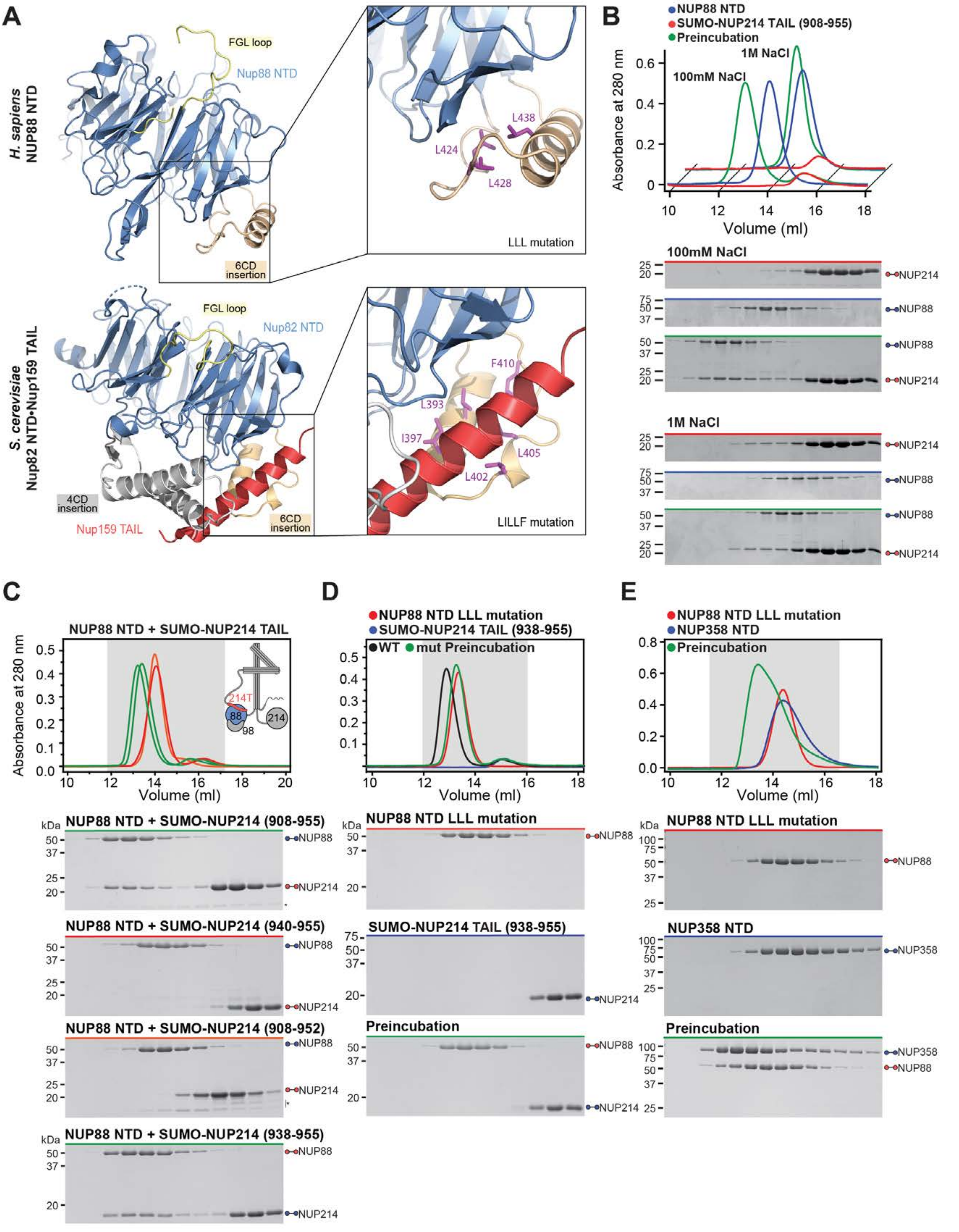
A
B
C
D
E
H. sapiens NUP88 NTD
FGL loop
Nup88 NTD
6CD insertion
L438
L424
L428
LLL mutation
S. cerevisiae Nup82 NTD•Nup159 TAIL
Nup82 NTD
4CD insertion
Nup159 TAIL
F410
L393
I397
L405
L402
LILLF mutation
NUP88 NTD
SUMO-NUP214 TAIL (908-955)
Preincubation
1M NaCl
100mM NaCl
Absorbance at 280 nm
Volume (ml)
NUP214
NUP88
NUP88 NTD + SUMO-NUP214 TAIL
214T
88
98
214
kDa
NUP88 NTD + SUMO-NUP214 (908-955)
NUP88 NTD + SUMO-NUP214 (940-955)
NUP88 NTD + SUMO-NUP214 (908-952)
NUP88 NTD + SUMO-NUP214 (938-955)
NUP88 NTD LLL mutation
SUMO-NUP214 TAIL (938-955)
WT
mut Preincubation
NUP358 NTD
NUP358

**Fig. S78. Evolutionary conservation of NUP214 TAIL binding site on NUP88$^{NTD}$.** (**A**) Cartoon representation of *H. sapiens* NUP88$^{NTD}$•NUP98$^{APD}$ and *S. cerevisiae* Nup82$^{NTD}$•Nup159$^{TAIL}$•Nup116$^{CTD}$ (PDB ID 3PBP) (*59*), colored according to Fig. 9. The insets indicate the location of the Nup82$^{NTD}$•Nup159$^{TAIL}$ interface and the corresponding region in NUP88$^{NTD}$ that are magnified on the right. The five scNup82$^{NTD}$ residues of the previously identified LILLF (L393A, I397A, L402A, L405A, and F410A) combination mutant, located in the 6CD loop *(59)*, and the corresponding three residues (L424, L428, and L438) of NUP88$^{NTD}$ are shown in ball-and-stick representation (purple). For clarity, NUP98$^{APD}$ Nup116$^{CTD}$ have been omitted. (**B**) SEC interaction analysis of NUP88$^{NTD}$ with SUMO-tagged NUP214$^{TAIL}$ in buffers containing 100 mM or 1 M NaCl, establishing the salt-stable nature of the interaction. (**C**) SEC interaction analysis of NUP88$^{NTD}$ with the indicated SUMO-tagged NUP214$^{TAIL}$ truncations, identifying a minimal NUP214$^{TAIL}$ fragment (minimal TAIL; residues 938-955), which is sufficient for NUP88$^{NTD}$ binding. (**D**) SEC interaction analyses of SUMO-tagged NUP214$^{minimal\ TAIL}$ with wildtype NUP88$^{NTD}$ and the NUP88$^{NTD}$ LLL (L424A, L428A, and L438A) combination mutant. The NUP88$^{NTD}$ LLL mutation abolishes the NUP214$^{minimal\ TAIL}$ interaction, identifying the NUP214$^{minimal\ TAIL}$ binding site on the NUP88$^{NTD}$ surface and suggesting a binding mode that is evolutionarily conserved from yeast to human. (**E**) SEC interaction analysis of the NUP88$^{NTD}$ LLL combination mutant with NUP358$^{NTD}$, demonstrating that the NUP88$^{NTD}$ LLL mutation does not affect the NUP358$^{NTD}$ interaction. SEC profiles were obtained using a Superdex 200 10/300 GL column. Gray boxes indicate fractions resolved on SDS-PAGE gels and visualized by Coomassie brilliant blue staining.

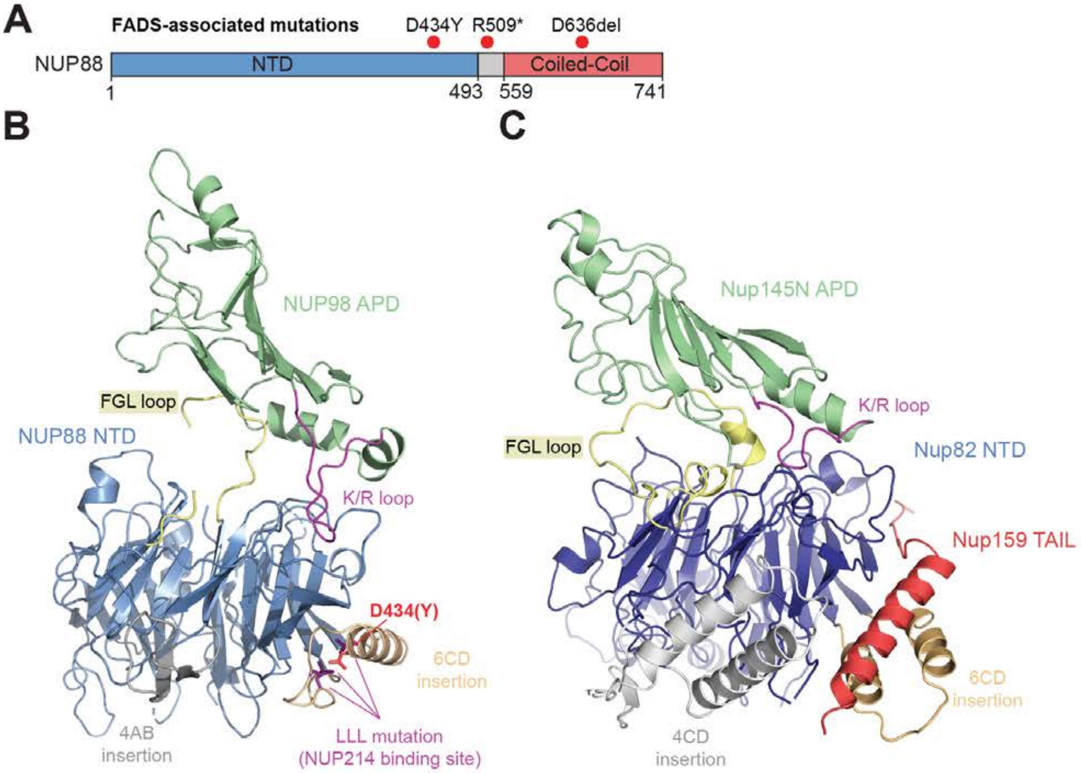

**NUP88 NTD•NUP98 APD**
**(*H.sapiens*)**

**Nup82 NTD•Nup145N APD•Nup159 TAIL**
**(*C. thermophilum*)**

**Fig. S79. FADS-associated NUP88 D434Y mutation is located at the NUP214$^{TAIL}$ binding site.** (**A**) NUP88 domain architecture indicating locations of the three mutations associated with fetal akinesia deformation sequence (FADS) (*99*). The D434Y mutant is located in the 6CD loop insertion of NUP88$^{NTD}$, directly adjacent to the identified NUP214$^{TAIL}$ binding site (L424, L428, and L438). The nonsense mutation R509* truncates the C-terminal coiled-coil domain, whereas the single residue deletion of D636 falls within the predicted coiled-coil domain. (**B**) Co-crystal structure of NUP88$^{NTD}$•NUP98$^{APD}$ is shown in cartoon representation and colored according to Fig 9, indicating the location of D434 and the adjacent L424, L428 and L438 of the LLL mutation of NUP88$^{NTD}$. (**C**) Crystal structure of *C. thermophilum* Nup82$^{NTD}$•Nup145N$^{APD}$•Nup159$^{TAIL}$ (PDB ID 5CWW) (*11*) shown in cartoon representation, illustrating the involvement of the 6CD loop of Nup159$^{NTD}$ in anchoring Nup159$^{TAIL}$.

**A**

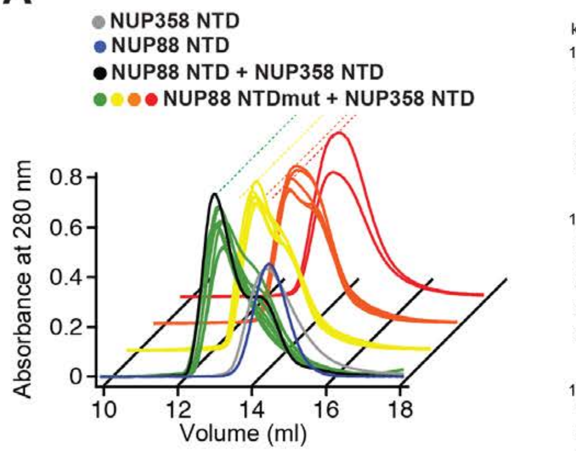

| Location | NUP88 mutation | NUP358 binding |
|---|---|---|
| L1 | D10A, E12A | No effect (green) |
| α2-1A | E29A, E38A, E40A | No effect (green) |
| 1BC/1CD | E74A, E75A, E91A, E92A | No effect (green) |
| 3D4A | E216A, E218A, E219A, E220A | No effect (green) |
| 3D4A/4AB insertion | E236A, E272A, E275A | No effect (green) |
| 4D5A | E304A, D305A, D310A | Mild (yellow) |
| 5BC/5D6A | E329A, E370A, E372A | No effect (green) |
| 5CD | E340A, E342A, E343A, E344A, D345A, D346A | Moderate (orange) |
| 5CD | E340A, E342A, E343A | Moderate (orange) |
| 5CD | E343A, E344A | Mild (yellow) |
| 5CD | E344A, D345, D346A | Moderate (orange) |
| 5CD | E350A, D354A, D358A | No effect (green) |
| 5D6A | E381A, D382A, D383A | No effect (green) |
| 6CD insertion-A | D431A, E432A, E433A, D434A | Moderate (orange) |
| 6CD insertion-B | D431A, E432A | Mild (yellow) |
| 6CD insertion-C | E433A, D434A | Moderate (orange) |
| 6CD insertion-D | D433A | Mild (yellow) |
| 6CD insertion-E | D434A | Mild (yellow) |
| 6CD insertion-F | D434Y | No effect (green) |
| 6CD insertion-G | D436A, E440A, E444A | Mild (yellow) |
| 5CD + 6CD insertion | E340A, E342A, E343A, E344A, D345A, D346A, D431A, E432A, E433A, D434A | Severe (red) |
| 5CD + 6CD insertion (5E3D) | E340A, E342A, E343A, E344A, D345A, D346A, E433A, D434A | Severe (red) |

Effect on Binding: No effect (green) Mild (yellow) Moderate (orange) Severe (red)

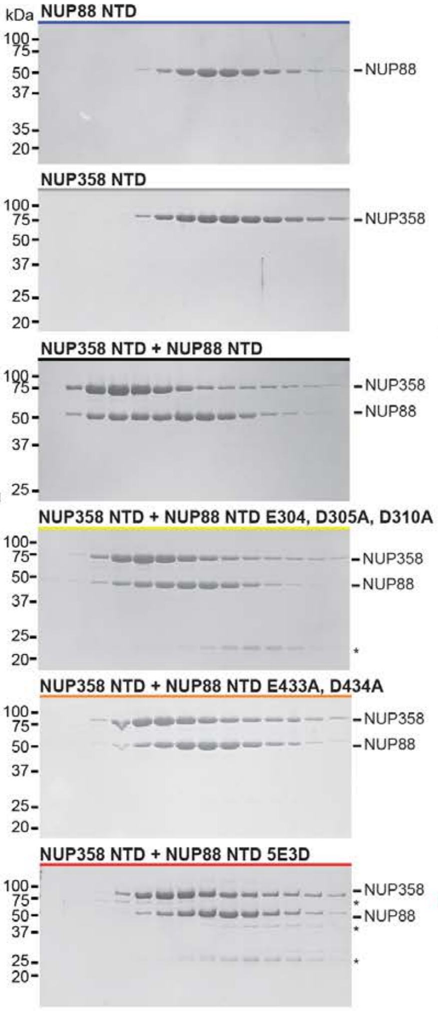

**B**

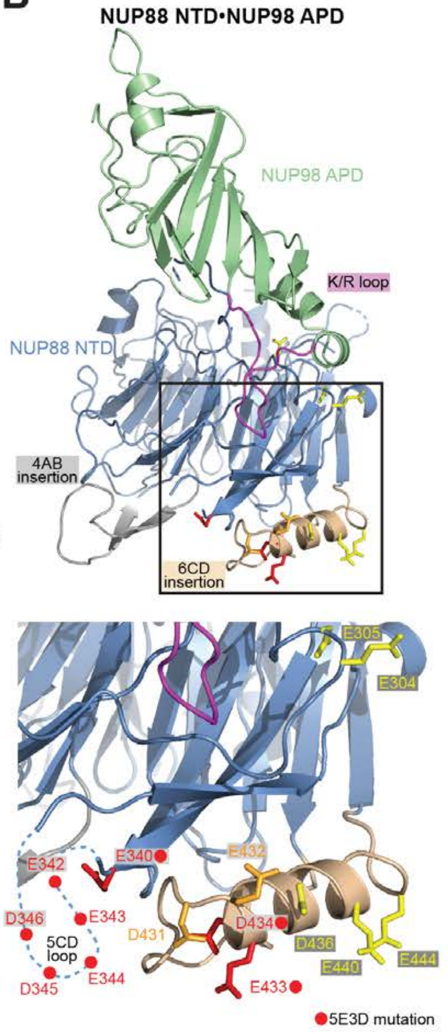

**Fig. S80. Identification of the NUP88$^{NTD}$ 5E3D mutation interrupting NUP358$^{NTD}$ binding.** (**A**) SEC interaction analyses of the indicated NUP88$^{NTD}$ mutants with NUP358$^{NTD}$. Stacked preincubation curves are shown grouped by effect category; no effect (green, front), mild effect (yellow), moderate effect (orange), and severe effect (red) on NUP358$^{NTD}$ binding. SDS-PAGE gels of representative preincubation peak fractions are shown. (**B**) Structure of NUP88$^{NTD}$•NUP98$^{APD}$ shown in cartoon representation. Residues that affect NUP358$^{NTD}$ binding are shown in ball-and-stick representation. The inset marks the location of the mutation that affects NUP358$^{NTD}$ binding. A dashed line indicates the approximate location of the unresolved acidic 5CD loop. Red circles indicate alanine substitutions in the 5E3D mutant.

A

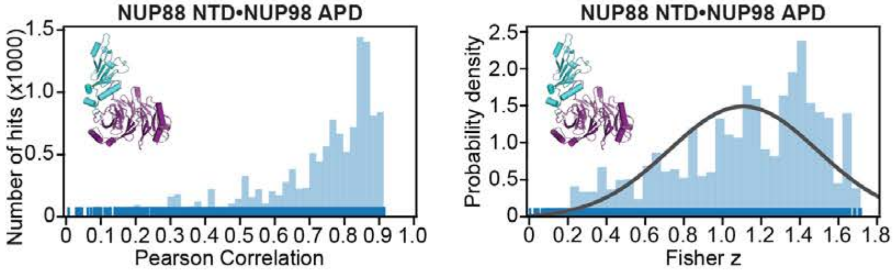

**Subtracted cryo-ET map fitting statistics**

| solution rank | Pearson correlation | Fisher z | p-value |
|---|---|---|---|
| 1 | 0.908 | 1.51 | 0.057 |
| 2 | 0.907 | 1.51 | 0.058 |
| 3 | 0.903 | 1.49 | 0.066 |
| 4 | 0.898 | 1.46 | 0.076 |
| 5 | 0.898 | 1.46 | 0.078 |

B

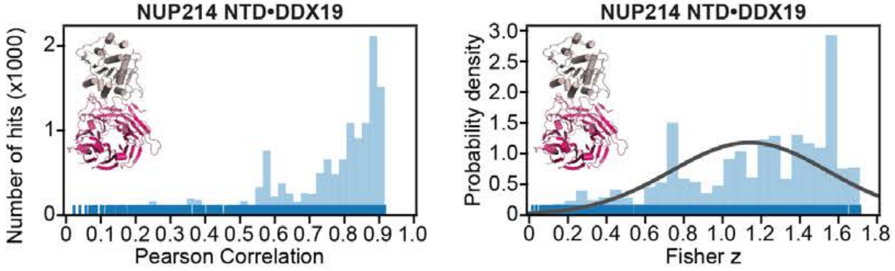

**Subtracted cryo-ET map fitting statistics**

| solution rank | Pearson correlation | Fisher z | p-value |
|---|---|---|---|
| 1 | 0.907 | 1.51 | 0.079 |
| 2 | 0.907 | 1.51 | 0.079 |
| 3 | 0.906 | 1.51 | 0.081 |
| 4 | 0.904 | 1.50 | 0.086 |
| 5 | 0.902 | 1.48 | 0.092 |

C

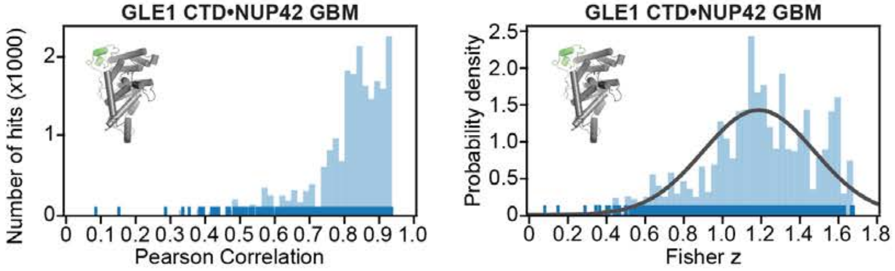

**Subtracted cryo-ET map fitting statistics**

| solution rank | Pearson correlation | Fisher z | p-value |
|---|---|---|---|
| 1 | 0.931 | 1.66 | 0.045 |
| 2 | 0.930 | 1.65 | 0.048 |
| 3 | 0.925 | 1.62 | 0.062 |
| 4 | 0.923 | 1.61 | 0.066 |
| 5 | 0.921 | 1.60 | 0.073 |

D

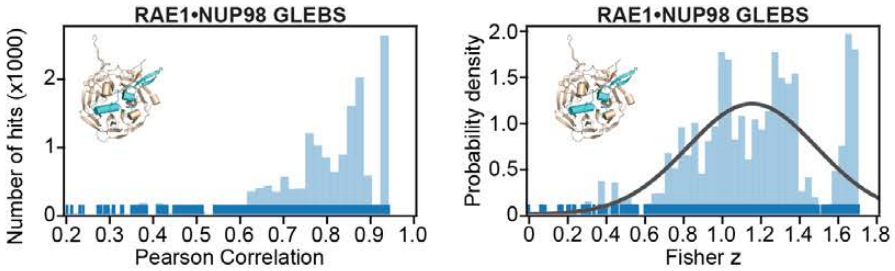

**Subtracted cryo-ET map fitting statistics**

| solution rank | Pearson correlation | Fisher z | p-value |
|---|---|---|---|
| 1 | 0.935 | 1.69 | 0.050 |
| 2 | 0.933 | 1.68 | 0.054 |
| 3 | 0.932 | 1.68 | 0.055 |
| 4 | 0.932 | 1.67 | 0.057 |
| 5 | 0.932 | 1.67 | 0.058 |

E

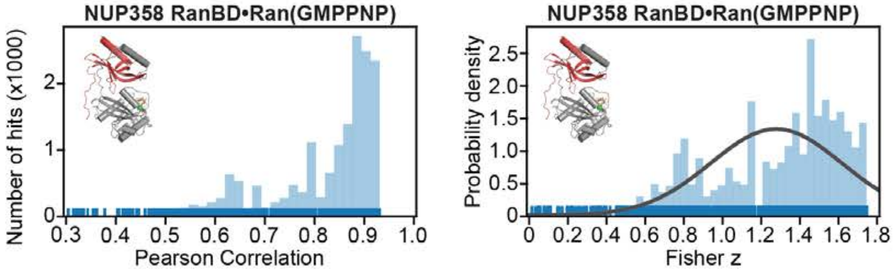

**Subtracted cryo-ET map fitting statistics**

| solution rank | Pearson correlation | Fisher z | p-value |
|---|---|---|---|
| 1 | 0.927 | 1.64 | 0.083 |
| 2 | 0.927 | 1.64 | 0.084 |
| 3 | 0.925 | 1.62 | 0.093 |
| 4 | 0.923 | 1.61 | 0.098 |
| 5 | 0.922 | 1.61 | 0.099 |

F

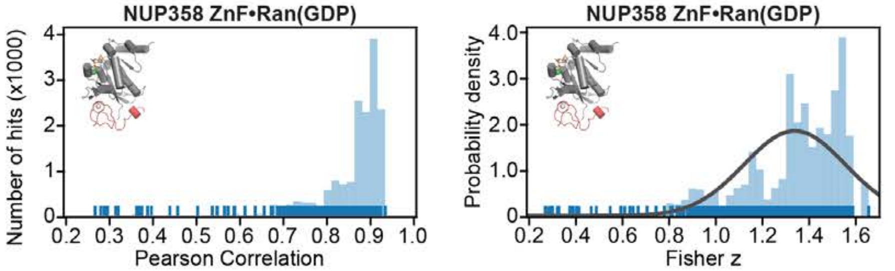

**Subtracted cryo-ET map fitting statistics**

| solution rank | Pearson correlation | Fisher z | p-value |
|---|---|---|---|
| 1 | 0.929 | 1.65 | 0.072 |
| 2 | 0.920 | 1.59 | 0.124 |
| 3 | 0.919 | 1.59 | 0.126 |
| 4 | 0.919 | 1.58 | 0.129 |
| 5 | 0.919 | 1.58 | 0.129 |

G

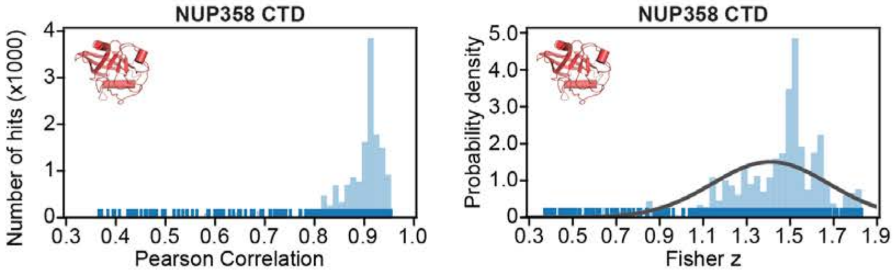

**Subtracted cryo-ET map fitting statistics**

| solution rank | Pearson correlation | Fisher z | p-value |
|---|---|---|---|
| 1 | 0.950 | 1.83 | 0.058 |
| 2 | 0.949 | 1.82 | 0.066 |
| 3 | 0.948 | 1.81 | 0.068 |
| 4 | 0.947 | 1.80 | 0.075 |
| 5 | 0.946 | 1.80 | 0.076 |

**Sup Fig. S81**

**Fig. S81. Quantitative docking searches for CF subunits.** Resolution-matched densities simulated from the crystal structures of (**A**) $NUP88^{NTD}$•$NUP98^{APD}$, (**B**) $NUP214^{NTD}$•DDX19 (PDB ID 3FMO) (*55*), (**C**) $GLE1^{CTD}$•$NUP42^{GBM}$ (PDB ID 6B4F) (*63*), (**D**) RAE1•$NUP98^{GLEBS}$ (PDB ID 3MMY) (*57*), (**E**) $NUP358^{RanBD\text{-}II}$•Ran, (**F**) $NUP358^{ZnF3}$•Ran, and (**G**) $NUP358^{CTD}$ (PDB ID 4I9Y) (45) were quantitatively docked into the ~12 Å sub-tomogram averaged cryo-ET map of the intact human NPC (EMD-XXXX) (*46*) from which cryo-ET density corresponding to all placed nucleoporins was subtracted. Rug plots (blue) and histograms (light blue) of Pearson correlation scores (*left*) and derived Fisher z scores fit with a normalized Gaussian curve (black) from a global search with 1 million random initial placements (*middle*) are shown. A tabular summary of the solutions fitting statistics, along with one-tailed p-values calculated from the Fisher z score distribution, is shown (*right*). No docking solution segregated to high confidence.

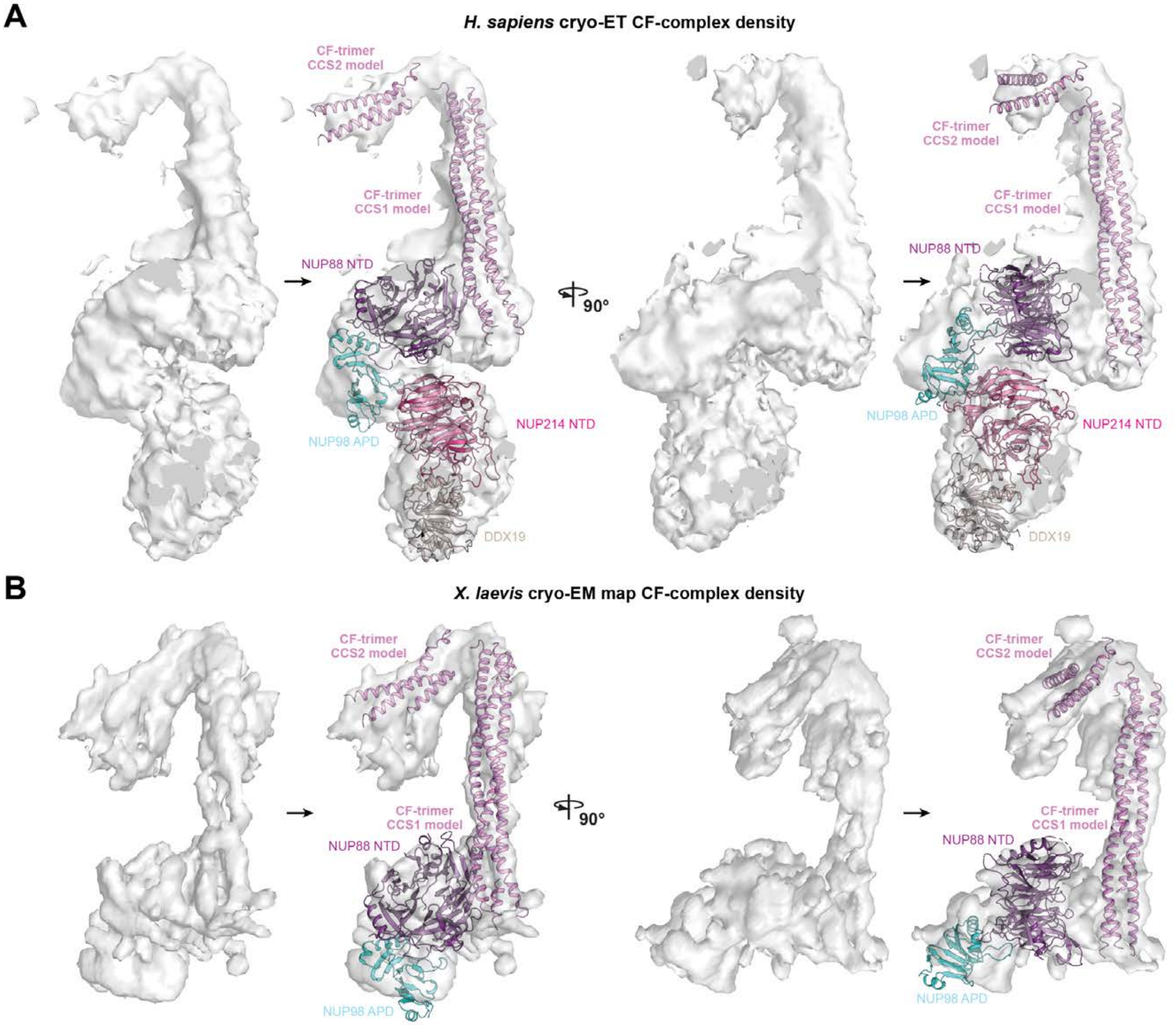

**Sup Fig. S82**

Hoelz et al., 2021b

**Fig. S82. Docking and modeling of the *H. sapiens* and *X. laevis* CFNC complex.** The CFNC complex was modeled by manually docking and rigid body refining structures against (**A**) the ~12 Å cryo-ET map of the intact human NPC (EMD-XXXX) (*46*) and (**B**) a ~8 Å region of the anisotropic composite single particle cryo-EM map of the *X. laevis* cytoplasmic outer ring protomer (EMD-0909) (*45*). In lieu of an experimental structure of the CFNC-hub coiled-coils, poly-alanine models of the CFNC-hub coiled-coil segments CCS1 and CCS2 (pink) were derived from the *X. laevis* NUP54•NUP58•NUP62 (PDB ID 5C3L) (*10*) and *C. thermophilum* Nup49•Nup57•Nsp1 (PDB ID 5CWS) (*11*) complexes and fit into the tube-like densities. $NUP88^{NTD}$•$NUP98^{APD}$ (purple and cyan) was placed in a disk-like density at the base of the long $CFNC\text{-}hub^{CCS1}$ structure. The dumbbell-shaped density assigned to $NUP214^{NTD}$•DDX19 (bright pink and umber) in the human ~12 Å cryo-ET map was not present in the ~8 Å region of the anisotropic composite single particle cryo-EM map of the *X. laevis* cytoplasmic outer ring protomer. Cryo-EM and cryo-ET densities were carved out from larger maps for illustrative purposes and rendered as gray isosurface. Structures and models are shown as cartoons.

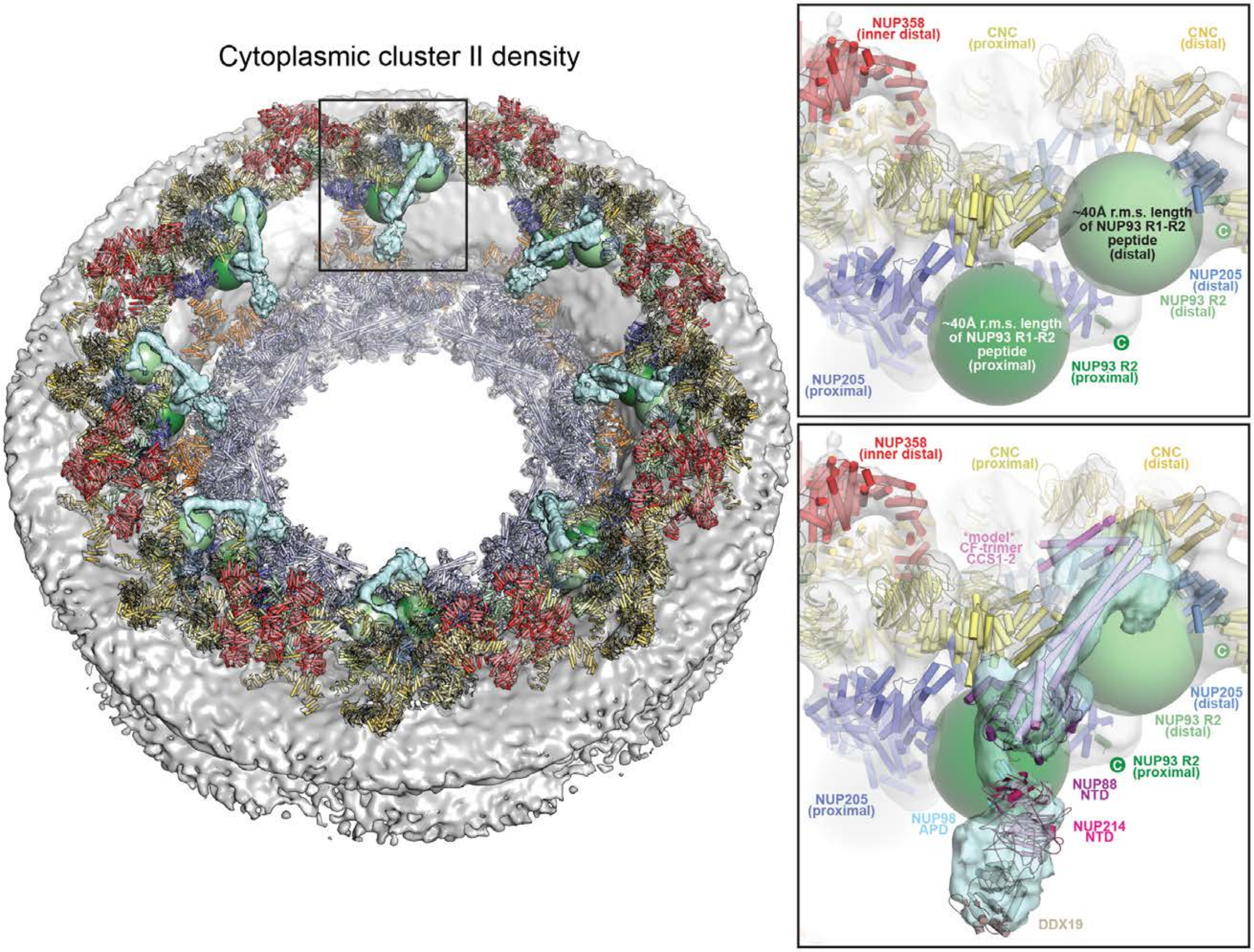

Cytoplasmic cluster II density
NUP358
(inner distal)
CNC
(proximal)
CNC
(distal)
~40Å r.m.s. length
of NUP93 R1-R2
peptide
(distal)
~40Å r.m.s. length
of NUP93 R1-R2
peptide
(proximal)
NUP205
(distal)
NUP93 R2
(distal)
NUP93 R2
(proximal)
NUP205
(proximal)
NUP358
(inner distal)
CNC
(proximal)
CNC
(distal)
*model*
CF-trimer
CCS1-2
NUP205
(distal)
NUP93 R2
(distal)
NUP93 R2
(proximal)
NUP88
NTD
NUP205
(proximal)
NUP98
APD
NUP214
NTD
DDX19

**Fig. S83. NUP93$^{R1\text{-}R2}$ peptide can span the distance between NUP205 and the CFNC-hub binding sites.** Composite structure of the human NPC shown as cartoon. The nuclear envelope (gray) and unassigned density cluster II, corresponding to the CFNC (cyan), are shown as isosurfaces (*left*). Root mean square (r.m.s.) end-to-end length estimates for the NUP93$^{R1\text{-}R2}$ peptide that links the NUP93$^{R2}$ region that binds to NUP205 and the NUP93$^{R1}$ region that binds to the CFNC-hub, are shown as spheres (proximal, green; distal, pale green). Magnified view of the region in the inset box shown in the absence (*right top*) and presence (*right bottom*) of the unassigned density cluster II and the CFNC-hub model docked in it. The distal NUP93$^{R1}$ region is expected in the vicinity of CFNC-hub coiled-coil segments CCS1, CCS2, and likely of the unresolved CCS3.

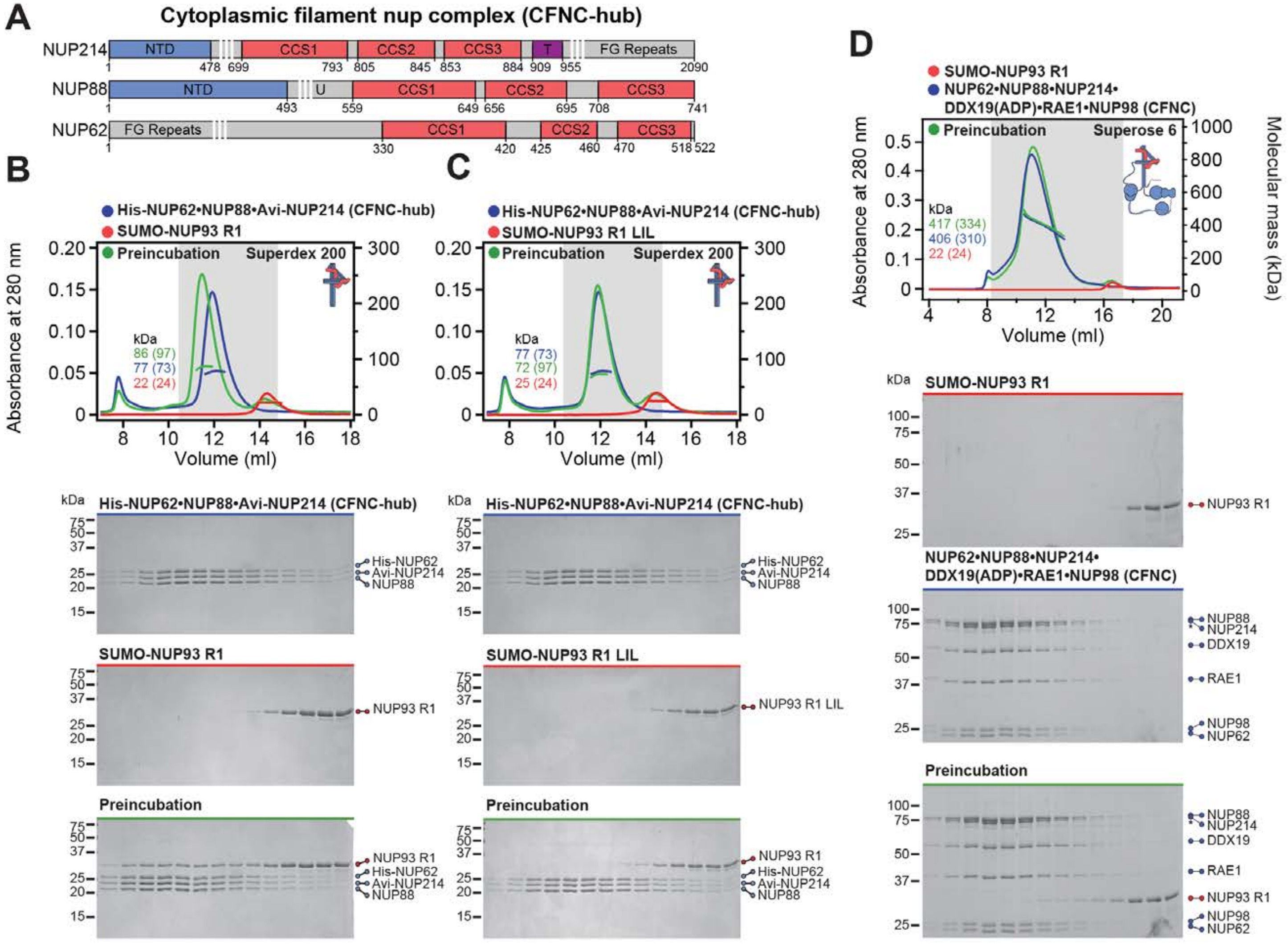

A
Cytoplasmic filament nup complex (CFNC-hub)
NUP214
NUP88
NUP62
B
His-NUP62•NUP88•Avi-NUP214 (CFNC-hub)
SUMO-NUP93 R1
Preincubation
Superdex 200
C
SUMO-NUP93 R1 LIL
D
NUP62•NUP88•NUP214•DDX19(ADP)•RAE1•NUP98 (CFNC)
Superose 6
Absorbance at 280 nm
Molecular mass (kDa)
Volume (ml)

**Fig. S84. NUP93 anchors the human CFNC to the NPC.** (**A**) Domain architectures of NUP214, NUP88, and NUP62 showing the predicted coiled-coil segments, CCS1, CCS2, and CSCS3, which form the CFNC-hub. (**B**) SEC-MALS interaction analysis of the human CFNC-hub with SUMO-NUP93$^{R1}$, demonstrating the formation of a stable stochiometric complex. (**C**) SEC-MALS interaction analyses of the human CFNC-hub with the NUP93$^{R1}$ LIL combination mutant, which we identify in the accompanying manuscript to abolish the interaction with the NUP54•NUP58•NUP62 channel nup hetero-trimer (CNT) (*42*), showing that the NUP93$^{R1}$ LIL mutant also abolishes the interaction with the human CFNC-hub. To resolve the three stoichiometric protein bands of the human CFNC-hub by SDS-PAGE, NUP62$^{CCS1\text{-}3}$ and NUP214$^{CCS1\text{-}3}$ were tagged with $His_6$-PreScission- and Avi-tag sequence overhangs, respectively. SEC interactions with the untagged proteins yielded identical results. (**D**) SEC-MALS interaction analysis of NUP93$^{R1}$ with human hetero-hexameric CFNC. SEC-MALS profiles of nups and nup complexes are shown individually (blue and red) and after their preincubation (green). SEC profiles were obtained using a Superdex 200 10/300 GL or a Superose 6 10/300 GL column, as indicated. Measured molecular masses are indicated, with the respective theoretical masses shown in parentheses. Inset schematics display tested protein samples colored to match the corresponding chromatogram trace. Gray boxes indicate fractions resolved on SDS-PAGE gels and visualized by Coomassie brilliant blue staining.

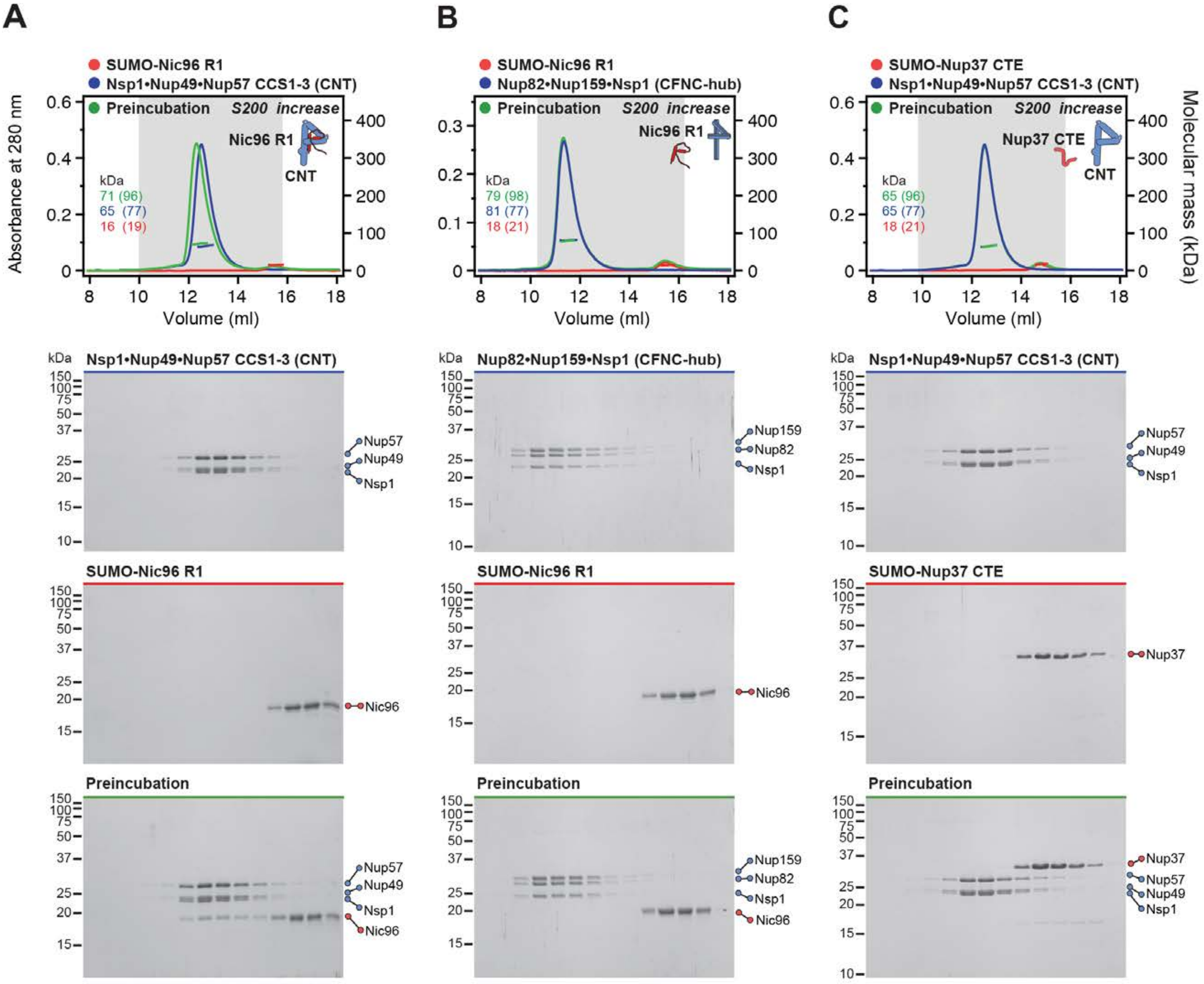
A
SUMO-Nic96 R1
Nsp1•Nup49•Nup57 CCS1-3 (CNT)
Preincubation
S200 increase
Nic96 R1
CNT
kDa
71 (96)
65 (77)
16 (19)
Absorbance at 280 nm
Volume (ml)
B
SUMO-Nic96 R1
Nup82•Nup159•Nsp1 (CFNC-hub)
Preincubation
S200 increase
Nic96 R1
kDa
79 (98)
81 (77)
18 (21)
C
SUMO-Nup37 CTE
Nsp1•Nup49•Nup57 CCS1-3 (CNT)
Preincubation
S200 increase
Nup37 CTE
CNT
kDa
65 (96)
65 (77)
18 (21)
Molecular mass (kDa)
Nsp1•Nup49•Nup57 CCS1-3 (CNT)
Nup82•Nup159•Nsp1 (CFNC-hub)
Nup57
Nup49
Nsp1
Nup159
Nup82
SUMO-Nic96 R1
SUMO-Nup37 CTE
Nic96
Nup37
Preincubation

**Fig. S85. Nic96 and Nup37 assembly sensors possess binding specificity for CNT and CFNC-hub.** SEC-MALS interaction analyses of (**A**) SUMO-Nic96$^{R1}$ with CNT, (**B**) SUMO-Nic96$^{R1}$ with CFNC-hub, and (**C**) SUMO-Nup37$^{CTE}$ with CNT. Whereas Nup37$^{CTE}$ exclusively binds the CFNC-hub, Nic96$^{R1}$ exclusively binds to the CNT, and not vice versa. SEC profiles of nups or nup complexes are shown individually (blue and red) and after their preincubation (green). SEC profiles were obtained using a Superdex 200 Increase 10/300 GL column. Measured molecular masses are indicated, with the respective theoretical masses shown in parentheses. Inset schematics display tested protein samples shaded to match corresponding chromatogram trace. Gray boxes indicate fractions resolved on SDS-PAGE gels and visualized by Coomassie brilliant blue staining.

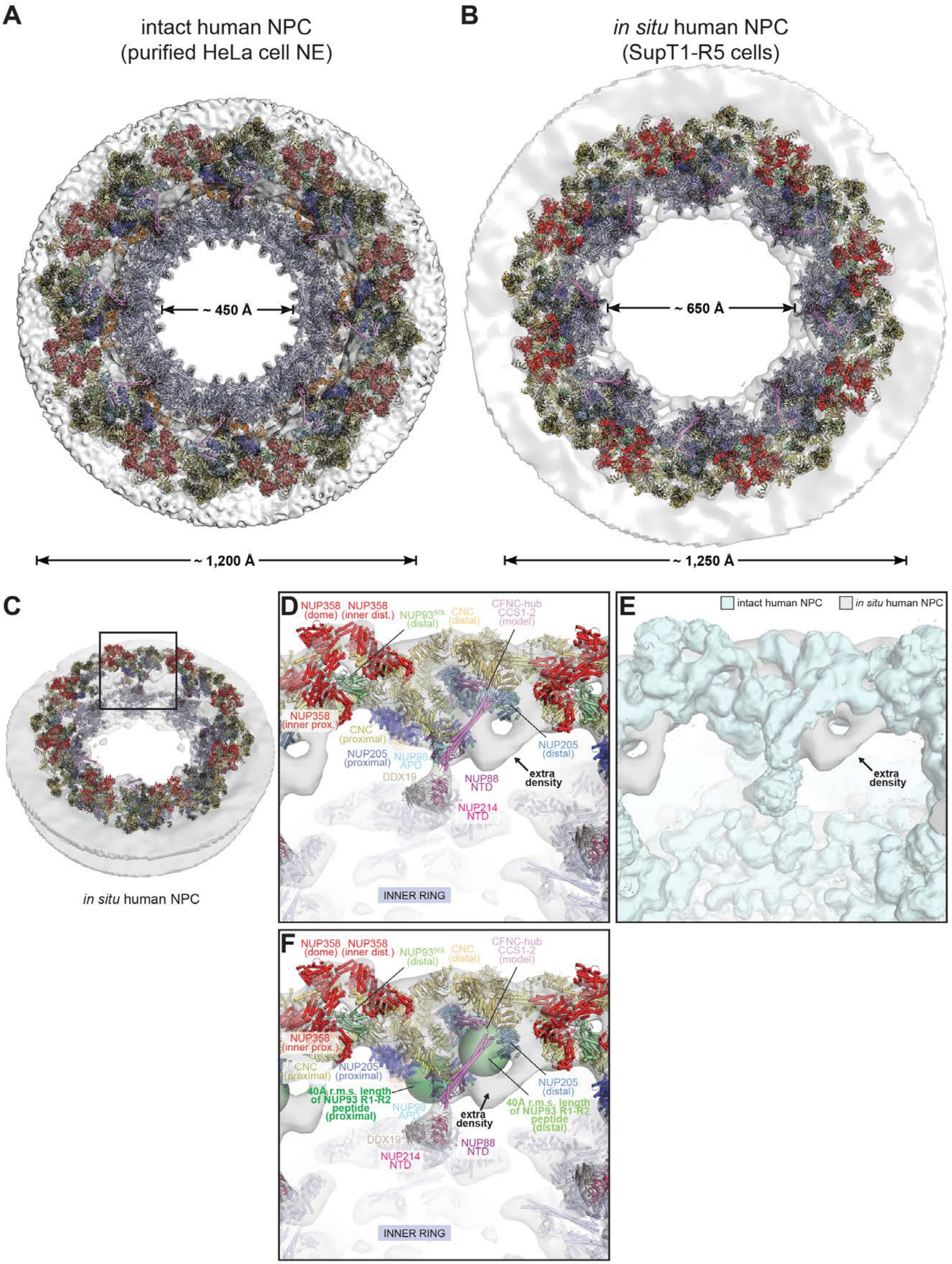
A
intact human NPC
(purified HeLa cell NE)
B
in situ human NPC
(SupT1-R5 cells)
~ 450 Å
~ 650 Å
~ 1,200 Å
~ 1,250 Å
C
in situ human NPC
D
NUP358 (dome)
NUP358 (inner dist.)
NUP93SOL (distal)
CNC (distal)
CFNC-hub CCS1-2 (model)
NUP358 (inner prox.)
CNC (proximal)
NUP205 (proximal)
NUP98 APD
NUP205 (distal)
extra density
DDX19
NUP88 NTD
NUP214 NTD
INNER RING
E
intact human NPC
in situ human NPC
extra density
F
NUP358 (dome)
NUP358 (inner dist.)
NUP93SOL (distal)
CNC (distal)
CFNC-hub CCS1-2 (model)
NUP358 (inner prox.)
CNC (proximal)
NUP205 (proximal)
40Å r.m.s. length of NUP93 R1-R2 peptide (proximal)
NUP205 (distal)
40Å r.m.s. length of NUP93 R1-R2 peptide (distal)
NUP98 APD
extra density
DDX19
NUP88 NTD
NUP214 NTD
INNER RING

**Fig. S86 Additional density in unassigned density cluster II from dilated *in situ* human NPC.** (**A**, **B**) Comparison of the cytoplasmic face of (A) the constricted human NPC imaged in purified HeLa cell nuclear envelopes and (B) the dilated human NPC imaged *in situ* in SupT1-R5 cells. Outer and inner ring spoke subcomplexes from the composite NPC structure built into the constricted ~12 Å cryo-ET sub-tomogram averaged human NPC map (EMD-XXXX) (*46*) were manually docked and locally fit as rigid units in the dilated *in situ* ~37 Å cryo-ET sub-tomogram averaged human NPC map (EMD-11967) (*101*). The composite NPC structures are shown as cartoons and superposed to isosurfaces of the constricted and dilated NPCs (gray). (**C**) A 60º-rotated overview of the dilated human NPC. The inset box indicates the region magnified in the following panels. (**D**) Magnified view of the docked structures and model of the CFNC-hub (lacking CCS3) are shown as cartoons superposed to the cryo-ET map of the dilated human NPC rendered as gray isosurface. (**E**) Magnified view of the cryo-ET map of the dilated human NPC (gray), superposed to the cryo-ET map of the constricted human NPC (cyan). Whereas the density of the constricted human NPC is entirely explained by the composite structure, additional unexplained density adjacent to the assigned CFNC-hub is observed in the map of the dilated human NPC (black arrow). (**F**) Magnified view as in panel (D) with green spheres indicating the root mean square (r.m.s) end-to-end length estimates for the $NUP93^{R1-R2}$ peptide connector, which links $NUP93^{R2}$•NUP205 with the $NUP93^{R1}$•CFNC-hub (proximal, green; distal, pale green). The unassigned rod-shaped density of the cryo-ET map of the dilated human NPC is in close proximity to NUP205•$NUP93^{R2}$, suggesting that the unexplained additional density may accommodate a second CFNC copy. The composite structures in all figure panels are colored according to Fig. 10.

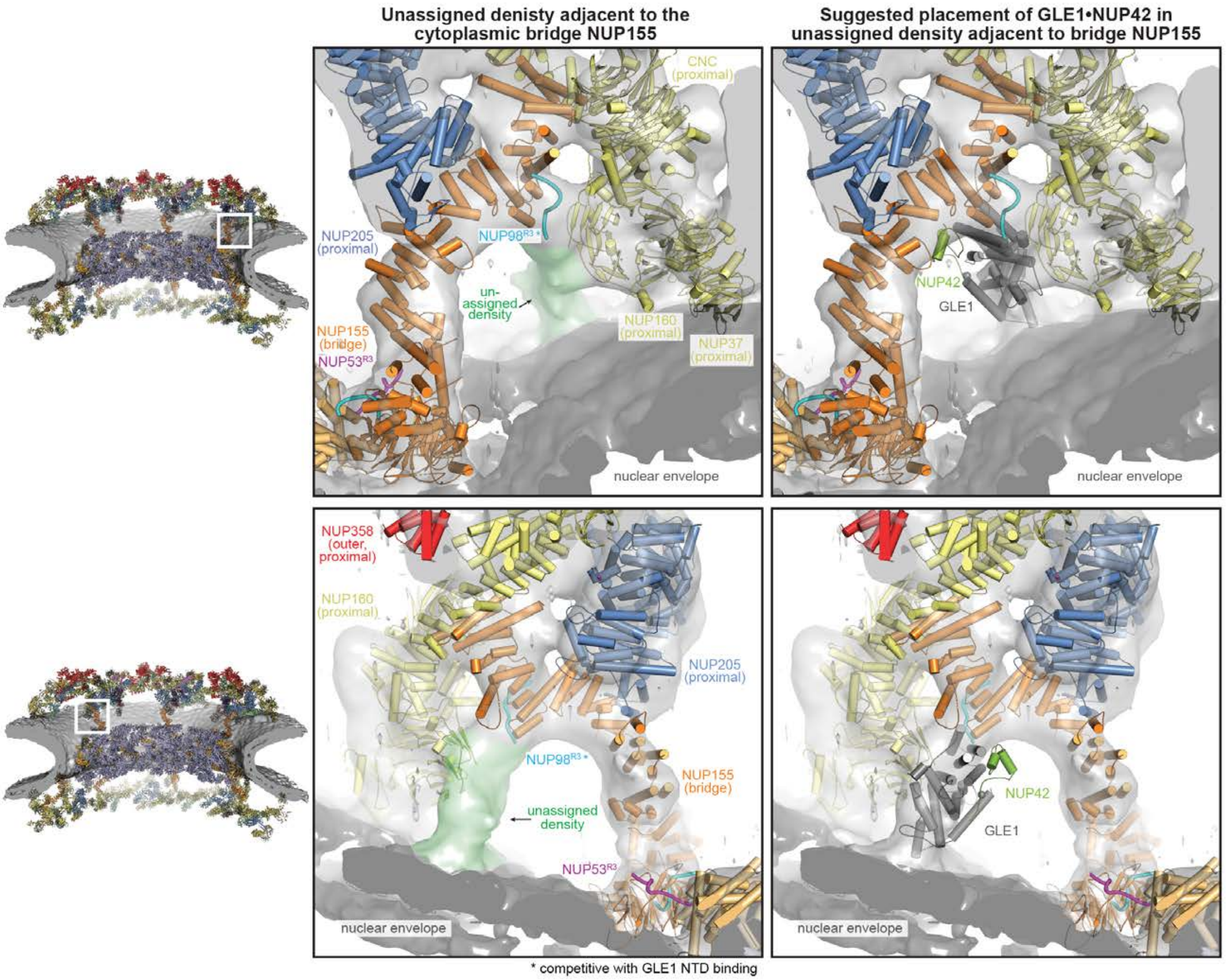

Unassigned denisty adjacent to the cytoplasmic bridge NUP155
Suggested placement of GLE1•NUP42 in unassigned density adjacent to bridge NUP155
CNC (proximal)
NUP205 (proximal)
NUP98[R3] *
un-assigned density
NUP160 (proximal)
NUP37 (proximal)
NUP155 (bridge)
NUP53[R3]
nuclear envelope
NUP42
GLE1
NUP358 (outer, proximal)
unassigned density
* competitive with GLE1 NTD binding

**Fig. S87. Suggested placement of GLE1$^{CTD}$•NUP42.** Overview of the composite structure of the human NPC (*left*). The inset boxes indicate regions magnified in the middle and right panels, harboring unassigned cytoplasmic density directly adjacent to the bridging NUP155 molecules in the ~12 Å cryo-ET map of the intact human NPC (EMD-XXXX) (*46*). Magnified views illustrating the globular shape and environment of the cytoplasmic unexplained density (green) shown from the front and the back (*middle*). As previously proposed (*63*), the manually placed GLE1$^{CTD}$•NUP42 structure (PDB ID 6B4F) accounts for the unexplained density and is consistent with previous biochemical analyses showing that GLE1$^{NTD}$ and NUP98$^{R3}$ bind to NUP155 in a mutually exclusive fashion (*right*). GLE1$^{CTD}$•NUP42 is shown in cartoon representation and colored in dark gray and green. The quantitatively docked nups in close proximity to the unassigned density are shown in cartoon representation and colored according to Fig. 10. Cryo-ET densities corresponding to protein and nuclear envelope are rendered as gray and dark gray isosurfaces, respectively.

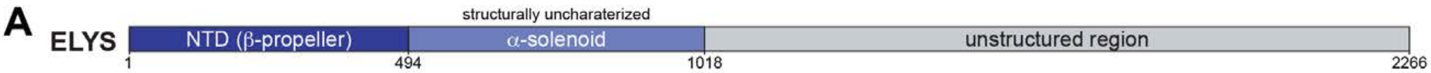

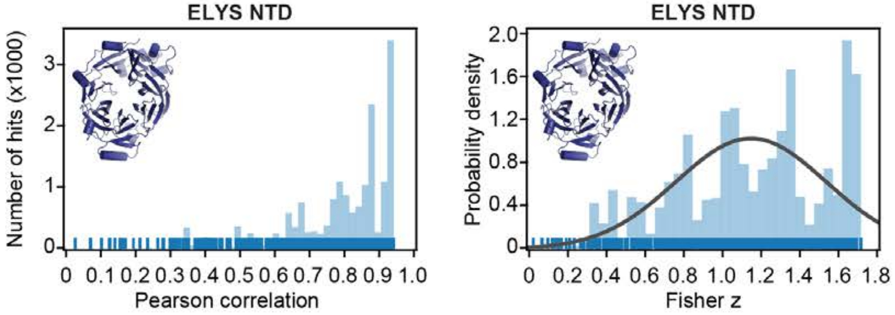

**Subtracted cryo-ET map fitting statistics**

| solution rank | Pearson correlation | Fisher z | p-value |
|---|---|---|---|
| 1 | 0.936 | 1.71 | 0.072 |
| 2 | 0.933 | 1.68 | 0.080 |
| 3 | 0.933 | 1.68 | 0.081 |
| 4 | 0.933 | 1.68 | 0.082 |
| 5 | 0.932 | 1.67 | 0.085 |

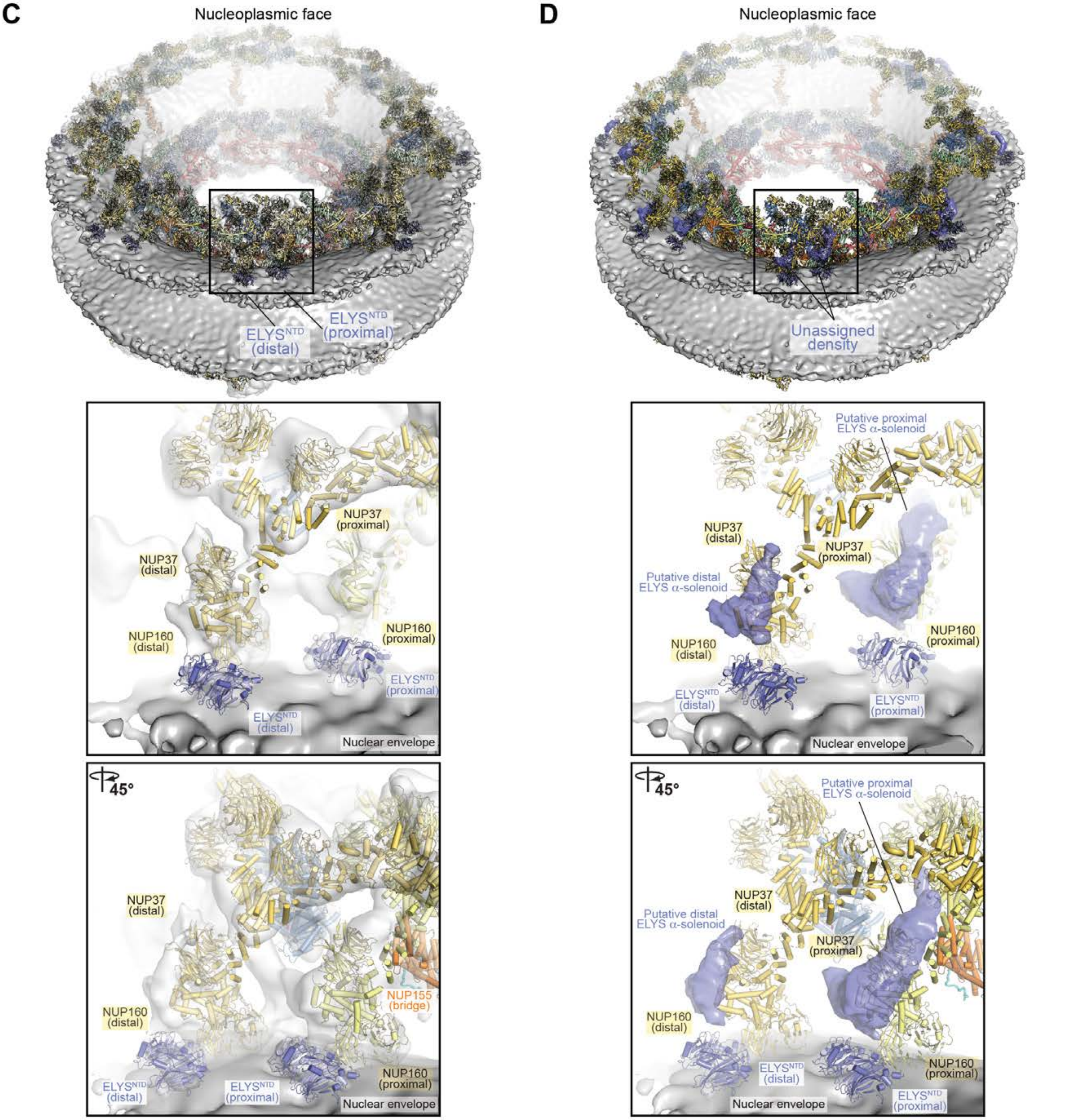

**Fig. S88. ELYS placement on nuclear face of the NPC.** (**A**) Domain architecture of human ELYS with residue numbers indicated. (**B**) Resolution-matched densities simulated from the *M. musculus* $ELYS^{NTD}$ crystal structure (PDB ID 4I0O) (*25*) were quantitatively docked into the ~12 Å sub-tomogram averaged cryo-ET map of the intact human NPC (EMD-XXXX) (*46*) from which cryo-ET density corresponding to all placed nups was subtracted. Rug plots (blue) and histograms (light blue) of Pearson correlation scores (*left*) and derived Fisher z scores fit with a normalized Gaussian curve (black) from a global search with 1 million random initial placements are shown (*middle*). A tabular summary of the solutions fitting statistics, along with one-tailed p-values calculated from the Fisher z score distribution (*right*). No docking solution segregated to high confidence. (**C**) Two magnified views (*middle and bottom*) of the region in the inset box (*top*), showing the placement of $ELYS^{NTD}$ copies into disk-shaped density adjacent to nuclear proximal and distal NUP160 copies and the nuclear envelope. Docked structures are shown in cartoon representation. Cryo-ET densities corresponding to protein and nuclear envelope are rendered as gray and dark gray isosurfaces, respectively. (**D**) Views as in (C), with unexplained crescent-shaped densities suggested to correspond to the ELYS $\alpha$-solenoid domain rendered as isosurface (dark blue).

#### **Table S1.** Materials and reagents

| REAGENT or RESOURCE | SOURCE | IDENTIFIER |
|---|---|---|
| ***C. thermophilum* and *H. sapiens* nucleoporin cDNAs** | | |
| Nup82 | This study | N/A |
| Nup159 | This study | N/A |
| Nsp1 | (*11*) | N/A |
| Dbp5 | (*63*) | N/A |
| Nup145N | (*11*) | N/A |
| Gle1 | This study | N/A |
| Gle2 | This study | N/A |
| Nup42 | (*63*) | N/A |
| Nic96 | (*11*) | N/A |
| Nup120 | (*38*) | N/A |
| Nup37 | (*38*) | N/A |
| Elys | (*38*) | N/A |
| Nup85 | (*38*) | N/A |
| Sec13 | (*38*) | N/A |
| Nup145C | (*38*) | N/A |
| Nup84 | (*38*) | N/A |
| Nup133 | (*38*) | N/A |
| NUP358 | (*80*) | N/A |
| NUP88 | Open Biosystems | MHS6278-202826062 |
| NUP214 | (*54*) | N/A |
| NUP62 | Open Biosystems | MHS6278-202806024 |
| NUP98 | Gift, Beatriz Fontoura (*105*) | N/A |
| RAE1 | (*57*) | N/A |
| DDX19 | (*55*) | N/A |
| GLE1 | (*63*) | N/A |
| NUP42 | Gift, Susan Wente (*106*) | N/A |
| NUP155 | Gift, Susan Wente (*106*) | N/A |
| GFP | (*76*) | N/A |
| H1B | (*76*) | N/A |
| Insulin | (*76*) | N/A |
| ALPP | DNASU | HsCD00641203 |
| IL6 | DNASU | HsCD00001047 |
| IL10 | DNASU | HsCD00001333 |
| RPL26 | Horizon | MHS6278-20280 |
| TNF$\alpha$ | DNASU | HsCD00001318 |
| **Molecular biology reagents** | | |
| SV Total RNA Isolation System | Promega | Cat# Z3100 |
| Superscript-III Reverse Transcriptase | Invitrogen | Cat# 18080044 |
| PfuUltra II Fusion HotStart DNA Polymerase | Agilent Technologies | Cat# 600674 |
| Phusion High-Fidelity DNA Polymerase | New England Biolabs | Cat# M0530L |
| AscI | New England Biolabs | Cat# R0558L |
| BamHI | New England Biolabs | Cat# R0136L |
| EcoRI | New England Biolabs | Cat# R0101L |
| HindIII | New England Biolabs | Cat# R0104L |
| KpnI | New England Biolabs | Cat# R0142L |
| NcoI | New England Biolabs | Cat# R0193L |
| NdeI | New England Biolabs | Cat# R0111L |
| NotI | New England Biolabs | Cat# R0189L |
| SalI | New England Biolabs | Cat# R0138L |
| SmaI | New England Biolabs | Cat# R0141L |
| Sbf1 | New England Biolabs | Cat# R3642L |
| XhoI | New England Biolabs | Cat# R0146L |
| T4 DNA Ligase | New England Biolabs | Cat# M0202L |
| QIAquick PCR Purification Kit | Qiagen | Cat# 28104 |
| QIAquick Miniprep Kit | Qiagen | Cat# 27104 |
| NucleoSpin buffer | Clontech Laboratories | Cat# 740953 |

| REAGENT or RESOURCE | SOURCE | IDENTIFIER |
| --- | --- | --- |
| ViaFect transfection reagent | Promega | Cat# E4982 |
| Nucleic acid miniprep columns | VitaScientific | Cat# DBOC50009 |
| Agarose | Invitrogen | Cat# 16500500 |
| **Bacterial and *C. thermophilum* culture reagents** | | |
| Dextrin heptahydrate | Fisher BioReagents | Cat# BP9725-5 |
| Peptone | Fisher BioReagents | Cat# BP9726-5 |
| Yeast extract | Fisher BioReagents | Cat# BP9727-5 |
| Luria-Bertani (LB) media | Fisher BioReagents | Cat# BP9723-5 |
| 2×YT media | Fisher BioReagents | Cat# BP97435 |
| Isopropyl-β-D-thiogalactopyranoside (IPTG) | Gold Biotechnology | Cat# 12481C-1KG |
| Ampicillin | Gold Biotechnology | Cat# A-301-100 |
| Kanamycin | Gold Biotechnology | Cat# K-120-100 |
| Chloramphenicol | Gold Biotechnology | Cat# C-105-100 |
| Dipotassium hydrogen phosphate trihydrate | Macron | Cat# 7088-06 |
| Iron (III) sulfate | Sigma-Aldrich | Cat# 307718 |
| Magnesium sulfate | Sigma-Aldrich | Cat# M2643 |
| Seleno-L-methionine (SeMet) | TCL | Cat# S0442 |
| **Insect cell culture reagents** | | |
| 5-bromo-3-indolyl-β-D-galactopyranoside (BluoGal) | Sigma-Aldrich | Cat# 97753-82-7 |
| Sf-900 III SFM (1x) | Gibco | Cat# 12658-019 |
| ESF 921 insect cell culture medium | Expression Systems | Cat# 96-001-01 |
| Production boost additive | Expression Systems | Cat# 95-006-100 |
| Cellfectin II reagent | Gibco | Cat# 10362-100 |
| **Protein purification reagents / general chemicals** | | |
| TRIS (trometamol, 2-amino-2-(hydroxymethyl)-1,3-propanediol) | Sigma-Aldrich | Cat# 93362 |
| MES (2-(N-morpholino)ethanesulfonic acid) | Gold Biotechnology | Cat# M-090-500 |
| Sodium chloride | Sigma-Aldrich | Cat# 71376-5KG |
| Potassium phosphate | Sigma-Aldrich | Cat# 60353-1KG |
| Zinc chloride | Fluka | Cat# 96468 |
| Magnesium chloride | Fluka | Cat# 63068 |
| Imidazole | Sigma-Aldrich | Cat# 56750 |
| Glutathione | Sigma-Aldrich | Cat# G4251 |
| Glycerol | Fisher Scientific | Cat# BP229-1 |
| 2-Mercaptoethanol (β-ME) | Sigma-Aldrich | Cat# M6250 |
| Dithiothreitol (DTT) | Gold Biotechnology | Cat# DTT100 |
| Aprotinin from bovine lung | Sigma-Aldrich | Cat# A6279 |
| Phenylmethylsulfonyl fluoride (PMSF) | Gold Biotechnology | Cat# P-470-50 |
| EDTA-free protease inhibitor cocktail | Roche | Cat# A32965 |
| DNase I | Sigma-Aldrich | Cat# D4527 |
| Inositol hexaphosphate ($IP_6$) | Sigma-Aldrich | Cat# P8810-25G |
| GMPPNP (5'-guanosine-(β,γ-imido)-triphosphate) | Jena Bioscience | Cat# NU-401-50 |
| AMPPNP (Adenosine 5′-(β,γ-imido)-triphosphate lithium salt hydrate) | Sigma-Aldrich | Cat# A2647 |
| Biotin | Pierce | Cat# 21335 |
| Ethylenediaminetetraacetic acid (EDTA) | Sigma-Aldrich | Cat# 03620-1KG |
| Ethanol | Sigma-Aldrich | Cat# 459844 |
| Alkaline phosphatase | Roche | Cat# 10 567 744 001 |
| Ubl-specific protease 1(ULP1) | Invitrogen | Cat# 12588018 |
| PreScission protease (Pres) | Sigma-Aldrich | Cat# GE27-0843-01 |
| Tabaco etch virus protease (TEV) | Sigma-Aldrich | Cat# T4455 |
| **Synthetic antibodies reagents** | | |
| M13-KO7 helper phage | New England Biolabs | Cat# N0315S |
| 96 well high binding microlon plates | Greiner Bio-one | Cat# 655081 |
| Neutravidin | Thermo Fisher Scientific | Cat# 34021 |
| Streptavidin MagneSphere | Promega | Cat# Z548C |
| 3,3′,5,5′-tetramethylbenzidine (TMB) substrate | Pierce | Cat# 34021 |
| BSA | Sigma-Aldrich | Cat# A9418-100G |
| Tween-20 | Anatrace | Cat# T1003 |

| REAGENT or RESOURCE | SOURCE | IDENTIFIER |
|---|---|---|
| **SDS-PAGE reagents** | | |
| 40% Acrylamide/Bis solution (37.5:1) | Bio-Rad | Cat# 1610148 |
| TEMED | Bio-Rad | Cat# 161-0801 |
| Glycine | Gold Biotechnology | Cat# G-630-5 |
| Sodium dodecyl sulfate | Sigma-Aldrich | Cat# L3771-1KG |
| Ammonium persulfate | Sigma-Aldrich | Cat# 09913-100g |
| Tricine | Gold Biotechnology | Cat# T-870-1 |
| Acetic acid | Sigma-Aldrich | Cat# 695092 |
| Brilliant Blue R | Sigma-Aldrich | Cat# B7920-50G |
| **LLPS interaction assay reagents** | | |
| Bodipy (boron-dipyrromethene) FL NHS Ester (succinimidyl ester) | Invitrogen | Cat# D2184 |
| 7-Hydroxycoumarin-3carboxylic acid NHS ester (succinimidyl ester) | Sigma-Aldrich | Cat# 55156 |
| Alexa Fluor 647 NHS Ester (succinimidyl ester) | Invitrogen | Cat# A37573 |
| Sodium bicarbonate | Sigma-Aldrich | Cat# S5761 |
| **RNA reagents** | | |
| T7 RNA polymerase | Invitrogen | Cat# AM1334 |
| T4 polynucleotide kinase | New England Biolabs | Cat# M0201L |
| Calf intestinal alkaline phosphatase | New England Biolabs | Cat# M0290 |
| Inorganic pyrophosphatase from baker’s yeast | Sigma-Aldrich | Cat# I1643 |
| Uridine 5’-Triphosphate trisodium salt | Sigma-Aldrich | Cat# U6625 |
| Cytidine 5’-Triphosphate disodium salt | Sigma-Aldrich | Cat# C1506 |
| Guanosine 5’-Triphosphate sodium salt | Sigma-Aldrich | Cat# G8877 |
| Adenosine 5’-Triphosphate disodium salt | Sigma-Aldrich | Cat# A7699 |
| $^{32}$P $\gamma$-ATP (3000Ci/mmol, 10mCi/ml) | Perkin Elmer | Cat# BLU002A250UC |
| Acid-phenol:chloroform:IAA (125:124:1) | Life technologies | cat# AM9722 |
| Spermidine-HCl | Sigma-Aldrich | Cat# 85578 |
| SUPERase-In | Invitrogen | Cat# AM2694 |
| SYBR Gold stain | Invitrogen | Cat# 1730968 |
| 10 x NEB Buffer 4 | New England Biolabs | Cat# B7004S |
| 10 x PNK Buffer | New England Biolabs | Cat# B0201S |
| Zeba Spin Desalting Columns | Thermo Scientific | Cat# 89882 |
| **Crystallization reagents** | | |
| Ammonium sulfate | MP Bio | Cat# 808211 |
| Bicine | Sigma-Aldrich | Cat# 14871 |
| Bis-TRIS | Sigma-Aldrich | Cat# 14879 |
| Citric acid | Sigma-Aldrich | Cat# 27487 |
| HEPES | Sigma-Aldrich | Cat# 54457-250G-F |
| Magnesium formate | Fluka | Cat# 793 |
| PEG 2,000 MME | Sigma-Aldrich | Cat# 202509 |
| PEG 3,350 | Sigma-Aldrich | Cat# 88276 |
| PEG 4,000 | Sigma-Aldrich | Cat# 95904-1kG-F |
| Potassium sodium tartrate tetrahydrate | Sigma-Aldrich | Cat# S6170 |
| Sodium acetate | Sigma-Aldrich | Cat# S3272 |
| Sodium citrate | Sigma-Aldrich | Cat# 71405 |
| Sodium formate | Sigma-Aldrich | Cat# 71539 |
| Succinic acid | Sigma-Aldrich | Cat# 14079 |
| 100% glycerol | Sigma-Aldrich | Cat# 49767 |
| 100% Ethylene glycol | Fluka | Cat# 03750 |
| Paratone oil | Hampton Research | Cat# HR2-043 |
| **Cell culture reagents** | | |
| McCoy’s 5A medium | ATCC | Cat# 30-2007 |
| Dulbecco’s Modified Eagle Medium (DMEM) | Gibco | Cat# 11965-092 |
| Fetal bovine serum (FBS) | Gibco | Cat# 16000044 |
| Penicillin and streptomycin | Gibco | Cat# 15140-122 |
| Hygromycin | Gibco | Cat# 10687010 |
| 0.25% trypsin/0.05% EDTA | Gibco | Cat# 25200-056 |
| Lipofectamine 2000 | Sigma-Aldrich | Cat# 25988-63-0 |

| REAGENT or RESOURCE | SOURCE | IDENTIFIER |
|---|---|---|
| Opti-MEM | Gibco | Cat# 3185-062 |
| Poly-L-lysine | Sigma-Aldrich | Cat# P8920 |
| Paraformaldehyde | Polysciences | Cat# 04018 |
| TritonX-100 | Thermo-Fisher | Cat# 041092 |
| Saponin | Sigma-Aldrich | Cat# 47036-50G-F |
| Nocodazole | Sigma-Aldrich | Cat# M1404-10MG |
| 3-Indoleacetic acid (auxin) | Sigma-Aldrich | Cat# I3750 |
| DMSO | Sigma-Aldrich | Cat# D2650-5 x 5ML |
| Sucrose | Sigma-Aldrich | Cat# 821721 |
| Calcium dicholoride | Sigma-Aldrich | Cat# 529575 |
| Superblock Blocking Buffer | Thermo-Fisher | Cat# 37517 |
| Immobilon P transfer membrane | Millipore | Cat# IPVH00010 |
| Click-iT RNA Alexa Fluor 594 Imaging Kits | Invitrogen | Cat# C10330 |
| **Commercial crystallization screens** | | |
| Crystal Screen | Hampton Research | Cat# HR2-110 |
| Crystal Screen 2 | Hampton Research | Cat# HR2-112 |
| Index Screen | Hampton Research | Cat# HR2-144 |
| PEG/Ion Screen | Hampton Research | Cat# HR2-126 |
| PEG/Ion 2 Screen | Hampton Research | Cat# HR2-098 |
| ProPlex Screen | Molecular Dimensions | Cat# BN043 |
| PEGs Suite | Qiagen | Cat# 130704 |
| PEGs II Suite | Qiagen | Cat# 130716 |
| **Antibodies** | | |
| Rat monoclonal anti-NUP98 | Abcam | Cat# ab50610;RRID:AB_881769 |
| Rabbit polyclonal anti-NUP214 | Abcam | Cat# ab70497; RRID:AB_1269607 |
| Rabbit polyclonal anti-GLE1 | Abcam | Cat# ab96007; RRID:AB_10678755NUP |
| Rabbit monoclonal Anti-LaminA/C | Abcam | Cat# ab108595; RRID:AB_10866185 |
| Mouse monoclonal anti-TBP | Abcam | Cat# ab51841; RRID:AB_945758 |
| Rabbit polyclonal anti-NUP358 | Bethyl Laboratories | Cat# A301-797A; RRID:AB_1211503 |
| Rabbit polyclonal anti-NUP96 | Bethyl Laboratories | Cat# A301-784A; RRID:AB_1211487 |
| Rabbit polyclonal anti-NUP133 | Bethyl Laboratories | Cat# A302-386A; RRID:AB_1907269 |
| Mouse monoclonal mAb414 antibody | BioLegend | Cat# 902907; RRID:AB_2734672 |
| Mouse monoclonal anti-HA | BioLegend | Covance Cat# MMS-101P; RRID:AB_231467 |
| Mouse monoclonal anti-Phospho-Histone H3 (Ser10) | Millipore | Cat# 05-806; RRID:AB_310016 |
| Mouse monoclonal anti-Histone H3 | Millipore | Cat# 05-499; RRID:AB_309763 |
| Mouse monoclonal anti-NUP88 | Santa Cruz Biotechnology | Cat# sc-365868; RRID:AB_10842170 |
| Mouse monoclonal anti-NUP93 | Santa Cruz Biotechnology | Cat# sc-374399;RRID:AB_10990113 |
| Mouse monoclonal anti-CyclinB1 | Santa Cruz Biotechnology | Cat# sc-245; RRID:AB_627338) |
| Mouse monoclonal anti-CyclinD1 | Santa Cruz Biotechnology | Cat# sc-8396; RRID:AB_627344 |
| Mouse monoclonal anti-CyclinE | Santa Cruz Biotechnology | Cat# sc-247; RRID:AB_627357 |
| Rabbit polyclonal anti-NUP160 | St John's Lab | Cat# STJ94576 |
| Mouse monoclonal anti-FLAG | Sigma-Aldrich | Cat# F1804; RRID:AB_262044 |
| Mouse monoclonal anti-Tubulin | Sigma-Aldrich | Cat# T6199; RRID:AB_477583 |
| Rabbit anti-NUP43 | Gift, Valerie Doye (*142*) | N/A |
| Rabbit anti-NUP205 | Gift, Ulrike Kutay (*143*) | N/A |
| Rabbit anti-ELYS | Gift, Iain Mattaj (*144*) | N/A |
| Mouse monoclonal anti-HA, Alexa Fluor 488 | Invitrogen | Cat#A-21287; RRID:AB_2535829 |
| Mouse monoclonal anti-M13 phage coat protein, HRP | Invitrogen | Cat# MA5-36126 |
| Donkey anti-Rabbit IgG(H+L), Alexa Fluor 488 | Invitrogen | Cat# A-21206; RRID:AB_2535792 |
| Goat anti-Mouse IgG(H+L), Alexa Fluor 568 | Invitrogen | Cat# A-11004; RRID:AB_2534072 |
| Goat anti-Rat IgG (H+L), Alexa Fluor 488 | Invitrogen | Cat# A-11006; RRID:AB_2534074 |
| IRDye 800CW Goat anti-Mouse IgG | LI-COR | P/N 925-32210; RRID: AB_2687825 |
| IRDye 800CW Goat anti-Rat IgG | LI-COR | P/N 926-32219; RRID: AB_2721932 |
| IRDye 800CW Goat anti-Rabbit IgG | LI-COR | P/N 926-32211; RRID: AB_2651127 |
| **Software** | | |
| DALI protein structure comparison server | (*82*) | http://ekhidna2.biocenter.helsinki.fi/dali/ |
| XDS | (*117*) | https://xds.mr.mpg.de/ |
| DIALS | (*118*) | http://dials.github.io/ |

| REAGENT or RESOURCE | SOURCE | IDENTIFIER |
|---|---|---|
| RESOLVE | (*124*) | http://resolve.lanl.gov/ |
| COOT | (*126*) | http://www2.mrc-lmb.cam.ac.uk/personal/pemsley/coot/ |
| PHENIX | (*127*) | http://phenix-online.org/ |
| MolProbity | (*128*) | http://molprobity.biochem.duke.edu/ |
| Fiji | (*137*) | http://imagej.net/software/fiji/ |
| ALSCRIPT | (*139*) | http://www.cryst.bbk.ac.uk/CCSG/info/software/prot-sequ/alscript/alscript.html |
| ASTRA 6 | Wyatt Technology | http://www.wyatt.com/products/software/astra.html |
| ClustalX | (*138*) | http://www.clustal.org/clustal2/ |
| Chimera fitmap tool | (*129*) | http://www.cgl.ucsf.edu/chimera/ |
| CRISPR Design Tools | (*133*) | http://crispr.mit.edu:8079<br>http://figshare.com/articles/ CRISPR_Design_Tool/1117899 |
| Prism | GraphPad Software | http://www.graphpad.com/scientific-software/prism/ |
| PYMOL | Schrödinger | http://pymol.org/2/ |
| IGOR Pro | WaveMetrics | http://www.wavemetrics.com/ |
| Adobe illustrator | Adobe | http://www.adobe.com/products/illustrator.html |
| Adobe photoshop | Adobe | http://www.adobe.com/products/photoshop.html |
| **Expression vectors** | | |
| pET24a | Addgene | Cat# 69749-3 |
| pET28a-SUMO | (*108*) | N/A |
| pET28a-SUMO-Avi | This study | N/A |
| pET28a-His-PreS-Avi | (*27*) | N/A |
| pET28a-His-PreS-Avi-SUMO | This study | N/A |
| pET28a-PreS | (*107*) | N/A |
| pGEX-6P1 | GE Healthcare | Cat# 28-9546-48 |
| pETDuet1 | Novagen | Cat# 71146-3 |
| pETDuet1-SUMO | This study | N/A |
| pETDuet1-SUMO2 | This study | N/A |
| pETDuet1-SUMO-Avi | This Study | N/A |
| pET-MCN | (*109*) | N/A |
| pET-MCN-SUMO | (*11*) | N/A |
| pET-MCN-PreS | This study | N/A |
| pcDNA3.1 | Invitrogen | Cat# V79020 |
| pX330 | Addgene | Cat# 42230 |
| pRH2.2 | Sidhu Lab (*110*) | N/A |
| pFastBac-HTB | Invitrogen | Cat# 10360014 |
| pFastBac-Dual | Invitrogen | Cat# 10712024 |
| **Columns and resins** | | |
| HiLoad 16/60 Superdex 200 PG | GE Healthcare | N/A |
| HiLoad 16/60 Superdex 75 PG | GE Healthcare | N/A |
| Superdex 200 10/300 GL | GE Healthcare | N/A |
| Superdex 75 10/300 GL | GE Healthcare | N/A |
| Superdex 200 Increase 10/300 GL | GE Healthcare | N/A |
| Superose 6 10/300 GL | GE Healthcare | N/A |
| HiTrap Q HP | GE Healthcare | N/A |
| HiTrap SP HP | GE Healthcare | N/A |
| MonoQ 10/100 GL | GE Healthcare | N/A |
| HiTrap Heparin HP | GE Healthcare | N/A |
| HiPrep 26/20 Desalting | GE Healthcare | N/A |
| HiTrap Desalting 5 mL | GE Healthcare | N/A |
| HiTrap MabSelect SuRe | GE Healthcare | N/A |
| Glutathione Sepharose 4 fast flow | GE Healthcare | Cat# 17-5132-02 |
| Ni-NTA agarose | Qiagen | Cat# 30230 |
| **Cell lines** | | |
| $HCT116^{AID::NUP358}$ | This study | N/A |
| $DLD1^{AID::NUP358}$ | This study | N/A |
| $DLD1^{NUP160::NG\text{-}AID}$ | This study | N/A |
| *E. coil* BL21-CodonPlus (DE3)-RIL | Agilent Technologies | Cat# 230245 |
| *E. coli* Max Efficiency DH5$\alpha$ | Invitrogen | Cat# 18258012 |
| *E. coli* XL1-Blue cells | Agilent Technologies | Cat# 200249 |

| REAGENT or RESOURCE | SOURCE | IDENTIFIER |
|---|---|---|
| *E. coli* BL21-Gold | Agilent Technologies | Cat# 230130 |
| DH10Bac cells | Gibco | Cat# 10361012 |
| *T. ni* Hi5 cells | Gibco | Cat# B5502 |
| *S. frugiperda* Sf9 cells | Gibco | Cat# 11496015 |
| *C. thermophilum* strain | Gift, Ed Hurt (*145*) | DSM-1495 |

#### **Table S2.** Bacterial expression constructs and conditions

| # | Protein | Residues | Expression vector | Restriction sites 5', 3' | N-terminal overhang | C-terminal overhang | Expression conditions |
|---|---|---|---|---|---|---|---|
| 1 | NUP88 | 1-741 | pET28a-SUMO | BamHI, NotI | S | AAALEHHHHHH | 18 °C, 18 hours |
| 2[♦] | NUP88 NTD | 1-493 | pET28a-SUMO | BamHI, NotI | S | none | 18 °C, 18 hours |
| 3 | NUP62<br>NUP214 | 317-522<br>(1-450)-(699-955) | pETDuet1-SUMO2 | NcoI, NotI<br>NdeI, XhoI | S<br>S | none<br>none | 18 °C, 18 hours |
| 4 | NUP62 | 317-522 | pET28a-PreS | NdeI, XhoI | MGSSHHHHHHSS<br>GLEVLFQGPH | none | 18 °C, 18 hours |
| 5 | NUP214 CCS1-3<br>NUP88 CCS1-3 | 699-888<br>559-741 | pETDuet1-SUMO | BamHI, NotI<br>NdeI/XhoI | S<br>none | none<br>none | 18 °C, 18 hours |
| 6 | AVI-NUP214 CCS1-3<br><br><br>NUP88 CCS1-3 | 699-888<br><br><br>559-741 | pETDuet1-SUMO-Avi | BamHI, NotI<br><br><br>Nde, XhoI | SSGLNDIFEAQKIE<br>WHEGSAGGSGGS<br>M | none<br><br><br>none | 18 °C, 18 hours |
| 7 | NUP98 GLEBS-APD | (157-213)-GSGSGSHM-(715-863) | pET28a-Pres | NdeI, XhoI | GPHM | none | 18 °C, 18 hours |
| 8[♦] | NUP98 APD | 715-863 | pET28a-PreS | NdeI, XhoI | GPHM | none | 18 °C, 18 hours |
| 9 | DDX19 | 1-479 | pET28a-PreS | NdeI, NotI | none | none | 18 °C, 18 hours |
| 10 | DDX19 E243Q | 1-479 E243Q | pET28a-PreS | NdeI, NotI | GPHM | none | 18 °C, 18 hours |
| 11 | GLE1 CTD<br>NUP42 GBM | 382-698<br>379-423 | pETDuet1-PreS | NdeI, XhoI<br>BamHI, NotI | M<br>GPSGS | none<br>none | 18 °C, 18 hours |
| 12 | NUP42 ZnF | 2-26 | pET28a-SUMO | BamHI, XhoI | S | none | 37 °C, 2 hours |
| 13 | NUP214 NTD | 1-450 | pET28a-PreS | NdeI, NotI | GPHM | none | 18 °C, 18 hours |
| 14 | SUMO-NUP214 TAIL | 908-955 | pET28a-SUMO | BamHI, NotI | none | none | 18 °C, 18 hours |
| 15 | SUMO-NUP214 (908-952) | 908-952 | pET28a-SUMO | BamHI, NotI | none | none | 18 °C, 18 hours |
| 16 | SUMO-NUP214 (938-955) | 938-955 | pET28a-SUMO | BamHI, NotI | none | none | 18 °C, 18 hours |
| 17 | SUMO-NUP214 (940-955) | 940-955 | pET28a-SUMO | BamHI, NotI | none | none | 18 °C, 18 hours |
| 18 | NUP155 CTD | 870-1391 | pET28a-PreS | NdeI, NotI | GPHM | none | 18 °C, 18 hours |
| 19 | SUMO-NUP93 R1 | 2-93 | pET28a-SUMO | BamHI, NotI | S | none | 18 °C, 18 hours |
| 20 | SUMO-NUP93 R1 LIL | 2-93<br>L33A, I36A, L43A | pET28a-SUMO | BamHI, NotI | S | none | 18 °C, 18 hours |
| 21[♦] | NUP358 NTD | 1-752 I599M | pET28a-SUMO | BamHI, NotI | S | none | 18 °C, 18 hours |
| 22 | NUP358 OE | 802-832 | pET28a-SUMO | BamHI, NotI | S | none | 18 °C ,18 hours |
| 23[♦] | NUP358 OE | 805-832 | pET28a-SUMO | BamHI, NotI | S | none | 18 °C ,18 hours |
| 24 | NUP358 NTD-OE | 1-832 I599M | pET28a-SUMO | BamHI, NotI | S | AAALEHHHHHH | 18 °C ,18 hours |
| 25 | NUP358 OE LIQIML | 802-832<br>L811A, I814A, Q817A, I821A, M825A, L828A | pET28a-SUMO | BamHI, NotI | S | none | 18 °C, 18 hours |
| 26 | NUP358 NTD-OE LIQIML | 1-832 I599M<br>L811A, I814A, Q817A, I821A, M825A, L828A | pET28a-SUMO | BamHI, NotI | S | AAALEHHHHHH | 18 °C, 18 hours |
| 27 | NUP358 TPR | 1-145 | pGEX-6P1 | BamHI, NotI | GPLGS | none | 18 °C, 18 hours |
| 28 | NUP358 NTD ΔTPR | 145-752 I599M | pET28a-SUMO | BamHI, NotI | S | none | 18 °C, 18 hours |
| 29 | NUP358 1-673 | 1-673 I599M | pET28a-SUMO | BamHI, NotI | S | none | 18 °C, 18 hours |
| 30[♦] | NUP358 145-673 | 145-673 I599M | pET28a-SUMO | BamHI, NotI | S | none | 18 °C, 18 hours |
| 31[#] | AVI-NUP358 145-673 | 145-673 | pET28a-SUMO-Avi | BamHI, NotI | SSGLNDIFEAQKIE<br>WHEGSAGGSGGS | none | 18 °C, 18 hours |
| 32[♦] | NUP358 RBD-I | 1171-1306 | pGEX-6P1 | BamHI, NotI | GPLGS | none | 18 °C, 18 hours |
| 33[♦] | NUP358 RBD-II | 2012-2148 | pGEX-6P1 | BamHI, NotI | GPLGS | none | 18 °C, 18 hours |
| 34[♦] | NUP358 RBD-III | 2309-2444 | pGEX-6P1 | BamHI, NotI | GPLGS | none | 18 °C, 18 hours |
| 35 | NUP358 RBD-IV | 2911-3045 | pGEX-6P1 | BamHI, NotI | GPLGS | none | 18 °C, 18 hours |
| 36[♦] | NUP358 RBD-IV | 2911-3045<br>W2962Q, H2963N, T2964Y, M2965D, K2966N, N2967K, Y2968Q, Y2969V | pGEX-6P1 | BamHI, NotI | GPLGS | none | 18 °C, 18 hours |
| 37 | NUP358 ZnF1 | 1343-1380 | pGEX-6P1 | BamHI, NotI | GPLGS | none | 37 °C, 2 hours |
| 38[♦] | NUP358 ZnF2 | 1416-1443 | pGEX-6P1 | BamHI, NotI | GPLGS | none | 37 °C, 2 hours |
| 39[♦] | NUP358 ZnF3 | 1471-1507 | pGEX-6P1 | BamHI, NotI | GPLGS | none | 37 °C, 2 hours |
| 40[♦] | NUP358 ZnF4 | 1535-1571 | pGEX-6P1 | BamHI, NotI | GPLGS | none | 37 °C, 2 hours |
| 41[♦] | NUP358 ZnF5/6 | 1607-1634 | pGEX-6P1 | BamHI, NotI | GPLGS | none | 37 °C, 2 hours |
| 42[♦] | NUP358 ZnF7 | 1716-1752 | pGEX-6P1 | BamHI, NotI | GPLGS | none | 37 °C, 2 hours |
| 43[♦] | NUP358 ZnF8 | 1774-1809 | pGEX-6P1 | BamHI, NotI | GPLGS | none | 37 °C, 2 hours |
| 44 | NUP358 ZnF1 ΔNTE | 1352-1380 | pGEX-6P1 | BamHI, NotI | GPLGS | none | 37 °C, 2 hours |
| 45 | NUP358 ZnF2 ΔNTE | 1416-1443 | pGEX-6P1 | BamHI, NotI | GPLGS | none | 37 °C, 2 hours |
| 46 | NUP358 ZnF3 ΔNTE | 1480-1507 | pGEX-6P1 | BamHI, NotI | GPLGS | none | 37 °C, 2 hours |
| 47 | NUP358 ZnF4 ΔNTE | 1480-1507 | pGEX-6P1 | BamHI, NotI | GPLGS | none | 37 °C, 2 hours |
| 48 | NUP358 ZnF5/6 ΔNTE | 1607-1634 | pGEX-6P1 | BamHI, NotI | GPLGS | none | 37 °C, 2 hours |
| 49 | NUP358 ZnF7 ΔNTE | 1725-1752 | pGEX-6P1 | BamHI, NotI | GPLGS | none | 37 °C, 2 hours |
| 50 | NUP358 ZnF8 ΔNTE | 1782-1809 | pGEX-6P1 | BamHI, NotI | GPLGS | none | 37 °C, 2 hours |
| 51 | NUP358 ZFD | 1343-1890 | pGEX-6P1 | BamHI, NotI | GPLGS | none | 37 °C, 2 hours |
| 52 | NUP358 CTD | 3062-3224 | pET28a-PreS | NdeI, NotI | GPHM | none | 18 °C, 18 hours |
| 53 | NUP358 NTD (R2E/R3E) | 1-752 I599M | pET28a-SUMO | BamHI, NotI | S | none | 18 °C, 18 hours |
| 54 | NUP358 NTD (R2A/R3A) | 1-752 I599M | pET28a-SUMO | BamHI, NotI | S | none | 18 °C, 18 hours |
| 55 | NUP358 NTD (K5E) | 1-752 I599M | pET28a-SUMO | BamHI, NotI | S | none | 18 °C, 18 hours |

| # | Protein | Residues | Expression vector | Restriction sites 5’, 3’ | N-terminal overhang | C-terminal overhang | Expression conditions |
|---|---|---|---|---|---|---|---|
| 56 | NUP358 NTD (K5A) | 1-752 I599M | pET28a-SUMO | BamHI, NotI | S | none | 18 °C, 18 hours |
| 57 | NUP358 NTD (R10E) | 1-752 I599M | pET28a-SUMO | BamHI, NotI | S | none | 18 °C, 18 hours |
| 58 | NUP358 NTD (R10A) | 1-752 I599M | pET28a-SUMO | BamHI, NotI | S | none | 18 °C, 18 hours |
| 59 | NUP358 NTD (K23E) | 1-752 I599M | pET28a-SUMO | BamHI, NotI | S | none | 18 °C, 18 hours |
| 60 | NUP358 NTD (K23A) | 1-752 I599M | pET28a-SUMO | BamHI, NotI | S | none | 18 °C, 18 hours |
| 61 | NUP358 NTD (K25E) | 1-752 I599M | pET28a-SUMO | BamHI, NotI | S | none | 18 °C, 18 hours |
| 62 | NUP358 NTD (K25A) | 1-752 I599M | pET28a-SUMO | BamHI, NotI | S | none | 18 °C, 18 hours |
| 63 | NUP358 NTD (K28E) | 1-752 I599M | pET28a-SUMO | BamHI, NotI | S | none | 18 °C, 18 hours |
| 64 | NUP358 NTD (K28A) | 1-752 I599M | pET28a-SUMO | BamHI, NotI | S | none | 18 °C, 18 hours |
| 65 | NUP358 NTD (K28E/R58E) | 1-752 I599M | pET28a-SUMO | BamHI, NotI | S | none | 18 °C, 18 hours |
| 66 | NUP358 NTD (K34E) | 1-752 I599M | pET28a-SUMO | BamHI, NotI | S | none | 18 °C, 18 hours |
| 67 | NUP358 NTD (K34A) | 1-752 I599M | pET28a-SUMO | BamHI, NotI | S | none | 18 °C, 18 hours |
| 68 | NUP358 NTD (K40E) | 1-752 I599M | pET28a-SUMO | BamHI, NotI | S | none | 18 °C, 18 hours |
| 69 | NUP358 NTD (K40A) | 1-752 I599M | pET28a-SUMO | BamHI, NotI | S | none | 18 °C, 18 hours |
| 70 | NUP358 NTD (K34A/K40A) | 1-752 I599M | pET28a-SUMO | BamHI, NotI | S | none | 18 °C, 18 hours |
| 71 | NUP358 NTD (K34A/K40A/K61A/R64A) | 1-752 I599M | pET28a-SUMO | BamHI, NotI | S | none | 18 °C, 18 hours |
| 72 | NUP358 NTD (K34A/K40A/R58A/K61A/R64A) | 1-752 I599M | pET28a-SUMO | BamHI, NotI | S | none | 18 °C, 18 hours |
| 73 | NUP358 NTD 2R5K (K34A/K40A/R58A/K61A/R64A/ K500A/K502A) | 1-752 I599M | pET28a-SUMO | BamHI, NotI | S | none | 18 °C, 18 hours |
| 74 | NUP358 NTD (K40E/K61E/R64E) | 1-752 I599M | pET28a-SUMO | BamHI, NotI | S | none | 18 °C, 18 hours |
| 75 | NUP358 NTD (K40A/K61A/R64A) | 1-752 I599M | pET28a-SUMO | BamHI, NotI | S | none | 18 °C, 18 hours |
| 76 | NUP358 NTD (K40A/R58A/K61A/R64A) | 1-752 I599M | pET28a-SUMO | BamHI, NotI | S | none | 18 °C, 18 hours |
| 77 | NUP358 NTD (K40A/R58A/K61A/R64A/H303A) | 1-752 I599M | pET28a-SUMO | BamHI, NotI | S | none | 18 °C, 18 hours |
| 78 | NUP358 NTD (K46/K47E) | 1-752 I599M | pET28a-SUMO | BamHI, NotI | S | none | 18 °C, 18 hours |
| 79 | NUP358 NTD (K47A) | 1-752 I599M | pET28a-SUMO | BamHI, NotI | S | none | 18 °C, 18 hours |
| 80 | NUP358 NTD (R58E) | 1-752 I599M | pET28a-SUMO | BamHI, NotI | S | none | 18 °C, 18 hours |
| 81 | NUP358 NTD (R58A) | 1-752 I599M | pET28a-SUMO | BamHI, NotI | S | none | 18 °C, 18 hours |
| 82 | NUP358 NTD (K61E) | 1-752 I599M | pET28a-SUMO | BamHI, NotI | S | none | 18 °C, 18 hours |
| 83 | NUP358 NTD (K61A) | 1-752 I599M | pET28a-SUMO | BamHI, NotI | S | none | 18 °C, 18 hours |
| 84 | NUP358 NTD (R64E) | 1-752 I599M | pET28a-SUMO | BamHI, NotI | S | none | 18 °C, 18 hours |
| 85 | NUP358 NTD (R64A) | 1-752 I599M | pET28a-SUMO | BamHI, NotI | S | none | 18 °C, 18 hours |
| 86 | NUP358 NTD (K61E/R64E) | 1-752 I599M | pET28a-SUMO | BamHI, NotI | S | none | 18 °C, 18 hours |
| 87 | NUP358 NTD (K61A/R64A) | 1-752 I599M | pET28a-SUMO | BamHI, NotI | S | none | 18 °C, 18 hours |
| 88 | NUP358 NTD (K78E) | 1-752 I599M | pET28a-SUMO | BamHI, NotI | S | none | 18 °C, 18 hours |
| 89 | Nup358 NTD (R84E/R85E) | 1-752 I599M | pET28a-SUMO | BamHI, NotI | S | none | 18 °C, 18 hours |
| 90 | NUP358 NTD (K94E) | 1-752 I599M | pET28a-SUMO | BamHI, NotI | S | none | 18 °C, 18 hours |
| 91 | NUP358 NTD (K94A) | 1-752 I599M | pET28a-SUMO | BamHI, NotI | S | none | 18 °C, 18 hours |
| 92 | NUP358 NTD (K99E) | 1-752 I599M | pET28a-SUMO | BamHI, NotI | S | none | 18 °C, 18 hours |
| 93 | NUP358 NTD (K106E) | 1-752 I599M | pET28a-SUMO | BamHI, NotI | S | none | 18 °C, 18 hours |
| 94 | NUP358 NTD (R113E) | 1-752 I599M | pET28a-SUMO | BamHI, NotI | S | none | 18 °C, 18 hours |
| 95 | NUP358 NTD (R113A) | 1-752 I599M | pET28a-SUMO | BamHI, NotI | S | none | 18 °C, 18 hours |
| 96 | NUP358 NTD (K115E) | 1-752 I599M | pET28a-SUMO | BamHI, NotI | S | none | 18 °C, 18 hours |
| 97 | NUP358 NTD (R120E) | 1-752 I599M | pET28a-SUMO | BamHI, NotI | S | none | 18 °C, 18 hours |
| 98 | NUP358 NTD (K123E) | 1-752 I599M | pET28a-SUMO | BamHI, NotI | S | none | 18 °C, 18 hours |
| 99 | NUP358 NTD (K133E) | 1-752 I599M | pET28a-SUMO | BamHI, NotI | S | none | 18 °C, 18 hours |
| 100 | NUP358 NTD (K133A) | 1-752 I599M | pET28a-SUMO | BamHI, NotI | S | none | 18 °C, 18 hours |
| 101 | NUP358 NTD (K135E) | 1-752 I599M | pET28a-SUMO | BamHI, NotI | S | none | 18 °C, 18 hours |
| 102 | NUP358 NTD (K135A) | 1-752 I599M | pET28a-SUMO | BamHI, NotI | S | none | 18 °C, 18 hours |
| 103 | NUP358 NTD (K149E) | 1-752 I599M | pET28a-SUMO | BamHI, NotI | S | none | 18 °C, 18 hours |
| 104 | NUP358 NTD (R161E) | 1-752 I599M | pET28a-SUMO | BamHI, NotI | S | none | 18 °C, 18 hours |
| 105 | NUP358 NTD (R161A) | 1-752 I599M | pET28a-SUMO | BamHI, NotI | S | none | 18 °C, 18 hours |
| 106 | NUP358 NTD (R170E) | 1-752 I599M | pET28a-SUMO | BamHI, NotI | S | none | 18 °C, 18 hours |
| 107 | NUP358 NTD (R170A) | 1-752 I599M | pET28a-SUMO | BamHI, NotI | S | none | 18 °C, 18 hours |
| 108 | NUP358 NTD (R176E) | 1-752 I599M | pET28a-SUMO | BamHI, NotI | S | none | 18 °C, 18 hours |
| 109 | NUP358 NTD (K179E/R180E) | 1-752 I599M | pET28a-SUMO | BamHI, NotI | S | none | 18 °C, 18 hours |
| 110 | NUP358 NTD (K179A) | 1-752 I599M | pET28a-SUMO | BamHI, NotI | S | none | 18 °C, 18 hours |
| 111 | NUP358 NTD (R180A) | 1-752 I599M | pET28a-SUMO | BamHI, NotI | S | none | 18 °C, 18 hours |
| 112 | NUP358 NTD (K182E) | 1-752 I599M | pET28a-SUMO | BamHI, NotI | S | none | 18 °C, 18 hours |
| 113 | NUP358 NTD (K182A) | 1-752 I599M | pET28a-SUMO | BamHI, NotI | S | none | 18 °C, 18 hours |
| 114 | NUP358 NTD (R193E) | 1-752 I599M | pET28a-SUMO | BamHI, NotI | S | none | 18 °C, 18 hours |
| 115 | NUP358 NTD (R198E) | 1-752 I599M | pET28a-SUMO | BamHI, NotI | S | none | 18 °C, 18 hours |
| 116 | NUP358 NTD (R198A) | 1-752 I599M | pET28a-SUMO | BamHI, NotI | S | none | 18 °C, 18 hours |
| 117 | NUP358 NTD (K212E) | 1-752 I599M | pET28a-SUMO | BamHI, NotI | S | none | 18 °C, 18 hours |
| 118 | NUP358 NTD (K212A) | 1-752 I599M | pET28a-SUMO | BamHI, NotI | S | none | 18 °C, 18 hours |
| 119 | NUP358 NTD (K225E) | 1-752 I599M | pET28a-SUMO | BamHI, NotI | S | none | 18 °C, 18 hours |
| 120 | NUP358 NTD (K225A) | 1-752 I599M | pET28a-SUMO | BamHI, NotI | S | none | 18 °C, 18 hours |
| 121 | NUP358 NTD (R229E) | 1-752 I599M | pET28a-SUMO | BamHI, NotI | S | none | 18 °C, 18 hours |
| 122 | NUP358 NTD (R250E) | 1-752 I599M | pET28a-SUMO | BamHI, NotI | S | none | 18 °C, 18 hours |

| # | Protein | Residues | Expression vector | Restriction sites 5’, 3’ | N-terminal overhang | C-terminal overhang | Expression conditions |
|---|---|---|---|---|---|---|---|
| 123 | NUP358 NTD (R256E) | 1-752 I599M | pET28a-SUMO | BamHI, NotI | S | none | 18 °C, 18 hours |
| 124 | NUP358 NTD (K270E) | 1-752 I599M | pET28a-SUMO | BamHI, NotI | S | none | 18 °C, 18 hours |
| 125 | NUP358 NTD (K286E) | 1-752 I599M | pET28a-SUMO | BamHI, NotI | S | none | 18 °C, 18 hours |
| 126 | NUP358 NTD (K286A) | 1-752 I599M | pET28a-SUMO | BamHI, NotI | S | none | 18 °C, 18 hours |
| 127 | NUP358 NTD (K299E) | 1-752 I599M | pET28a-SUMO | BamHI, NotI | S | none | 18 °C, 18 hours |
| 128 | NUP358 NTD (H303A) | 1-752 I599M | pET28a-SUMO | BamHI, NotI | S | none | 18 °C, 18 hours |
| 129 | NUP358 NTD (R310E) | 1-752 I599M | pET28a-SUMO | BamHI, NotI | S | none | 18 °C, 18 hours |
| 130 | NUP358 NTD (R328E) | 1-752 I599M | pET28a-SUMO | BamHI, NotI | S | none | 18 °C, 18 hours |
| 131 | NUP358 NTD (K330E/K332E) | 1-752 I599M | pET28a-SUMO | BamHI, NotI | S | none | 18 °C, 18 hours |
| 132 | NUP358 NTD (K335E) | 1-752 I599M | pET28a-SUMO | BamHI, NotI | S | none | 18 °C, 18 hours |
| 133 | NUP358 NTD (R363E/K365E) | 1-752 I599M | pET28a-SUMO | BamHI, NotI | S | none | 18 °C, 18 hours |
| 134 | NUP358 NTD (K370E) | 1-752 I599M | pET28a-SUMO | BamHI, NotI | S | none | 18 °C, 18 hours |
| 135 | NUP358 NTD (K379E) | 1-752 I599M | pET28a-SUMO | BamHI, NotI | S | none | 18 °C, 18 hours |
| 136 | NUP358 NTD (K396E) | 1-752 I599M | pET28a-SUMO | BamHI, NotI | S | none | 18 °C, 18 hours |
| 137 | NUP358 NTD (R412E) | 1-752 I599M | pET28a-SUMO | BamHI, NotI | S | none | 18 °C, 18 hours |
| 138 | NUP358 NTD (R421E) | 1-752 I599M | pET28a-SUMO | BamHI, NotI | S | none | 18 °C, 18 hours |
| 139 | NUP358 NTD (R428E) | 1-752 I599M | pET28a-SUMO | BamHI, NotI | S | none | 18 °C, 18 hours |
| 140 | NUP358 NTD (R454E/K458E) | 1-752 I599M | pET28a-SUMO | BamHI, NotI | S | none | 18 °C, 18 hours |
| 141 | NUP358 NTD (K455E) | 1-752 I599M | pET28a-SUMO | BamHI, NotI | S | none | 18 °C, 18 hours |
| 142 | NUP358 NTD (R470E) | 1-752 I599M | pET28a-SUMO | BamHI, NotI | S | none | 18 °C, 18 hours |
| 143 | NUP358 NTD (K500E/K502E) | 1-752 I599M | pET28a-SUMO | BamHI, NotI | S | none | 18 °C, 18 hours |
| 144 | NUP358 NTD (K500A/K502A) | 1-752 I599M | pET28a-SUMO | BamHI, NotI | S | none | 18 °C, 18 hours |
| 145 | NUP358 NTD (K521E) | 1-752 I599M | pET28a-SUMO | BamHI, NotI | S | none | 18 °C, 18 hours |
| 146 | NUP358 NTD (R527E/K529E) | 1-752 I599M | pET28a-SUMO | BamHI, NotI | S | none | 18 °C, 18 hours |
| 147 | NUP358 NTD (R527A/K529A) | 1-752 I599M | pET28a-SUMO | BamHI, NotI | S | none | 18 °C, 18 hours |
| 148 | NUP358 NTD (R541E/K542E) | 1-752 I599M | pET28a-SUMO | BamHI, NotI | S | none | 18 °C, 18 hours |
| 149 | NUP358 NTD (R550E/R552E) | 1-752 I599M | pET28a-SUMO | BamHI, NotI | S | none | 18 °C, 18 hours |
| 150 | NUP358 NTD (R563) | 1-752 I599M | pET28a-SUMO | BamHI, NotI | S | none | 18 °C, 18 hours |
| 151 | NUP358 NTD (K567E) | 1-752 I599M | pET28a-SUMO | BamHI, NotI | S | none | 18 °C, 18 hours |
| 152 | NUP358 NTD (K584E) | 1-752 I599M | pET28a-SUMO | BamHI, NotI | S | none | 18 °C, 18 hours |
| 153 | NUP358 NTD (R596E) | 1-752 I599M | pET28a-SUMO | BamHI, NotI | S | none | 18 °C, 18 hours |
| 154 | NUP358 NTD (R601) | 1-752 I599M | pET28a-SUMO | BamHI, NotI | S | none | 18 °C, 18 hours |
| 155 | NUP358 NTD (K614E) | 1-752 I599M | pET28a-SUMO | BamHI, NotI | S | none | 18 °C, 18 hours |
| 156 | NUP358 NTD (K607E K608E) | 1-752 I599M | pET28a-SUMO | BamHI, NotI | S | none | 18 °C, 18 hours |
| 157 | NUP358 NTD (K617E/K618E/K619E) | 1-752 I599M | pET28a-SUMO | BamHI, NotI | S | none | 18 °C, 18 hours |
| 158 | NUP358 NTD (K631E) | 1-752 I599M | pET28a-SUMO | BamHI, NotI | S | none | 18 °C, 18 hours |
| 159 | NUP358 NTD (K675E) | 1-752 I599M | pET28a-SUMO | BamHI, NotI | S | none | 18 °C, 18 hours |
| 160 | NUP358 NTD (K690E) | 1-752 I599M | pET28a-SUMO | BamHI, NotI | S | none | 18 °C, 18 hours |
| 161 | NUP358 NTD (R689E/K690E) | 1-752 I599M | pET28a-SUMO | BamHI, NotI | S | none | 18 °C, 18 hours |
| 162 | NUP358 NTD (K708E) | 1-752 I599M | pET28a-SUMO | BamHI, NotI | S | none | 18 °C, 18 hours |
| 163 | NUP358 NTD (R712E) | 1-752 I599M | pET28a-SUMO | BamHI, NotI | S | none | 18 °C, 18 hours |
| 164 | NUP358 NTD (R715E) | 1-752 I599M | pET28a-SUMO | BamHI, NotI | S | none | 18 °C, 18 hours |
| 165 | NUP358 NTD (K713E/K720E) | 1-752 I599M | pET28a-SUMO | BamHI, NotI | S | none | 18 °C, 18 hours |
| 166 | NUP358 NTD (K720E) | 1-752 I599M | pET28a-SUMO | BamHI, NotI | S | none | 18 °C, 18 hours |
| 167 | NUP358 NTD (K733E K734E) | 1-752 I599M | pET28a-SUMO | BamHI, NotI | S | none | 18 °C, 18 hours |
| 168 | NUP358 NTD (K743E) | 1-752 I599M | pET28a-SUMO | BamHI, NotI | S | none | 18 °C, 18 hours |
| 169• | NUP358 NTD (D140M) | 1-752 I599M | pET28a-SUMO | BamHI, NotI | S | none | 18 °C, 18 hours |
| 170• | NUP358 NTD (D152M) | 1-752 I599M | pET28a-SUMO | BamHI, NotI | S | none | 18 °C, 18 hours |
| 171• | NUP358 NTD (I171M) | 1-752 I599M | pET28a-SUMO | BamHI, NotI | S | none | 18 °C, 18 hours |
| 172• | NUP358 NTD (I322M) | 1-752 I599M | pET28a-SUMO | BamHI, NotI | S | none | 18 °C, 18 hours |
| 173• | NUP358 NTD (I427M) | 1-752 I599M | pET28a-SUMO | BamHI, NotI | S | none | 18 °C, 18 hours |
| 174• | NUP358 NTD (I453M) | 1-752 I599M | pET28a-SUMO | BamHI, NotI | S | none | 18 °C, 18 hours |
| 175• | NUP358 NTD (I539M) | 1-752 I599M | pET28a-SUMO | BamHI, NotI | S | none | 18 °C, 18 hours |
| 176• | NUP358 NTD (I559M) | 1-752 I599M | pET28a-SUMO | BamHI, NotI | S | none | 18 °C, 18 hours |
| 177• | NUP358 NTD (I615M) | 1-752 I599M | pET28a-SUMO | BamHI, NotI | S | none | 18 °C, 18 hours |
| 178• | NUP358 NTD (I626M) | 1-752 I599M | pET28a-SUMO | BamHI, NotI | S | none | 18 °C, 18 hours |
| 179• | NUP358 NTD (I652M) | 1-752 I599M | pET28a-SUMO | BamHI, NotI | S | none | 18 °C, 18 hours |
| 180• | NUP358 NTD (L401M) | 1-752 I599M | pET28a-SUMO | BamHI, NotI | S | none | 18 °C, 18 hours |
| 181• | NUP358 NTD (D693M) | 1-752 I599M | pET28a-SUMO | BamHI, NotI | S | none | 18 °C, 18 hours |
| 182• | NUP358 NTD (N709M) | 1-752 I599M | pET28a-SUMO | BamHI, NotI | S | none | 18 °C, 18 hours |
| 183• | NUP358 NTD (Q137M) | 1-752 I599M | pET28a-SUMO | BamHI, NotI | S | none | 18 °C, 18 hours |
| 184• | NUP358 NTD (T92M) | 1-752 I599M | pET28a-SUMO | BamHI, NotI | S | none | 18 °C, 18 hours |
| 185• | NUP358 NTD (V668M) | 1-752 I599M | pET28a-SUMO | BamHI, NotI | S | none | 18 °C, 18 hours |
| 186♦ | NUP358 NTD ANE1 T585M | 1-752 I599M | pET28a-SUMO | BamHI, NotI | S | none | 18 °C, 18 hours |
| 187♦ | NUP358 NTD ANE1 T653I | 1-752 I599M | pET28a-SUMO | BamHI, NotI | S | none | 18 °C, 18 hours |
| 188♦ | NUP358 NTD ANE1 I656V | 1-752 I599M | pET28a-SUMO | BamHI, NotI | S | none | 18 °C, 18 hours |
| 189 | NUP358 NTD ANE1 W681C | 1-752 I599M | pET28a-SUMO | BamHI, NotI | S | none | 18 °C, 18 hours |
| 190• | NUP358 145-673 (I652M) | 145-673 I599M | pET28a-SUMO | BamHI, NotI | S | none | 18 °C, 18 hours |
| 191• | NUP358 145-673 (I615M) | 145-673 I599M | pET28a-SUMO | BamHI, NotI | S | none | 18 °C, 18 hours |
| 192• | NUP358 145-673 (L401M) | 145-673 I599M | pET28a-SUMO | BamHI, NotI | S | none | 18 °C, 18 hours |
| 193• | NUP358 145-673 (I169M) | 145-673 I599M | pET28a-SUMO | BamHI, NotI | S | none | 18 °C, 18 hours |

| # | Protein | Residues | Expression vector | Restriction sites 5’, 3’ | N-terminal overhang | C-terminal overhang | Expression conditions |
|---|---|---|---|---|---|---|---|
| 194• | NUP358 145-673 (L385M) | 145-673 I599M | pET28a-SUMO | BamHI, NotI | S | none | 18 °C, 18 hours |
| 195• | NUP358 145-673 (I372M) | 145-673 I599M | pET28a-SUMO | BamHI, NotI | S | none | 18 °C, 18 hours |
| 196 | NUP88 NTD (D10A/E12A) | 1-493 | pET28a-SUMO | BamHI, NotI | S | none | 18 °C, 18 hours |
| 197 | NUP88 NTD (E29A, E38A, E40A) | 1-493 | pET28a-SUMO | BamHI, NotI | S | none | 18 °C, 18 hours |
| 198 | NUP88 NTD (E236A, E272A, E275) | 1-493 | pET28a-SUMO | BamHI, NotI | S | none | 18 °C, 18 hours |
| 199 | NUP88 NTD (E304A/D305/ D310A) | 1-493 | pET28a-SUMO | BamHI, NotI | S | none | 18 °C, 18 hours |
| 200 | NUP88 NTD (E340A/E342A/E343/ E344A/ D345A/D346A) | 1-493 | pET28a-SUMO | BamHI, NotI | S | none | 18 °C, 18 hours |
| 201 | NUP88 NTD (E350A/E354A/E358A) | 1-493 | pET28a-SUMO | BamHI, NotI | S | none | 18 °C, 18 hours |
| 202 | NUP88 NTD (E381A/D382A/D383A) | 1-493 | pET28a-SUMO | BamHI, NotI | S | none | 18 °C, 18 hours |
| 203 | NUP88 NTD (D436A/E440A/E444A) | 1-493 | pET28a-SUMO | BamHI, NotI | S | none | 18 °C, 18 hours |
| 204 | NUP88 NTD (E216A/E218A/E219A/E220A) | 1-493 | pET28a-SUMO | BamHI, NotI | S | none | 18 °C, 18 hours |
| 205 | NUP88 NTD (E74A/E75A/E91A/E92A) | 1-493 | pET28a-SUMO | BamHI, NotI | S | none | 18 °C, 18 hours |
| 206 | NUP88 NTD (E329A/E370A/E372A) | 1-493 | pET28a-SUMO | BamHI, NotI | S | none | 18 °C, 18 hours |
| 207 | NUP88 NTD (D434Y) | 1-493 | pET28a-SUMO | BamHI, NotI | S | none | 18 °C, 18 hours |
| 208 | NUP88 NTD (D434A) | 1-493 | pET28a-SUMO | BamHI, NotI | S | none | 18 °C, 18 hours |
| 209 | NUP88 NTD (E433A) | 1-493 | pET28a-SUMO | BamHI, NotI | S | none | 18 °C, 18 hours |
| 210 | NUP88 NTD (E340A/E342A/E343A/E344A/ D345A/D346A/D431A/E432A/ E433A/D434A) | 1-493 | pET28a-SUMO | BamHI, NotI | S | none | 18 °C, 18 hours |
| 211 | NUP88 NTD (E340A/E342A/E343A) | 1-493 | pET28a-SUMO | BamHI, NotI | S | none | 18 °C, 18 hours |
| 212 | NUP88 NTD (E344A/D345A/D346A) | 1-493 | pET28a-SUMO | BamHI, NotI | S | none | 18 °C, 18 hours |
| 213 | NUP88 NTD (D431A/E432A) | 1-493 | pET28a-SUMO | BamHI, NotI | S | none | 18 °C, 18 hours |
| 214 | NUP88 NTD (E433A/D434A) | 1-493 | pET28a-SUMO | BamHI, NotI | S | none | 18 °C, 18 hours |
| 215 | NUP88 NTD (E343A/E344A) | 1-493 | pET28a-SUMO | BamHI, NotI | S | none | 18 °C, 18 hours |
| 216 | NUP88 NTD (E340A/E342A/E343A/E344A/ D345A/D346A/E433A/D434A) | 1-493 | pET28a-SUMO | BamHI, NotI | S | none | 18 °C, 18 hours |
| 217 | NUP88 NTD LLL | 1-493 L424A, L428A, L438A | pET28a-SUMO | BamHI, NotI | S | none | 18 °C, 18 hours |
| 218 | NUP88 NTD MNY | 1-493 M299A, N306A, Y307A | pET28a-SUMO | BamHI, NotI | S | none | 18 °C, 18 hours |
| 219 | NUP88 NTD EMNY | 1-493 E275A, M299A, N306A, Y307A | pET28a-SUMO | BamHI, NotI | S | none | 18 °C, 18 hours |
| 220 | NUP88 NTD ∆FGL loop | 1-493 ∆228-233 | pET28a-SUMO | BamHI, NotI | S | none | 18 °C, 18 hours |
| 221 | TAP<br>P15 | 117-554<br>1-140 | pET-MCN-SUMO | BamHI, NotI<br>NdeI, XhoI | S<br>none | none<br>none | 18 °C, 18 hours |
| 222♦ | RAN | 1-216 | pET28a-SUMO | BamHI, NotI | S | none | 18 °C, 18 hours |
| 223 | RAN Q69L | 1-216 Q69L | pET28a-SUMO | BamHI, NotI | S | none | 18 °C, 18 hours |
| 224♦ | RAN F35S | 1-216 | pET28a-SUMO | BamHI, NotI | S | none | 18 °C, 18 hours |
| 225♦ | Thrombin Ran F35 | 1-216 | pET28a | NdeI, NotI | MGSSHHHHHHSS GLVPRGSS | none | 18 °C, 18 hours |
| 226♦ | NUP50 RBD | 336-468 | pET28a-SUMO | BamHI, NotI | S | none | 18 °C, 18 hours |
| 227♦ | sAB14-LC<br>sAB14-HC | 1-217<br>1-263 | pRH2.2 | HindIII, SalI | none | none | 37 °C, 2 hours |
| 228♦ | rnNup153 ZnF1 | 649-687 | pGEX-6P1 | BamHI, NotI | GPLGS | none | 37 °C, 2 hours |
| 229♦ | rnNup153 ZnF2 | 713-749 | pGEX-6P1 | BamHI, NotI | GPLGS | none | 37 °C, 2 hours |
| 230♦ | rnNup153 ZnF3 | 781-817 | pGEX-6P1 | BamHI, NotI | GPLGS | none | 37 °C, 2 hours |
| 231♦ | rnNup153 ZnF4 | 839-874 | pGEX-6P1 | BamHI, NotI | GPLGS | none | 37 °C, 2 hours |
| 232 | rnNup153 ZnF1∆NTE | 659-687 | pGEX-6P1 | BamHI, NotI | GPLGS | none | 37 °C, 2 hours |
| 233 | rnNup153 ZnF2∆NTE | 723-749 | pGEX-6P1 | BamHI, NotI | GPLGS | none | 37 °C, 2 hours |
| 234 | rnNup153 ZnF3∆NTE | 791-817 | pGEX-6P1 | BamHI, NotI | GPLGS | none | 37 °C, 2 hours |
| 235 | rnNup153 ZnF4∆NTE | 848-874 | pGEX-6P1 | BamHI, NotI | GPLGS | none | 37 °C, 2 hours |
| 236 | Nup82 | 1-882 | pET28a-SUMO | BamHI, NotI | S | AAALEHHHHHH | 18 °C, 18 hours |
| 237 | Nup82 NTD | 1-595 | pET28a-PreS | NdeI, NotI | GPHM | none | 18 °C, 18 hours |
| 238 | Nup82 CCS1 | 661-784 | pET28a-SUMO | BamHI, XhoI | S | none | 18 °C, 18 hours |
| 239 | Nup82 CCS2-3 | 793-882 | pET28a-SUMO | BamHI, XhoI | S | none | 18 °C, 18 hours |
| 240 | Nup82 CCS1-3 | 661-882 | pET28a-SUMO | BamHI, XhoI | S | none | 18 °C, 18 hours |

| # | Protein | Residues | Expression vector | Restriction sites 5’, 3’ | N-terminal overhang | C-terminal overhang | Expression conditions |
|---|---|---|---|---|---|---|---|
| 241 | Nup159 ΔNTD | 1109-1367 | pET28a-SUMO | BamHI, NotI | S | none | 18 °C, 18 hours |
| 242 | Nup159 NTD | 1-444 | pET28a-SUMO | BamHI, XhoI | S | none | 18 °C, 18 hours |
| 243 | Nup159 CCS1 | 1109-1259 | pET28a-SUMO | BamHI, XhoI | S | none | 18 °C, 18 hours |
| 244 | Nup159 CCS2-3 | 1262-1381 | pET28a-SUMO | BamHI, NotI | S | none | 18 °C, 18 hours |
| 245 | Nup159 CCS1-3 | 1109-1381 | pET28a-SUMO | BamHI, NotI | S | none | 18 °C, 18 hours |
| 246 | Nsp1 CCS1 | 467-557 | pET28a-SUMO | BamHI, XhoI | S | none | 18 °C, 18 hours |
| 247 | Nsp1 CCS2-3 | 569-674 | pET28a-SUMO | BamHI, XhoI | S | none | 18 °C, 18 hours |
| 248 | Nsp1 CCS1-3 | 467-678 | pET28a-SUMO | BamHI, XhoI | S | none | 18 °C, 18 hours |
| 249 | Nsp1<br>Nup159 | 467-678<br>1-456 1109-1481 | pETDuet1 | NcoI, NotI<br>NdeI, XhoI | none<br>M | none<br>none | 18 °C, 18 hours |
| 250 | Nsp1<br>Nup82 CCS1-3 | 467-678<br>661-879 | pETDuet1 | NcoI, NotI<br>NdeI, XhoI | none<br>M | none<br>none | 18 °C, 18 hours |
| 251 | Nsp1<br>Nup82ΔNTD | 467-678<br>661-879 | pETDuet1 | NcoI, NotI<br>NdeI, XhoI | none<br>M | none<br>HHHHHH | 18 °C, 18 hours |
| 252 | Nsp1<br>Nup159ΔNTD | 467-678<br>1109-1367 | pETDuet-SUMO | BamHI, NotI<br>NdeI/XhoI | M<br>S | none<br>none | 18 °C, 18 hours |
| 253 | Nup159 TAIL | 1440-1481 | pET28a-SUMO | BamHI, NotI | S | none | 18 °C, 18 hours |
| 254 | Nup159ΔTAIL | 1-456 1109-1367 | pET28a-SUMO | BamHI, NotI | S | AAALEHHHHHH | 18 °C, 18 hours |
| 255 | Nup145N GLEBS-APD | 171-221 868-994 | pET28a-SUMO | BamHI, NotI | S | none | 23 °C, 18 hours |
| 256 | Nup145N APD | 868-994 | pET28a-PreS | NdeI, XhoI | GPHM | none | 23 °C, 18 hours |
| 257 | Nup145N GLEBS | 171-221 | pET28a-SUMO | BamHI, NotI | S | none | 18 °C, 18 hours |
| 258 | Dbp5 | 1-477 | pET-MCN-SUMO | BamHI, NotI | S | none | 18 °C, 18 hours |
| 259 | Gle1 | 1-478 | pET28a-PreS | NdeI, NotI | GPHM | none | 37 °C, 3 hours |
| 260 | Gle1 CTD | 216-478 | pET28a-PreS | NdeI, NotI | GPHM | none | 37 °C, 3 hours |
| 261 | Gle1 NTD | 1-216 | pET28a-SUMO | BamHI, NotI | S | none | 18 °C, 18 hours |
| 262 | Nup42 GBM | 494-558 | pETDuet-PreS | NdeI, NotI | MGSSHHHHHHSS<br>GLEGSGSGPSGS | None | 30 °C, 3 hours |
| 263 | Nup120 | 1-1262 | pET28a-SUMO | BamHI, NotI | S | none | 37 °C, 2 hours |
| 264 | Elys | 1-299 | pET-MCN-SUMO | BamHI, NotI | S | none | 37 °C, 2 hours |
| 265 | Nup37 | 1-751 | pET-MCN-SUMO | BamHI, NotI | S | none | 37 °C, 2 hours |
| 266 | Nup37 ΔCTE | 1-692 | pET-MCN-SUMO | BamHI, NotI | S | none | 37 °C, 2 hours |
| 267 | Nup120<br>Nup37 NTD<br>ELYS | 1-1262<br>1-610<br>1-299 | pET-MCN-SUMO | NdeI, BamHI<br>NdeI, BamHI<br>BamHI, NotI | GPHMHHHHHH<br>Q<br>S | none<br>none<br>none | 37 °C, 2 hours |
| 268 | SUMO-Nup37 CTE | 693-751 | pET28a-SUMO | BamHI, NotI | S | none | 18 °C, 18 hours |
| 269 | AVI-SUMO-Nup37 CTE | 693-751 | pET28a-His-PreS-Avi-SUMO | BamHI, NotI | GPLMSGLNDIFEA<br>QKIEWHEGSAGGS<br>GHM | none | 18 °C, 18 hours |
| 270 | SUMO-Nup37 (723-751) | 723-751 | pET28a-SUMO | BamHI, NotI | S | none | 18 °C, 18 hours |
| 271 | SUMO-Nup37 (728-751) | 728-751 | pET28a-SUMO | BamHI, NotI | S | none | 18 °C, 18 hours |
| 272 | SUMO-Nup37 (693-746) | 693-746 | pET28a-SUMO | BamHI, NotI | S | none | 18 °C, 18 hours |
| 273 | SUMO-Nup37 (693-741) | 693-741 | pET28a-SUMO | BamHI, NotI | S | none | 18 °C, 18 hours |
| 274 | SUMO-Nup37 (693-736) | 693-736 | pET28a-SUMO | BamHI, NotI | S | none | 18 °C, 18 hours |
| 275 | Nup85 | 230-1169 | pET24a | NdeI, BamHI | MGHHHHHH | none | 37 °C, 2 hours |
| 276 | Nup85 NTE | 1-230 | pET28a-PreS | NdeI, BamHI | M | none | 18 °C, 18 hours |
| 277 | Nup84 | 1-939 | pET24a | NdeI, BamHI | MGHHHHHH | none | 18 °C, 18 hours |
| 278 | Nup133 | 1-1364 | pET-MCN-SUMO | BamHI, NotI | S | none | 18 °C, 18 hours |
| 279 | Nup133 ΔNTE | 105-1364 | pET-MCN-SUMO | BamHI, NotI | S | none | 18 °C, 18 hours |
| 280 | Nup133 NTE | 1-114 | pET28a-SUMO | BamHI, NotI | S | none | 18 °C, 18 hours |
| 281 | Sec13-Nup145C | 1-309 1-800 | pET-MCN-PreS | NdeI, NotI | GPHM | none | 30 °C, 3 hours |
| 282 | Sec13-Nup145C ΔNTE | 1-309 272-800 | pET-MCN-PreS | NdeI, NotI | GPHM | none | 30 °C, 3 hours |
| 283 | Nup145C NTE | 1-272 | pET28a-SUMO | BamHI, NotI | S | AAALEHHHHHH | 18 °C, 18 hours |
| 284 | AVI-SUMO-Nup145C NTE | 2-272 | pET28a-His-PreS-Avi-SUMO | BamHI, NotI | GPLMSGLNDIFEA<br>QKIEWHEGSAGGS<br>GHM | none | 18 °C, 18 hours |
| 285 | Nup49 | 246-470 | pET28a-SUMO | BamHI, NotI | S | none | 18 °C, 18 hours |
| 286 | Nsp1<br>Nup57 | 467-674<br>74-319 | pETDuet1 | NcoI, NotI<br>NdeI, XhoI | none<br>M | none<br>none | 18 °C, 18 hours |
| 287 | Nic96 R1 | 139-180 | pET28a-SUMO | BamHI, NotI | S | none | 18 °C, 18 hours |

Constructs #8 (*60*), #9, #10, #11, #18, #258, and #260 (*63*), #27 (*80*), #52 (*74*), #237, #253, #256, #285, #286, and #287 (*11*), and #263, #264, #267, #275, #277, #278, and #279 (*38*) were previously reported.

♦ Constructs that were used for crystallization

### Constructs that were used for sAB and MB selection

• Constructs that were used for ascertaining sequence register by Se-Met labeling

**Table S3.** Baculovirus/insect cell expression constructs and conditions

| # | Protein | Residues | Expression vector | Restriction sites 5', 3' | N-terminal overhang | C-terminal overhang | Expression conditions |
|---|---|---|---|---|---|---|---|
| 288 | RAE1 | 1-368 | pFastBac-Dual | SmaI, KpnI | none | HHHHHHH | Described in methods |
| 289 | RAE1<br>NUP98 GLEBS | 1-368<br>157-213 | pFastBac-Dual | SmaI, KpnI<br>BamHI, EcoRI | none | HHHHHHH | Described in methods |
| 290 | Gle2 | 1-357 | pFastBac-HTB | BamHI, XhoI | GAMGS | none | Described in methods |

Constructs #288 and #289 were reported previously in (*57*).

#### **Table S4.** Mammalian cell expression constructs

| Name | Protein | Residues (Mutations) | Vector | Restriction sites 5', 3' | N-terminal tag | C-terminal tag |
|---|---|---|---|---|---|---|
| WT | NUP358 | 1-3224 | pcDNA3.1-3×HA | BamHI, NotI | 3×HA | none |
| TPR | NUP358 | 1-145 | pcDNA3.1-3×HA | BamHI, NotI | 3×HA | none |
| NTD-ZFD | NUP358 | 1-1809 | pcDNA3.1-3×HA | BamHI, NotI | 3×HA | none |
| ΔNTD-ZFD | NUP358 | 1810-3224 | pcDNA3.1-3×HA | BamHI, NotI | 3×HA | none |
| NTD-RanBD-I | NUP358 | 1-1306 | pcDNA3.1-3×HA | BamHI, NotI | 3×HA | none |
| ΔNTD-RanBD-I | NUP358 | 1308-3224 | pcDNA3.1-3×HA | BamHI, NotI | 3×HA | none |
| NTD-OE | NUP358 | 1-832 | pcDNA3.1-3×HA | BamHI, NotI | 3×HA | none |
| ΔNTD-OE | NUP358 | 833-3224 | pcDNA3.1-3×HA | BamHI, NotI | 3×HA | none |
| NTD | NUP358 | 1-752 | pcDNA3.1-3×HA | BamHI, NotI | 3×HA | none |
| OE | NUP358 | 753-832 | pcDNA3.1-3×HA | BamHI, NotI | 3×HA | none |
| FL (LIQIML) | NUP358 | 1-3224 (L811A, I814A, Q817A, I828A, M825A, L828A) | pcDNA3.1-3×HA | BamHI, NotI | 3×HA | none |
| NTD-OE (LIQIML) | NUP358 | 1-832 (L811A, I814A, Q817A, I828A, M825A, L828A) | pcDNA3.1-3×HA | BamHI, NotI | 3×HA | none |
| FL (2R5K) | NUP358 | 1-3224 (K34A, K40A, R58A, K61A, R64A, K500A, K502A) | pcDNA3.1-3×HA | BamHI, NotI | 3×HA | none |
| NTD-OE (2R5K) | NUP358 | 1-832 (K34A, K40A, R58A, K61A, R64A, K500A, K502A) | pcDNA3.1-3×HA | BamHI, NotI | 3×HA | none |
| T585M | NUP358 | 1-3224 (T585M) | pcDNA3.1-3×HA | BamHI, NotI | 3×HA | none |
| T653I | NUP358 | 1-3224 (T653I) | pcDNA3.1-3×HA | BamHI, NotI | 3×HA | none |
| I656V | NUP358 | 1-3224 (I656V) | pcDNA3.1-3×HA | BamHI, NotI | 3×HA | none |
| W681C | NUP358 | 1-3224 (W681C) | pcDNA3.1-3×HA | BamHI, NotI | 3×HA | none |
| Insulin | Insulin | 1-110 | pcDNA3.1-3×C-FLAG | Kpn1/Kpn1 | none | 3×FLAG |
| ALPP | ALPP | 1-535 | pcDNA3.1-3×C-FLAG | Kpn1/Kpn1 | none | 3×FLAG |
| IL6 | IL6 | 1-184 | pcDNA3.1-3×C-FLAG | Kpn1/Kpn1 | none | 3×FLAG |
| IL10 | IL10 | 1-145 | pcDNA3.1-3×C-FLAG | Kpn1/Kpn1 | none | 3×FLAG |
| RPL26 | RPL26 | 1-145 | pcDNA3.1-3×C-FLAG | Kpn1/Kpn1 | none | 3×FLAG |
| GFP | GFP | 1-239 | pcDNA3.1-3×C-FLAG | Kpn1/Kpn1 | none | 3×FLAG |
| H1B | H1B | 1-219 | pcDNA3.1-3×C-FLAG | Kpn1/Kpn1 | none | 3×FLAG |
| TNFα | TNFα | 1-233 | pcDNA3.1-3×C-FLAG | Kpn1/Kpn1 | none | 3×FLAG |

3×HA: N-MYPYDVPDYAGGGYPYDVPDYAGGGYPYDVPDYA-C
3×FLAG: N-MDYKDHDGDYKDHDIDYKDDDDK-C

**Table S5.** Protein purification protocols

| Protein(s) | Expression constructs | Purification step | Buffer A | Buffer B |
|---|---|---|---|---|
| NUP62•NUP88•NUP214 | Co-expression<br>(# 1 and 3) | 1. Ni-NTA<br>2. Dialysis/Cleavage<br>3. Mono Q<br>4. Superose 6 10/300 GL | 1. Ni-A1, 5 mM β-ME, 5% glycerol<br>2. IEX-A1, pH 8.0, 5 mM DTT, 5% glycerol / ULP1<br>3. IEX-A1, pH 8.0, 5 mM DTT, 5% glycerol<br>4. SEC-A, 5 mM DTT, 5% Glycerol | 1. Ni-B1, 5 mM β-ME, 5% glycerol<br>2. N/A<br>3. IEX-B1, pH 8.0, 5 mM DTT, 5% glycerol<br>4. N/A |
| NUP88 NTD<br>Wild-type and mutants | Individual<br>(#2, 196-220) | 1. Ni-NTA<br>2. HiPrep 26/20 Desalting/Cleavage<br>3. HiTrap Q HP<br>4. HiLoad Superdex 200 16/60 PG | 1. Ni-A1, 5 mM β-ME<br>2. IEX-A1, pH 8.0, 5 mM β-ME / ULP1<br>3. IEX-A1, pH 8.0, 1 mM DTT<br>4. SEC-A, 1 mM DTT | 1. Ni-B1, 5 mM β-ME<br>2. N/A<br>3. IEX-B1, pH 8.0, 1 mM DTT<br>4. N/A |
| NUP62•NUP88•NUP214 CCS1-3 | Co-expression<br>(#4 and 5) | 1. Ni-NTA<br>2. HiPrep 26/20 Desalting<br>3. HiTrap Q HP<br>4. HiLoad Superdex 200 16/60 PG<br>5. Cleavage<br>6. MonoQ | 1. Ni-A1, 5 mM β-ME<br>2. IEX-A1, pH 8.0, 5 mM DTT<br>3. IEX-A1, pH 8.0, 5 mM DTT<br>4. SEC-A, 5 mM DTT<br>5. N/A<br>6. IEX-A1, pH 8.0, 5 mM DTT | 1. Ni-B1, 5 mM β-ME<br>2. N/A<br>3. IEX-B1, pH 8.0, 5 mM DTT<br>4. N/A<br>5. N/A<br>6. IEX-B1, pH 8.0, 5 mM DTT |
| NUP62•NUP88•AVI-NUP214 CCS1-3 | Co-expression<br>(#4 and 6) | 1. Ni-NTA<br>2. HiPrep 26/20 Desalting<br>3. HiTrap Q HP<br>4. HiLoad Superdex 200 16/60 PG<br>5. Cleavage<br>6. MonoQ | 1. Ni-A1, 5 mM β-ME<br>2. IEX-A1, pH 8.0, 5 mM DTT<br>3. IEX-A1, pH 8.0, 5 mM DTT<br>4. SEC-A, 5 mM DTT<br>5. N/A<br>6. IEX-A1, pH 8.0, 5 mM DTT | 1. Ni-B1, 5 mM β-ME<br>2. N/A<br>3. IEX-B1, pH 8.0, 5 mM DTT<br>4. N/A<br>5. N/A<br>6. IEX-B1, pH 8.0, 5 mM DTT |
| NUP98 GLEBS-APD | Individual<br>(#7) | 1. Ni-NTA<br>2. Dialysis/Cleavage<br>3. Ni-NTA<br>4. HiTrap Q HP<br>5. HiLoad Superdex 75 16/60 PG | 1. Ni-A1, 5 mM β-ME<br>2. IEX-A1, pH 8.0, 5 mM β-ME / Pres<br>3. Ni-A2, 5 mM β-ME<br>4. IEX-A1, pH 8.0, 5 mM DTT<br>5. SEC-A, 5 mM DTT | 1. Ni-B1, 5 mM β-ME<br>2. N/A<br>3. Ni-B1, 5 mM β-ME<br>4. IEX-B1, pH 8.0, 5 mM DTT<br>5. N/A |
| NUP98 APD | Individual<br>(#8) | 1. Ni-NTA<br>2. Dialysis/Cleavage<br>3. Ni-NTA<br>4. HiTrap Q HP<br>5. HiLoad Superdex 75 16/60 PG | 1. Ni-A1, 5 mM β-ME<br>2. IEX-A1, pH 8.0, 5 mM β-ME / Pres<br>3. Ni-A2, 5 mM β-ME<br>4. IEX-A1, pH 8.0, 5 mM DTT<br>5. SEC-A, 5 mM DTT | 1. Ni-B1, 5 mM β-ME<br>2. N/A<br>3. Ni-B1, 5 mM β-ME<br>4. IEX-B1, pH 8.0, 5 mM DTT<br>5. N/A |
| DDX19 | Individual<br>(#9) | 1. Ni-NTA<br>2. Dialysis/Cleavage<br>3. Ni-NTA<br>4. HiTrap Q HP<br>5. HiLoad Superdex 200 16/60 PG | 1. Ni-A1, 5 mM β-ME, 5% glycerol<br>2. IEX-A1, pH 8.0, 5 mM β-ME, 5% glycerol / Pres<br>3. Ni-A2, 5 mM β-ME, 5% glycerol<br>4. IEX-A1, pH 8.0, 5 mM DTT, 5% glycerol<br>5. SEC-A, 5 mM DTT, 5% glycerol | 1. Ni-B1, 5 mM β-ME, 5% glycerol<br>2. N/A<br>3. Ni-B1, 5 mM β-ME, 5% glycerol<br>4. IEX-B1, pH 8.0, 5 mM DTT, 5% Glycerol<br>5. N/A |
| DDX19 E243Q | Individual<br>(#10) | 1. Ni-NTA<br>2. Dialysis/Cleavage<br>3. Ni-NTA<br>4. HiTrap Q HP<br>5. HiLoad Superdex 200 16/60 PG | 1. Ni-A1, 5 mM β-ME, 5% glycerol<br>2. IEX-A1, pH 8.0, 5 mM β-ME, 5% glycerol / Pres<br>3. Ni-A2, 5 mM β-ME, 5% glycerol<br>4. IEX-A1, pH 8.0, 5 mM DTT, 5% glycerol<br>5. SEC-A, 5 mM DTT, 5% glycerol | 1. Ni-B1, 5 mM β-ME, 5% glycerol<br>2. N/A<br>3. Ni-B1, 5 mM β-ME, 5% glycerol<br>4. IEX-B1, pH 8.0, 5 mM DTT, 5% Glycerol<br>5. N/A |
| GLE1 CTD•NUP42 GBM | Co-expression<br>(#11) | 1. Ni-NTA<br>2. Dialysis/Cleavage<br>3. Ni-NTA<br>4. HiTrap Heparin HP<br>5. HiLoad Superdex 200 16/60 PG | 1. Ni-A1, 5 mM β-ME<br>2. IEX-A1, pH 8.0, 5 mM β-ME / Pres<br>3. Ni-A2, 5 mM β-ME<br>4. IEX-A1, pH 8.0, 5 mM DTT<br>5. SEC-A, 5 mM DTT | 1. Ni-B1, 5 mM β-ME<br>2. N/A<br>3. Ni-B1, 5 mM β-ME<br>4. IEX-B1, pH 8.0, 5 mM DTT<br>5. N/A |
| SUMO-NUP42 ZnF | Individual<br>(#12) | 1. Ni-NTA<br>2. HiPrep 26/20 Desalting<br>3. HiTrap Q HP<br>4. HiLoad Superdex 75 16/60 PG | 1. Ni-A1, 5 mM β-ME<br>2. IEX-A1, pH 8.0, 5 mM DTT<br>3. IEX-A1, pH 8.0, 5 mM DTT<br>4. SEC-A, 5 mM DTT | 1. Ni-B1, 5 mM β-ME<br>2. N/A<br>3. IEX-B1, pH 8.0, 5 mM DTT<br>4. N/A |
| NUP214 NTD | Individual<br>(#13) | 1. Ni-NTA<br>2. Dialysis/Cleavage<br>3. Ni-NTA<br>4. HiTrap Q HP<br>5. HiLoad Superdex 200 16/60 PG | 1. Ni-A1, 5 mM β-ME<br>2. IEX-A1, pH 8.0, 5 mM β-ME / Pres<br>3. Ni-A2, 5 mM β-ME<br>4. IEX-A1, pH 8.0, 5 mM DTT<br>5. SEC-A, 5 mM DTT | 1 Ni-B1, 5 mM β-ME<br>2. N/A<br>3. Ni-B1, 5 mM β-ME<br>4. IEX-B1, pH 8.0, 5 mM DTT<br>5. N/A |
| SUMO-NUP214TAIL<br>WT and truncations | Individual<br>(#14-17) | 1. Ni-NTA<br>2. HiPrep 26/20 Desalting<br>3. HiTrap SP HP<br>4. HiLoad Superdex 75 16/60 PG | 1. Ni-A1, 5 mM β-ME<br>2. IEX-A1, pH 8.0, 5 mM DTT<br>3. IEX-A1, pH 8.0, 5 mM DTT<br>4. SEC-A, 5 mM DTT | 1. Ni-B1, 5 mM β-ME<br>2. N/A<br>3. IEX-B1, pH 7.0, 5 mM DTT<br>4. N/A |
| NUP155 CTD | Individual<br>(#18) | 1. Ni-NTA<br>2. HiPrep 26/20 Desalting/Cleavage<br>3. Ni-NTA<br>4. HiTrap Q HP<br>5. HiLoad Superdex 200 16/60 PG | 1. Ni-A1, 5 mM β-ME, 5% glycerol<br>2. IEX-A1, pH 8.0, 5 mM β-ME, 5% glycerol / Pres<br>3. Ni-A2, 5 mM β-ME, 5% glycerol<br>4. IEX-A1, pH 8.0, 5 mM DTT, 5% glycerol<br>5. SEC-B, 5 mM DTT, 5% glycerol | 1. Ni-B1, 5 mM β-ME, 5% glycerol<br>2. N/A<br>3. Ni-B1, 5 mM β-ME, 5% glycerol<br>4. IEX-B1, pH 8.0, 5 mM DTT, 5% Glycerol<br>5.N/A |
| NUP93 R1<br>Wildtype and mutants | Individual<br>(#19-20) | 1. Ni-NTA<br>2. HiPrep 26/20 Desalting/Cleavage<br>3. HiTrap Q HP<br>4. HiLoad Superdex 200 16/60 PG | 1. Ni-A1, 5 mM β-ME<br>2. IEX-A1, pH 8.0, 5 mM DTT/ ULP1<br>3. IEX-A1, pH 8.0, 5 mM DTT<br>4. SEC-C, 5 mM DTT | 1. Ni-B1, 5 mM β-ME<br>2. N/A<br>3. IEX-B1, pH 8.0, 5 mM DTT<br>4. N/A |
| SUMO-NUP93 R1 | Individual<br>(#19) | 1. Ni-NTA<br>2. HiPrep 26/20 Desalting<br>3. HiTrap Q HP<br>4. HiLoad Superdex 200 16/60 PG | 1. Ni-A1, 5 mM β-ME<br>2. IEX-A1, pH 8.0, 5 mM DTT<br>3. IEX-A1, pH 8.0, 5 mM DTT<br>4. SEC-C, 5 mM DTT | 1. Ni-B1, 5 mM β-ME<br>2. N/A<br>3. IEX-B1, pH 8.0, 5 mM DTT<br>4. N/A |
| NUP358NTD<br>Wild-type and mutants | Individual<br>(#21, 53-189) | 1. Ni-NTA<br>2. HiPrep 26/20 Desalting/Cleavage<br>3. HiTrap Q HP<br>4. HiLoad Superdex 200 16/60 PG | 1. Ni-A1, 5 mM β-ME<br>2. IEX-A1, pH 8.0, 5 mM β-ME / ULP1<br>3. IEX-A1, pH 8.0, 1 mM DTT<br>4. SEC-A, 1 mM DTT | 1. Ni-B1, 5 mM β-ME<br>2. N/A<br>3. IEX-B1, pH 8.0, 1 mM DTT<br>4. N/A |
| NUP358 OE and OE LIQIML | Individual<br>(#22, 23, 25) | 1. Ni-NTA<br>2. HiPrep 26/20 Desalting/Cleavage<br>3. Ni-NTA<br>4. HiTrap Q HP<br>5. HiLoad Superdex 200 16/60 PG | 1. Ni-A1, 5 mM β-ME<br>2. IEX-A1, pH 8.0, 5 mM β-ME / Pres<br>3. Ni-A2, 5 mM β-ME<br>4. IEX-A1, pH 8.0, 5 mM DTT<br>5. SEC-A, 5 mM DTT | 1. Ni-B1, 5 mM β-ME<br>2. N/A<br>3. Ni-B1, 5 mM β-ME<br>4. IEX-B1, pH 8.0, 5 mM DTT<br>5. N/A |

| Protein(s) | Expression constructs | Purification step | Buffer A | Buffer B |
|---|---|---|---|---|
| NUP358 NTD-OE | Individual<br>(#24) | 1. Ni-NTA<br>2. HiPrep 26/20 Desalting/Cleavage<br>3. HiLoad Superdex 200 16/60 PG<br>4. HiLoad Superdex 200 16/60 PG | 1. Ni-A1, 5 mM β-ME<br>2. SEC-E, 5mM DTT / ULP1<br>3. SEC-E, 5mM DTT<br>4. SEC-E, 5mM DTT | 1. Ni-B1, 5 mM β-ME<br>2. N/A<br>3. N/A<br>4. N/A |
| NUP358 NTD-OE<br>LIQIML mutant | Individual<br>(#26) | 1. Ni-NTA<br>2. HiPrep 26/20 Desalting/Cleavage<br>3. HiTrap Q HP<br>4. HiLoad Superdex 200 16/60 PG | 1. Ni-A1, 5 mM β-ME<br>2. IEX-A1, pH 8.0, 5 mM β-ME / ULP1<br>3. IEX-A1, pH 8.0, 1 mM DTT<br>4. SEC-A, 1 mM DTT | 1. Ni-B1, 5 mM β-ME<br>2. N/A<br>3. IEX-B1, pH 8.0, 1 mM DTT<br>4. N/A |
| NUP358 TPR | Individual<br>(#27) | 1. GST Affinity<br>2. Dialysis/Cleavage<br>3. GST Affinity<br>4. HiTrap Q HP<br>5. HiLoad Superdex 75 16/60 PG | 1. GST-A, 5 mM DTT<br>2. IEX-A1, pH 8.0, 5 mM DTT / Pres<br>3. IEX-A1, pH 8.0, 5 mM DTT<br>4. IEX-A1, pH 8.0, 5 mM DTT<br>5. SEC-A, 5 mM DTT | 1. GST-B, 5 mM DTT<br>2. N/A<br>3. GST-B, 5 mM DTT<br>4. IEX-B1, pH 8.0, 5 mM DTT<br>5. N/A |
| NUP358 NTDΔTPR | Individual<br>(#28) | 1. Ni-NTA<br>2. HiPrep 26/20 Desalting/Cleavage<br>3. HiTrap Q HP<br>4. HiLoad Superdex 200 16/60 PG | 1. Ni-A1, 5 mM β-ME<br>2. IEX-A1, pH 8.0, 5 mM β-ME / ULP1<br>3. IEX-A1, pH 8.0, 1 mM DTT<br>4. SEC-A, 1 mM DTT | 1. Ni-B1, 5 mM β-ME<br>2. N/A<br>3. IEX-B1, pH 8.0, 1 mM DTT<br>4. N/A |
| NUP358 1-673 | Individual<br>(#29) | 1. Ni-NTA<br>2. HiPrep 26/20 Desalting/Cleavage<br>3. HiTrap Q HP<br>4. HiLoad Superdex 200 16/60 PG | 1. Ni-A1, 5 mM β-ME<br>2. IEX-A1, pH 8.0, 5 mM β-ME / ULP1<br>3. IEX-A1, pH 8.0, 1 mM DTT<br>4. SEC-A, 1 mM DTT | 1. Ni-B1, 5 mM β-ME<br>2. N/A<br>3. IEX-B1, pH 8.0, 1 mM DTT<br>4. N/A |
| NUP358 145-673 | Individual<br>(#30, 190-195) | 1. Ni-NTA<br>2. HiPrep 26/20 Desalting/Cleavage<br>3. HiTrap Q HP<br>4. HiLoad Superdex 200 16/60 PG | 1. Ni-A1, 5 mM β-ME<br>2. IEX-A1, pH 8.0, 5 mM β-ME / ULP1<br>3. IEX-A1, pH 8.0, 1 mM DTT<br>4. SEC-A, 1 mM DTT | 1. Ni-B1, 5 mM β-ME<br>2. N/A<br>3. IEX-B1, pH 8.0, 1 mM DTT<br>4. N/A |
| AVI-NUP358 145-673 | Individual<br>(#31) | 1. Ni-NTA<br>2. HiPrep 26/20 Desalting/Cleavage<br>3. HiTrap Q HP<br>4. HiLoad Superdex 200 16/60 PG | 1. Ni-A1, 5 mM β-ME<br>2. IEX-A1, pH 8.0, 5 mM β-ME / ULP1<br>3. IEX-A1, pH 8.0, 1 mM DTT<br>4. SEC-A, 1 mM DTT | 1. Ni-B1, 5 mM β-ME<br>2. N/A<br>3. IEX-B1, pH 8.0, 1 mM DTT<br>4. N/A |
| NUP358 RBD | Individual<br>(#32-36) | 1. GST Affinity<br>2. Dialysis/Cleavage<br>3. GST Affinity<br>4. HiTrap Q HP<br>5. HiLoad Superdex 75 16/60 PG | 1. GST-A, 5 mM DTT<br>2. IEX-A1, pH 8.0, 5 mM DTT / Pres<br>3. IEX-A1, pH 8.0, 5 mM DTT<br>4. IEX-A1, pH 8.0, 5 mM DTT<br>5. SEC-A, 5 mM DTT | 1. GST-B, 5 mM DTT<br>2. N/A<br>3. GST-B, 5 mM DTT<br>4. IEX-B1, pH 8.0, 5 mM DTT<br>5. N/A |
| NUP358 ZnFs | Individual<br>(#37-50) | 1. GST Affinity<br>2. Dialysis/Cleavage<br>3. GST Affinity<br>4. HiTrap Q HP<br>5. HiLoad Superdex 75 16/60 PG | 1. GST-A, 5 mM DTT<br>2. IEX-A1, pH 8.0, 5 mM DTT / Pres<br>3. IEX-A1, pH 8.0, 5 mM DTT<br>4. IEX-A1, pH 8.0, 5 mM DTT<br>5. SEC-A, 5 mM DTT | 1. GST-B, 5 mM DTT<br>2. N/A<br>3. GST-B, 5 mM DTT<br>4. IEX-B1, pH 8.0, 5 mM DTT<br>5. N/A |
| NUP358 ZFD | Individual<br>(#51) | 1. GST Affinity<br>2. Dialysis/Cleavage<br>3. GST Affinity<br>4. HiTrap Q HP<br>5. HiLoad Superdex 200 16/60 PG | 1. GST-A, 5 mM DTT<br>2. IEX-A1, pH 8.0, 5 mM DTT / Pres<br>3. IEX-A1, pH 8.0, 5 mM DTT<br>4. IEX-A1, pH 8.0, 5 mM DTT<br>5. SEC-A, 5 mM DTT | 1. GST-B, 5 mM DTT<br>2. N/A<br>3. GST-B, 5 mM DTT<br>4. IEX-B1, pH 8.0, 5 mM DTT<br>5. N/A |
| NUP358 CTD | Individual<br>(#52) | 1. Ni-NTA<br>2. HiPrep 26/20 Desalting/Cleavage<br>3. Ni-NTA<br>4. HiLoad Superdex 75 16/60 PG | 1. Ni-A1, 5 mM β-ME<br>2. IEX-A1, pH 8.0, 5 mM β-ME / Pres<br>3. Ni-A2, 5 mM β-ME<br>4. SEC-A, 5 mM DTT | 1. Ni-B1, 5 mM β-ME<br>2. N/A<br>3. Ni-B1, 5 mM β-ME<br>4. N/A |
| P15 TAP | Co-expression<br>(# 221) | 1. Ni-NTA<br>2. HiPrep 26/20 Desalting/Cleavage<br>3. Ni-NTA<br>4. HiTrap Heparin<br>5. HiLoad Superdex 200 16/60 PG | 1. Ni-A1, 5 mM β-ME<br>2. IEX-A1, pH 8.0, 5 mM β-ME / ULP1<br>3. Ni-A2, 5 mM β-ME<br>4. IEX-A1, pH 8.0, 5 mM DTT<br>5. SEC-F, 5 mM DTT | 1. Ni-B1, 5 mM β-ME<br>2. N/A<br>3. Ni-B1, 5 mM β-ME<br>4. IEX-B1, pH 8.0, 5 mM DTT<br>5. N/A |
| Ran<br>Wild-type and mutants | Individual<br>(#222-224) | 1. Ni-NTA<br>2. HiPrep 26/20 Desalting/Cleavage<br>3. Ni-NTA<br>4. HiTrap Q HP<br>5. HiLoad Superdex 75 16/60 PG | 1. Ni-A1, 5 mM β-ME<br>2. IEX-A1, pH 8.0, 5 mM β-ME / ULP1<br>3. Ni-A2, 5 mM β-ME<br>4. IEX-A1, pH 8.0, 5 mM DTT<br>5. SEC-A, 5 mM DTT | 1. Ni-B1, 5 mM β-ME<br>2. N/A<br>3. Ni-B1, 5 mM β-ME<br>4. IEX-B1, pH 8.0, 5 mM DTT<br>5. N/A |
| Thrombin Ran F35S | Individual<br>(#225) | 1. Ni-NTA<br>2. HiPrep 26/20 Desalting<br>3. HiTrap Q HP<br>4. HiLoad Superdex 200 16/60 PG | 1. Ni-A1, 5 mM β-ME<br>2. IEX-A1, pH 8.0, 5 mM β-ME<br>3. IEX-A1, pH 8.0, 1 mM DTT<br>4. SEC-A, 1 mM DTT | 1. Ni-B1, 5 mM β-ME<br>2. N/A<br>3. IEX-B1, pH 8.0, 1 mM DTT<br>4. N/A |
| NUP50 RanBD | Individual<br>(#226) | 1. Ni-NTA<br>2. HiPrep 26/20 Desalting/Cleavage<br>3. Ni-NTA<br>4. HiTrap Q HP<br>5. HiLoad Superdex 75 16/60 PG | 1. Ni-A1, 5 mM β-ME<br>2. IEX-A1, pH 8.0, 5 mM β-ME / ULP1<br>3. Ni-A2, 5 mM β-ME<br>4. IEX-A1, pH 8.0, 5 mM DTT<br>5. SEC-A, 5 mM DTT | 1. Ni-B1, 5 mM β-ME<br>2. N/A<br>3. Ni-B1, 5 mM β-ME<br>4. IEX-B1, pH 8.0, 5 mM DTT<br>5. N/A |
| sAB14 | Individual<br>(#227) | Detailed description in SI methods | | |
| rnNUP153 ZnFs | Individual<br>(#228-235) | 1. GST Affinity<br>2. Dialysis/Cleavage<br>3. GST Affinity<br>4. HiTrap Q HP<br>5. HiLoad Superdex 75 16/60 PG | 1. GST-A, 5 mM DTT<br>2. IEX-A1, pH 8.0, 5 mM DTT / Pres<br>3. IEX-A1, pH 8.0, 5 mM DTT<br>4. IEX-A1, pH 8.0, 5 mM DTT<br>5. SEC-A, 5 mM DTT | 1. GST-B, 5 mM DTT<br>2. N/A<br>3. GST-B, 5 mM DTT<br>4. IEX-B1, pH 8.0, 5 mM DTT<br>5. N/A |
| Nup82•Nsp1•Nup159 | Co-expression<br>(#236<br>and 249) | 1. Ni-NTA<br>2. HiPrep 26/20 Desalting /Cleavage<br>3. MonoQ 10/100 GL<br>4. Superose 6 10/300 GL | 1. Ni-A3, 5 mM β-ME<br>2. IEX-A3, pH 8.0, 5 mM DTT / ULP1<br>3. IEX-A3, pH 8.0, 5 mM DTT<br>4. SEC-C, pH 8.0, 5 mM DTT | 1. Ni-B3, 5 mM β -ME<br>2. N/A<br>3. IEX-B2, pH 8.0, 5 mM DTT<br>4. N/A |
| Nup82•Nsp1•<br>Nup159ΔNTD | Co-expression<br>(#236<br>and 252) | 1. Ni-NTA<br>2. HiPrep 26/20 Desalting /Cleavage<br>3. MonoQ 10/100 GL<br>4. Superose 6 10/300 GL | 1. Ni-A3, 5 mM β -ME<br>2. IEX-A3, pH 8.0, 5 mM DTT / ULP1<br>3. IEX-A3, pH 8.0, 5 mM DTT<br>4. SEC-C, pH 8.0, 5 mM DTT | 1. Ni-B3, 5 mM β-ME<br>2. N/A<br>3. IEX-B2, pH 8.0, 5 mM DTT<br>4. N/A |
| Nup82 NTD•<br>Nup159 TAIL | Co-expression<br>(# 237<br>and 253) | 1. Ni-NTA<br>2. Dialysis/Cleavage<br>3. Ni-NTA<br>4. MonoQ 10/100 GL<br>5. HiLoad Superdex 200 16/60 PG | 1. Ni-A3, 5 mM β-ME<br>2. Ni-A4, 5 mM β-ME / ULP1 / PreS<br>3. Ni-A4, 5 mM β-ME<br>4. IEX-A2, pH 8.0, 5 mM DTT<br>5. SEC-C, pH 8.0, 5 mM DTT | 1. Ni-B3, 5 mM β-ME<br>2. N/A<br>3. Ni-B3, 5 mM β-ME<br>4. IEX-B2, pH 8.0, 5 mM DTT<br>5. N/A |

| Protein(s) | Expression constructs | Purification step | Buffer A | Buffer B |
|---|---|---|---|---|
| Nup82•Nup159•Nsp1<br>CCS1 | Co-lysis<br>(#238, 243,<br>and 246) | 1. Ni-NTA<br>2. Desalting/Cleavage<br>3. Ni-NTA<br>4. HiTrap Q HP<br>5. HiLoad Superdex 200 16/60 PG | 1. Ni-A3, 5 mM β-ME<br>2. Ni-A4, 5 mM β-ME / ULP1<br>3. Ni-A4, 5 mM β-ME<br>4. IEX-A2, pH 8.0, 5 mM DTT<br>5. SEC-C, pH 8.0, 5 mM DTT | 1. Ni-B3, 5 mM β-ME<br>2. N/A<br>3. Ni-B3, 5 mM β-ME<br>4. IEX-B2, pH 8.0, 5 mM DTT<br>5. N/A |
| Nup82•Nup159•Nsp1<br>CCS2-3 | Co-lysis<br>(#239, 244,<br>and 247) | 1. Ni-NTA<br>2. HiPrep 26/20 Desalting/Cleavage<br>3. Ni-NTA<br>4. HiTrap S HP<br>5. HiLoad Superdex 200 16/60 PG | 1. Ni-A3, 5 mM β-ME<br>2. Ni-A4, 5 mM β-ME / ULP1<br>3. Ni-A4, 5 mM β-ME<br>4. IEX-A2, pH 8.0, 5 mM DTT<br>5. SEC-C, pH 8.0, 5 mM DTT | 1. Ni-B3, 5 mM β-ME<br>2. N/A<br>3. Ni-B3, 5 mM β-ME<br>4. IEX-B2, pH 8.0, 5 mM DTT<br>5. N/A |
| Nup82•Nsp1•Nup159<br>CFNC-hub | Co-expression<br>(#245<br>and 250) | 1. Ni-NTA<br>2. HiPrep 26/20 Desalting/Cleavage<br>3. HiTrap Q HP<br>4. HiLoad Superdex 200 16/60 PG | 1. Ni-A3, 5 mM β-ME<br>2. IEX-A3, pH 8.0, 5 mM DTT / ULP1<br>3. IEX-A3, pH 8.0, 5 mM DTT<br>4. SEC-C, pH 8.0, 5 mM DTT | 1. Ni-B3, 5 mM β-ME<br>2. N/A<br>3. IEX-B2, pH 8.0, 5 mM DTT<br>4. N/A |
| Nup159 NTD | Individual<br>(#242) | 1. Ni-NTA<br>2. HiPrep 26/20 Desalting/Cleavage<br>3. Ni-NTA<br>4. HiTrap Q HP<br>5. HiLoad Superdex 200 16/60 PG | 1. Ni-A3, 5 mM β-ME<br>2. Ni-A4, 5 mM β-ME / ULP1<br>3. Ni-A4, 5 mM β-ME<br>4. IEX-A2, pH 8.0, 5 mM DTT<br>5. SEC-C, pH 8.0, 5 mM DTT | 1. Ni-B3, 5 mM β-ME<br>2. N/A<br>3. Ni-B3, 5 mM β-ME<br>4. IEX-B2, pH 8.0, 5 mM DTT<br>5. N/A |
| Nup82ΔNTD•Nsp1•<br>Nup159ΔTAIL | Co-expression<br>(#251<br>and 254) | 1. Ni-NTA<br>2. HiPrep 26/20 Desalting/Cleavage<br>3. MonoQ 10/100 GL<br>4. Superose 6 10/300 GL | 1. Ni-A3, 5 mM β-ME<br>2. IEX-A3, pH 8.0, 5 mM DTT / ULP1<br>3. IEX-A3, pH 8.0, 5 mM DTT<br>4. SEC-C, pH 8.0, 5 mM DTT | 1. Ni-B3, 5 mM β-ME<br>2. N/A<br>3. IEX-B2, pH 8.0, 5 mM DTT<br>4. N/A |
| Nup145N GLEBS-APD | Individual<br>(#255) | 1. Ni-NTA<br>2. HiPrep 26/20 Desalting/Cleavage<br>3. Ni-NTA<br>4. HiTrap SP HP<br>5. HiLoad Superdex 200 16/60 PG | 1. Ni-A3, 5 mM β-ME<br>2. Ni-A4, 5 mM β-ME / ULP1<br>3. Ni-A4, 5 mM β-ME<br>4. IEX-A2, pH 6.5, 5 mM DTT<br>5. SEC-C, pH 8.0, 5 mM DTT | 1. Ni-B3, 5 mM β-ME<br>2. N/A<br>3. Ni-B3, 5 mM β-ME<br>4. IEX-B2, pH 6.5, 5 mM DTT<br>5. N/A |
| Nup145N APD | Individual<br>(#256) | 1. Ni-NTA<br>2. Dialysis/Cleavage<br>3. Ni-NTA<br>4. HiTrap SP HP<br>5. HiLoad Superdex 200 16/60 PG | 1. Ni-A3, 5 mM β-ME<br>2. Ni-A4, 5 mM β-ME / PreS<br>3. Ni-A4, 5 mM β-ME<br>4. IEX-A2, pH 6.5, 5 mM DTT<br>5. SEC-C, pH 8.0, 5 mM DTT | 1. Ni-B3, 5 mM β-ME<br>2. N/A<br>3. Ni-B3, 5 mM β-ME<br>4. IEX-B2, pH 6.5, 5 mM DTT<br>5. N/A |
| Nup145N GLEBS | Individual<br>(#257) | 1. Ni-NTA<br>2. HiPrep 26/20 Desalting/Cleavage<br>3. Ni-NTA<br>4. HiTrap Q HP<br>5. HiLoad Superdex 200 16/60 PG | 1. Ni-A3, 5 mM β-ME<br>2. Ni-A4, 5 mM β-ME / ULP1<br>3. Ni-A4, 5 mM β-ME<br>4. IEX-A2, pH 8.0, 5 mM DTT<br>5. SEC-C, pH 8.0, 5 mM DTT | 1. Ni-B3, 5 mM β -ME<br>2. N/A<br>3. Ni-B3, 5 mM β-ME<br>4. IEX-B2, pH 8.0, 5 mM DTT<br>5. N/A |
| Dbp5 | Individual<br>(#258) | 1. Ni-NTA<br>2. HiPrep 26/20 Desalting/Cleavage<br>3. HiTrap Q HP<br>4. HiLoad Superdex 200 16/60 PG | 1. Ni-A3, 5 mM β-ME<br>2. IEX-A2, pH 8.0, 5 mM DTT / ULP1<br>3. IEX-A2, pH 8.0, 5 mM DTT<br>4. SEC-C, pH 8.0, 5 mM DTT | 1. Ni-B3, 5 mM β-ME<br>2. N/A<br>3. IEX-B2, pH 8.0, 5 mM DTT<br>4. N/A |
| Gle1•Nup42 GBM | Co-expression<br>(#259<br>and 262) | 1. Ni-NTA<br>2. Dialysis/Cleavage<br>3. HiTrap SP HP<br>4. HiLoad Superdex 200 16/60 PG | 1. Ni-A3, 5 mM β-ME<br>2. IEX-A3, pH 8.0, 5 mM DTT / ULP1<br>3. IEX-A3, pH 6.5, 5 mM DTT<br>4. SEC-C, pH 8.0, 5 mM DTT | 1. Ni-B3, 5 mM β-ME<br>2. N/A<br>3. IEX-B2, pH 8.0, 5 mM DTT<br>4. N/A |
| Gle1 CTD•Nup42 GBM | Co-expression<br>(#260<br>and 262) | 1. Ni-NTA<br>2. Dialysis/Cleavage<br>3. HiTrap SP HP<br>4. HiLoad Superdex 200 16/60 PG | 1. Ni-A3, 5 mM β-ME<br>2. IEX-A3, pH 8.0, 5 mM DTT / ULP1<br>3. IEX-A3, pH 6.5, 5 mM DTT<br>4. SEC-C, pH 8.0, 5 mM DTT | 1. Ni-B3, 5 mM β-ME<br>2. N/A<br>3. IEX-B2, pH 8.0, 5 mM DTT<br>4. N/A |
| Gle1 NTD | Individual<br>(# 261) | 1. Ni-NTA<br>2. HiPrep 26/20 Desalting/Cleavage<br>3. HiTrap Q HP<br>4. HiLoad Superdex 200 16/60 PG | 1. Ni-A3, 5 mM β-ME<br>2. IEX-A2, pH 8.0, 5 mM DTT / ULP1<br>3. IEX-A2, pH 8.0, 5 mM DTT<br>4. SEC-C, pH 8.0, 5 mM DTT | 1. Ni-B3, 5 mM β-ME<br>2. N/A<br>3. IEX-B2, pH 8.0, 5 mM DTT<br>4. N/A |
| Nup120<br>Nup37<br>Elys<br>Nup85 | Co-expression<br>(# 263<br>and 265)<br>co-lysis with<br>(# 264<br>and 275) | 1. Ni-NTA<br>2. Dialysis/Cleavage<br>3. HiTrap Q HP<br>4. HiLoad Superdex 200 16/60 PG | 1. Ni-A3, 5 mM β-ME<br>2. IEX-A4, pH 8.0, 5 mM DTT / ULP1<br>3. IEX-A4, pH 8.0, 5 mM DTT<br>4. SEC-D, 5 mM DTT | 1. Ni-B1, 5 mM β-ME, 5 % glycerol<br>2. N/A<br>3. IEX-B2, pH 8.0, 5 mM DTT<br>4. N/A |
| Nup120<br>Nup37ΔCTE<br>Elys<br>Nup85 | Co-expression<br>(# 267<br>and 275) | 1. Ni-NTA<br>2. Dialysis/Cleavage<br>3. HiTrap Q HP<br>4. HiLoad Superdex 200 16/60 PG | 1. Ni-A3, 5 mM β-ME<br>2. IEX-A4, pH 8.0, 5 mM DTT / ULP1<br>3. IEX-A4, pH 8.0, 5 mM DTT<br>4. SEC-D, 5 mM DTT | 1. Ni-B1, 5 mM β-ME, 5 % glycerol<br>2. N/A<br>3. IEX-B2, pH 8.0, 5 mM DTT<br>4. N/A |
| Nup37 | Individual<br>(#265) | 1. Ni-NTA<br>2. HiPrep 26/20 Desalting/Cleavage<br>3. Ni-NTA<br>4. MonoQ 10/100 GL<br>5. HiLoad Superdex 200 16/60 PG | 1. Ni-A3, 5 mM β-ME<br>2. Ni-A4, 5 mM β-ME / ULP1<br>3. Ni-A4, 5 mM β-ME<br>4. IEX-A2, pH 8.0, 5 mM DTT<br>5. SEC-C, pH 8.0, 5 mM DTT | 1. Ni-B3, 5 mM β-ME<br>2. N/A<br>3. Ni-B3, 5 mM β-ME<br>4. IEX-B2, pH 8.0, 5 mM DTT<br>5. N/A |
| Nup37ΔCTE | Individual<br>(#266) | 1. Ni-NTA<br>2. HiPrep 26/20 Desalting/Cleavage<br>3. Ni-NTA<br>4. MonoQ 10/100 GL<br>5. HiLoad Superdex 200 16/60 PG | 1. Ni-A3, 5 mM β-ME<br>2. Ni-A4, 5 mM β-ME / ULP1<br>3. Ni-A4, 5 mM β-ME<br>4. IEX-A2, pH 8.0, 5 mM DTT<br>5. SEC-C, pH 8.0, 5 mM DTT | 1. Ni-B3, 5 mM β-ME<br>2. N/A<br>3. Ni-B3, 5 mM β-ME<br>4. IEX-B2, pH 8.0, 5 mM DTT<br>5. N/A |
| SUMO-Nup37 CTE<br>wildtype and truncation | Individual<br>(#268, 270-<br>274) | 1. Ni-NTA<br>2. Dialysis<br>3. HiTrap Q HP<br>4. HiLoad Superdex 75 16/60 PG | 1. Ni-A3, 5 mM β-ME<br>2. Ni-A4, 5 mM β-ME<br>3. IEX-A2, pH 8.0, 5 mM DTT<br>4. SEC-C, pH 8.0, 5 mM DTT | 1. Ni-B3, 5 mM β-ME<br>2. N/A<br>3. IEX-B2, pH 8.0, 5 mM DTT<br>4. N/A |
| AVI-SUMO-Nup37 CTE | Individual<br>(#269) | 1. Ni-NTA<br>2. HiPrep 26/20 Desalting<br>3. HiTrap Q HP<br>4. HiLoad Superdex 75 16/60 PG | 1. Ni-A3, 5 mM β-ME<br>2. Ni-A4, 5 mM β-ME<br>3. IEX-A2, pH 8.0, 5 mM DTT<br>4. SEC-C, pH 8.0, 5 mM DTT | 1. Ni-B3, 5 mM β-ME<br>2. N/A<br>3. IEX-B2, pH 8.0, 5 mM DTT<br>4. N/A |
| Nup85 NTE | Individual<br>(#276) | 1. Ni-NTA<br>2. Dialysis/Cleavage<br>3. Ni-NTA<br>4. HiTrap Q HP<br>5. HiLoad Superdex 200 16/60 PG | 1. Ni-A3, 5 mM β-ME<br>2. Ni-A4, 5 mM β-ME / PreS<br>3. Ni-A4, 5 mM β-ME<br>4. IEX-A2, pH 8.0, 5 mM DTT<br>5. SEC-C, pH 8.0, 5 mM DTT | 1. Ni-B3, 5 mM β-ME<br>2. N/A<br>3. Ni-B3, 5 mM β-ME<br>4. IEX-B2, pH 8.0, 5 mM DTT<br>5. N/A |

| Protein(s) | Expression constructs | Purification step | Buffer A | Buffer B |
|---|---|---|---|---|
| Nup84•Nup133 | Co-expression (#277-278) | 1. Ni-NTA<br>2. HiPrep 26/20 Desalting/Cleavage<br>3. MonoQ 10/100 GL<br>4. HiLoad Superdex 200 16/60 PG | 1. Ni-A3, 5 mM β-ME<br>2. IEX-A3, pH 8.0, 5 mM DTT / ULP1<br>3. IEX-A3, pH 8.0, 5 mM DTT<br>4. SEC-D, pH 8.0, 5 mM DTT | 1. Ni-B3, 5 mM β-ME<br>2. N/A<br>3. IEX-B2, pH 8.0, 5 mM DTT<br>4. N/A |
| Nup84•Nup133ΔNTE | Co-expression (#277 and 279) | 1. Ni-NTA<br>2. HiPrep 26/20 Desalting/Cleavage<br>3. MonoQ 10/100 GL<br>4. HiLoad Superdex 200 16/60 PG | 1. Ni-A3, 5 mM β-ME<br>2. IEX-A3, pH 8.0, 5 mM DTT / ULP1<br>3. IEX-A3, pH 8.0, 5 mM DTT<br>4. SEC-D, pH 8.0, 5 mM DTT | 1. Ni-B3, 5 mM β-ME<br>2. N/A<br>3. IEX-B2, pH 8.0, 5 mM DTT<br>4. N/A |
| Nup133 NTE | Individual (#280) | 1. Ni-NTA<br>2. Dialysis/Cleavage<br>3. Ni-NTA<br>4. HiTrap SP HP<br>5. HiLoad Superdex 75 16/60 PG | 1. Ni-A3, 5 mM β-ME<br>2. Ni-A4, 5 mM β-ME / PreS<br>3. Ni-A4, 5 mM β-ME<br>4. IEX-A2, pH 6.5, 5 mM DTT<br>5. SEC-C, pH 8.0, 5 mM DTT | 1. Ni-B3, 5 mM β-ME<br>2. N/A<br>3. Ni-B3, 5 mM β-ME<br>4. IEX-B2, pH 6.5, 5 mM DTT<br>5. N/A |
| Sec13-Nup145C | Individual (#281) | 1. Ni-NTA<br>2. Dialysis/Cleavage<br>3. Ni-NTA<br>4. MonoQ 10/100 GL<br>5. HiLoad Superdex 200 16/60 PG | 1. Ni-A3, 5 mM β-ME<br>2. Ni-A4, 5 mM β-ME / PreS<br>3. Ni-A4, 5 mM β-ME<br>4. IEX-A2, pH 8.0, 5 mM DTT<br>5. SEC-C, pH 8.0, 5 mM DTT | 1. Ni-B3, 5 mM β-ME<br>2. N/A<br>3. Ni-B3, 5 mM β-ME<br>4. IEX-B2, pH 8.0, 5 mM DTT<br>5. N/A |
| Sec13-Nup145CΔNTE | Individual (#282) | 1. Ni-NTA<br>2. Dialysis/Cleavage<br>3. Ni-NTA<br>4. MonoQ 10/100 GL<br>5. HiLoad Superdex 200 16/60 PG | 1. Ni-A3, 5 mM β-ME<br>2. Ni-A4, 5 mM β-ME / PreS<br>3. Ni-A4, 5 mM β-ME<br>4. IEX-A2, pH 8.0, 5 mM DTT<br>5. SEC-C, pH 8.0, 5 mM DTT | 1. Ni-B3, 5 mM β-ME<br>2. N/A<br>3. Ni-B3, 5 mM β-ME<br>4. IEX-B2, pH 8.0, 5 mM DTT<br>5. N/A |
| Nup145C NTE | Individual (#283) | 1. Ni-NTA<br>2. HiPrep 26/20 Desalting/Cleavage<br>3. HiTrap Q HP<br>4. Ni-NTA<br>5. HiLoad Superdex 200 16/60 PG | 1. Ni-A3, 5 mM β-ME<br>2. Ni-A4, 5 mM β-ME / ULP1<br>3. IEX-A2, pH 8.0, 5 mM β-ME<br>4. IEX-A2, pH 8.0, 5 mM β-ME<br>5. SEC-C, pH 8.0, 5 mM DTT | 1. Ni-B3, 5 mM β-ME<br>2. N/A<br>3. IEX-B2, pH 8.0, 5 mM DTT<br>4. Ni-B3, 5 mM β-ME<br>5. N/A |
| AVI-SUMO-Nup145C NTE | Individual (#284) | 1. Ni-NTA<br>2. HiPrep 26/20 Desalting<br>3. HiTrap Q HP<br>5. HiLoad Superdex 200 16/60 PG | 1. Ni-A3, 5 mM β-ME<br>2. IEX-A2, pH8.0, 5 mM β-ME<br>3. IEX-A2, pH 8.0, 5 mM β-ME<br>5. SEC-C, pH 8.0, 5 mM DTT | 1. Ni-B3, 5 mM β-ME<br>2. N/A<br>3. IEX-B2, pH 8.0, 5 mM DTT<br>5. N/A |
| Nup49•Nup57•Nsp1 | Co-expression (# 285 and 286) | 1. Ni-NTA<br>2. HiPrep 26/20 Desalting/Cleavage<br>3. Ni-NTA<br>4. MonoQ 10/100 GL<br>5. HiLoad Superdex 200 16/60 PG | 1. Ni-A3, 5 mM β-ME<br>2. Ni-A3, 5 mM β-ME / ULP1<br>3. Ni-A3, 5 mM β-ME<br>4. IEX-A2, pH 8.0, 5 mM DTT<br>5. SEC-C, pH 8.0, 5 mM DTT | 1. Ni-B3, 5 mM β-ME<br>2. N/A<br>3. Ni-B3, 5 mM β-ME<br>4. IEX-B2, pH 8.0, 5 mM DTT<br>5. N/A |
| Nic96 R1 | Individual (#287) | 1. Ni-NTA<br>2. HiPrep 26/20 Desalting<br>3. HiTrap Q HP<br>4. HiLoad Superdex 75 16/60 PG | 1. Ni-A3, 5 mM β-ME<br>2. IEX-A3, pH 8.0, 5 mM DTT<br>3. IEX-A3, pH 8.0, 5 mM DTT<br>4. SEC-C, 5 mM DTT | 1. Ni-B3, 5 mM β-ME<br>2. N/A<br>3. IEX-B2, pH 8.0, 5 mM DTT<br>4. N/A |
| RAE1 | Individual (#288) | 1. Ni-NTA<br>2. Dialysis/Cleavage<br>3. HiTrap SP HP<br>4. HiLoad Superdex 75 16/60 PG | 1. Ni-A1, 5 mM β-ME<br>2. IEX-A3, pH 8.0, 5 mM DTT/TEV<br>3. IEX-A3, pH 6.5, 5 mM DTT<br>4. SEC-D, 5 mM DTT | 1. Ni-B1, 5 mM β-ME<br>2. N/A<br>3. IEX-B1, pH 6.50, 5 mM DTT<br>4. N/A |
| RAE1•NUP98GLEBS | Co-expression (#289) | 1. Ni-NTA<br>2. Dialysis/Cleavage<br>3. HiTrap SP HP<br>4. HiLoad Superdex 75 16/60 PG | 1. Ni-A1, 5 mM β-ME<br>2. IEX-A3, pH 8.0, 5 mM DTT/TEV<br>3. IEX-A3, pH 6.5, 5 mM DTT<br>4. SEC-D, 5 mM DTT | 1. Ni-B1, 5 mM β-ME<br>2. N/A<br>3. IEX-B1, pH 6.5, 5 mM DTT<br>4. N/A |
| Gle2 | Individual (#290) | 1. Ni-NTA<br>2. Dialysis/Cleavage<br>3. HiTrap SP HP<br>4. HiLoad Superdex 75 16/60 PG | 1. Ni-A1, 5 mM β-ME<br>2. IEX-A3, pH 8.0, 5 mM DTT/TEV<br>3. IEX-A3, pH 6.5, 5 mM DTT<br>4. SEC-D, 5 mM DTT | 1. Ni-B1, 5 mM β-ME<br>2. N/A<br>3. IEX-B1, pH 6.50, 5 mM DTT<br>4. N/A |

Ni-A1: 20 mM TRIS (pH 8.0), 500 mM NaCl, 20 mM imidazole
Ni-A2: 20 mM TRIS (pH 8.0), 100 mM NaCl, 20 mM imidazole
Ni-A3: 20 mM TRIS (pH 8.0), 500 mM NaCl, 20 mM imidazole, 5% (v/v) glycerol
Ni-A4: 20 mM TRIS (pH 8.0), 100 mM NaCl, 20 mM imidazole, 5% (v/v) glycerol
Ni-B1: 20 mM TRIS (pH 8.0), 500 mM NaCl, 500 mM imidazole
Ni-B2: 20 mM TRIS (pH 8.0), 100 mM NaCl, 500 mM imidazole
Ni-B3: 20 mM TRIS (pH 8.0), 500 mM NaCl, 500 mM imidazole, 5% (v/v) glycerol
GST-A: 20 mM TRIS (pH 8.0), 300 mM NaCl
GST-B: 20 mM TRIS (pH 8.0), 300 mM NaCl, 20 mM glutathione
IEX-A1: 20 mM TRIS (pH 8.0), 100 mM NaCl
IEX-A2: 20 mM TRIS (pH 8.0), 100 mM NaCl, 5% (v/v) glycerol
IEX-A3: 20 mM TRIS (pH 8.0), 150 mM NaCl, 5% (v/v) glycerol
IEX-A4: 20 mM TRIS (pH 8.0), 200 mM NaCl, 5% (v/v) glycerol
IEX-B1: 20 mM TRIS, 2.0 M NaCl
IEX-B2: 20 mM TRIS, 2.0 M NaCl, 5% (v/v) glycerol
SEC-A: 20 mM TRIS (pH 8.0), 100 mM NaCl
SEC-B: 20 mM TRIS (pH 8.0), 200 mM NaCl
SEC-C: 20 mM TRIS (pH 8.0), 100 mM NaCl, 5% (v/v) glycerol
SEC-D: 20 mM TRIS (pH 8.0), 200 mM NaCl, 5% (v/v) glycerol
SEC-E: 20 mM TRIS (pH 8.0), 350 mM NaCl
SEC-F: 20 mM TRIS (pH 8.0), 150 mM NaCl

**Table S6.** SEC-MALS analysis

| Figure(s) | Nucleoporin or nucleoporin complex | Experimental mass (kDa) | Theoretical mass (kDa) | Stoichiometry | Buffer |
|---|---|---|---|---|---|
| Fig. 1D<br>Fig. S4A | Nup82•Nup159•Nsp1<br>Gle2•Nup145N GLEBS-APD<br>Dbp5<br>Preincubation | 202<br>60<br>47<br>283 | 211<br>62<br>53<br>326 | equimolar<br>equimolar<br>n/a<br>equimolar | SEC-A |
| Fig. 1E<br>Fig. S20A | Nup82•Nup159•Nsp1•Dbp5•Nup145N•Gle2 (CFNC)<br>Gle1•Nup42 GBM +$IP_6$<br>Preincubation | 258<br>62<br>275 | 326<br>68<br>394 | equimolar<br>equimolar<br>equimolar | SEC-A<br>200 $\mu$M $IP_6$ |
| Fig. 1F<br>Fig. S23A | Nup82•Nup159•Nsp1•Dbp5•Nup145N•Gle2 (CFNC)<br>Nup120•Nup37•Elys•Nup85•Sec13-Nup145C•Nup84•Nup133$\Delta$NTE(CNC)<br>Gle1•Nup42GBM +$IP_6$<br>Preincubation | 255<br>791<br>63<br>1163 | 326<br>730<br>68<br>1124 | equimolar<br>equimolar<br>equimolar<br>equimolar | SEC-A<br>200 $\mu$M $IP_6$ |
| Fig. S4B | Nup82 $\Delta$NTD•Nup159 $\Delta$TAIL•Nsp1<br>Dbp5<br>Preincubation | 119<br>51<br>153 | 125<br>52<br>177 | equimolar<br>n/a<br>equimolar | SEC-A |
| Fig. S4C | Nup82•Nup159 $\Delta$NTD•Nsp1<br>Dbp5<br>Preincubation | 141<br>51<br>137 | 150<br>52<br>202 | equimolar<br>n/a<br>no interaction | SEC-A |
| Fig. S4D | Nup82•Nup159 $\Delta$NTD•Nsp1<br>Gle2•Nup145N GLEBS-APD<br>Preincubation | 141<br>60<br>182 | 150<br>62<br>212 | equimolar<br>equimolar<br>equimolar | SEC-A |
| Fig. S4E | Nup82 $\Delta$NTD•Nup159 $\Delta$TAIL•Nsp1<br>Gle2•Nup145N GLEBS-APD<br>Preincubation | 119<br>60<br>121 | 125<br>62<br>187 | equimolar<br>equimolar<br>no interaction | SEC-A |
| Fig. S5A | Nup82•Nup159•Nsp1 (CFNC-hub)<br>Nup159 NTD•Dbp5<br>Preincubation | 81<br>80<br>75 | 77<br>100<br>177 | equimolar<br>equimolar<br>no interaction | SEC-A |
| Fig. S5B | Nup82•Nup159•Nsp1 (CFNC-hub)<br>Gle2•Nup145N GLEBS<br>Preincubation | 81<br>38<br>76 | 77<br>47<br>124 | equimolar<br>equimolar<br>no interaction | SEC-A |
| Fig. S5C | Nup82•Nup159•Nsp1 (CFNC-hub)<br>Nup82 NTD•Nup145N APD•Nup159 TAIL<br>Preincubation | 81<br>57<br>75 | 77<br>85<br>162 | equimolar<br>equimolar<br>no interaction | SEC-A |
| Fig. S5D | Nup82•Nup159•Nsp1 (CFNC-hub)<br>Nup159 NTD•Dbp5<br>Nup82 NTD•Nup145N APD•Nup159 TAIL<br>Gle2•Nup145N GLEBS<br>Preincubation | 80<br>82<br>61<br>43<br>79/80/60/40 | 77<br>100<br>85<br>47<br>309 | equimolar<br>equimolar<br>equimolar<br>equimolar<br>no interaction | SEC-A |
| Fig. S8A | Nup82•Nup159•Nsp1•Dbp5•Nup145N•Gle2 (CFNC)<br>Nup120•Nup37•Elys•Nup85•Sec13-Nup145C•Nup84• Nup133$\Delta$NTE<br>Preincubation | 275<br>734<br>1016 | 326<br>730<br>1056 | equimolar<br>equimolar<br>equimolar | SEC-A |
| Fig. S8B | Nup159NTD•Dbp5<br>Nup120•Nup37•Elys•Nup85•Sec13-Nup145C•Nup84•Nup133$\Delta$NTE<br>Preincubation | 88<br>765<br>762 | 100<br>730<br>830 | equimolar<br>equimolar<br>no interaction | SEC-A |
| Fig. S8C | Gle2•Nup145N GLEBS<br>Nup120•Nup37•Elys•Nup85•Sec13-Nup145C•Nup84•Nup133$\Delta$NTE<br>Preincubation | 42<br>765<br>779 | 47<br>730<br>777 | equimolar<br>equimolar<br>no interaction | SEC-A |
| Fig. S8D | Nup82 NTD•Nup145N APD•Nup159 TAIL<br>Nup120•Nup37•Elys•Nup85•Sec13-Nup145C•Nup84•Nup133$\Delta$NTE<br>Preincubation | 63<br>765<br>780 | 85<br>730<br>815 | equimolar<br>equimolar<br>no interaction | SEC-A |
| Fig. S8E | Nup82•Nup159•Nsp1 (CFNC-hub)<br>Nup120•Nup37•Elys•Nup85•Sec13-Nup145C•Nup84•Nup133$\Delta$NTE<br>Preincubation | 76<br>765<br>819 | 77<br>730<br>806 | equimolar<br>equimolar<br>equimolar | SEC-A |
| Fig. S9A | Nup82•Nup159•Nsp1 (CFNC-hub)<br>Nup133 NTE<br>Preincubation | 81<br>26<br>87 | 77<br>26<br>98 | equimolar<br>n/a<br>no interaction | SEC-A |
| Fig. S9B | Nup82•Nup159•Nsp1 (CFNC-hub)<br>Nup85 NTE<br>Preincubation | 83<br>23<br>83 | 77<br>24<br>101 | equimolar<br>n/a<br>no interaction | SEC-A |
| Fig. S9C | Nup82•Nup159•Nsp1 (CFNC-hub)<br>Nup145C NTE<br>Preincubation | 82<br>28<br>87 | 77<br>30<br>107 | equimolar<br>n/a<br>weak interaction | SEC-A |
| Fig. S9D | Nup82•Nup159•Nsp1 (CFNC-hub)<br>SUMO-Nup37 CTE<br>Preincubation | 81<br>19<br>87 | 77<br>21<br>98 | equimolar<br>n/a<br>equimolar | SEC-A |
| Fig. S10A | Nup82•Nup159•Nsp1 (CFNC-hub)<br>Nup37<br>Preincubation | 81<br>74<br>143 | 77<br>80<br>157 | equimolar<br>n/a<br>equimolar | SEC-A |
| Fig. S10B | Nup82•Nup159•Nsp1 (CFNC-hub)<br>Nup37$\Delta$CTE<br>Preincubation | 81<br>66<br>81 | 77<br>64<br>141 | equimolar<br>n/a<br>no interaction | SEC-A |

| Figure(s) | Nucleoporin or nucleoporin complex | Experimental mass (kDa) | Theoretical mass (kDa) | Stoichiometry | Buffer |
|---|---|---|---|---|---|
| Fig. S10C | Nup82•Nup159•Nsp1 (CFNC-hub)<br>Nup120•Nup37 NTD•Elys•Nup85•Sec13-Nup145C•Nup84•Nup133ΔNTE<br>Preincubation | 76<br>756<br>737 | 77<br>712<br>789 | equimolar<br>equimolar<br>no interaction | SEC-A |
| Fig. S11A | Nup82•Nup159•Nsp1•Dbp5•Nup145N•Gle2 (CFNC)<br>Nup120•Nup37 NTD•Elys•Nup85•Sec13-Nup145C•Nup84• Nup133ΔNTE<br>Preincubation | 275<br>792<br>905 | 326<br>712<br>1037 | equimolar<br>equimolar<br>weak interaction | SEC-A |
| Fig. S11B | Nup82•Nup159•Nsp1•Dbp5•Nup145N•Gle2 (CFNC)<br>Nup120•Nup37•Elys•Nup85•Sec13-Nup145CΔNTE•Nup84•<br>TE<br>Preincubation | 275<br>711<br>942 | 326<br>699<br>1024 | equimolar<br>equimolar<br>equimolar | SEC-A |
| Fig. S11C | Nup82•Nup159•Nsp1•Dbp5•Nup145N•Gle2 (CFNC)<br>Nup120•Nup37<br>Nup85•Sec13-Nup145CΔNTE•Nup84•Nup133ΔNTE<br>Preincubation | 275<br>724<br>829 | 326<br>683<br>1008 | equimolar<br>equimolar<br>weak interaction | SEC-A |
| Fig. S12A | Nup82•Nup159•Nsp1 (CFNC-hub)<br>Nup120•Nup37•Elys•Nup85<br>Preincubation | 76<br>424<br>553 | 77<br>360<br>437 | equimolar<br>equimolar<br>equimolar | SEC-A |
| Fig. S12B | Nup82•Nup159•Nsp1 (CFNC-hub)<br>Sec13-Nup145C<br>Preincubation | 76<br>118<br>120 | 77<br>124<br>201 | equimolar<br>equimolar<br>weak interaction | SEC-A |
| Fig. S12C | Nup82•Nup159•Nsp1 (CFNC-hub)<br>Nup84•Nup133ΔNTE<br>Preincubation | 76<br>226<br>183 | 77<br>246<br>323 | equimolar<br>equimolar<br>no interaction | SEC-A |
| Fig. S13A | Nup82•Nup159•Nsp1<br>Nup120•Nup37<br>Nup85•Sec13-Nup145CΔNTE•Nup84•Nup133ΔNTE<br>Preincubation | 201<br>724<br>719 | 211<br>683<br>894 | equimolar<br>equimolar<br>no interaction | SEC-A |
| Fig. S13B | Nup82•Nup159•Nsp1<br>Nup120•Nup37<br>Nup85•Sec13-Nup145CΔNTE•Nup84•Nup133ΔNTE<br>Dbp5<br>Preincubation | 201<br>724<br>53<br>733 | 211<br>683<br>53<br>947 | equimolar<br>equimolar<br>n/a<br>no interaction | SEC-A |
| Fig. S13C | Nup82•Nup159•Nsp1<br>Nup120•Nup37<br>Nup85•Sec13-Nup145CΔNTE•Nup84•Nup133ΔNTE<br>Nup145N APD<br>Preincubation | 201<br>724<br>15<br>721 | 211<br>683<br>15<br>909 | equimolar<br>equimolar<br>n/a<br>no interaction | SEC-A |
| Fig. S13D | Nup82•Nup159•Nsp1<br>Nup120•Nup37<br>Nup85•Sec13-Nup145CΔNTE•Nup84•Nup133ΔNTE<br>Gle2•Nup145N GLEBS-APD<br>Preincubation | 201<br>724<br>59<br>846 | 211<br>683<br>62<br>956 | equimolar<br>equimolar<br>n/a<br>weak interaction | SEC-A |
| Fig. S16A | Nup82•Nup159•Nsp1 (CFNC-hub) 100 mM NaCl<br>SUMO-Nup37 CTE 100 mM NaCl<br>Preincubation 100 mM NaCl<br>Preincubation 200 mM NaCl<br>Preincubation 300 mM NaCl<br>Preincubation 500 mM NaCl<br>Preincubation 1000 mM NaCl | 78<br>17<br>87<br>84<br>83<br>82<br>80 | 77<br>21<br>98<br>98<br>98<br>98<br>98 | equimolar<br>n/a<br>equimolar<br>equimolar<br>equimolar<br>equimolar<br>equimolar | SEC-A<br><br>100 mM NaCl<br>200 mM NaCl<br>300 mM NaCl<br>500 mM NaCl<br>1000 mM NaCl |
| Fig. S16B | Nup82•Nup159•Nsp1 (CFNC-hub) 100 mM NaCl<br>Nup145C-NTE 100 mM NaCl<br>Preincubation 100 mM NaCl<br>Preincubation 200 mM NaCl<br>Preincubation 300 mM NaCl<br>Preincubation 500 mM NaCl<br>Preincubation 1000 mM NaCl | 78<br>17<br>84<br>80<br>80<br>77<br>79 | 77<br>28<br>105<br>105<br>105<br>105<br>105 | equimolar<br>n/a<br>weak interaction<br>weak interaction<br>weak interaction<br>weak interaction<br>weak interaction | SEC-A<br><br>100 mM NaCl<br>200 mM NaCl<br>300 mM NaCl<br>500 mM NaCl<br>1000 mM NaCl |
| Fig. S16C | Nup82•Nup159•Nsp1 (CFNC-hub)<br>SUMO-Nup37 CTE<br>Nup145C NTE<br>Preincubation | 79<br>19<br>27<br>87 | 77<br>21<br>30<br>107/98 | equimolar<br>n/a<br>n/a<br>equimolar | SEC-A |
| Fig. S17A | Nup82•Nup159•Nsp1•Dbp5•Nup145N•Gle2 (CFNC)<br>SUMO-Nup37 CTE<br>Preincubation | 264<br>20<br>283 | 326<br>21<br>347 | equimolar<br>equimolar<br>equimolar | SEC-A |
| Fig. S17B | Nup82•Nup159•Nsp1•Dbp5•Nup145N•Gle2 (CFNC)<br>Nup145 NTE<br>Preincubation | 264<br>29<br>289 | 326<br>30<br>394 | equimolar<br>equimolar<br>weak interaction | SEC-A |
| Fig. S19B | Nup82•Nup159•Nsp1•Dbp5•Nup145N•Gle2 (CFNC)<br>Gle1•Nup42 GBM -$IP_6$<br>Preincubation | 275<br>71<br>277 | 326<br>68<br>394 | equimolar<br>equimolar<br>weak interaction | SEC-A<br>200 μM $IP_6$ |
| Fig. S19C | Nup82•Nup159•Nsp1•Dbp5•Nup145N•Gle2 (CFNC)<br>Gle1 NTD<br>Preincubation | 258<br>33<br>271 | 326<br>23<br>349 | equimolar<br>n/a<br>weak interaction | SEC-A |

| Figure(s) | Nucleoporin or nucleoporin complex | Experimental mass (kDa) | Theoretical mass (kDa) | Stoichiometry | Buffer |
|---|---|---|---|---|---|
| Fig. S19D | Nup82•Nup159•Nsp1•Dbp5•Nup145N•Gle2 (CFNC)<br>Gle1 CTD•Nup42 GBM -$IP_6$<br>Preincubation | 264<br>43<br>275 | 326<br>44<br>370 | equimolar<br>equimolar<br>no interaction | SEC-A<br>200 $\mu$M $IP_6$ |
| Fig. S19E | Nup82•Nup159•Nsp1•Dbp5•Nup145N•Gle2 (CFNC)<br>Gle1 CTD•Nup42 GBM +$IP_6$<br>Preincubation | 258<br>43<br>258 | 326<br>44<br>370 | equimolar<br>equimolar<br>no interaction | SEC-A<br>200 $\mu$M $IP_6$ |
| Fig. S20A | Dbp5<br>Gle1•Nup42 GBM +$IP_6$<br>Preincubation | 45<br>63<br>62 | 53<br>68<br>121 | n/a<br>equimolar<br>weak interaction | SEC-C<br>200 $\mu$M $IP_6$ |
| Fig. S20B | Nup159 NTD•Dbp5<br>Gle1•Nup42 GBM +$IP_6$<br>Preincubation | 83<br>63<br>62 | 100<br>68<br>168 | equimolar<br>equimolar<br>weak interaction | SEC-A<br>200 $\mu$M $IP_6$ |
| Fig. S20C | Dbp5<br>Gle1 CTD•Nup42 GBM +$IP_6$<br>Preincubation | 45<br>40<br>50 | 53<br>44<br>97 | n/a<br>equimolar<br>weak interaction | SEC-A<br>200 $\mu$M $IP_6$ |
| Fig. S20D | Nup159 NTD•Dbp5<br>Gle1•Nup42 GBM +$IP_6$<br>Preincubation | 83<br>40<br>81 | 100<br>44<br>144 | equimolar<br>equimolar<br>weak interaction | SEC-A<br>200 $\mu$M $IP_6$ |
| Fig. S21A | Nup159NTD•Dbp5<br>Gle1•Nup42 GBM<br>Preincubation | 82<br>65<br>80 | 100<br>68<br>168 | equimolar<br>equimolar<br>no interaction | SEC-A |
| Fig. S21B | Gle2•Nup145N GLEBS<br>Gle1•Nup42 GBM<br>Preincubation | 43<br>65<br>63 | 47<br>68<br>115 | equimolar<br>equimolar<br>no interaction | SEC-A |
| Fig. S21C | Nup82 NTD•Nup145N APD•Nup159 TAIL<br>Gle1•Nup42 GBM<br>Preincubation | 61<br>65<br>63 | 85<br>68<br>115 | equimolar<br>equimolar<br>no interaction | SEC-A |
| Fig. S21D | Nup82•Nup159•Nsp1 (CFNC-hub)<br>Gle1•Nup42 GBM<br>Preincubation | 79<br>65<br>78 | 77<br>68<br>145 | equimolar<br>equimolar<br>no interaction | SEC-A |
| Fig. S21E | Nup82•Nup159•Nsp1 (CFNC-hub)<br>Nup159NTD•Dbp5<br>Gle1•Nup42 GBM<br>Gle2•Nup145N GLEBS<br>Nup82 NTD•Nup145N APD•Nup159 TAIL<br>Preincubation | 79<br>83<br>65<br>40<br>60<br>80/82/72/42/60 | 77<br>100<br>68<br>68<br>47<br>145 | equimolar<br>equimolar<br>equimolar<br>equimolar<br>equimolar<br>no interaction | SEC-A |
| Fig. S23B | Nup82•Nup159•Nsp1•Dbp5•Nup145N•Gle2 (CFNC)<br>Nup120•Nup37•Elys•Nup85•Sec13-Nup145C•Nup84•Nup133ΔNTE<br>Gle1•Nup42GBM -$IP_6$<br>Preincubation | 275<br>734<br>71<br>2002-1273 | 326<br>730<br>68<br>1124 | equimolar<br>equimolar<br>equimolar<br>equimolar<br>imer | SEC-A<br>200 $\mu$M $IP_6$ |
| Fig. S23C | Nup82•Nup159•Nsp1•Dbp5•Nup145N•Gle2 (CFNC)<br>Nup120•Nup37•Elys•Nup85•Sec13-Nup145C•Nup84•Nup133ΔNTE<br>Gle1CTD•Nup42GBM -$IP_6$<br>Preincubation | 275<br>775<br>53<br>1625 | 326<br>730<br>44<br>1100 | equimolar<br>equimolar<br>equimolar<br>equimolar | SEC-A<br>200 $\mu$M $IP_6$ |
| Fig. S23D | Nup82•Nup159•Nsp1•Dbp5•Nup145N•Gle2 (CFNC)<br>Nup120•Nup37•Elys•Nup85•Sec13-Nup145C•Nup84•Nup133ΔNTE<br>Gle1NTD<br>Preincubation | 275<br>775<br>21<br>1625 | 326<br>730<br>23<br>1100 | equimolar<br>equimolar<br>n/a<br>weak interaction | SEC-A |
| Fig. S24A | Gle1•Nup42 GBM -$IP_6$<br>Nup120•Nup37•Elys•Nup85•Sec13-Nup145C•Nup84•Nup133ΔNTE<br>Preincubation | 67<br>765<br>822 | 68<br>730<br>798 | equimolar<br>equimolar<br>equimolar | SEC-A<br>200 $\mu$M $IP_6$ |
| Fig. S24B | Gle1•Nup42 GBM +$IP_6$<br>Nup120•Nup37•Elys•Nup85•Sec13-Nup145C•Nup84•Nup133ΔNTE<br>Preincubation | 67<br>765<br>736 | 68<br>730<br>798 | equimolar<br>equimolar<br>no interaction | SEC-A<br>200 $\mu$M $IP_6$ |
| Fig. S25A | Gle1 NTD -$IP_6$<br>Nup120•Nup37•Elys•Nup85•Sec13-Nup145C•Nup84•Nup133ΔNTE<br>Preincubation | 22<br>743<br>716 | 23<br>730<br>753 | equimolar<br>equimolar<br>no interaction | SEC-A<br>200 $\mu$M $IP_6$ |
| Fig. S25B | Gle1 CTD•Nup42 GBM -$IP_6$<br>Nup120•Nup37•Elys•Nup85•Sec13-Nup145C•Nup84•Nup133ΔNTE<br>Preincubation | 57<br>743<br>672 | 44<br>730<br>774 | equimolar<br>equimolar<br>equimolar | SEC-A<br>200 $\mu$M $IP_6$ |
| Fig. S25C | Gle1 CTD•Nup42 GBM +$IP_6$<br>Nup120•Nup37•Elys•Nup85•Sec13-Nup145C•Nup84•Nup133ΔNTE<br>Preincubation | 40<br>743<br>683 | 44<br>730<br>774 | equimolar<br>equimolar<br>no interaction | SEC-A<br>200 $\mu$M $IP_6$ |
| Fig. 2C<br>Fig. S29A | NUP62•NUP88•NUP214 (CFNC-hub)<br>DDX19 (ADP)<br>RAE1•NUP98<br>Preincubation | 216<br>51<br>57<br>337 | 188<br>54<br>67<br>310 | equimolar<br>n/a<br>equimolar<br>equimolar | SEC-A |
| Fig. S30A | NUP358 NTD<br>NUP88 NTD<br>Preincubation | 89<br>49<br>118 | 86<br>54<br>140 | n/a<br>n/a<br>weak interaction | SEC-B |

| Figure(s) | Nucleoporin or nucleoporin complex | Experimental mass (kDa) | Theoretical mass (kDa) | Stoichiometry | Buffer |
|---|---|---|---|---|---|
| Fig. S30B | NUP358 NTD<br>NUP88 NTD•NUP98 APD<br>Preincubation | 89<br>65<br>95 | 86<br>72<br>158 | n/a<br>equimolar<br>weak interaction | SEC-B |
| Fig. S30C | NUP358 NTD<br>NUP88 NTD•NUP98 APD•NUP214 TAIL<br>Preincubation | 106<br>71<br>94 | 86<br>77<br>163 | n/a<br>equimolar<br>weak interaction | SEC-B |
| Fig. S30D | NUP358 NTD<br>NUP88 NTD•NUP214 TAIL<br>Preincubation | 106<br>56<br>123 | 86<br>60<br>146 | n/a<br>equimolar<br>weak interaction | SEC-B |
| Fig. S30E | NUP358 NTD<br>NUP98 APD<br>Preincubation | 89<br>19<br>93 | 86<br>17<br>103 | n/a<br>n/a<br>no interaction | SEC-B |
| Fig. S30F | NUP358 NTD<br>SUMO-NUP214 TAIL<br>Preincubation | 93<br>23<br>92 | 86<br>17<br>103 | n/a<br>n/a<br>no interaction | SEC-B |
| Fig. S30G | NUP358 NTD<br>NUP214 NTD<br>Preincubation | 93<br>48<br>76 | 86<br>50<br>136 | n/a<br>n/a<br>no interaction | SEC-B |
| Fig. S30H | NUP358 NTD<br>NUP62•NUP88•NUP214 (CFNC-hub)<br>Preincubation | 93<br>69<br>88 | 86<br>69<br>155 | n/a<br>equimolar<br>no interaction | SEC-B |
| Fig. S30I | NUP358 NTD<br>GLE1 CTD•NUP42 GBM<br>Preincubation | 96<br>39<br>95 | 86<br>42<br>128 | n/a<br>equimolar<br>no interaction | SEC-B |
| Fig. S30J | NUP358 NTD<br>DDX19 (ADP)<br>Preincubation | 99<br>51<br>72 | 86<br>54<br>140 | n/a<br>n/a<br>no interaction | SEC-B<br>2 mM $MgCl_2$ |
| Fig. S30K | NUP358 NTD<br>DDX19 E243Q (AMPPNP)<br>Preincubation | 99<br>51<br>101 | 86<br>54<br>140 | n/a<br>n/a<br>no interaction | SEC-B<br>2 mM $MgCl_2$ |
| Fig. S30L | NUP358 NTD<br>SUMO-NUP42 ZnF<br>Preincubation | 99<br>17<br>100 | 86<br>17<br>103 | n/a<br>n/a<br>no interaction | SEC-B<br>2 mM $MgCl_2$<br>1 $\mu$M $ZnCl_2$ |
| Fig. S30M | NUP358 NTD<br>RAE1•NUP98 GLEBS<br>Preincubation | 92<br>46<br>95 | 86<br>48<br>134 | n/a<br>equimolar<br>no interaction | SEC-B |
| Fig. S30N | NUP358 NTD<br>NUP358 OE<br>Preincubation | 99<br>9<br>92 | 86<br>3.6<br>89.5 | n/a<br>n/a<br>no interaction | SEC-B |
| Fig. S31A | NUP88 NTD<br>NUP358 OE<br>Preincubation | 52<br>10<br>52 | 55<br>3.6<br>58.6 | n/a<br>n/a<br>no interaction | SEC-B |
| Fig. S31B | NUP358 OE<br>NUP98 APD<br>Preincubation | 11<br>18<br>17 | 3.6<br>17<br>20.6 | n/a<br>n/a<br>no interaction | SEC-B |
| Fig. S31C | NUP358 OE<br>SUMO-NUP214 TAIL<br>Preincubation | 10<br>23<br>21 | 3.6<br>17<br>20.6 | n/a<br>n/a<br>no interaction | SEC-B |
| Fig. S31D | NUP358 OE<br>NUP214 NTD<br>Preincubation | 10<br>48<br>48 | 3.6<br>50<br>53.6 | n/a<br>n/a<br>no interaction | SEC-B |
| Fig. S31E | NUP358 OE<br>NUP62•NUP88•NUP214 (CFNC-hub)<br>Preincubation | 10<br>69<br>69 | 3.6<br>69<br>72.6 | n/a<br>n/a<br>no interaction | SEC-B |
| Fig. S31F | NUP358 OE<br>DDX19 (ADP)<br>Preincubation | 10<br>51<br>51 | 3.6<br>54<br>57.6 | n/a<br>n/a<br>no interaction | SEC-B<br>2 mM $MgCl_2$ |
| Fig. S31G | NUP358 OE<br>DDX19 E243Q (AMPPNP)<br>Preincubation | 10<br>51<br>53 | 3.6<br>54<br>57.6 | n/a<br>n/a<br>no interaction | SEC-B<br>2 mM $MgCl_2$ |
| Fig. S31H | NUP358 OE<br>GLE1 CTD•NUP42 GBM<br>Preincubation | 10<br>39<br>39 | 3.6<br>42<br>45.6 | n/a<br>equimolar<br>no interaction | SEC-B |
| Fig. S31I | NUP358 OE<br>SUMO-NUP42 ZnF<br>Preincubation | 10<br>17<br>17 | 3.6<br>17<br>20.6 | n/a<br>n/a<br>no interaction | SEC-B |
| Fig. S31J | NUP358 OE<br>RAE1•NUP98 GLEBS<br>Preincubation | 10<br>46<br>46 | 3.6<br>48<br>51.6 | n/a<br>equimolar<br>no interaction | SEC-B |
| Fig. S32A | NUP88 NTD<br>NUP98 APD<br>Preincubation | 49<br>19<br>71 | 54<br>17<br>72 | n/a<br>n/a<br>equimolar | SEC-B |

| Figure(s) | Nucleoporin or nucleoporin complex | Experimental mass (kDa) | Theoretical mass (kDa) | Stoichiometry | Buffer |
|---|---|---|---|---|---|
| Fig. S32B | NUP88 NTD<br>NUP214 TAIL<br>Preincubation | 52<br>23<br>71 | 55<br>17<br>72 | n/a<br>n/a<br>equimolar | SEC-B |
| Fig. S32C | NUP88 NTD<br>NUP214 NTD<br>Preincubation | 51<br>48<br>50 | 55<br>50<br>105 | n/a<br>n/a<br>no interaction | SEC-B |
| Fig. S32D | NUP88 NTD<br>NUP62•NUP88•NUP214 (CFNC-hub)<br>Preincubation | 51<br>69<br>68 | 55<br>69<br>124 | n/a<br>equimolar<br>no interaction | SEC-B |
| Fig. S32E | NUP88 NTD<br>DDX19 (ADP)<br>Preincubation | 51<br>51<br>53 | 55<br>54<br>109 | n/a<br>n/a<br>no interaction | SEC-B<br>2 mM $MgCl_2$ |
| Fig. S32F | NUP88 NTD<br>DDX19 E243Q (AMPPNP)<br>Preincubation | 51<br>51<br>56 | 55<br>54<br>109 | n/a<br>n/a<br>no interaction | SEC-B<br>2 mM $MgCl_2$ |
| Fig. S32G | NUP88 NTD<br>GLE1 CTD•NUP42 GBM<br>Preincubation | 51<br>39<br>50 | 55<br>42<br>97 | n/a<br>equimolar<br>no interaction | SEC-B |
| Fig. S32H | NUP88 NTD<br>SUMO-NUP42 ZnF<br>Preincubation | 51<br>17<br>51 | 55<br>17<br>72 | n/a<br>n/a<br>no interaction | SEC-B<br>2 mM $MgCl_2$<br>1 $\mu$M $ZnCl_2$ |
| Fig. S32I | NUP88 NTD<br>RAE1•NUP98 GLEBS<br>Preincubation | 52<br>46<br>52 | 55<br>48<br>115 | n/a<br>equimolar<br>no interaction | SEC-B |
| Fig. S33A | NUP98 APD<br>SUMO-NUP214 TAIL<br>Preincubation | 17<br>21<br>21 | 17<br>17<br>34 | n/a<br>n/a<br>no interaction | SEC-B |
| Fig. S33B | NUP98 APD<br>NUP214 NTD<br>Preincubation | 17<br>48<br>47 | 17<br>50<br>67 | n/a<br>n/a<br>no interaction | SEC-B |
| Fig. S33C | NUP98 APD<br>NUP62•NUP88•NUP214 (CFNC-hub)<br>Preincubation | 17<br>69<br>67 | 17<br>69<br>86 | n/a<br>equimolar<br>no interaction | SEC-B |
| Fig. S33D | NUP98 APD<br>DDX19 (ADP)<br>Preincubation | 17<br>51<br>51 | 17<br>54<br>71 | n/a<br>n/a<br>no interaction | SEC-B<br>2 mM $MgCl_2$ |
| Fig. S33E | NUP98 APD<br>DDX19 E243Q (AMPPNP)<br>Preincubation | 17<br>52<br>52 | 17<br>54<br>71 | n/a<br>n/a<br>no interaction | SEC-B<br>2 mM $MgCl_2$ |
| Fig. S33F | NUP98 APD<br>GLE1 CTD•NUP42 GBM<br>Preincubation | 17<br>40<br>52 | 17<br>42<br>59 | n/a<br>equimolar<br>no interaction | SEC-B |
| Fig. S33G | NUP98 APD<br>SUMO-NUP42 ZF<br>Preincubation | 19<br>17<br>17 | 17<br>17<br>34 | n/a<br>n/a<br>no interaction | SEC-B<br>2 mM $MgCl_2$<br>1 $\mu$M $ZnCl_2$ |
| Fig. S33H | NUP98 APD<br>NUP155 CTD<br>Preincubation | 17<br>57<br>57 | 17<br>60<br>77 | n/a<br>n/a<br>no interaction | SEC-B |
| Fig. S33I | NUP98 APD<br>RAE1•NUP98 GLEBS<br>Preincubation | 17<br>46<br>44 | 17<br>48<br>65 | n/a<br>equimolar<br>no interaction | SEC-B |
| Fig. S34A | SUMO-NUP214 TAIL<br>NUP214 NTD<br>Preincubation | 23<br>48<br>48 | 17<br>50<br>65 | n/a<br>n/a<br>no interaction | SEC-B |
| Fig. S34B | SUMO-NUP214 TAIL<br>NUP62•NUP88•NUP214 (CFNC-hub)<br>Preincubation | 21<br>69<br>70 | 17<br>69<br>86 | n/a<br>equimolar<br>no interaction | SEC-B |
| Fig. S34C | SUMO-NUP214 TAIL<br>DDX19 (ADP)<br>Preincubation | 21<br>51<br>51 | 17<br>54<br>71 | n/a<br>n/a<br>no interaction | SEC-B<br>2 mM $MgCl_2$ |
| Fig. S34D | SUMO-NUP214 TAIL<br>DDX19 E243Q (AMPPNP)<br>Preincubation | 17<br>51<br>50 | 17<br>54<br>71 | n/a<br>n/a<br>no interaction | SEC-B<br>2 mM $MgCl_2$ |
| Fig. S34E | SUMO-NUP214 TAIL<br>GLE1 CTD•NUP42 GBM<br>Preincubation | 21<br>40<br>24 | 17<br>42<br>59 | n/a<br>equimolar<br>no interaction | SEC-B |
| Fig. S34F | SUMO-NUP214 TAIL<br>SUMO-NUP42 ZnF<br>Preincubation | 21<br>17<br>21 | 17<br>17<br>34 | n/a<br>n/a<br>no interaction | SEC-B<br>2 mM $MgCl_2$<br>1 $\mu$M $ZnCl_2$ |
| Fig. S34G | SUMO-NUP214 TAIL<br>RAE1•NUP98 GLEBS<br>Preincubation | 20<br>46<br>24 | 17<br>48<br>65 | n/a<br>equimolar<br>no interaction | SEC-B |

| Figure(s) | Nucleoporin or nucleoporin complex | Experimental mass (kDa) | Theoretical mass (kDa) | Stoichiometry | Buffer |
|---|---|---|---|---|---|
| Fig. S35A | NUP214 NTD<br>NUP62•NUP88•NUP214 (CFNC-hub)<br>Preincubation | 48<br>69<br>69 | 50<br>69<br>119 | n/a<br>equimolar<br>no interaction | SEC-B |
| Fig. S35B | NUP214 NTD<br>DDX19 (ADP)<br>Preincubation | 48<br>51<br>99 | 50<br>54<br>119 | n/a<br>n/a<br>equimolar | SEC-B<br>2 mM $MgCl_2$ |
| Fig. S35C | NUP214 NTD<br>DDX19 E243Q (AMPPNP)<br>Preincubation | 48<br>51<br>98 | 50<br>54<br>119 | n/a<br>n/a<br>equimolar | SEC-B<br>2 mM $MgCl_2$ |
| Fig. S35D | NUP214 NTD<br>GLE1 CTD•NUP42 GBM<br>Preincubation | 48<br>40<br>47 | 50<br>42<br>92 | n/a<br>equimolar<br>no interaction | SEC-B |
| Fig. S35E | NUP214 NTD<br>SUMO-NUP42 ZnF<br>Preincubation | 48<br>17<br>47 | 50<br>17<br>67 | n/a<br>n/a<br>no interaction | SEC-B<br>2 mM $MgCl_2$<br>1 µM $ZnCl_2$ |
| Fig. S35F | NUP214 NTD<br>RAE1•NUP98 GLEBS<br>Preincubation | 47<br>46<br>47 | 50<br>48<br>92 | n/a<br>equimolar<br>no interaction | SEC-B |
| Fig. S36A | NUP62•NUP88•NUP214 (CFNC-hub)<br>DDX19 (ADP)<br>Preincubation | 69<br>51<br>69 | 69<br>54<br>123 | equimolar<br>n/a<br>no interaction | SEC-B<br>2 mM $MgCl_2$ |
| Fig. S36B | NUP62•NUP88•NUP214 (CFNC-hub)<br>DDX19 E243Q (AMPPNP)<br>Preincubation | 69<br>51<br>71 | 69<br>54<br>123 | equimolar<br>n/a<br>no interaction | SEC-B<br>2 mM $MgCl_2$ |
| Fig. S36C | NUP62•NUP88•NUP214 (CFNC-hub)<br>GLE1 CTD•NUP42 GBM<br>Preincubation | 69<br>40<br>69 | 69<br>42<br>111 | equimolar<br>equimolar<br>no interaction | SEC-B |
| Fig. S36D | NUP62•NUP88•NUP214 (CFNC-hub)<br>SUMO-NUP42 ZnF<br>Preincubation | 69<br>16<br>77 | 69<br>17<br>86 | equimolar<br>n/a<br>no interaction | SEC-B<br>2 mM $MgCl_2$<br>1 µM $ZnCl_2$ |
| Fig. S36E | NUP62•NUP88•NUP214 (CFNC-hub)<br>RAE1•NUP98 GLEBS<br>Preincubation | 69<br>46<br>69 | 69<br>48<br>117 | equimolar<br>equimolar<br>no interaction | SEC-B |
| Fig. S37A | DDX19 (ADP)<br>GLE1 CTD•NUP42 GBM<br>Preincubation | 51<br>69<br>58 | 54<br>42<br>96 | n/a<br>equimolar<br>weak interaction | SEC-B<br>2 mM $MgCl_2$ |
| Fig. S37B | DDX19 E243Q (AMPPNP)<br>GLE1 CTD•NUP42 GBM<br>Preincubation | 51<br>40<br>59 | 54<br>42<br>96 | n/a<br>equimolar<br>weak interaction | SEC-B<br>2 mM $MgCl_2$ |
| Fig. S37C | DDX19 (APD)<br>SUMO-NUP42 ZnF<br>Preincubation | 53<br>17<br>53 | 54<br>17<br>71 | n/a<br>n/a<br>no interaction | SEC-B<br>2 mM $MgCl_2$<br>1 µM $ZnCl_2$ |
| Fig. S37D | DDX19 E243Q (AMPPNP)<br>SUMO-NUP42 ZnF<br>Preincubation | 48<br>17<br>53 | 54<br>17<br>71 | n/a<br>n/a<br>no interaction | SEC-B<br>2 mM $MgCl_2$<br>1 µM $ZnCl_2$ |
| Fig. S37E | DDX19 (ADP)<br>RAE1•NUP98 GLEBS<br>Preincubation | 52<br>46<br>52 | 54<br>48<br>102 | n/a<br>equimolar<br>no interaction | SEC-B<br>2 mM $MgCl_2$ |
| Fig. S37F | DDX19 E243Q (AMPPNP)<br>RAE1•NUP98 GLEBS<br>Preincubation | 52<br>46<br>53 | 54<br>48<br>102 | n/a<br>equimolar<br>no interaction | SEC-B<br>2 mM $MgCl_2$ |
| Fig. S83A | GLE1 CTD•NUP42 GBM<br>SUMO-NUP42 ZnF<br>Preincubation | 41<br>17<br>24 | 42<br>17<br>59 | equimolar<br>n/a<br>no interaction | SEC-B<br>2 mM $MgCl_2$<br>1 µM $ZnCl_2$ |
| Fig. S38B | GLE1 CTD•NUP42 GBM<br>RAE1•NUP98 GLEBS<br>Preincubation | 40<br>46<br>45 | 42<br>48<br>90 | equimolar<br>equimolar<br>no interaction | SEC-B |
| Fig. S39A | SUMO-NUP42 ZnF<br>RAE1•NUP98 GLEBS<br>Preincubation | 16<br>47<br>29 | 17<br>48<br>65 | n/a<br>equimolar<br>no interaction | SEC-B<br>2 mM $MgCl_2$<br>1 µM $ZnCl_2$ |
| Fig. 4B<br>Fig. S41B | NUP358 NTD 2 µM 100 mM NaCl<br>NUP358 NTD 5 µM 100 mM NaCl<br>NUP358 NTD 10 µM 100 mM NaCl<br>NUP358 NTD 20 µM 100 mM NaCl<br>NUP358 NTD 30 µM 100 mM NaCl | 94<br>92<br>95<br>102<br>142 | 86<br>86<br>86<br>86<br>86 | monomer<br>monomer<br>monomer<br>monomer<br>monomer/dimer | SEC-B |
| Fig. 4B<br>Fig. S41C | NUP358 NTD 2 µM 350 mM NaCl<br>NUP358 NTD 5 µM 350 mM NaCl<br>NUP358 NTD 10 µM 350 mM NaCl<br>NUP358 NTD 20 µM 350 mM NaCl<br>NUP358 NTD 30 µM 350 mM NaCl | 81<br>82<br>81<br>82<br>84 | 86<br>86<br>86<br>86<br>86 | monomer<br>monomer<br>monomer<br>monomer<br>monomer | SEC-C |

| Figure(s) | Nucleoporin or nucleoporin complex | Experimental mass (kDa) | Theoretical mass (kDa) | Stoichiometry | Buffer |
|---|---|---|---|---|---|
| Fig. 4B<br>Fig. S41D | NUP358 NTDΔTPR 2 μM 100 mM NaCl<br>NUP358 NTDΔTPR 5 μM 100 mM NaCl<br>NUP358 NTDΔTPR 10 μM 100 mM NaCl<br>NUP358 NTDΔTPR 20 μM 100 mM NaCl<br>NUP358 NTDΔTPR 30 μM 100 mM NaCl<br>NUP358 NTDΔTPR 50 μM 100 mM NaCl | 68<br>67<br>67<br>69<br>69<br>70 | 69<br>69<br>69<br>69<br>69<br>69 | monomer<br>monomer<br>monomer<br>monomer<br>monomer<br>monomer | SEC-B |
| Fig. 4B<br>Fig. S41E | NUP358 NTDΔTPR 2 μM 350 mM NaCl<br>NUP358 NTDΔTPR 5 μM 350 mM NaCl<br>NUP358 NTDΔTPR 10 μM 350 mM NaCl<br>NUP358 NTDΔTPR 20 μM 350 mM NaCl<br>NUP358 NTDΔTPR 30 μM 350 mM NaCl<br>NUP358 NTDΔTPR 50 μM 350 mM NaCl | 65<br>66<br>67<br>68<br>68<br>68 | 69<br>69<br>69<br>69<br>69<br>69 | monomer<br>monomer<br>monomer<br>monomer<br>monomer<br>monomer | SEC-C |
| Fig. 4C<br>Fig. S41A | NUP358 NTD-OE (1-832) 2 μM 350 mM NaCl<br>NUP358 NTD-OE (1-832) 5 μM 350 mM NaCl<br>NUP358 NTD-OE (1-832) 10 μM 350 mM NaCl | 396<br>435<br>449 | 97<br>97<br>97 | tetramer<br>tetramer/pentamer<br>tetramer/pentamer | SEC-C |
| Fig. 4G<br>Fig. S42B | NUP358 OE 150 μM 100 mM NaCl<br>NUP358 OE 375 μM 100 mM NaCl<br>NUP358 OE 750 μM 100 mM NaCl<br>NUP358 OE 1500 μM 100 mM NaCl | 8.1<br>10.4<br>11.6<br>12.5 | 3.8<br>3.8<br>3.8<br>3.8 | dimer<br>dimer/trimer<br>trimer<br>trimer/tetramer | SEC-B |
| Fig. 4G<br>Fig. S42C | NUP358 OE 150 μM 350 mM NaCl<br>NUP358 OE 375 μM 350 mM NaCl<br>NUP358 OE 750 μM 350 mM NaCl<br>NUP358 OE 1500 μM 350 mM NaCl | 9.7<br>11.2<br>12<br>12.7 | 3.8<br>3.8<br>3.8<br>3.8 | dimer/trimer<br>trimer<br>trimer/tetramer<br>trimer/tetramer | SEC-C |
| Fig. 4G<br>Fig. S42D | NUP358 OE LIQIML 150 μM 100 mM NaCl<br>NUP358 OE LIQIML 375 μM 100 mM NaCl<br>NUP358 OE LIQIML 750 μM 100 mM NaCl<br>NUP358 OE LIQIML 1500 μM 100 mM NaCl | 3.4<br>3.4<br>3.4<br>3.4 | 3.8<br>3.8<br>3.8<br>3.8 | monomer<br>monomer<br>monomer<br>monomer | SEC-B |
| Fig. 4G<br>Fig. S42E | NUP358 OE LIQIML 150 μM 350 mM NaCl<br>NUP358 OE LIQIML 375 μM 350 mM NaCl<br>NUP358 OE LIQIML 750 μM 350 mM NaCl<br>NUP358 OE LIQIML 1500 μM 350 mM NaCl | 3.1<br>3.5<br>3.4<br>3.3 | 3.8<br>3.8<br>3.8<br>3.8 | monomer<br>monomer<br>monomer<br>monomer | SEC-C |
| Fig. S42F | NUP358 NTD-OE (1-832) LIQIML 1 μM 100 mM NaCl<br>NUP358 NTD-OE (1-832) LIQIML 2 μM 100 mM NaCl<br>NUP358 NTD-OE (1-832) LIQIML 5 μM 100 mM NaCl<br>NUP358 NTD-OE (1-832) LIQIML 10 μM 100 mM NaCl | 112<br>118<br>123<br>138 | 96<br>96<br>96<br>96 | monomer/dimer<br>monomer/dimer<br>monomer/dimer<br>monomer/dimer | SEC-B |
| Fig. S42G | NUP358 NTD-OE (1-832) LIQIML 1 μM 350 mM NaCl<br>NUP358 NTD-OE (1-832) LIQIML 2 μM 350 mM NaCl<br>NUP358 NTD-OE (1-832) LIQIML 5 μM 350 mM NaCl<br>NUP358 NTD-OE (1-832) LIQIML 10 μM 350 mM NaCl<br>NUP358 NTD-OE (1-832) LIQIML 20 μM 350 mM NaCl<br>NUP358 NTD-OE (1-832) LIQIML 30 μM 350 mM NaCl<br>NUP358 NTD-OE (1-832) LIQIML 50 μM 350 mM NaCl | 90<br>92<br>91<br>93<br>95<br>97<br>102 | 96<br>96<br>96<br>96<br>96<br>96<br>96 | monomer<br>monomer<br>monomer<br>monomer<br>monomer<br>monomer<br>monomer | SEC-C |
| Fig. S54A | Ran (GMPPNP)<br>NUP358 RBD-I<br>Preincubation | 27<br>15<br>38 | 24<br>16<br>40 | n/a<br>n/a<br>equimolar | SEC-B<br>2 mM $MgCl_2$ |
| Fig. S54B | Ran (GMPPNP)<br>NUP358 RBD-II<br>Preincubation | 27<br>16<br>37 | 24<br>16<br>40 | n/a<br>n/a<br>equimolar | SEC-B<br>2 mM $MgCl_2$ |
| Fig. S54C | Ran (GMPPNP)<br>NUP358 RBD-III<br>Preincubation | 27<br>22<br>40 | 24<br>17<br>41 | n/a<br>n/a<br>equimolar | SEC-B<br>2 mM $MgCl_2$ |
| Fig. 4IJ<br>Fig. S54D | Ran (GMPPNP)<br>NUP358 RBD-IV<br>Preincubation | 27<br>20<br>39 | 24<br>16<br>40 | n/a<br>n/a<br>equimolar | SEC-B<br>2 mM $MgCl_2$ |
| Fig. S54E | Ran (GMPPNP)<br>NUP50 RBD<br>Preincubation | 27<br>14<br>26 | 24<br>15<br>39 | n/a<br>n/a<br>equimolar | SEC-B<br>2 mM $MgCl_2$ |
| Fig. 10G<br>Fig. S84F | NUP62•NUP88•NUP214•DDX19 (ADP)•RAE1•NUP98<br>SUMO-NUP93 R1<br>Preincubation | 406<br>22<br>417 | 310<br>24<br>334 | equimolar<br>n/a<br>equimolar | SEC-A |
| Fig. 10H<br>Fig. S84B | His-NUP62•NUP88•AVI-NUP214 CFNC-hub<br>SUMO-NUP93 R1<br>Preincubation | 77<br>22<br>86 | 73<br>24<br>97 | equimolar<br>n/a<br>equimolar | SEC-A |
| Fig. 10I<br>Fig. S84C | His-NUP62•NUP88•AVI-NUP214 CFNC-hub<br>SUMO-NUP93 R1 LIL<br>Preincubation | 77<br>23<br>72 | 73<br>24<br>97 | equimolar<br>n/a<br>no interaction | SEC-A |
| Fig. S84D | NUP62•NUP88•NUP214 CFNC-hub<br>SUMO-NUP93 R1<br>Preincubation | 73<br>22<br>84 | 69<br>24<br>93 | equimolar<br>n/a<br>equimolar | SEC-A |
| Fig. S84E | NUP62•NUP88•NUP214 CFNC-hub<br>SUMO-NUP93 R1 LIL mutation<br>Preincubation | 73<br>30<br>72 | 69<br>24<br>93 | equimolar<br>n/a<br>no interaction | SEC-A |

| Figure(s) | Nucleoporin or nucleoporin complex | Experimental mass (kDa) | Theoretical mass (kDa) | Stoichiometry | Buffer |
|---|---|---|---|---|---|
| Fig. S85A | Nsp1•Nup49•Nup57 CCS1-3 (CNT)<br>SUMO-Nic96 R1<br>Preincubation | 65<br>16<br>86 | 77<br>19<br>97 | equimolar<br>n/a<br>equimolar | SEC-A |
| Fig. S85B | Nup82•Nup159•Nsp1 CFNC-hub<br>SUMO-Nic96 R1<br>Preincubation | 81<br>18<br>79 | 77<br>21<br>98 | equimolar<br>n/a<br>no interaction | SEC-A |
| Fig. S85C | Nsp1•Nup49•Nup57 CCS1-3 (CNT)<br>SUMO-Nup37 CTE<br>Preincubation | 65<br>18<br>65 | 77<br>21<br>96 | equimolar<br>n/a<br>no interaction | SEC-A |

SEC-A: 20 mM TRIS (pH 8.0), 100 mM NaCl, 5% (v/v) glycerol, 5 mM DTT
SEC-B: 20 mM TRIS (pH 8.0), 100 mM NaCl, 5 mM DTT
SEC-C: 20 mM TRIS (pH 8.0), 350 mM NaCl, 5 mM DTT

#### **Table S7.** Crystallization and cryoprotection conditions

| Protein(s) | Concentration | Crystallization condition | Cryo protection condition |
|---|---|---|---|
| NUP358 145-673 | 20 mg/ml | 0.1 M HEPES (pH 7.0)<br>0.8 M succinic acid (pH 7.0)<br>1% (w/v) PEG 2,000 MME | 20% (v/v) ethylene glycol |
| NUP358 145-673<br>SeMet | 20 mg/ml | 0.1 M HEPES (pH 7.0)<br>0.8 M succinic acid (pH 7.0)<br>1% (w/v) PEG 2,000 MME | 20% (v/v) ethylene glycol |
| NUP358 NTD•sAB-14 | 5 mg/ml | 0.1 M sodium citrate (pH 4.6)<br>0.15 M sodium acetate<br>3% (w/v) PEG 4,000 | 25% (v/v) ethylene glycol |
| NUP358 NTD•sAB-14<br>SeMet | 5 mg/ml | 0.1 M sodium citrate (pH 4.6)<br>0.15 M sodium acetate<br>3% (w/v) PEG 4,000 | 25% (v/v) ethylene glycol |
| NUP358 NTD•sAB-14 T585M | 5 mg/ml | 0.1 M sodium citrate (pH 4.9)<br>0.15 M sodium acetate<br>2.5% (w/v) PEG 4,000 | 20% (v/v) ethylene glycol |
| NUP358 NTD•sAB-14 T653I | 5 mg/ml | 0.1 M sodium citrate (pH 4.5)<br>0.15 M sodium acetate<br>4% (w/v) PEG 4,000 | 20% (v/v) ethylene glycol |
| NUP358 NTD•sAB-14 I656V | 5 mg/ml | 0.1 M sodium citrate (pH 4.6)<br>0.15 M sodium acetate<br>3.5% (w/v) PEG 4,000 | 20% (v/v) ethylene glycol |
| NUP358 NTD•sAB-14 (D140M/I599M), (I322M/I599M), (I427M/I599M), (I453M/I599M), (I539M/I599M), (I615M/I599M), (I626M/I599M), (I652M/I599M), (L401M/I599M), (N709M/I599M), (Q137M/I599M), (T92M/I599M), (V668M/I599M)<br>SeMet | 5 mg/ml | 0.1 M sodium citrate (pH 4.9)<br>0.15 M sodium acetate<br>2.5% (w/v) PEG 4,000 | 20% (v/v) ethylene glycol |
| NUP358 NTD•sAB-14 I152M I599M<br>SeMet | 5 mg/ml | 0.1 M sodium citrate (pH 4.5)<br>0.15 M sodium acetate<br>4% (w/v) PEG 4,000 | 20% (v/v) ethylene glycol |
| NUP358 NTD•sAB-14 D693M I599M<br>SeMet | 5 mg/ml | 0.1 M sodium citrate (pH 4.7)<br>0.15 M sodium acetate<br>4% (w/v) PEG 4,000 | 20% (v/v) ethylene glycol |
| NUP358 NTD•sAB-14 I171M I599M<br>SeMet | 5 mg/ml | 0.1 M sodium citrate (pH 4.4)<br>0.15 M sodium acetate<br>2.5% (w/v) PEG 4,000 | 20% (v/v) ethylene glycol |
| NUP358 NTD•sAB-14 I559M I599M<br>SeMet | 5 mg/ml | 0.1 M sodium citrate (pH 4.7)<br>0.15 M sodium acetate<br>6% (w/v) PEG 4,000 | 20% (v/v) ethylene glycol |
| NUP358 OE | 20 mg/ml | 0.1 M citric acid (pH 3.5)<br>2 M ammonium sulfate | 20% (v/v) ethylene glycol |
| NUP358 ZnF2•Ran(GDP) F35S | 15 mg/ml | 0.1M Bis-TRIS (pH 6.4)<br>17 % (w/v) PEG 3,350 | 20% (v/v) glycerol |
| NUP358 ZnF2•Ran(GDP) F35S (Thrombin) | 15 mg/ml | 0.1 M Bis-TRIS (pH 6.5)<br>21% (w/v) PEG 3350 | 20% (v/v) glycerol |
| NUP358 ZnF3•Ran(GDP) F35S (Thrombin) | 15 mg/ml | 0.1 M Bis-TRIS (pH 6.5)<br>19% (w/v) PEG 3350 | 20% (v/v) glycerol |
| NUP358 ZnF4•Ran(GDP) F35S (Thrombin) | 30 mg/ml | 0.1 M Bis-TRIS (pH 6.5)<br>18 % (w/v) PEG 3,350 | 20% (v/v) glycerol |
| NUP358 ZnF5/6•Ran(GDP) F35S (Thrombin) | 30 mg/ml | 0.1 M Bis-TRIS (pH 6.5)<br>19 % (w/v) PEG 3,350 | 20% (v/v) glycerol |
| NUP358 ZnF7•Ran(GDP) F35S (Thrombin) | 15 mg/ml | 0.1 M Bis-TRIS (pH 6.5)<br>19% (w/v) PEG 3350 | 20% (v/v) glycerol |
| NUP358 ZnF8•Ran(GDP) F35S (Thrombin) | 15 mg/ml | 0.1 M Bis-TRIS (pH 6.7)<br>19% (w/v) PEG 3350 | 20% (v/v) glycerol |
| rnNUP153 ZnF1•Ran(GDP) F35S | 15 mg/ml | 0.1 M Bis-TRIS (pH 6.0)<br>22 % (w/v) PEG 3,350 | 20% (v/v) glycerol |
| rnNUP153 ZnF2•Ran(GDP) F35S | 15 mg/ml | 0.1 M Bis-TRIS (pH 6.3)<br>19 % (w/v) PEG 3,350 | 20% (v/v) glycerol |
| rnNUP153 ZnF3•Ran(GDP) F35S | 10 mg/ml | 0.1 M Bis-TRIS (pH 7.0)<br>20 % (w/v) PEG 3,350 | 20% (v/v) glycerol |
| rnNUP153 ZnF4•Ran(GDP) F35S | 15 mg/ml | 0.1 M Bis-TRIS (pH 6.2)<br>22 % (w/v) PEG 3,350 | 20% (v/v) glycerol |
| rnNUP153 ZnF1•Ran(GDP) F35S (Thrombin) | 15 mg/ml | 0.1 M Bis-TRIS (pH 6.5)<br>20% (w/v) PEG 3350 | 20% (v/v) glycerol |
| rnNUP153 ZnF2•Ran(GDP) F35S (Thrombin) | 15 mg/ml | 0.1 M Bis-TRIS (pH 6.5)<br>19% (w/v) PEG 3350 | 20% (v/v) glycerol |
| rnNUP153 ZnF3•Ran(GDP) F35S (Thrombin) | 15 mg/ml | 0.1 M Bis-TRIS (pH 6.5)<br>19% (w/v) PEG 3350 | 20% (v/v) glycerol |
| rnNUP153 ZnF4•Ran(GDP) F35S (Thrombin) | 15 mg/ml | 0.1 M Bis-TRIS (pH 6.5)<br>19% (w/v) PEG 3350 | 20% (v/v) glycerol |
| NUP358 RanBD-I•Ran(GMPPNP) | 8 mg/ml | 0.2 M potassium sodium tartrate tetrahydrate<br>20 % (w/v) PEG 3,350 | 20% (v/v) glycerol |

| Protein(s) | Concentration | Crystallization condition | Cryo protection condition |
|---|---|---|---|
| NUP358 RanBD-II•Ran(GMPPNP) | 20 mg/ml | 0.125 M magnesium formate<br>15 % (w/v) PEG 3,350 | 20% (v/v) ethylene glycol |
| NUP358 RanBD-III•Ran(GMPPNP) | 5 mg/ml | 0.2 M sodium formate<br>20 % (w/v) PEG 3,350 | 25% (v/v) ethylene glycol |
| NUP358 RanBD-IV QNYDNKQV•Ran(GMPPNP) | 3.5 mg/ml | 0.1 M HEPES (pH 7.0)<br>0.1 M ammonium sulfate<br>20 % (w/v) PEG 3,350 | 25% (v/v) glycerol |
| NUP50 RanBD•Ran(GMPPNP) | 20 mg/ml | 0.1 M HEPES (pH 7.5)<br>0.2 M ammonium sulfate<br>25 % (w/v) PEG 3,350 | 20% (v/v) ethylene glycol |
| NUP88 NTD•NUP98 APD | 15 mg/ml | 0.1 M TRIS (pH 8.3)<br>0.22 M NaCl<br>20% (w/v) PEG 4000 | Paratone-N |

**Table S8.** X-ray crystallographic analysis of NUP358$^{145\text{-}673}$

| | | | | |
|---|---|---|---|---|
| **Data collection** | | | | |
| Protein 1 | NUP358$^{145\text{-}673}$ | NUP358$^{145\text{-}673}$ | NUP358$^{145\text{-}673}$ | NUP358$^{145\text{-}673}$ |
| Variant | Wildtype | Wildtype | Wildtype | Wildtype |
| Stoichiometry | 3 | 3 | 3 | 3 |
| PDB code | 7MNJ | | | |
| Synchrotron | SSRL[a] | SSRL | SSRL | SSRL |
| Beamline | BL 12-2 | BL 12-2 | BL 12-2 | BL 12-2 |
| Space group | $P4_32_12$ | $P4_32_12$ | $P4_32_12$ | $P4_32_12$ |
| Cell parameters | | | | |
| *a* (Å) | 133.5 | 136.4 | 136.6 | 136.3 |
| *b* (Å) | 133.5 | 136.4 | 136.6 | 136.3 |
| *c* (Å) | 286.9 | 289.6 | 289.6 | 288.9 |
| α (°) | 90 | 90 | 90 | 90 |
| β (°) | 90 | 90 | 90 | 90 |
| γ (°) | 90 | 90 | 90 | 90 |
| | *Native* | *Se peak* | *Se inflection* | *Se remote* |
| Wavelength (Å) | 1.03320 | 0.97920 | 0.97940 | 0.95370 |
| Resolution (Å) | 30.0 – 3.8 | 30.0 – 4.2 | 30.0 – 4.3 | 30.0 – 4.4 |
| $R_{merge}$ (%)[c] | 8.7 (148.6) | 13.7 (176.4) | 14.3 (163.9) | 16.2 (196.3) |
| $R_{meas}$ (%)[c] | 9.0 (155.0) | 14.3 (183.2) | 14.9 (170.4) | 16.8 (205.0) |
| $R_{pim}$ (%)[c] | 2.4 (43.6) | 3.9 (49.3) | 4.0 (46.2) | 4.6 (58.6) |
| $CC_{1/2}$ (%)[c] | 99.9 (93.5) | 100.0 (84.5) | 100.0 (85.6) | 100.0 (75.5) |
| $\langle I \rangle / \langle \sigma I \rangle$[c] | 18.3 (2.1) | 15.6 (2.3) | 15.3 (2.5) | 14.1 (1.9) |
| Completeness (%)[c] | 100.0 (98.8) | 100.0 (100.0) | 100.0 (100.0) | 100.0 (100.0) |
| No. observations[c] | 654,682 (60,258) | 517,269 (52,138) | 481,005 (47,613) | 444,858 (40,060) |
| No. unique reflections[c,d] | 26,310 (2,545) | 20,692 (2,008) | 19,360 (1,872) | 17,981 (1,752) |
| Redundancy[c] | 24.9 (23.7) | 25.0 (26.0) | 24.8 (25.4) | 24.7 (22.9) |
| Wilson B factor | 155 | 171 | 172 | 182 |
| **Refinement** | | | | |
| Resolution (Å) | 30.0 – 3.8 | | | |
| No. reflections total | 26,172 | | | |
| No. reflections test set | 1,292 (4.9%) | | | |
| $R_{work}$ / $R_{free}$ (%) | 20.8 / 24.2 | | | |
| No. atoms | 12,695 | | | |
| Protein | 12,695 | | | |
| Ligand | 0 | | | |
| Water | 0 | | | |
| *B*-factors | 210 | | | |
| Protein | 210 | | | |
| Ligand | | | | |
| Water | | | | |
| R.m.s. deviations | | | | |
| Bond lengths (Å) | 0.002 | | | |
| Bond angles (°) | 0.4 | | | |
| **Ramachandran plot**[e] | | | | |
| Favored (%) | 98.29 | | | |
| Additionally allowed (%) | 1.71 | | | |
| Outliers (%) | 0 | | | |

[a] SSRL, Stanford Synchrotron Radiation Lightsource
[b] APS, Advanced Photon Source
[c] Highest-resolution shell is shown in parentheses
[d] Unique reflections distinguish anomalous Friedel pairs
[e] As determined by MolProbity (*128*)

**Table S9.** X-ray crystallographic analysis of NUP358$^{145\text{-}673}$ seleno-L-methionine mutations

| **Data collection** | | | | | | | |
|---|---|---|---|---|---|---|---|
| Protein | NUP358$^{145\text{-}673}$ I652M | NUP358$^{145\text{-}673}$ I615M/I599M | NUP358$^{145\text{-}673}$ I599M | NUP358$^{145\text{-}673}$ L401M/I599M | NUP358$^{145\text{-}673}$ I169M/I481M | NUP358$^{145\text{-}673}$ L385M/I599M | NUP358$^{145\text{-}673}$ I372M/I599M |
| Variant | I652M | I615M/I599M | I599M | L401M/I599M | I169M/I481M | L385M/I599M | I372M/I599M |
| Stoichiometry | 3 | 3 | 3 | 3 | 3 | 3 | 3 |
| PDB code | | | | | | | |
| Synchrotron | SSRL[a] | APS[b] | SSRL | SSRL | SSRL | SSRL | APS |
| Beamline | BL 12-2 | 23 ID-D | BL 12-2 | BL 12-2 | BL 12-2 | BL 12-2 | 23 ID-D |
| Space group | $P4_32_12$ | $P4_32_12$ | $P4_32_12$ | $P4_32_12$ | $P4_32_12$ | $P4_32_12$ | $P4_32_12$ |
| Cell parameters | | | | | | | |
| *a* (Å) | 133.6 | 135.6 | 134.0 | 136.2 | 133.8 | 133.1 | 135.5 |
| *b* (Å) | 133.6 | 135.6 | 134.0 | 136.2 | 133.8 | 133.1 | 135.5 |
| *c* (Å) | 291.3 | 289.8 | 288.3 | 291.6 | 291.5 | 286.8 | 289.6 |
| α (°) | 90 | 90 | 90 | 90 | 90 | 90 | 90 |
| β (°) | 90 | 90 | 90 | 90 | 90 | 90 | 90 |
| γ (°) | 90 | 90 | 90 | 90 | 90 | 90 | 90 |
| | *Se peak* | *Se peak* | *Se peak* | *Se peak* | *Se peak* | *Se peak* | *Se peak* |
| Wavelength (Å) | 0.97910 | 0.97936 | 0.97950 | 0.97920 | 0.97950 | 0.97920 | 0.97936 |
| Resolution (Å) | 30.0 – 4.5 | 30.0 – 5.2 | 30.0 – 6.3 | 30.0 – 6.3 | 30.0 – 4.7 | 30.0 – 5.5 | 30.0 – 5.0 |
| $R_{merge}$ (%)[c] | 24.3 (388.5) | 28.9 (502.0) | 27.6 (258.0) | 16.9 (527.2) | 18.3 (152.9) | 19.3 (375.4) | 27.5 (517.9) |
| $R_{meas}$ (%)[c] | 24.8 (396.2) | 29.6 (512.9) | 28.3 (264.1) | 17.2 (538.9) | 19.0 (159.3) | 19.6 (382.9) | 28.1 (529.1) |
| $R_{pim}$ (%)[c] | 4.7 (76.1) | 6.1 (105.5) | 6.0 (56.1) | 3.3 (109.9) | 5.2 (44.5) | 3.8 (72.6) | 5.8 (108.6) |
| $CC_{1/2}$ (%)[c] | 99.9 (60.3) | 99.9 (73.2) | 99.7 (86.8) | 99.9 (70.5) | 99.9 (89.4) | 99.9 (72.6) | 100.0 (71.2) |
| $\langle I \rangle / \langle \sigma I \rangle$[c] | 15.3 (1.6) | 16.0 (1.3) | 11.7 (1.4) | 20.7 (1.3) | 9.7 (1.0) | 19.3 (1.9) | 16.8 (1.3) |
| Completeness (%)[c] | 100.0 (100.0) | 100.0 (100.0) | 100.0 (100.0) | 99.0 (88.3) | 100.0 (100.0) | 100.0 (99.8) | 100.0 (100.0) |
| No. observations[c] | 833,414 (77,052) | 469,578 (46,501) | 232,212 (23,396) | 299,622 (23,303) | 350,720 (33,574) | 437,310 (43,864) | 527,226 (52,764) |
| No. unique reflections[c,d] | 16,277 (1,572) | 10,962 (1,061) | 6,065 (589) | 6,259 (549) | 14,414 (1,405) | 8,883 (857) | 12,270 (1,196) |
| Redundancy[c] | 51.2 (49.0) | 42.8 (43.8) | 38.3 (39.7) | 47.9 (42.4) | 24.3 (23.9) | 49.2 (51.2) | 43.0 (44.1) |
| Wilson B factor | 225 | 246 | 308 | 440 | 214 | 317 | 238 |

[a] SSRL, Stanford Synchrotron Radiation Lightsource
[b] APS, Advanced Photon Source
[c] Highest-resolution shell is shown in parentheses
[d] Unique reflections distinguish anomalous Friedel pairs
[e] As determined by MolProbity (*128*)

**Table S10.** X-ray crystallographic analysis of NUP358$^{NTD}$ and NUP358$^{NTD}$ ANE1 mutants

| | | | | |
|---|---|---|---|---|
| **Data collection** | | | | |
| Protein 1 | NUP358$^{1\text{-}752}$ | NUP358$^{1\text{-}752}$ T585M | NUP358$^{1\text{-}752}$ T653I | NUP358$^{1\text{-}752}$ I656V |
| Variant | Wildtype | ANE1 T585M | ANE1 T653I | ANE1 I656V |
| Protein 2 | SAB-14 | SAB-14 | SAB-14 | SAB-14 |
| Stoichiometry | 2:2 | 2:2 | 2:2 | 2:2 |
| PDB code | 7MNL | 7MNM | 7MNN | 7MNO |
| Synchrotron | APS[a] | APS[a] | SSRL[b] | SSRL[b] |
| Beamline | 23 ID-B | 23 ID-B | BL 12-2 | BL 12-2 |
| Space group | $P6_522$ | $P6_522$ | $P6_522$ | $P6_522$ |
| Cell parameters | | | | |
| *a* (Å) | 161.4 | 160.4 | 161.5 | 161.6 |
| *b* (Å) | 161.4 | 160.4 | 161.5 | 161.6 |
| *c* (Å) | 644.7 | 642.2 | 647.4 | 645.4 |
| α (°) | 90.0 | 90.0 | 90.0 | 90.0 |
| β (°) | 90.0 | 90.0 | 90.0 | 90.0 |
| γ (°) | 120.0 | 120.0 | 120.0 | 120.0 |
| | *Native* | *Native* | *Native* | *Native* |
| Wavelength (Å) | 1.03314 | 1.03318 | 0.97946 | 1.03317 |
| Resolution (Å) | 20.0 – 3.95 | 30.0 – 4.7 | 30.0 – 6.7 | 30.0 – 6.7 |
| $R_{merge}$ (%)[c] | 37.9 (588.1) | 58.6 (481.8) | 77.9 (637.8) | 55.4 (321.5) |
| $R_{meas}$ (%)[c] | 38.4 (595.9) | 59.4 (488.1) | 78.3 (640.9) | 56.2 (325.9) |
| $R_{pim}$ (%)[c] | 6.0 (95.1) | 9.4 (77.8) | 7.7 (62.1) | 9.1 (52.9) |
| $CC_{1/2}$ (%)[c] | 99.9 (45.5) | 99.9 (58.6) | 99.6 (67.5) | 99.3 (62.1) |
| $\langle I \rangle / \langle \sigma I \rangle$[c] | 10.3 (1.0) | 6.6 (1.1) | 11.0 (1.5) | 8.5 (1.6) |
| Completeness (%)[c] | 99.2 (99.8) | 99.6 (100.0) | 98.9 (100.0) | 98.6 (98.9) |
| No. observations[c] | 1,772,720 (176,902) | 1,037,121 (182,414) | 955,034 (275,234) | 348,970 (95,598) |
| No. unique reflections[c,d] | 44,627 (4,585) | 26,575 (4,697) | 9,674 (2,677) | 9,487 (2,593) |
| Redundancy[c] | 39.7 (38.6) | 39.0 (38.8) | 98.7 (102.8) | 36.8 (36.9) |
| Wilson B factor | 181 | 295 | 549 | 398 |
| **Refinement** | | | | |
| Resolution (Å) | 20.0 – 3.95 | 30.0 – 4.7 | 30.0 – 6.7 | 30.0 – 6.7 |
| No. reflections total | 44,483 | 26,442 | 9,591 | 9,417 |
| No. reflections test set | 2,213 (5.0%) | 1,304 (4.9%) | 456 (4.8%) | 445 (4.7%) |
| $R_{work}$ / $R_{free}$ (%) | 19.2 / 24.1 | 20.4 / 25.4 | 21.7 / 26.3 | 20.5 / 25.4 |
| No. atoms | 17,358 | 17,453 | 17,499 | 17,474 |
| Protein | 17,358 | 17,453 | 17,499 | 17,474 |
| Ligand | 0 | 0 | 0 | 0 |
| Water | 0 | 0 | 0 | 0 |
| *B*-factors | 204 | 246 | 331 | 317 |
| Protein | 204 | 246 | 331 | 317 |
| Ligand | | | | |
| Water | | | | |
| R.m.s. deviations | | | | |
| Bond lengths (Å) | 0.002 | 0.002 | 0.003 | 0.002 |
| Bond angles (°) | 0.5 | 0.5 | 0.6 | 0.6 |
| **Ramachandran plot**[e] | | | | |
| Favored (%) | 98.68 | 98.33 | 98.38 | 98.33 |
| Additionally allowed (%) | 1.32 | 1.67 | 1.62 | 1.67 |
| Outliers (%) | 0 | 0 | 0 | 0 |

[a] APS, Advanced Photon Source
[b] SSRL, Stanford Synchrotron Radiation Lightsource
[c] Highest-resolution shell is shown in parentheses
[d] Unique reflections distinguish anomalous Friedel pairs
[e] As determined by MolProbity (*128*)

**Table S11.** X-ray crystallographic analysis of NUP358$^{NTD}$ seleno-L-methionine mutations

| **Data collection** | | | | | | | | | |
|---|---|---|---|---|---|---|---|---|---|
| Protein 1 | NUP358$^{1-752}$ D140M I599M | NUP358$^{1-752}$ D152M/I599M | NUP358$^{1-752}$ D693M/I599M | NUP358$^{1-752}$ I171M/I599M | NUP358$^{1-752}$ I322M/I599M | NUP358$^{1-752}$ I427M/I599M | NUP358$^{1-752}$ I453M/I599M | NUP358$^{1-752}$ I539M/I599M | NUP358$^{1-752}$ I559M/I599M |
| Variant | D140M/I599M | D152M/I599M | D693M/I599M | I171M/I599M | I322M/I599M | I427M/I599M | I453M/I599M | I539M/I599M | I559M/I599M |
| Protein 2 | SAB-14 | SAB-14 | SAB-14 | SAB-14 | SAB-14 | SAB-14 | SAB-14 | SAB-14 | SAB-14 |
| Stoichiometry | 2:2 | 2:2 | 2:2 | 2:2 | 2:2 | 2:2 | 2:2 | 2:2 | 2:2 |
| PDB code | | | | | | | | | |
| Synchrotron | APS[a] | APS[a] | SSRL[b] | APS[a] | SSRL[b] | SSRL[b] | SSRL[b] | SSRL[b] | SSRL[b] |
| Beamline | 23 ID-B | 23 ID-B | BL 12-2 | 23 ID-D | BL 12-2 | BL 12-2 | BL 12-2 | BL 12-2 | BL 12-2 |
| Space group | $P6_522$ | $P6_522$ | $P6_522$ | $P6_522$ | $P6_522$ | $P6_522$ | $P6_522$ | $P6_522$ | $P6_522$ |
| Cell parameters | | | | | | | | | |
| *a* (Å) | 160.2 | 161.0 | 160.5 | 160.6 | 160.9 | 161.8 | 160.4 | 161.9 | 160.8 |
| *b* (Å) | 160.2 | 161.0 | 160.5 | 160.6 | 160.9 | 161.8 | 160.4 | 161.9 | 160.8 |
| *c* (Å) | 639.9 | 644.0 | 643.7 | 641.9 | 642.7 | 645.4 | 642.9 | 646.4 | 643.7 |
| $\alpha$ (°) | 90.0 | 90.0 | 90.0 | 90.0 | 90.0 | 90.0 | 90.0 | 90.0 | 90.0 |
| $\beta$ (°) | 90.0 | 90.0 | 90.0 | 90.0 | 90.0 | 90.0 | 90.0 | 90.0 | 90.0 |
| $\gamma$ (°) | 120.0 | 120.0 | 120.0 | 120.0 | 120.0 | 120.0 | 120.0 | 120.0 | 120.0 |
| | *Se peak* | *Se peak* | *Se peak* | *Se peak* | *Se peak* | *Se peak* | *Se peak* | *Se peak* | *Se peak* |
| Wavelength (Å) | 0.97923 | 0.97922 | 0.97910 | 0.97937 | 0.97944 | 0.97921 | 0.97920 | 0.97929 | 0.97949 |
| Resolution (Å) | 30.0 – 6.0 | 30.0 – 7.0 | 30.0 – 4.7 | 30.0 – 7.5 | 30.0 – 5.7 | 30.0 – 6.3 | 30.0 – 5.3 | 30.0 – 4.25 | 30.0 – 4.3 |
| $R_{merge}$ (%)[c] | 35.1 (267.6) | 47.9 (256.3) | 30.3 (322.9) | 45.5 (115.4) | 42.1 (345.3) | 40.9 (247.2) | 30.8 (305.0) | 37.5 (389.4) | 49.3 (365.4) |
| $R_{meas}$ (%)[c] | 35.7 (271.4) | 48.6 (259.8) | 30.7 (327.1) | 46.6 (117.9) | 42.7 (349.7) | 41.5 (250.3) | 31.3 (309.3) | 38.0 (394.3) | 49.9 (370.3) |
| $R_{pim}$ (%)[c] | 6.0 (44.6) | 8.0 (42.0) | 4.9 (51.4) | 9.9 (24.2) | 6.9 (55.0) | 6.7 (39.4) | 5.3 (51.1) | 6.0 (61.4) | 7.9 (59.0) |
| $CC_{1/2}$ (%)[c] | 99.7 (77.2) | 99.3 (76.7) | 99.9 (62.6) | 98.4 (90.9) | 99.7 (63.5) | 99.5 (77.9) | 99.8 (78.8) | 99.8 (62.3) | 99.8 (65.8) |
| $\langle I\rangle / \langle\sigma I\rangle$[c] | 9.5 (1.9) | 7.3 (1.8) | 12.3 (1.6) | 6.0 (3.0) | 9.8 (1.6) | 10.2 (2.1) | 11.9 (1.7) | 9.7 (1.4) | 8.6 (1.5) |
| Completeness (%)[c] | 99.1 (99.6) | 98.7 (100.0) | 99.4 (99.1) | 98.4 (100.0) | 99.1 (99.4) | 99.0 (99.9) | 98.8 (98.9) | 99.7 (99.7) | 99.7 (99.8) |
| No. observations[c] | 440,163 (129,500) | 303,958 (86,803) | 1,020,322 (184,556) | 145,273 (43,416) | 572,998 (166,519) | 430,299 (125,907) | 630,389 (151,852) | 1,411,824 (173,500) | 1,352,239 (173,737) |
| No. unique reflections[c,d] | 13,015 (3,602) | 8,352 (2,310) | 26,547 (4,651) | 6,820 (1,890) | 15,278 (4,225) | 11,444 (3,181) | 18,541 (4,290) | 36,253 (4,269) | 34,750 (4,507) |
| Redundancy[c] | 33.8 (36.0) | 36.4 (37.6) | 38.4 (39.7) | 21.3 (23.0) | 37.5 (39.4) | 37.6 (39.6) | 34.0 (35.4) | 38.9 (40.6) | 38.9 (38.5) |
| Wilson B factor | 387 | 662 | 324 | 516 | 359 | 364 | 349 | 238 | 242 |

[a] APS, Advanced Photon Source
[b] SSRL, Stanford Synchrotron Radiation Lightsource
[c] Highest-resolution shell is shown in parentheses
[d] Unique reflections distinguish anomalous Friedel pairs
[e] As determined by MolProbity (*128*)

**Table S12.** X-ray crystallographic analysis of NUP358$^{NTD}$ seleno-L-methionine mutations

| **Data collection** | | | | | | | | |
|---|---|---|---|---|---|---|---|---|
| Protein 1 | NUP358$^{1-752}$ I615M/I599M | NUP358$^{1-752}$ I626M/I599M | NUP358$^{1-752}$ I652M/I599M | NUP358$^{1-752}$ L401M/I599M | NUP358$^{1-752}$ N709M/I599M | NUP358$^{1-752}$ Q137M/I599M | NUP358$^{1-752}$ T92M/I599M | NUP358$^{1-752}$ V668M/I599M |
| Variant | I615M/I599M | I626M/I599M | I652M/I599M | L401M/I599M | N709M/I599M | Q137M/I599M | T92M/I599M | V668M/I599M |
| Protein 2 | SAB-14 | SAB-14 | SAB-14 | SAB-14 | SAB-14 | SAB-14 | SAB-14 | SAB-14 |
| Stoichiometry | 2:2 | 2:2 | 2:2 | 2:2 | 2:2 | 2:2 | 2:2 | 2:2 |
| PDB code | | | | | | | | |
| Synchrotron | SSRL[b] | SSRL[b] | SSRL[b] | SSRL[b] | SSRL[b] | SSRL[b] | SSRL[b] | SSRL[b] |
| Beamline | BL 12-2 | BL 12-2 | BL 12-2 | BL 12-2 | BL 12-2 | BL 12-2 | BL 12-2 | BL 12-2 |
| Space group | $P6_522$ | $P6_522$ | $P6_522$ | $P6_522$ | $P6_522$ | $P6_522$ | $P6_522$ | $P6_522$ |
| Cell parameters | | | | | | | | |
| *a* (Å) | 160.5 | 160.5 | 160.8 | 161.3 | 161.6 | 161.2 | 160.9 | 160.2 |
| *b* (Å) | 160.5 | 160.5 | 160.8 | 161.3 | 161.6 | 161.2 | 160.9 | 160.2 |
| *c* (Å) | 650.7 | 645.1 | 645.8 | 644.2 | 636.4 | 645.2 | 648.1 | 647.1 |
| α (°) | 90.0 | 90.0 | 90.0 | 90.0 | 90.0 | 90.0 | 90.0 | 90.0 |
| β (°) | 90.0 | 90.0 | 90.0 | 90.0 | 90.0 | 90.0 | 90.0 | 90.0 |
| γ (°) | 120.0 | 120.0 | 120.0 | 120.0 | 120.0 | 120.0 | 120.0 | 120.0 |
| | *Se peak* | *Se peak* | *Se peak* | *Se peak* | *Se peak* | *Se peak* | *Se peak* | *Se peak* |
| Wavelength (Å) | 0.97947 | 0.97949 | 0.97951 | 0.97929 | 0.97922 | 0.97922 | 0.97949 | 0.97910 |
| Resolution (Å) | 30.0 – 6.5 | 30.0 – 6.4 | 30.0 – 5.8 | 30.0 – 5.8 | 30.0 – 5.9 | 30.0 – 6.5 | 30.0 – 5.6 | 30.0 – 4.5 |
| $R_{merge}$ (%)[c] | 38.6 (262.6) | 35.3 (203.3) | 44.5 (244.5) | 39.2 (331.8) | 36.6 (286.8) | 32.3 (210.3) | 39.8 (329.3) | 36.5 (322.9) |
| $R_{meas}$ (%)[c] | 39.1 (266.3) | 35.8 (206.1) | 45.6 (250.1) | 39.7 (336.0) | 37.1 (290.8) | 33.0 (214.5) | 40.3 (333.5) | 37.0 (326.8) |
| $R_{pim}$ (%)[c] | 6.3 (43.5) | 5.8 (33.2) | 9.9 (52.0) | 6.4 (52.9) | 6.2 (47.8) | 6.4 (41.3) | 6.4 (52.0) | 5.8 (50.2) |
| $CC_{1/2}$ (%)[c] | 99.6 (75.2) | 99.7 (81.9) | 99.3 (67.0) | 99.7 (65.3) | 99.6 (66.8) | 99.6 (7.6) | 99.7 (72.4) | 99.9 (67.8) |
| <*I*> / <σ*I*>[c] | 9.9 (1.9) | 10.4 (2.3) | 5.8 (1.5) | 10.0 (1.5) | 9.2 (1.8) | 8.9 (1.9) | 9.3 (1.6) | 10.7 (1.6) |
| Completeness (%)[c] | 98.7 (98.9) | 98.8 (99.2) | 99.2 (99.6) | 98.9 (99.1) | 99.1 (100.0) | 99.0 (100.0) | 99.3 (99.9) | 99.3 (99.9) |
| No. observations[c] | 385,426 (105,437) | 397,949 (111,545) | 300,981 (90,375) | 539,571 (156,204) | 481,226 (138,712) | 270,195 (75,397) | 613,880 (178,204) | 1,180,062 (177,326) |
| No. unique reflections[c,d] | 10,447 (2,890) | 10,815 (2,968) | 14,546 (4,014) | 14,449 (3,971) | 13,790 (3,819) | 10,376 (2,885) | 16,129 (4,450) | 30,272 (4,318) |
| Redundancy[c] | 36.9 (36.5) | 36.8 (37.6) | 20.7 (22.5) | 37.3 (39.3) | 34.9 (36.3) | 26.0 (26.1) | 38.1 (40.0) | 39.0 (41.1) |
| Wilson B factor | 366 | 448 | 334 | 419 | 414 | 479 | 356 | 268 |

[a] APS, Advanced Photon Source
[b] SSRL, Stanford Synchrotron Radiation Lightsource
[c] Highest-resolution shell is shown in parentheses
[d] Unique reflections distinguish anomalous Friedel pairs
[e] As determined by MolProbity (*128*)

**Table S13.** X-ray crystallographic analysis of NUP358$^{OE}$

| | | |
|---|---|---|
| **Data collection** | | |
| Protein 1 | NUP358$^{805-832}$ OE | NUP358$^{805-832}$ OE |
| Protein 2 | | |
| Stoichiometry | 4 | 4 |
| PDB code | 7MNK | |
| Synchrotron | SSRL[b] | SSRL[b] |
| Beamline | BL 12-2 | BL 12-2 |
| Space group | I422 | I422 |
| Cell parameters | | |
| *a* (Å) | 99.9 | 99.9 |
| *b* (Å) | 99.9 | 99.9 |
| *c* (Å) | 60.2 | 60.2 |
| α (°) | 90.0 | 90.0 |
| β (°) | 90.0 | 90.0 |
| γ (°) | 90.0 | 90.0 |
| | *Native* | *S peak* |
| Wavelength (Å) | 0.90000 | 1.30000 |
| Resolution (Å) | 30.0 – 1.1 | 20.0 – 1.45 |
| $R_{merge}$ (%)[c] | 6.0 (294.8) | 5.2 (16.2) |
| $R_{meas}$ (%)[c] | 6.3 (307.6) | 5.4 (17.9) |
| $R_{pim}$ (%)[c] | 1.7 (86.8) | 1.5 (7.5) |
| $CC_{1/2}$ (%)[c] | 100.0 (55.3) | 100.0 (98.9) |
| $\langle I \rangle / \langle \sigma I \rangle$[c] | 24.1 (1.1) | 27.0 (9.5) |
| Completeness (%)[c] | 100.0 (99.9) | 99.9 (87.4) |
| No. observations[c] | 1,558,137 (144,840) | 51,021 (2,360) |
| No. unique reflections[c,d] | 61,560 (6,077) | 26,904 (1,123) |
| Redundancy[c] | 25.3 (23.8) | 1.9 (1.9) |
| Wilson B factor | 14 | 11 |
| **Refinement** | | |
| Resolution (Å) | 30.0 – 1.1 | |
| No. reflections total | 61,541 | |
| No. reflections test set | 2,090 (3.4%) | |
| $R_{work}$ / $R_{free}$ (%) | 15.5 / 16.7 | |
| No. atoms | 1,487 | |
| Protein | 1,294 | |
| Ligand | 55 | |
| Water | 138 | |
| *B*-factors | 23 | |
| Protein | 21 | |
| Ligand | 49 | |
| Water | 36 | |
| R.m.s. deviations | | |
| Bond lengths (Å) | 0.012 | |
| Bond angles (°) | 1.3 | |
| **Ramachandran plot**[e] | | |
| Favored (%) | 100.00 | |
| Additionally allowed (%) | 0 | |
| Outliers (%) | 0 | |

[a] APS, Advanced Photon Source
[b] SSRL, Stanford Synchrotron Radiation Lightsource
[c] Highest-resolution shell is shown in parentheses
[d] Unique reflections distinguish anomalous Friedel pairs
[e] As determined by MolProbity (*128*)

#### **Table S14.** X-ray crystallographic analysis of NUP358 ZnF•Ran(GDP) complexes

| | | | | | | | |
|---|---|---|---|---|---|---|---|
| **Data collection** | | | | | | | |
| Protein 1 | Ran(GDP) F35S | Ran(GDP) F35S | Ran(GDP) F35S | Ran(GDP) F35S | Ran(GDP) F35S | Ran(GDP) F35S | Ran(GDP) F35S |
| Expression construct | Sumo | Thrombin | Thrombin | Thrombin | Thrombin | Thrombin | Thrombin |
| Protein 2 | $NUP358^{1407-1443}$ ZnF2 | $NUP358^{1407-1443}$ ZnF2 | $NUP358^{1471-1507}$ ZnF3 | $NUP358^{1535-1571}$ ZnF4 | $NUP358^{1598-1634}$ ZnF5 $NUP358^{1657-1693}$ ZnF6 | $NUP358^{1716-1752}$ ZnF7 | $NUP358^{1774-1809}$ ZnF8 |
| Stoichiometry | 2:2 | 1:1 | 1:1 | 1:1 | 2:2 | 1:1 | 1:1 |
| PDB code | 7MNP | 7MNQ | 7MNR | 7MNS | 7MNT | 7MNU | 7MNV |
| Synchrotron | SSRL[a] | SSRL[a] | APS[b] | SSRL[a] | APS[b] | SSRL[a] | SSRL[a] |
| Beamline | BL12-2 | BL12-2 | 23 ID-B | BL12-2 | 23 ID-B | BL12-2 | BL12-2 |
| Space group | $P2_12_12_1$ | $P2_12_12_1$ | $P2_12_12$ | $P6_522$ | $P2_12_12_1$ | $P6_522$ | $P2_12_12$ |
| Cell parameters | | | | | | | |
| *a* (Å) | 60.2 | 59.8 | 59.6 | 92.5 | 53.0 | 92.7 | 60.5 |
| *b* (Å) | 79.8 | 78.7 | 80.2 | 92.5 | 94.3 | 92.7 | 80.5 |
| *c* (Å) | 108.8 | 56.3 | 54.8 | 194.8 | 108.2 | 196.0 | 58.3 |
| α (°) | 90.0 | 90.0 | 90.0 | 90.0 | 90.0 | 90.0 | 90.0 |
| β (°) | 90.0 | 90.0 | 90.0 | 90.0 | 90.0 | 90.0 | 90.0 |
| γ (°) | 90.0 | 90.0 | 90.0 | 120.0 | 90.0 | 120.0 | 90.0 |
| | *Native* | *Zn peak* | *Zn peak* | *Zn peak* | *Zn peak* | *Zn peak* | *Zn peak* |
| Wavelength (Å) | 1.03317 | 1.28291 | 1.28238 | 1.28210 | 1.28283 | 1.28275 | 1.28247 |
| Resolution (Å) | 30.0 – 2.05 | 30.0 – 2.05 | 30.0 – 1.8 | 30.0 – 2.1 | 30.0 – 2.45 | 30.0 – 2.0 | 30.0 – 1.8 |
| $R_{merge}$ (%)[c] | 11.5 (131.6) | 9.4 (191.6) | 7.8 (98.6) | 17.2 (223.5) | 11.0 (317.2) | 15.5 (221.4) | 10.8 (174.1) |
| $R_{meas}$ (%)[c] | 12.6 (147.0) | 10.2 (209.7) | 9.2 (124.2) | 17.9 (232.0) | 11.4 (329.4) | 16.4 (235.4) | 11.7 (189.4) |
| $R_{pim}$ (%)[c] | 5.2 (63.9) | 3.9 (83.7) | 4.9 (74.6) | 4.8 (62.0) | 3.1 (88.6) | 5.4 (79.6) | 4.5 (73.7) |
| $CC_{1/2}$ (%)[c] | 99.9 (74.1) | 99.9 (60.3) | 99.8 (57.5) | 99.9 (66.4) | 99.9 (54.1) | 99.9 (63.2) | 99.9 (58.5) |
| $\langle I \rangle / \langle \sigma I \rangle$[c] | 12.5 (1.6) | 13.9 (1.2) | 13.7 (1.3) | 21.2 (1.8) | 21.8 (1.6) | 14.0 (1.4) | 14.0 (1.2) |
| Completeness (%)[c] | 99.4 (95.3) | 99.4 (95.6) | 99.7 (95.8) | 100.0 (99.9) | 100.0 (100.0) | 100.0 (100.0) | 99.6 (96.4) |
| No. observations[c] | 326,611 (28,669) | 215,148 (17,443) | 153,861 (9,866) | 754,508 (73,905) | 524,547 (52,237) | 584,705 (55,080) | 341,183 (31,307) |
| No. unique reflections[c,d] | 33,217 (3,140) | 17,126 (1,596) | 24,915 (2,354) | 29,596 (2,860) | 20,617 (2,020) | 34,531 (3,346) | 26,934 (2,561) |
| Redundancy[c] | 10.9 (9.1) | 12.6 (10.9) | 6.2 (4.2) | 25.5 (25.8) | 25.4 (25.9) | 16.9 (16.5) | 12.7 (12.2) |
| Wilson B factor | 30 | 40 | 23 | 32 | 66 | 31 | 25 |
| **Refinement** | | | | | | | |
| Resolution (Å) | 30.0 – 2.05 | 30.0 – 2.05 | 30.0 – 1.8 | 30.0 – 2.1 | 30.0 – 2.45 | 30.0 – 2.0 | 30.0 – 1.8 |
| No. reflections total | 33,084 | 17,055 | 24,867 | 29,526 | 20,568 | 34,459 | 26,877 |
| No. reflections test set | 3,045 (4.88%) | 891 (5.22%) | 1,230 (4.95%) | 1,477 (5.00 %) | 1,019 (4.95%) | 1,784 (5.18%) | 1,422 (5.29%) |
| $R_{work}$ / $R_{free}$ | 19.4 / 23.1 | 19.0 / 20.7 | 16.0 / 19.6 | 17.5 / 20.3 | 21.2 / 25.7 | 20.1 / 22.4 | 15.9 / 18.6 |
| No. atoms | 4,194 | 2,009 | 2,173 | 2,234 | 3,882 | 2,157 | 2,181 |
| Protein | 3,868 | 1,914 | 1,931 | 2,045 | 3,795 | 1,931 | 1,945 |
| Ligand | 60 | 30 | 30 | 30 | 60 | 30 | 36 |
| Water | 266 | 65 | 212 | 159 | 27 | 196 | 200 |
| *B*-factors | 45 | 65 | 37 | 48 | 98 | 45 | 40 |
| Protein | 46 | 66 | 37 | 48 | 99 | 45 | 40 |
| Ligand | 34 | 51 | 26 | 34 | 84 | 32 | 35 |
| Water | 42 | 54 | 40 | 51 | 75 | 48 | 43 |
| R.m.s. deviations | | | | | | | |
| Bond lengths (Å) | 0.002 | 0.002 | 0.011 | 0.004 | 0.003 | 0.002 | 0.009 |
| Bond angles (°) | 0.6 | 0.6 | 1.1 | 0.7 | 0.6 | 0.6 | 0.9 |
| **Ramachandran plot**[e] | | | | | | | |
| Favored (%) | 97.67 | 98.26 | 98.74 | 98.31 | 98.30 | 98.29 | 97.89 |
| Additionally allowed (%) | 2.33 | 1.74 | 1.26 | 1.69 | 1.70 | 1.71 | 2.11 |
| Outliers (%) | 0 | 0 | 0 | 0 | 0 | 0 | 0 |

[a] SSRL, Stanford Synchrotron Radiation Lightsource
[b] APS, Advanced Photon Source
[c] Highest-resolution shell is shown in parentheses
[d] Unique reflections distinguish anomalous Friedel pairs
[e] As determined by MolProbity (*128*)

**Table S15.** X-ray crystallographic analysis of NUP153 ZnF•Ran(GDP) complexes

| | | | | | | | | | |
|---|---|---|---|---|---|---|---|---|---|
| **Data collection** | | | | | | | | | |
| Protein 1 | Ran(GDP) F35S | Ran(GDP) F35S | Ran(GDP) F35S | Ran(GDP) F35S | Ran(GDP) F35S | Ran(GDP) F35S | Ran(GDP) F35S | Ran(GDP) F35S | Ran(GDP) F35S |
| Expression construct | Sumo | Thrombin | Sumo | Thrombin | Sumo | Sumo | Thrombin | Sumo | Thrombin |
| Protein 2 | NUP153$^{649-687}$ ZnF1 | NUP153$^{651-687}$ ZnF1 | NUP153$^{713-749}$ ZnF2 | NUP153$^{713-749}$ ZnF2 | NUP153$^{781-817}$ ZnF3 | NUP153$^{781-817}$ ZnF3 | NUP153$^{781-817}$ ZnF3 | NUP153$^{838-874}$ ZnF4 | NUP153$^{838-874}$ ZnF4 |
| Stoichiometry | 1:1 | 1:1 | 2:2 | 1:1 | 2:2 | 2:2 | 2:2 | 1:1 | 1:1 |
| PDB code | 7MO1 | | 7MO2 | | 7MO3 | 7MO4 | | 7MO5 | |
| Synchrotron | SSRL[a] | SSRL | SSRL | SSRL | SSRL | SSRL | SSRL | SSRL | APS[b] |
| Beamline | BL12-2 | BL12-2 | BL12-2 | BL12-2 | BL12-2 | BL12-2 | BL12-2 | BL12-2 | 23 ID-B |
| Space group | $P2_12_12$ | $P2_12_12$ | $P2_1$ | $P2_12_12$ | $P2_1$ | $P2_1$ | $P2_12_12_1$ | $P2_12_12$ | $P2_12_12$ |
| Cell parameters | | | | | | | | | |
| *a* (Å) | 60.3 | 60.0 | 58.1 | 61.0 | 69.0 | 67.7 | 61.1 | 59.8 | 60.2 |
| *b* (Å) | 79.6 | 79.8 | 60.4 | 79.7 | 62.6 | 61.2 | 80.8 | 80.1 | 78.7 |
| *c* (Å) | 55.1 | 54.8 | 79.4 | 57.7 | 70.6 | 69.8 | 114.4 | 54.5 | 54.8 |
| $\alpha$ (°) | 90.0 | 90.0 | 90.0 | 90.0 | 90.0 | 90.0 | 90.0 | 90.0 | 90.0 |
| $\beta$ (°) | 90.0 | 90.0 | 90.1 | 90.0 | 105.1 | 104.0 | 90.0 | 90.0 | 90.0 |
| $\gamma$ (°) | 90.0 | 90.0 | 90.0 | 90.0 | 90.0 | 90.0 | 90.0 | 90.0 | 90.0 |
| | *Native* | *Zn peak* | *Native* | *Zn peak* | *Native* | *Native* | *Zn peak* | *Native* | *Zn peak* |
| Wavelength (Å) | 1.03317 | 1.28272 | 1.03317 | 1.28272 | 1.03317 | 0.97946 | 1.28272 | 1.03317 | 1.28279 |
| Resolution (Å) | 30.0 – 1.6 | 30.0 – 1.6 | 30.0 – 1.65 | 30.0 – 1.9 | 30.0 – 2.05 | 30.0 – 2.4 | 30.0 – 1.6 | 30.0 – 1.55 | 30.0 – 1.85 |
| $R_{merge}$ (%)[c] | 7.8 (171.7) | 5.8 (89.8) | 7.3 (107.9) | 8.6 (161.3) | 4.7 (169.4) | 7.9 (135.2) | 5.1 (115.8) | 6.8 (173.1) | 6.7 (134.2) |
| $R_{meas}$ (%)[c] | 8.1 (179.0) | 6.3 (97.8) | 7.9 (118.6) | 9.4 (175.6) | 5.5 (197.7) | 8.6 (146.3) | 5.5 (126.0) | 7.4 (187.4) | 7.3 (148.2) |
| $R_{pim}$ (%)[c] | 3.0 (49.8) | 2.4 (38.0) | 3.0 (48.1) | 3.6 (68.4) | 2.8 (101.3) | 3.3 (55.2) | 2.1 (49.1) | 2.8 (71.1) | 2.8 (61.5) |
| $CC_{1/2}$ (%)[c] | 100.0 (65.3) | 99.9 (87.3) | 99.9 (49.3) | 99.9 (77.8) | 99.9 (57.8) | 99.8 (75.1) | 100.0 (74.3) | 99.9 (58.8) | 99.9 (63.5) |
| $\langle I \rangle / \langle \sigma I \rangle$[c] | 19.0 (1.5) | 24.6 (2.9) | 13.8 (1.5) | 13.5 (1.2) | 15.7 (1.1) | 11.3 (1.3) | 23.2 (2.0) | 15.3 (2.3) | 20.1 (1.7) |
| Completeness (%)[c] | 100.0 (99.5) | 99.0 (97.5) | 98.5 (96.6) | 99.9 (99.4) | 93.2 (93.7) | 98.6 (98.5) | 99.0 (97.1) | 99.9 (99.9) | 100.0 (100.0) |
| No. observations[c] | 467,759 (22,144) | 445,449 (42,096) | 444,419 (18,540) | 282,478 (27,077) | 247,248 (24,296) | 145,075 (14,427) | 961,259 (88,740) | 498,936 (50,078) | 291,124 (23,395) |
| No. unique reflections[c,d] | 35,851 (1,762) | 35,072 (3,423) | 65,381 (3,167) | 22,744 (2,221) | 34,158 (3,415) | 21,520 (2,104) | 74,712 (7,263) | 38,670 (50,078) | 22,891 (2,238) |
| Redundancy[c] | 13.0 (12.6) | 6.7 (6.4) | 68 (5.9) | 12.4 (12.2) | 7.2 (7.1) | 6.7 (6.9) | 12.9 (12.2) | 12.9 (13.1) | 12.7 (10.5) |
| Wilson B factor | 21 | 23 | 21 | 38 | 49 | 63 | 23 | 21 | 31 |
| **Refinement** | | | | | | | | | |
| Resolution (Å) | 30.0 – 1.6 | | 30.0 – 1.65 | | 30.0 – 2.05 | 30.0 – 2.4 | | 30.0 – 1.55 | |
| No. reflections total | 67,816 | | 127,956 | | 66,213 | 41,648 | | 38,623 | |
| No. reflections test set | 3,328 (4.91%) | | 6,340 (5.03%) | | 3,136 (4.74%) | 2,102 (5.05%) | | 3,536 (4.83%) | |
| $R_{work}$ / $R_{free}$ | 15.1 / 17.5 | | 21.0 / 24.0 | | 20.4 / 22.8 | 20.4 / 23.7 | | 16.2 / 19.0 | |
| No. atoms | 2,235 | | 4,320 | | 4,004 | 3,914 | | 2,176 | |
| Protein | 1,983 | | 3,832 | | 3,861 | 3,828 | | 1,918 | |
| Ligand | 30 | | 60 | | 60 | 60 | | 30 | |
| Water | 222 | | 428 | | 83 | 26 | | 228 | |
| *B*-factors | 35 | | 38 | | 89 | 98 | | 36 | |
| Protein | 35 | | 38 | | 90 | 98 | | 35 | |
| Ligand | 23 | | 29 | | 64 | 79 | | 24 | |
| Water | 39 | | 37 | | 65 | 75 | | 40 | |
| R.m.s. deviations | | | | | | | | | |
| Bond lengths (Å) | 0.013 | | 0.012 | | 0.013 | 0.012 | | 0.013 | |
| Bond angles (°) | 1.2 | | 0.9 | | 0.9 | 0.8 | | 1.2 | |
| **Ramachandran plot**[e] | | | | | | | | | |
| Favored (%) | 97.84 | | 97.66 | | 97.70 | 97.90 | | 97.86 | |
| Additionally allowed (%) | 2.16 | | 2.34 | | 2.30 | 2.10 | | 2.14 | |
| Outliers (%) | 0.0 | | 0.0 | | 0.0 | 0.0 | | 0.0 | |

[a] SSRL, Stanford Synchrotron Radiation Lightsource
[b] APS, Advanced Photon Source
[c] Highest-resolution shell is shown in parentheses
[d] Unique reflections distinguish anomalous Friedel pairs
[e] As determined by MolProbity (*128*)

**Table S16.** X-ray crystallographic analysis of NUP358/50 RanBD•Ran(GMPPNP) complexes

| | | | | | |
|---|---|---|---|---|---|
| **Data collection** | | | | | |
| Protein 1 | NUP358$^{1171\text{-}1306}$ RanBD1 | NUP358$^{2011\text{-}2148}$ RanBD2 | NUP358$^{2309\text{-}2443}$ RanBD3 | NUP358$^{2911\text{-}3045}$ RanBD4 | NUP50$^{337\text{-}468}$ RanBD |
| Variant | Wildtype | Wildtype | Wildtype | WHTMKNYY/QNYDNKQV | Wildtype |
| Protein 2 | Ran(GMPPNP) | Ran(GMPPNP) | Ran(GMPPNP) | Ran(GMPPNP) | Ran(GMPPNP) |
| Stoichiometry | 4:4 | 6:6 | 6:6 | 6:6 | 2:2 |
| PDB code | 7MNW | 7MNX | 7MNY | 7MNZ | 7MO0 |
| Synchrotron | APS[a] | SSRL[b] | APS[a] | SSRL | APS[a] |
| Beamline | 23 ID-B | BL 12-2 | 23 ID-B | BL 12-2 | 23 ID-D |
| Space group | I222 | $P2_12_12_1$ | $P2_12_12_1$ | $P3_1$ | $P2_12_12_1$ |
| Cell parameters | | | | | |
| *a* (Å) | 137.4 | 94.2 | 110.2 | 142.7 | 66.6 |
| *b* (Å) | 137.8 | 134.0 | 136.0 | 142.7 | 73.2 |
| *c* (Å) | 172.9 | 182.5 | 158.8 | 96.7 | 152.9 |
| α (°) | 90.0 | 90.0 | 90.0 | 90.0 | 90 |
| β (°) | 90.0 | 90.0 | 90.0 | 90.0 | 90 |
| γ (°) | 90.0 | 90.0 | 90.0 | 120.0 | 90 |
| | *Native* | *Native* | *Native* | *Native* | *Se peak* |
| Wavelength (Å) | 1.00000 | 0.97946 | 1.00000 | 1.03317 | 0.97942 |
| Resolution (Å) | 30.0 – 2.40 | 30.0 – 2.4 | 30.0 – 2.70 | 30.0 – 2.35 | 30.0 – 2.45 |
| $R_{merge}$ (%)[c] | 28.6 (267.8) | 16.1 (230.4) | 21.7 (174.4) | 14.1 (164.5) | 11.0 (250.5) |
| $R_{meas}$ (%)[c] | 30.8 (288.2) | 17.4 (251.3) | 23.4 (188.3) | 15.8 (185.2) | 11.9 (270.8) |
| $R_{pim}$ (%)[c] | 11.5 (106.1) | 6.6 (98.8) | 8.8 (70.9) | 6.9 (84.1) | 4.5 (102.3) |
| $CC_{1/2}$ (%)[c] | 99.7 (47.0) | 99.7 (54.7) | 99.6 (69.9) | 99.8 (59.9) | 99.8 (55.2) |
| $<I> / <\sigma I>$[c] | 6.7 (1.3) | 9.7 (1.1) | 8.4 (1.5) | 12.4 (1.8) | 15.3 (1.1) |
| Completeness (%)[c] | 99.8 (98.2) | 98.8 (96.9) | 100.0 (99.8) | 99.9 (99.8) | 100.0 (99.6) |
| No. observations[c] | 880,269 (87,689) | 1,189,731 (107,541) | 900,779 (88,275) | 935,004 (86,810) | 369,132 (36,629) |
| No. unique reflections[c,d] | 64,146 (6,219) | 89,794 (8,716) | 66,178 (6,497) | 91,578 (9,170) | 28,242 (2,762) |
| Redundancy[c] | 13.7 (14.1) | 13.2 (12.3) | 7.1 (7.0) | 10.2 (9.5) | 13.1 (13.3) |
| Wilson B factor | 36 | 46 | 44 | 44 | 56 |
| **Refinement** | | | | | |
| Resolution (Å) | 30.0 – 2.4 | 30.0 – 2.4 | 30.0 – 2.7 | 30.0 – 2.35 | 30.0 – 2.45 |
| No. reflections total | 64,143 | 89,660 | 66,076 | 91,515 | 28,183 |
| No. reflections test set | 3,185 (4.97%) | 2,353 (2.62%) | 3,310 (5.01%) | 4,426 (4.84%) | 2,618 (4.94%) |
| $R_{work}$ / $R_{free}$ (%) | 20.2 / 24.1 | 19.2 / 23.0 | 21.4 / 26.2 | 18.3 / 22.5 | 21.2 / 23.9 |
| No. atoms | 11,709 | 16,979 | 16,688 | 17,360 | 5,503 |
| Protein | 11,177 | 16,397 | 16,223 | 16,669 | 5,369 |
| Ligand | 132 | 198 | 198 | 270 | 71 |
| Water | 400 | 384 | 267 | 473 | 63 |
| *B*-factors | 56 | 69 | 76 | 60 | 78 |
| Protein | 56 | 70 | 77 | 61 | 79 |
| Ligand | 45 | 47 | 50 | 42 | 67 |
| Water | 44 | 55 | 53 | 51 | 59 |
| R.m.s. deviations | | | | | |
| Bond lengths (Å) | 0.004 | 0.004 | 0.003 | 0.002 | 0.004 |
| Bond angles (°) | 0.7 | 0.7 | 0.6 | 0.5 | 0.7 |
| **Ramachandran plot**[e] | | | | | |
| Favored (%) | 98.30 | 98.10 | 98.00 | 97.45 | 98.00 |
| Additionally allowed (%) | 1.70 | 1.90 | 2.00 | 2.55 | 2.00 |
| Outliers (%) | 0 | 0 | 0 | 0 | 0 |

[a] APS, Advanced Photon Source
[b] SSRL, Stanford Synchrotron Radiation Lightsource
[c] Highest-resolution shell is shown in parentheses
[d] Unique reflections distinguish anomalous Friedel pairs
[e] As determined by MolProbity (*128*)

**Table S17.** X-ray crystallographic analysis of NUP88$^{NTD}$•NUP98$^{APD}$

| | |
|---|---|
| **Data collection** | |
| Protein 1 | NUP88$^{1\text{-}493}$ NTD |
| Protein 2 | NUP98$^{732\text{-}880}$ APD |
| Stoichiometry | 2:2 |
| PDB code | 7MNI |
| Synchrotron | APS[a] |
| Beamline | 23 ID-B |
| Space group | P1 |
| Cell parameters | |
| *a* (Å) | 59.8 |
| *b* (Å) | 72.1 |
| *c* (Å) | 73.0 |
| α (°) | 92.5 |
| β (°) | 103.7 |
| γ (°) | 100.4 |
| | *Native* |
| Wavelength (Å) | 1.00003 |
| Resolution (Å) | 20.0 – 2.0 |
| $R_{merge}$ (%)[c] | 13.1 (98.4) |
| $R_{meas}$ (%)[c] | 14.3 (115.7) |
| $R_{pim}$ (%)[c] | 5.6 (60.4) |
| $CC_{1/2}$ (%)[c] | 99.6 (51.3) |
| $\langle I \rangle / \langle \sigma I \rangle$[c] | 7.5 (1.3) |
| Completeness (%)[c] | 97.8 (96.6) |
| No. observations[c] | 456,229 (16,182) |
| No. unique reflections[c,d] | 76,633 (4,472) |
| Redundancy[c] | 6.0 (3.6) |
| Wilson B factor | 36 |
| **Refinement** | |
| Resolution (Å) | 20.0 – 2.0 |
| No. reflections total | 76,489 |
| No. reflections test set | 3,639 (4.8%) |
| $R_{work}$ / $R_{free}$ (%) | 17.5 / 20.8 |
| No. atoms | 9,442 |
| Protein | 9,024 |
| Ligand | 0 |
| Water | 418 |
| *B*-factors | 56 |
| Protein | 56 |
| Ligand | |
| Water | 49 |
| R.m.s. deviations | |
| Bond lengths (Å) | 0.002 |
| Bond angles (°) | 0.5 |
| **Ramachandran plot**[e] | |
| Favored (%) | 98.63 |
| Additionally allowed (%) | 1.28 |
| Outliers (%) | 0.09 |

[a] APS, Advanced Photon Source
[b] SSRL, Stanford Synchrotron Radiation Lightsource
[c] Highest-resolution shell is shown in parentheses
[d] Unique reflections distinguish anomalous Friedel pairs
[e] As determined by MolProbity (*128*)
